# Multiscale mapping of transcriptomic signatures for cardiotoxic drugs

**DOI:** 10.1101/2021.11.02.466774

**Authors:** Jens Hansen, Yuguang Xiong, Priyanka Dhanan, Bin Hu, Arjun S. Yadaw, Gomathi Jayaraman, Rosa Tolentino, Yibang Chen, Kristin G. Beaumont, Robert Sebra, Dusica Vidovic, Stephan C. Schürer, Joseph Goldfarb, James Gallo, Marc R. Birtwistle, Eric A. Sobie, Evren U. Azeloglu, Seth Berger, Angel Chan, Christoph Schaniel, Nicole C. Dubois, Ravi Iyengar

**Affiliations:** Mount Sinai Institute for Systems Biomedicine, Icahn School of Medicine at Mount Sinai, New York, NY 10029, USA; Department of Pharmacological Sciences, Icahn School of Medicine at Mount Sinai, New York, NY 10029, USA; Department of Genetics and Genomic Sciences, Icahn School of Medicine at Mount Sinai, New York, NY 10029, USA; Institute for Data Science and Computing, University of Miami, Coral Gables, FL 33146, USA; School of Pharmacy and Pharmaceutical Sciences, University of Buffalo SUNY System, Buffalo NY 14260; Chemical and Biomolecular Engineering, Clemson University, Clemson, SC, 29634, USA; Department of Medicine, Division of Nephrology, Icahn School of Medicine at Mount Sinai, New, York, NY 10029, USA; Departments of Pediatrics and Genomics & Precision Medicine, George Washington University School of medicine and Health Sciences, Washington DC USA, 20012; Cardiology Division, Department of Medicine, Memorial Sloan Kettering Cancer Center New York NY 10065; Department of Medicine, Division of Hematology and Medical Oncology, Tisch Cancer Institute, Icahn School of Medicine at Mount Sinai, New York, NY 10029, USA; Department of Cell, Developmental and Regenerative Biology, Icahn School of Medicine at Mount Sinai, New York, NY 10029, USA

## Abstract

Drug-induced gene expression profiles can identify potential mechanisms of toxicity. We focused on obtaining signatures for cardiotoxicity of FDA-approved tyrosine kinase inhibitors (TKIs) in human induced pluripotent stem cell-derived cardiomyocytes. Using bulk transcriptomics profiles, we applied singular value decomposition to identify drug-selective patterns in cell lines obtained from multiple healthy human subjects. Cellular pathways affected by highly cardiotoxic TKIs include energy metabolism, contractile, and extracellular matrix dynamics. Projecting these pathways to single cell expression profiles indicates that TKI responses can be evoked in both cardiomyocytes and fibroblasts. Whole genome sequences of the cell lines, using outlier responses enabled us to correctly reidentify a genomic variant associated with anthracycline cardiotoxicity and predict genomic variants potentially associated with TKI cardiotoxicity. We conclude that mRNA expression profiles when integrated with publicly available genomic, pathway, and single cell transcriptomic datasets, provide multiscale predictive understanding of cardiotoxicity for drug development and patient stratification.

**One sentence summary:** Genes, pathways, and cell types of the human heart associated with antineoplastic drug cardiotoxicity.

## Introduction

Adverse side-effects of therapeutically useful drugs continue to be a substantial problem ^1^, post-approval pharmacovigilance studies often revealing adverse events that lead to warning labels mandated by the FDA ^2^. Early indications of a potential for adverse events will be useful in drug development ^3,4^ and the use of whole genome sequence data from individuals can predict who might be susceptible to adverse events. These assertions are based on the premise that various functions at the molecular and cellular levels drive adverse events in different cell types ^5^. Preclinical studies at the molecular level are useful, as has been demonstrated by HERG channel protein interacting drugs and the potential for arrhythmias ^6,7^. Drug-related adverse events are often organ-selective. Many efficacious antineoplastic drugs, such as tyrosine kinase inhibitors (TKI) that are used for targeted therapy, are associated with cardiac insufficiencies and development of heart failure ^8,9^. We have shown that drug-induced transcriptomic profiles in adult human heart cells can be associated with clinical adverse event propensity as assessed from FDA pharmacovigilance data ^10^. However, a systematic understanding of the molecular pathways, cell origins and genomic determinants associated with drug therapy-related cardiotoxicity are still not well understood.

Human induced pluripotent stem cell (hiPSC)-derived cardiomyocytes ^11^ have been useful for understanding cardiotoxicity ^12^. Hence, transcriptional profiles in human cardiac cells can form starting points for studies focused on mechanism-based drug signatures that could be used for prediction of cardiotoxicity potential. Here, we have used six hiPSC-derived cardiac cell lines from healthy individuals to study 54 FDA-approved drugs and identify drug-selective transcriptomic signatures across the different human subject lines by singular value decomposition (SVD)-based analysis of total transcriptomic responses. Single cell analyses of hiPSC-derived “ventricular cardiomyocytes” indicated that they are composed of multiple clusters. They include a cluster very similar to adult cardiomyocytes. Additional clusters show varying levels of similarity to different cell types in the human heart, including fibroblasts. Projections of drug affected pathways inferred from bulk transcriptomic data of hiPSC-derived cardiomyocytes on to single cell expression profiles of human hearts from healthy and heart failure patients indicate that TKIs can affect both cardiomyocytes as well as other cell types to produce adverse events, and that the drug-induced pathways are similar to those altered in hiPSC-derived cardiomyocytes from patients with failing hearts ^13,14^. Using outlier analyses and Whole Genome Sequencing (WGS) from the cell lines used for the transcriptomic studies, we correctly re-identified the genomic variant associated with anthracycline cardiotoxicity that was originally discovered in genome-wide association studies (GWAS) ^15^. We used this approach to identify and predict potential effects of genomic variants associated with TKI action as identified by the transcriptomic signature. These findings indicate that integration of experimental data with public data sets can be powerful drivers of multiscale understanding of organ level adverse events associated with drug therapy.

## Results

To identify pathway activities and potential genomic variants associated with TKI-induced cardiotoxicity (TIC), we treated six hiPSC-derived cardiomyocyte cell lines from six healthy individuals ^16^ (Suppl. Figures 1A/B) (Suppl. Table 1) with one of 27 TKIs, 4 anthracyclines and 26 other cardiac and noncardiac acting drugs (Suppl. Table 2) for 48 hours (Suppl. Table 3 ^16^). All drugs are FDA-approved. The group of TKIs contained 23 small molecule TKIs and 4 monoclonal antibodies against TKs. They could be separated into 10 cardiotoxic and 17 non-cardiotoxic TKIs based on the results of clinical studies (Suppl. Table 4) and FAERS data analysis (Suppl. Figure 2). Bulk transcriptomic analysis of control ^16^ and treated cell lines generated 266 lists of differentially expressed genes (DEGs) (Suppl. Table 5) (Suppl. Table 6 shows averaged DEGs). All lists of DEGs, each representing the response observed for one sample, i.e. a unique cell line/drug combination (Suppl. Figure 3A), were subjected to pairwise correlation and hierarchical clustering (Suppl. Figure 3B).

### Singular value decomposition to unmask drug-selective expression responses

To identify drug-selective expression responses we developed a computational pipeline that uses Singular Value Decomposition (SVD) to search for shared gene expression components in multiple cell lines treated with the same drug (Figure 1A, Suppl. Figure 4). Clustering of the transcriptomic data only grouped a few samples treated with the same drugs into the same cluster (Figure 1B, Suppl. Figure 3B). On the contrary, the treated cell line or the amplitude of the drug response, i.e., the number of significant DEGs, mainly determined the clustering results (Suppl. Figure 5A). To quantify drug-selective clustering, we calculated one F1 score for each drug that documents how close all cell lines treated with that drug cluster together (Suppl. Figure 5B). With possible values from 0 to 1, the median F1-score of 0.116 documents very low drug-selective clustering efficiencies (Figure 1C).

**Figure 1.**
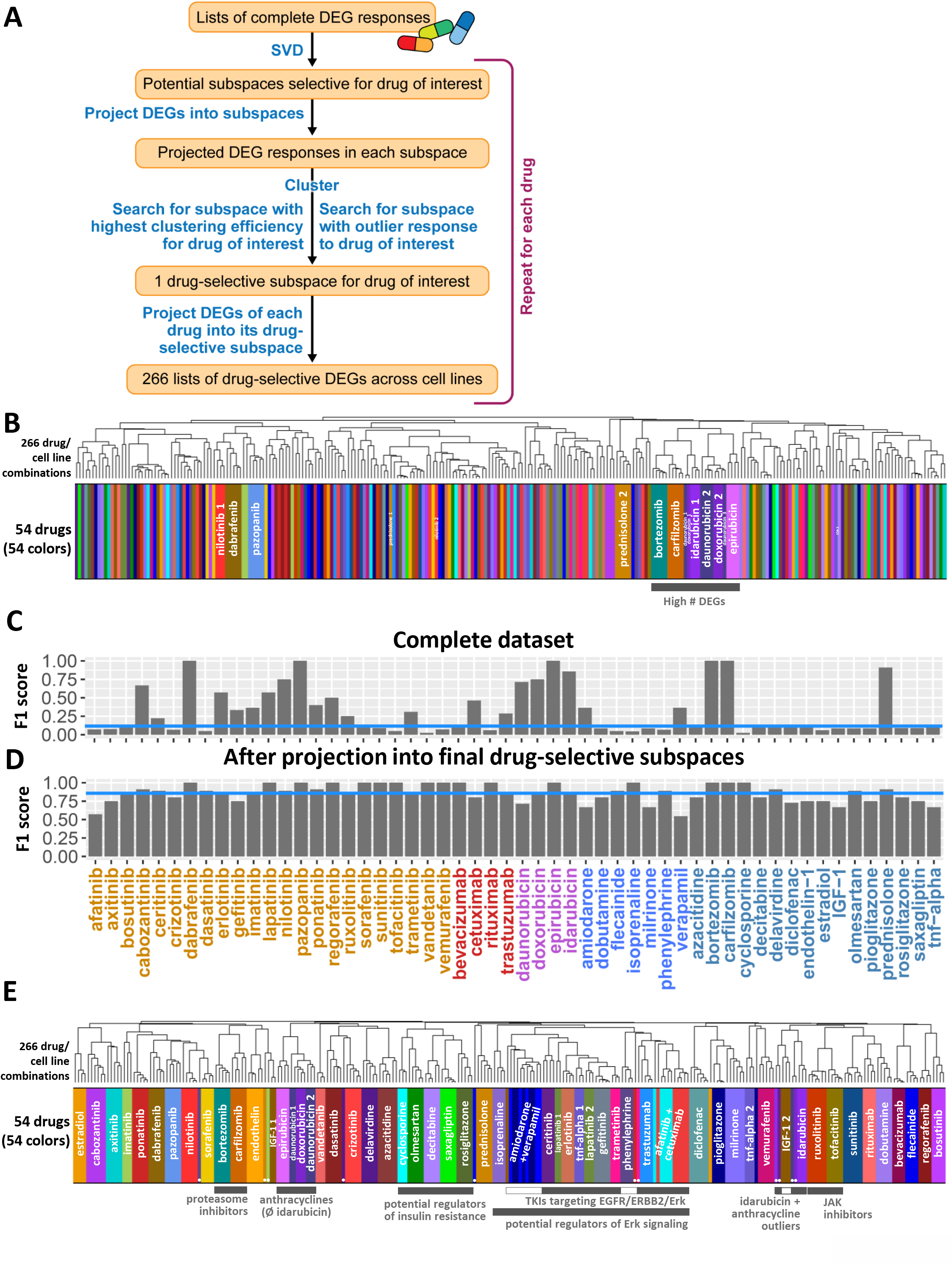
Singular value decomposition identifies drug-selective gene expression responses. 266 samples, each representing a unique combination of one out of three to six hiPSC-derived cardiomyocyte cell lines treated with one out of 54 drugs, were subjected to bulk RNAseq analysis, followed by calculation of 266 lists of drug-induced, differentially expressed genes (DEGs). **(A)** Our computational pipeline uses singular value decomposition (SVD) to identify drug-selective gene expression responses that are components of the complete responses. See methods section and Suppl. Figure 4 for details. **(B)** Pairwise correlation analysis followed by hierarchical clustering reveals that most complete DEGs are dominated by cell-line-selective effects hiding drug-selective effects. See Suppl. Figure 5 for larger dendrogram. **(C)** We used F1 score statistics to document the clustering efficiency, i.e. how close samples treated with the same drug cluster together. Low clustering efficiencies quantitatively describe the finding that only a few complete DEG responses are dominated by drug-selective effects. orange: Small molecule kinase inhibitors (KI), red: monoclonal antibodies against KIs, purple: anthracyclines, blue: cardiac acting drugs, gray-blue: noncardiac acting drugs. **(D)** Projection of the complete DEG responses into each of the identified 54 drug-selective subspaces greatly increases the clustering efficiencies for all 54 drugs. **(E)** Pairwise correlation of 266 merged drug-selective responses, followed by hierarchical clustering, documents that SVD allows identification of components induced in all cell lines treated with the same drug. Clusters that contain drugs with similar mechanisms are labeled with gray bars. White insets indicate drugs in those clusters that are not part of the outlined mechanisms written below the bars. White circles indicate outlier samples that were identified by our pipeline and cluster as outliers in the merged dataset as well. See Suppl. Figure 13 for larger dendrogram.

SVD (Suppl. Figure 6A) of all 266 samples identified 266 eigenarrays (Suppl. Figure 6B) whose linear combination using sample-specific coefficients gives the complete gene expression profiles. The first eigenarray correlates (Suppl. Figure 6C) with the amplitude of the response (Suppl. Figure 6D), and its most prominent genes (Suppl. Table 7) enrich for muscle contraction (Suppl. Figure 6E, Suppl. Table 8). These results indicate that the first eigenarray describes a general response to perturbation that masks drug-selective effects. Removal of the first eigenarray (Suppl. Table 9) disrupts the clustering by the number of significant DEGs (Suppl. Figure 7). Since the clustering is still dominated by cell-line-selective effects, the removal did not markedly improve the drug-selective clustering efficiencies (Suppl. Figure 8).

We ranked the remaining eigenarrays by their ability to separate samples of each drug or cell line from all other samples (Suppl. Figure 9A). Our algorithm allowed a clear separation of cell-line and drug-selective effects (Suppl. Figure 9B). For each drug, we combined unique sets of drug-selective eigenarrays to drug-selective subspaces with the intention to optimize the drug’s clustering efficiency (Suppl. Figure 10A) while preserving a maximum of the original information, as quantified by cosine similarity. Changing the relative contribution of F1 scores (Suppl. Figure 10B) and cosine similarities (Suppl. Figure 10C) we screened the potential subspaces for outlier responses where one cell line showed a significantly different response to the drug of interest than all other cell lines (Suppl. Figures 10D, 11, 12). Such outlier responses will be linked to our genomic analysis at a later stage. For 24 drugs, we could identify subspaces with outlier responses (Suppl. Figure 10E), for the other 30 drugs, we selected subspaces based on high relative contribution of the F1 score (95%).

### Drug-selective gene expression profiles are similar for drugs with similar mechanisms

Projection of gene expression profiles into the final drug-selective subspaces generated drug-selective gene expression profiles (Suppl. Table 10) (averaged DEGs in Suppl. Table 11) with high clustering efficiencies (Figure 1D, Suppl. Figure 10F). Drug-selective DEGs were combined into a new matrix. Clustering of this matrix identifies clusters that mostly contained samples that were treated with the same drug and documented preservation of outlier characteristics (Figure 1E, Suppl. Figure 13). Eleven of the 24 identified outlier samples are clearly separated from the cell lines treated with the same drug (closed circles in Figure 1E and Suppl. Figure 13). Another eight outlier samples are grouped together with the samples treated with the same drug in a larger cluster (open circles in Suppl. Figure 13), but are separated from those, once that cluster is subclustered. Only four identified outliers reside in the same cluster (open rectangles) as the other cell lines treated with the same drug. Comparison of Figures 1B and 1E in terms of the breadth of the colored bars allows for visualization of this drug selective clustering.

Our algorithm identified drug-selective gene expression profiles for each drug independently. Nevertheless, drugs with similar mechanisms and overlapping targets still cluster together (Figure 1E, Suppl. Figure 13). All eight TKIs targeting EGFR signaling, either by inhibiting the receptor (EGFR, ERBB2) or its intracellular ERK signaling cascade (MAP2K1, MAP2K2) ^17^ are part of the same cluster. One of the four non-TKI drugs in this cluster, phenylephrine, and isoprenaline which is part of a slightly bigger cluster, stimulate adrenergic signaling that can cross-activate ERK signaling in the heart ^18^, as directly documented for phenylephrine in the perfused rat heart ^19^. Two additional non-TKI drugs in this cluster, verapamil and amiodarone, influence adrenergic signaling as antagonists ^20-22^ and verapamil has been shown to antagonize ERK signaling in rat cardiomyocytes ^23^. Similarly, the JAK inhibitors ruxolitinib and tofacitinib and the proteasome inhibitors bortezomib and carflizomib are part of two independent clusters that contain no other drugs. Two of three antidiabetics, rosiglitazone and saxagliptin are part of a smaller cluster that first merges with a cluster containing decitabine, a drug with hyperglycemia as a major side effect ^24^ and then with a cluster containing cyclosporine and olmesartan. Cyclosporine increases insulin resistance in humans ^25^ and olmesartan -/+ amlodipine ameliorates it in rats ^26^ and patients ^27^, respectively. The anthracyclines epirubicin, daunorubicin and doxorubicin are part of larger cluster, while the anthracycline idarubicin clusters together with their identified outlier samples.

In summary, our SVD pipeline uncovers similar expression profiles induced by similar drugs. Since it does not actively search for such similarities, but only aims at maximizing the overlap between DEGs induced by the same drug, this finding serves as internal validation for the extraction of drug-selective components by our computational pipeline.

The grouping of drugs with similar mechanisms into higher-level clusters explains eleven non-outlier samples that are separated from the other samples treated with the same drug (closed rhombuses), leaving only four samples that are not part of drug-selective clusters for no obvious biological reason (open rhombuses)(Suppl. Fig. 13).

### SVD enables identification of subcellular processes relevant to TKI cardiotoxicity

Complete and decomposed DEGs were subjected to pathway enrichment analysis using the Molecular Biology of the Cell Ontology (MBCO) (20) (Suppl. Figure 14A). MBCO subcellular processes (SCPs) are organized in three to four levels where higher-level SCPs (i.e., levels with lower numbers) describe more general, and lower-level SCPs more detailed, cell biological processes. Predicted up- or downregulated SCPs of each drug/cell line combination and SCP level were ranked by significance (Suppl. Tables 12/13/14). We will refer to these ranks as enrichment ranks in the following.

Overall comparison of the enrichment results before and after decomposition documents a great increase in the number of overlapping SCPs in different cell lines treated with the same drug (Suppl. Figures 14B/C/D/E, 15A).

In addition, our decomposition pipeline allowed identification of SCPs that were masked in the complete dataset. For example, we could document downregulation of SCPs involved in single protein degradation as an anthracycline-specific effect (Suppl. Figure 15B), results that we did not obtain using the complete DEGs (Suppl. Figure 15C). For a discussion of these SCPs, their clinical relevance for supportive treatment and examples that demonstrate agreement with prior knowledge from small-scale experiments see Supplemental Information.

### Subcellular processes and cell types indicative of cardiotoxic response to TKI treatment

To identify signatures associated with TKI-induced cardiotoxicity we searched for SCPs that are almost exclusively up- or downregulated at higher enrichment ranks by cardiotoxic TKIs as compared to noncardiotoxic TKIs, using F1 score and Area under the Curve (AUC) statistics (Figure 2A, Suppl. Figure 16). SCPs were ranked by decreasing AUC. The top 10, 10, 25 and 10 level-1, -2, -3 and -4 SCPs were grouped by functional similarities (Figure 2B, Suppl. Figure 18) and integrated into the MBCO hierarchy (Figure 2C, Suppl. Figure 19). In addition, we mapped identified SCPs back to the cardiotoxic TKIs that up- or downregulate them (Suppl. Figure 20).

**Figure 2.**
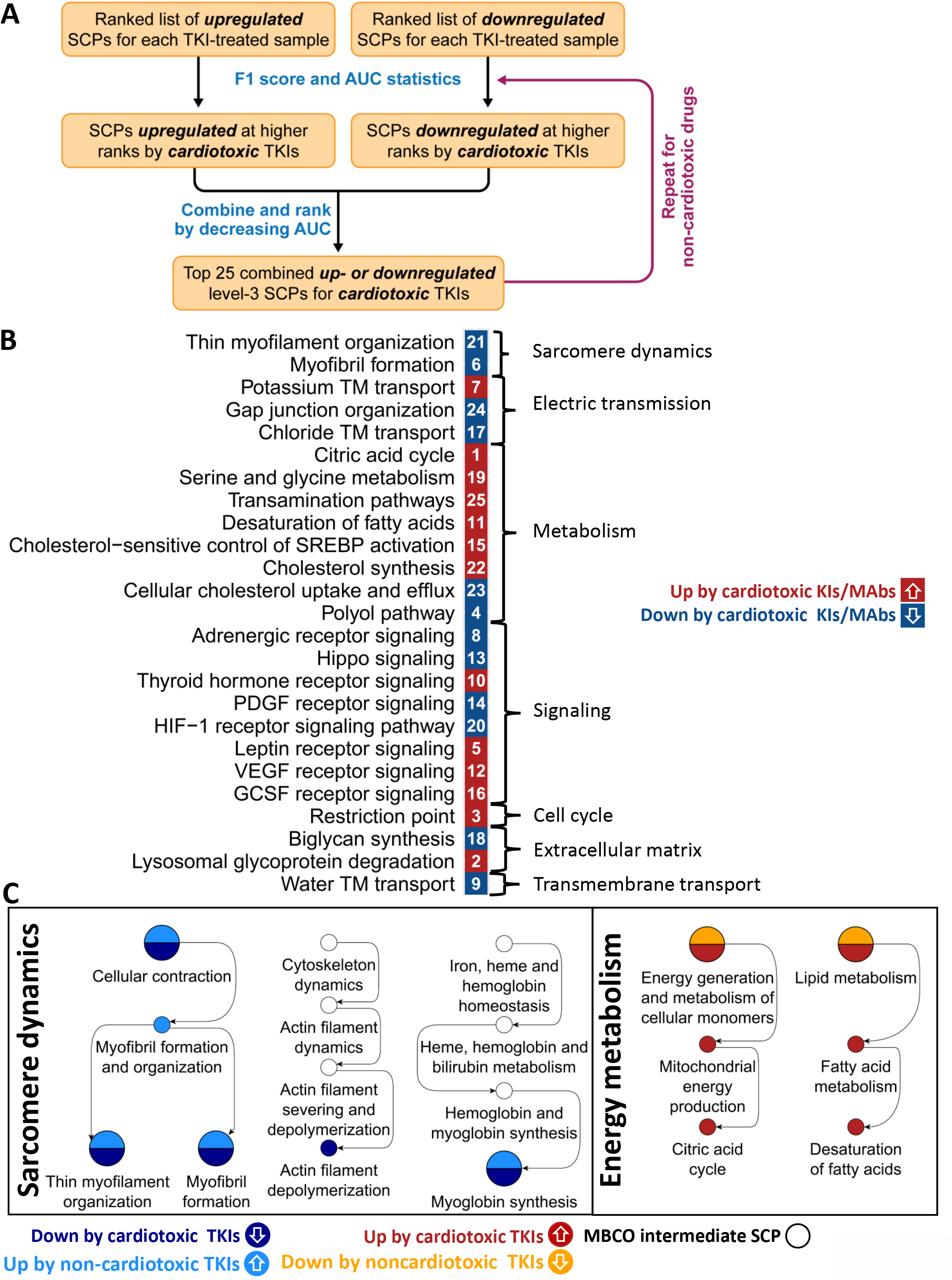
Potential subcellular processes indicative of drug induced cardiotoxicity. Up- and downregulated genes among the top 600 drug-selective gene expression profiles were subjected to pathway enrichment analysis using the Molecular Biology of the Cell Ontology (MBCO). Significantly up- or downregulated subcellular processes (SCPs) (p-value ≤ 0.05) were ranked separately by significance for each sample and SCP level. **(A)** We developed a computational pipeline that screens for SCPs that are up- or downregulated at higher ranks by cardiotoxic (or non-cardiotoxic) TKIs. Up- and downregulated SCPs are ranked based on F1 and Area under the Curve statistics. See methods for details. **(B)** Top 25 level-3 SCPs predicted for the cardiotoxic TKIs were grouped based on the higher-level functions. **(C)** SCPs identified for cardiotoxic and non-cardiotoxic TKIs for all levels were integrated into the MBCO hierarchy. Selected branches are shown.

Besides being elevated or decreased above or below a threshold that determines an SCP’s association with a cardiotoxic response, an SCP’s activity might already be beyond that threshold at baseline level, i.e., before TKI treatment. Candidates for this group should be those SCPs that are regulated by non-cardiotoxic TKIs. A non-cardiotoxic TKI might change the baseline SCP activity from sufficient to insufficient for a cardiotoxic response. Consequently, our second focus was the investigation of the SCPs that our algorithm identified, if applied to non-cardiotoxic TKIs in the same manner (Suppl. Figures 18, 19, 21).

Single cell RNA sequencing of four of our six cardiomyocyte cell lines identified at least two major subtypes – adult cardiomyocytes and an epicardium derived cell type that is similar to fibroblasts ^16^ (Suppl. Figure 22A-D, Suppl. Tables 15,16A/B). The functions of SCPs we identified by bulk transcriptomics often represent canonical functions of cell types other than cardiomyocytes. We determined how the top SCPs in our data map to our subtypes (Suppl. Figure 23, Suppl. Tables 16C-F), and to the major cell types of the adult human heart analyzed by others using single cell transcriptomics ^28^ (Figure 3A, Suppl. Figure 23, Suppl. Tables 17, 18A-D).

**Figure 3.**
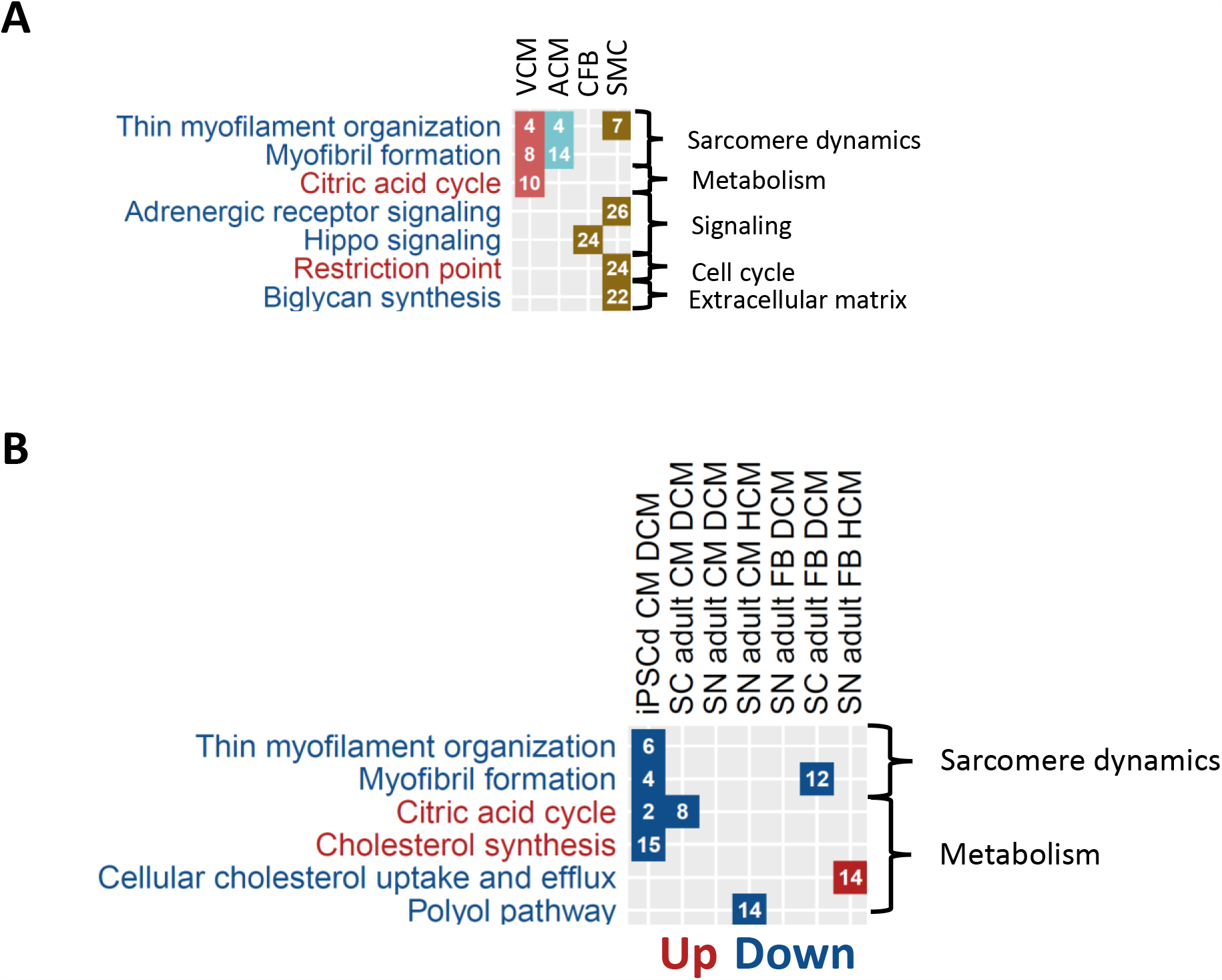
SCPs can be mapped to cellular subtypes and known cardiomyopathy disease mechanisms. **(A)** We subjected marker genes for ventricular and atrial cardiomyocytes (VCM and ACM, respectively), cardiac fibroblasts (CFB) and smooth muscle cells (SMC) obtained from single nucleus RNAseq of the adult human heart ^28^ to pathway enrichment analysis using MBCO. Significant SCPs of each cell type (p-value ≤ 0.05) were ranked by significance (numbers in the diagram). Names of SCPs whose higher and lower activities favor a cardiotoxic response are colored red and blue, respectively. **(B)** Differentially expressed genes (DEGs) in heart cells obtained by single cell (SC) ^13^ or nucleus (SN) ^14^ RNAseq from patients with Dilated or Hypertrophic Cardiomyopathy (DCM or HCM, respectively) as well as in hiPSC-derived cardiomyocytes obtained from an infant patient with DCM (GSE184899) were subjected to pathway enrichment analysis. Significantly up- (red fields) or downregulated (blue fields) SCPs of each cell type (p-value ≤ 0.05) were ranked by significance (numbers in the diagram).

Additionally, we mapped our SCPs to SCPs in hiPSC-derived cardiomyocytes obtained from infants with Dilated Cardiomyopathy (DCM) (GSE184899) (Suppl. Table 19) or in adult heart cells of patients with DCM or Hypertrophic Cardiomyopathy (HCM) ^13,14^ (Figure 3A, Suppl. Figure 24, Suppl. Tables 20A/B/C). Many of the identified SCPs described well-known biology involved in inherited and acquired cardiomyopathies that develop independently of chemotherapy and map to multiple cell types of the adult human heart. Our results suggest that cardiotoxic TKIs might mimic these mechanisms by early (within 48 hrs) transcriptional regulation.

Four SCPs involved in muscle contraction and sarcomere renewal were simultaneously identified for the cardiotoxic and noncardiotoxic drugs (Figure 2B, Suppl. Figure 18). Another two SCPs were identified for each group independently. These SCPs that span all four SCP levels (Figure 2C, Suppl. Figure 19) are regulated by the two TKI toxicity groups in the opposite directions. The difference in directionality suggests insufficient sarcomere renewal as relevant for TKI-induced cardiotoxicity. All cardiotoxic TKIs, except vandetinib and trastuzumab downregulate (Suppl. Figure 20), while 13 of 17 non-cardiotoxic TKIs upregulate (Suppl. Figure 21) at least one of these SCPs, respectively. The SCPs map to cardiomyocyte clusters in cell lines and the adult human heart (Figure 3B, Suppl. Figure 23). In agreement with our data, proper heart functioning depends on a continuous turnover of cardiomyocyte sarcomere proteins ^29^. About half of hypertrophic and dilated cardiomyopathies are associated with mutations in sarcomeric proteins ^30^. Further supporting this line of reasoning, we found downregulation of muscle contraction-related SCPs in adult and hiPSC-derived DCM cardiomyocytes (Figure 3C, Suppl. figure 24). These SCPs might serve as a starting point for supportive therapy. Four of the six regulated genes of the SCP ‘Thin myofilament organization’ (Suppl. Table 12) are involved in blocking the binding of the myosin head to the thin myofilament during muscle contraction. This mechanism is targeted by the new drug mavacamten that was recently approved by the FDA to treat obstructive Hypertrophic Cardiomyopathy (HCM) ^31^.

Our data associates the upregulation of SCPs involved in the citric acid cycle and mitochondrial energy generation with a cardiotoxic response (Figures 2B/C, Suppl. Figures 18/19). They are almost exclusively upregulated by pazopanib (Suppl. Figure 20), a TKI with a high rate of cardiotoxicity (>10%) (Suppl. Table 4). Many studies document an overall reduction in oxidative phosphorylation during heart failure ^32^ and identified SCPs are downregulated in adult DCM and HCM cardiomyocytes as well as in hiPSC-derived DCM cardiomyocytes (Figure 3B, Suppl. Figure 24). Nevertheless, in support of our findings, compensatory upregulation of oxidative phosphorylation was suggested for patients belonging to a large DCM subgroup that is caused by truncating titin variants ^33-35^.

Increasing evidence links ferroptosis, an iron-dependent accumulation of lipid peroxides that triggers cell death, to heart failure ^36^. Some identified SCPs in our data might converge on ferroptosis as an endpoint. The SCP ‘Desaturation of fatty acids’ was associated with a cardiotoxic response (Figures 2B/C, Suppl. Figures 18/19) due to its upregulation by pazopanib (Suppl. Figure 20). Upregulated genes mapping to this SCP (Suppl. Table 12) can generate polyunsaturated fatty acids (PUVAs) ^37,38^ that are precursors of lipid peroxides ^36^. A lower activity of the level-2 SCP ‘Cellular antioxidant systems’ and its level-1 parent ‘Cellular redox homeostasis’, pathways offering protection against ferroptosis ^36^, is associated with a cardiotoxic response (Suppl. Figures 18/19). They were downregulated by the cardiotoxic TKIs vandetinib and bevacizumab (Suppl. Figure 20). In addition, downregulation of ‘Cellular iron storage’ by the anthracyclines doxorubicin and daunorubicin identified by us (Suppl. Figures 14D, 15B) and others ^39^ suggests stimulation of ferroptosis by an increase of intracellular free iron ^39^. In agreement, supportive therapy with iron chelators protects against anthracycline-induced cardiotoxicity ^40^.

The clinical relevance of our findings is further supported by identification of SCPs involved in cholesterol metabolism and natriuretic peptide signaling (Figure 2B, Suppl. Figure 18). This is supported by recommended treatment of cardiomyopathy and drug-induced cardiomyopathy with statins ^41^ and neprilysin ^42,43^, respectively. Prediction of extracellular collagen-crosslinking (Suppl. Figure 18) agrees with histopathological observations that correlate with disease progression ^44,45^. Involvement of identified signaling pathways in TIC i.e., PDGF, HIF-1alpha, Oncostatin M and Hippo signaling (Figure 2B, Suppl. Figure 18) is supported by their involvement in drug-independent cardiomyopathy. These and other findings regarding SCPs are discussed in further detail in the supplemental information.

Taken together the data from our experiments, when integrated with data from the literature, indicate the TKI are likely to have their cardiotoxic effects by regulating SCPs in multiple cell types of the heart.

### Candidates for genomic variants associated with drug-induced cardiotoxicity

Using whole genome sequencing of the six cell lines ^16^, our next analysis step focused on the identification of potential genomic variants that might be associated with a higher or lower risk for drug- or more specifically TKI-induced cardiotoxicity (DIC, TIC, respectively) (Figure 4A, Suppl. Figure 25). Cardiotoxic responses to TKI treatment are observed in less than 10% to 20% of the treated patients (Suppl. Table 4). Consequently, we hypothesized that the population-wide frequency of a potential monogenic allele associated with DIC should be in a similar range. Relevance for cardiac tissue was assumed for those variants that map to gene coding regions or are part of cis-expression (e) or -splicing (s) QTLs in the adult human heart ^46^. We hypothesized that potentially cardiotoxic or -protective variants could interfere with a drug’s pharmacokinetics (PK) or -dynamics (PD) or with pathways that are targeted by that drug.

**Figure 4.**
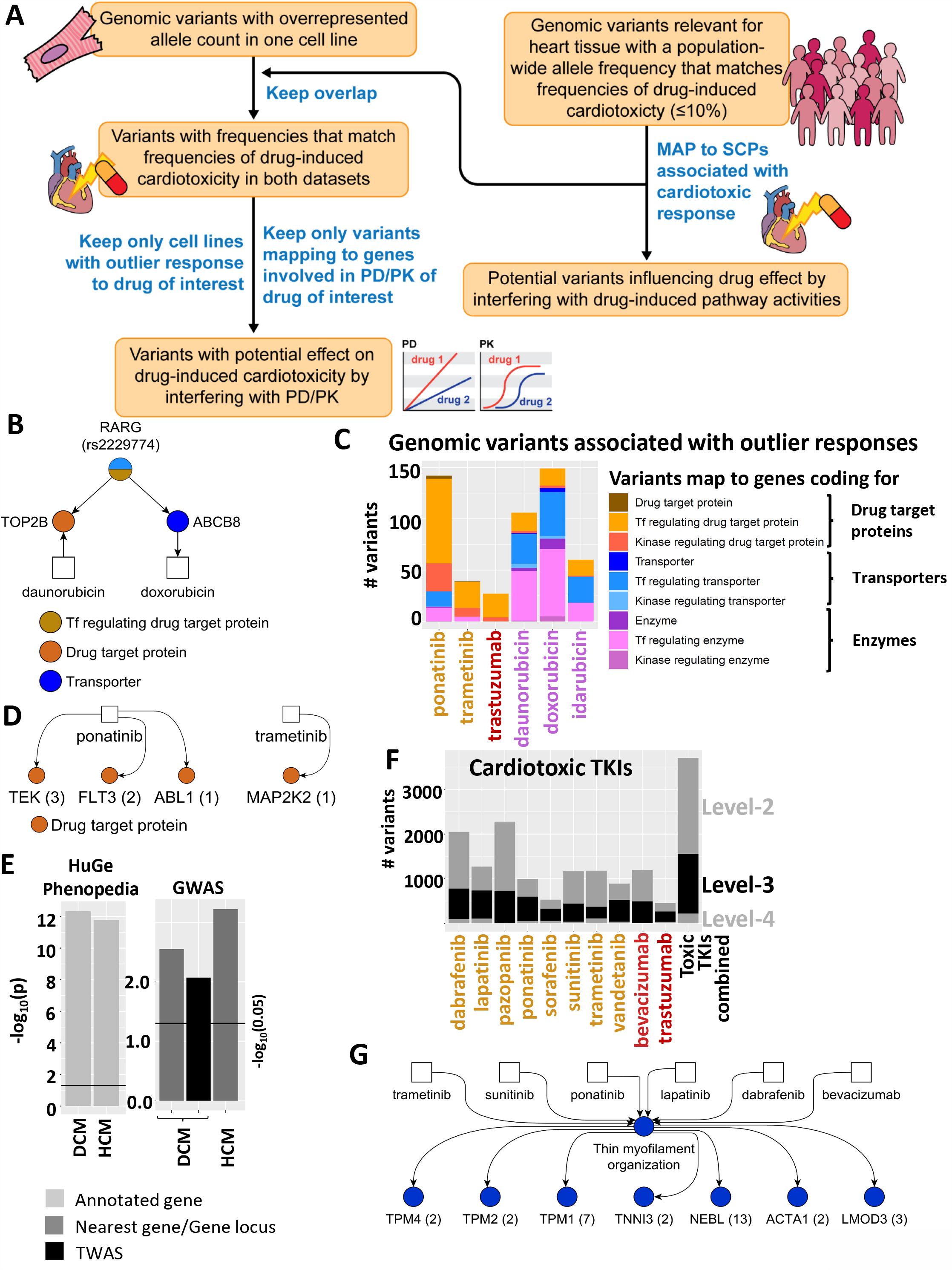
Identification of genomic variants that are potentially associated with a cardiotoxic response. Whole genome sequencing of our six cell lines ^16^ was used to identify alleles in our cell lines at known variant positions. **(A)** Pipeline for identification of potential genomic variants involved in PK/PD or induced SCP activities for a drug of interest. **(B)** The identified variant rs2229774 in cell line MSN09 maps to the coding region of the transcription factor RARG regulating the expression of TOP2B and ABCB8, both involved in PK/PD of daunorubicin and doxorubicin that induce an outlier response in this cell line. **(C)** In total, we identified 213 and 201 potential variants associated with TKI- or anthracycline-induced cardiotoxicity by interference with PK/PD mechanisms, respectively. Variants mapping to multiple gene classes are split equally among them to prevent double counting. **(D)** Numbers in brackets indicate how many variants map to individual drug target proteins of cardiotoxic TKIs. **(E)** We compared the overlap of identified SCP genes associated with a cardiotoxic or non-cardiotoxic response to genes associated with inherited DCM or HCM, either within the HuGe Phenopedia database or identified in GWAS. In the DCM, GWAS identified variants were associated with either the nearest genes or genes with the connected strongest cis-regulated expression as predicted by transcription-wide-association studies (TWAS). **(F)** Variants with a low frequency in the whole population that are part of e/sQTLs or lie within coding regions were mapped to up- or downregulated level -2, -3 and -4 SCPs that we predicted as indicative for TKI-induced cardiotoxicity. Variants that map to identified SCPs of multiple levels are only counted for the lowest level SCPs (higher level numbers) to prevent double counting. **(G)** Up- and downregulated SCPs associated with a cardiotoxic response were mapped back to the cardiotoxic TKIs that induce them. Numbers in brackets show identified variants for each SCP gene.

Variants of the first interference group should induce a different transcriptomic response, allowing us to link cell-line-specific variants to identified 24 outlier responses. Since, based on statistical likelihoods, either none or only one of our six healthy volunteers should suffer from cardiotoxicity induced by a drug of interest, we focused on variants that are overexpressed in an outlier cell line for a drug of interest, if compared to the other five lines. Mechanistic knowledge is added by considering only variants that map to genes with potential involvement in a drug’s PK/PD.

Variants of the second group should map to pathways that either contain drug target proteins or are induced or repressed by the drug of interest on the transcriptional level. Using the results of our transcriptomic analysis, we focused on the second group with a particular interest in those SCPs that are associated with a cardiotoxic or non-cardiotoxic response.

### Identification of genomic variants interfering with a drug’s PK/PD

Our PK/PD algorithm allowed re-identification of the variant *rs2229774* (Figure 4B, Suppl. Table 21) within the coding region of the *RARG* gene ^15^, one out of three genomic variants with the strongest evidence for anthracycline-induced cardiotoxicity (AIC) ^47^. The suppressive activity of the transcription factor RARG on the expression of the anthracycline target TOP2B is reduced by this variant, leading to increased AIC ^48^. Pathway enrichment analysis (Suppl. Figure 14D) suggests missing upregulation of WNT receptor signaling as a potential mechanism of *rs2229774* -triggered AIC (Suppl. Figures 14D, 26A). In agreement, inhibition or activation of WNT signaling enhances or mitigates AIC, respectively ^49-51^.

In total, our outlier-based screening approach identified 213 and 201 potential variants that map to 133 and 129 genes involved in the PK/PD of three anthracyclines and three cardiotoxic TKIs, respectively (Figure 4C, Suppl. Figure 26B, Suppl. Table 21). Identified variant candidates could trigger or protect from AIC or TIC. Seven of the variants map to drug target proteins of ponatinib and trametinib (Figure 4D), TKIs with medium (1-10%) and high (>10%) rates of TIC (Suppl. Table 4).

Considering all 24 drugs with documented outlier responses for this analysis we identified 847 variants (Suppl. Figure 26B) mapping to 378 genes (Suppl. Figure 26C) (Suppl. Table 21).

### SCP-based identification of genomic variants associated with TKI -induced cardiotoxicity

Identified level-2, -3 and -4 SCPs associated with the response to cardiotoxic and non-cardiotoxic TKIs enrich for genes that are mapped to inherited DCM or HCM, either within the HuGe Phenopedia database ^52^ or obtained from GWAS ^53,54^ (Figure 4E). We left out level-1 SCPs from this analysis, since they were predicted based on genes mapping to a sub function of the general cellular functions they describe. The significant overlap supports our observation that TKIs induce cardiotoxicity by mimicking mechanisms involved in inherited or acquired cardiomyopathy, as discussed above and in the supplemental information.

To identify a second set of potential variants interfering with TIC we mapped all variants underrepresented in the general population to identified SCPs. In total, we identified 2383, 1340 and 219 non-overlapping variants mapping to level-2, -3 and -4 SCPs that are up- or downregulated by cardiotoxic TKIs (Figure 4F). Mapping identified SCPs back to the drugs that induce them (Suppl. Figure 20) enables identification of variants that might interfere with cardiotoxicity of a drug of interest (Suppl. Table 22). For example, we predicted 31 variants mapping to seven genes of the SCP ‘Thin myofilament organization’ that might interfere with the cardiac response to sunitinib and ponatinib downregulating this SCP in six (enrichment ranks 2×3, 4, 2×7) and five (5×1) of six treated cell lines, respectively (Suppl. Figure 20), but also to lapatinib, dabrafenib and bevacizumab (Figure 4G). Four of those genes are involved in the same function that is targeted by mavacamten, as described above.

Most variants identified in GWAS that map to our SCPs are too frequent to meet our cutoff criteria that aim at identification of single variants triggering TIC in a monogenetic manner. Inherited HCM and DCM exist as both monogenic and polygenic diseases. Similarly, single or multiple variants might be related to TIC, requiring development of more sophisticated genomic algorithms. Nevertheless, identified SCPs are a good starting point to reduce the burden of multiple testing hypothesis by restricting the focus on genes with a functional implication in TIC.

## Discussion

A key question that arises in identifying transcriptional signatures from short term treatment of cell lines with drugs is whether early transcriptional events in a 1-2-day time frame can be predictive of later physiological (pathophysiological) states of an organ that takes weeks or months to manifest when a drug is used clinically. While it is not possible to answer this question for multiple organs and drug classes, in the case of TKI drugs and heart the answer is yes. Two types of relationships support this conclusion. We previously showed that transcriptomic signatures of TKI with differing toxicity profiles in FAERS could be associated with these profiles ^55^. In this study, the pathway analyses of drug-selective gene expression lists across cell lines show that many TKIs with adverse event propensity selectively regulate pathways that have also been shown to be involved in different types of cardiomyopathies ^56^. Additionally, the identity of the pathways themselves such as contraction dynamics and extracellular matrix are directly relevant to development of cardiac insufficiency. It is noteworthy that mavacamten, a drug approved in 2022 by the FDA to treat hypertrophic cardiomyopathy ^31^ regulates the level of actin-myosin bridge formation and thus controls contractile dynamics. Taken together, it appears reasonable to conclude that the pathways regulated by cardiotoxic drugs can serve as pathway signatures for cardiotoxicity. Such signatures could be used in drug development. Transcriptomic profiles of drug candidates could predict risk of cardiotoxicity from early cell-based study (Figure5A). Additionally, this approach can also be used to predict new drug targets that could mitigate TKI-induced cardiotoxicity (Figure 5B) by reversing relevant pathway activities ^57^.

**Figure 5.**
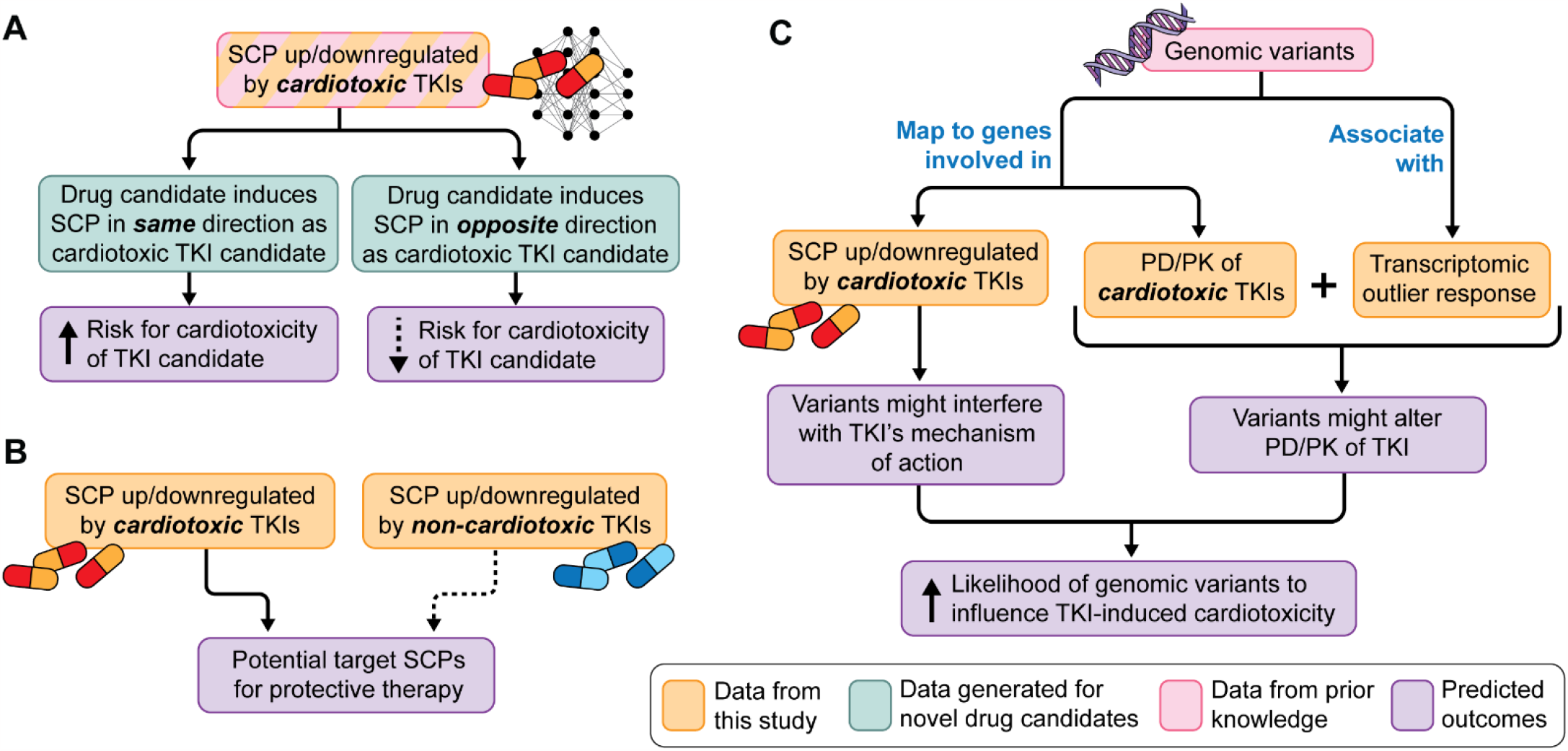
Potential use of cell based transcriptomic data for drug therapy induced adverse events. The flowcharts summarize the how integration of experimentally gathered transcriptomic data with publicly available pathway and genomic data bases can **(A**) help predict toxicity of drug candidates, (**B**) identify potential new drug targets to mitigate cardiotoxicity and (**C)** enable design of clinical studies to associate genomic variants with cardiotoxicity propensity.

Our method for identifying drug selective DEGs could have wider applications since both bulk and single cell expression profiles often represent the combination of multiple DEG sets induced by different actions on the same cell type or mixtures of cell types (i.e., tissue). Our computational pipeline decomposes these mixed gene expression profiles into individual effect-selective DEG sets and identifies the DEG sets involved in the response of interest. For example, grouping patients with the same disease into subgroups based on shared disease clinical manifestations can enable identification of gene expression profiles associated with each selected subgroup that can lead to tailored treatment options. A particular advantage of our pipeline is that patient groupings do not need to be mutually exclusive, i.e. patients can be assigned to each disease subgroup independently of the other assignments.

A serendipitous finding from our studies on the hiPSC derived cardiomyocytes is that single cell RNA-Seq allowed us to resolve these cells into multiple clusters ^16^. As single cell transcriptomic data from normal ^28^ and diseased ^13,14^ adult human heart became available in 2022 we compared the groups in our data to the different cell types in the adult human heart. This comparison allowed us to identify the group that is most like ventricular cardiomyocytes and to identify a group related to fibroblasts. Several of the top ranked SCPs are most likely expressed in multiple cell types, since they are not specifically enriched in any of them. Thus, the integration of bulk transcriptomic and single cell transcriptomic data allows us to project the SCPs onto different cell types of the heart. These integrated analyses suggest that the cardiotoxic drugs are likely to have their pathophysiological effects by affecting multiple cell types. The effects on fibroblasts and extracellular matrix dynamics ^58^ can readily affect the elasticity of the cardiac muscle and it has long been known that extracellular collagen-crosslinking is directly related to heart failure ^44,45^. It appears likely that the effects of TKIs on fibroblasts can contribute in part to cardiotoxic effects. In evaluating drug candidates for cardiotoxicity, it may be necessary to consider the drugs’ effects not only on cardiomyocytes but also on cardiac fibroblasts and other cell types in the heart.

Using the common differential expression patterns for each drug across cell lines, it is possible to identify outlier responses in a particular cell line (i.e., human subject). Such responses could provide information about genomic characteristics that could be associated with adverse events since these events occur in typically 1-20% of the population. Genomic variants associated with the proteins involved in PK/PD are likely candidates. We tested the validity of this assumption and found that a genomic variant associated with RARG can be identified by integration of whole genome wide sequencing of our six cell lines and outlier analysis of the anthracycline drugs. GWAS ^15^ and follow-up studies ^48^ had previously identified RARG to be causally linked to anthracycline-induced cardiotoxicity, since RARG regulates the expression of the anthracycline target topoisomerase 2B. Based on this validation we searched for genomic variants associated with cardiotoxic TKIs. We considered PK, PD, and key pathways regulated by these drugs and developed a list of variants whose occurrence resemble those of cardiotoxic events. Here data integration identifies genomic variants that could serve as hypotheses for targeted genomic studies to identify risk determinants in different populations (Figure 5C). Overall, our study shows that integration of transcriptomic and genomic data with various publicly available datasets can provide an integrated understanding and serve as a powerful hypothesis generator for different laboratory and clinic-based studies to detect and avoid adverse events associated with drug therapy.

## Data Availability

Transcriptomic and genomic data are available at NCBI GEO and dbGAP under the accession numbers GSE174773, GSE217421 and phs002088.v1.p1, respectively.

## Acknowledgement

This research was supported by a grant from NIH Common Fund 5U54HG008098 Drug Combination Signatures for Prediction and Mitigation of Toxicity as part of the LINCS program. We thank Jill Gregory for help with the flowchart figures.

