## Supplemental Materials for "Multiscale mapping of transcriptomic signatures for cardiotoxic drugs"

Jens Hansen,[1,2] Yuguang Xiong,[1,2] Priyanka Dhanan,[1,2] Bin Hu,[1,2] Arjun S. Yadaw,[1,2] Gomathi Jayaraman,[1,2] Rosa Tolentino,[1,2] Yibang Chen,[1,2] Kristin G. Beaumont,[3] Robert Sebra,[3] Dusica Vidovic,[4] Stephan C. Schürer,[4] Joseph Goldfarb,[1, 2] James Gallo,[1,2,5] Marc R. Birtwistle,[1,2,6] Eric A. Sobie,[1,2] Evren U. Azeloglu,[1,2,7] Seth Berger,[8] Angel Chan,[1,2,9] Christoph Schaniel,[1,10] Nicole C. Dubois, [1,11] Ravi Iyengar[1,2]

1 Mount Sinai Institute for Systems Biomedicine, Icahn School of Medicine at Mount Sinai, New York, NY 10029, USA
2 Department of Pharmacological Sciences, Icahn School of Medicine at Mount Sinai, New York, NY 10029, USA
3 Department of Genetics and Genomic Sciences, Icahn School of Medicine at Mount Sinai, New York, NY 10029, USA
4 Institute for Data Science and Computing, University of Miami, Coral Gables, FL 33146, USA
5 School of Pharmacy and Pharmaceutical Sciences, University of Buffalo SUNY System, Buffalo NY 14260
6 Chemical and Biomolecular Engineering, Clemson University, Clemson, SC, 29634, USA
7 Department of Medicine, Division of Nephrology, Icahn School of Medicine at Mount Sinai, New, York, NY 10029, USA
8 Departments of Pediatrics and Genomics & Precision Medicine, George Washington University School of medicine and Health Sciences, Washington DC USA, 20012
9 Cardiology Division, Department of Medicine, Memorial Sloan Kettering Cancer Center New York NY 10065
10 Department of Medicine, Division of Hematology and Medical Oncology, Tisch Cancer Institute, Icahn School of Medicine at Mount Sinai, New York, NY 10029, USA
11 Department of Cell, Developmental and Regenerative Biology, Icahn School of Medicine at Mount Sinai, New York, NY 10029, USA

Address correspondence to

Jens Hansen –
or
Ravi Iyengar -

### Experimental Methods

**Materials** The sources of all materials are listed in Supplemental Table 3 that is taken from Schaniel et al. [16], since the two studies were conducted concurrently. It is also available as detailed Protocols associated with the LINCS project.

**Healthy human subject iPSC lines.** Skin fibroblasts were obtained from six healthy volunteers, dedifferentiated into human induced pluripotent stem cells (hIPSC) and differentiated into cardiomyocytes, as described previously [16]. Descriptions of the cell lines used in this study are provided in Supplemental Table 1.

**Drug treatment and Bulk Transcriptomics.** Bulk transcriptomic data of untreated control iPSC-derived cardiomyocytes has been published previously [16] (see Suppl. Figure S4B in our previous publication for a PCA plot) and was deposited at NCBI GEO under the accession number GSE174773. Here, we stimulated six cardiomyocyte cell lines with 54 different single drugs for 48 hours. The sources and concentrations of the drugs used for the treatments are in Supplemental Table 3. Typically, each cell line/drug combination had quadruplicate measurements. After treatment, for each replicate, cells were harvested, and RNA was extracted. Only RNA cell line/drug combinations with RIN> 8 were used for library preparation. Details have been previously described [16]. Pooled libraries were clustered in either an S1 flowcell or a full NovaSeq 6000. The cell line/drug combinations were then sequenced using a 1x100bp Single End configuration, and typically 24 million reads were mapped to the reference genome for each cell line/drug combination.

### Computational Methods

#### Curation of cardiotoxic risk profiles from FAERS database

We curated data from the FDA adverse event reporting system (FAERS) database to calculate reporting odds ratios of drugs that act on cardiac systems. We downloaded data from the FDA website from (2004_Q1 to 2019_Q4) which contains over 10 million adverse event reports in AERS (2004 _Q1 to 2012_Q3) & FAERS (2012_Q4 to 2019_Q4). The quarterly data files include the following: a) demographic and administrative information including imaging reports, b) each patient identified by an id number, c) drug information from the case report, d) adverse event information from the report, f) patient outcome information from the report, g) information on the source of the report, h) information on the date of report. To parse FAERS (and AERS) data into a relational database (by default it is sqlite), we downloaded the ASCII zip file for each quarter and used the FAERS toolkit (https://github.com/kylechua/faers-toolkit). For further analysis, we used the python program to parse raw data from text files and identify the frequency of occurrence of adverse events of interest. Finally, we used a reporting odds ratios (mean and 95% confidence interval) formula for each drug of interest.

#### Identification of differentially expressed genes

Control transcriptomic data [16] and transcriptomic data generated by treatment of six cardiomyocyte cell lines with 54 drugs was analyzed, as described previously [59], except that we used human reference genome hg38 in this study. Briefly, we aligned the bulk transcriptomic data to hg38 using STAR [60], followed by mapping of read counts to genes using the RefGene reference annotation and feature counts [61]. Pairwise correlation, followed by hierarchical clustering, identified replicates with deviating gene expression profiles that were consequently removed from the analysis. Only drug-treated or control cell lines with at least three remaining replicates were subjected to differential expression analysis using edgeR [62], yielding 266 lists of differentially expressed genes, each describing the response observed in a unique cell line/drug combination. Genes with a False Discovery Rate (FDR) of at max 10% were defined as significant. Each of these lists contained significance p-values for 16,345 genes, ranging from zero to one. We transformed these p-values into $-\log_{10}$(p-values) and added a negative sign in case of downregulated genes (or kept the positive sign in case of upregulated genes), to document the direction of change. P-values that were zero were replaced by $10^{-64}$ before transformation, since this is the lowest non-zero p-value that is generated by our edgeR pipeline.

#### Pairwise correlation and hierarchical clustering

All generated signed $-\log_{10}$(p-values) were integrated into a matrix where the columns represent the 266 treated cell line/drug combinations and the rows the 16,345 genes. We calculated the pairwise correlation coefficients between all rows and all columns, followed by hierarchical clustering of the rows and columns based on the pairwise correlation coefficients (distance function: (1-cor(x)) / 2). Rows and columns of the original matrix were rearranged based on the clustering results before visualization. To document how efficient the clustering algorithm groups cell line/drug combinations that represented different cell lines treated with the same drug into the same cluster, we calculated an F1 score for each cluster drug combination. Every cluster that could be obtained by cutting the dendrogram at any height and contained at least two cell line/drug combinations was analyzed. The F1 score is the harmonic mean of precision and recall (we selected beta = 1). Here, precision documents the fraction of cell line/drug combinations in a particular cluster that were treated with a particular drug, recall documents the fraction of all cell line/drug combinations treated with that drug that were in that cluster. For each drug, we determined the cluster with the highest F1 score and saved that F1 score for that drug.

**Singular value decomposition and identification of drug-selective subspaces and gene expression profiles**

The gene expression matrix filled with the signed $-\log_{10}$(p-values) was subjected to our computational pipeline for the identification of drug-selective responses (Suppl. Figure 4). It is based on singular value decomposition (SVD) and implemented in the programming language R. SVD decomposes the input matrix into a matrix of orthonormal left singular vectors, a diagonal matrix of singular values and a matrix of orthonormal right singular vectors. Applied to gene expression data, the left singular vectors are often referred to as eigenarrays [63], because each gene expression profile (columns in the input matrix) is a linear combination of all eigenarrays (Suppl. Figure 6A). The right singular vectors contain the cell line/drug combination-specific coefficients for this linear combination. The eigenarray-specific singular values document how much of the initial variance can be explained by each eigenarray and need to be added as additional factors to the terms in the linear combination to reconstruct the original values.

To further characterize the eigenarrays, we correlated the cell line/drug combination-specific coefficients associated with each eigenarray with the number of significantly differentially expressed genes (FDR ≤ 0.1) in each cell line/drug combination, using Pearson correlation (Suppl. Figure 6C). The first eigenarray was removed from the gene expression matrix and the new matrix subjected to the same pipeline. We used a two-tailed t-test to analyze whether the cell line/drug combination-specific coefficients of each eigenarray significantly differed between all cell line/drug combinations of the same drug and all other cell line/drug combinations (Suppl. Figure 9A). Similarly, we analyzed whether the cell line/drug combination-specific coefficients of each eigenarray significantly differed between all cell line/drug combinations of the same cell line and all other cell line/drug combinations. As a result, we obtained one p-value for each drug and eigenarray combination or cell line and eigenarray combination. P-values were transformed into $-\log_{10}$(p-values) and integrated into a matrix where the columns represent the eigenarrays and the rows the drugs or cell lines. The matrix was subjected to pairwise correlation analysis and hierarchical clustering as described above.

Any combination of eigenarrays spans a subspace that contains a particular fraction of the complete DEG profiles. For each drug, we ranked all eigenarrays by significance and generated 263 potential drug-selective subspaces spanned by the top three to 265 ranked eigenarrays. We then projected all 266 full gene expression profiles into each of these subspaces, followed by pairwise correlation analysis and hierarchical clustering. Each potential drug-selective subspace can be characterized by two different measures. The first measure is the maximum F1 score that documents how close all cell line/drug combinations treated with the drug of interest cluster together. The second measure is the median cosine similarity that documents how much of the gene expression profiles, after removal of the first eigenarray induced by the drug of interest, is still preserved in that subspace. The maximum F1 score was determined as described above for the complete dataset. For each subspace we calculated a final selection score that is the weighted mean between the maximum F1 score and median cosine similarity. The relative contribution of the F1 score and the median cosine similarity to the final selection score was defined by the F1 score weight. High and low F1 score weights favor F1 score or median cosine similarity, respectively. Varying the F1 score weights from 0.00 to 0.95 in steps of 0.05 and searching for the subspace with the highest selection score yielded 20 potential drug-selective subspaces for each drug.

Besides our interest in the characterization of drug-selective gene expression profiles, we were also interested in the identification of cell lines that showed a significantly different response from the other cell lines to treatment with the same drug. To identify cell lines with such outlier responses we screened each of the 20 potential drug-selective subspaces for clustering results that indicated an outlier response, i.e., a single cell line that does not cluster together with the other cell lines treated with the same drug. To find such outlier responses, we calculated cell line/drug combination-specific F1 scores using the same approach described above, except that the cell line/drug combination of interest has to be in that cluster. Dixon's Q test of cell line/drug combination-specific F1 scores was used to identify outlier responses (adj. p-value ≤ 0.05). Identified outliers were only accepted if the mean F1 score of all non-outlier cell line/drug combinations was larger than 0.5. If an outlier response was identified in any of the 20 potential drug-selective subspaces, we ranked the potential subspaces by Dixon's Q test adjusted p-value, followed by decreasing mean F1 score of all non-outlier cell line/drug combinations and increasing F1 score weight. The top ranked subspace was defined as the drug-selective subspace. If no outlier was identified, we selected that subspace with the highest selection score based on an F1 score weight of 0.95.

Projection of gene expression profiles into the final drug-selective subspaces generated drug-selective gene expression profiles that were combined into a new matrix.

### Pathway enrichment analysis

We subjected the complete set of DEGs, the DEGs after removal of the first eigenarray and the drug-selective DEGs to pathway enrichment analysis (Suppl. Figure 14A). For each group of DEGs, up- and downregulated genes among the top 600 most significant DEGs (as indicated by complete or decomposed absolute $-\log_{10}$(p-values)) were separately subjected to enrichment analysis using the Molecular Biology of the Cell Ontology (MBCO) [64] (www.mbc-ontology.org) (https://github.com/SBCNY/Molecular-Biology-of-the-Cell) and Fisher's Exact test. Only SCPs predicted with a maximum p-value of 0.05 were considered. For each treatment, predicted up- or downregulated level-1, -2, -3 and -4 SCPs were separately ranked by significance, generating eight lists of ranked SCPs per treatment. Pathway enrichment analysis was implemented in the programming language C#.

#### Identification of SCPs associated with cardiotoxic and non-cardiotoxic responses

To identify SCPs that are associated with cardiotoxic and non-cardiotoxic TKI responses, we counted, at each significance rank cutoff, how many cardiotoxic or non-cardiotoxic kinase inhibitors and monoclonal antibodies up- or downregulate an SCP of interest with a significance rank below or equal to the current cutoff (Suppl. Figure 16). Counting results were used to calculate precision and recall of each SCP at each rank to be either up- or downregulated by either the cardiotoxic or non-cardiotoxic TKIs. F1 scores were calculated with a high emphasis on the precision (beta = 0.25). To ensure consistent results across multiple ranks, we calculated the area under the F1 score curve (AUC) between the ranks 1 and 20 for the levels 1, 2 and 4 and 1 and 30 for SCPs of level 3. Different rank cutoffs were selected due to the different number of SCPs of each level. Consequently, we obtained four AUCs for each SCP (up- or downregulated by cardiotoxic or non-cardiotoxic TKIs). To enable cross-comparison of AUCs across different levels, we normalized the AUCs by calculation of the percent of the maximal available area covered by each AUC. To emphasize identification of SCPs that are either up- or downregulated by cardiotoxic or non-cardiotoxic TKIs, we introduced a penalty on each AUC by subtracting half of the AUC calculated for the same group but based on the DEGs changing in the opposite direction. Final AUCs of the same toxicity group were ranked (independently of the direction of change) and the level-1, -2, -3 and -4 SCPs associated with the top 10, 10, 25 and 10 ranked AUCs further investigated. Different rank cutoffs were selected due to the different number of SCPs of each level. This pipeline was implemented in C# and R.

#### Single cell and nucleus RNAseq dataset analysis

Single cell RNASeq data of four hiPSC-derived cell lines that were also used in this study were obtained from Schaniel et al. [16] (GSE175761). We reanalyzed the data using an updated Seurat package (version 4.1.3) [65]. Cells with feature counts below 500 and expression of mitochondrial genes of more than 50% were removed during quality control. The Seurat functionality 'SCTransform' was used to normalize, scale, identify the top 2000 features, and regress out mitochondrial gene counts in each individual dataset. The four datasets were integrated using the Seurat 'IntegrateData' pipeline. We used the top 30 principal components for dimensionality reduction. After identification of cell neighborhoods using 'FindNeighbors' we clustered the combined integrated Seurat object using a resolution of 0.085, since this resolution allowed identification of six different clusters that can be mapped to reasonable cardiac cell types. Cluster marker genes were calculated using ‘FindMarkers’ and the ‘RNA’-assay and log-normalized ‘data’-slot, using a significance cutoff of an adjusted p-value ≤ 0.05. To annotate clusters to cell types we subjected the top 500 significant marker genes to pathway enrichment analysis using Fisher’s Exact test and cell type genes identified in single cell and single nucleus RNAseq of the adult [28] or the developing human heart [66]. To identify cluster-specific SCPs we subjected the marker genes to enrichment analysis using MBCO [64].

Suppl. Figure 22A shows UMAPs with cluster-cell type annotations that are similar to our previous annotations published Suppl. Figure S4D in Schaniel et al [16]. Suppl. Figure 22B shows a similar distribution of cells among the different clusters in each of the four cell lines as Suppl. Figure S4E in our previous publication. Cell type annotations in current Suppl. Figure 22A match those in our previous publication shown in Suppl. Figure S4G[16].

Cell type marker genes in the adult human heart [28] used above were calculated using a published Seurat object and FindMarkers as described above. The top 500 marker genes of each cell type were subjected to enrichment analysis using MBCO.

Single cell RNAseq of hiPSC-derived cardiomyocytes from patients with dilated cardiomyopathy and healthy controls was obtained from NCBI GEO (GSE184899). Cells with feature counts below 400 or above 9000 and expression of mitochondrial genes of more than 50% were removed during quality control. Cells were processed as described above. The Seurat functionality ‘FindMarkers’ of the Seurat package was used to calculate pseudo-bulk DEGs between DCM and healthy cardiomyocytes using the log-normalized RNA assay data slot. Up- and downregulated genes among the top 600 most significant DEGs with a maximum adjusted p-value of 0.05 were subjected to enrichment analysis using MBCO.

Up- and downregulated genes among the top 600 most significant DEGs in each cell type obtained by single cell [13] or single nucleus RNAseq [14] from patients with dilated and/or hypertrophic cardiomyopathy were subjected to enrichment analysis using MBCO as well.

### Whole genome sequencing

Whole genome sequencing of the six cell lines used in this study has been previously published [16] and is deposited on dbGAP under the accession id phs002088.v2.p1. Here, we reanalyzed the data as described previously [16], except that we used the hg38 reference genome. Sequencing reads were aligned to hg38 reference using bwa-mem2. Aligned bam-files were called to g.vcf files using deepvariant and a joint vcf of calls were generated using GLnexus. Jointly the vcf file was annotated using ANNOVAR to provide RefSeq gene-based annotations, gnomAD allele frequencies, CADD 1.6 scores, SpliceAI scores, ClinVar annotations, dbSNP annotations, GTEx annotations for atrial and ventricular expression and splice QTLs downloaded from the GTEx portal. Variants reported in the NHGRI-EBI Catalog of human genome-wide association studies were annotated.

### Identification of genomic variants potentially associated with drug-induced cardiotoxicity

Building on our transcriptomic analysis, we implemented a pipeline in C# and R that searches for potential variants of drug-induced cardiotoxicity (Suppl. Figure 25). We searched for variants that either interfere with a drug's pharmacokinetics (PK) or -dynamics (PD), or map to SCPs identified to be associated with a cardiotoxic or noncardiotoxic response by our transcriptomic analysis.
In both cases, we initially filtered variants based on population-wide statistics. Cardiotoxic responses to kinase inhibitor treatment are normally observed in 1% to 20% of the treated patients (Suppl. Table 4). Consequently, the population-wide frequency of an allele in a genomic variant associated with drug-induced cardiotoxicity should be in a similar range or even lower, so we focused on variant alleles with frequencies of 10% or less. In addition, we assumed that the least frequent allele, minor allele in the following, should be the most likely allele associated with a deviating, i.e., cardiotoxic (or cardioprotective) response. Relevance for cardiac tissue should be assigned for those variants that either map to gene coding regions or are part of a cis-expression (e) or -splicing (s) QTL in the adult heart that we obtained from the GTEx Portal (GTEX v8) [46]. For further analysis, we only considered variants that met these filter criteria.

For the identification of variants that interfere with PK/PD of a given drug we integrated the results of our transcriptomic outlier analysis with the WGS results of our cell lines. With cardiotoxic responses in 1- 20% of treated patients (Suppl. Table 4), either none or only one of our six healthy volunteers that donated skin fibroblasts for hiPSC generation and cardiomyocyte differentiation should suffer from cardiotoxicity induced by a drug of interest. Consequently, we considered only those variants in each cell line that show a higher count for the minor allele (as identified in the population-wide analysis) in that cell line compared to the other five cell lines. For the last selection step, we hypothesized that interference with the drug's PK/PD induces a different transcriptomic response, allowing us to link drug-related outlier responses identified in our transcriptomic analysis to variants overrepresented in our cell lines. To link the remaining variants to a drug's PK/PD, we curated all human drug target proteins, transporters and enzymes of drugs with an identified outlier response from the Drugbank database [21]. In addition to the genes directly involved in each drug's PK/PD, we were also interested in mechanistic regulators of those genes, i.e. transcription factors or kinases. We downloaded the libraries 'Chea_2016' [67], 'Encode_tf_chip_seq_2015', 'Transfac_and_jaspar_pwms', 'TRRUST_transcription_factors_2019', 'Kea_2015' [68] from the enrichR website [69] and curated all transcription factors and kinases that regulate any target proteins, transporters or enzymes. Only variants mapping to genes involved in a drug's PK/PD were suggested as potential variants interfering with the PK/PD of a drug of interest.

For the identification of variants with cardiotoxic or cardioprotective effects by interference with transcriptionally regulated SCPs that are associated with a cardiotoxic or non-cardiotoxic response, we mapped all variants identified in the population-wide analysis to identified SCPs. To prevent double counting we removed any variants mapping to genes that are annotated to multiple SCPs in parent-child relationships from all ancestor SCPs before visualizing the results as stacked bar diagrams.

**Comparison of identified variant genes with prior knowledge**

To compare our predictions with genes involved in inherited cardiomyopathies, we extracted all genes annotated to "Cardiomyopathy, Hypertrophic" or "Cardiomyopathy, Dilated" in the HuGE Phenopedia [52] (downloaded on 2020 June 04) and genes associated with DCM [53] or HCM [54] in GWAS. In the DCM study [53], we focused on those genes annotated to the MRI phenotypes 'LVEF', 'LVESV' and 'LVESVi', since DCM was the disease most strongly associated with these phenotypes, as stated by the authors. We compared both the genes listed in the column 'Nearest Gene' and 'TWAS Gene of Suppl. data file 3 with our results. In case of the HCM [54] study, we used all genes listed in the column 'Locus Name' in their Suppl. Table 2.

We merged all predicted genes that contain potential variants and map to level-2, -3 and -4 SCPs of the MBCO ontology [64] identified to be up- or downregulated by cardiotoxic or non-cardiotoxic drugs. Here, we ignored genes annotated to level-1 SCPs, since level-1 SCPs cover multiple subfunctions only a few of which might contribute to the prediction of the level-1 SCPs by our algorithm. Consequently, level-1 SCPs might contain genes that are unrelated to the subfunctions that enabled their prediction. Since our algorithm can only identify genes annotated to any level-2, -3 and -4 SCPs, we selected those genes as the background set of genes for a Fisher's exact test. For statistical accuracy, we removed all genes associated with inherited DCM and HCM in the datasets described above that were not part of the background gene list. A right-tailed Fisher's exact test was used to calculate the significance of the overlap between our genes and the published gene lists.

### Supplemental Text

#### Drug-selective gene expression profiles allow for identification of SCPs which agree with prior knowledge of SCPs associated with cardiac diseases and drug effects

Up- and downregulated genes in each cell line/drug combination were subjected to enrichment analysis using the Molecular Biology of the Cell Ontology (MBCO) [64]. Predicted subcellular processes (SCPs) were ranked by decreasing significance for each list of genes and SCP level. We will refer to these ranks as enrichment ranks.

As outlined in the main text, the SVD-based identification of drug-selective gene expression profiles greatly increased the number of predicted SCPs that are up- or downregulated in at least two of three, three of four, four of five or four of six cell lines (i.e., ≥ 66%) by the same drugs (Suppl. Figure 15A). The median overlap between downregulated level-1, -2, -3, -4 SCPs among the top 5, 5, 10 and 5 predicted SCPs increased from 2 to 3.5, 1 to 3, 1 to 4 and 0 to 1, respectively. The median overlap between upregulated level-1, -2, -3, -4 SCPs among the top 5, 5, 10 and 5 predicted SCPs increased from 1 to 3, 0 to 3, 1 to 4 and 0 to 1, respectively. These results depend on the complete SVD-pipeline, since removal of the first eigenarray alone improved the median number of drug-selective overlapping up- or downregulated SCPs in only one case (upregulated level-2 SCP, from 0 to 1).

Analysis of drug-selective gene expression profiles enabled identification of enrichment patterns that were obscured within the complete gene expression profiles. For example, our decomposition pipeline allowed identification of SCPs that were specifically downregulated by the group of anthracyclines that have high rates of cardiotoxicity [70]. These SCPs are involved in iron metabolism and single protein turn-over (Suppl. Figure 14/B/C/D/E). They form parent-child relationships within three MBCO branches (Suppl. Figure 15B). The observed downregulation of 'Cellular iron storage' is in agreement with the suggested interference of anthracyclines with iron metabolism [71] and ferroptosis as a disease mechanism involved in heart failure [36]. The clinical relevance of our unbiased findings from transcriptomic data is documented by supportive therapy with iron chelators [40]. Identified down-regulation of SCPs involved in protein degradation might explain cardiotoxicity by interference with sarcomere turn-over [29], as supported by the cardiotoxic effects of proteasome inhibitors [72]. The almost exclusive downregulation of these SCPs by anthracyclines could not be documented using the complete gene expression profiles (Suppl. Figure 15C).

SCPs predicted from the drug-selective gene expression profiles of other drugs often describe functions that are supported by prior knowledge as well, based on transcriptomic and orthogonal methodologies.

For example, enrichment analysis of nilotinib-selective gene expression profiles leads to a more consistent prediction of upregulated SCPs involved in centrosome and mitotic spindle dynamics as well as mitosis compared to analysis of nilotinib's complete gene expression profiles (Suppl. Figure 14B/C/D). Similar results were obtained for the kinase inhibitors imatinib, ponatinib, sorafenib and regorafenib. In agreement, previous morphological analysis documents centrosome aberrations and mitotic spindle defects induced by treatment of primary human fibroblasts with imatinib or nilotinib [73]. Centrosome defects were also observed in disease-unaffected cells from patients treated with imatinib, nilotinib, sorafenib, sunitinib, dasatinib and bosutinib [74].

There are two MBCO level-3 SCPs that refer to increased cholesterol synthesis activity, 'Cholesterol synthesis' and 'Cholesterol-sensitive control of SREBP activation'. Though multiple drugs upregulate these SCPs at varying ranks, analysis of lapatinib-selective gene expression profiles predicts upregulation of both SCPs at top enrichment ranks (5x rank 1 and ranks 2, 3, 4, 5, 6, respectively) (Suppl. Figure 14D). In agreement, mevalonate pathway activity was found to be increased in lapatinib-resistant and lapatinib + trastuzumab–resistant cells [75]. Resistance was reversed by statin treatment that inhibits cholesterol synthesis.

**SCPs associated with cardiotoxic and non-cardiotoxic responses to TKI treatment describe cellular dysfunctions involved in cardiomyopathy development**

In the main text, we briefly described SCPs involved in muscle contraction and sarcomere renewal, energy metabolism and ferroptosis. Here we discuss identified and additional pathways in more detail.

Our F1 score and AUC statistics generate lists of SCPs that are up- or downregulated by cardiotoxic or non-cardiotoxic TKIs. Within each toxicity group combined up- and downregulated SCPs were ranked by decreasing AUC. We will refer to these ranks as AUC ranks to distinguish them from the enrichment ranks that are described above.

As already indicated in the main results section, we distinguish between SCPs whose activity for an association with a cardiotoxic response reaches sufficient levels after TKI treatment or is already sufficient at baseline levels as evidenced by a response to non-cardiotoxic drugs compared to cardiotoxic drugs. The first set of SCPs was identified by screening for those SCPs that are up- or downregulated at higher ranks by cardiotoxic TKIs, as compared to non-cardiotoxic TKIs. Treatment with a cardiotoxic TKI might change the SCP activity beyond a threshold that separates association or non-association with a cardiotoxic response. Treatment with a non-cardiotoxic TKI might move the SCP activity across the threshold in the other direction, changing its status from sufficient to insufficient for association with a cardiotoxic response. Both sets of SCPs can be organized based on the SCP activity that favors a cardiotoxic response. SCPs upregulated by cardiotoxic TKIs and downregulated by non-cardiotoxic TKIs are SCPs whose higher activity favors a cardiotoxic response. In contrast, SCPs downregulated by cardiotoxic TKIs and upregulated by non-cardiotoxic TKIs are SCPs whose lower activity favors a cardiotoxic response. We use this classification in comparing the inferred SCPs from this data set to previously published data, since presence or absence of sufficient SCP activity is associated with presence or absence of cardiomyopathy development. We followed this principle in our figures as well. For SCPs that are upregulated by cardiotoxic and downregulated by non-cardiotoxic TKIs we use the colors red and orange, respectively (Suppl. Figures 17/18). For SCPs downregulated by cardiotoxic and upregulated by non-cardiotoxic TKIs we use the colors blue and light blue, respectively. The same color scheme was applied in our network-based visualization that integrates identified SCPs into the MBCO parent-child hierarchy and groups SCPs participating in similar functions into the same module (Suppl. Figure 19).

For both TKI groups we focused on the top 10 level-1, 10 level-2, 25 level-3 and 10 level-4 SCPs based on increasing AUC ranks (Suppl. Figure 18). Six level-1, two level-2, eleven level-3 and four level-4 SCPs were predicted to be relevant for cardiotoxicity and regulated by both TKI groups in the opposite direction. For those SCPs, both TKI groups link the same SCP activity levels (i.e., higher or lower) to be associated with a cardiotoxic response. This approach was mostly successful. We only obtained conflicting results for one level-4 SCP that was downregulated by both TKI groups.

Identified SCPs were mapped back to the TKIs that increase or decrease their activity them. To document which TKIs contributed to the identification of an SCP we show which cardiotoxic (Suppl. Figures 20) or non-cardiotoxic (Suppl. Figures 21) TKIs up- or downregulate an identified SCP at specified enrichment ranks.

**Sarcomere dynamics**

SCPs involved in muscle contractility were discussed in the main section. Here we want to add some more details. Identified level-1 SCP ‘Cellular contraction’, its level-3 grandchild SCPs ‘Thin myofilament organization’, ‘Myofibril formation’ and the level-4 SCPs ‘Actin filament depolymerization’ and ‘Myoglobin synthesis’ (Suppl. Figure 19), are all involved in dynamics and turnover of the sarcomere, as already described in the main results section. The identified SCPs whose lower activities favor a cardiotoxic response, belong to those SCPs that were identified with evidence for both the cardiotoxic and non-cardiotoxic group: four of five SCPs were downregulated by cardiotoxic TKIs and at the same time upregulated by non-cardiotoxic TKIs (Suppl. Figure 18). In addition, the level-2 SCP ‘Myofibril formation and organization’, a child of ‘Cellular contraction’ is identified (Suppl. Figure 19), because it is downregulated by the non-cardiotoxic TKIs (Suppl. Figure 18). Overall, all cardiotoxic TKIs, except vandetinib and trastuzumab downregulate

(Suppl. Figure 20), while 13 of 17 non-cardiotoxic TKIs upregulate (Suppl. Figure 21) at least one of the contractility-related SCPs in one to six or one to five cell lines, respectively. Four of six SCP genes of the SCP ‘Thin myofilament organization’ that are inhibited by cardiotoxic and induced by noncardiotoxic TKIs are components of tropomyosin or inhibitory troponin. Both complexes interact to block the binding of the myosin head to the thin myofilament during muscle contraction. This mechanism is also targeted by the new drug mavacamten that was recently approved by the FDA to treat heart failure [31], supporting the clinical relevance of our findings.

**Electric transmission**

Four cardiotoxic TKIs, trametinib, trastuzumab, lapatinib and bevacizumab upregulate ‘Potassium transmembrane transport’ in three of four (enrichment ranks 2, 3, 9), two of three (7, 10), two of five (7, 10) and one of four (17) cell lines (Suppl. Figure 20), respectively. The SCP with an AUC rank of 7 (Figure 2B, Suppl. Figure 18) is predicted based on potassium channels, transporters and components of the sodium potassium ATPase (Suppl. Table 12). Among the SCP genes is the regulatory SUR2A subunit of the cardiac ATP-sensitive potassium channel involved in genetic DCM [76].

The level-3 SCP ‘Gap junction organization’ is predicted with an AUC rank of 24 for the cardiotoxic TKIs (Figure 2B, Suppl. Figure 18). It is downregulated by the cardiotoxic TKI dabrafenib in one of five (enrichment rank 24), pazopanib in two of six (1, 3), ponatinib in three of six (15, 16, 17), vandetinib in one of three (9) and vevacizumab in four of four (13, 2x14, 15) treated cell lines (Suppl. Figure 20). In agreement, multiple connexins that form the building block of gap junctions are downregulated in explanted heart from patients with idiopathic cardiomyopathy [77].

**Energy metabolism**

We identified multiple SCPs that suggest interference with cardiac energy metabolism as a major trigger for TKI-induced cardiotoxicity.

The level-2 SCPs ‘Mitochondrial energy production’, ‘Fatty acid metabolism’, ‘Carbohydrate metabolism and transport’ and ‘Post-translational protein modification in Mitochondria’ were upregulated by cardiotoxic TKIs with the AUC ranks one to four, respectively (Suppl. Figure 18). Their upregulated level-3 child SCPs ‘Citric acid cycle’ and ‘Desaturation of fatty acids’ were predicted with AUC ranks 1 and 11 (Suppl. Figures 18, 19). Three of these level-2 and -3 SCPs map to ventricular cardiomyocytes in the adult heart (Figure 3B, Suppl. Figure 22) as the cell type with the highest energy requirement. These SCPs that interact with each other to ensure energy supply were almost exclusively predicted based on their induction by pazopanib (Suppl. Figure 20), a TKI with a high rate of cardiotoxicity (>10%) (Suppl. Table 4). Genes of these SCPs induced by pazopanib are involved in the citric acid cycle and oxidative phosphorylation, fatty acid activation, elongation and desaturation as well as glucose import, release from glycogen and degradation (Suppl. Table 12).

Many studies document an overall reduction in oxidative phosphorylation during heart failure [32], in agreement with the enrichment results obtained in adult and hiPSC-derived DCM and adult HCM cardiomyocytes (Figure 3C, Suppl. Figure 24). Nevertheless, compensatory upregulation of oxidative phosphorylation was suggested for a large DCM subgroup that is caused by truncating titin variants [33-35]. Another prominent molecular phenotype observed in these studies is a shift from fatty acid oxidation towards glycolysis. Upregulated genes mapping to carbohydrate catabolism and fatty acid anabolism might support such interpretation of our data as well, though the level-3 and -4 child SCPs that specifically describe these functions, were not among the top predictions. For the noncardiotoxic TKIs, we predict the level-2 SCP ‘Carbohydrate metabolism and transport’ and its level-3 child ‘Glycolysis and Gluconeogenesis’ at ranks 1 and 25 (Suppl. Figure 18) based on a set of partly overlapping genes (Suppl. Table 12), further supporting glucose utilization as a cellular function whose higher activity favors a cardiotoxic response to TKI-treatment.

Polyunsaturated fatty acids (PUFAs) generated by genes mapping to the upregulated SCP “Fatty acid desaturation” **[37,38]** can be deoxygenized by lipoxygenases, resulting in cardiotoxic PUVA hydroperoxides [36]. PUVA hydroperoxides are a main stimulator of ferroptosis, a potentially major mechanism involved in heart failure.

The involvement of ferroptosis in drug-induced cardiomyopathy was also predicted for the cardiotoxic anthracyclines based on the downregulation of ‘Cellular iron storage’ (Suppl. Figures 14D, 15B), as discussed above. However, upregulated transporter activities involved in import of iron-containing molecules, such as transferrin or haptoglobin-hemoglobin complexes, are predicted as part of the level-3 SCP ‘Cellular iron uptake and export’ for the non-cardiotoxic TKIs with AUC rank 23 (Suppl. Figure 18). These seemingly contradictory results could indicate that it is the balance in cellular iron content rather than an increase or decrease of iron levels that is associated with cardiotoxicity.

The level-3 SCP ‘Serine and glycine metabolism’ was identified for cardiotoxic and noncardiotoxic TKIs as an SCP whose higher activity indicates a cardiotoxic response with the AUC ranks 19 and 5, respectively (Suppl. Figure 18). Seven cardiotoxic (Suppl. Figure 20) and twelve non-cardiotoxic (Suppl. Figure 21) TKIs up- and downregulate this SCP, respectively, in one to six cell lines. Dabrafenib, a TKI with a cardiotoxicity frequency between 1 and 10%, upregulated this SCP in all five cell lines with top enrichment ranks (3x1, 2, 3), followed by pazopanib (ranks 2x2, 3, 4, 5) (Suppl. Figure 20). Identified SCP genes are involved in serine and glycine biosynthesis (Suppl. Table 12). Stimulation of serine biosynthesis in hiPSC-derived cardiomyocytes from patients with genetic DCM rescues contractile dysfunction in-vitro by increasing the glucose flux into the citric acid cycle and oxidative phosphorylation [78]. These seemingly contradictory findings suggest that levels of serine could be a driver of energy balance in cardiomyocytes. This hypothesis needs to be experimentally verified.

Two level-3 SCPs involved in cholesterol synthesis and another SCP predicted based on genes involved in cholesterol export (Suppl. Table 12) were up- and downregulated by the cardiotoxic TKIs, with the AUC ranks 15, 22 and 23, respectively (Figure 2B, Suppl. Figure 18). Both synthesis SCPs were also downregulated by noncardiotoxic drugs, with the AUC ranks 3 and 24. Our results link higher intracellular cholesterol levels, either based on increased synthesis or decreased export, to a cardiotoxic response. While six cardiotoxic (Suppl. Figure 20) and eleven noncardiotoxic (Suppl. Figure 21) TKIs induce the described pathway activities in opposite directions with varying enrichment ranks, it is, in particular, the cardiotoxic TKI lapatinib that upregulates both cholesterol synthesis SCPs with top enrichment ranks (5x1 and 2, 3, 4, 5, 6). Our observations agree with the results of a meta-analysis documenting a cardioprotective effect of statin treatment against chemotherapy-induced cardiomyopathy [41], though statins might induce multiple cardioprotective mechanisms besides inhibition of HMG-CoA reductase, the rate-controlling enzyme in cholesterol synthesis [79]. Since lapatinib resistance of breast cancer cells due to increased cholesterol synthesis could be reversed *in-vitro* by statin treatment [75], our results might suggest a potential effect of statin treatment on lapatinib's cardiotoxicity as well. Potential detrimental effects of statin treatment in heart failure [80] and involvement of cholesterol biosynthesis intermediates in the antioxidant defense against ferroptosis [36] might indicate the need for correct balance of cholesterol homeostasis and TKI-selective supportive statin therapy.

#### Cellular antioxidant systems

The level-1 and -2 SCPs 'Cellular redox homeostasis' and 'Cellular antioxidant defense systems' in parent-child relationships (Suppl. Figure 19) were predicted with AUC ranks 2 and 5 (Suppl. Figure 18), respectively. Both were downregulated by vandetinib in three of three (enrichment ranks 3, 4, 5 and 3, 4, 7) and bevacizumab in two of four (6, 7 and 7, 8) treated cell lines (Suppl. Figure 20). Multiple animal models document involvement of oxidative stress in heart failure and oxidative stress biomarkers are increasingly used in the monitoring of heart failure patients [81]. Reduction of oxidative stress is a protective mechanism against ferroptosis [36], a cardiotoxic mechanism that is also suggested based on other identified SCPs.

**Posttranslational protein modification and translational quality control**

The cardiotoxic TKI dabrafenib upregulates the level-1 SCP ‘Posttranslational protein modification’ and its level-2 child SCP ‘Posttranslational protein modification and quality control during secretory pathway’ (Suppl. Figure 19) in two of five treated cell lines (both enrichment ranks 1, 2) (Suppl. Figure 20). Both SCPs were predicted for the cardiotoxic drugs with AUC ranks of 1 and 7, respectively (Suppl. Figure 18). Many of the upregulated SCP genes participate in protein folding, quality control and stress response in the endoplasmic reticulum (ER) (Suppl. Table 12). Their upregulation could be a response to protein accumulation in the ER that can be differentiated into two phases [82]. The initial physiological ER stress response of the heart addresses the accumulation of proteins in the ER, while in case of prolonged stress a pathological response triggers autophagy and apoptosis.

**Cellular signaling pathways**

Among the identified signaling pathways were PDGF, Natriuretic receptor, HIF-1 alpha, Oncostatin M and Hippo signaling (Figure 2B, Suppl. Figure 18).

PDGF signaling was identified as an SCP that favors a non-cardiotoxic response with the AUC rank 14 (Figure 2B, Suppl. Figure 18) and preferentially maps to cardiac fibroblasts and smooth muscle cells in the adult human heart (Figure 3B, Suppl. Figure 23). It is downregulated by seven cardiotoxic TKIs in one to six cell lines (Suppl. Figure 20). Ponatinib (cardiotoxicity 1-10%), trastuzumab (>10%) and sorafenib (1-10%) downregulated the SCP in all treated cell lines (enrichment ranks 4, 3x6, 13; 2, 4, 5, 11 and 5, 6, 10, 15, respectively). In agreement with our results, PDGF and PDGF signaling in the heart increase during disease states and are generally considered to play an important role during cardioprotection [83], as supported by the PDGF-induced increase in cell survival and contractility of engineered cardiac tissue [84,85].

The level-4 SCPs ’Atrial -’, ‘Brain-‘ and ‘C-type natriuretic receptor signaling’ and their level-3 parent SCP ‘Natriuretic peptide receptor signaling’ (Suppl. Figure 19) were identified to favor a non-cardiotoxic response (Suppl. Figures 18). They were upregulated by six, four, two and six overlapping non-cardiotoxic drugs in one to four cell lines, respectively (Suppl. Figure 21). The SCP ‘Atrial natriuretic peptide receptor signaling’ was additionally downregulated by the three cardiotoxic TKIs lapatinib, ponatinib and vandetanib in one of their treated cell lines (enrichment ranks 4, 4 and 2, respectively). Induced genes contain the ligands *NPPA*, *NPPB* and *NPPC* as well as receptor genes (Suppl. Table 12). Though our data was generated *in vitro*, the classification into cardiotoxic and non-cardiotoxic drugs is based on clinical data. A potential explanation for the identified association could be that both secreted ANP and BNP have a protective effect on cardiac preload, afterload and cardiovascular growth [86]. The endopeptidase neprilysin degrades several endogenous vasoactive peptides including natriuretic peptides [87]. Treatment schemes of patients with chronic heart failure that combine neprilysin and angiotensin II receptor inhibition achieve better results than treatment with an angiotensin converting enzyme (ACE) inhibitor [42,43], supporting the clinical relevance of our findings [87].

We identified genes involved in the inhibition of HIF-1 alpha signaling (Suppl. Table 12) as associated with a non-cardiotoxic response with the AUC rank 20 (Figure 2B, Suppl. Figure 18). They were downregulated by the two cardiotoxic TKIs dabrafenib and pazopanib in two (enrichment ranks 8, 8) and three (1, 6, 6) of five treated cell lines, respectively (Suppl. Figure 20). In agreement, only short-term HIF-1 signaling in acute stress situations has a cardioprotective effect, while continuously active HIF-1 signaling might be harmful [88], suggesting an explanation of why its inhibition under long-term TKI treatment might improve cardiac outcomes.

A similar effect was described for Oncostatin M (OSM) receptor signaling that favors a cardiotoxic response (Suppl. Figure 18), since it is downregulated by two non-cardiotoxic TKIs in two and four cell lines (Suppl. Figure 21). Stimulation of OSM signaling cascades mitigates cardiac damage in acute stress conditions, while its chronic activation contributes to the development of heart failure [89].

The level-3 SCP 'Hippo signaling' was downregulated by cardiotoxic TKIs with an AUC rank of 13 (Figure 2D, Suppl. Figure 18). Pazopanib downregulates this SCP in three of five treated cell lines (enrichment ranks 1, 2x6) (Suppl. Figure 20). It preferentially maps to cardiac fibroblasts in the adult human heart (Suppl. Figure 23). Five of six downregulated genes inhibit the Hippo downstream transcription factor YAP1 (Suppl. Table 12). Consistent with this finding, increased YAP activity and expression of hypertrophic target genes was also observed in HCM patient tissue cell line/drug combinations and a HCM murine model [90].

**Extracellular matrix organization**

Fibrillar collagen constitutes the most abundant protein in the cardiac extracellular matrix (ECM) which provides structural organization for the correct alignment of cardiomyocytes, generates myocardial stiffness and helps in force transmission [91]. Diffuse interstitial fibrosis, a histopathological feature observed in non-ischemic cardiomyopathies or hypertensive heart disease is characterized by excessive deposition of type I and III collagen, by changing ratios of type I to type III collagen and by an increase in collagen crosslinking. The degree of collagen crosslinking, but not total collagen deposition correlates with surrogate parameters (e.g., myocardial stiffness) or hospitalization in patients with hypertensive heart disease [44,45]. In agreement, ‘Collagen fiber cross-linking’ is downregulated by eleven non-cardiotoxic TKIs in one to six cell lines (Suppl. Figure 21), identified with the AUC rank 7 and consequently categorized as an SCP that favors a cardiotoxic response (Suppl. Figures 18). The downregulation of the SCP ‘Elastin cross-linking and assembly’ (AUC rank 16) is predicted based on genes shared with the collagen-cross linking SCP. The SCP ‘Collagen fibril organization by fibril-associated bridges’ that contains the FACIT collagens is upregulated by eleven noncardiotoxic TKIs in one to six cell lines (Suppl. Figure 21) and was identified with the AUC rank 17 as an SCP whose higher activity favors a noncardiotoxic response (Suppl. Figure 18). The reduced expression of FACIT collagen during progressive liver cirrhosis is associated with a loss of flexibility [92]. Whether FACIT collagen can exhibit a similar effect on the cardiac wall needs to be investigated.

**Water transmembrane transport**

Down- and upregulation of aquaporins that are the main components of the level-3 SCP ‘Water transmembrane transport’ is predicted with AUC ranks of 9 and 2 for cardiotoxic and noncardiotoxic TKIs, respectively (Figure 2B, Suppl. Figure 18). Three cardiotoxic drugs, dabrafenib, pazopanib and ponatinib, downregulate the SCP in two of three (enrichment ranks 6, 7), one of five (4) and two of six (6, 9) treated cell lines (Suppl. Figure 20). The induced and repressed SCP genes, aquaporins 1, 3, 7 and 10 (Suppl. Table 12C) might be related to their function in transmembrane transport of water (AQP1, 3, 7, 10), CO2 and NO (AQP1) or urea and the energy substrate glycerol (AQP 3, 7, 10) [93].

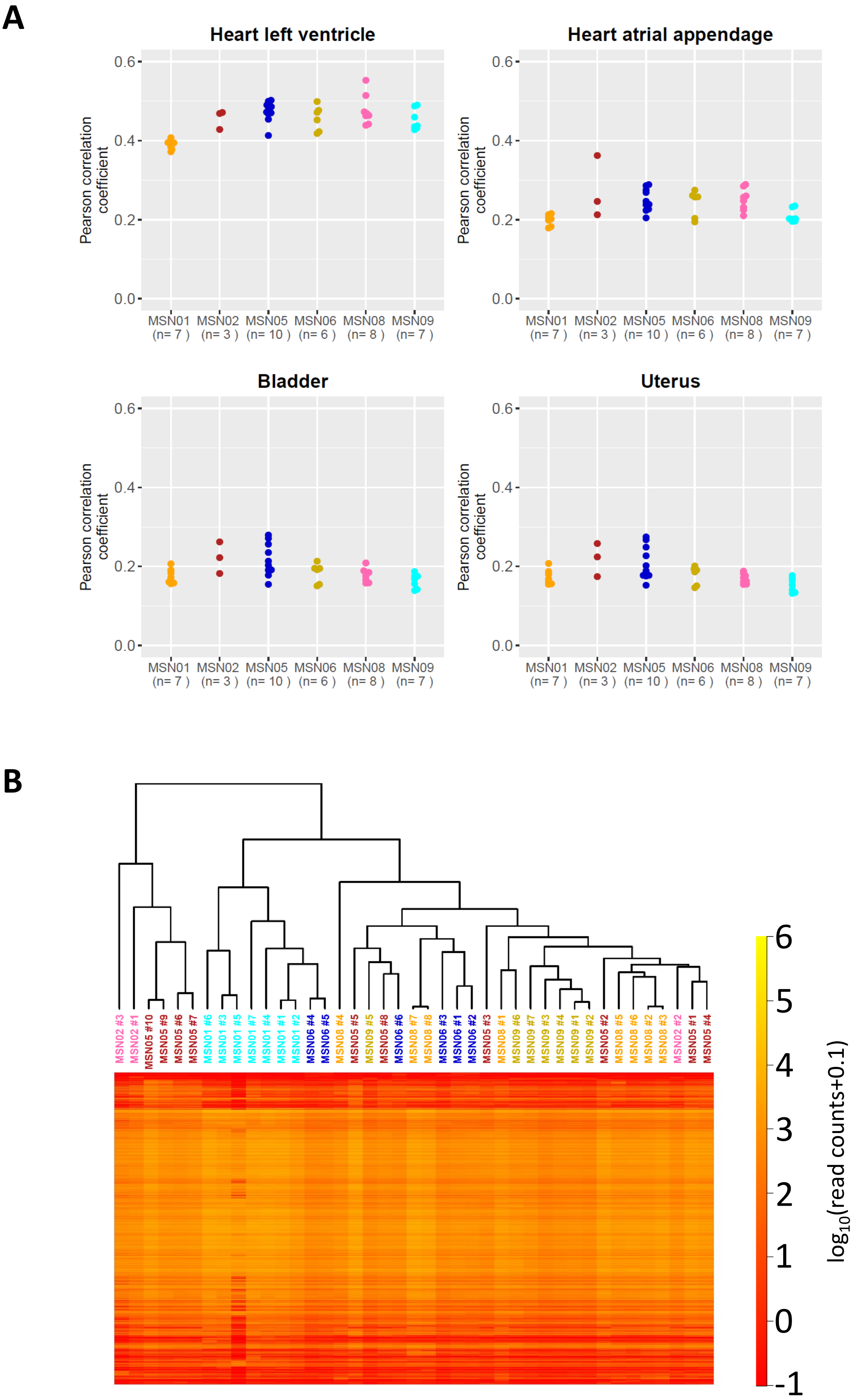


**Supplemental figure 1**

**Supplemental Figure 1. Basal gene expression of six hiPSC-derived cardiomyocyte cell lines map to the human heart and show cell line specificity. (A)** Raw read counts of vehicle-treated replicates [16] were correlated with median gene expression levels for each tissue in the GTEx database. Pearson correlation coefficients are shown for each replicate of all cell lines and the top four tissues with the highest correlation coefficients. Numbers of replicates are provided in parentheses. **(B)** Gene expression raw counts obtained in all replicates of vehicle-treated cell lines [16] were subjected to pairwise correlation analysis, followed by hierarchical clustering. Visualized matrix shows the $\log_{10}$(read counts + 0.1). Rows and columns were re-arranged according to clustering results. Replicates of the same cell line are colored with the same color.

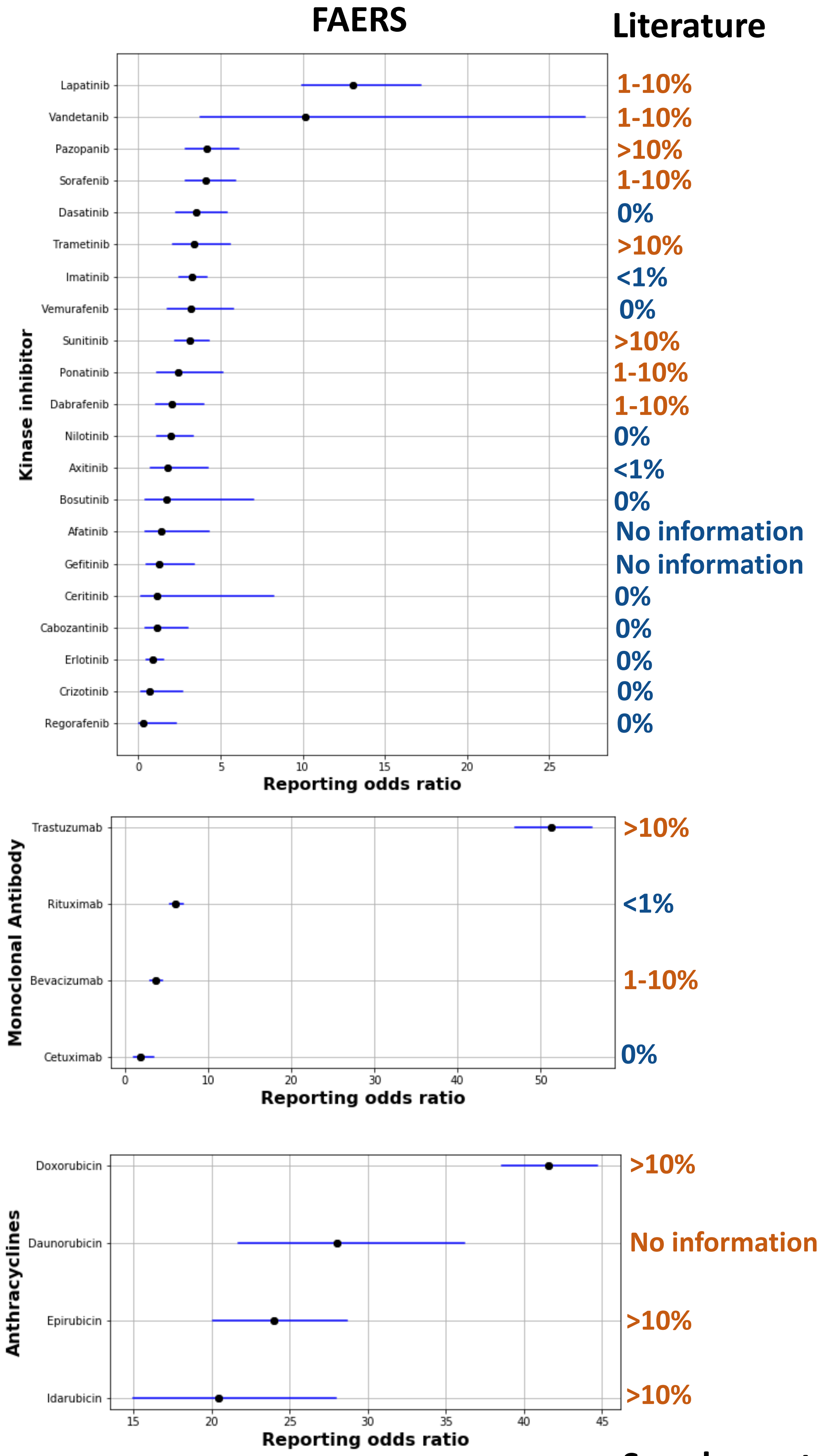


Supplemental figure 2

**Supplemental Figure 2. Cardiac toxicity of kinase inhibitors and monoclonal antibodies curated from the FAERS database.** Risk profiles were curated from the FAERS database. Horizontal lines indicate 95% confidence intervals. Blue and orange comments describe published cardiotoxicity levels as outlined in Suppl. Table 4.

**A**

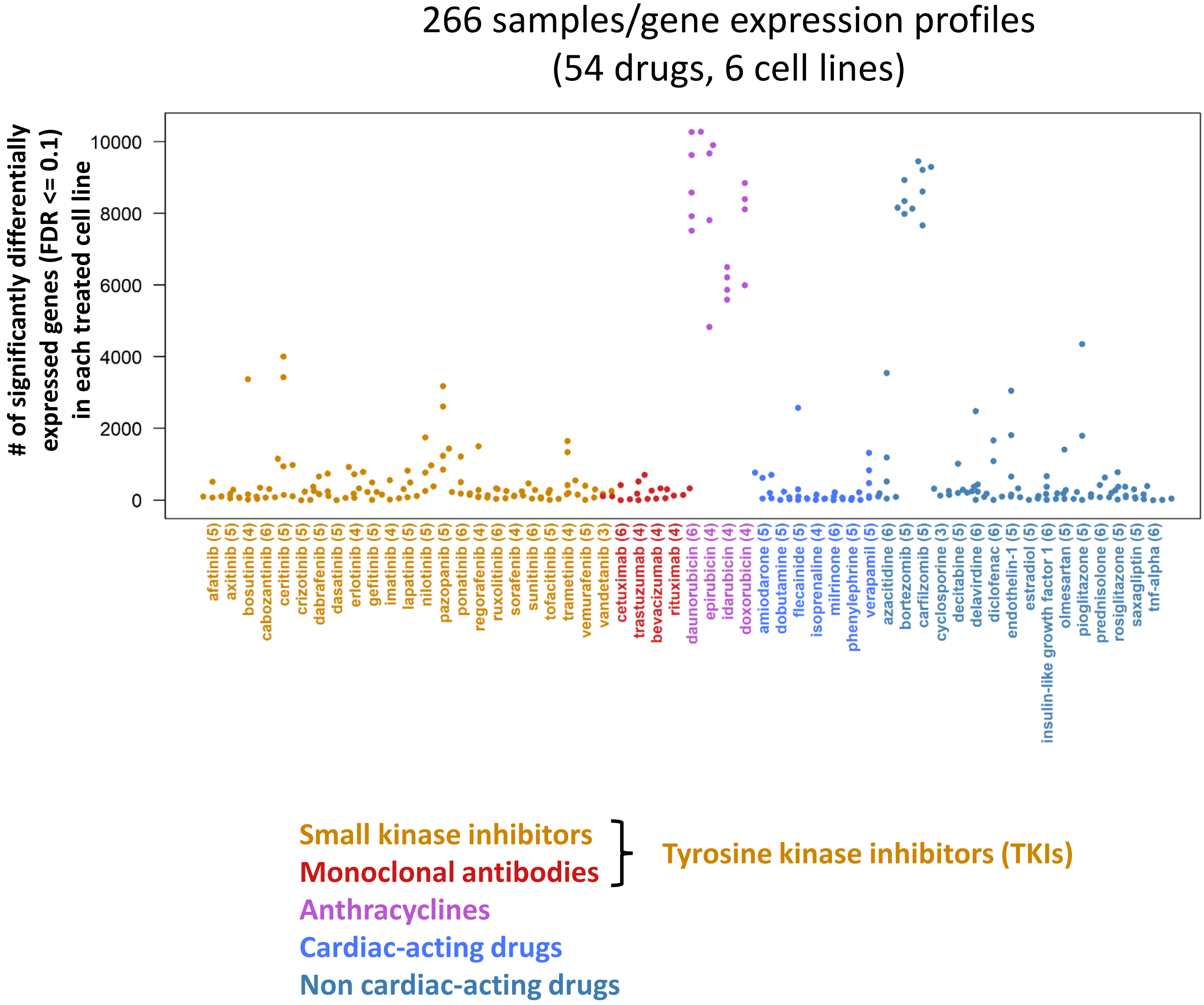


**Supplemental figure 3**

B

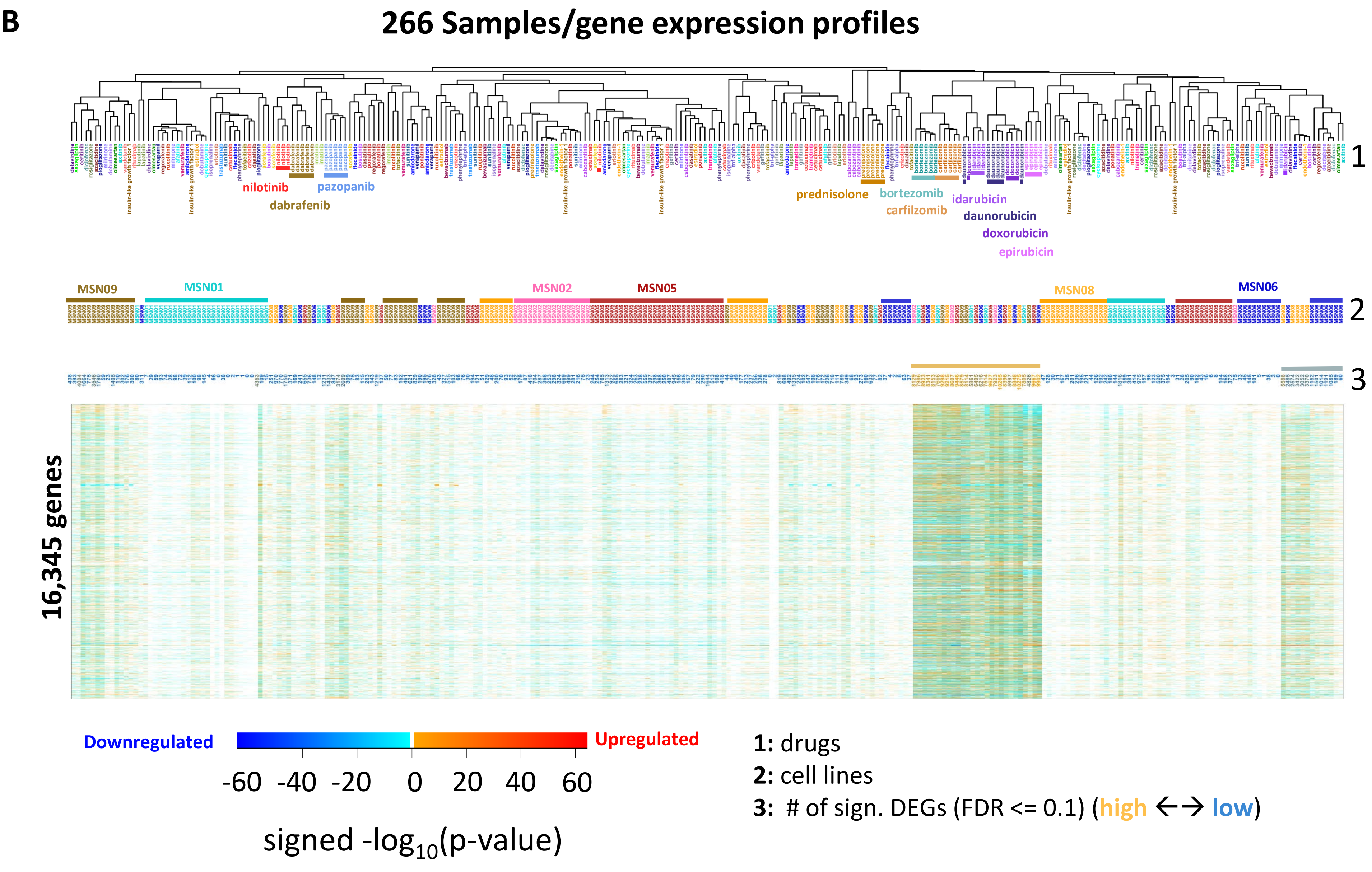


Supplemental figure 3

**Supplemental Figure 3. Drug-induced gene expression profiles in six hiPSC-derived cardiomyocyte cell lines. (A)** Three to six hiPSC-derived cardiomyocyte cell lines were stimulated with one out of 54 drugs or vehicle for 48h, followed by bulk RNAseq and identification of 266 lists of differentially expressed genes (DEGs). The number of DEGs induced by the different drugs in the different cell lines showed great variation (FDR <= 10%). Total numbers of treated cell lines for each drug are shown in parentheses next to the drug labels. **(B)** Significance p-values were transformed into -$\log_{10}$(p-values) and defined to be positive or negative for up- or downregulated genes, respectively. Pairwise correlation analysis followed by hierarchical clustering, documents that only a few gene expression profiles are determined by the drug used for treatment (1), while most profiles are determined by the treated cell line (2) or the number of significant DEGs (3). Figure 1B shows the same dendrogram.

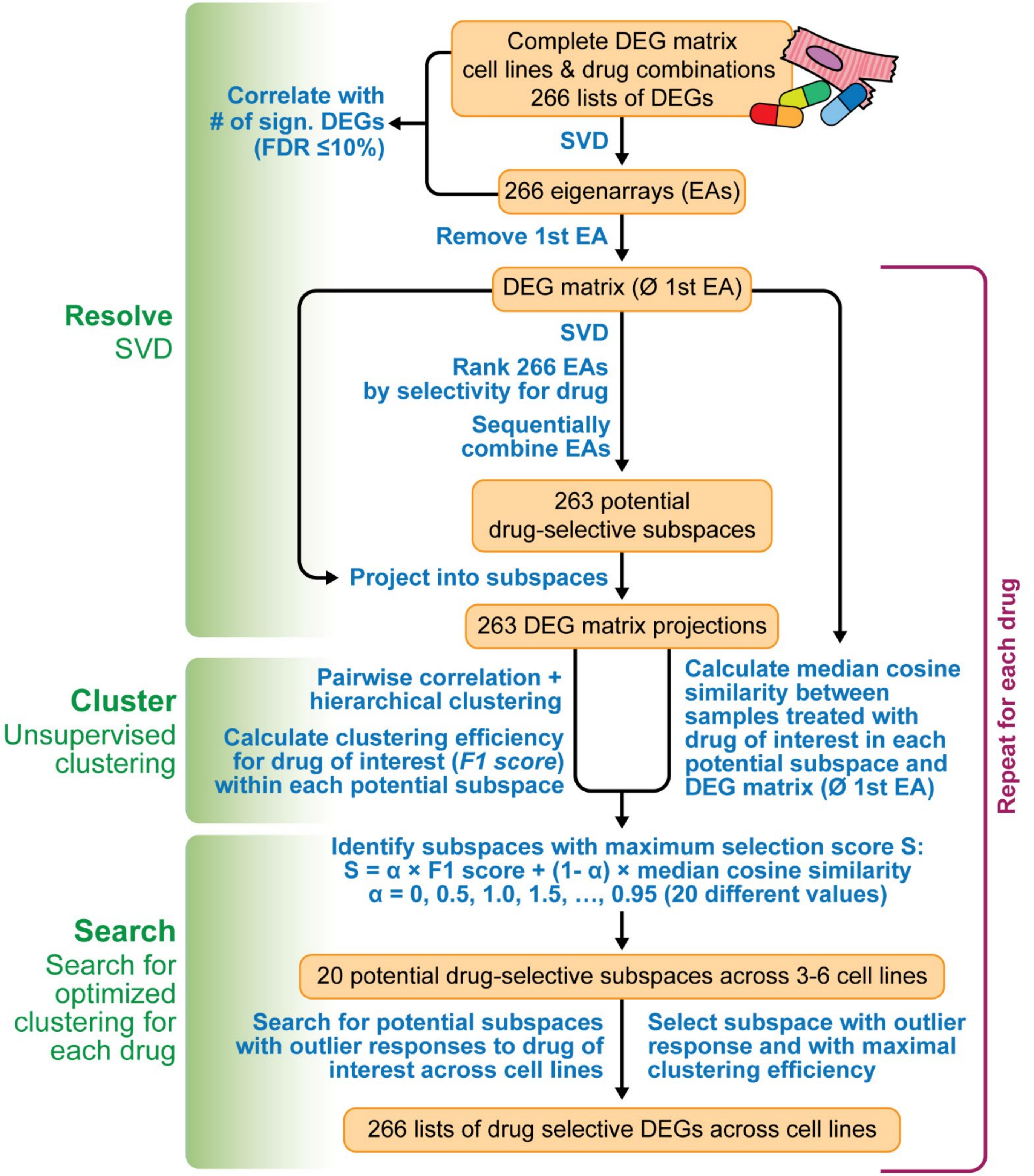


**Supplemental figure 4**

**Supplemental Figure 4. Computational pipeline for the identification of drug-selective and outlier gene expression responses.** Our pipeline subjects drug-induced differentially expressed genes (DEGs) to Singular Value Decomposition (SVD) to identify drug-selective gene expression profiles and cell lines that respond differently to a drug of interest than the other cell lines. See methods for detail.

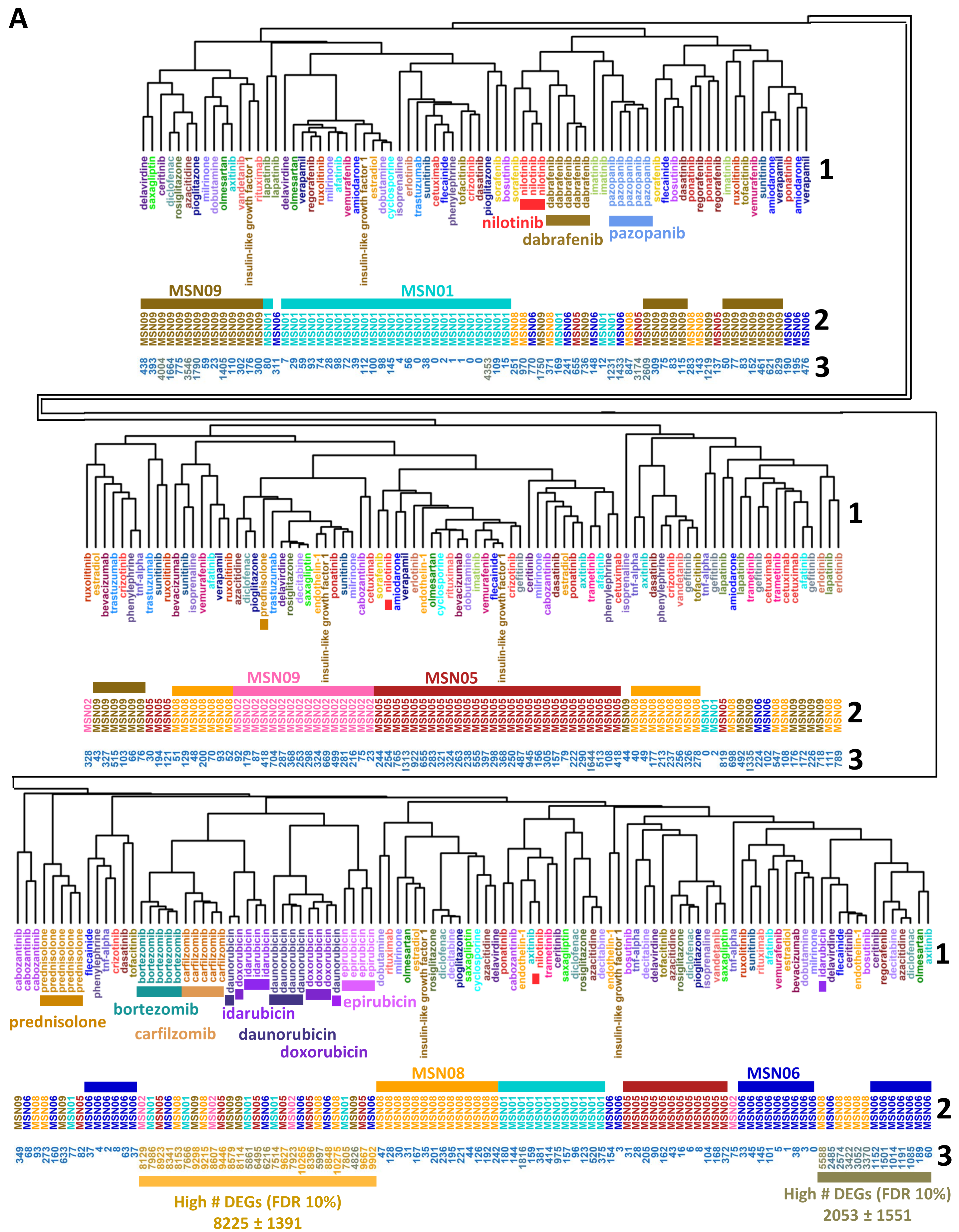


Supplemental figure 5

**B**

**dasatinib cluster in complete gene expression profiles**

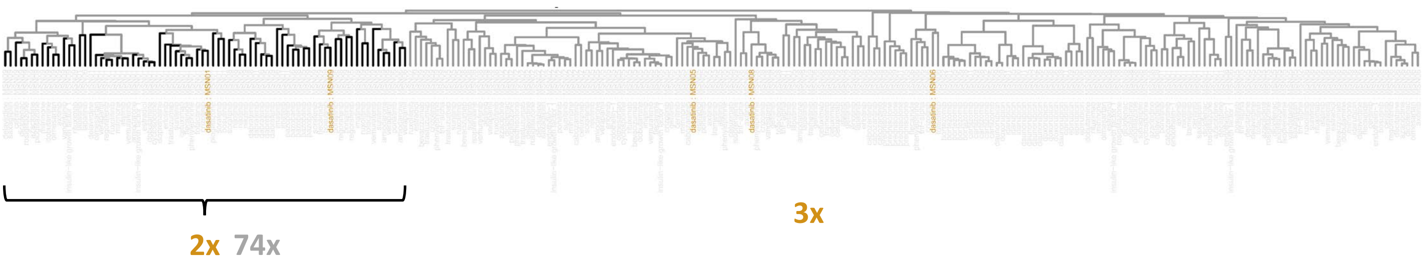


**Cluster with highest F1 score for dasatinib-treated cell lines**
**Precision = 2/74, Recall = 2/5, F1-score = 0.05**

**Supplemental figure 5**

**Supplemental Figure 5. Clustering efficiency of drug-induced gene expression profiles in six hiPSC-derived cardiomyocyte cell lines. (A)** To allow detailed investigation of the clustering results we visualized dendrogram and dendrogram labels of Suppl. Figure 3B at larger sizes. Figure 1B shows the same dendrogram with a focus on the drugs. **(B)** For each drug, we calculated one F1 score for each cluster that can be obtained by cutting the dendrogram at any height and contains at least two cell line/drug combinations. The F1 score is the harmonic mean of the precision (how many cell line/drug combinations within a cluster were treated with the drug) and the recall (how many cell line/drug combinations treated with the drug were in that cluster). The cluster with the highest F1 score that was selected for further analysis in this example is labeled black.

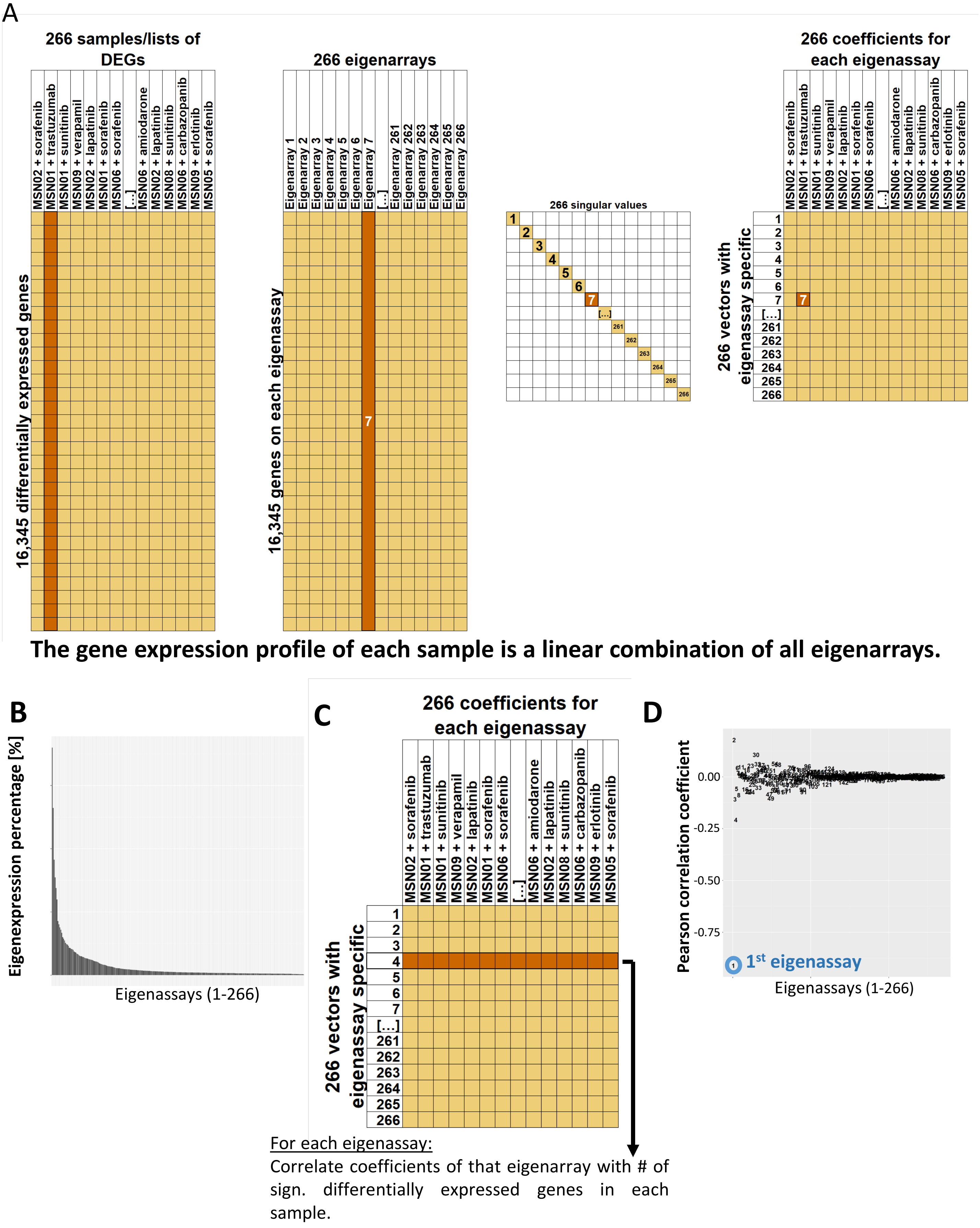


Supplemental figure 6

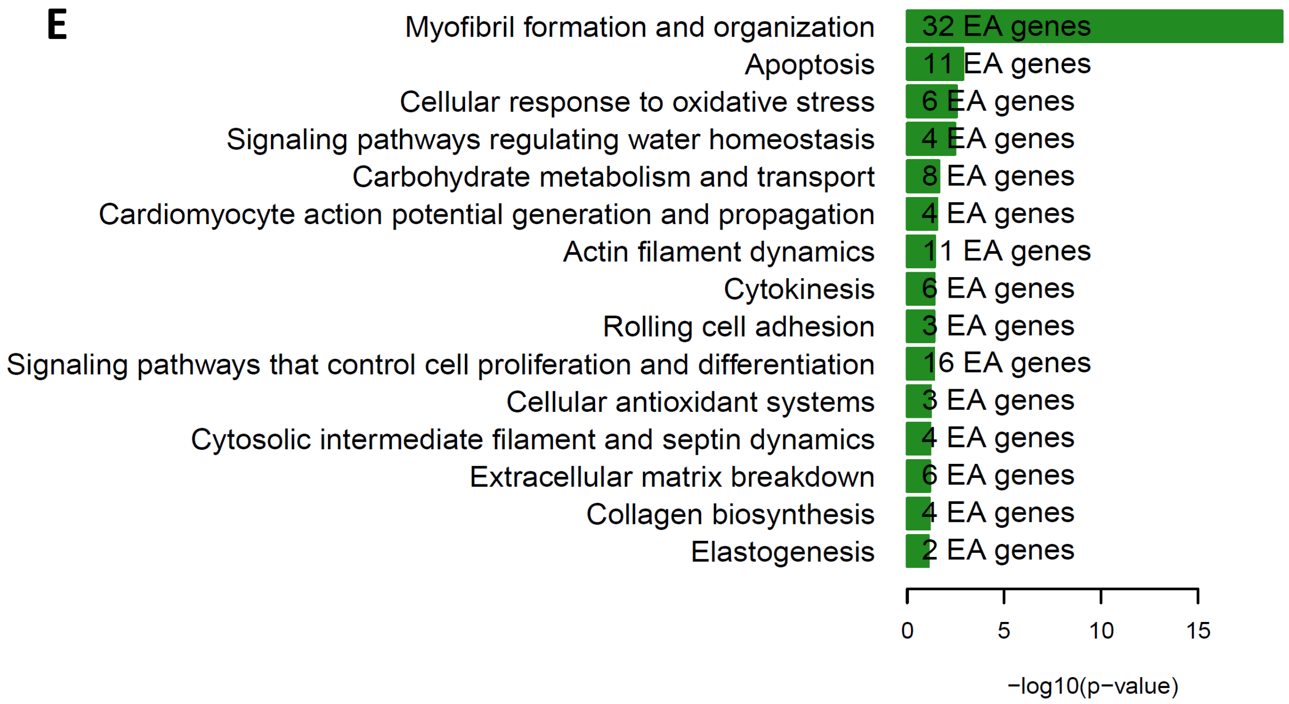


**Supplemental figure 6**

**Supplemental Figure 6. Identification of drug-selective gene expression responses using Singular Value Decomposition. (A)** Singular value decomposition (SVD) decomposes the input data matrix into a matrix of left singular vectors or eigenarrays, a diagonal matrix of singular or eigenexpression values and a matrix of right singular vectors. Each cell line/drug combination gene expression vector in the full matrix is a linear combination of all eigenarrays. Cell line/drug combination specific coefficients of this linear combination are documented in the matrix of right singular vectors. The eigenexpression values in the diagonal document how much each eigenarray contributes to the complete gene expression dataset of all cell line/drug combinations and needs to be considered for the linear combination as well. To calculate the contribution of the seventh eigenarray to the complete gene expression profile induced by trametinib in cell line MSN09 it must be multiplied with highlighted eigenexpression value and the highlighted coefficient, both labeled with seven. **(B)** SVD of the gene expression matrix identified 266 orthonormal eigenarrays that are sorted by their relative contribution to the total variance. **(C)** For each eigenarray, we calculated the Pearson correlation between the cell line/drug combination-specific coefficients and the number of significantly differentially expressed genes (DEGs) in the corresponding full gene expression profiles. **(D)** Our results document a high correlation with the number of significant DEGs for the first eigenarray. **(E)** Pathway enrichment analysis of the top 600 genes of the first eigenarray identifies muscle contraction.

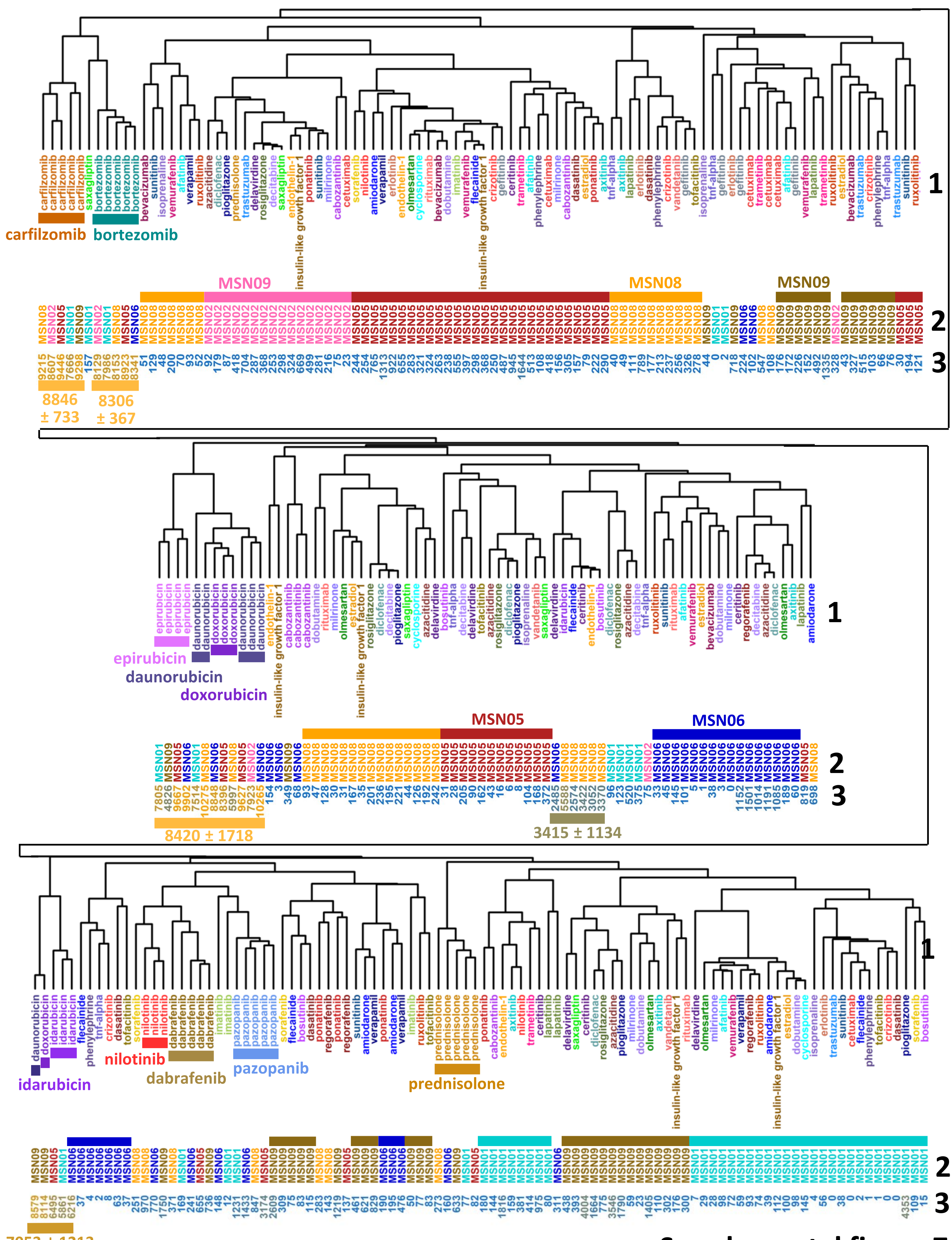


Supplemental figure 7

**Supplemental Figure 7. Clustering of DEGs after removal of first eigenarray.** Removal of the first eigenarray from the complete DEG matrix disrupts hierarchical clustering by the number of significant DEGs (3).

.

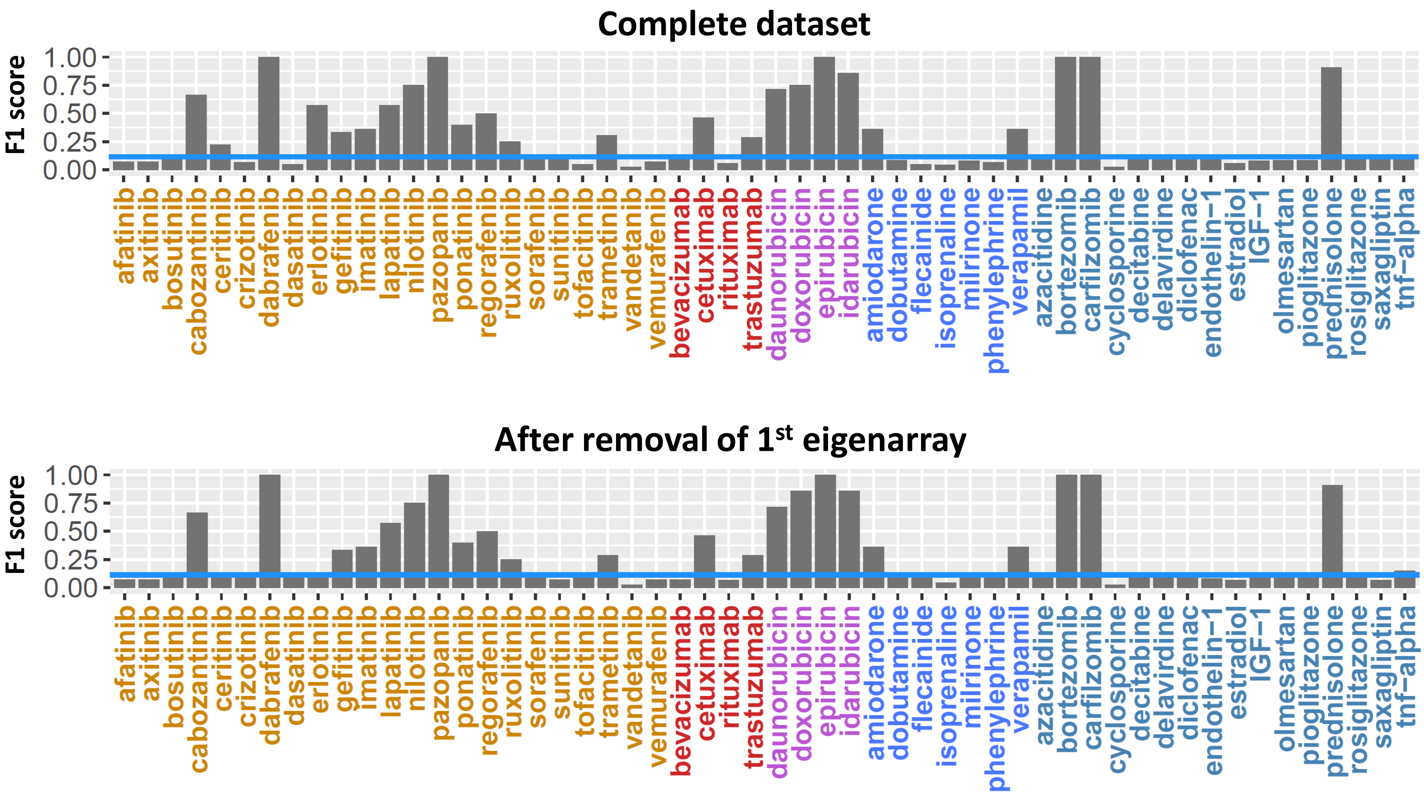


**Supplemental figure 8**

**Supplemental Figure 8. Clustering efficiency after removal of the first eigenarray.** DEG matrix after removal of the first eigenarray was subjected to pairwise correlation analysis and hierarchical clustering, followed by calculation of the highest F1 scores for each drug (bottom figure). To allow easier comparison, we added the F1 scores calculated for each drug using the complete DEG profiles (top figure) that is also shown in main Figure 1C.

**A**

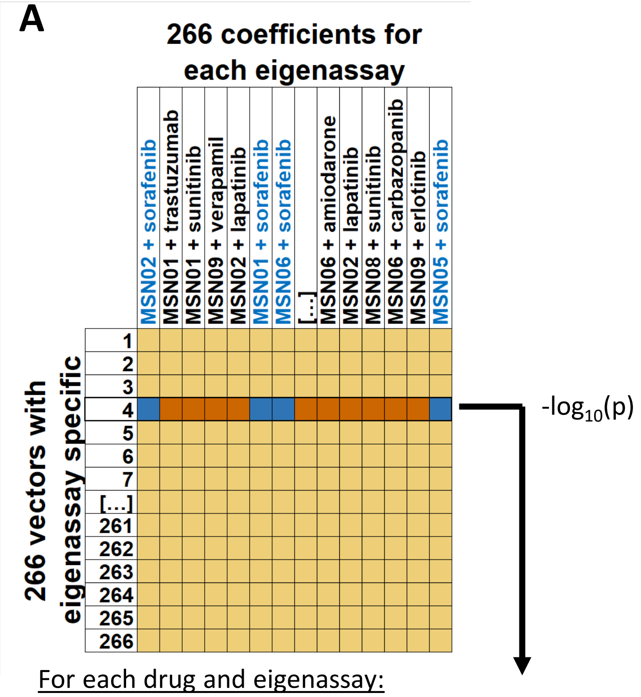


For each drug and eigenassay:

Use student's t-test to investigate if the coefficients for an eigenarray of interest significantly differ between all samples of a drug of interest and all other samples.

**Supplemental figure 9**

B

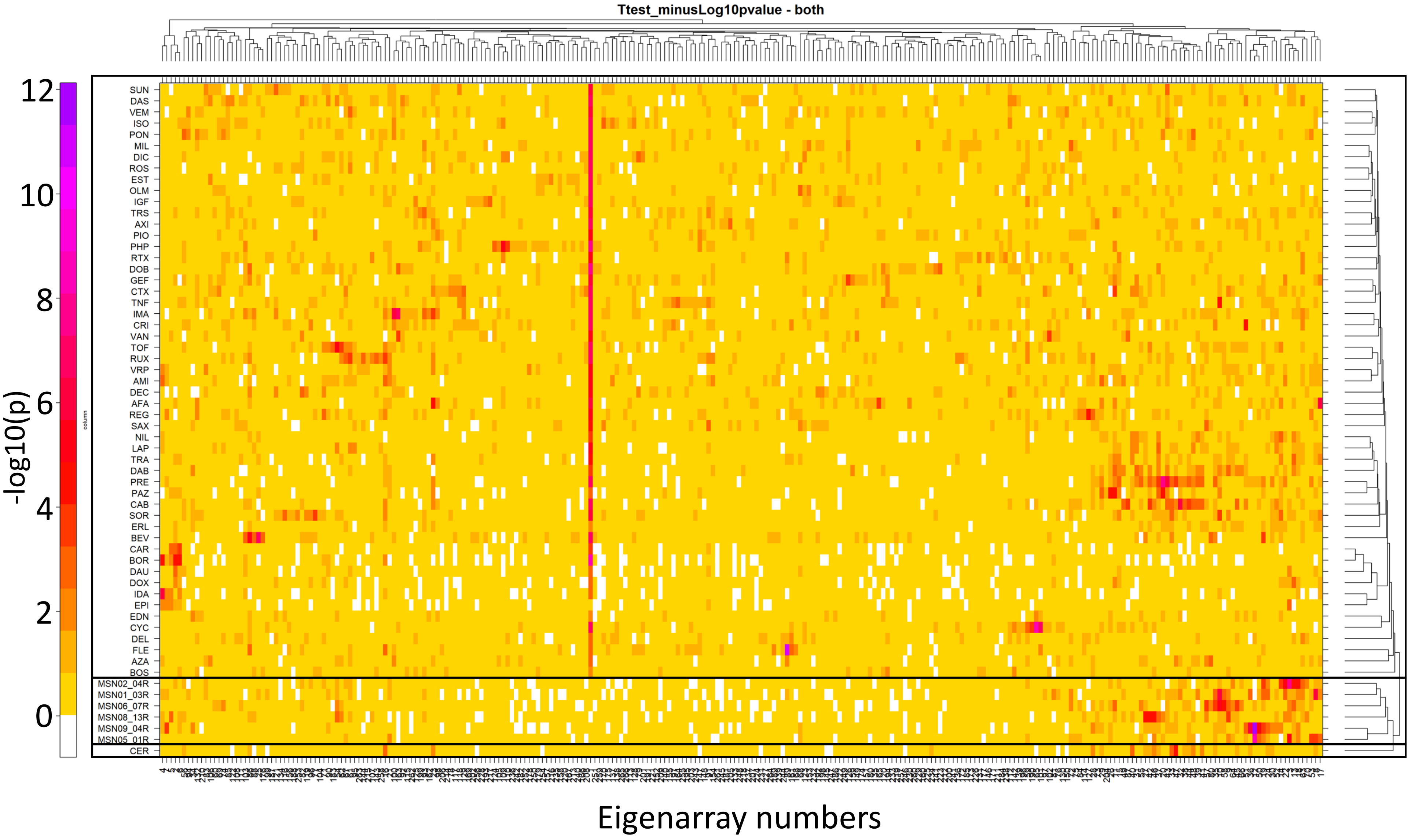


Supplemental figure 9

**Supplemental Figure 9. Cell-line- and drug-selective effects are captured by different eigenarrays. (A)** For each eigenarray and drug, we analyzed if the coefficients that are related to the gene expression profiles for that drug on that eigenarray significantly differ from all other coefficients on that eigenarray. Consequently, we calculated one p-value for each drug-eigenarray combination. Similarly, we calculated one p-value for each cell line-eigenarray combination. **(B)** All p-values were transformed into -$log_{10}$(p-values) and used to calculate pairwise correlation coefficients between all drugs and cell lines, followed by hierarchical clustering. The initial heatmap of -$log_{10}$(p-values) was rearranged according to the clustering results. Grouping of the six cell lines into a single separated cluster (boxed) suggests that the eigenarray decomposition allows differentiation of cell-line-specific effects from drug-specific effects.

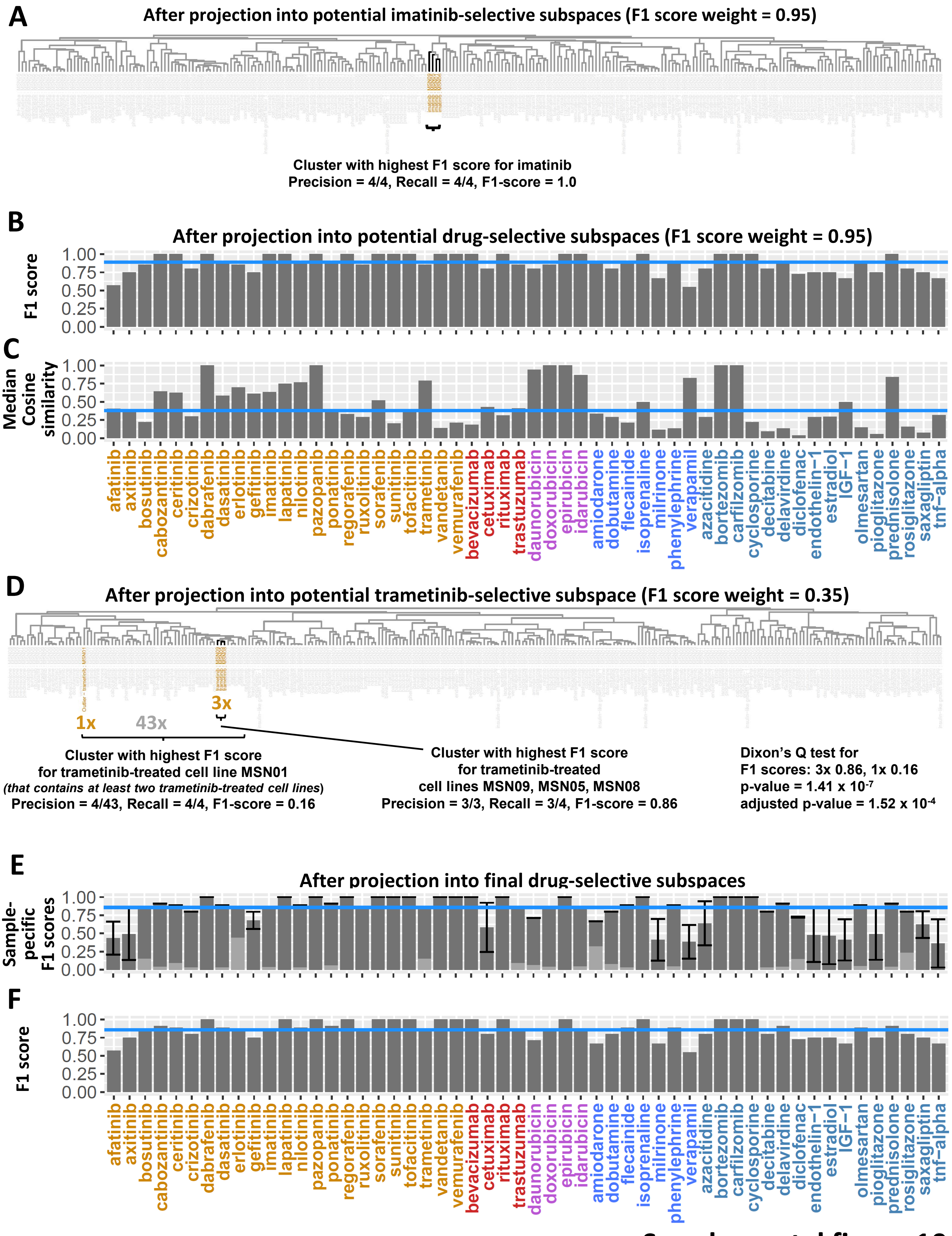


Supplemental figure 10

**Supplemental Figure 10. Identification of drug-selective gene expression profiles and outlier responses. (A)** For each drug, we ranked all eigenarrays by their ability to separate the coefficients associated with cell line/drug combinations treated with that drug from all other coefficients, i.e. we ranked them by increasing p-values. The top 3 to 266 eigenarrays were combined to yield 264 potential drug-selective subspaces. Gene expression profiles after removal of the first eigenarray were projected into the subspaces, followed by pairwise correlation, hierarchical clustering and F1 score calculation for the drug of interest. The example shows the F1 score calculation for imatinib after projection of the data into one of the potential imatinib-selective subspaces. **(B/C)** Median cosine similarities between the projected gene expression profiles and the gene expression profiles after removal of the first eigenarray were calculated for each drug. For each potential drug-selective subspace, we calculated weighted averages between the F1 score and the median cosine similarity with changing relative contributions as defined by 20 different F1 score weights (ranging from 0.00 to 0.95 in steps of 0.05). Each F1 score weight allowed us to select one drug-selective subspace, i.e., that subspace with the highest weighted mean or selection score. Shown are **(B)** F1 scores and **(C)** median cosine similarities obtained based on an F1 score weight of 0.95. Blue lines indicate median values. **(D)** For each drug, we screened all 20 potential drug-selective subspaces (that are defined by different F1 score weights) for subspaces where one cell line/drug combination shows a different transcriptomic response to the drug of interest than all other cell line/drug combinations. We calculated cell line/drug combination-specific F1 scores, using the same approach described above, except that the cell line/drug combination of interest has to be part of the corresponding cluster. Dixon's Q test applied to cell line/drug combination-specific F1 scores was used to identify outliers (adj. p-value = 0.05). **(E)** Cell line/drug combination-specific F1-scores of non-outlier cell line/drug combinations in the final drug-selective subspaces were averaged (dark gray bars). Error bars show standard deviations. F1 scores identified for outlier cell line/drug combinations are visualized separately (light gray bars). The blue line indicates the median height of dark gray bars. **(F)** Projection of gene expression profiles into the final drug-selective subspaces leads to a great increase in drug-specific F1 scores. Notice that the F1 scores are the maximum cell line/drug combination-specific F1-scores shown in E. This figure is the same as figure 1D.

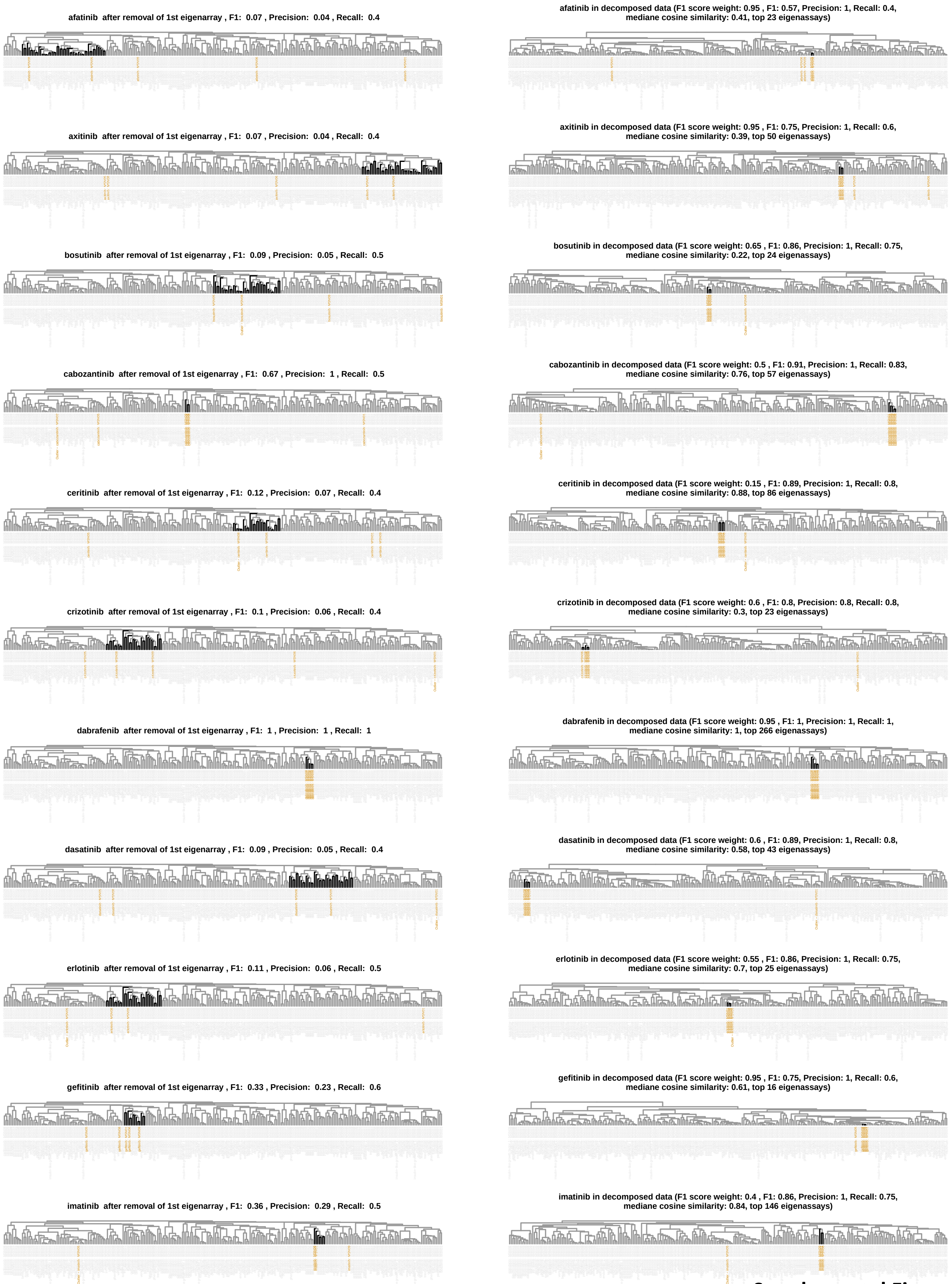


**Supplemental Figure 11**

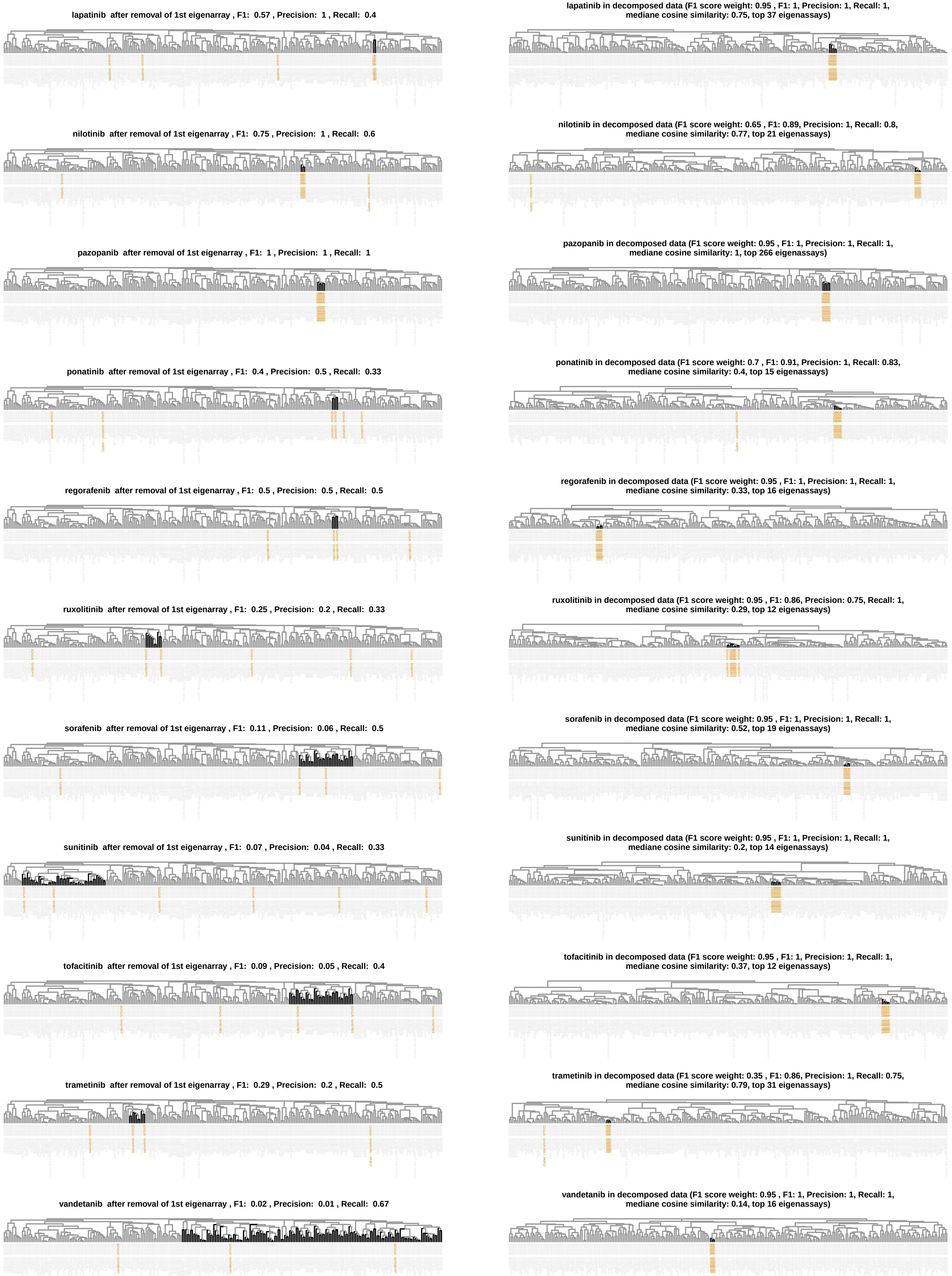


**Supplemental Figure 11**

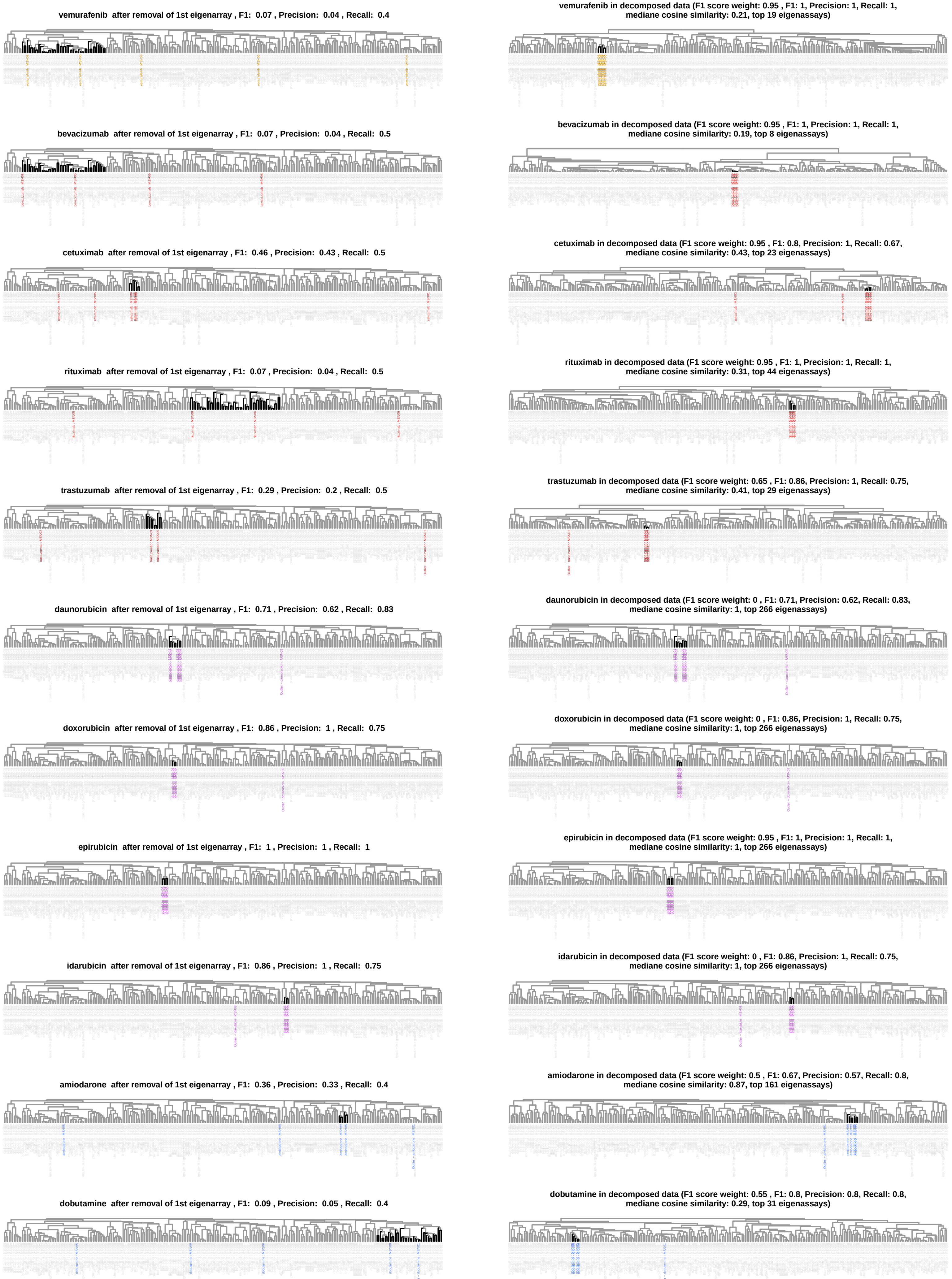


**Supplemental Figure 11**

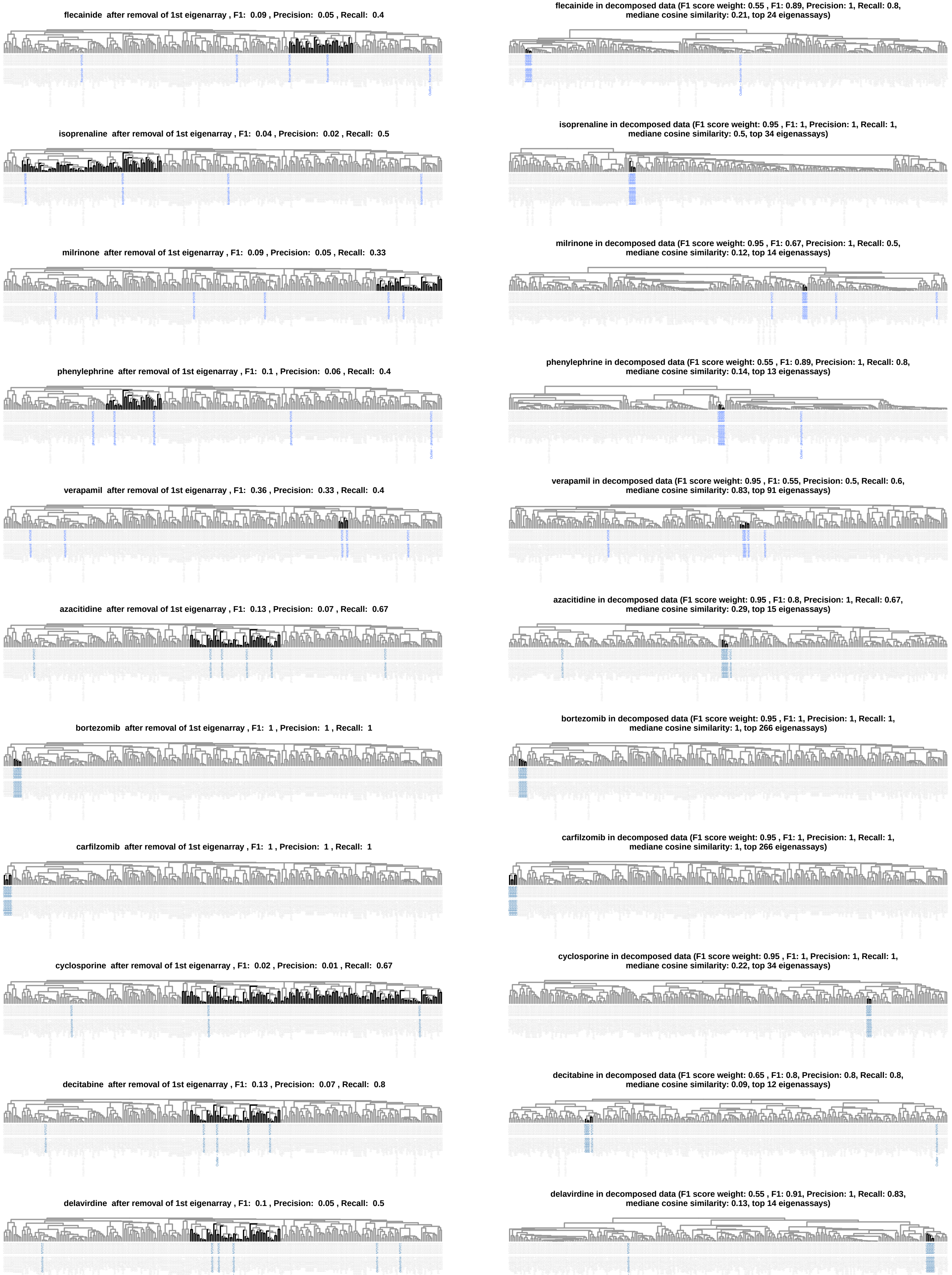


**Supplemental Figure 11**

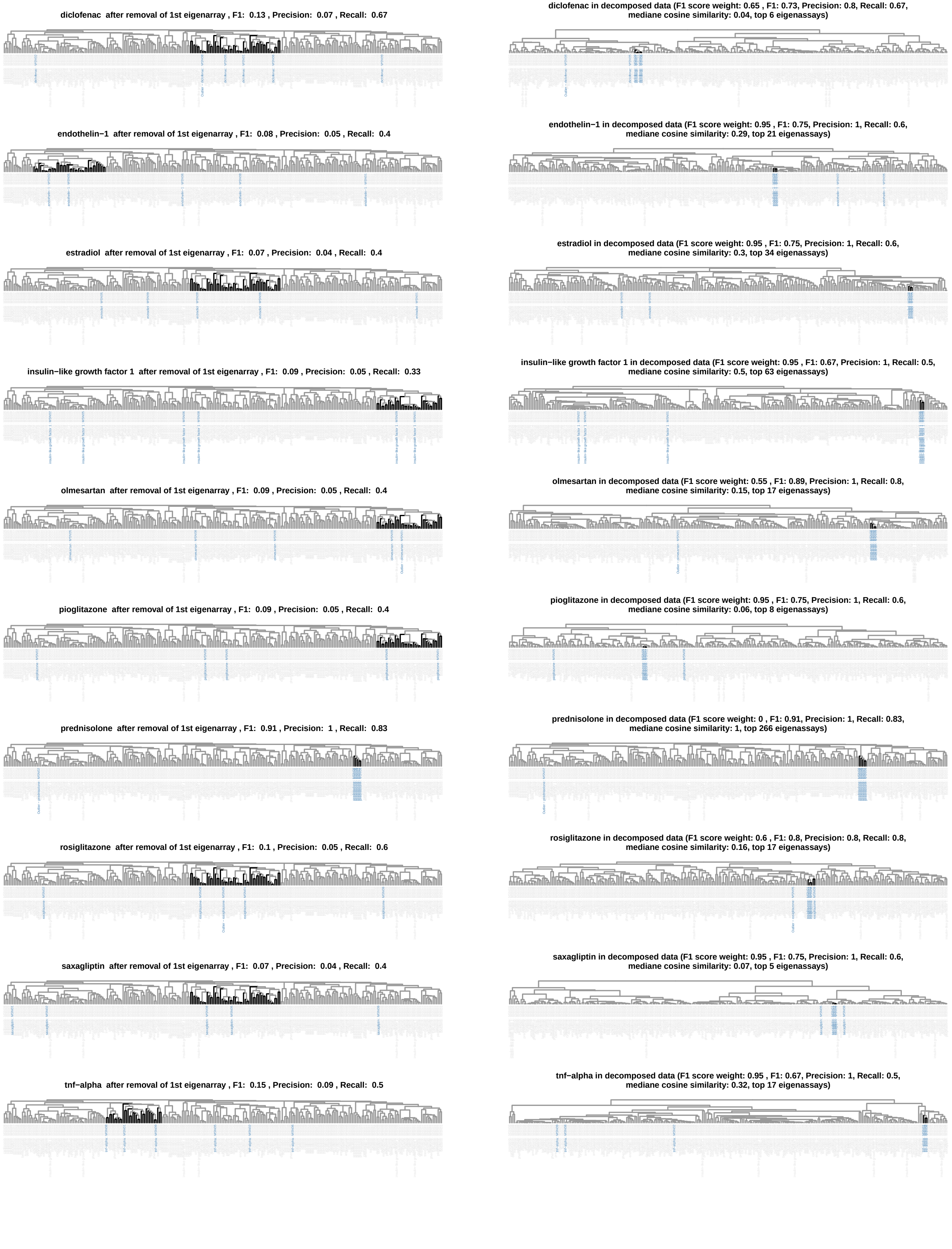


**Supplemetnal Figure 11**

**Supplemental Figure 11. Clustering results for each drug after removal of the first eigenarray and in the final drug-selective subspaces.** Clusters with the highest F1 scores are labeled black. Small molecule kinase inhibitors, monoclonal antibodies, anthracyclines, cardiac acting and non-cardiac acting drugs are labeled orange, red, purple, blue and gray-blue. The cluster dendrogram shown in Suppl. Figure 10D is part of this figure set as well.

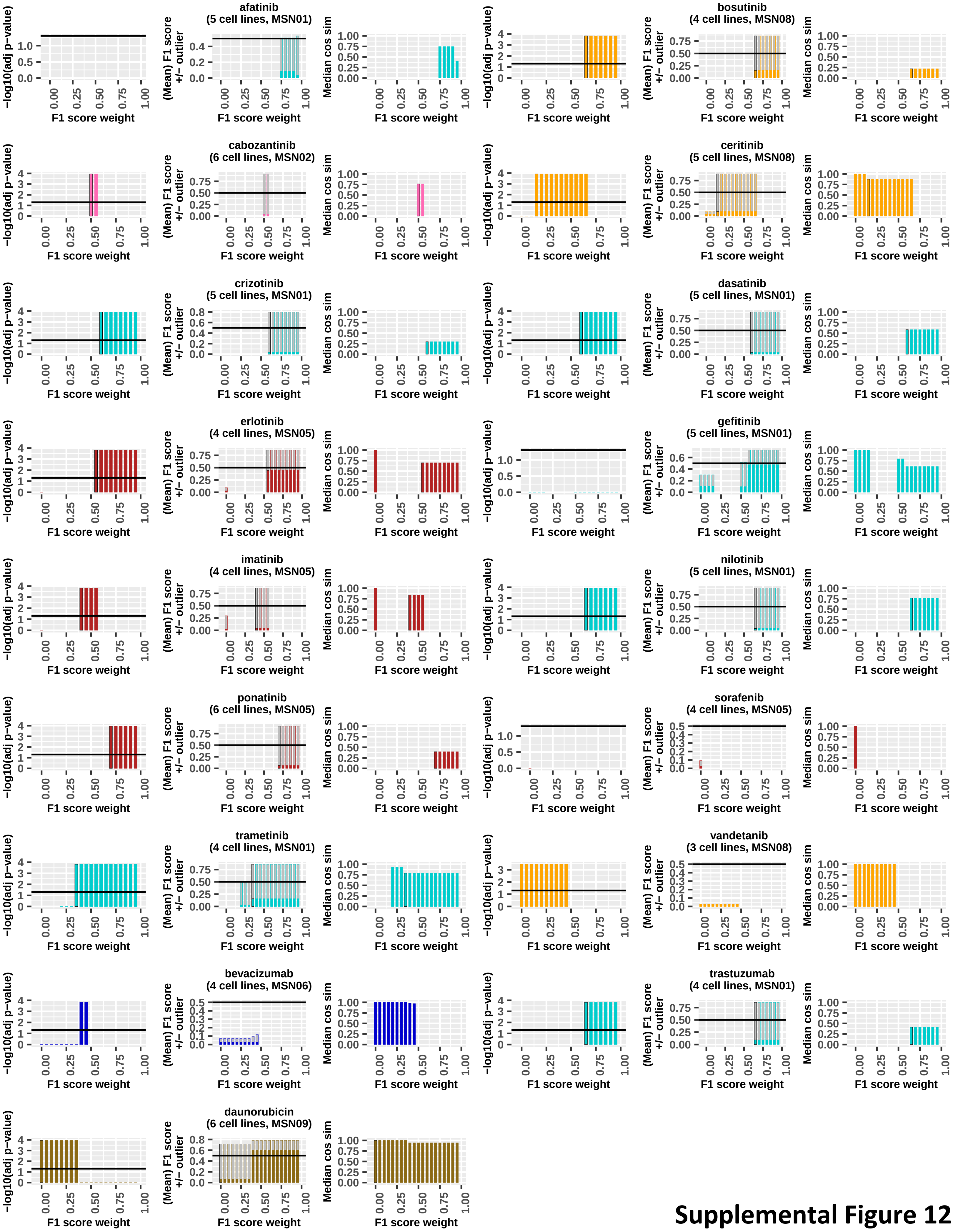


Supplemental Figure 12

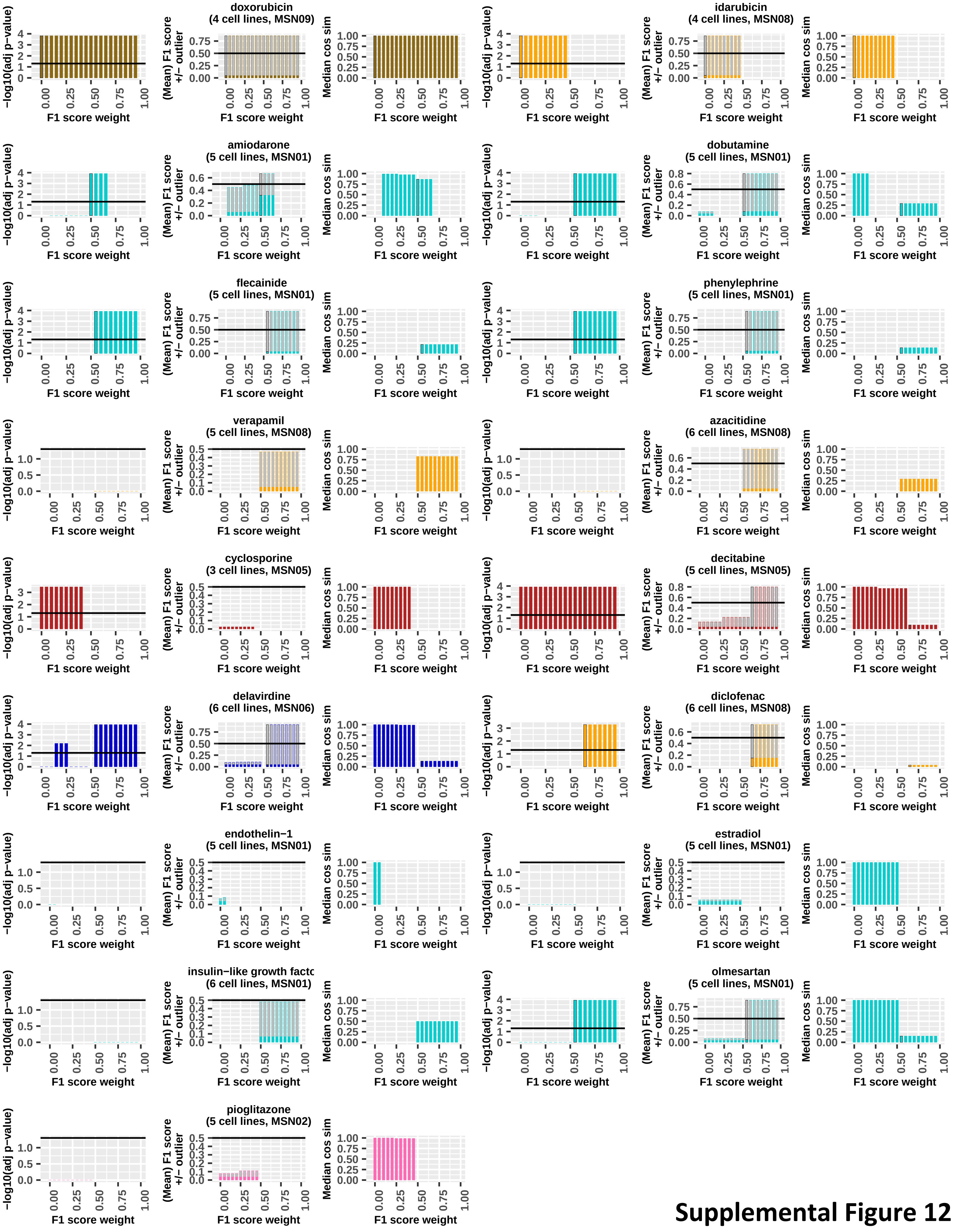


Supplemental Figure 12

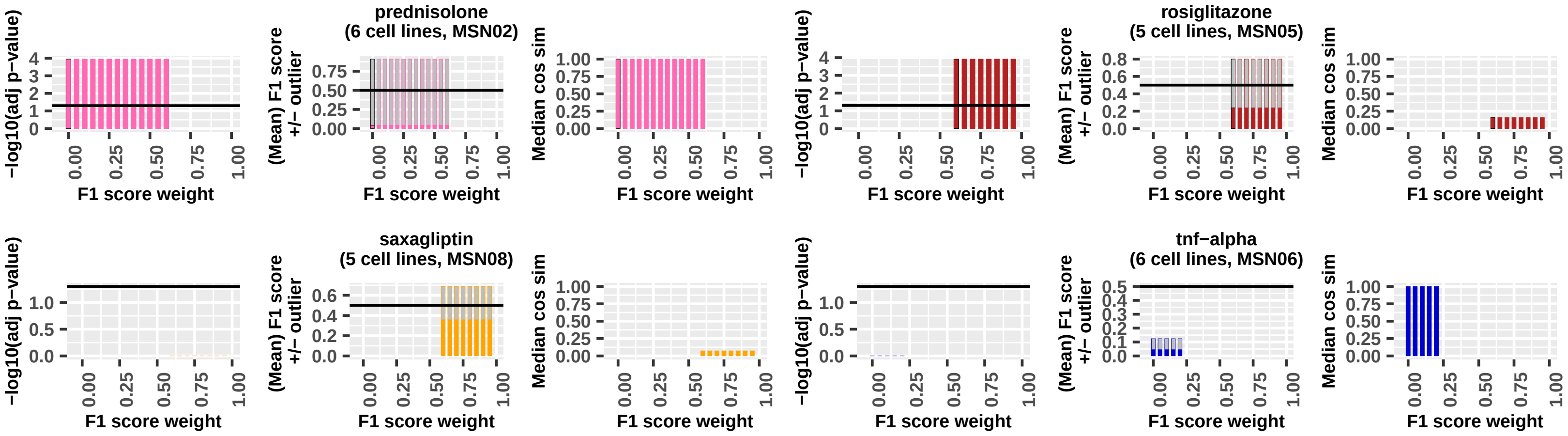


**Supplemental Figure 12**

**Supplemental Figure 12. Identification of outlier responses.** For each drug, we screened all 20 potential drug-selective subspaces (that are defined by different F1 score weights) for subspaces where one cell line/drug combination shows a different transcriptomic response to the drug of interest than all other cell line/drug combinations. As shown for an example in Suppl. Figure 10D, we calculated cell line/drug combination-specific F1 scores, using the same approach described above, except that the cell line/drug combination of interest has to be part of the corresponding cluster. Dixon's Q test of cell line/drug combination-specific F1 scores was used to identify outliers (adj. p-value = 0.05). Identified outliers were only accepted, if the mean F1 score of all non-outlier cell line/drug combinations was larger than 0.5 (empty bars in middle figure). We selected that subspace with the most significant adjusted p-value as the final drug-selective subspace (black frame). Mean F1 scores of all non-outlier cell line/drug combinations and decreasing F1 score weight were used as first and second tiebreakers, respectively. If no outlier was identified, we selected that subspace with the highest selection score based on an F1 score weight of 0.95.

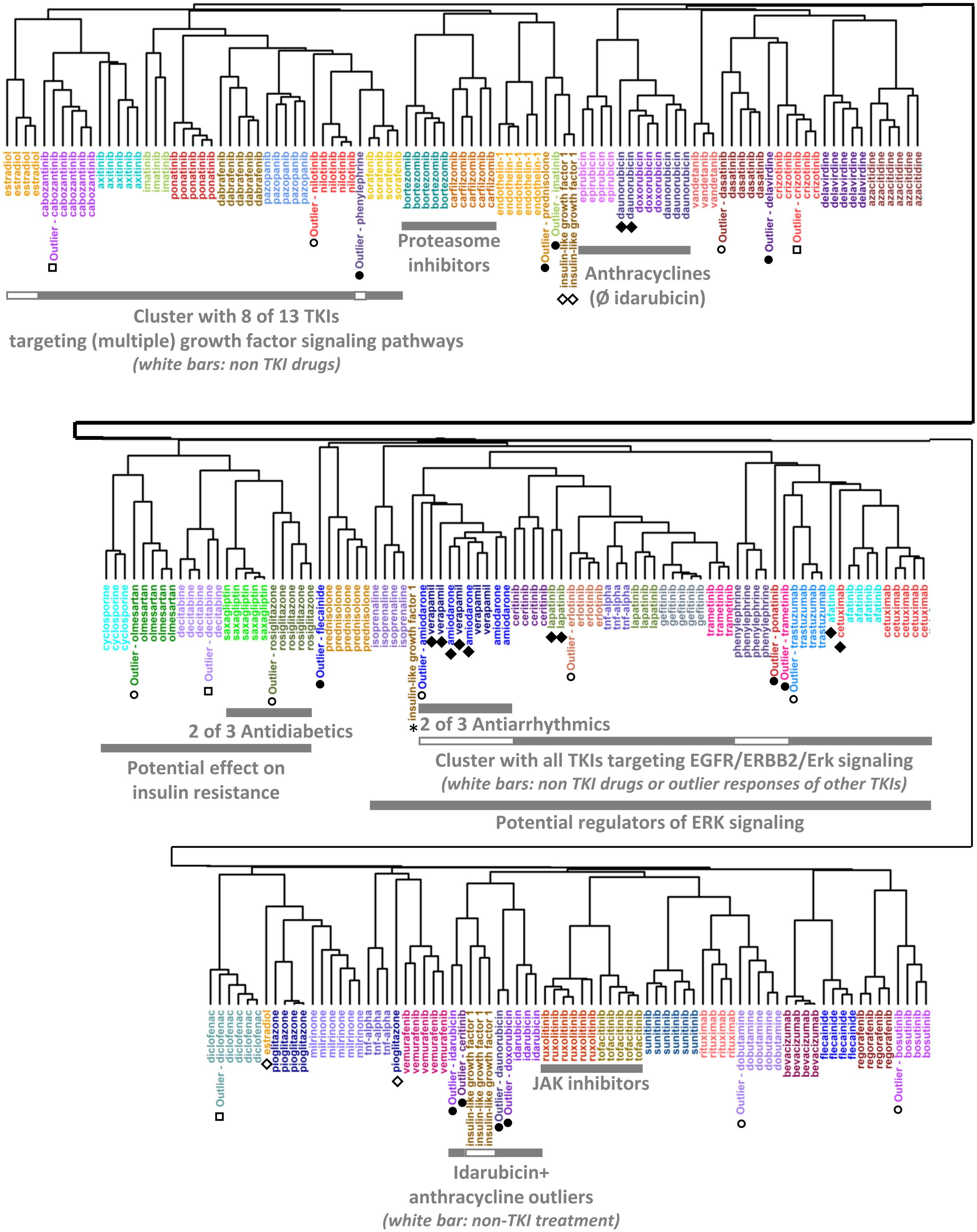


**Supplemental figure 13**

**Supplemental Figure 13. Clustering of merged drug-selective gene expression profiles.** Drug-selective gene expression profiles obtained after projection of the complete gene expression matrix into drug-selective subspaces were merged, followed by pairwise correlation and hierarchical clustering. Outlier cell line/drug combinations are labeled with ‘Outlier’. Open circles label outlier responses that show great outlier characteristics in this dendrogram as well, and do not cluster together with the other cell line/drug combinations treated with the same drugs. Closed circles label outlier responses with minor outlier characteristics in this dendrogram, i.e. ones that are grouped together with the cell line/drug combinations treated with the same drug in a larger cluster, but get separated from those after sub-clustering. Closed squares label outlier responses that are grouped together with the cell line/drug combinations treated with the same drug. Asterisks label cell line/drug combinations that do not cluster together with the other cell line/drug combinations treated with the same drug and were not identified as outlier responses. Brackets label clusters that are composed of cell line/drug combinations from different drugs with closely related potential mechanisms. Figure 1E shows the same dendrogram, drug labels and most bars grouping drugs with similar mechanisms.

A

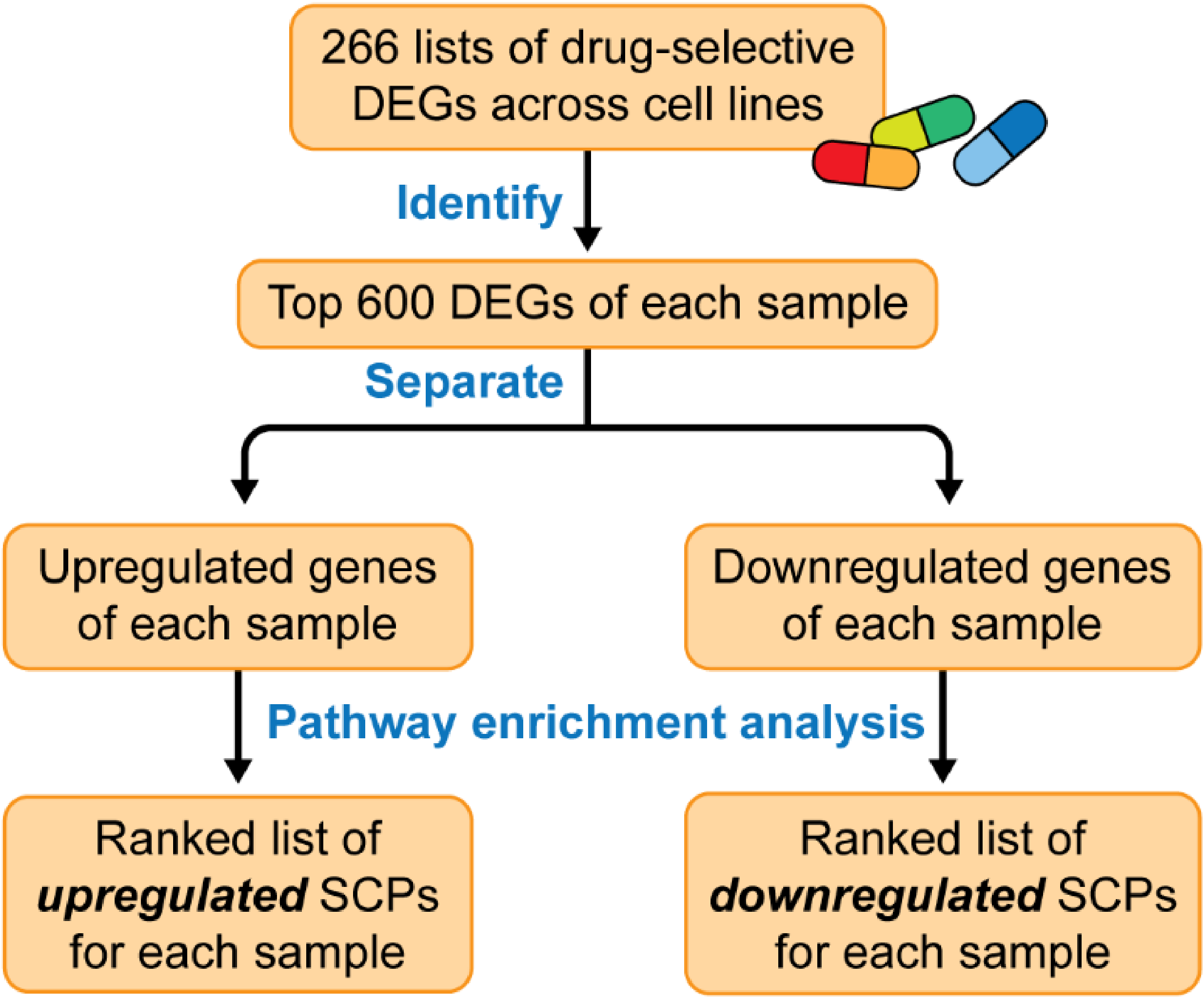


**Supplemental figure 14**

**Supplemental figure 14B/C/D/E: See additional PDFs**

**Supplemental Figure 14. Top Subcellular Processes predicted from complete gene expression profiles, after removal of first eigenarray and from drug-selective gene expression profiles. (A)** Complete, decomposed gene expression profiles and gene expression profiles after removal of the first eigenarray were subjected to pathway enrichment analysis using the Molecular Biology of the Cell Ontology and Fisher's Exact Test to identify up- and downregulated subcellular processes (SCPs). Significant up- or downregulated **(B)** level-1, **(C)** -2, **(D)** -3 and **(E)** -4 SCPs (p-value<=0.05) were separately ranked by significance for each cell line/drug combination. SCPs predicted for each drug are shown if they are among the top five ranked SCPs for at least one cell line. Numbers indicate ranks, '>' indicates that an SCP was not predicted or predicted with a rank above 99. Cell lines with identified outlier responses to treatment with a drug of interest are colored purple.

**A**

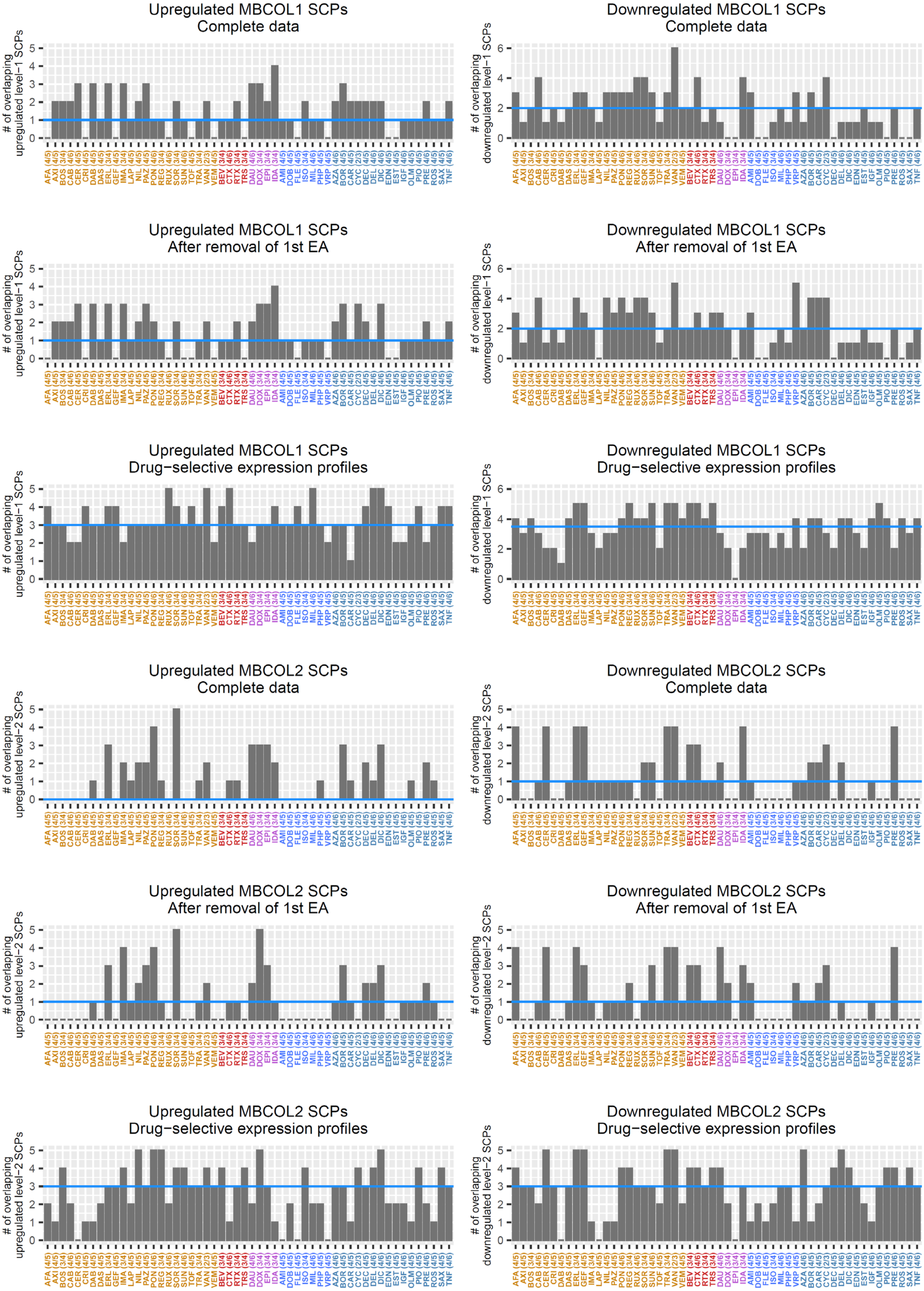


**Supplemental figure 15**

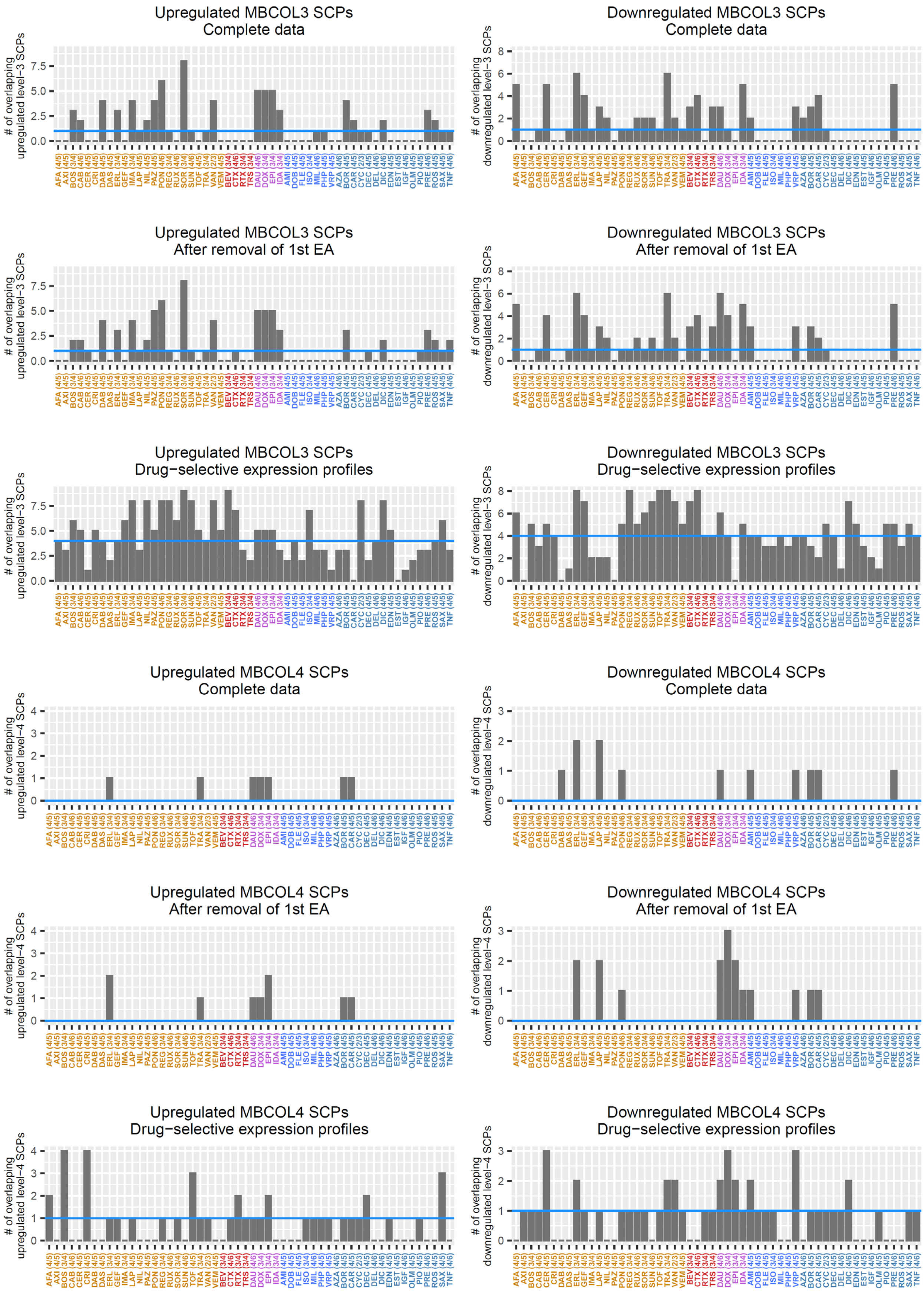


**Supplemental figure 15**

B

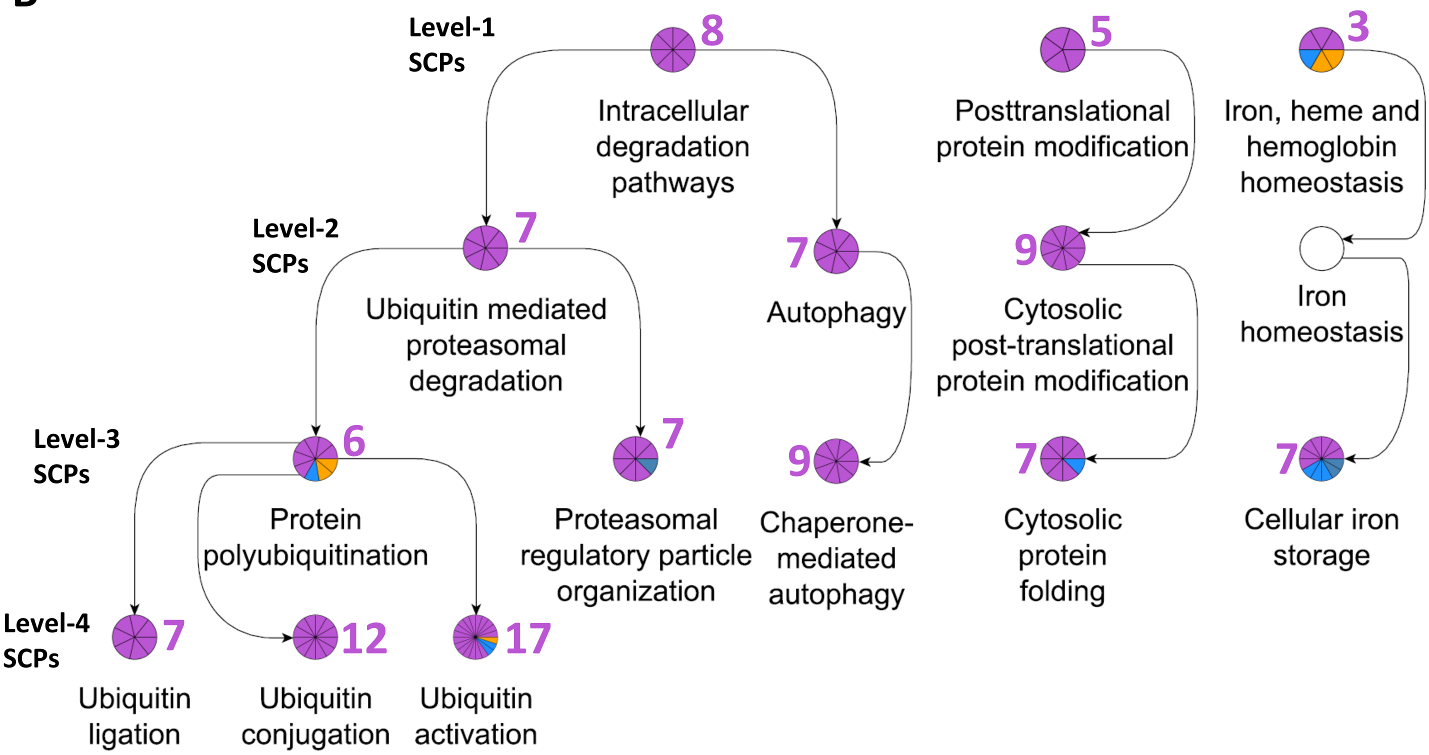


C

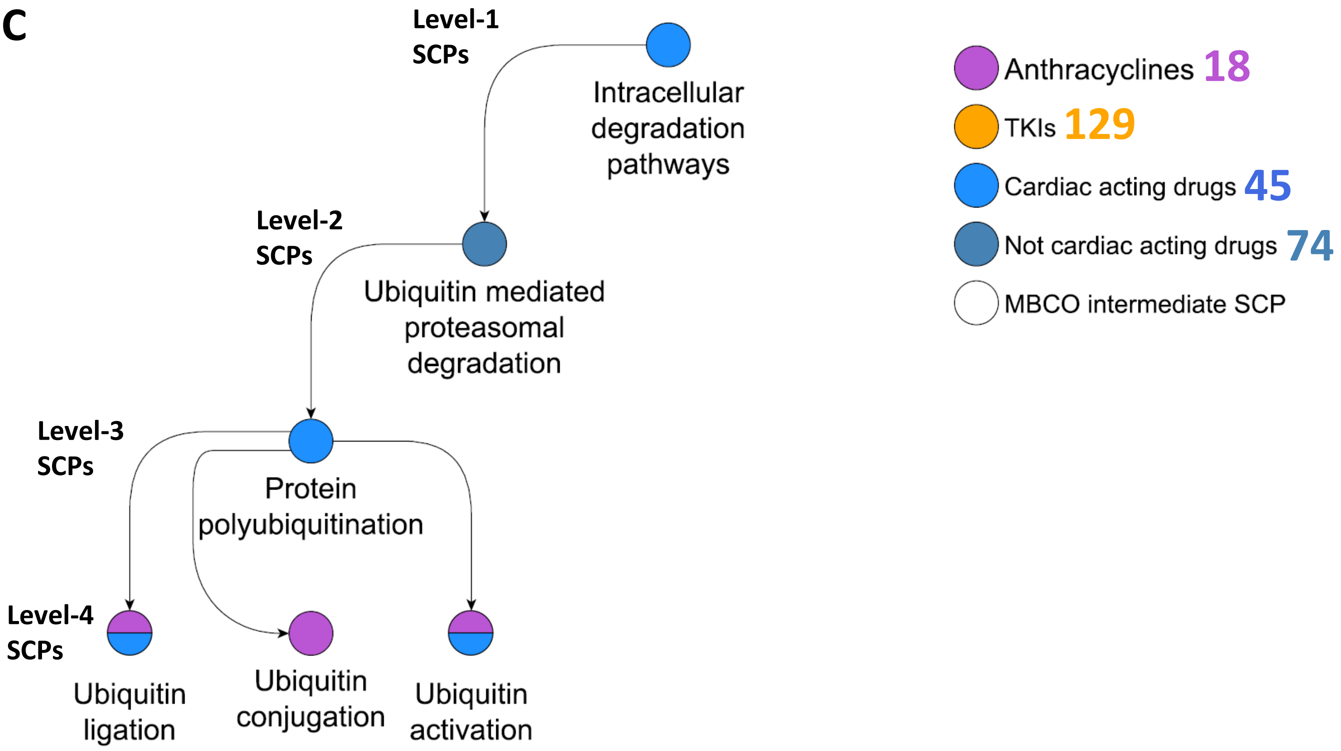


**Supplemental figure 15**

**Supplemental Figure 15. SVD decomposition increases the consistency of identified SCPs by the same drug across different cell lines and reveals potentially cardiotoxic SCPs. (A)** We analyzed, for each drug, how many up- or downregulated level-1, -2, -3 and -4 SCPs were predicted in at least 66% of all treated cell lines with a maximum rank of five, five, ten and five, respectively. The minimum numbers that equal or exceed 66% and the total numbers of treated cell lines are given as numerators and denominators, respectively, in brackets after the drug abbreviations. The blue line documents the median of overlapping SCP counts. The subfigures 'Down-' and 'Upregulated MBCOL3 SCPs, Drug-selective expression profiles' are also shown as main figures 2A/B. **(B)** The top five, five, ten and five level-1, -2, -3 and -4 SCPs predicted from downregulated genes in the drug-selective gene expression profiles were integrated into the MBCO hierarchy. Selected SCPs in parent-child relationships (arrows) are shown. Any drug that downregulates an SCP is added as a new pie slice colored according to the drug's class. Purple numbers next to the SCPs indicate the number of anthracycline-treated cell line/drug combinations for which the SCP was predicted (i.e., the number of purple slices). Numbers next to the drug classes in the legend indicate how many cell line/drug combinations were treated with drugs of that class in total. **(C)** The top five, five, ten and five level-1, -2, -3 and -4 SCPs predicted from downregulated genes in the complete gene expression profiles were integrated into the MBCO hierarchy, as described in B.

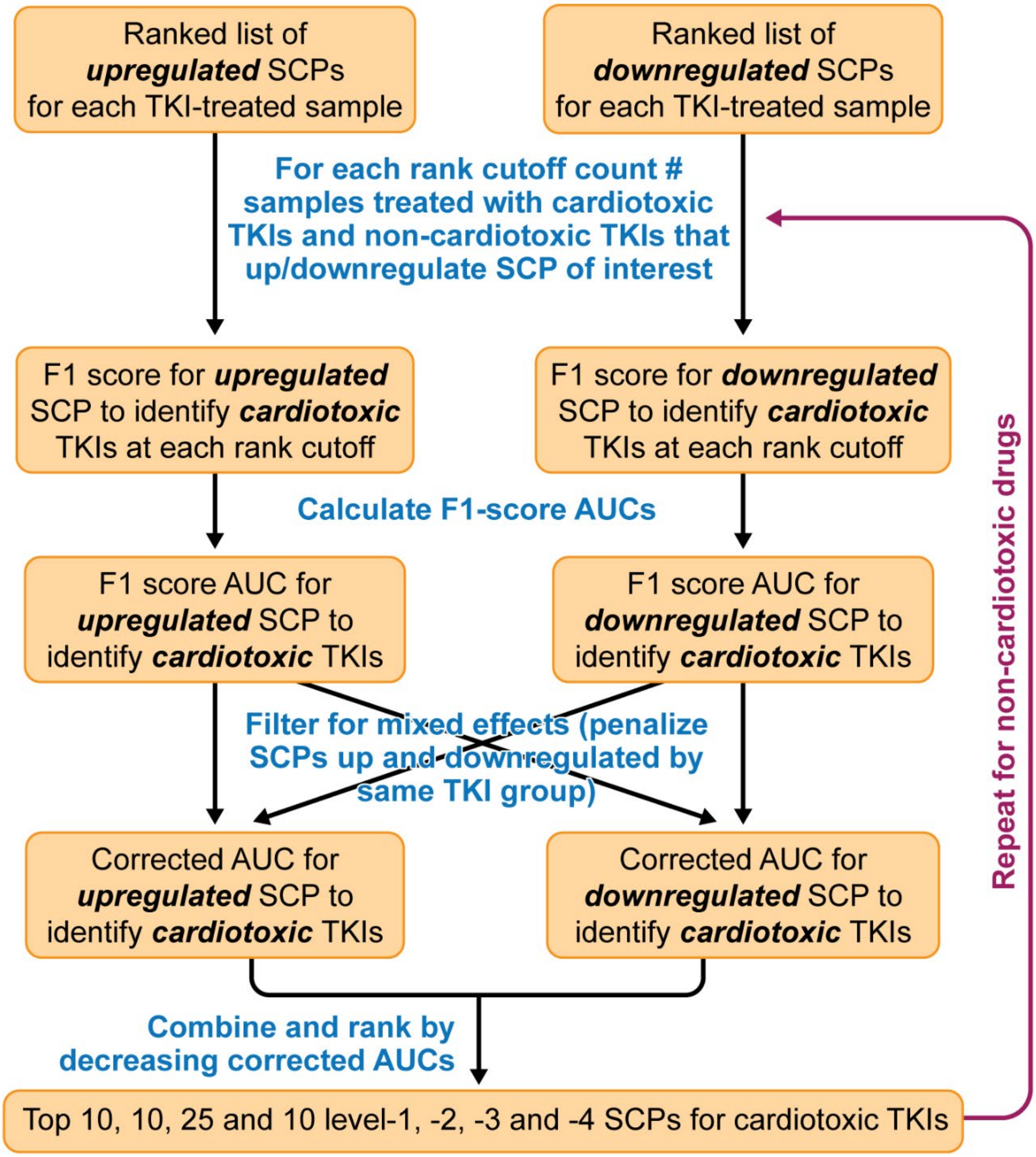


**Supplemental figure 16**

**Supplemental Figure 16. Identification of SCPs associated with cardiotoxic and non-cardiotoxic TKIs.** SCPs that were predicted from pathway enrichment analysis of drug-selective gene expression profiles were subjected to our computational pipeline that searches for SCPs that are indicative of a cardiotoxic or non-cardiotoxic TKI (i.e., small molecule tyrosine kinase inhibitors and monoclonal antibodies against tyrosine kinases). See methods for details.

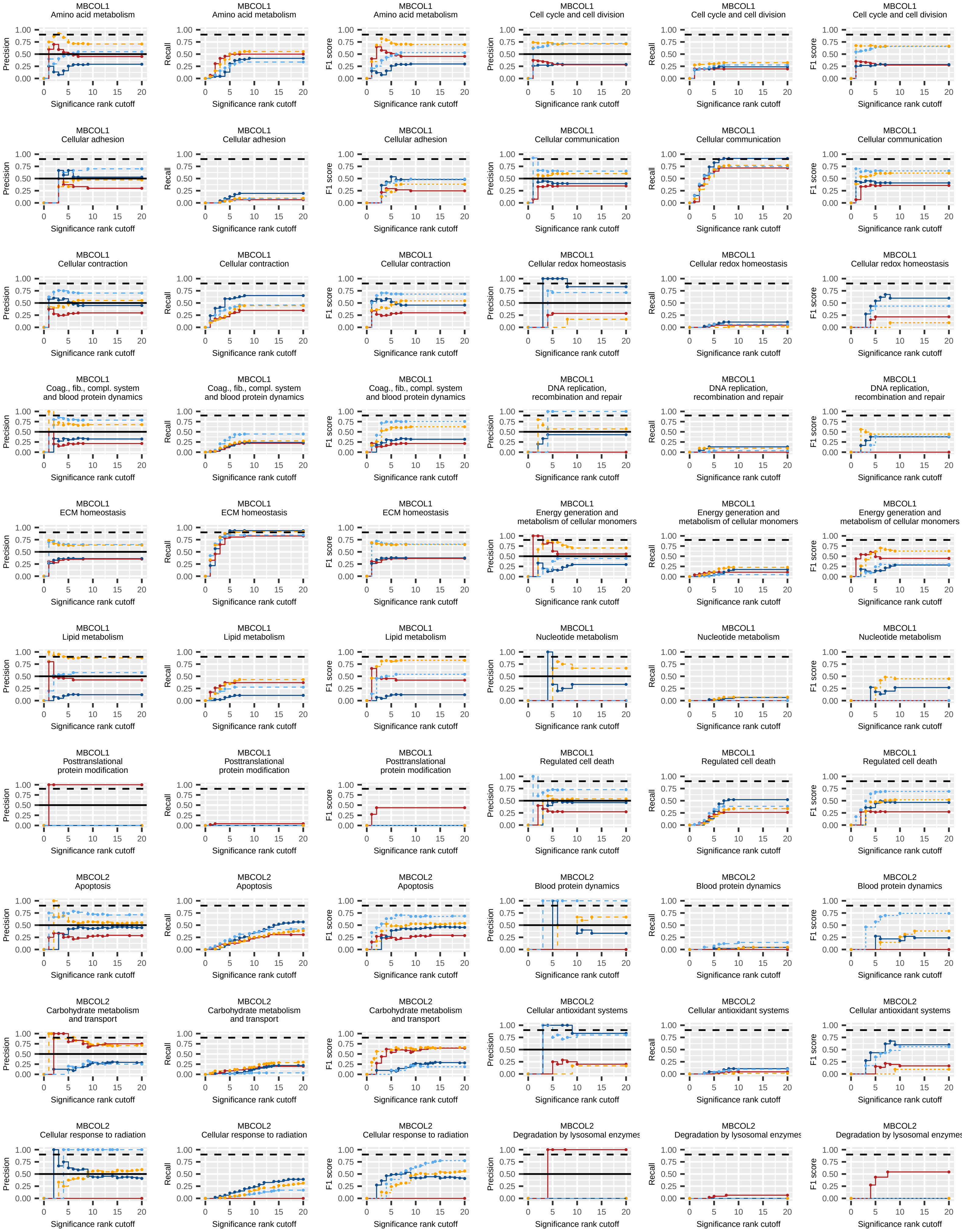


**Supplemental Figure 17**

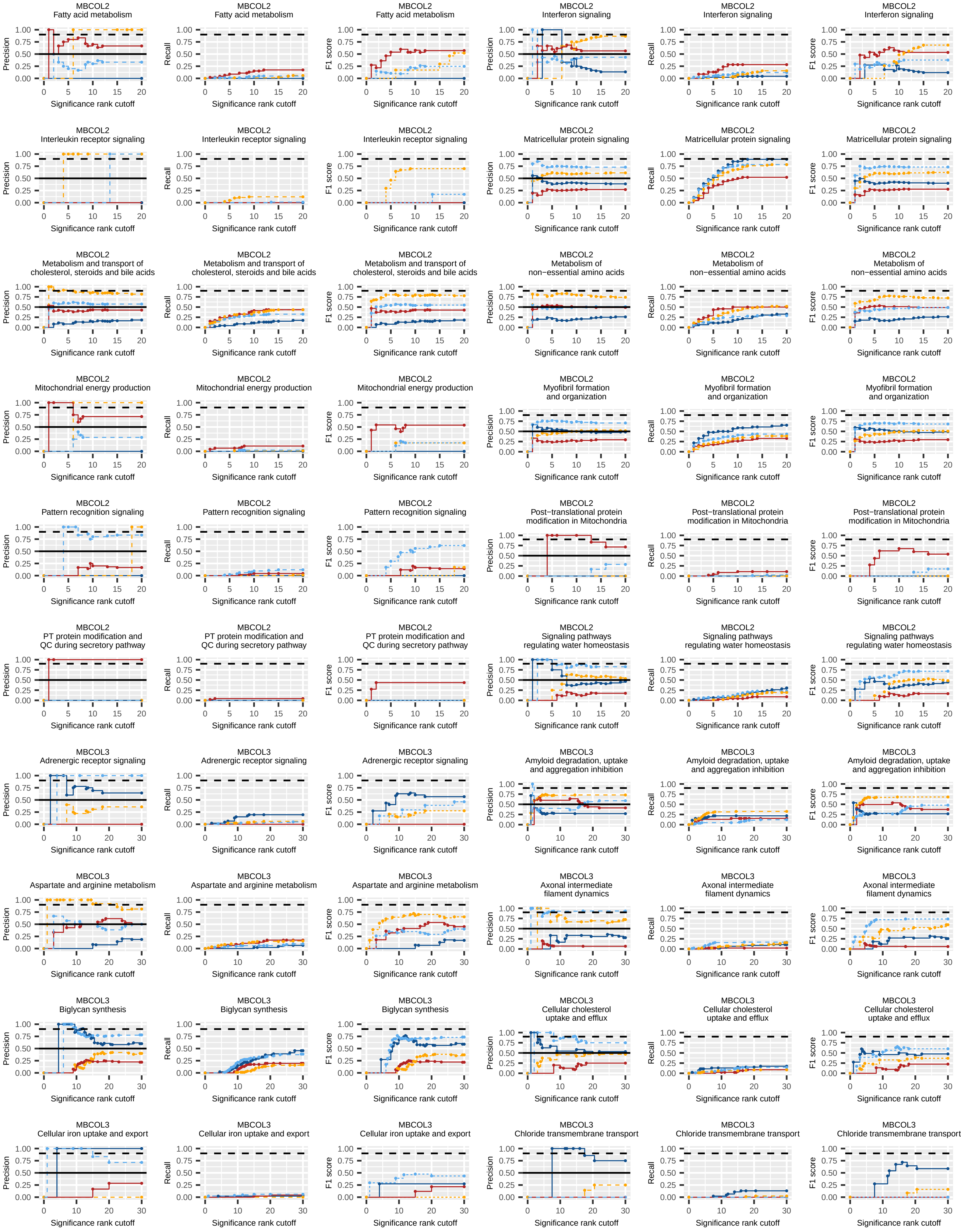


**Supplemental Figure 17**

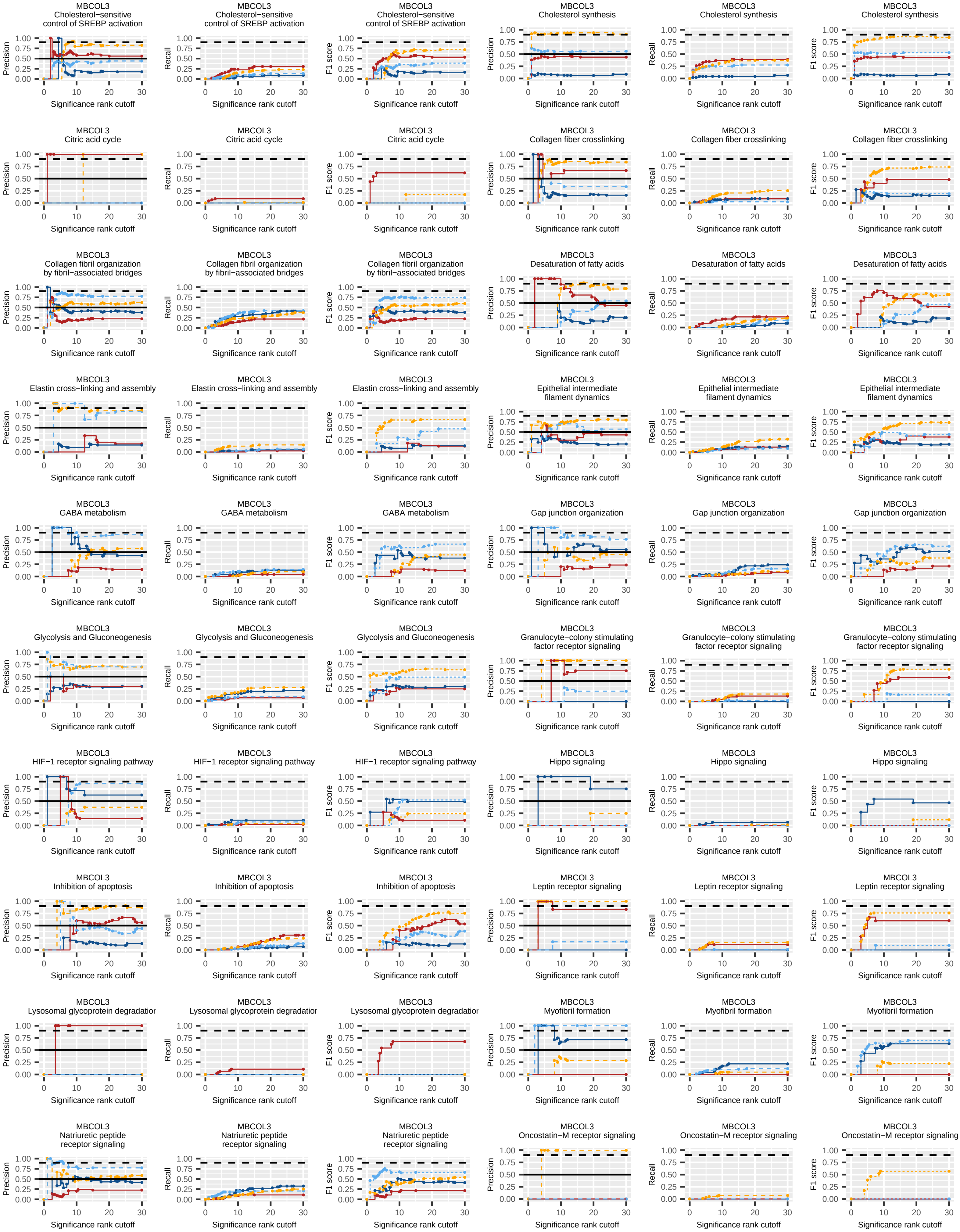


**Supplemental Figure 17**

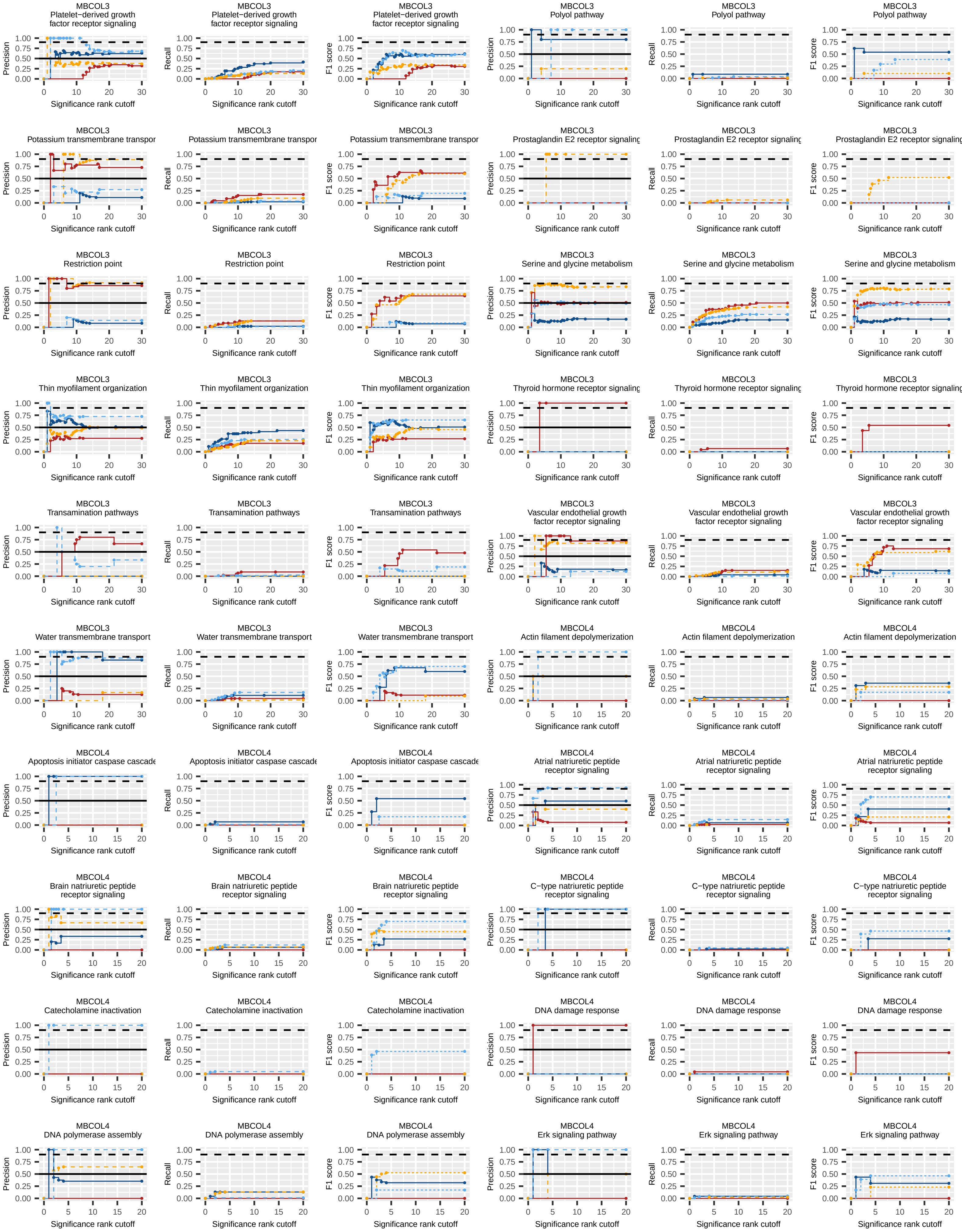


Supplemental Figure 17

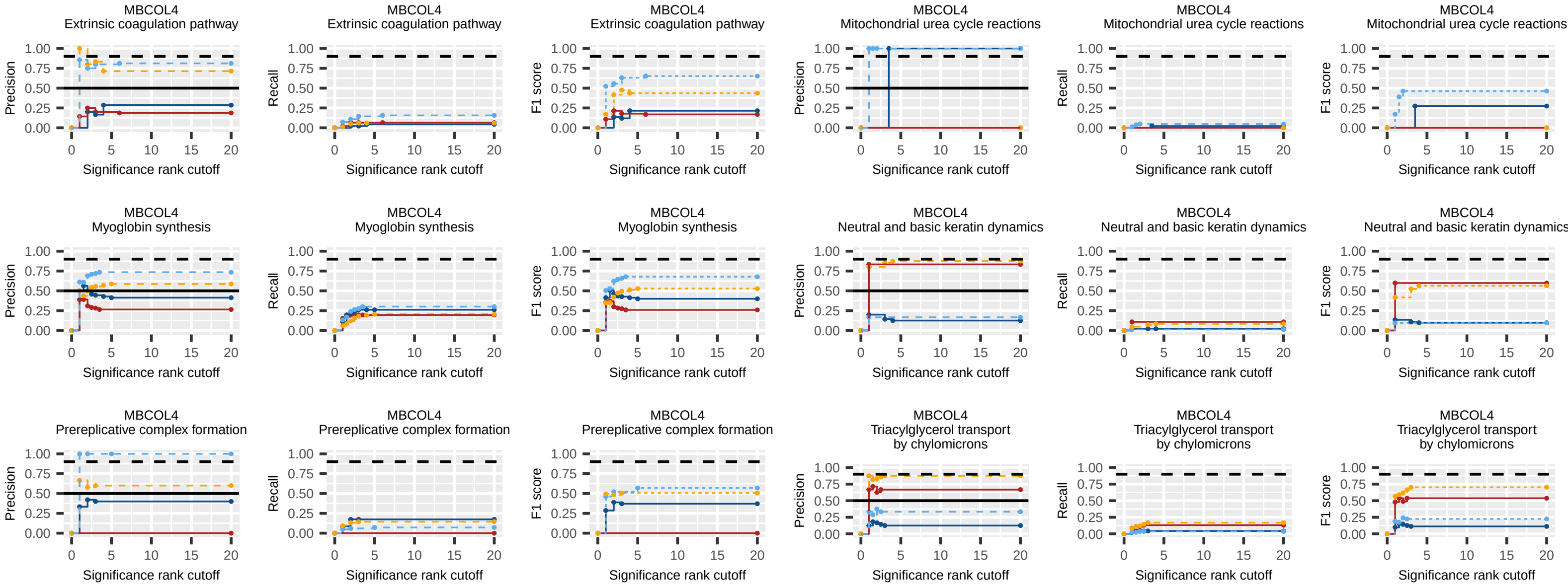


**Supplemental Figure 17**

**Supplemental Figure 17. F1 score and area under the curve statistics.** At each significance rank cutoff we counted how many cardiotoxic or non-cardiotoxic TKIs up- or down-regulate a particular SCP with a significance rank below or equal to the current cutoff. Results were used to calculate precision, recall and F1 score (beta = 0.2) of each SCP at each rank to be either up- or downregulated by either the cardiotoxic or non-cardiotoxic drugs. Shown are the results for the SCPs that our algorithm selected to be associated with a cardiotoxic or non-cardiotoxic response. See methods for description of the algorithm. Solid lines indicate results for the SCP, if up- (red) or downregulated (dark blue) by cardiotoxic TKIs, dashed lines indicate results for the SCP, if up- (light blue) or downregulated (orange) by non-cardiotoxic TKIs. Color combinations were selected to indicate if the higher (red, orange) or lower (dark blue, light blue) activity of an SCP favors a cardiotoxic response.

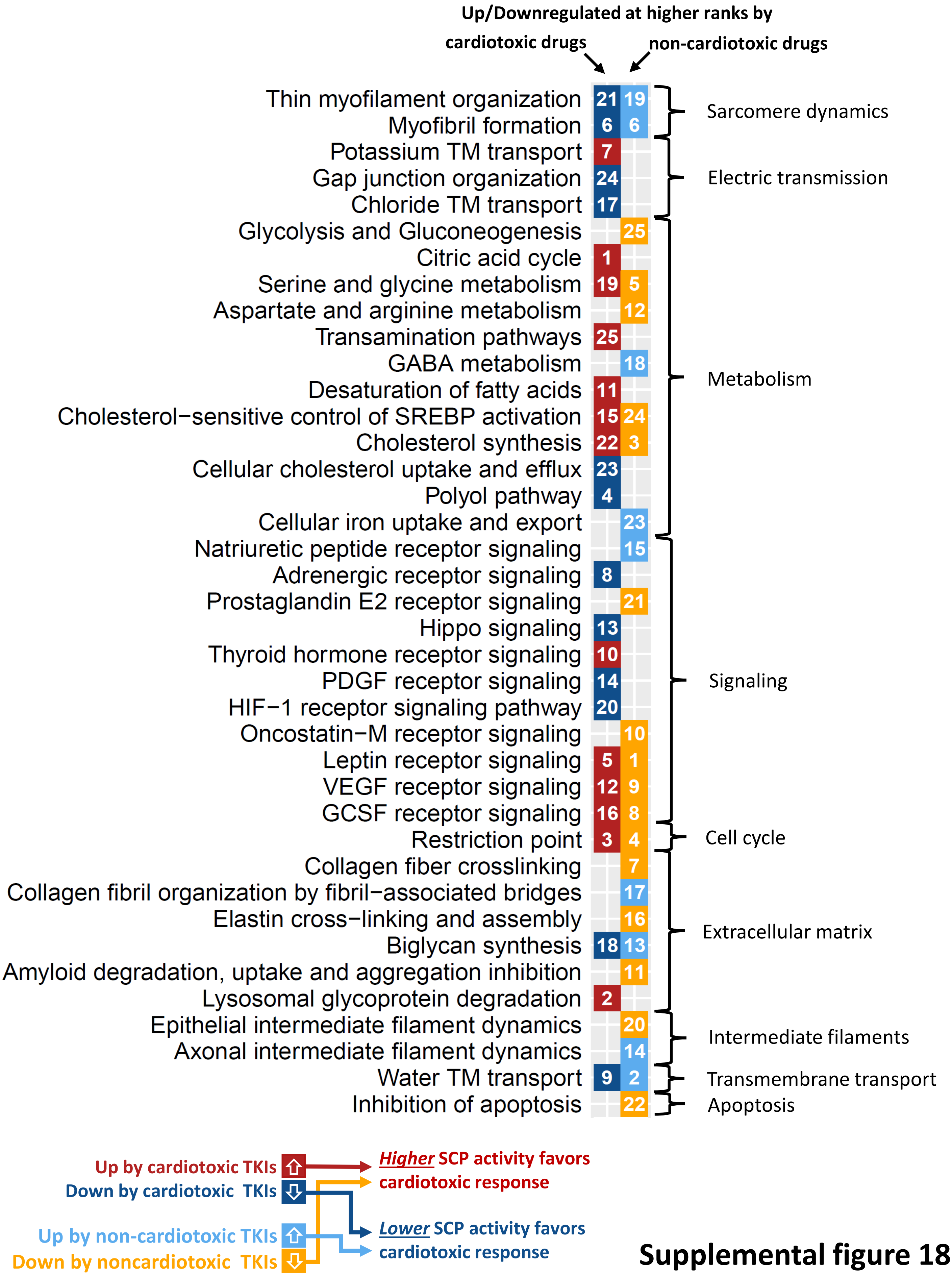


**Supplemental figure 18**

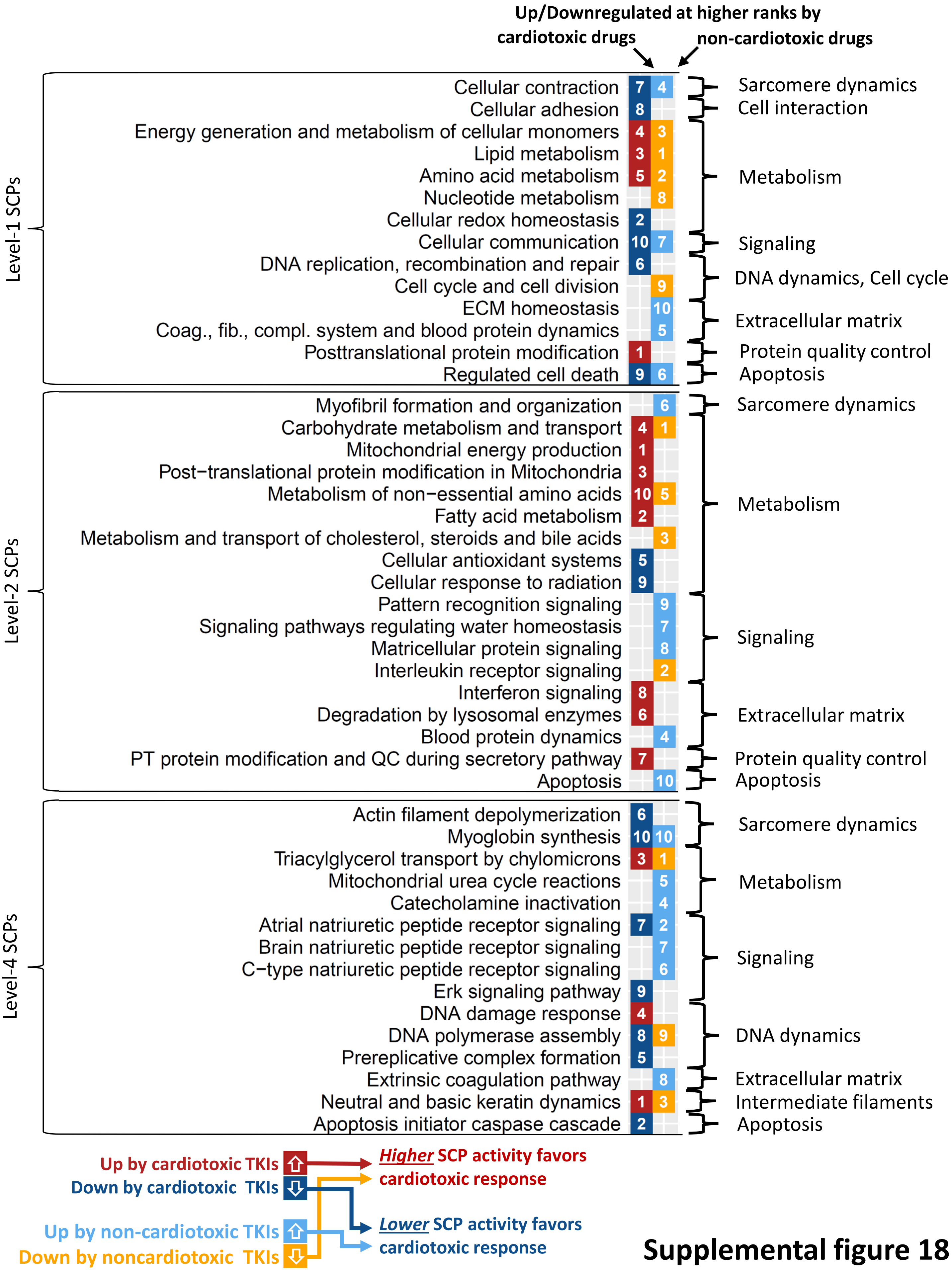


**Supplemental figure 18**

**Supplemental Figure 18. SCPs associated with cardiotoxic and non-cardiotoxic responses.** Predicted up- and downregulated subcellular processes (SCPs) of the same level were ranked by significance for each drug and cell line. We searched for those SCPs that are up- or downregulated at higher significance ranks by cardiotoxic or non-cardiotoxic TKIs. Identified SCPs were ranked by their selectivity for either cardiotoxic or non-cardiotoxic drugs (white numbers). To simplify our findings, we defined that SCPs that are upregulated by cardiotoxic drugs (red) or downregulated by non-cardiotoxic drugs (orange) are associated with a cardiotoxic response after upregulation or at baseline level, respectively. For these SCPs a higher activity favors a cardiotoxic response. Similarly, we defined that SCPs downregulated by cardiotoxic (dark blue) or upregulated by non-cardiotoxic drugs (light blue) are associated with a non-cardiotoxic response after downregulation or at baseline level, respectively. For these SCPs a lower activity favors a cardiotoxic response. Shown are the top 25, 10, 10 and 10 predicted level-3, -1, -2 and -4 SCPs for cardiotoxic and non-cardiotoxic TKIs. The level-3 SCPs that are up- or downregulated by cardiotoxic TKIs are also shown in main figure 2B.

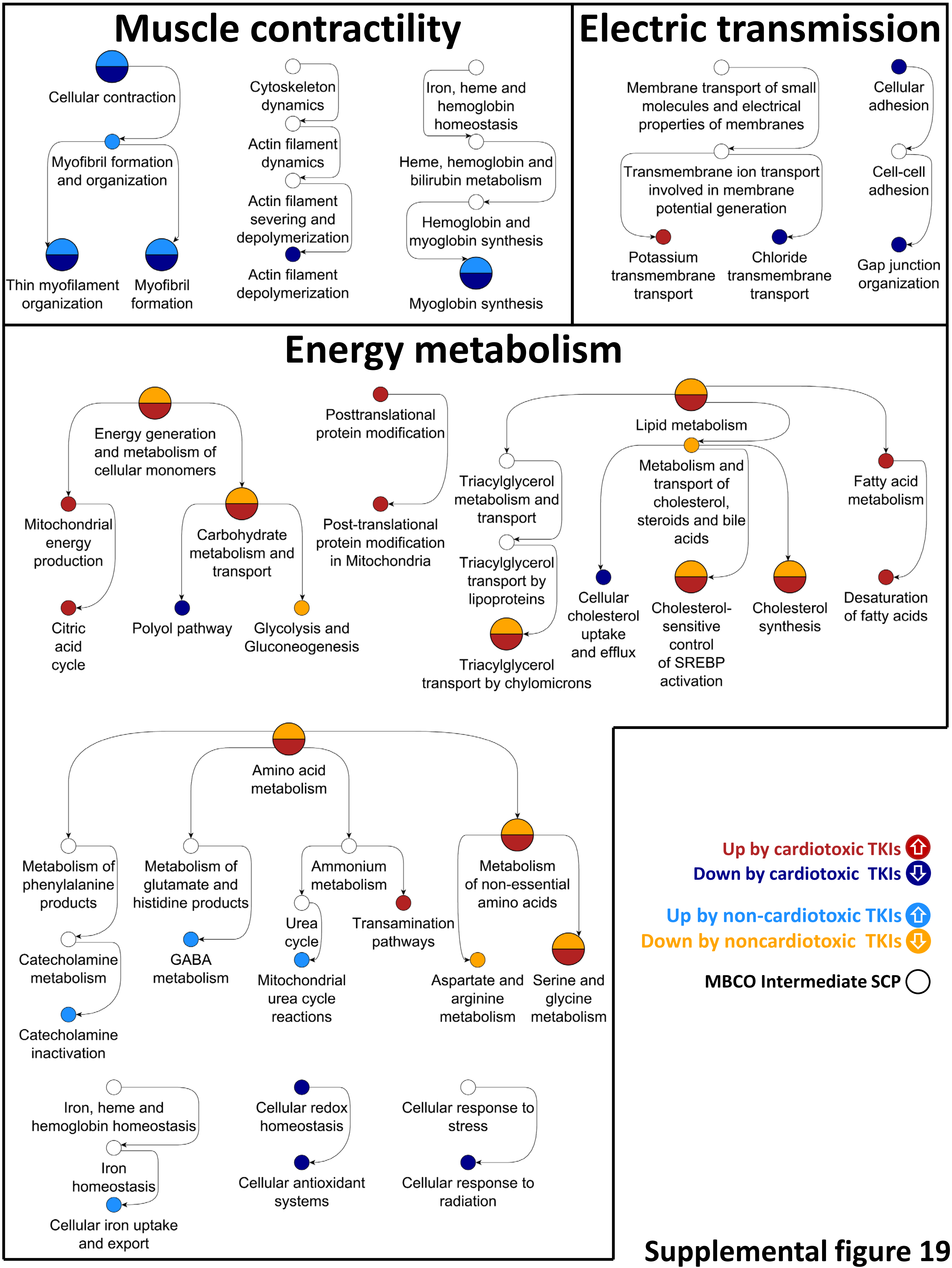


**Supplemental figure 19**

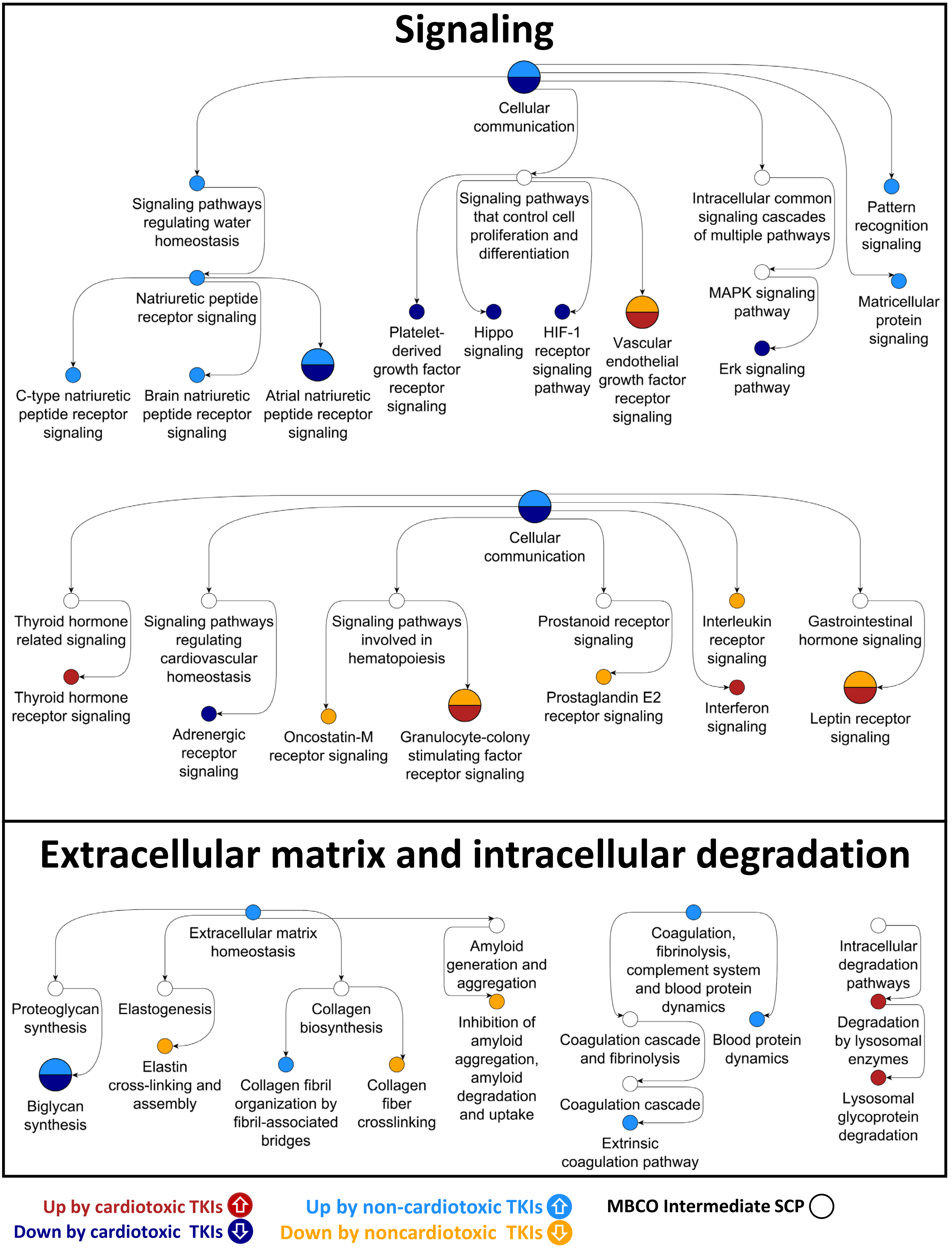


**Supplemental figure 19**

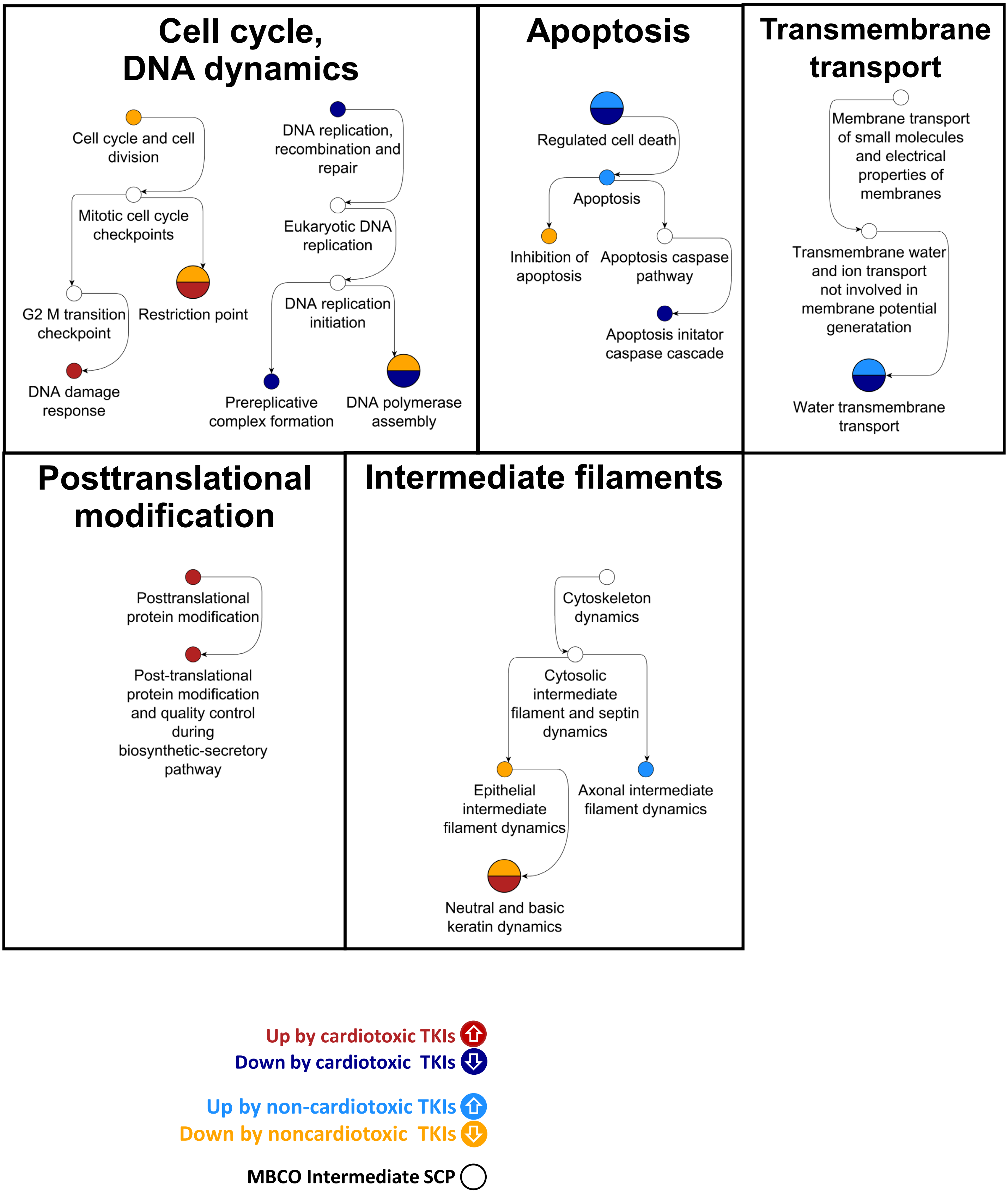


Supplemental figure 19

**Supplemental Figure 19. Integration of identified SCPs into the MBCO hierarchy.** Up- and downregulated SCPs associated with a cardiotoxic or non-cardiotoxic response were integrated into the MBCO hierarchy. Arrows point from parent to child SCPs. Each tree starts with a level-1 SCP and then consecutively connects it to predicted level-2, -3 and -4 SCPs. Non-predicted SCPs that are ancestors of predicted SCPs are in white. Red/orange: SCPs whose higher activity favors a cardiotoxic response, Dark blue/light blue: SCPs whose lower activity favors a cardiotoxic response. The muscle contractility SCPs, and selected SCPs involved in Energy metabolism are also shown in main figure 2C.

Level-3 SCPs

| | dabrafenib (5x) | lapatinib (5x) | pazopanib (5x) | ponatinib (6x) | sorafenib (4x) | sunitinib (6x) | trametinib (4x) | vandetanib (3x) | bevacizumab (4x) | trastuzumab (4x) | # variants |
|---|---|---|---|---|---|---|---|---|---|---|---|
| Thin myofilament organization | 5 | 7;12<br>17 | | 1;1;1<br>1;1 | | 3;3;4<br>7;7;7 | 22 | | 3;3<br>5;7 | | 31 |
| Myofibril formation | 10<br>10 | | 9 | 3;4;8<br>12;12 | 8 | | | | 8 | | 53 |
| Potassium transmembrane transport | | 7<br>10 | | | | | 2;3<br>9 | | 17 | 7<br>10 | 123 |
| Gap junction organization | 24 | | 1;3 | 15;16<br>17 | | | | 9 | 13;14<br>14;15 | | 72 |
| Chloride transmembrane transport | | | | 14;14<br>16 | | 12<br>12 | | 8 | | | 86 |
| Citric acid cycle | | | 1;1<br>2;3 | | | | | | | | 60 |
| Serine and glycine metabolism | 1;1;1<br>2;3 | | 1;1;3<br>4;5 | 5;5;5<br>5;6 | 2;11<br>19;21 | 12 | 4 | 12<br>14 | | | 47 |
| Transamination pathways | | | 6;10<br>10;11 | | | | | | | | 30 |
| Desaturation of fatty acids | 3 | 3;4 | 2;7<br>10 | | 7;8<br>14;14 | | | | | | 30 |
| Cholesterol−sensitive control of SREBP activation | | 2;3;4<br>5;6 | 4;5;8<br>8;8 | 15<br>15 | 14 | 7 | | | | | 21 |
| Cholesterol synthesis | 1;4<br>5 | 1;1;1<br>1;1 | 2;2<br>13 | | 2;4 | 1;1<br>3;8 | | 2 | | | 63 |
| Cellular cholesterol uptake and efflux | 4 | 4<br>22 | | | | 3;3<br>7 | | 1<br>15 | | | 118 |
| Polyol pathway | | | | | | | | | 1;1<br>1;1 | | 16 |
| Adrenergic receptor signaling | 9 | 9 | | 10;13<br>14 | | | | | 2;6<br>7;9 | | 134 |
| Hippo signaling | | | 3;5<br>7 | | | | | | | | 137 |
| Thyroid hormone receptor signaling | | | | | | | | | 4;4<br>6 | | 30 |
| Platelet−derived growth factor receptor signaling | | 20 | 9 | 4;6;6<br>6;13 | 5;6<br>10;15 | | 15<br>30 | 9<br>13 | | 2;4<br>5;11 | 57 |
| HIF−1 receptor signaling pathway | 8;8 | | 1;6<br>6 | | | | | | | | 42 |
| Leptin receptor signaling | | 3;5<br>5;6 | | | | | | | | 4 | 20 |
| Vascular endothelial growth factor receptor signaling | 7;7<br>10 | | 6;10<br>10;11 | | | | | | | | 46 |
| Granulocyte−colony stimulating factor receptor signaling | | 7;8<br>11;12 | | | | | | | 15 | 11 | 29 |
| Restriction point | | | | | | 2;3;4<br>6;11 | | 9 | | | 39 |
| Biglycan synthesis | 12 | 18;19<br>21;28 | 15 | 9;10;10<br>11;12;14 | 8 | | 26<br>28 | 5;8<br>10 | | 15;17<br>19 | 3 |
| Lysosomal glycoprotein degradation | | | 4;4;5<br>8;8 | | | | | | | | 21 |
| Water transmembrane transport | 6;7 | | 4 | 6;9 | | | | | | | 44 |

Up by cardiotoxic TKIs Down by cardiotoxic TKIs

**Supplemental figure 20**

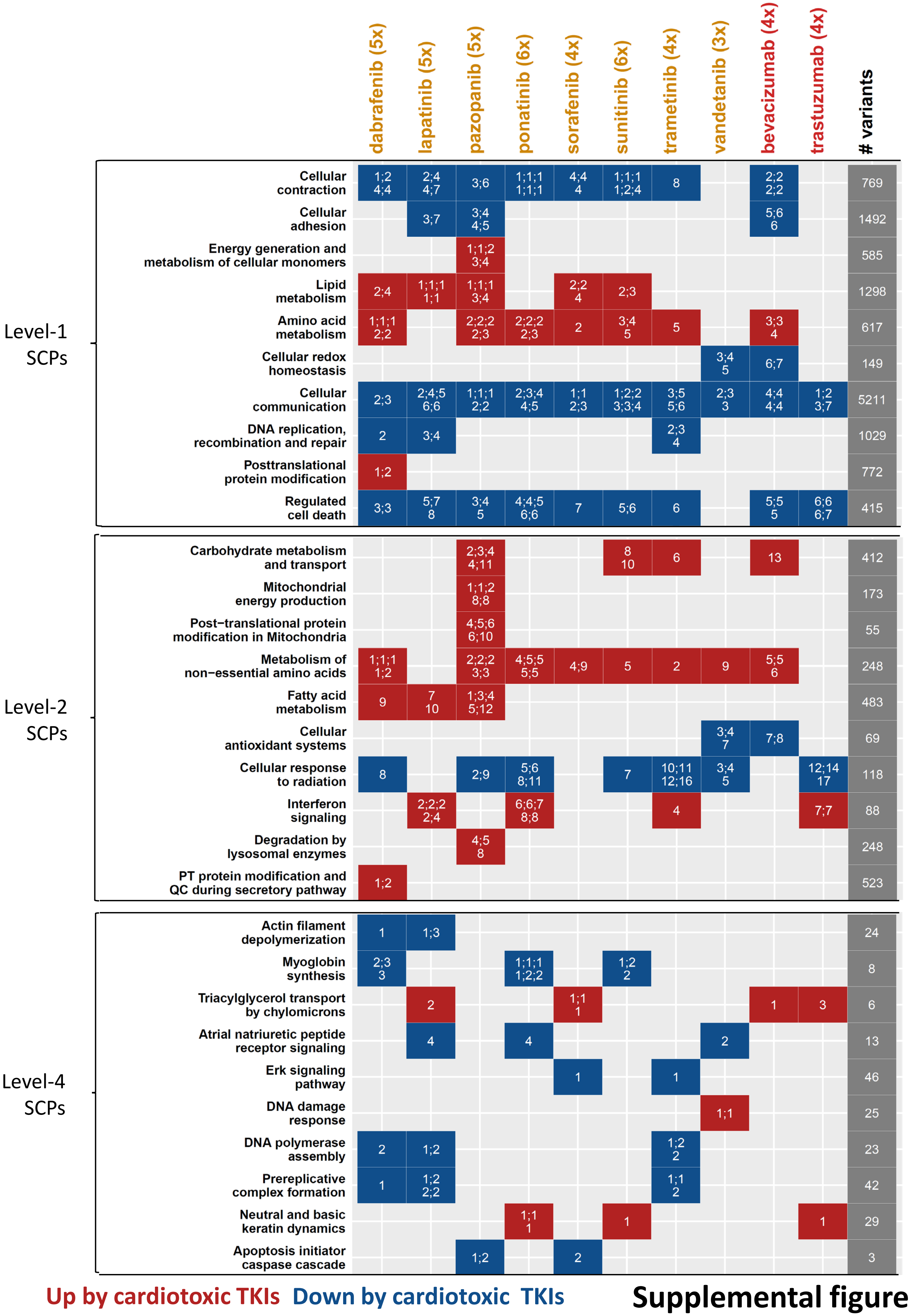


Supplemental figure 20

**Supplemental Figure 20. Regulation of identified SCPs by each cardiotoxic TKI.** SCPs that were up- or downregulated at higher ranks by cardiotoxic TKIs were mapped back to the individual drugs that upregulated (red) or repressed (dark blue) them. Numbers indicate significance ranks as shown in Suppl. Figures 14B/C/D/E for level-1, -2, -3 and -4 SCPs. Only ranks that were below the maximum rank cutoff in our F1 score and AUC statistics are shown (20, 20, 30, 20 for level-1, -2, -3 and -4 SCPs). Numbers in parentheses after drug labels indicate total numbers of treated cell lines for each TKI. Number of genomic variants that are underrepresented in the general population and map to SCP genes are shown in the last column. See methods for details. Note that this representation gives an estimation of the recall for each SCP, but does not allow conclusions about the precision that was favored during identification of SCPs associated with a cardiotoxic response. Drug labels of small molecule kinase inhibitors and monoclonal antibodies are colored orange and red, respectively.

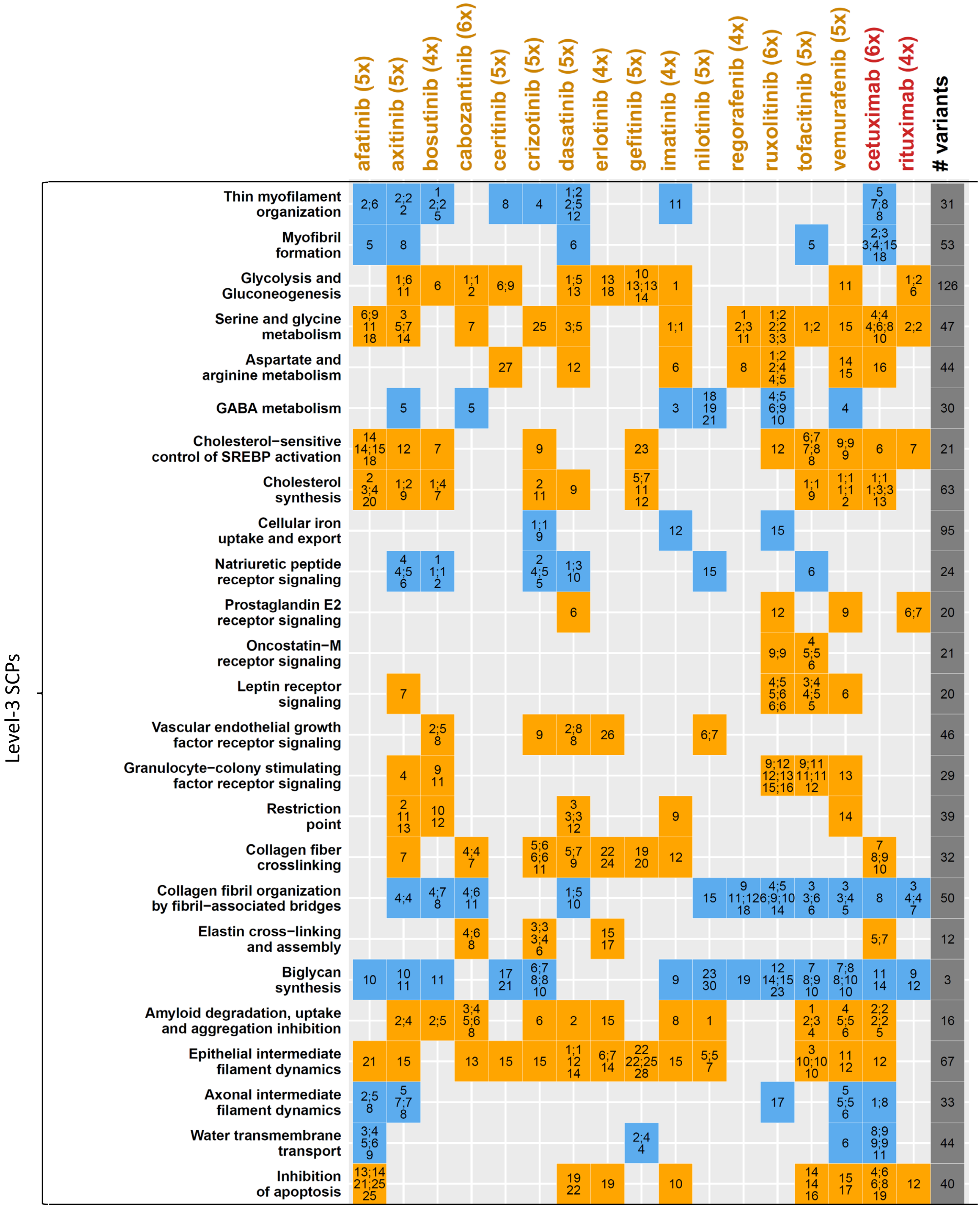


Up by non-cardiotoxic TKIs

Down by noncardiotoxic TKIs

**Supplemental figure 21**

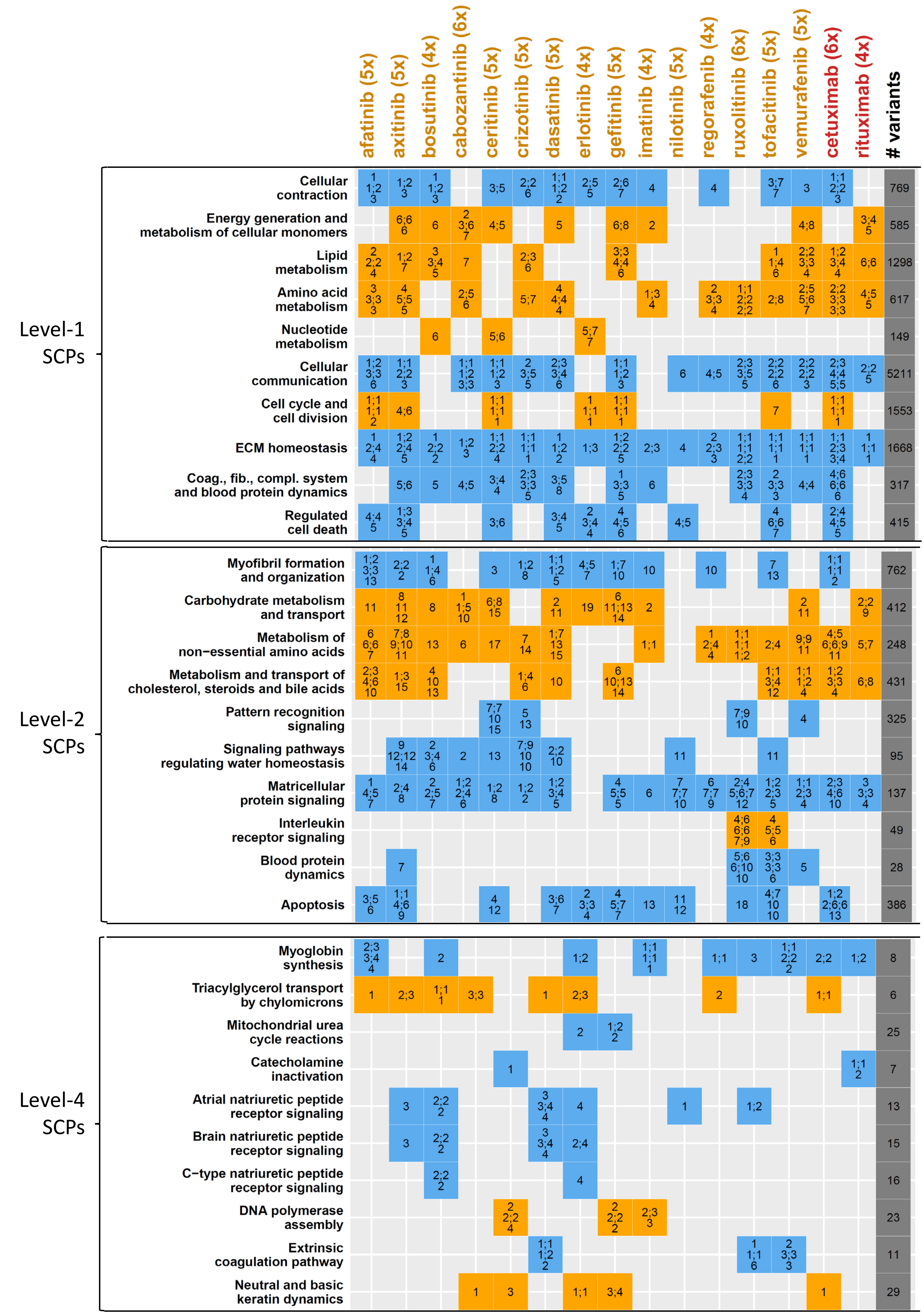


Up by non-cardiotoxic TKIs    Down by noncardiotoxic TKIs

**Supplemental figure 21**

**Supplemental Figure 21. Regulation of identified SCPs by each non-cardiotoxic TKI.** SCPs that were up- or downregulated at higher ranks by non-cardiotoxic TKIs were mapped back to the individual drugs that down- (orange) or upregulated (light blue) them. Numbers indicate significance ranks as shown in Suppl. Figures 14B/C/D/E for level-1, -2, -3 and -4 SCPs. Only ranks that were below the maximum rank cutoff in our F1 score and AUC statistics are shown (20, 20, 30, 20 for level-1, -2, -3 and -4 SCPs). Numbers after drug indicates total numbers of treated cell lines for each TKI. Number of genomic variants that are underrepresented in the general population and map to SCP genes are shown in the last column. See methods for details. Note that this representation gives an estimation of the recall for each SCP, but does not allow conclusions about the precision that was favored during identification of SCPs associated with a non-cardiotoxic response. Drug labels of small molecule kinase inhibitors and monoclonal antibodies are colored orange and red, respectively.

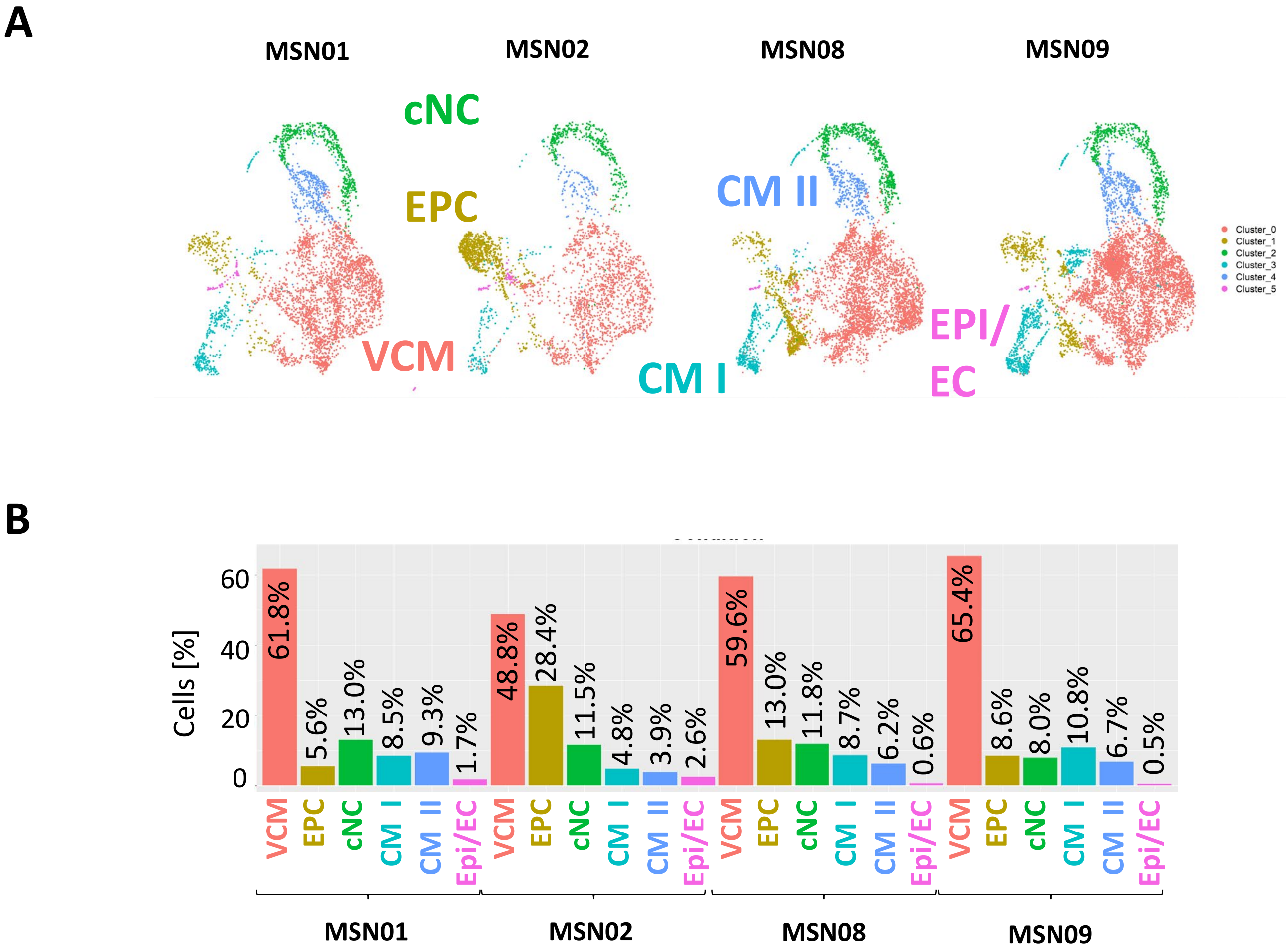


**Supplemental figure 22**

C

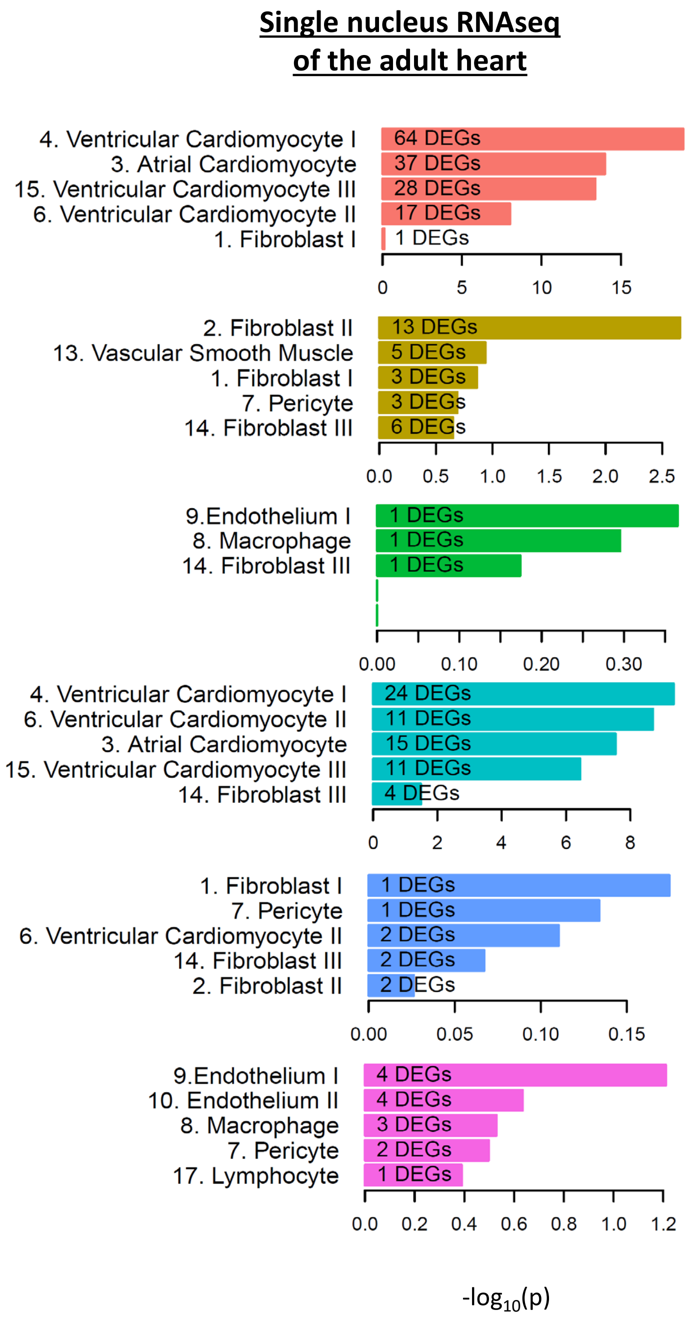


**Supplemental figure 22**

D

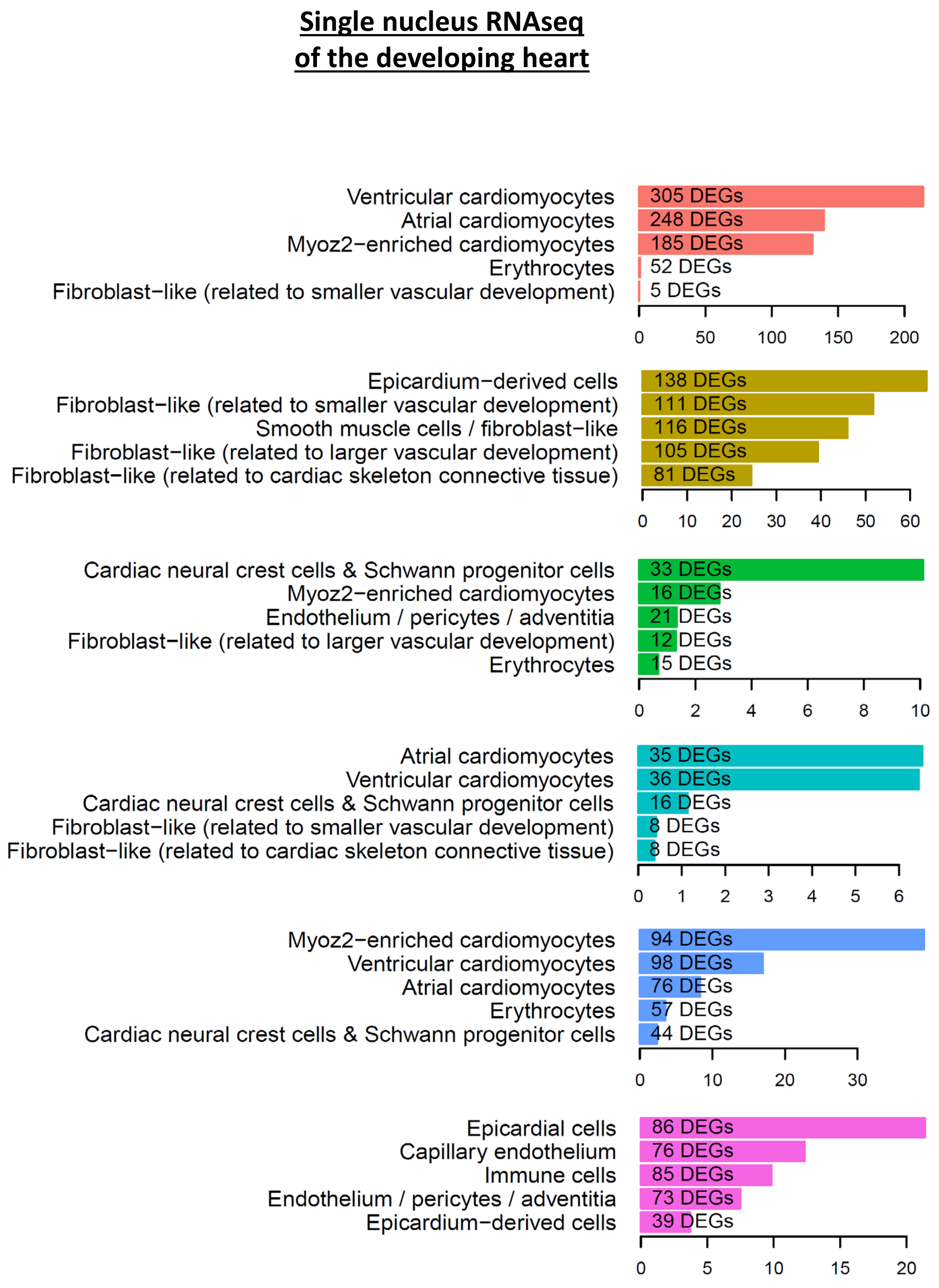


**Supplemental figure 22**

**Supplemental Figure 22. Single cell RNAseq identifies five different cellular subtypes. (A)** Single cell RNAseq analysis of four of our six different hiPSC-derived cardiomyocyte cell lines identifies one ventricular cardiomyocyte (VCM) subtype, two additional cardiomyocyte subtypes (CM I and CM II), one epicardial cell derived subtype (EPC), one cardiac neural crest (cNC) subtype and one epicardial (EPI) or endothelial (EC) cell subtype. **(B)** Cell counts of the identified subtypes document that most of our cells are ventricular cardiomyoyctes in all four cell lines. A and B are updated versions of two supplemental figures in our previous publication [16]. **(C)** Subtype-specific marker genes were subjected to pathway enrichment analysis using cell type marker genes identified from single nucleus RNAseq of the human adult heart or **(D)** cell type marker genes identified from single nucleus RNAseq of the human fetal heart. Enrichment results were used for cell type annotations shown in (A).

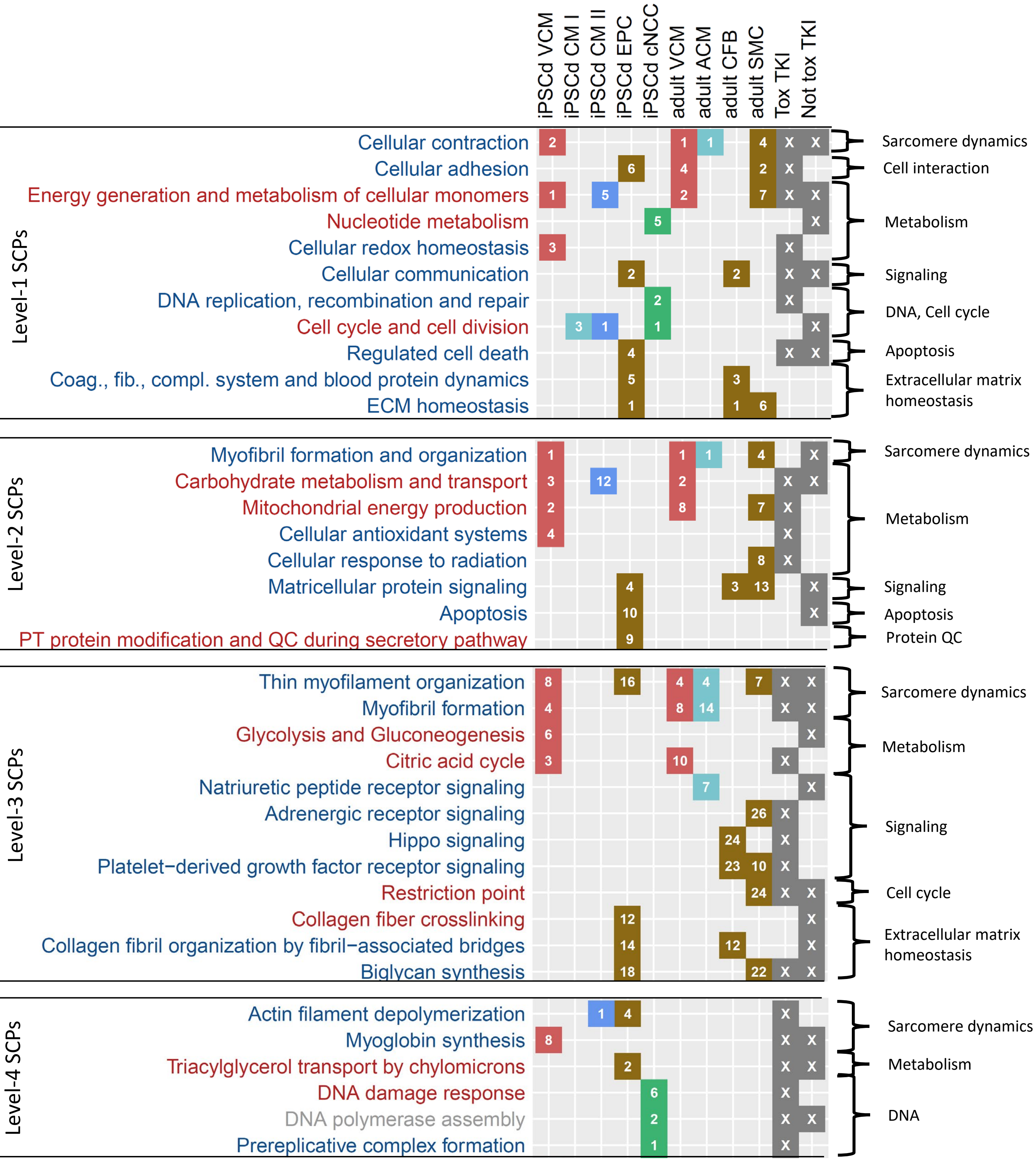


**Supplemental figure 23**

**Supplemental Figure 23. SCPs can be mapped to cellular cardiac cell types.** Subtype marker genes identified by single cell RNAseq analysis of our four cell lines [16] were subjected to enrichment analysis using MBCO. Significant SCPs ($p \leq 0.05$) were ranked by significance (numbers in the diagram). Similarly, we subjected cell type marker genes obtained from single nucleus RNAseq of the adult human heart [28] to pathway enrichment analysis. The last two columns indicate if the SCP was identified based on cardiotoxic and/or non-cardiotoxic TKIs. SCPs whose higher and lower activity is associated with a cardiotoxic response are in red and blue, respectively. Results for level-3 SCPs identified based on cardiotoxic TKIs are also shown in main figure 3A.

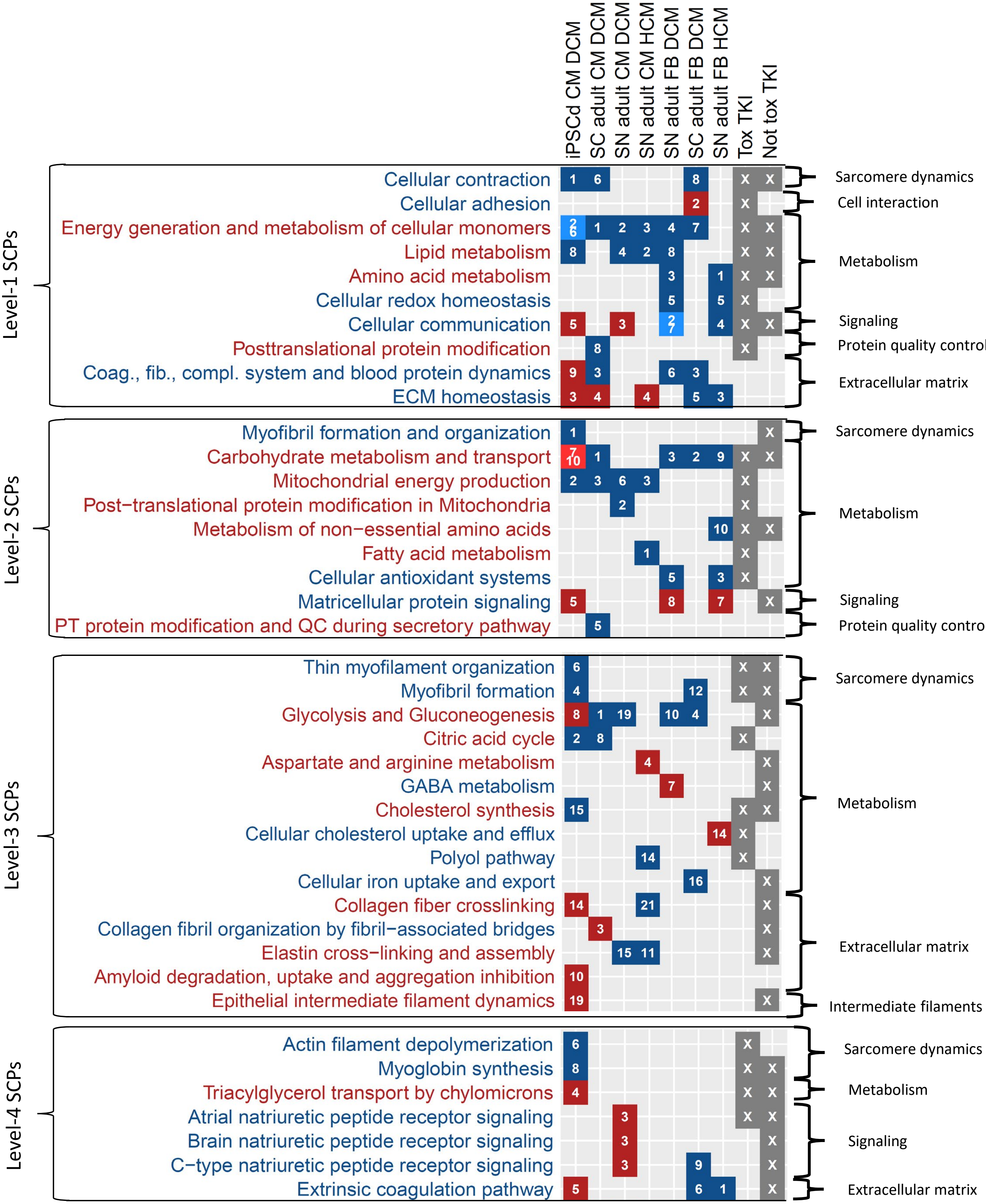


**Supplemental figure 24**

**Supplemental Figure 24. SCPs indicative of TKI-induced cardiotoxicity partially overlap with prior knowledge obtained from single cell and single nucleus RNAseq studies.** Differentially expressed genes in heart cells from patients with Dilated or Hypertrophic Cardiomyopathy (DCM or HCM, respectively) obtained by single cell (SC) [13] and nucleus (SN) [14] RNAseq as well as in hiPSC-derived cardiomyocytes from infant DCM patients (GSE184899) were subjected to pathway enrichment analysis using MBCO. Significantly up- or downregulated SCPs of each cell type ($p \leq 0.05$) were ranked by significance (numbers in the diagram). Only SCPs that overlap with SCPs for which higher (red) or lower (blue) activity favors a cardiotoxic response are shown. The last two columns indicate whether the SCP was identified based on cardiotoxic and/or non-cardiotoxic TKIs. iPSCd: iPSC-derived, CM: cardiomyocyte, FB: Cardiac fibroblast. Results for level-3 SCPs that were predicted based on cardiotoxic TKIs are also shown in main Figure 3B.

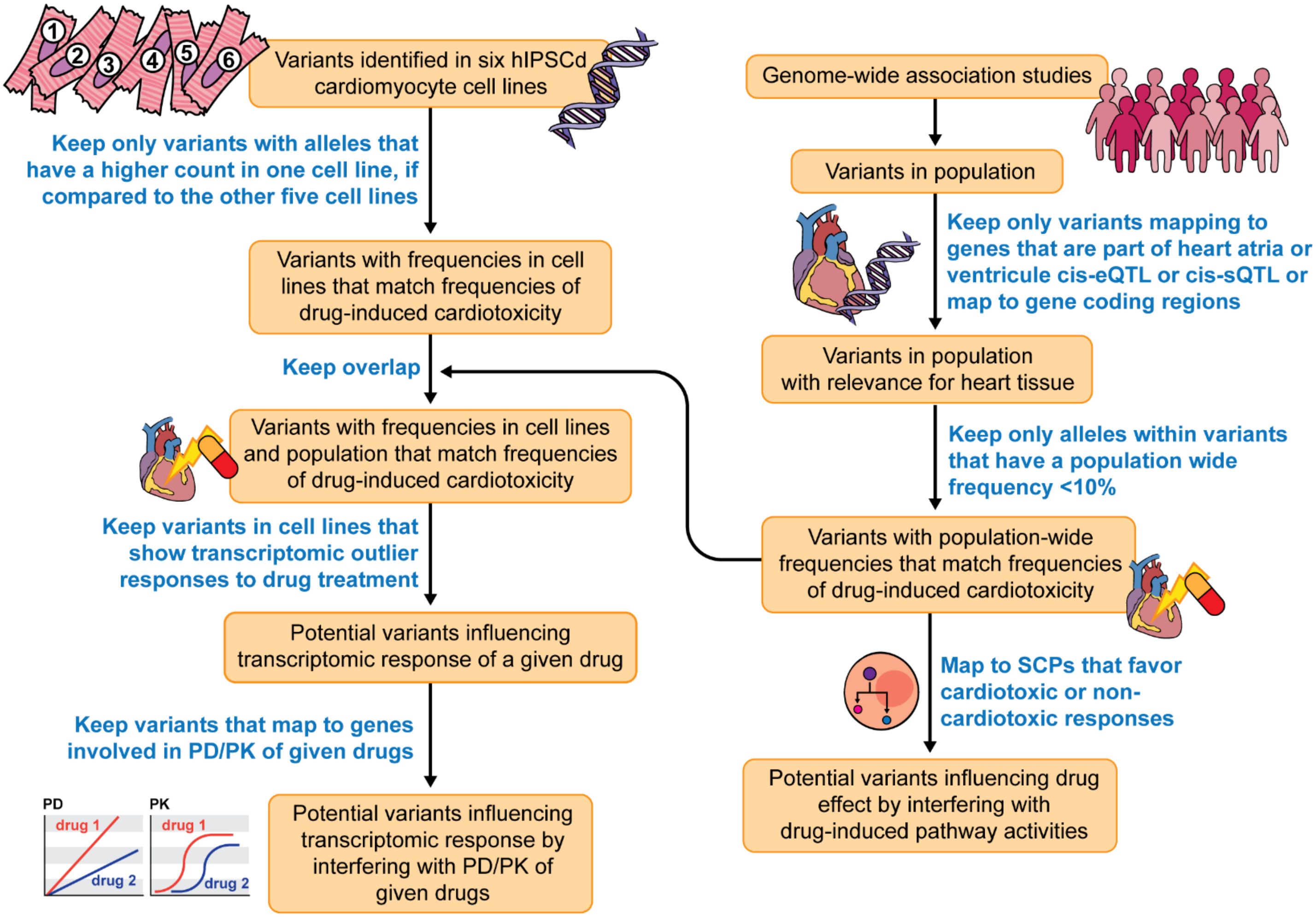


**Supplemental figure 25**

**Supplemental Figure 25. Computational pipeline for the identification of potential genomic variants associated with anthracycline- and TKI-induced cardiotoxicity.** The flow chart shows the steps involved in our pipeline for the identification of genomic variant candidates for drug-induced cardiotoxicity. See methods for details.

A

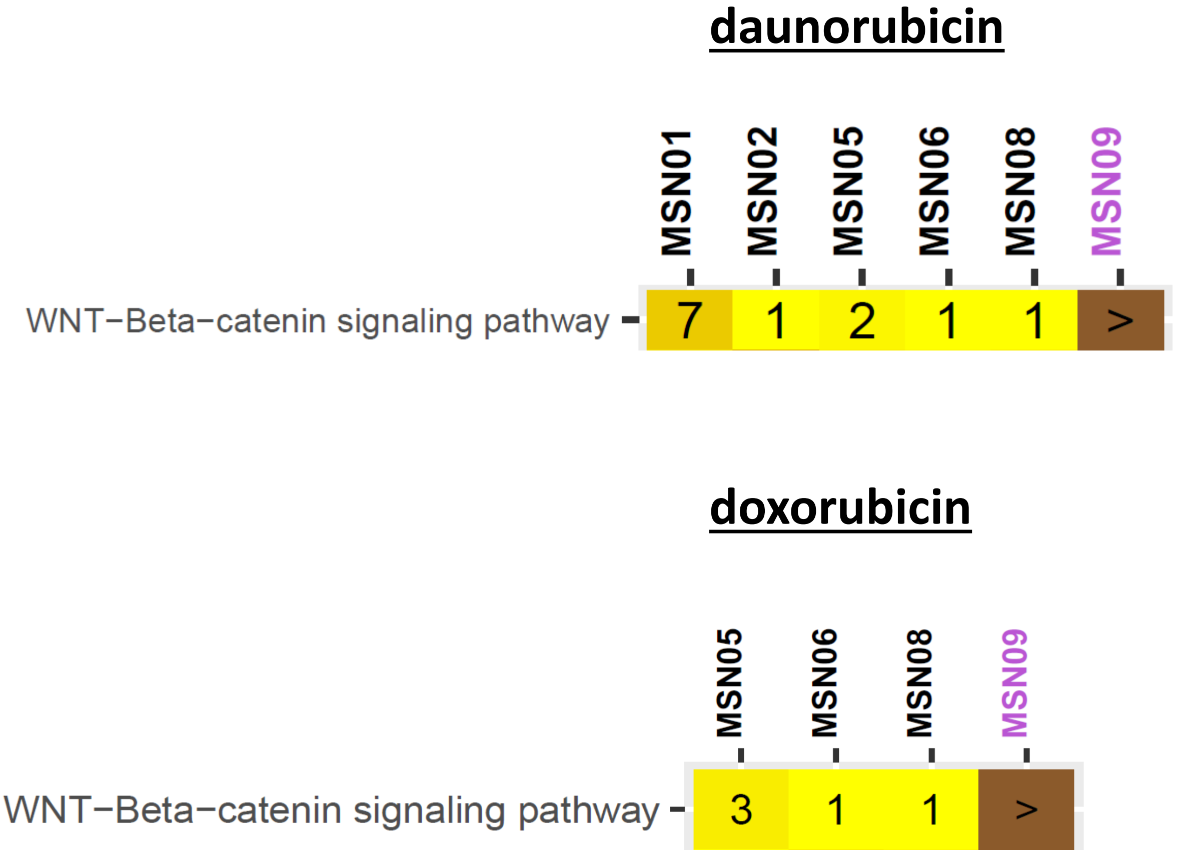


Supplemental figure 26

B

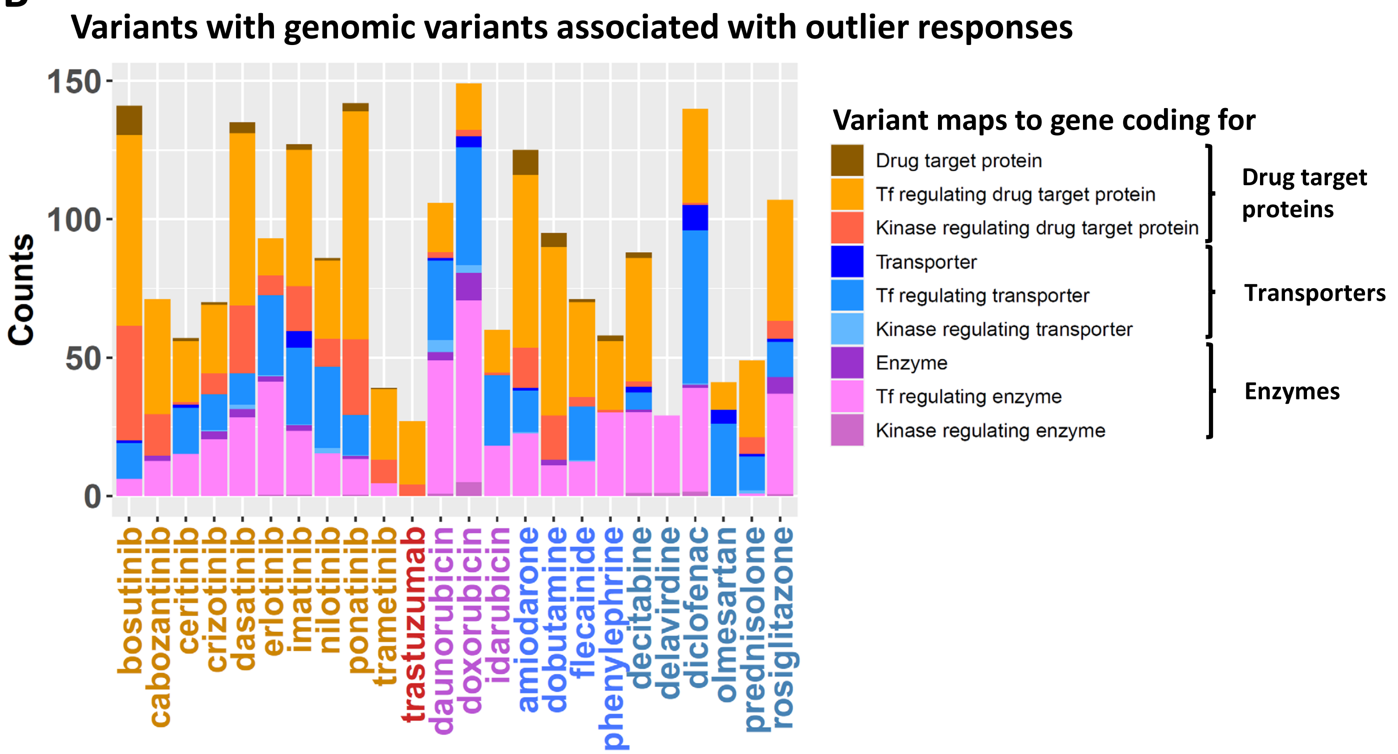


C

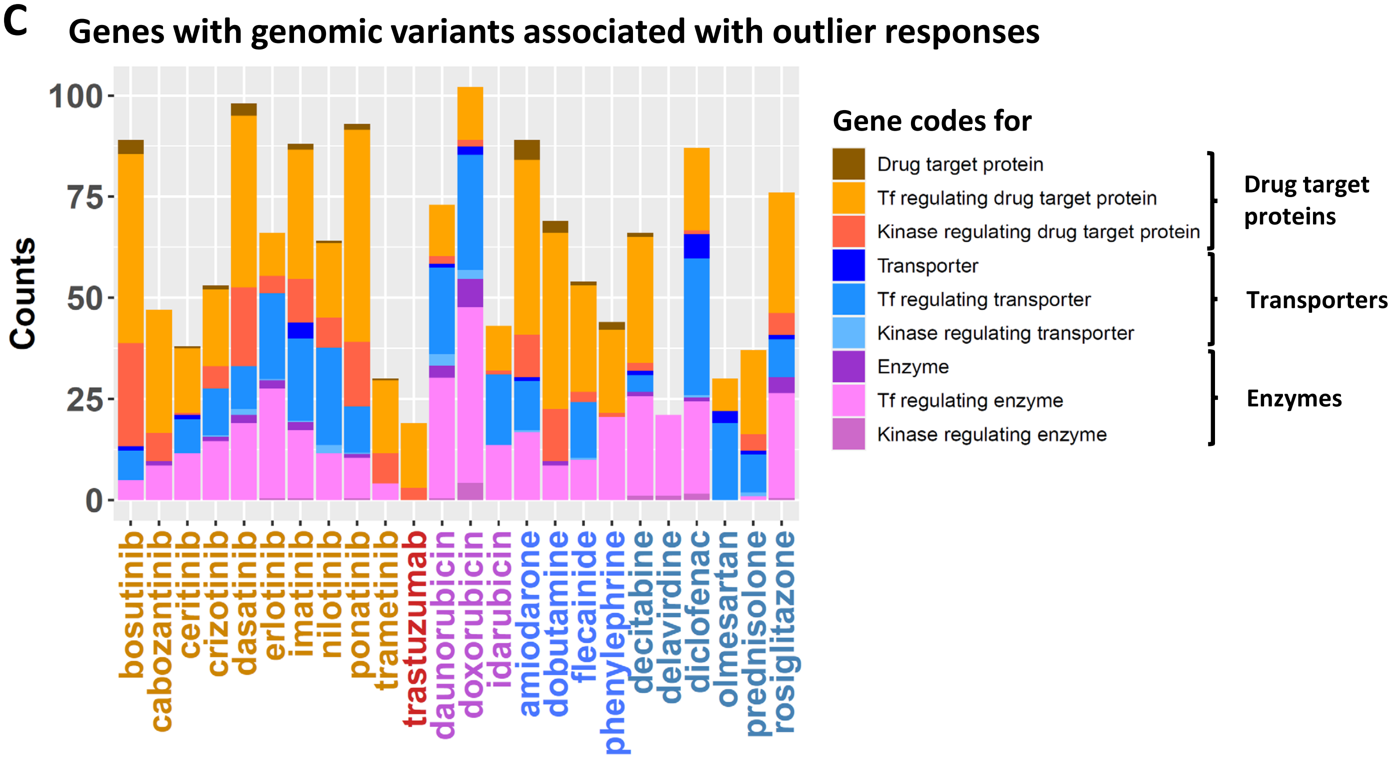


**Supplemental figure 26**

D

### Non-cardiotoxic TKIs

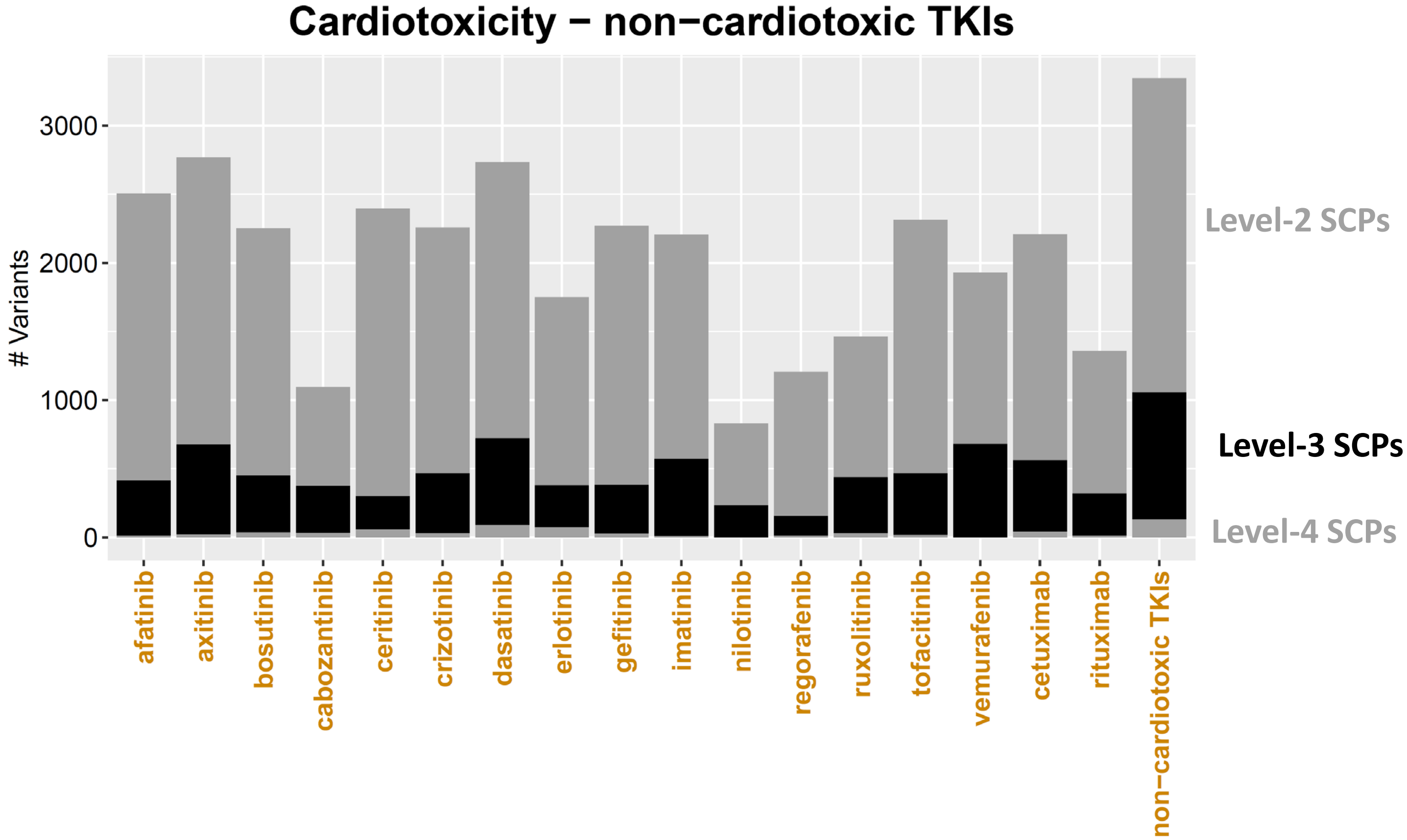


Supplemental figure 26

**Supplemental Figure 26. Identification of potential genomic variants associated with anthracycline- and TKI-induced cardiotoxicity. (A)** Results for the enrichment of upregulated genes induced by daunorubicin and doxorubicin for the 'Wnt-Beta-catenin signaling pathway' were extracted from Suppl. Figure 14D. The cell line MSN09 (purple) was identified for both drugs as an outlier cell line/drug combination and contains the rs2229774 mutation in the RARG gene that is linked to increased anthracycline-induced cardiotoxicity. **(B)** Genomic variants that our algorithm identified as potential regulators of genes involved in a drug's pharmacodynamics (PD) or -kinetics (PK) are shown for all drugs. The results for the cardiotoxic TKIs and anthracyclines are also shown in main figure 4C. **(C)** Since multiple variants map to the same genes, we also counted the number of genes with at least one variant. **(D)** Variants with a low frequency in the whole population that are part of e/sQTLs or lie within coding regions were mapped to up- or downregulated level-2, -3 and -4 SCPs that we predicted to be up- or downregulated at higher ranks by non-cardiotoxic TKIs. Variants that are part of identified SCPs of multiple levels are only counted for the lowest level SCPs (higher level number) to prevent double counting. See main Figure 4F for results obtained for cardiotoxic TKIs.

**Supplemental Tables.**

All tables are provided as tab delimited text files separately.

**Supplemental Table 1.** Characteristics of cell lines used in this study, as published previously [16].

**Supplemental Table 2.** Source and treatment concentrations for each drug used in this study.

**Supplemental Table 3.** Description of materials used in this study. This table is taken from the Supplemental Materials for Schaniel et al. [16] since the two studies were conducted concurrently.

**Supplemental Table 4.** Cardiotoxic and non-cardiotoxic small molecule tyrosine kinase inhibitors and monoclonal antibodies.

**Supplemental Table 5. Drug-induced differentially expressed genes in each cell line.** DEGs were ranked by significance. The spreadsheet contains the top 600 most significant DEGs induced by each drug in each treated cell line.

**Supplemental Table 6. Averaged drug-induced differentially expressed genes.**

**Supplemental Table 7. Top 600 genes of the first eigenarray.**

**Supplemental Table 8. Enrichment analysis of the first eigenarray.** The top 600 genes with the highest absolute expression values of the first eigenarray were subjected to pathway enrichment analysis using the Molecular Biology of the Cell Ontology MBCO level-2 SCPs.

**Supplemental Table 9. Drug-induced differentially expressed genes in each cell line after removal of the first eigenarray.** Significance p-values were transformed into $-\log_{10}$(p-values). We defined $-\log_{10}$(p-values) of up- and down-regulated genes as positive and negative, respectively. Signed $-\log_{10}$(p-values) were integrated into a gene expression matrix that was subjected to singular value decomposition (SVD). After removal of the first eigenarray the matrix contained decomposed $-\log_{10}$(p-values). Since our SVD pipeline acts on the $-\log_{10}$(p-values) and not on the $\log_2$(fold changes), we removed the $\log_2$(fold changes) from the gene expression profiles after any decomposition step.

**Supplemental Table 10. Drug-selective gene expression profiles after projection into drug-selective subspaces**. Drug-selective subspaces were identified using SVD. Projection of the complete gene expression profiles into drug-selective subspaces gave drug-selective gene expression profiles with decomposed $-\log_{10}$(p-values). As explained for Suppl. Table 9, these profiles do not contain $\log_2$(fold changes).

**Supplemental Table 11. Averaged drug-selective gene expression profiles.**

**Supplemental Table 12. Enrichment analysis results of drug-selective gene expression profiles.** After projection of gene expression profiles into drug-selective subspaces, we ranked the genes within each cell line/drug combination (i.e., unique cell line and drug combination) by decreasing absolute decomposed $-\log_{10}$(p-values). Up- and downregulated genes among the top 600 ranked genes were subjected to pathway enrichment analysis using Molecular Biology of the Cell Ontology **(A)** level-1, **(B)** -2, **(C)** -3 and **(D)** -4 SCPs.

**Supplemental Table 13. Enrichment analysis results of the complete gene expression profiles.** Up- and downregulated genes among the top 600 most significantly differentially expressed genes of the complete dataset were subjected to pathway enrichment analysis using Molecular Biology of the Cell Ontology **(A)** level-1, **(B)** -2, **(C)** -3 and **(D)** -4 SCPs.

**Supplemental Table 14. Enrichment analysis results of gene expression profiles after removal of the first eigenarray.** Up- and downregulated genes among the top 600 most significantly differentially expressed genes after removal of the first eigenarray were subjected to pathway enrichment analysis using Molecular Biology of the Cell Ontology **(A)** level-1, **(B)** -2, **(C)** -3 and **(D)** -4 SCPs.

**Supplemental Table 15. Cluster marker genes identified by single cell RNAseq of hiPSC-derived cardiomyocytes.** Single cell RNAseq analysis of four of our six different hiPSC-derived cardiomyocyte cell lines identified six different clusters. Cluster marker genes were calculated by comparing gene expression profiles in each cluster versus all other clusters.

**Supplemental Table 16. Enrichment analysis of cluster marker genes obtained by single cell RNAseq of hiPSC-derived cardiomycytes.** The top 500 marker genes of each cluster with a maximum adjusted p-value of 0.05 were subjected to enrichment analysis using **(A)** cell type marker genes of the adult human heart, **(B)** cell type marker genes of the developing human heart, MBCO **(C)** level-1, **(D)** -2, **(E)** -3 and **(F)** -4 SCPs.

**Supplemental Table 17. Cell type marker genes of the adult human heart.** Cell type marker genes were calculated for each cell type in the adult human heart.

**Supplemental Table 18. Enrichment analysis of cell type marker genes of the adult human heart.** The top 500 marker genes of each cell type in the adult human heart with a maximum adjusted p-value of 0.05 were subjected to enrichment analysis using MBCO **(A)** level-1, **(B)** -2, **(C)** -3 and **(D)** -4 SCPs.

**Supplemental Table 19. Differentially expressed genes in DCM vs hiPSC-derived cardiomyocytes from a healthy subject.** Pseudo-bulk analysis to identify differentially expressed genes in hiPSC-derived cardiomyocytes from an infant patient with DCM and a healthy subject.

**Supplemental Table 20. MBCO enrichment results of cell types obtained from DCM and HCM patients.** Up- and downregulated genes in **(A)** hiPSC-derived cardiomyocytes of an infant DCM patient, **(B)** in adult heart cells of DCM patients, and **(C)** in adult heart cells of DCM or HCM patients obtained from a different study, were subjected to enrichment analysis using MBCO.

**Supplemental Table 21: Genomic variants related to pharmacodynamics or pharmacokinetics.**

**Supplemental Table 22: Genomic variants mapping to SCPs up- or downregulated at higher ranks by cardiotoxic or non-cardiotoxic TKIs.**
