## Supplemental Figure 14B for "Multiscale mapping of transcriptomic signatures for cardiotoxic drugs"

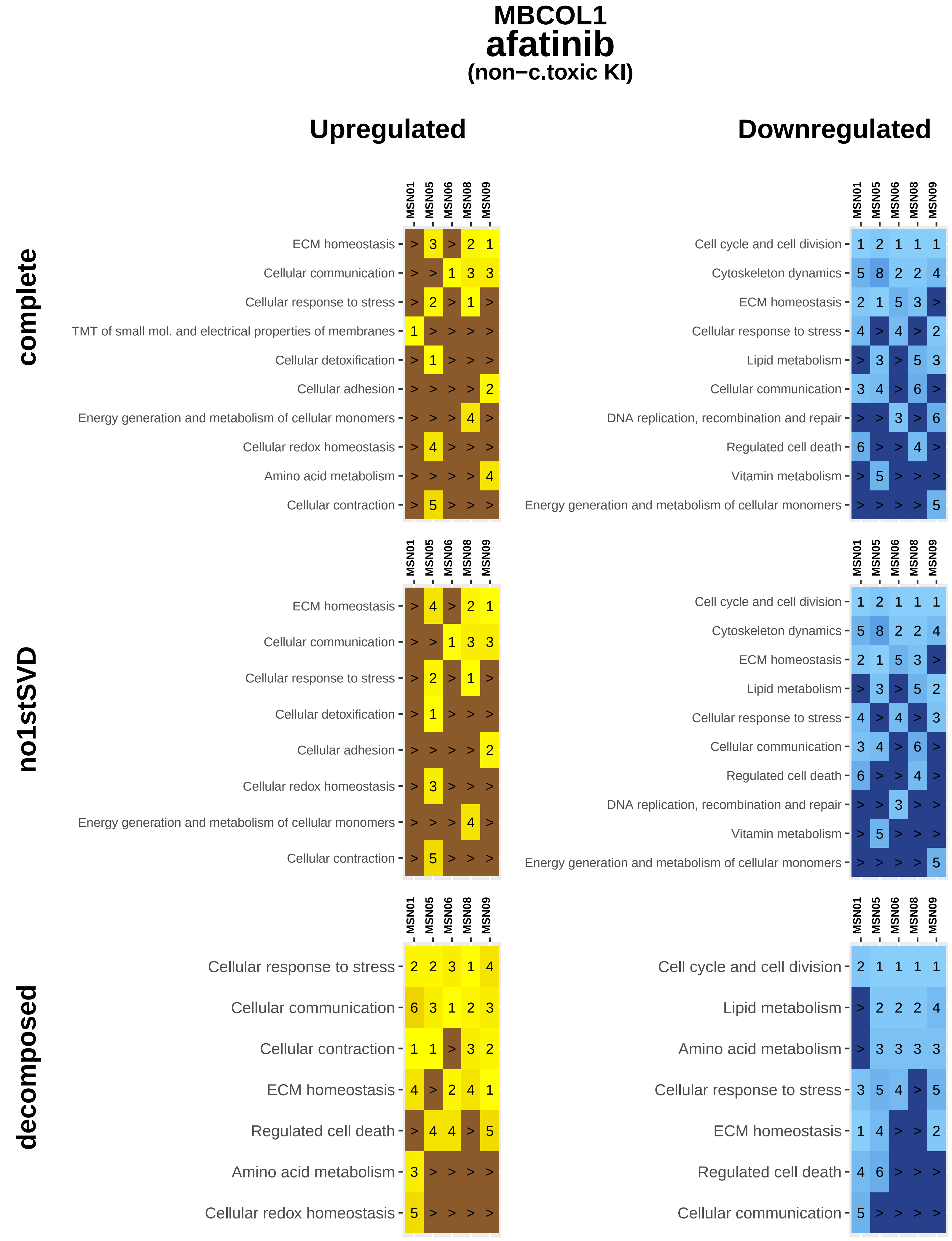
MBCOL1
afatinib
(non−c.toxic KI)
Upregulated
Downregulated
complete
no1stSVD
decomposed
MSN01 MSN05 MSN06 MSN08 MSN09
ECM homeostasis > 3 > 2 1
Cellular communication > > 1 3 3
Cellular response to stress > 2 > 1 >
TMT of small mol. and electrical properties of membranes 1 > > > >
Cellular detoxification > 1 > > >
Cellular adhesion > > > > 2
Energy generation and metabolism of cellular monomers > > > 4 >
Cellular redox homeostasis > 4 > > >
Amino acid metabolism > > > > 4
Cellular contraction > 5 > > >
Cell cycle and cell division 1 2 1 1 1
Cytoskeleton dynamics 5 8 2 2 4
ECM homeostasis 2 1 5 3 >
Cellular response to stress 4 > 4 > 2
Lipid metabolism > 3 > 5 3
Cellular communication 3 4 > 6 >
DNA replication, recombination and repair > > 3 > 6
Regulated cell death 6 > > 4 >
Vitamin metabolism > 5 > > >
Energy generation and metabolism of cellular monomers > > > > 5
ECM homeostasis > 4 > 2 1
Cellular communication > > 1 3 3
Cellular response to stress > 2 > 1 >
Cellular detoxification > 1 > > >
Cellular adhesion > > > > 2
Cellular redox homeostasis > 3 > > >
Energy generation and metabolism of cellular monomers > > > 4 >
Cellular contraction > 5 > > >
Cell cycle and cell division 1 2 1 1 1
Cytoskeleton dynamics 5 8 2 2 4
ECM homeostasis 2 1 5 3 >
Lipid metabolism > 3 > 5 2
Cellular response to stress 4 > 4 > 3
Cellular communication 3 4 > 6 >
Regulated cell death 6 > > 4 >
DNA replication, recombination and repair > > 3 > >
Vitamin metabolism > 5 > > >
Energy generation and metabolism of cellular monomers > > > > 5
Cellular response to stress 2 2 3 1 4
Cellular communication 6 3 1 2 3
Cellular contraction 1 1 > 3 2
ECM homeostasis 4 > 2 4 1
Regulated cell death > 4 4 > 5
Amino acid metabolism 3 > > > >
Cellular redox homeostasis 5 > > > >
Cell cycle and cell division 2 1 1 1 1
Lipid metabolism > 2 2 2 4
Amino acid metabolism > 3 3 3 3
Cellular response to stress 3 5 4 > 5
ECM homeostasis 1 4 > > 2
Regulated cell death 4 6 > > >
Cellular communication 5 > > > >

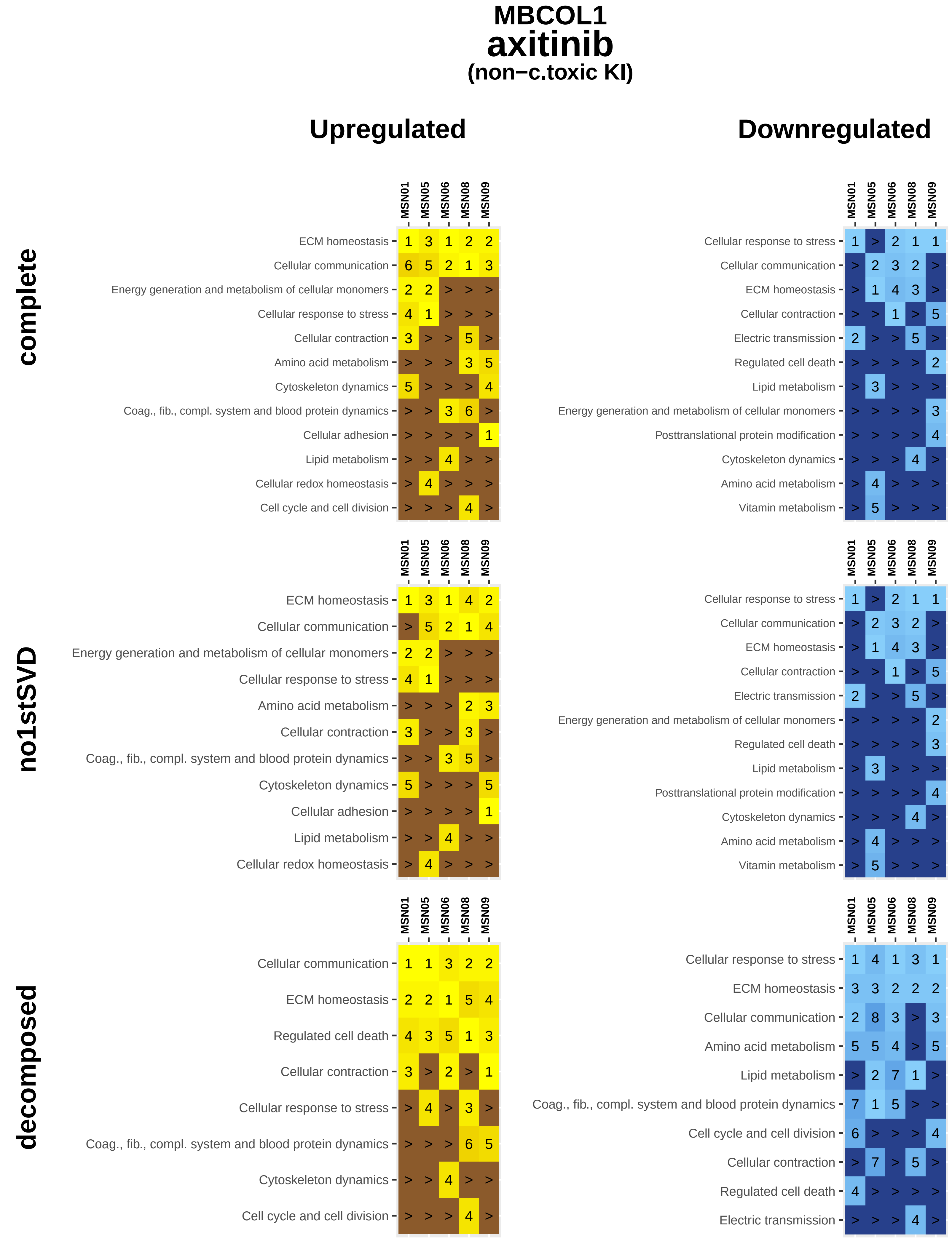
MBCOL1
axitinib
(non−c.toxic KI)
Upregulated
Downregulated
complete
MSN01 MSN05 MSN06 MSN08 MSN09
ECM homeostasis 1 3 1 2 2
Cellular communication 6 5 2 1 3
Energy generation and metabolism of cellular monomers 2 2 > > >
Cellular response to stress 4 1 > > >
Cellular contraction 3 > > 5 >
Amino acid metabolism > > > 3 5
Cytoskeleton dynamics 5 > > > 4
Coag., fib., compl. system and blood protein dynamics > > 3 6 >
Cellular adhesion > > > > 1
Lipid metabolism > > 4 > >
Cellular redox homeostasis > 4 > > >
Cell cycle and cell division > > > 4 >
MSN01 MSN05 MSN06 MSN08 MSN09
Cellular response to stress 1 > 2 1 1
Cellular communication > 2 3 2 >
ECM homeostasis > 1 4 3 >
Cellular contraction > > 1 > 5
Electric transmission 2 > > 5 >
Regulated cell death > > > > 2
Lipid metabolism > 3 > > >
Energy generation and metabolism of cellular monomers > > > > 3
Posttranslational protein modification > > > > 4
Cytoskeleton dynamics > > > 4 >
Amino acid metabolism > 4 > > >
Vitamin metabolism > 5 > > >
no1stSVD
MSN01 MSN05 MSN06 MSN08 MSN09
ECM homeostasis 1 3 1 4 2
Cellular communication > 5 2 1 4
Energy generation and metabolism of cellular monomers 2 2 > > >
Cellular response to stress 4 1 > > >
Amino acid metabolism > > > 2 3
Cellular contraction 3 > > 3 >
Coag., fib., compl. system and blood protein dynamics > > 3 5 >
Cytoskeleton dynamics 5 > > > 5
Cellular adhesion > > > > 1
Lipid metabolism > > 4 > >
Cellular redox homeostasis > 4 > > >
MSN01 MSN05 MSN06 MSN08 MSN09
Cellular response to stress 1 > 2 1 1
Cellular communication > 2 3 2 >
ECM homeostasis > 1 4 3 >
Cellular contraction > > 1 > 5
Electric transmission 2 > > 5 >
Energy generation and metabolism of cellular monomers > > > > 2
Regulated cell death > > > > 3
Lipid metabolism > 3 > > >
Posttranslational protein modification > > > > 4
Cytoskeleton dynamics > > > 4 >
Amino acid metabolism > 4 > > >
Vitamin metabolism > 5 > > >
decomposed
MSN01 MSN05 MSN06 MSN08 MSN09
Cellular communication 1 1 3 2 2
ECM homeostasis 2 2 1 5 4
Regulated cell death 4 3 5 1 3
Cellular contraction 3 > 2 > 1
Cellular response to stress > 4 > 3 >
Coag., fib., compl. system and blood protein dynamics > > > 6 5
Cytoskeleton dynamics > > 4 > >
Cell cycle and cell division > > > 4 >
MSN01 MSN05 MSN06 MSN08 MSN09
Cellular response to stress 1 4 1 3 1
ECM homeostasis 3 3 2 2 2
Cellular communication 2 8 3 > 3
Amino acid metabolism 5 5 4 > 5
Lipid metabolism > 2 7 1 >
Coag., fib., compl. system and blood protein dynamics 7 1 5 > >
Cell cycle and cell division 6 > > > 4
Cellular contraction > 7 > 5 >
Regulated cell death 4 > > > >
Electric transmission > > > 4 >

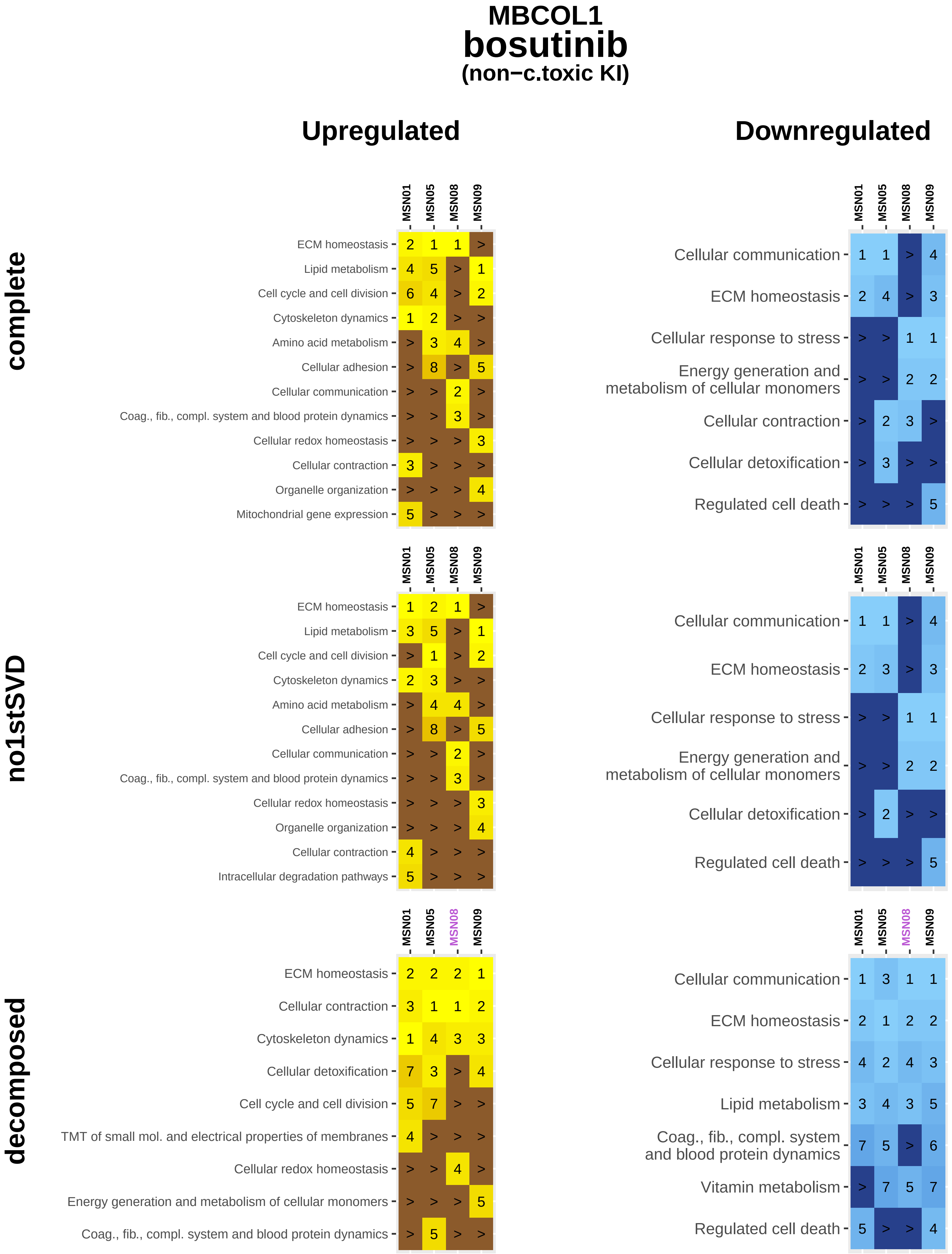
MBCOL1
bosutinib
(non−c.toxic KI)
Upregulated
Downregulated
complete
no1stSVD
decomposed
MSN01 MSN05 MSN08 MSN09
ECM homeostasis 2 1 1 >
Lipid metabolism 4 5 > 1
Cell cycle and cell division 6 4 > 2
Cytoskeleton dynamics 1 2 > >
Amino acid metabolism > 3 4 >
Cellular adhesion > 8 > 5
Cellular communication > > 2 >
Coag., fib., compl. system and blood protein dynamics > > 3 >
Cellular redox homeostasis > > > 3
Cellular contraction 3 > > >
Organelle organization > > > 4
Mitochondrial gene expression 5 > > >
Cellular communication 1 1 > 4
ECM homeostasis 2 4 > 3
Cellular response to stress > > 1 1
Energy generation and metabolism of cellular monomers > > 2 2
Cellular contraction > 2 3 >
Cellular detoxification > 3 > >
Regulated cell death > > > 5
ECM homeostasis 1 2 1 >
Lipid metabolism 3 5 > 1
Cell cycle and cell division > 1 > 2
Cytoskeleton dynamics 2 3 > >
Amino acid metabolism > 4 4 >
Cellular adhesion > 8 > 5
Cellular communication > > 2 >
Coag., fib., compl. system and blood protein dynamics > > 3 >
Cellular redox homeostasis > > > 3
Organelle organization > > > 4
Cellular contraction 4 > > >
Intracellular degradation pathways 5 > > >
Cellular communication 1 1 > 4
ECM homeostasis 2 3 > 3
Cellular response to stress > > 1 1
Energy generation and metabolism of cellular monomers > > 2 2
Cellular detoxification > 2 > >
Regulated cell death > > > 5
ECM homeostasis 2 2 2 1
Cellular contraction 3 1 1 2
Cytoskeleton dynamics 1 4 3 3
Cellular detoxification 7 3 > 4
Cell cycle and cell division 5 7 > >
TMT of small mol. and electrical properties of membranes 4 > > >
Cellular redox homeostasis > > 4 >
Energy generation and metabolism of cellular monomers > > > 5
Coag., fib., compl. system and blood protein dynamics > 5 > >
Cellular communication 1 3 1 1
ECM homeostasis 2 1 2 2
Cellular response to stress 4 2 4 3
Lipid metabolism 3 4 3 5
Coag., fib., compl. system and blood protein dynamics 7 5 > 6
Vitamin metabolism > 7 5 7
Regulated cell death 5 > > 4

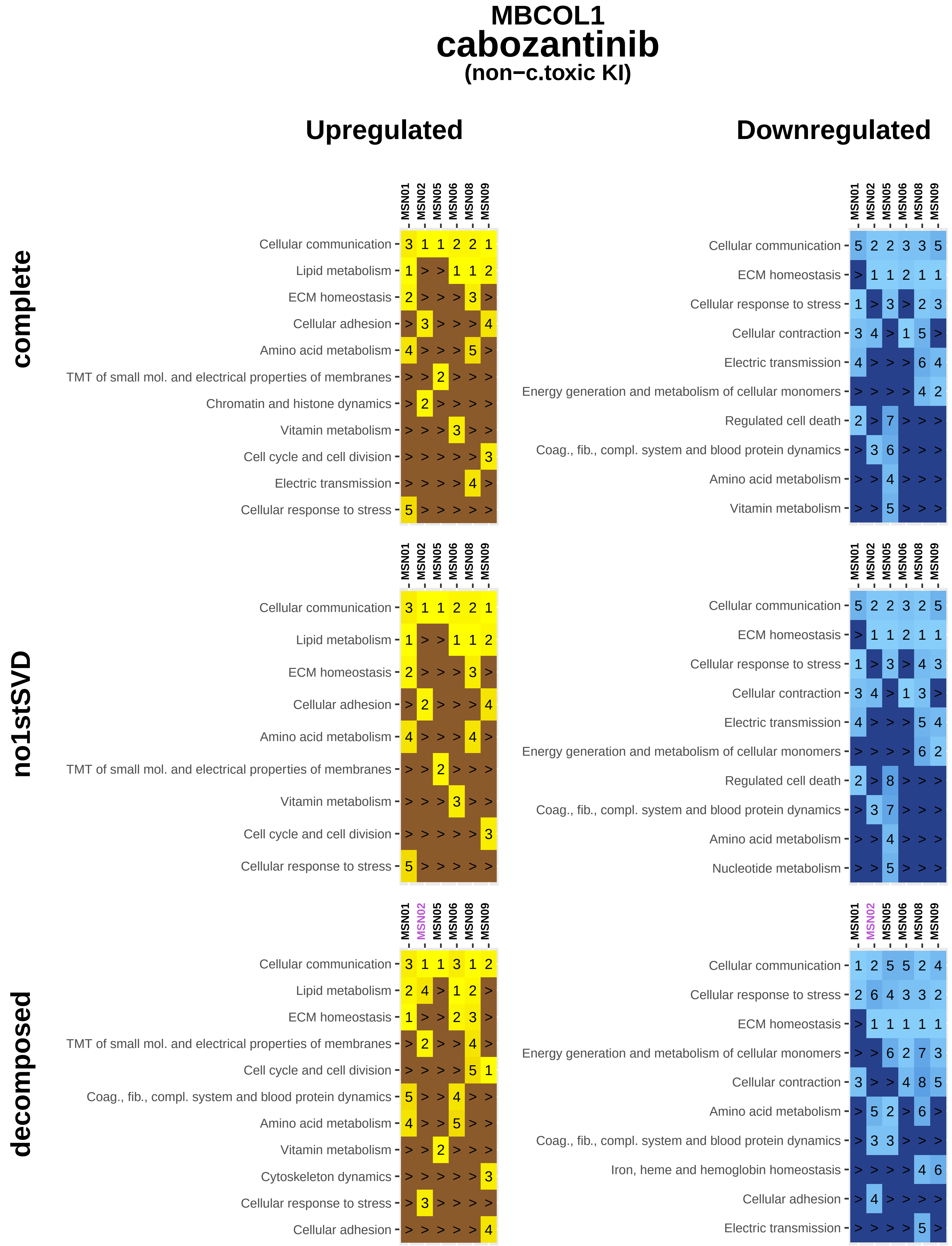

MBCOL1
cabozantinib
(non−c.toxic KI)
Upregulated
Downregulated
complete
no1stSVD
decomposed
MSN01
MSN02
MSN05
MSN06
MSN08
MSN09
Cellular communication
Lipid metabolism
ECM homeostasis
Cellular adhesion
Amino acid metabolism
TMT of small mol. and electrical properties of membranes
Chromatin and histone dynamics
Vitamin metabolism
Cell cycle and cell division
Electric transmission
Cellular response to stress
Cellular contraction
Energy generation and metabolism of cellular monomers
Regulated cell death
Coag., fib., compl. system and blood protein dynamics
Nucleotide metabolism
Cytoskeleton dynamics
Iron, heme and hemoglobin homeostasis

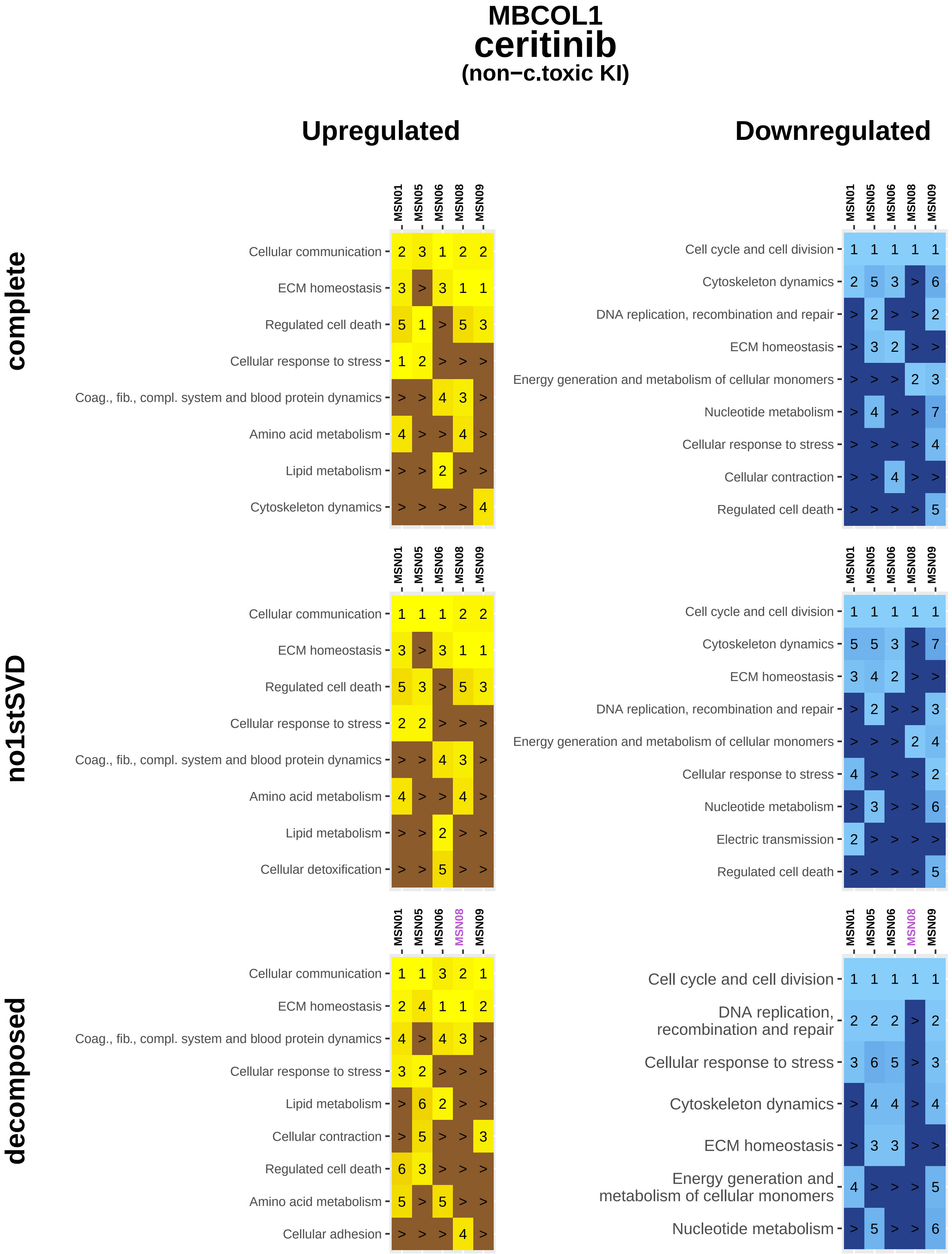
MBCOL1
ceritinib
(non−c.toxic KI)
Upregulated
Downregulated
complete
no1stSVD
decomposed
MSN01
MSN05
MSN06
MSN08
MSN09
Cellular communication
ECM homeostasis
Regulated cell death
Cellular response to stress
Coag., fib., compl. system and blood protein dynamics
Amino acid metabolism
Lipid metabolism
Cytoskeleton dynamics
Cellular detoxification
Cellular contraction
Cellular adhesion
Cell cycle and cell division
DNA replication, recombination and repair
Energy generation and metabolism of cellular monomers
Nucleotide metabolism
Electric transmission

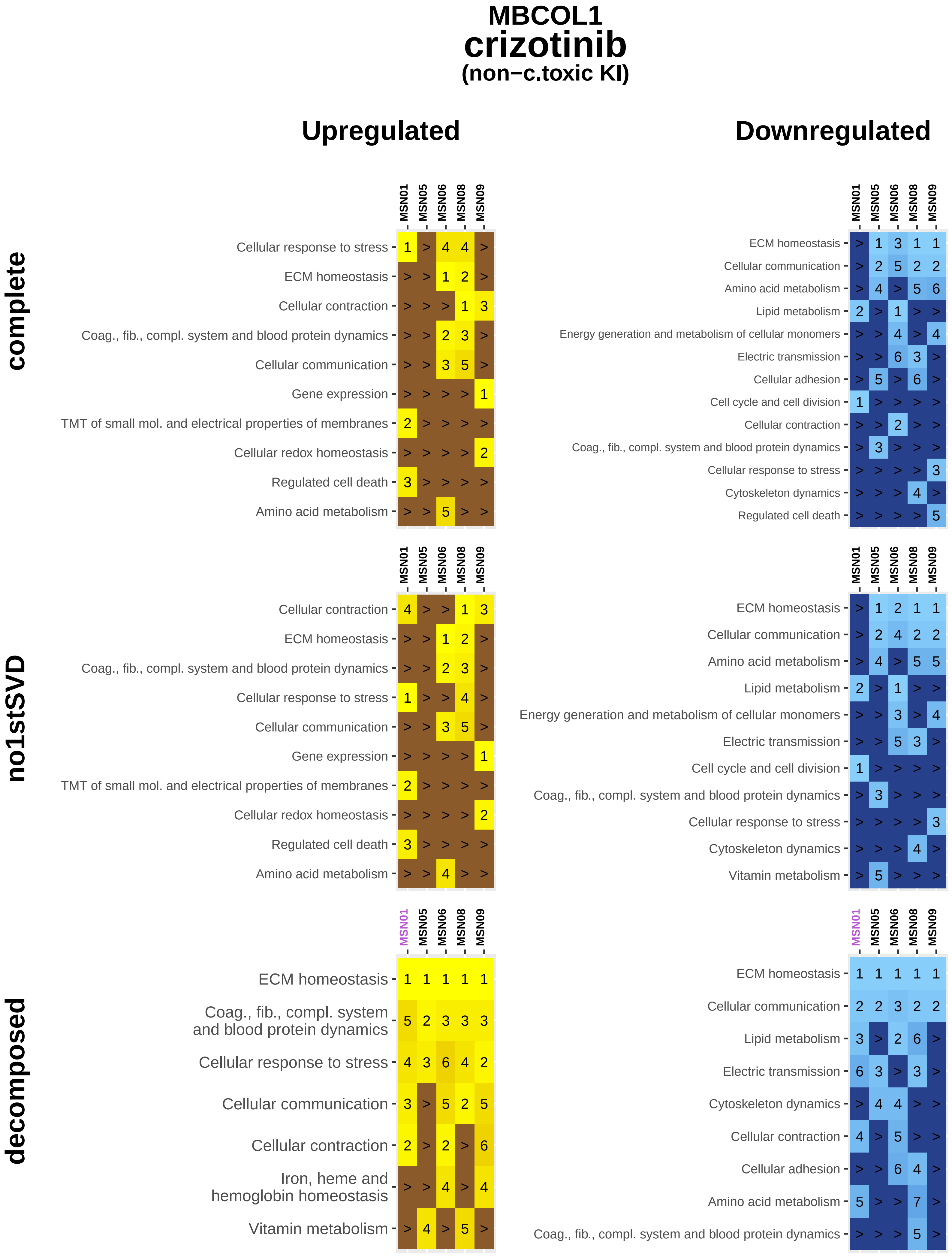
MBCOL1
crizotinib
(non−c.toxic KI)
Upregulated
Downregulated
complete
no1stSVD
decomposed
MSN01
MSN05
MSN06
MSN08
MSN09
Cellular response to stress
ECM homeostasis
Cellular contraction
Coag., fib., compl. system and blood protein dynamics
Cellular communication
Gene expression
TMT of small mol. and electrical properties of membranes
Cellular redox homeostasis
Regulated cell death
Amino acid metabolism
Lipid metabolism
Energy generation and metabolism of cellular monomers
Electric transmission
Cellular adhesion
Cell cycle and cell division
Cytoskeleton dynamics
Vitamin metabolism
Iron, heme and hemoglobin homeostasis

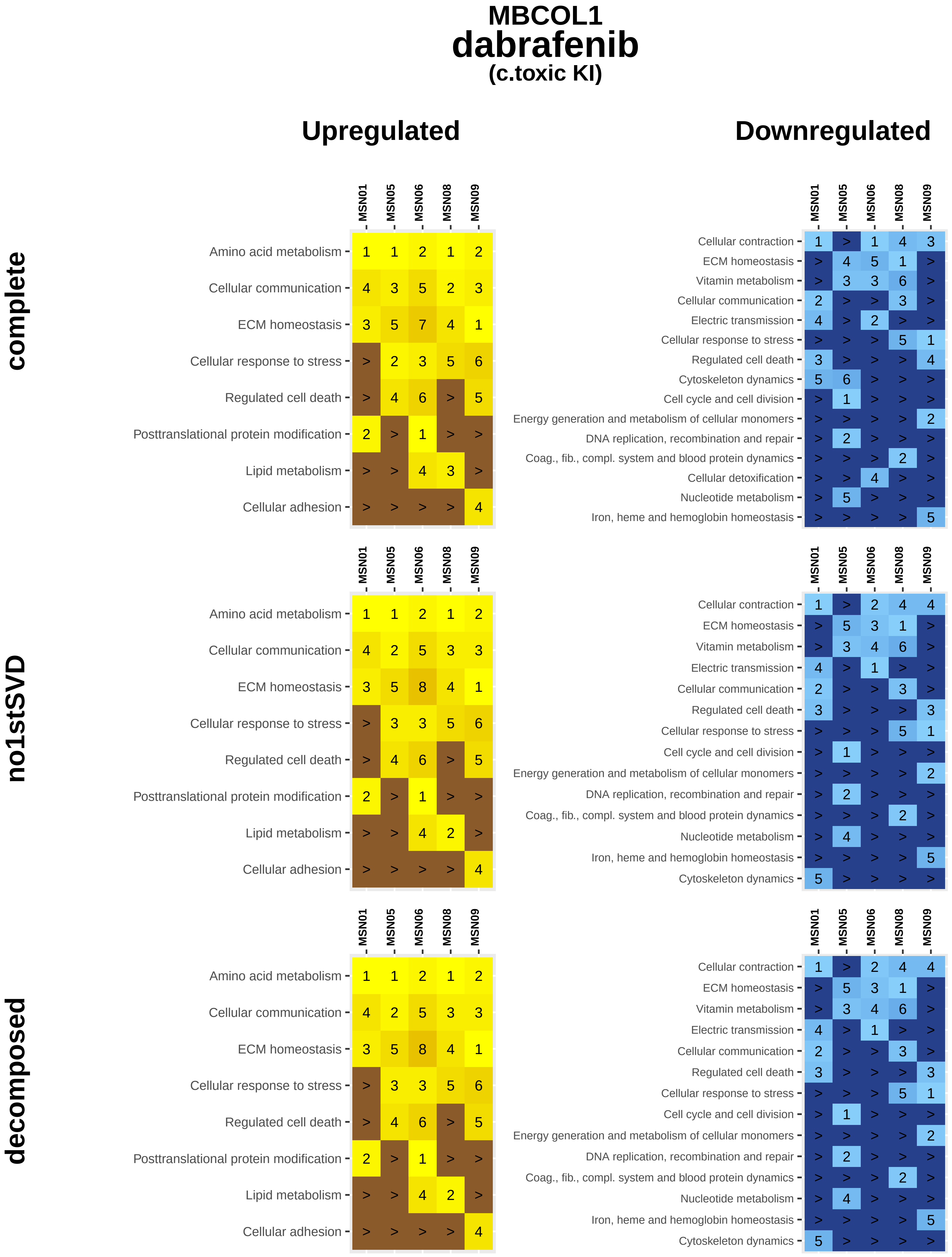
MBCOL1
dabrafenib
(c.toxic KI)
Upregulated
Downregulated
complete
no1stSVD
decomposed
MSN01 MSN05 MSN06 MSN08 MSN09
Amino acid metabolism 1 1 2 1 2
Cellular communication 4 3 5 2 3
ECM homeostasis 3 5 7 4 1
Cellular response to stress > 2 3 5 6
Regulated cell death > 4 6 > 5
Posttranslational protein modification 2 > 1 > >
Lipid metabolism > > 4 3 >
Cellular adhesion > > > > 4
Cellular contraction 1 > 1 4 3
ECM homeostasis > 4 5 1 >
Vitamin metabolism > 3 3 6 >
Cellular communication 2 > > 3 >
Electric transmission 4 > 2 > >
Cellular response to stress > > > 5 1
Regulated cell death 3 > > > 4
Cytoskeleton dynamics 5 6 > > >
Cell cycle and cell division > 1 > > >
Energy generation and metabolism of cellular monomers > > > > 2
DNA replication, recombination and repair > 2 > > >
Coag., fib., compl. system and blood protein dynamics > > > 2 >
Cellular detoxification > > 4 > >
Nucleotide metabolism > 5 > > >
Iron, heme and hemoglobin homeostasis > > > > 5
Amino acid metabolism 1 1 2 1 2
Cellular communication 4 2 5 3 3
ECM homeostasis 3 5 8 4 1
Cellular response to stress > 3 3 5 6
Regulated cell death > 4 6 > 5
Posttranslational protein modification 2 > 1 > >
Lipid metabolism > > 4 2 >
Cellular adhesion > > > > 4
Cellular contraction 1 > 2 4 4
ECM homeostasis > 5 3 1 >
Vitamin metabolism > 3 4 6 >
Electric transmission 4 > 1 > >
Cellular communication 2 > > 3 >
Regulated cell death 3 > > > 3
Cellular response to stress > > > 5 1
Cell cycle and cell division > 1 > > >
Energy generation and metabolism of cellular monomers > > > > 2
DNA replication, recombination and repair > 2 > > >
Coag., fib., compl. system and blood protein dynamics > > > 2 >
Nucleotide metabolism > 4 > > >
Iron, heme and hemoglobin homeostasis > > > > 5
Cytoskeleton dynamics 5 > > > >
Amino acid metabolism 1 1 2 1 2
Cellular communication 4 2 5 3 3
ECM homeostasis 3 5 8 4 1
Cellular response to stress > 3 3 5 6
Regulated cell death > 4 6 > 5
Posttranslational protein modification 2 > 1 > >
Lipid metabolism > > 4 2 >
Cellular adhesion > > > > 4
Cellular contraction 1 > 2 4 4
ECM homeostasis > 5 3 1 >
Vitamin metabolism > 3 4 6 >
Electric transmission 4 > 1 > >
Cellular communication 2 > > 3 >
Regulated cell death 3 > > > 3
Cellular response to stress > > > 5 1
Cell cycle and cell division > 1 > > >
Energy generation and metabolism of cellular monomers > > > > 2
DNA replication, recombination and repair > 2 > > >
Coag., fib., compl. system and blood protein dynamics > > > 2 >
Nucleotide metabolism > 4 > > >
Iron, heme and hemoglobin homeostasis > > > > 5
Cytoskeleton dynamics 5 > > > >

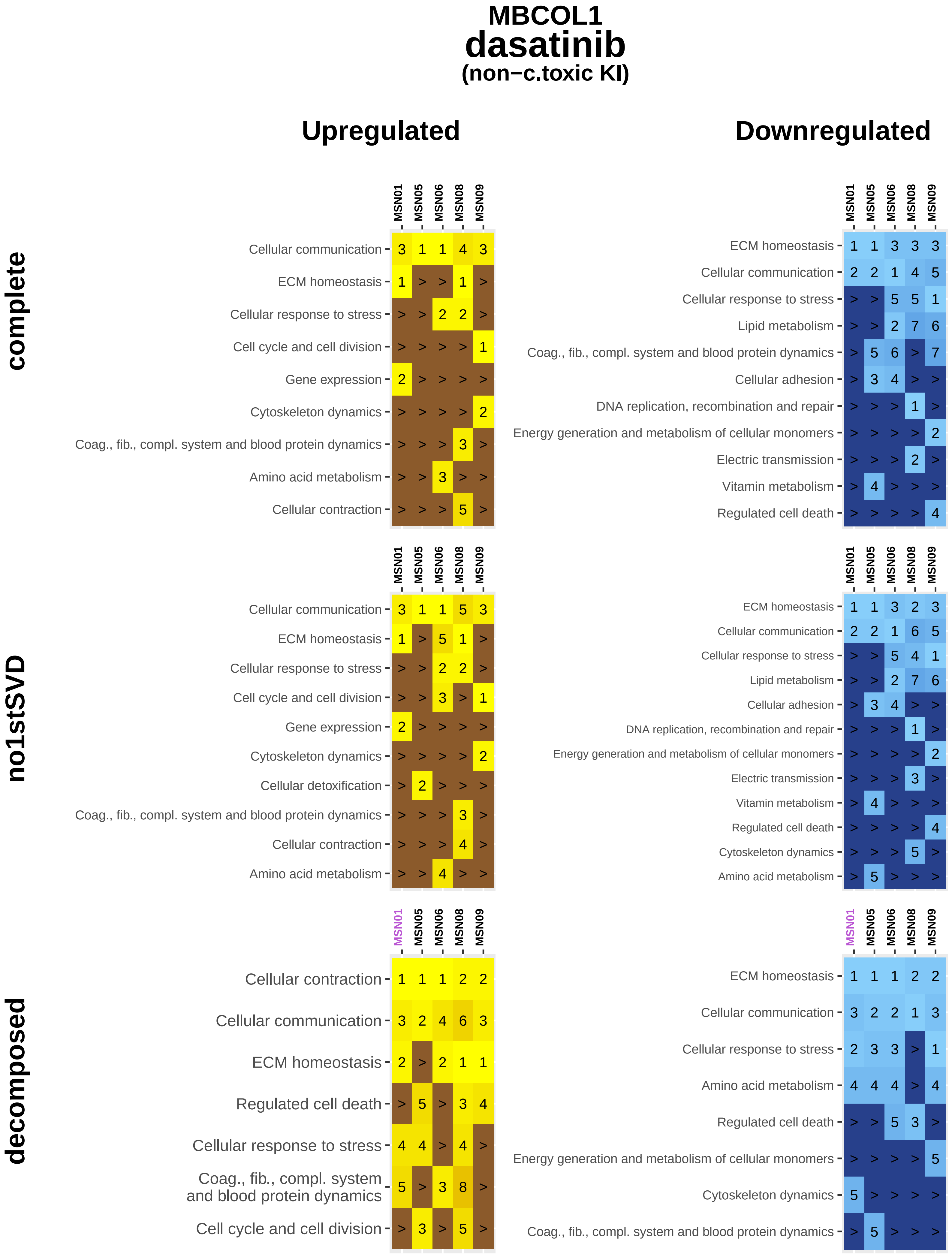

MBCOL1
dasatinib
(non−c.toxic KI)
Upregulated
Downregulated
complete
no1stSVD
decomposed
MSN01 MSN05 MSN06 MSN08 MSN09
Cellular communication 3 1 1 4 3
ECM homeostasis 1 > > 1 >
Cellular response to stress > > 2 2 >
Cell cycle and cell division > > > > 1
Gene expression 2 > > > >
Cytoskeleton dynamics > > > > 2
Coag., fib., compl. system and blood protein dynamics > > > 3 >
Amino acid metabolism > > 3 > >
Cellular contraction > > > 5 >
ECM homeostasis 1 1 3 3 3
Cellular communication 2 2 1 4 5
Cellular response to stress > > 5 5 1
Lipid metabolism > > 2 7 6
Coag., fib., compl. system and blood protein dynamics > 5 6 > 7
Cellular adhesion > 3 4 > >
DNA replication, recombination and repair > > > 1 >
Energy generation and metabolism of cellular monomers > > > > 2
Electric transmission > > > 2 >
Vitamin metabolism > 4 > > >
Regulated cell death > > > > 4
Cellular communication 3 1 1 5 3
ECM homeostasis 1 > 5 1 >
Cellular response to stress > > 2 2 >
Cell cycle and cell division > > 3 > 1
Gene expression 2 > > > >
Cytoskeleton dynamics > > > > 2
Cellular detoxification > 2 > > >
Coag., fib., compl. system and blood protein dynamics > > > 3 >
Cellular contraction > > > 4 >
Amino acid metabolism > > 4 > >
ECM homeostasis 1 1 3 2 3
Cellular communication 2 2 1 6 5
Cellular response to stress > > 5 4 1
Lipid metabolism > > 2 7 6
Cellular adhesion > 3 4 > >
DNA replication, recombination and repair > > > 1 >
Energy generation and metabolism of cellular monomers > > > > 2
Electric transmission > > > 3 >
Vitamin metabolism > 4 > > >
Regulated cell death > > > > 4
Cytoskeleton dynamics > > > 5 >
Amino acid metabolism > 5 > > >
Cellular contraction 1 1 1 2 2
Cellular communication 3 2 4 6 3
ECM homeostasis 2 > 2 1 1
Regulated cell death > 5 > 3 4
Cellular response to stress 4 4 > 4 >
Coag., fib., compl. system and blood protein dynamics 5 > 3 8 >
Cell cycle and cell division > 3 > 5 >
ECM homeostasis 1 1 1 2 2
Cellular communication 3 2 2 1 3
Cellular response to stress 2 3 3 > 1
Amino acid metabolism 4 4 4 > 4
Regulated cell death > > 5 3 >
Energy generation and metabolism of cellular monomers > > > > 5
Cytoskeleton dynamics 5 > > > >
Coag., fib., compl. system and blood protein dynamics > 5 > > >

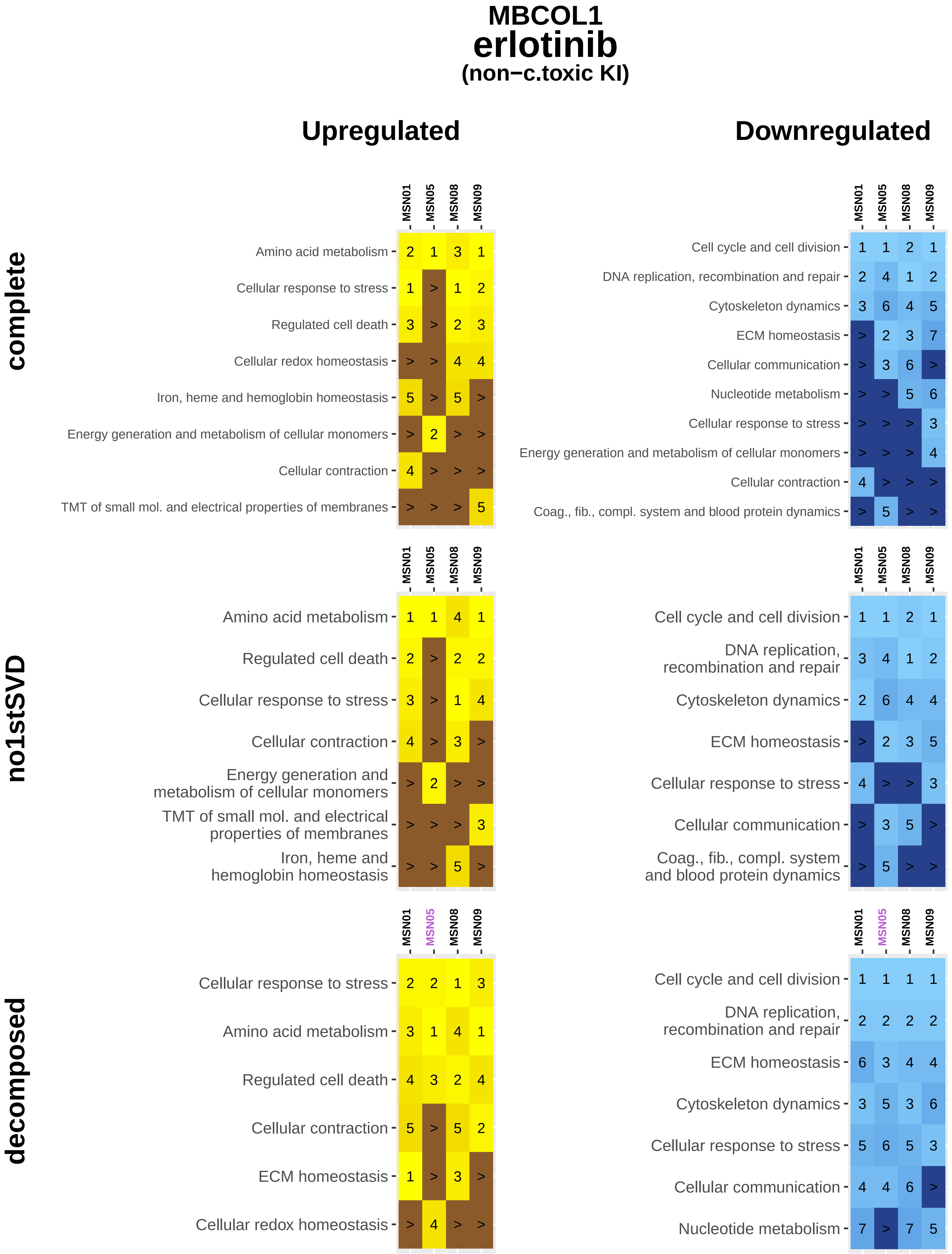

MBCOL1
erlotinib
(non−c.toxic KI)
Upregulated
Downregulated
complete
no1stSVD
decomposed
MSN01
MSN05
MSN08
MSN09
Amino acid metabolism
Cellular response to stress
Regulated cell death
Cellular redox homeostasis
Iron, heme and hemoglobin homeostasis
Energy generation and metabolism of cellular monomers
Cellular contraction
TMT of small mol. and electrical properties of membranes
Cell cycle and cell division
DNA replication, recombination and repair
Cytoskeleton dynamics
ECM homeostasis
Cellular communication
Nucleotide metabolism
Coag., fib., compl. system and blood protein dynamics

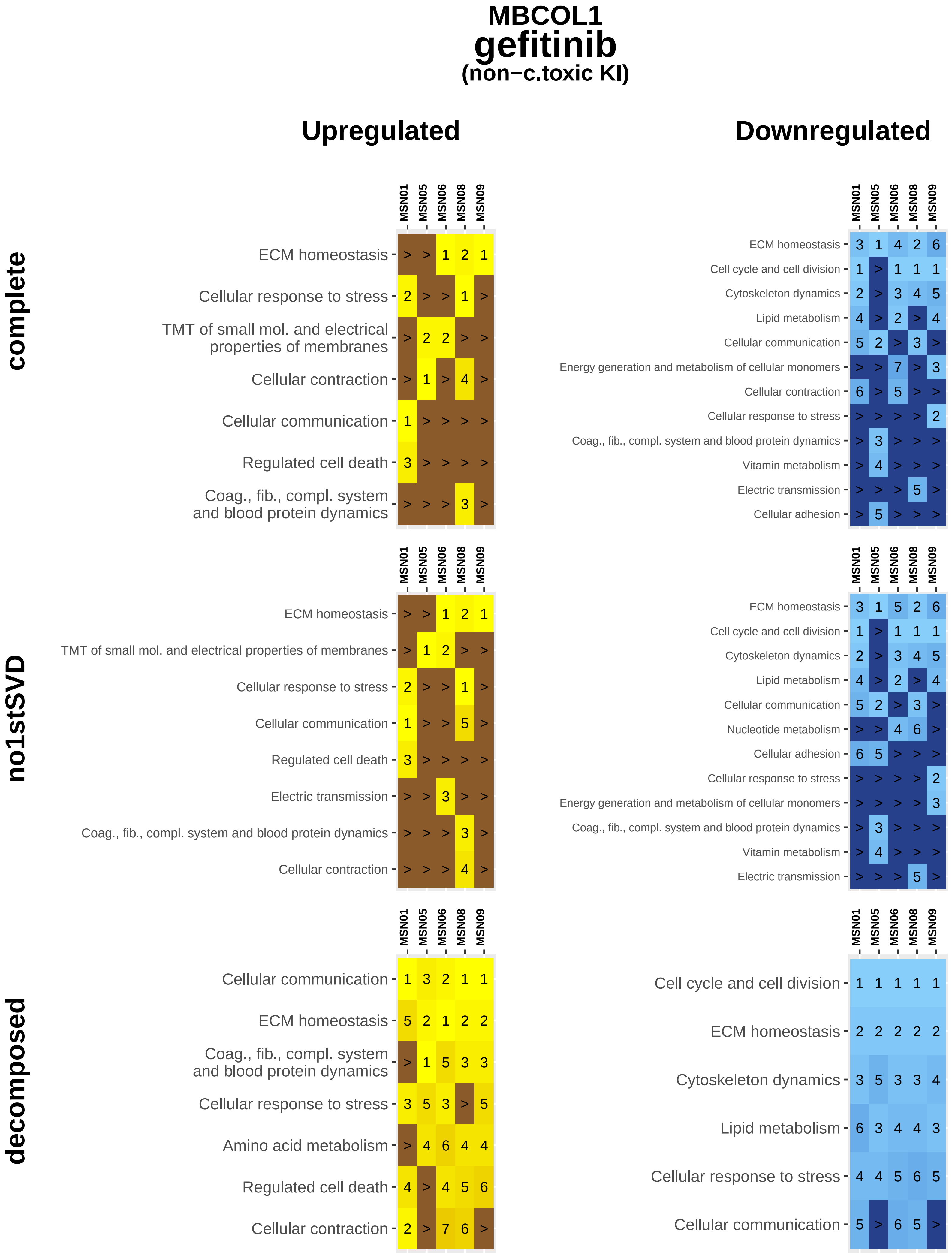
MBCOL1
gefitinib
(non−c.toxic KI)
Upregulated
Downregulated
complete
no1stSVD
decomposed
MSN01
MSN05
MSN06
MSN08
MSN09
ECM homeostasis
Cellular response to stress
TMT of small mol. and electrical properties of membranes
Cellular contraction
Cellular communication
Regulated cell death
Coag., fib., compl. system and blood protein dynamics
Cell cycle and cell division
Cytoskeleton dynamics
Lipid metabolism
Energy generation and metabolism of cellular monomers
Vitamin metabolism
Electric transmission
Cellular adhesion
Nucleotide metabolism
Amino acid metabolism

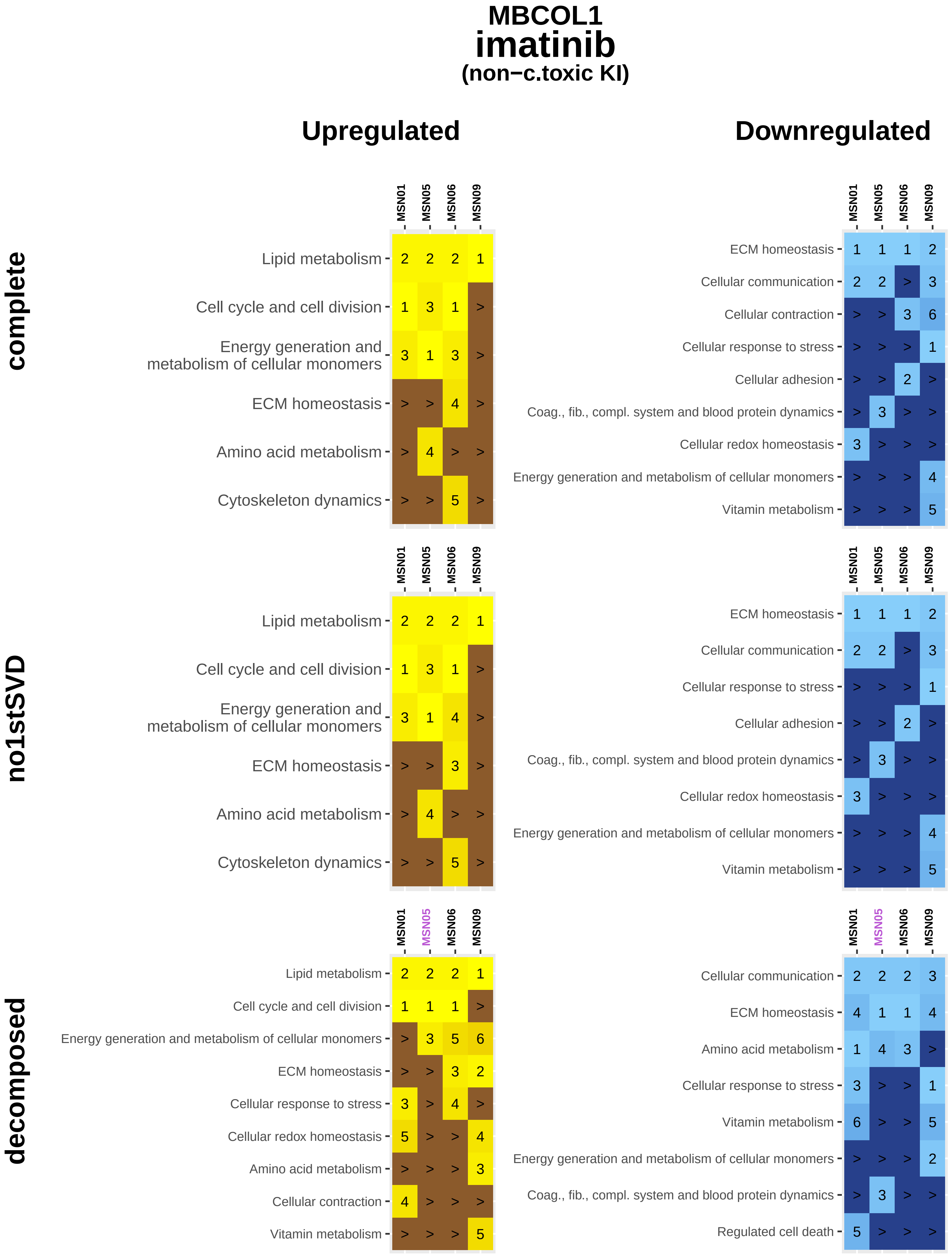
MBCOL1
imatinib
(non−c.toxic KI)
Upregulated
Downregulated
complete
no1stSVD
decomposed
MSN01
MSN05
MSN06
MSN09
Lipid metabolism
Cell cycle and cell division
Energy generation and metabolism of cellular monomers
ECM homeostasis
Amino acid metabolism
Cytoskeleton dynamics
Cellular communication
Cellular contraction
Cellular response to stress
Cellular adhesion
Coag., fib., compl. system and blood protein dynamics
Cellular redox homeostasis
Vitamin metabolism
Regulated cell death

**MBCOL1**

### lapatinib

**(c.toxic KI)**

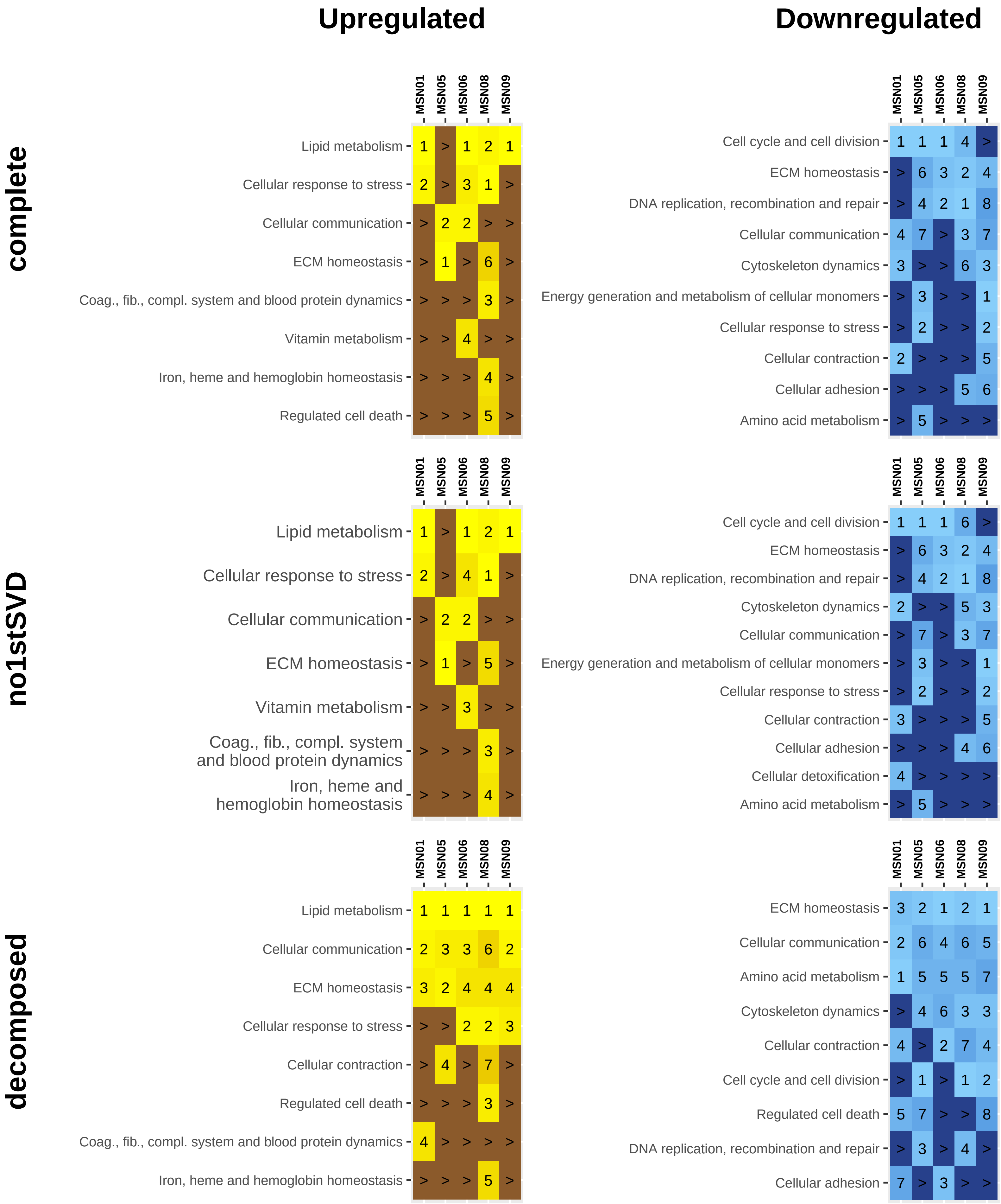

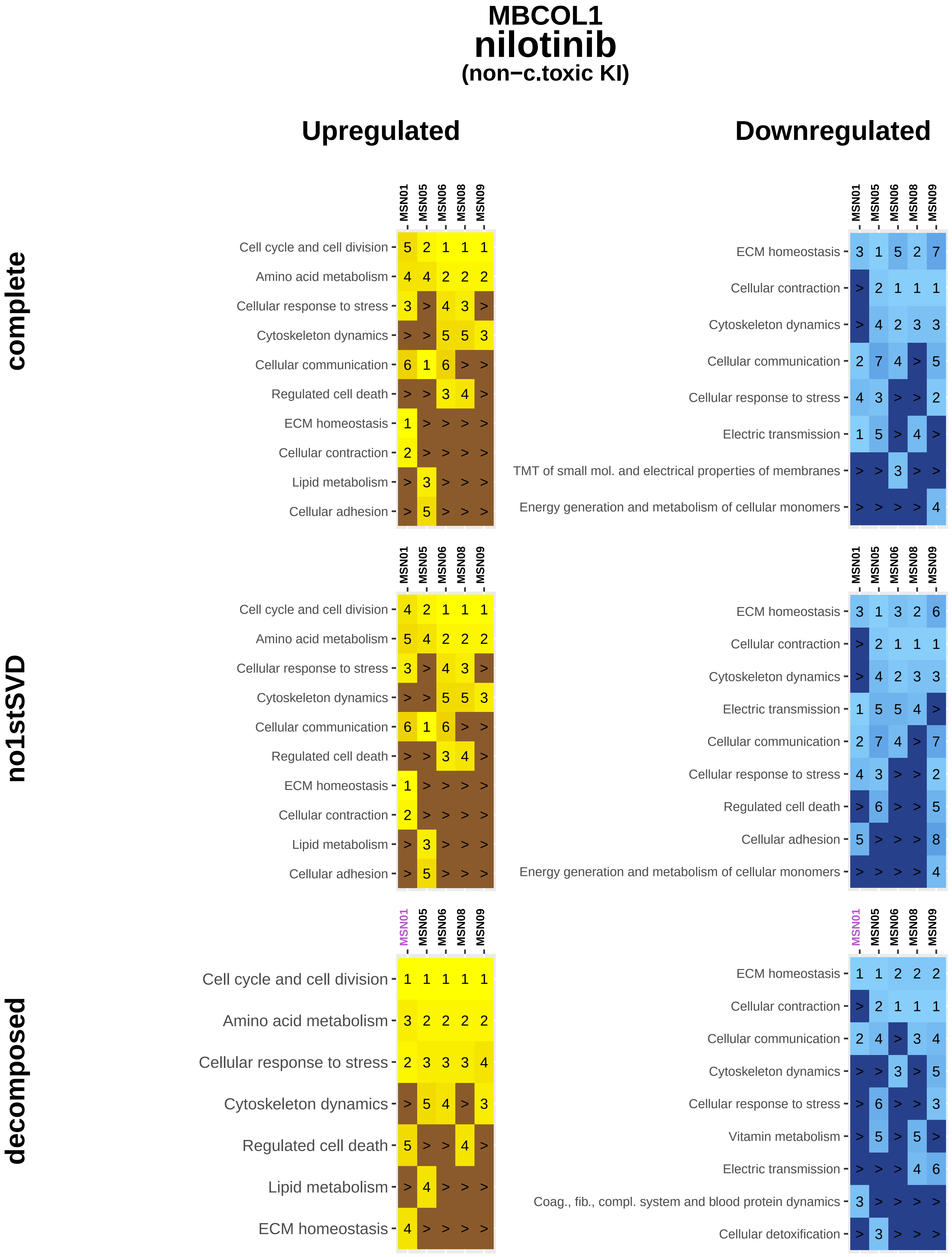
MBCOL1
nilotinib
(non−c.toxic KI)
Upregulated
Downregulated
complete
MSN01 MSN05 MSN06 MSN08 MSN09
Cell cycle and cell division 5 2 1 1 1
Amino acid metabolism 4 4 2 2 2
Cellular response to stress 3 > 4 3 >
Cytoskeleton dynamics > > 5 5 3
Cellular communication 6 1 6 > >
Regulated cell death > > 3 4 >
ECM homeostasis 1 > > > >
Cellular contraction 2 > > > >
Lipid metabolism > 3 > > >
Cellular adhesion > 5 > > >
MSN01 MSN05 MSN06 MSN08 MSN09
ECM homeostasis 3 1 5 2 7
Cellular contraction > 2 1 1 1
Cytoskeleton dynamics > 4 2 3 3
Cellular communication 2 7 4 > 5
Cellular response to stress 4 3 > > 2
Electric transmission 1 5 > 4 >
TMT of small mol. and electrical properties of membranes > > 3 > >
Energy generation and metabolism of cellular monomers > > > > 4
no1stSVD
MSN01 MSN05 MSN06 MSN08 MSN09
Cell cycle and cell division 4 2 1 1 1
Amino acid metabolism 5 4 2 2 2
Cellular response to stress 3 > 4 3 >
Cytoskeleton dynamics > > 5 5 3
Cellular communication 6 1 6 > >
Regulated cell death > > 3 4 >
ECM homeostasis 1 > > > >
Cellular contraction 2 > > > >
Lipid metabolism > 3 > > >
Cellular adhesion > 5 > > >
MSN01 MSN05 MSN06 MSN08 MSN09
ECM homeostasis 3 1 3 2 6
Cellular contraction > 2 1 1 1
Cytoskeleton dynamics > 4 2 3 3
Electric transmission 1 5 5 4 >
Cellular communication 2 7 4 > 7
Cellular response to stress 4 3 > > 2
Regulated cell death > 6 > > 5
Cellular adhesion 5 > > > 8
Energy generation and metabolism of cellular monomers > > > > 4
decomposed
MSN01 MSN05 MSN06 MSN08 MSN09
Cell cycle and cell division 1 1 1 1 1
Amino acid metabolism 3 2 2 2 2
Cellular response to stress 2 3 3 3 4
Cytoskeleton dynamics > 5 4 > 3
Regulated cell death 5 > > 4 >
Lipid metabolism > 4 > > >
ECM homeostasis 4 > > > >
MSN01 MSN05 MSN06 MSN08 MSN09
ECM homeostasis 1 1 2 2 2
Cellular contraction > 2 1 1 1
Cellular communication 2 4 > 3 4
Cytoskeleton dynamics > > 3 > 5
Cellular response to stress > 6 > > 3
Vitamin metabolism > 5 > 5 >
Electric transmission > > > 4 6
Coag., fib., compl. system and blood protein dynamics 3 > > > >
Cellular detoxification > 3 > > >

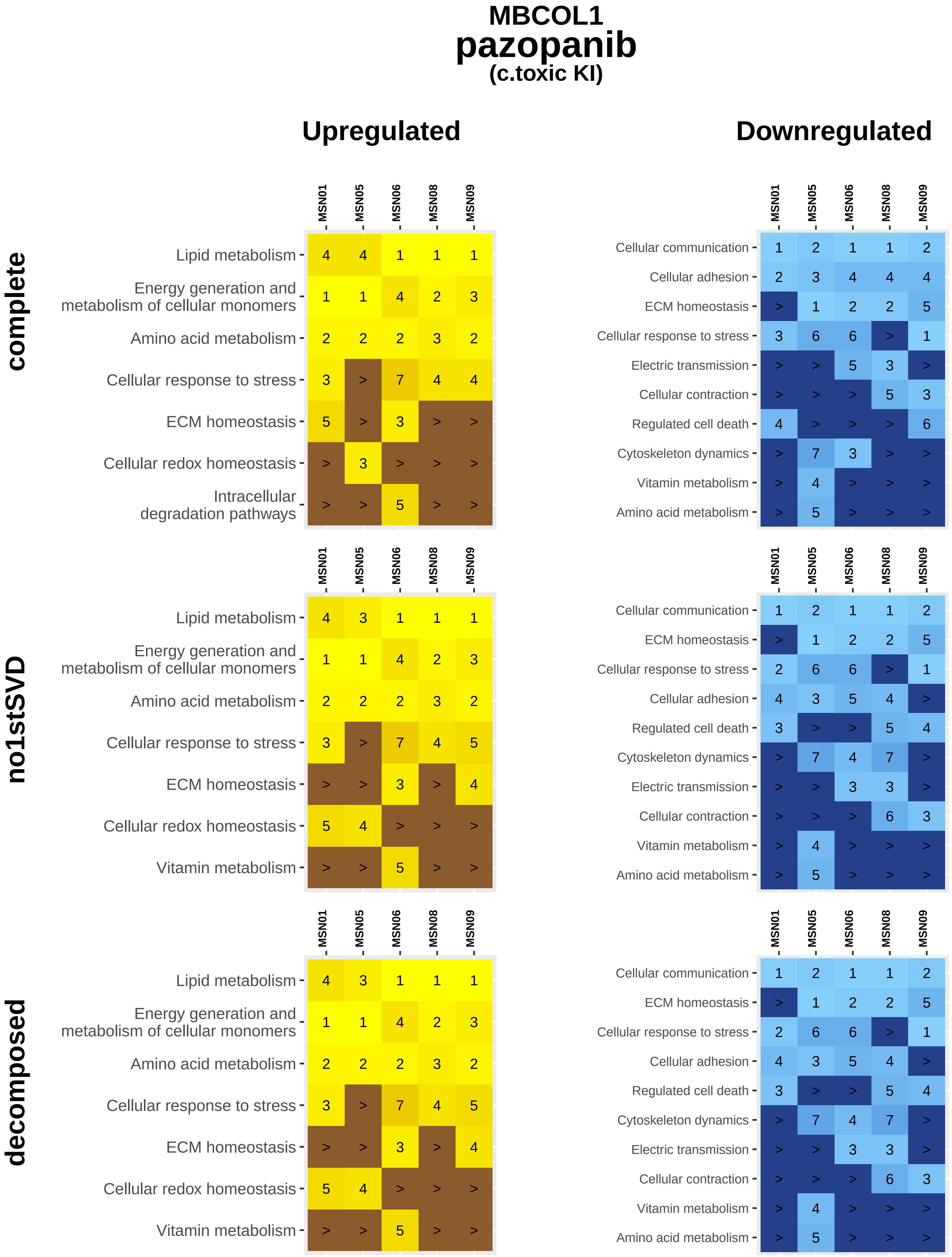
MBCOL1
pazopanib
(c.toxic KI)
Upregulated
Downregulated
complete
no1stSVD
decomposed
MSN01 MSN05 MSN06 MSN08 MSN09
Lipid metabolism 4 4 1 1 1
Energy generation and metabolism of cellular monomers 1 1 4 2 3
Amino acid metabolism 2 2 2 3 2
Cellular response to stress 3 > 7 4 4
ECM homeostasis 5 > 3 > >
Cellular redox homeostasis > 3 > > >
Intracellular degradation pathways > > 5 > >
Cellular communication 1 2 1 1 2
Cellular adhesion 2 3 4 4 4
ECM homeostasis > 1 2 2 5
Cellular response to stress 3 6 6 > 1
Electric transmission > > 5 3 >
Cellular contraction > > > 5 3
Regulated cell death 4 > > > 6
Cytoskeleton dynamics > 7 3 > >
Vitamin metabolism > 4 > > >
Amino acid metabolism > 5 > > >
Lipid metabolism 4 3 1 1 1
Energy generation and metabolism of cellular monomers 1 1 4 2 3
Amino acid metabolism 2 2 2 3 2
Cellular response to stress 3 > 7 4 5
ECM homeostasis > > 3 > 4
Cellular redox homeostasis 5 4 > > >
Vitamin metabolism > > 5 > >
Cellular communication 1 2 1 1 2
ECM homeostasis > 1 2 2 5
Cellular response to stress 2 6 6 > 1
Cellular adhesion 4 3 5 4 >
Regulated cell death 3 > > 5 4
Cytoskeleton dynamics > 7 4 7 >
Electric transmission > > 3 3 >
Cellular contraction > > > 6 3
Vitamin metabolism > 4 > > >
Amino acid metabolism > 5 > > >
Lipid metabolism 4 3 1 1 1
Energy generation and metabolism of cellular monomers 1 1 4 2 3
Amino acid metabolism 2 2 2 3 2
Cellular response to stress 3 > 7 4 5
ECM homeostasis > > 3 > 4
Cellular redox homeostasis 5 4 > > >
Vitamin metabolism > > 5 > >
Cellular communication 1 2 1 1 2
ECM homeostasis > 1 2 2 5
Cellular response to stress 2 6 6 > 1
Cellular adhesion 4 3 5 4 >
Regulated cell death 3 > > 5 4
Cytoskeleton dynamics > 7 4 7 >
Electric transmission > > 3 3 >
Cellular contraction > > > 6 3
Vitamin metabolism > 4 > > >
Amino acid metabolism > 5 > > >

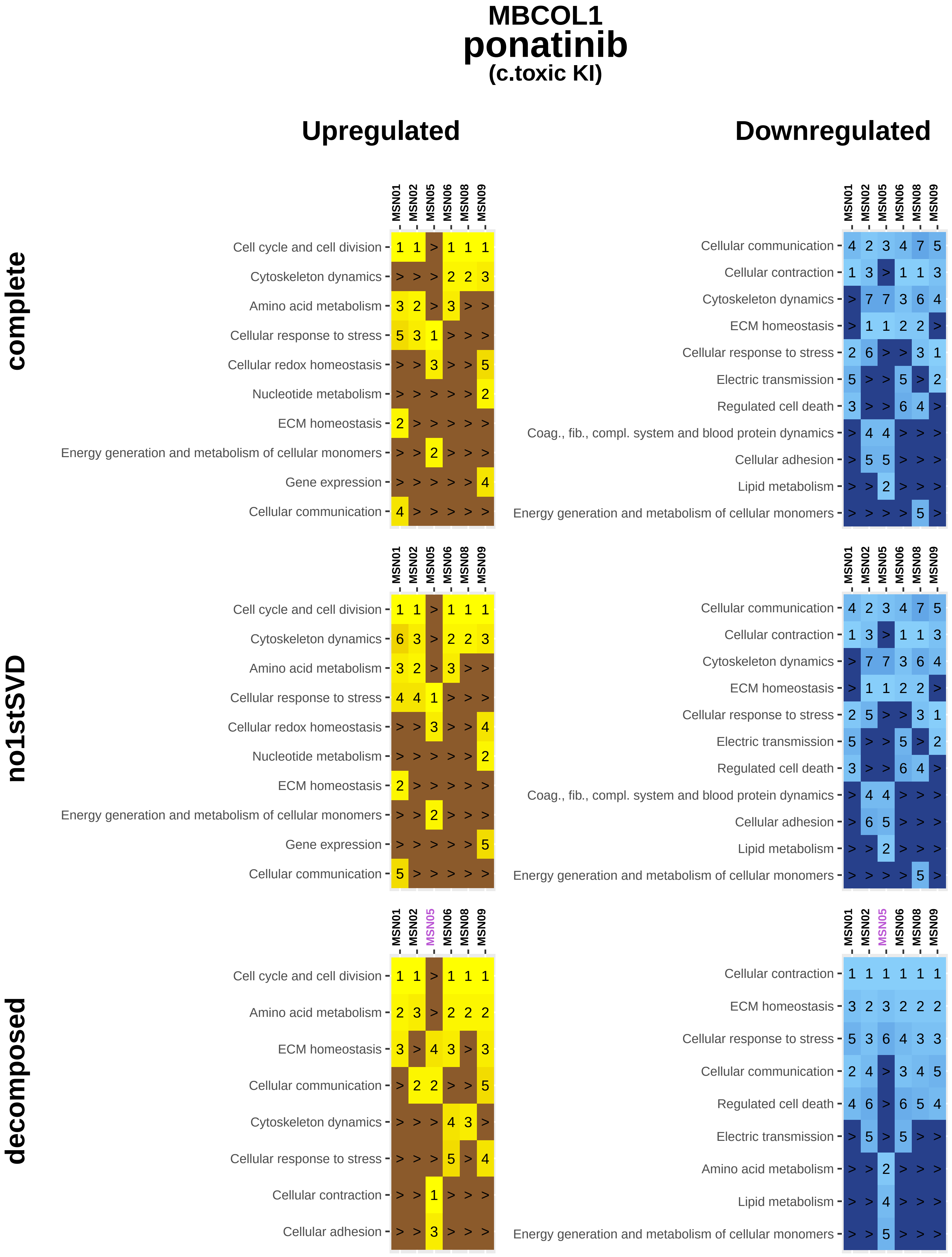
MBCOL1
ponatinib
(c.toxic KI)
Upregulated
Downregulated
complete
no1stSVD
decomposed
MSN01 MSN02 MSN05 MSN06 MSN08 MSN09
Cell cycle and cell division 1 1 > 1 1 1
Cytoskeleton dynamics > > > 2 2 3
Amino acid metabolism 3 2 > 3 > >
Cellular response to stress 5 3 1 > > >
Cellular redox homeostasis > > 3 > > 5
Nucleotide metabolism > > > > > 2
ECM homeostasis 2 > > > > >
Energy generation and metabolism of cellular monomers > > 2 > > >
Gene expression > > > > > 4
Cellular communication 4 > > > > >
MSN01 MSN02 MSN05 MSN06 MSN08 MSN09
Cellular communication 4 2 3 4 7 5
Cellular contraction 1 3 > 1 1 3
Cytoskeleton dynamics > 7 7 3 6 4
ECM homeostasis > 1 1 2 2 >
Cellular response to stress 2 6 > > 3 1
Electric transmission 5 > > 5 > 2
Regulated cell death 3 > > 6 4 >
Coag., fib., compl. system and blood protein dynamics > 4 4 > > >
Cellular adhesion > 5 5 > > >
Lipid metabolism > > 2 > > >
Energy generation and metabolism of cellular monomers > > > > 5 >
MSN01 MSN02 MSN05 MSN06 MSN08 MSN09
Cell cycle and cell division 1 1 > 1 1 1
Cytoskeleton dynamics 6 3 > 2 2 3
Amino acid metabolism 3 2 > 3 > >
Cellular response to stress 4 4 1 > > >
Cellular redox homeostasis > > 3 > > 4
Nucleotide metabolism > > > > > 2
ECM homeostasis 2 > > > > >
Energy generation and metabolism of cellular monomers > > 2 > > >
Gene expression > > > > > 5
Cellular communication 5 > > > > >
MSN01 MSN02 MSN05 MSN06 MSN08 MSN09
Cellular communication 4 2 3 4 7 5
Cellular contraction 1 3 > 1 1 3
Cytoskeleton dynamics > 7 7 3 6 4
ECM homeostasis > 1 1 2 2 >
Cellular response to stress 2 5 > > 3 1
Electric transmission 5 > > 5 > 2
Regulated cell death 3 > > 6 4 >
Coag., fib., compl. system and blood protein dynamics > 4 4 > > >
Cellular adhesion > 6 5 > > >
Lipid metabolism > > 2 > > >
Energy generation and metabolism of cellular monomers > > > > 5 >
MSN01 MSN02 MSN05 MSN06 MSN08 MSN09
Cell cycle and cell division 1 1 > 1 1 1
Amino acid metabolism 2 3 > 2 2 2
ECM homeostasis 3 > 4 3 > 3
Cellular communication > 2 2 > > 5
Cytoskeleton dynamics > > > 4 3 >
Cellular response to stress > > > 5 > 4
Cellular contraction > > 1 > > >
Cellular adhesion > > 3 > > >
MSN01 MSN02 MSN05 MSN06 MSN08 MSN09
Cellular contraction 1 1 1 1 1 1
ECM homeostasis 3 2 3 2 2 2
Cellular response to stress 5 3 6 4 3 3
Cellular communication 2 4 > 3 4 5
Regulated cell death 4 6 > 6 5 4
Electric transmission > 5 > 5 > >
Amino acid metabolism > > 2 > > >
Lipid metabolism > > 4 > > >
Energy generation and metabolism of cellular monomers > > 5 > > >

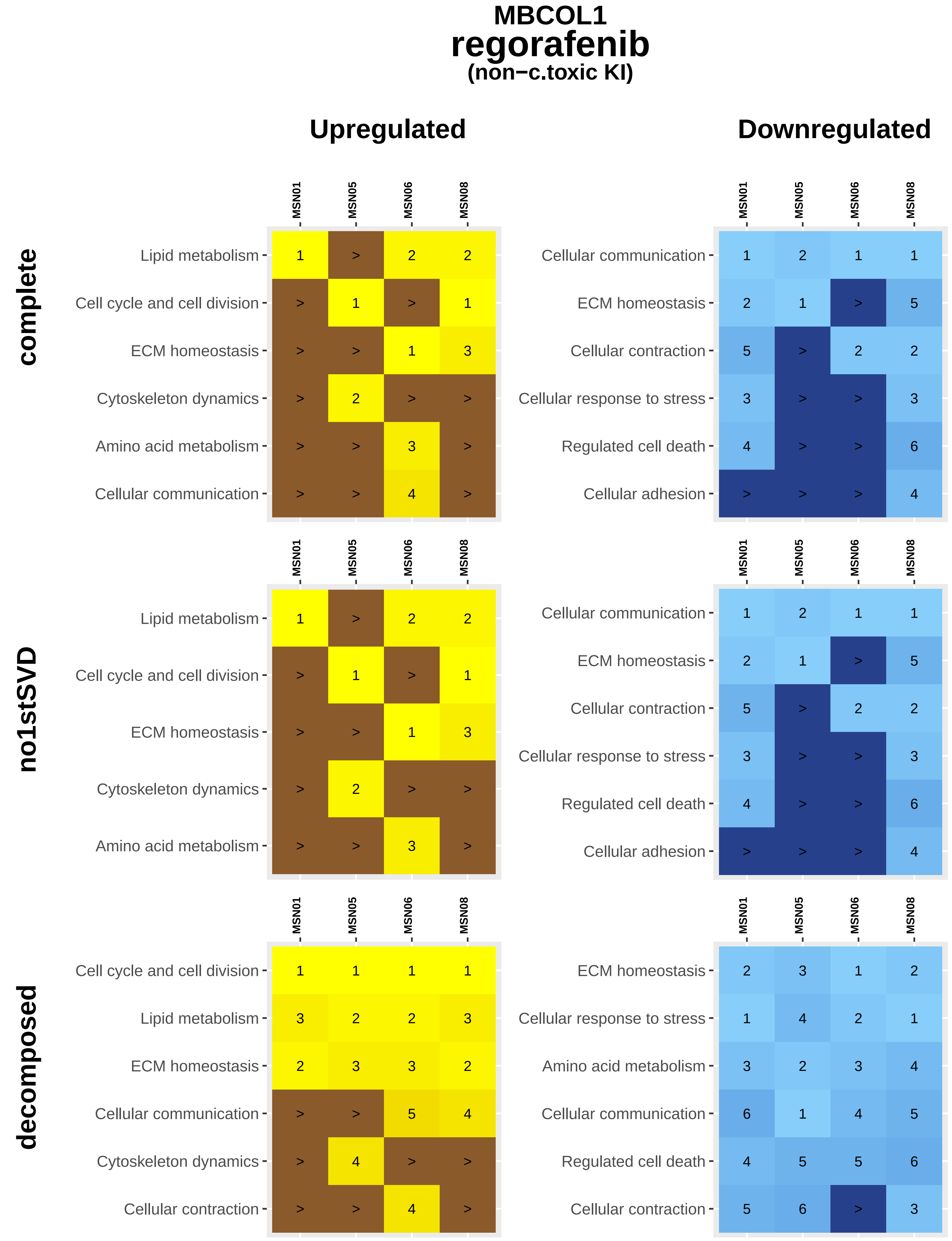
MBCOL1
regorafenib
(non−c.toxic KI)
Upregulated
Downregulated
complete
no1stSVD
decomposed
MSN01
MSN05
MSN06
MSN08
Lipid metabolism 1 > 2 2
Cell cycle and cell division > 1 > 1
ECM homeostasis > > 1 3
Cytoskeleton dynamics > 2 > >
Amino acid metabolism > > 3 >
Cellular communication > > 4 >
Cellular communication 1 2 1 1
ECM homeostasis 2 1 > 5
Cellular contraction 5 > 2 2
Cellular response to stress 3 > > 3
Regulated cell death 4 > > 6
Cellular adhesion > > > 4
Lipid metabolism 1 > 2 2
Cell cycle and cell division > 1 > 1
ECM homeostasis > > 1 3
Cytoskeleton dynamics > 2 > >
Amino acid metabolism > > 3 >
Cellular communication 1 2 1 1
ECM homeostasis 2 1 > 5
Cellular contraction 5 > 2 2
Cellular response to stress 3 > > 3
Regulated cell death 4 > > 6
Cellular adhesion > > > 4
Cell cycle and cell division 1 1 1 1
Lipid metabolism 3 2 2 3
ECM homeostasis 2 3 3 2
Cellular communication > > 5 4
Cytoskeleton dynamics > 4 > >
Cellular contraction > > 4 >
ECM homeostasis 2 3 1 2
Cellular response to stress 1 4 2 1
Amino acid metabolism 3 2 3 4
Cellular communication 6 1 4 5
Regulated cell death 4 5 5 6
Cellular contraction 5 6 > 3

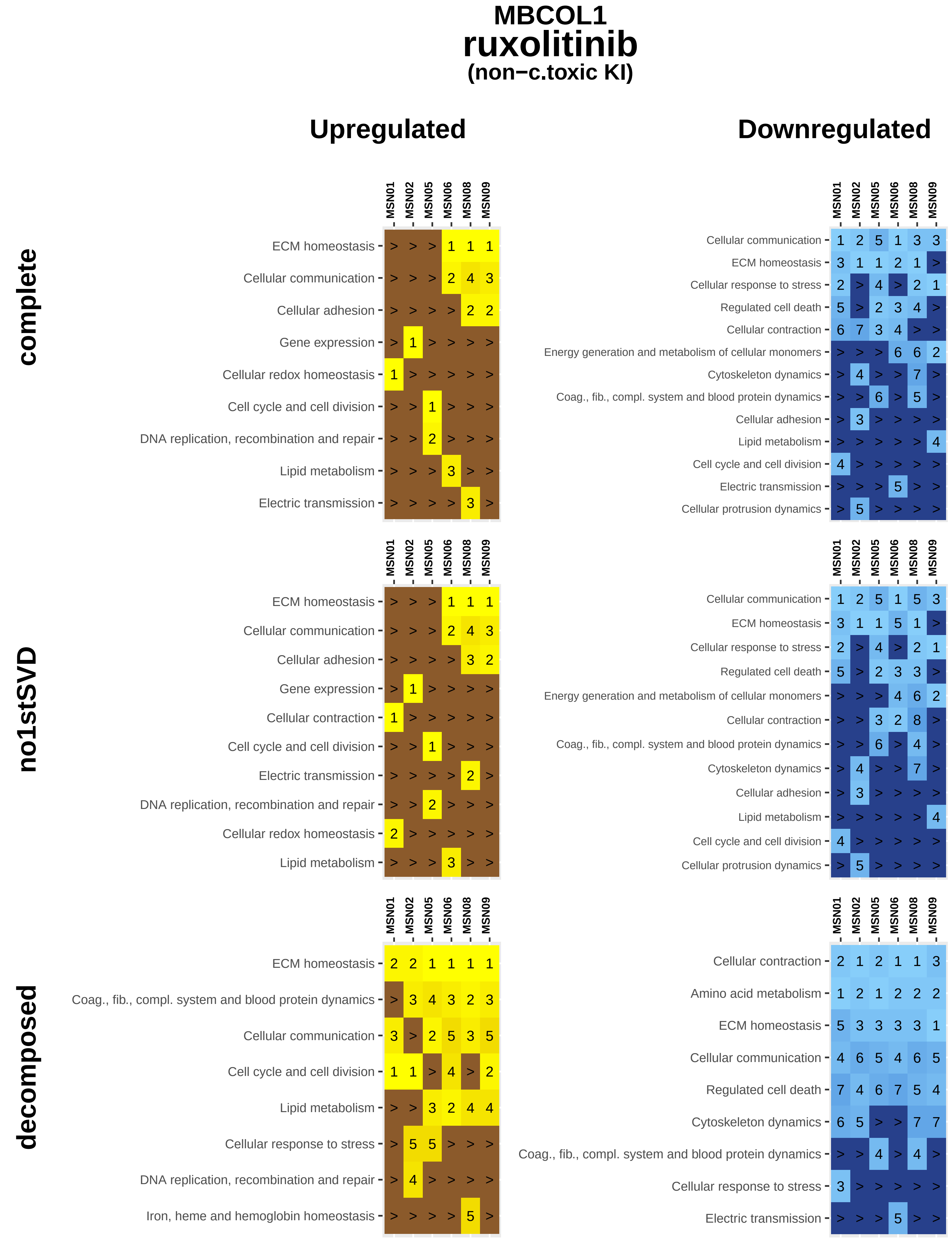
MBCOL1
ruxolitinib
(non−c.toxic KI)
Upregulated
Downregulated
complete
no1stSVD
decomposed
MSN01
MSN02
MSN05
MSN06
MSN08
MSN09
ECM homeostasis
Cellular communication
Cellular adhesion
Gene expression
Cellular redox homeostasis
Cell cycle and cell division
DNA replication, recombination and repair
Lipid metabolism
Electric transmission
Cellular response to stress
Regulated cell death
Cellular contraction
Energy generation and metabolism of cellular monomers
Cytoskeleton dynamics
Coag., fib., compl. system and blood protein dynamics
Cellular protrusion dynamics
Amino acid metabolism
Iron, heme and hemoglobin homeostasis

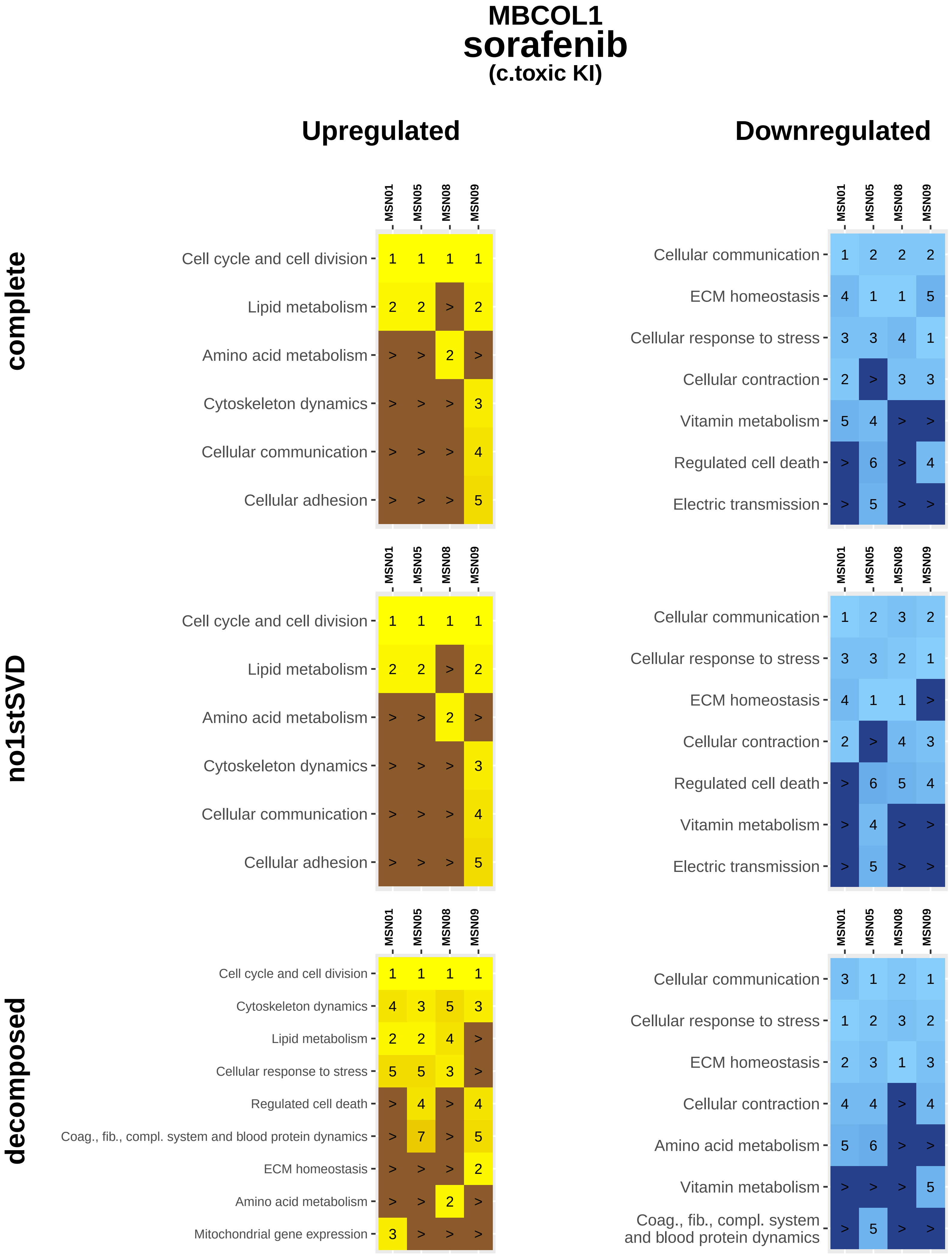
MBCOL1
sorafenib
(c.toxic KI)
Upregulated
Downregulated
complete
no1stSVD
decomposed
MSN01
MSN05
MSN08
MSN09
Cell cycle and cell division
Lipid metabolism
Amino acid metabolism
Cytoskeleton dynamics
Cellular communication
Cellular adhesion
Cellular response to stress
Regulated cell death
Coag., fib., compl. system and blood protein dynamics
ECM homeostasis
Mitochondrial gene expression
Cellular contraction
Vitamin metabolism
Electric transmission

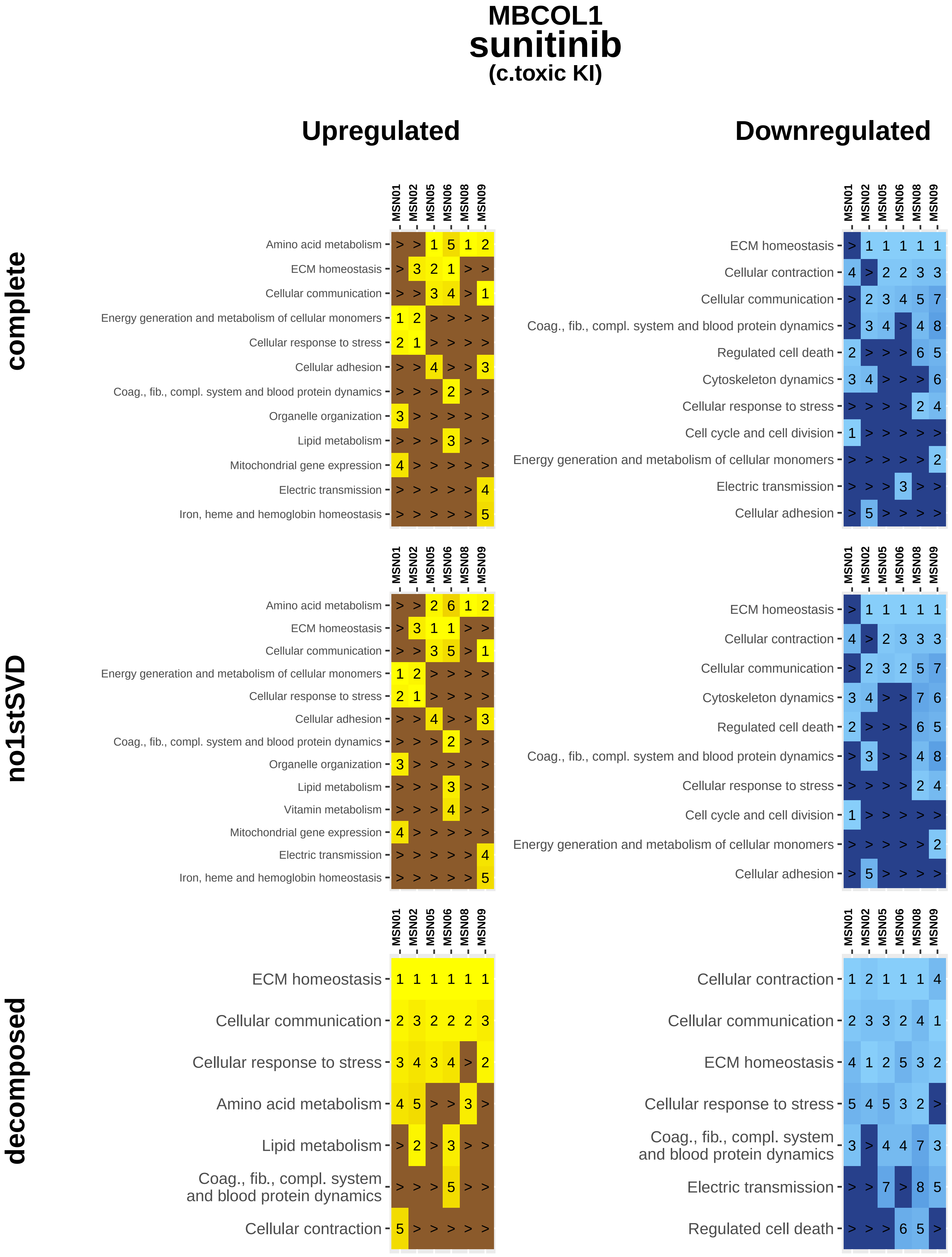

MBCOL1
sunitinib
(c.toxic KI)
Upregulated
Downregulated
complete
no1stSVD
decomposed
MSN01
MSN02
MSN05
MSN06
MSN08
MSN09
Amino acid metabolism
ECM homeostasis
Cellular communication
Energy generation and metabolism of cellular monomers
Cellular response to stress
Cellular adhesion
Coag., fib., compl. system and blood protein dynamics
Organelle organization
Lipid metabolism
Mitochondrial gene expression
Electric transmission
Iron, heme and hemoglobin homeostasis
Vitamin metabolism
Cellular contraction
Regulated cell death
Cytoskeleton dynamics
Cell cycle and cell division

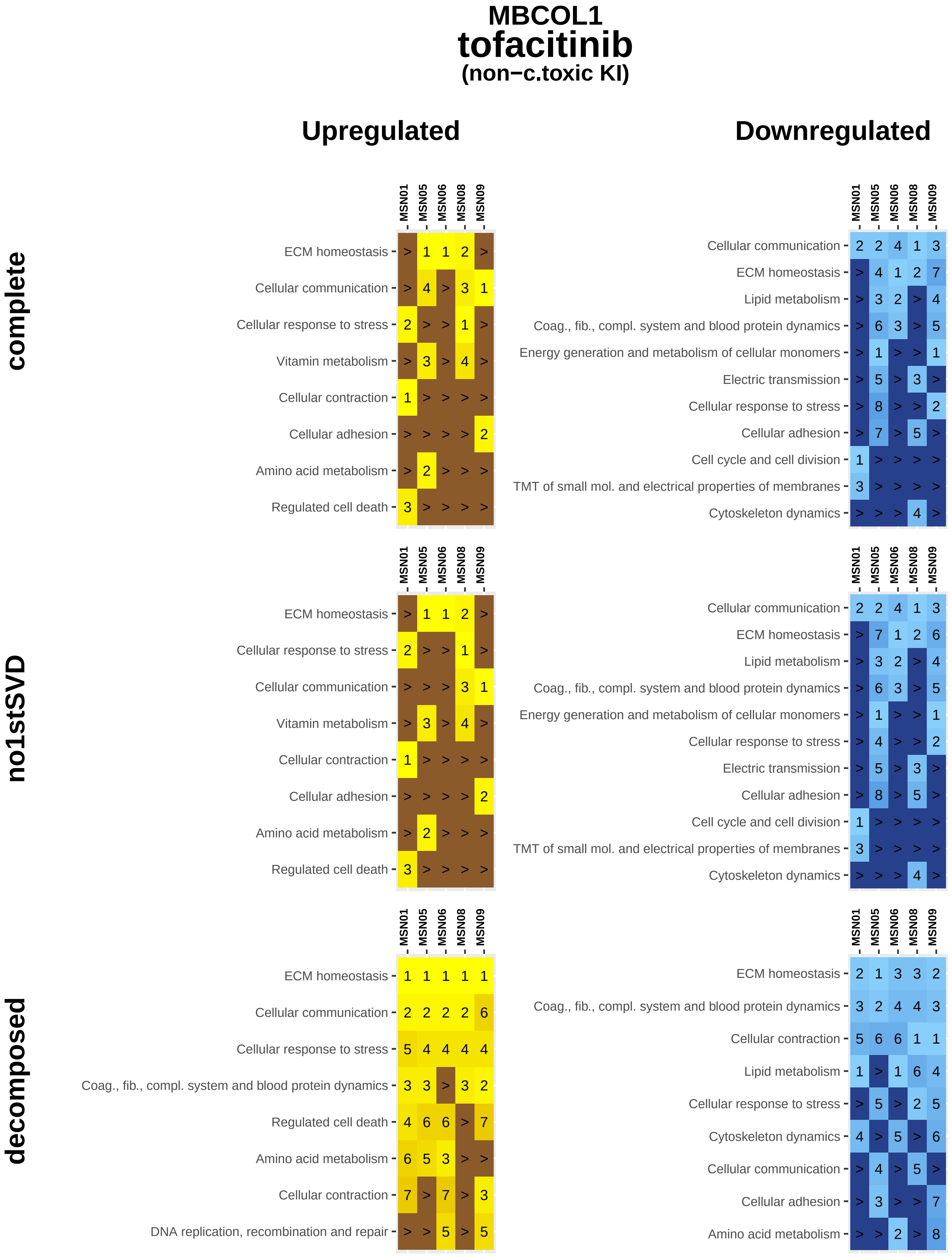
MBCOL1
tofacitinib
(non−c.toxic KI)
Upregulated
Downregulated
complete
no1stSVD
decomposed
MSN01
MSN05
MSN06
MSN08
MSN09
ECM homeostasis
Cellular communication
Cellular response to stress
Vitamin metabolism
Cellular contraction
Cellular adhesion
Amino acid metabolism
Regulated cell death
Lipid metabolism
Coag., fib., compl. system and blood protein dynamics
Energy generation and metabolism of cellular monomers
Electric transmission
Cell cycle and cell division
TMT of small mol. and electrical properties of membranes
Cytoskeleton dynamics
DNA replication, recombination and repair

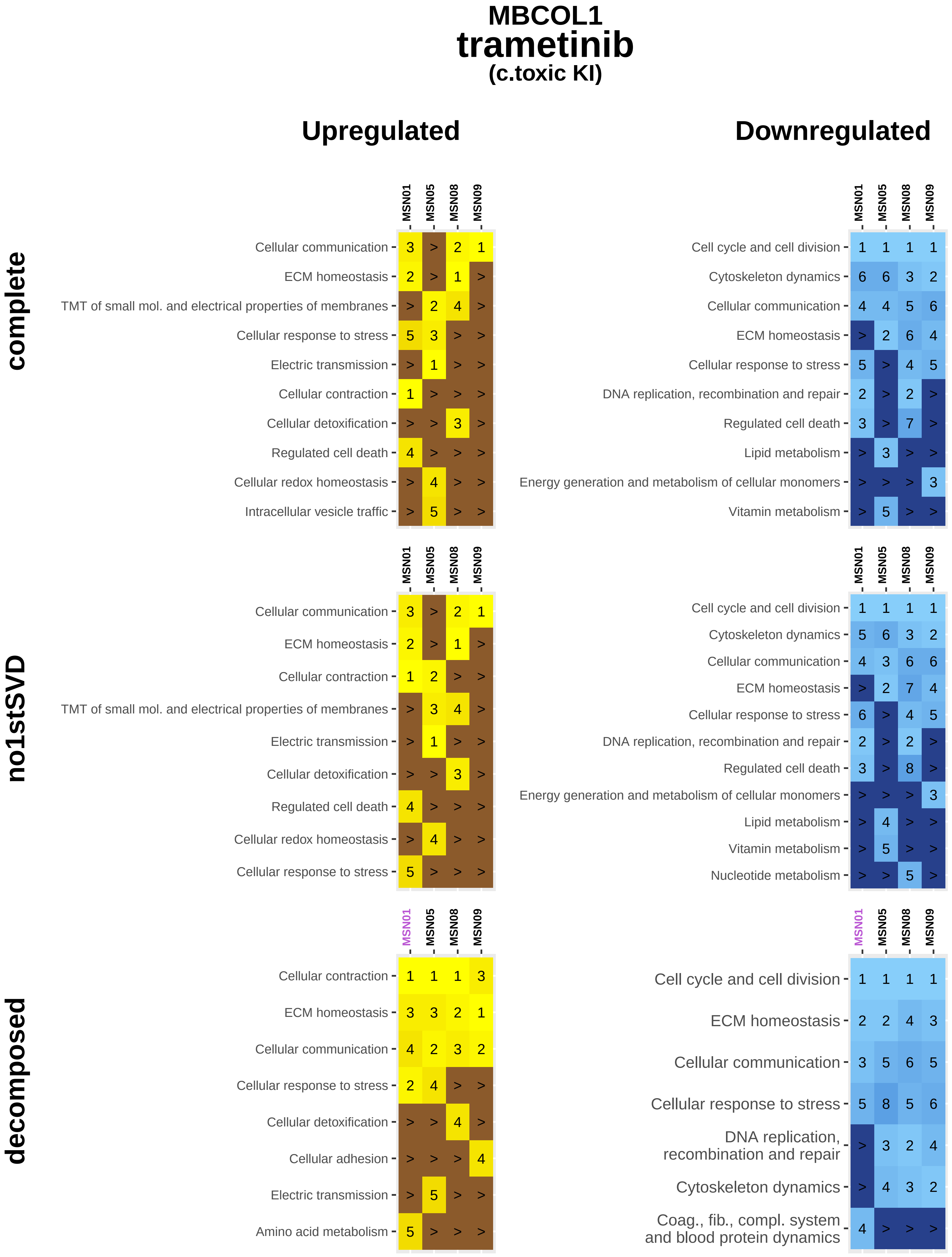

MBCOL1
trametinib
(c.toxic KI)
Upregulated
Downregulated
complete
no1stSVD
decomposed
MSN01
MSN05
MSN08
MSN09
Cellular communication
ECM homeostasis
TMT of small mol. and electrical properties of membranes
Cellular response to stress
Electric transmission
Cellular contraction
Cellular detoxification
Regulated cell death
Cellular redox homeostasis
Intracellular vesicle traffic
Cell cycle and cell division
Cytoskeleton dynamics
DNA replication, recombination and repair
Lipid metabolism
Energy generation and metabolism of cellular monomers
Vitamin metabolism
Nucleotide metabolism
Cellular adhesion
Amino acid metabolism
Coag., fib., compl. system and blood protein dynamics

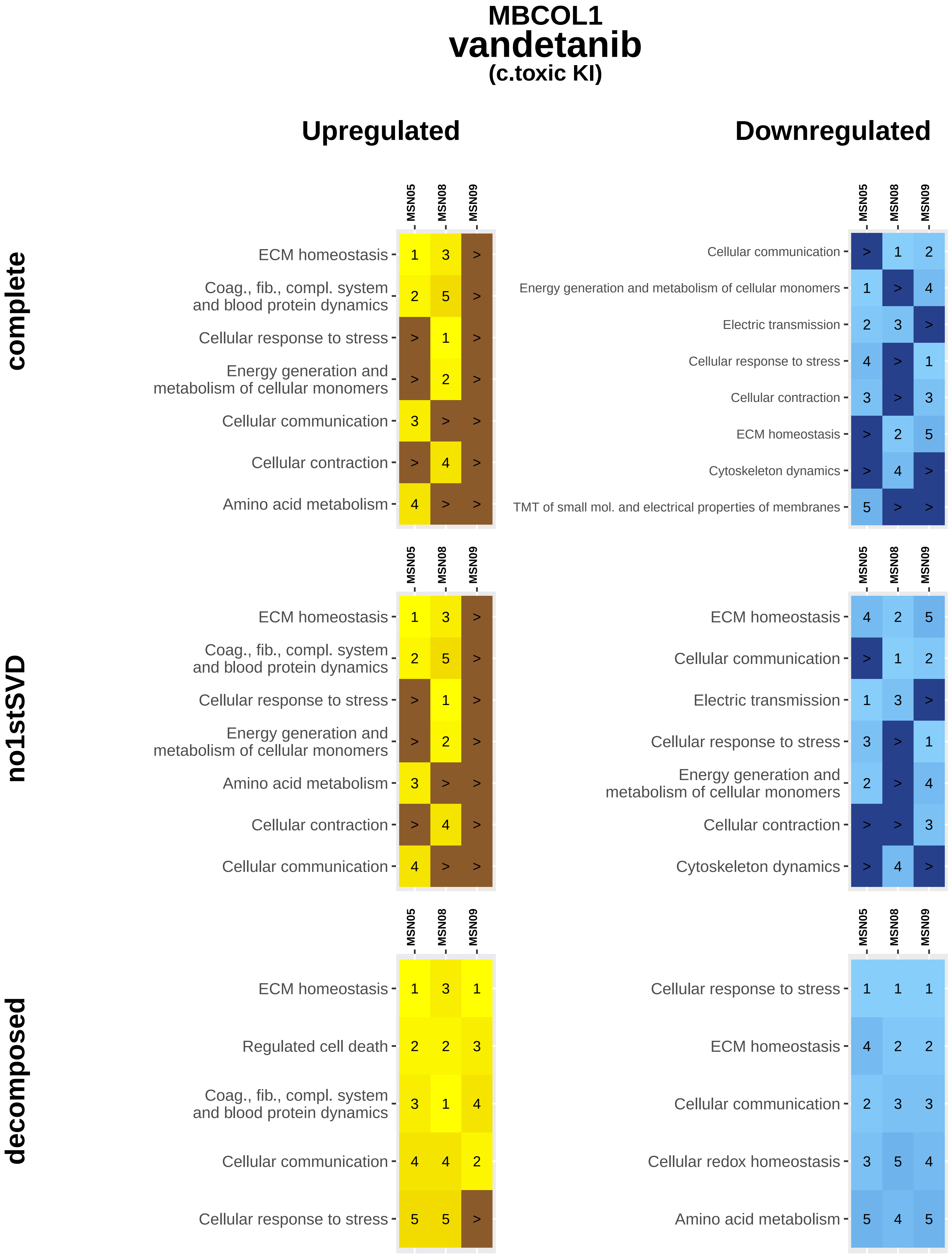

MBCOL1
vandetanib
(c.toxic KI)
Upregulated
Downregulated
complete
no1stSVD
decomposed
MSN05
MSN08
MSN09
ECM homeostasis
Coag., fib., compl. system and blood protein dynamics
Cellular response to stress
Energy generation and metabolism of cellular monomers
Cellular communication
Cellular contraction
Amino acid metabolism
Energy generation and metabolism of cellular monomers
Electric transmission
Cytoskeleton dynamics
TMT of small mol. and electrical properties of membranes
Regulated cell death
Cellular redox homeostasis

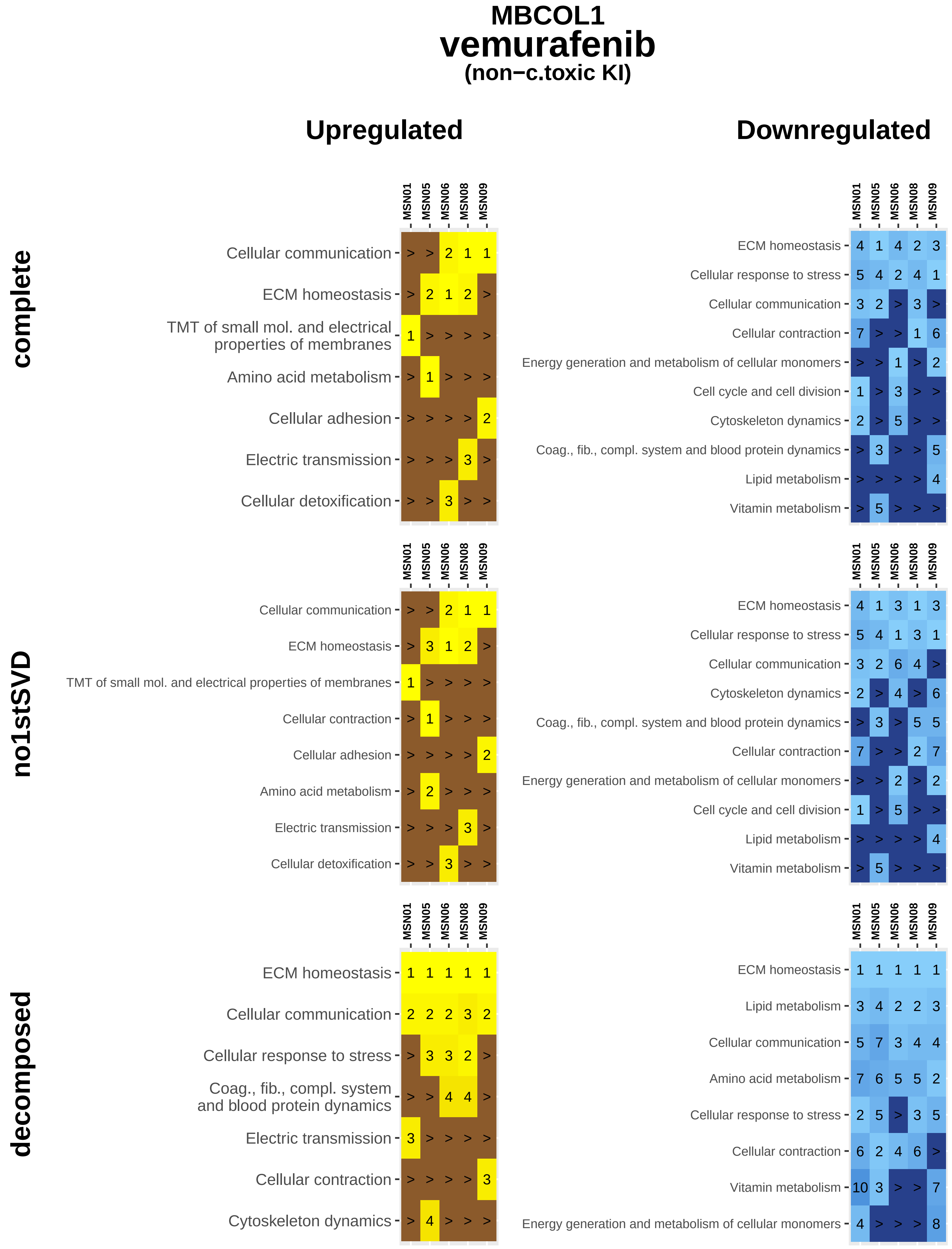
MBCOL1
vemurafenib
(non−c.toxic KI)
Upregulated
Downregulated
complete
no1stSVD
decomposed
MSN01
MSN05
MSN06
MSN08
MSN09
Cellular communication > > 2 1 1
ECM homeostasis > 2 1 2 >
TMT of small mol. and electrical properties of membranes 1 > > > >
Amino acid metabolism > 1 > > >
Cellular adhesion > > > > 2
Electric transmission > > > 3 >
Cellular detoxification > > 3 > >
ECM homeostasis 4 1 4 2 3
Cellular response to stress 5 4 2 4 1
Cellular communication 3 2 > 3 >
Cellular contraction 7 > > 1 6
Energy generation and metabolism of cellular monomers > > 1 > 2
Cell cycle and cell division 1 > 3 > >
Cytoskeleton dynamics 2 > 5 > >
Coag., fib., compl. system and blood protein dynamics > 3 > > 5
Lipid metabolism > > > > 4
Vitamin metabolism > 5 > > >
Cellular communication > > 2 1 1
ECM homeostasis > 3 1 2 >
TMT of small mol. and electrical properties of membranes 1 > > > >
Cellular contraction > 1 > > >
Cellular adhesion > > > > 2
Amino acid metabolism > 2 > > >
Electric transmission > > > 3 >
Cellular detoxification > > 3 > >
ECM homeostasis 4 1 3 1 3
Cellular response to stress 5 4 1 3 1
Cellular communication 3 2 6 4 >
Cytoskeleton dynamics 2 > 4 > 6
Coag., fib., compl. system and blood protein dynamics > 3 > 5 5
Cellular contraction 7 > > 2 7
Energy generation and metabolism of cellular monomers > > 2 > 2
Cell cycle and cell division 1 > 5 > >
Lipid metabolism > > > > 4
Vitamin metabolism > 5 > > >
ECM homeostasis 1 1 1 1 1
Cellular communication 2 2 2 3 2
Cellular response to stress > 3 3 2 >
Coag., fib., compl. system and blood protein dynamics > > 4 4 >
Electric transmission 3 > > > >
Cellular contraction > > > > 3
Cytoskeleton dynamics > 4 > > >
ECM homeostasis 1 1 1 1 1
Lipid metabolism 3 4 2 2 3
Cellular communication 5 7 3 4 4
Amino acid metabolism 7 6 5 5 2
Cellular response to stress 2 5 > 3 5
Cellular contraction 6 2 4 6 >
Vitamin metabolism 10 3 > > 7
Energy generation and metabolism of cellular monomers 4 > > > 8

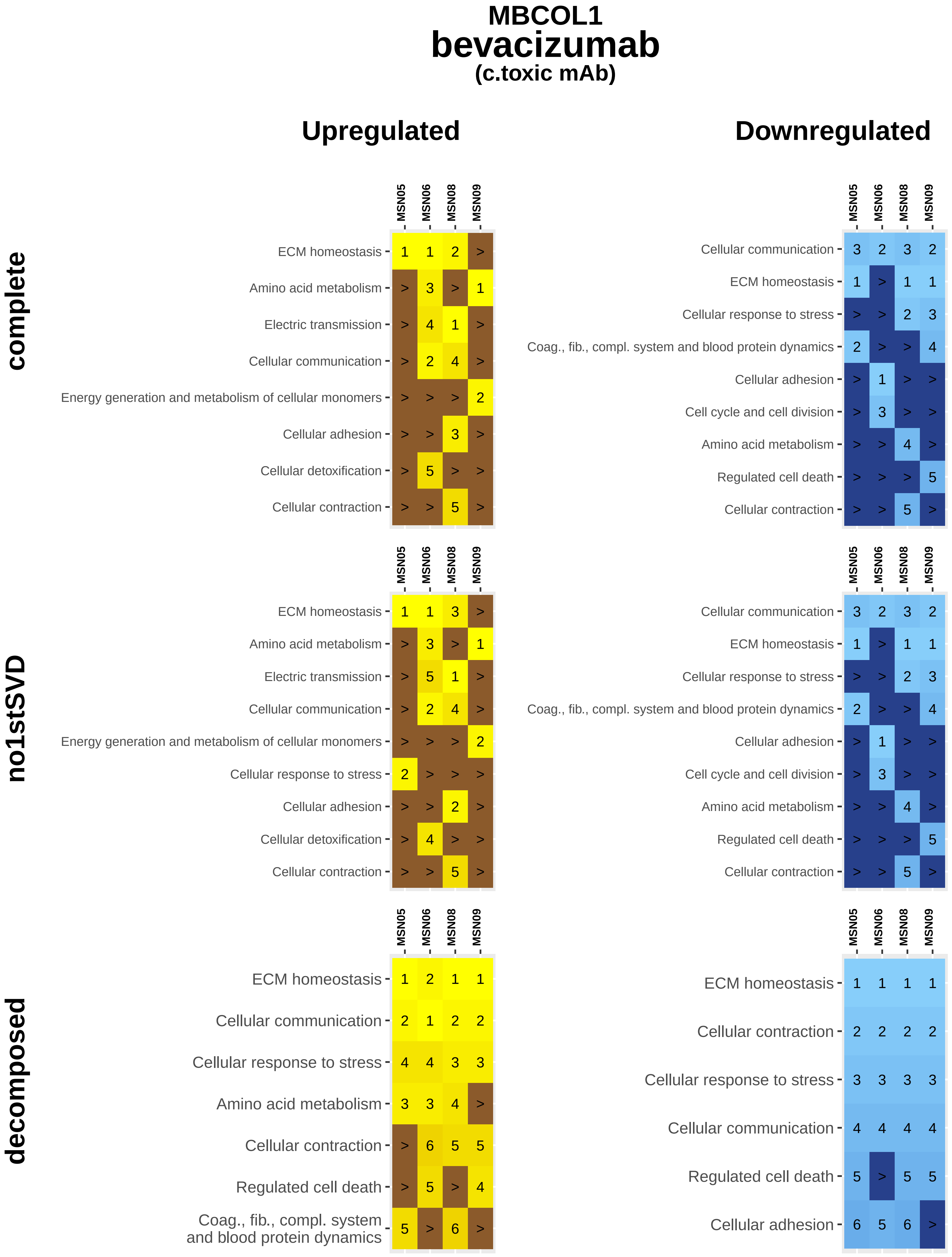
MBCOL1
bevacizumab
(c.toxic mAb)
Upregulated
Downregulated
complete
no1stSVD
decomposed
MSN05
MSN06
MSN08
MSN09
ECM homeostasis 1 1 2 >
Amino acid metabolism > 3 > 1
Electric transmission > 4 1 >
Cellular communication > 2 4 >
Energy generation and metabolism of cellular monomers > > > 2
Cellular adhesion > > 3 >
Cellular detoxification > 5 > >
Cellular contraction > > 5 >
Cellular communication 3 2 3 2
ECM homeostasis 1 > 1 1
Cellular response to stress > > 2 3
Coag., fib., compl. system and blood protein dynamics 2 > > 4
Cellular adhesion > 1 > >
Cell cycle and cell division > 3 > >
Amino acid metabolism > > 4 >
Regulated cell death > > > 5
Cellular contraction > > 5 >
ECM homeostasis 1 1 3 >
Amino acid metabolism > 3 > 1
Electric transmission > 5 1 >
Cellular communication > 2 4 >
Energy generation and metabolism of cellular monomers > > > 2
Cellular response to stress 2 > > >
Cellular adhesion > > 2 >
Cellular detoxification > 4 > >
Cellular contraction > > 5 >
Cellular communication 3 2 3 2
ECM homeostasis 1 > 1 1
Cellular response to stress > > 2 3
Coag., fib., compl. system and blood protein dynamics 2 > > 4
Cellular adhesion > 1 > >
Cell cycle and cell division > 3 > >
Amino acid metabolism > > 4 >
Regulated cell death > > > 5
Cellular contraction > > 5 >
ECM homeostasis 1 2 1 1
Cellular communication 2 1 2 2
Cellular response to stress 4 4 3 3
Amino acid metabolism 3 3 4 >
Cellular contraction > 6 5 5
Regulated cell death > 5 > 4
Coag., fib., compl. system and blood protein dynamics 5 > 6 >
ECM homeostasis 1 1 1 1
Cellular contraction 2 2 2 2
Cellular response to stress 3 3 3 3
Cellular communication 4 4 4 4
Regulated cell death 5 > 5 5
Cellular adhesion 6 5 6 >

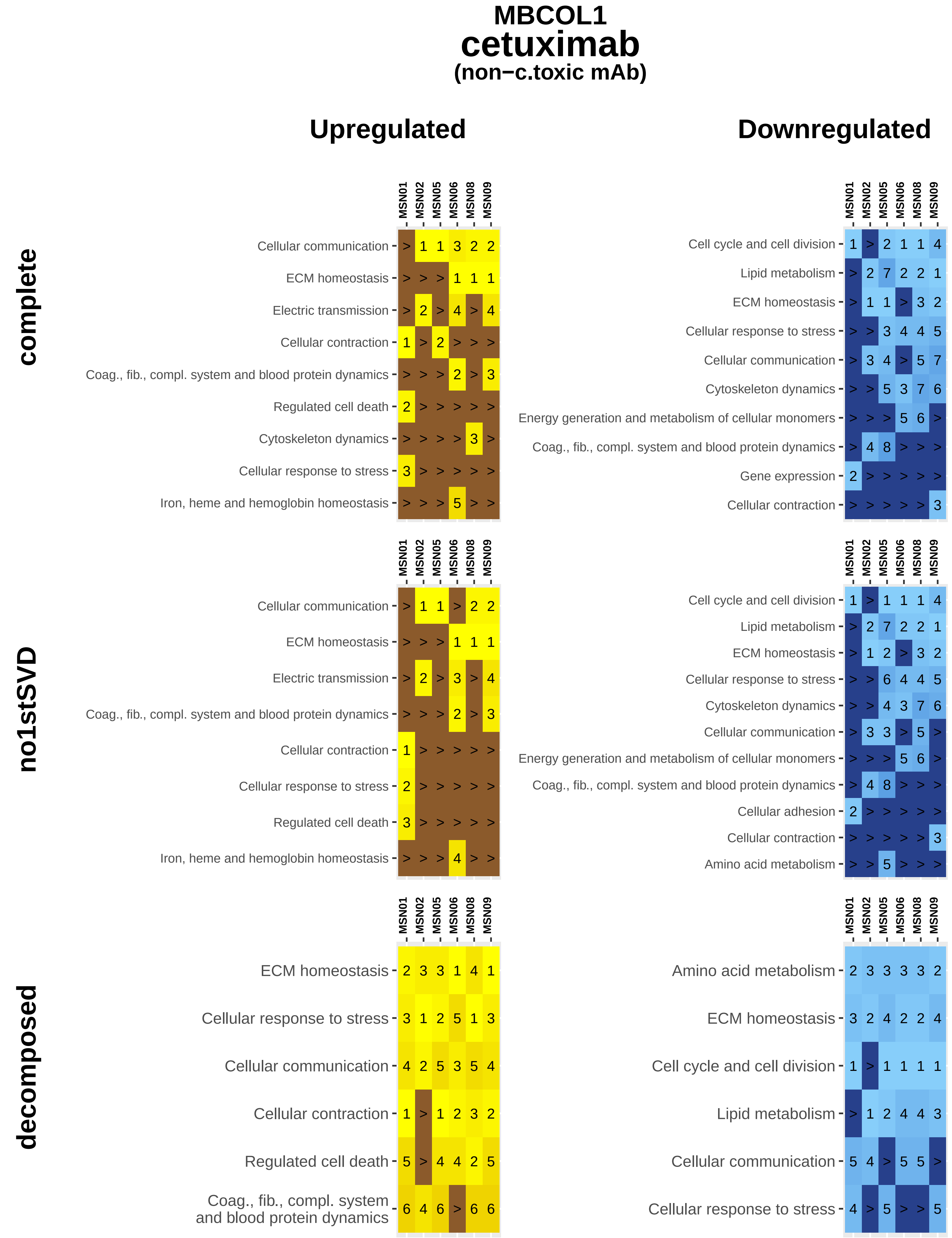
MBCOL1
cetuximab
(non−c.toxic mAb)
Upregulated
Downregulated
complete
MSN01 MSN02 MSN05 MSN06 MSN08 MSN09
Cellular communication > 1 1 3 2 2
ECM homeostasis > > > 1 1 1
Electric transmission > 2 > 4 > 4
Cellular contraction 1 > 2 > > >
Coag., fib., compl. system and blood protein dynamics > > > 2 > 3
Regulated cell death 2 > > > > >
Cytoskeleton dynamics > > > > 3 >
Cellular response to stress 3 > > > > >
Iron, heme and hemoglobin homeostasis > > > 5 > >
MSN01 MSN02 MSN05 MSN06 MSN08 MSN09
Cell cycle and cell division 1 > 2 1 1 4
Lipid metabolism > 2 7 2 2 1
ECM homeostasis > 1 1 > 3 2
Cellular response to stress > > 3 4 4 5
Cellular communication > 3 4 > 5 7
Cytoskeleton dynamics > > 5 3 7 6
Energy generation and metabolism of cellular monomers > > > 5 6 >
Coag., fib., compl. system and blood protein dynamics > 4 8 > > >
Gene expression 2 > > > > >
Cellular contraction > > > > > 3
no1stSVD
MSN01 MSN02 MSN05 MSN06 MSN08 MSN09
Cellular communication > 1 1 > 2 2
ECM homeostasis > > > 1 1 1
Electric transmission > 2 > 3 > 4
Coag., fib., compl. system and blood protein dynamics > > > 2 > 3
Cellular contraction 1 > > > > >
Cellular response to stress 2 > > > > >
Regulated cell death 3 > > > > >
Iron, heme and hemoglobin homeostasis > > > 4 > >
MSN01 MSN02 MSN05 MSN06 MSN08 MSN09
Cell cycle and cell division 1 > 1 1 1 4
Lipid metabolism > 2 7 2 2 1
ECM homeostasis > 1 2 > 3 2
Cellular response to stress > > 6 4 4 5
Cytoskeleton dynamics > > 4 3 7 6
Cellular communication > 3 3 > 5 >
Energy generation and metabolism of cellular monomers > > > 5 6 >
Coag., fib., compl. system and blood protein dynamics > 4 8 > > >
Cellular adhesion 2 > > > > >
Cellular contraction > > > > > 3
Amino acid metabolism > > 5 > > >
decomposed
MSN01 MSN02 MSN05 MSN06 MSN08 MSN09
ECM homeostasis 2 3 3 1 4 1
Cellular response to stress 3 1 2 5 1 3
Cellular communication 4 2 5 3 5 4
Cellular contraction 1 > 1 2 3 2
Regulated cell death 5 > 4 4 2 5
Coag., fib., compl. system and blood protein dynamics 6 4 6 > 6 6
MSN01 MSN02 MSN05 MSN06 MSN08 MSN09
Amino acid metabolism 2 3 3 3 3 2
ECM homeostasis 3 2 4 2 2 4
Cell cycle and cell division 1 > 1 1 1 1
Lipid metabolism > 1 2 4 4 3
Cellular communication 5 4 > 5 5 >
Cellular response to stress 4 > 5 > > 5

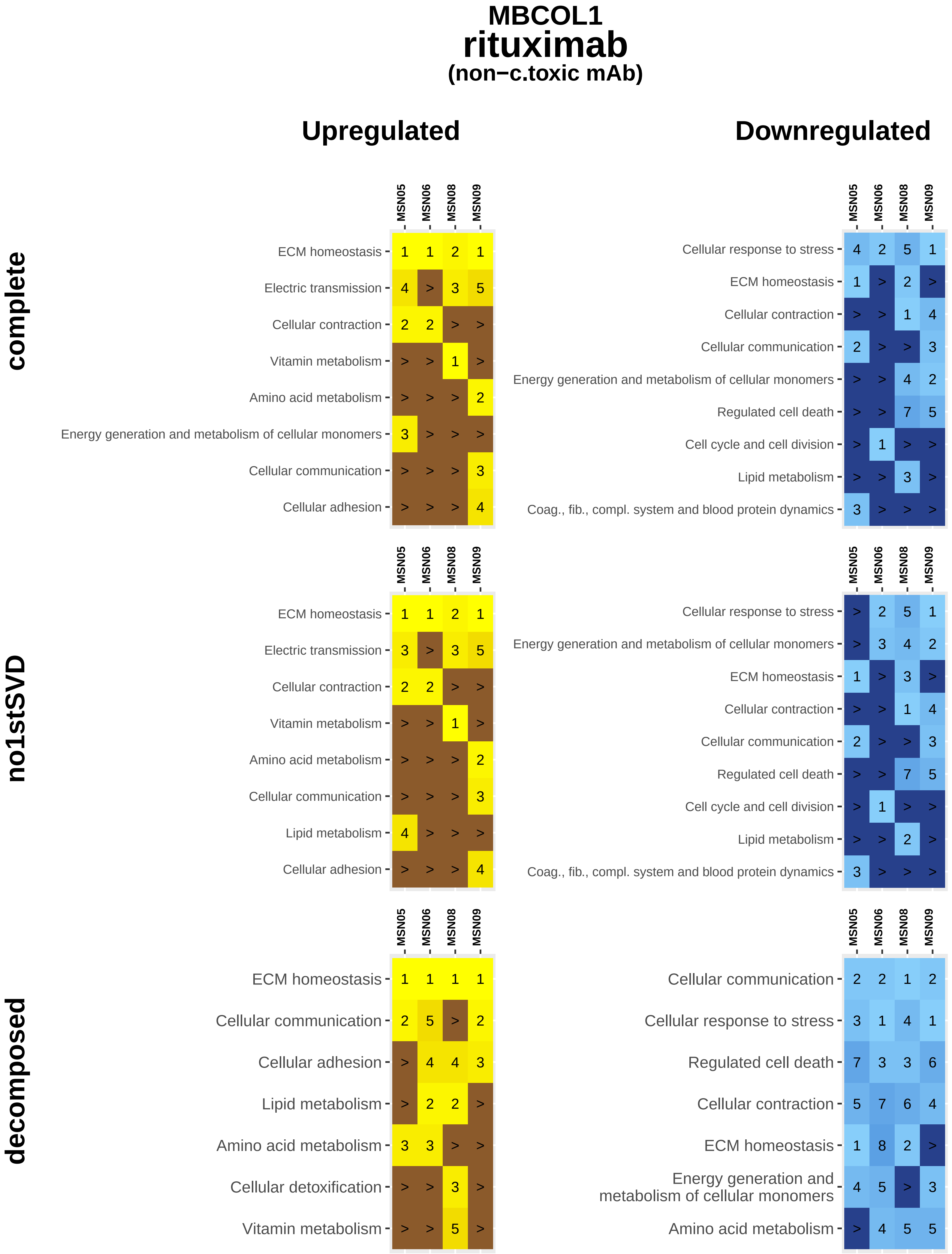
MBCOL1
rituximab
(non−c.toxic mAb)
Upregulated
Downregulated
complete
no1stSVD
decomposed
MSN05
MSN06
MSN08
MSN09
ECM homeostasis 1 1 2 1
Electric transmission 4 > 3 5
Cellular contraction 2 2 > >
Vitamin metabolism > > 1 >
Amino acid metabolism > > > 2
Energy generation and metabolism of cellular monomers 3 > > >
Cellular communication > > > 3
Cellular adhesion > > > 4
Cellular response to stress 4 2 5 1
ECM homeostasis 1 > 2 >
Cellular contraction > > 1 4
Cellular communication 2 > > 3
Energy generation and metabolism of cellular monomers > > 4 2
Regulated cell death > > 7 5
Cell cycle and cell division > 1 > >
Lipid metabolism > > 3 >
Coag., fib., compl. system and blood protein dynamics 3 > > >
ECM homeostasis 1 1 2 1
Electric transmission 3 > 3 5
Cellular contraction 2 2 > >
Vitamin metabolism > > 1 >
Amino acid metabolism > > > 2
Cellular communication > > > 3
Lipid metabolism 4 > > >
Cellular adhesion > > > 4
Cellular response to stress > 2 5 1
Energy generation and metabolism of cellular monomers > 3 4 2
ECM homeostasis 1 > 3 >
Cellular contraction > > 1 4
Cellular communication 2 > > 3
Regulated cell death > > 7 5
Cell cycle and cell division > 1 > >
Lipid metabolism > > 2 >
Coag., fib., compl. system and blood protein dynamics 3 > > >
ECM homeostasis 1 1 1 1
Cellular communication 2 5 > 2
Cellular adhesion > 4 4 3
Lipid metabolism > 2 2 >
Amino acid metabolism 3 3 > >
Cellular detoxification > > 3 >
Vitamin metabolism > > 5 >
Cellular communication 2 2 1 2
Cellular response to stress 3 1 4 1
Regulated cell death 7 3 3 6
Cellular contraction 5 7 6 4
ECM homeostasis 1 8 2 >
Energy generation and metabolism of cellular monomers 4 5 > 3
Amino acid metabolism > 4 5 5

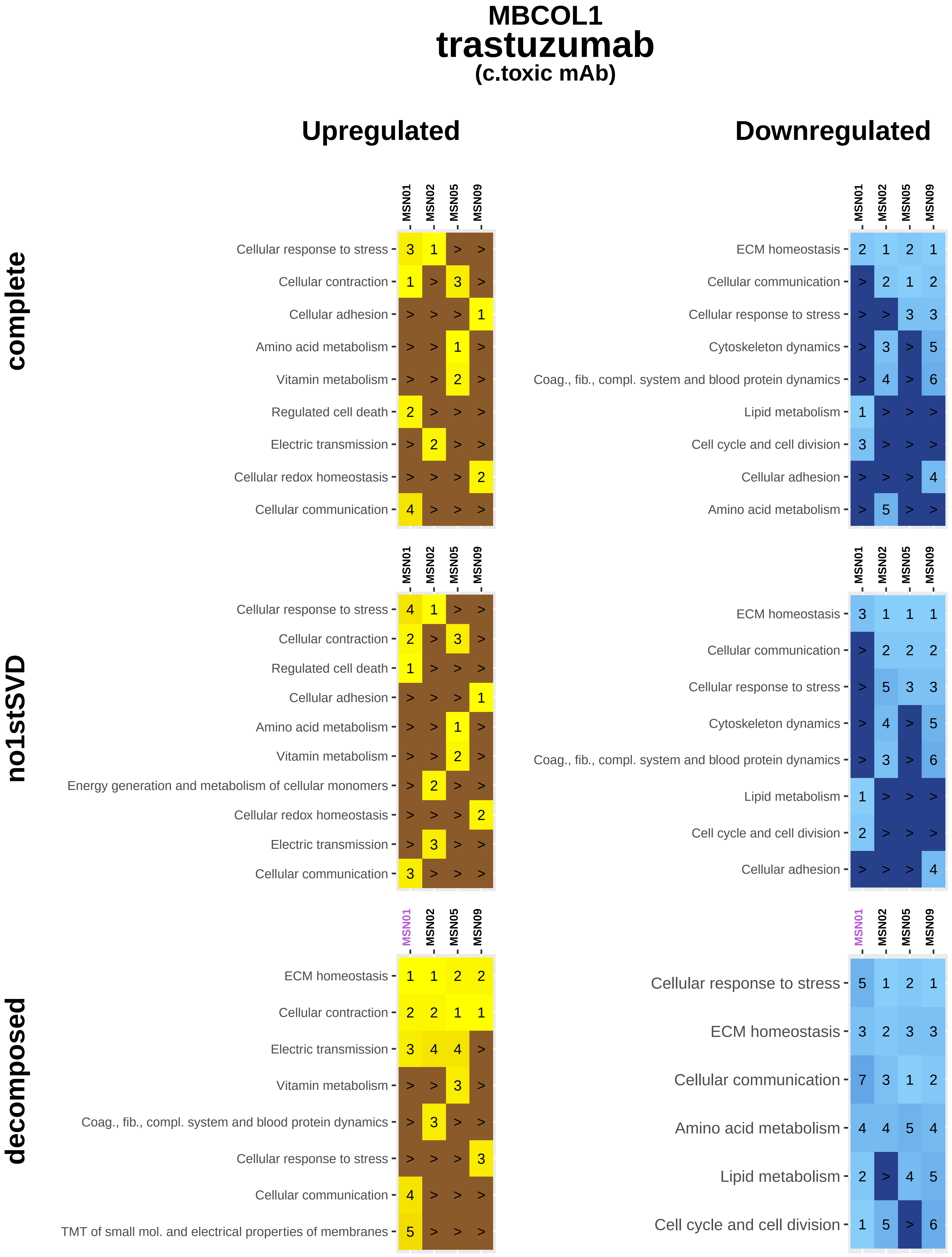
MBCOL1
trastuzumab
(c.toxic mAb)
Upregulated
Downregulated
complete
no1stSVD
decomposed
MSN01
MSN02
MSN05
MSN09
Cellular response to stress
Cellular contraction
Cellular adhesion
Amino acid metabolism
Vitamin metabolism
Regulated cell death
Electric transmission
Cellular redox homeostasis
Cellular communication
ECM homeostasis
Cytoskeleton dynamics
Coag., fib., compl. system and blood protein dynamics
Lipid metabolism
Cell cycle and cell division
Energy generation and metabolism of cellular monomers
TMT of small mol. and electrical properties of membranes

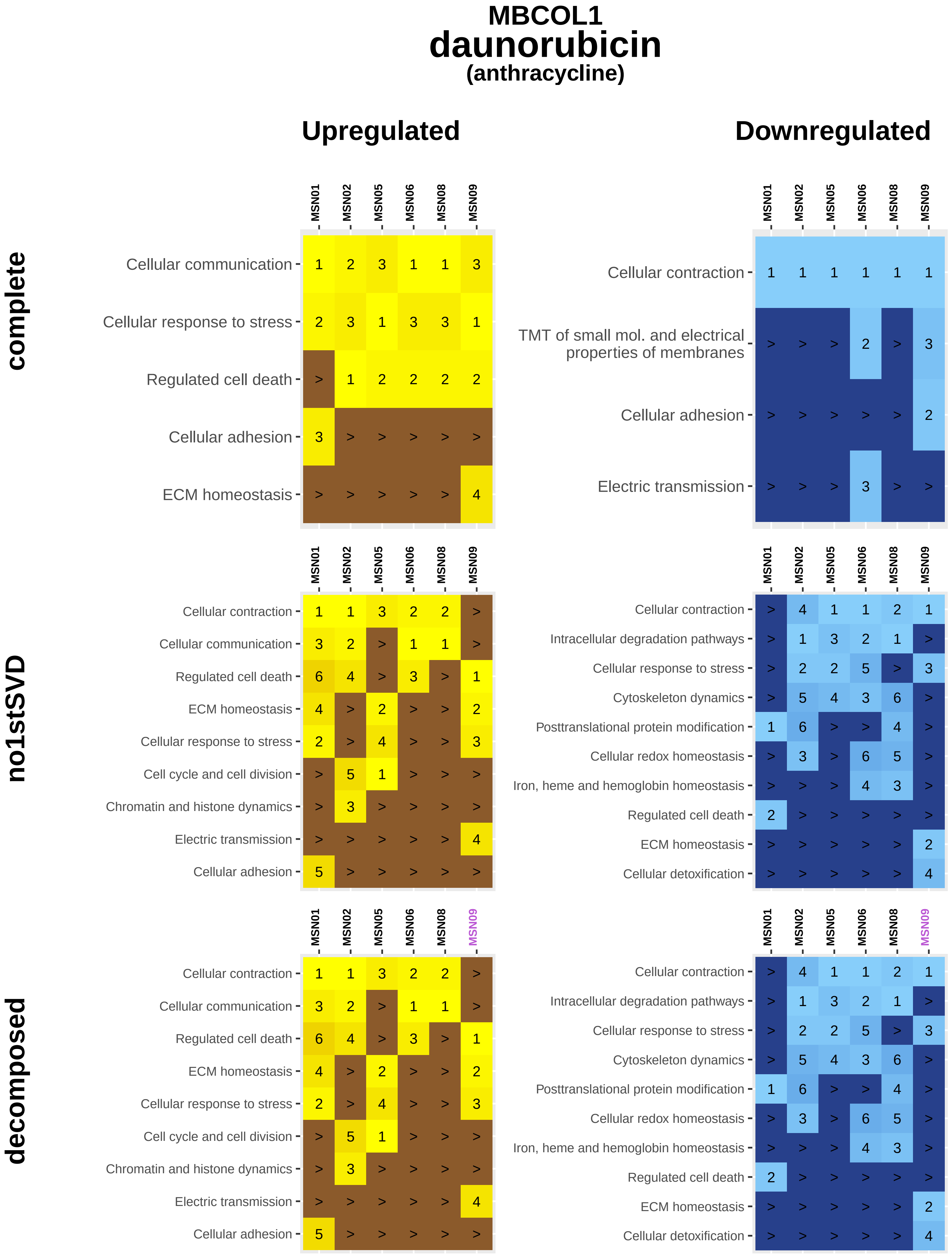
MBCOL1
daunorubicin
(anthracycline)
Upregulated
Downregulated
complete
no1stSVD
decomposed
MSN01
MSN02
MSN05
MSN06
MSN08
MSN09
Cellular communication
Cellular response to stress
Regulated cell death
Cellular adhesion
ECM homeostasis
Cellular contraction
TMT of small mol. and electrical properties of membranes
Electric transmission
Cell cycle and cell division
Chromatin and histone dynamics
Intracellular degradation pathways
Cytoskeleton dynamics
Posttranslational protein modification
Cellular redox homeostasis
Iron, heme and hemoglobin homeostasis
Cellular detoxification

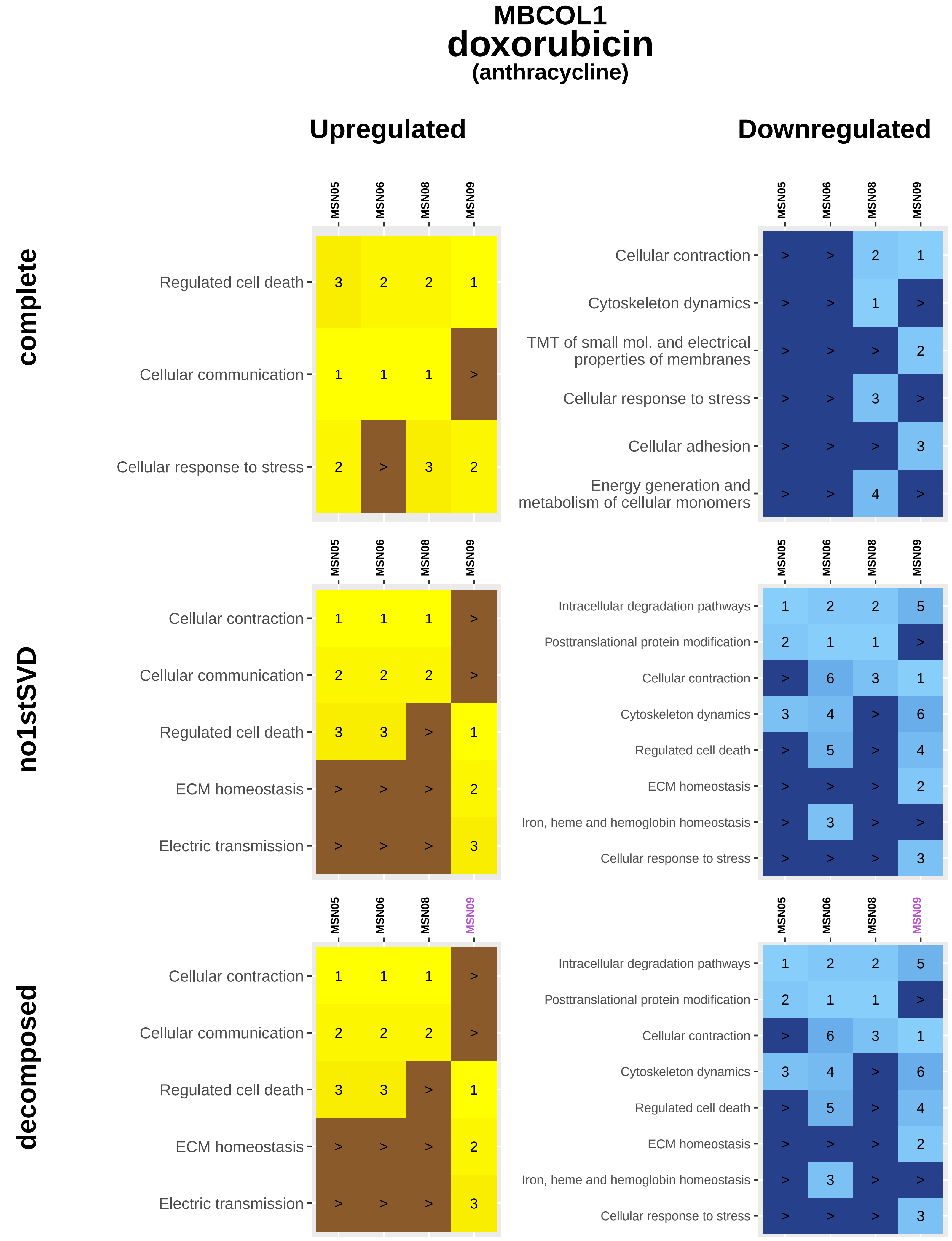

MBCOL1
doxorubicin
(anthracycline)
Upregulated
Downregulated
complete
no1stSVD
decomposed
MSN05
MSN06
MSN08
MSN09
Regulated cell death
Cellular communication
Cellular response to stress
Cellular contraction
Cytoskeleton dynamics
TMT of small mol. and electrical properties of membranes
Cellular adhesion
Energy generation and metabolism of cellular monomers
ECM homeostasis
Electric transmission
Intracellular degradation pathways
Posttranslational protein modification
Iron, heme and hemoglobin homeostasis

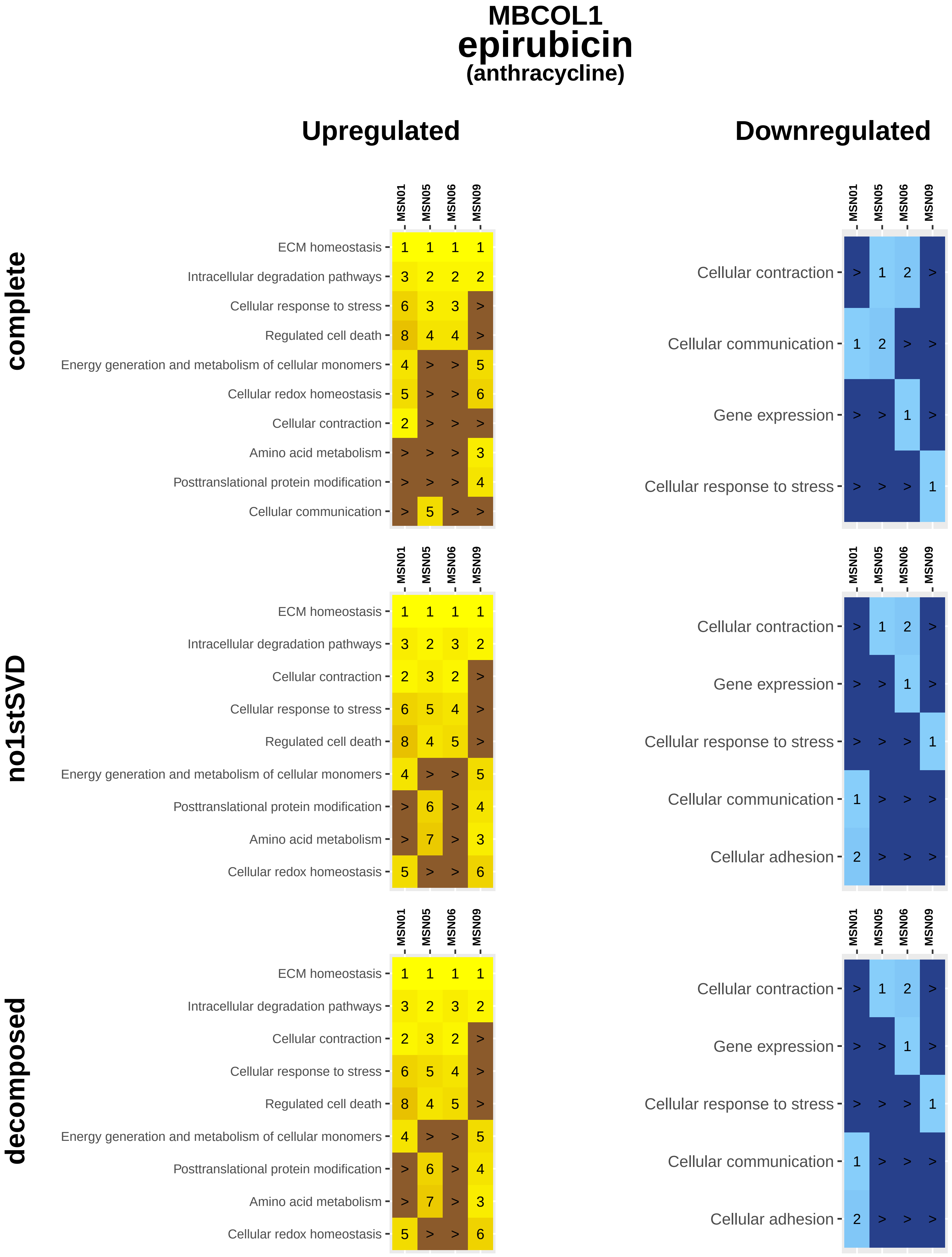
MBCOL1
epirubicin
(anthracycline)
Upregulated
Downregulated
complete
no1stSVD
decomposed
MSN01
MSN05
MSN06
MSN09
ECM homeostasis
Intracellular degradation pathways
Cellular response to stress
Regulated cell death
Energy generation and metabolism of cellular monomers
Cellular redox homeostasis
Cellular contraction
Amino acid metabolism
Posttranslational protein modification
Cellular communication
Gene expression
Cellular adhesion

### MBCOL1
### idarubicin
#### (anthracycline)

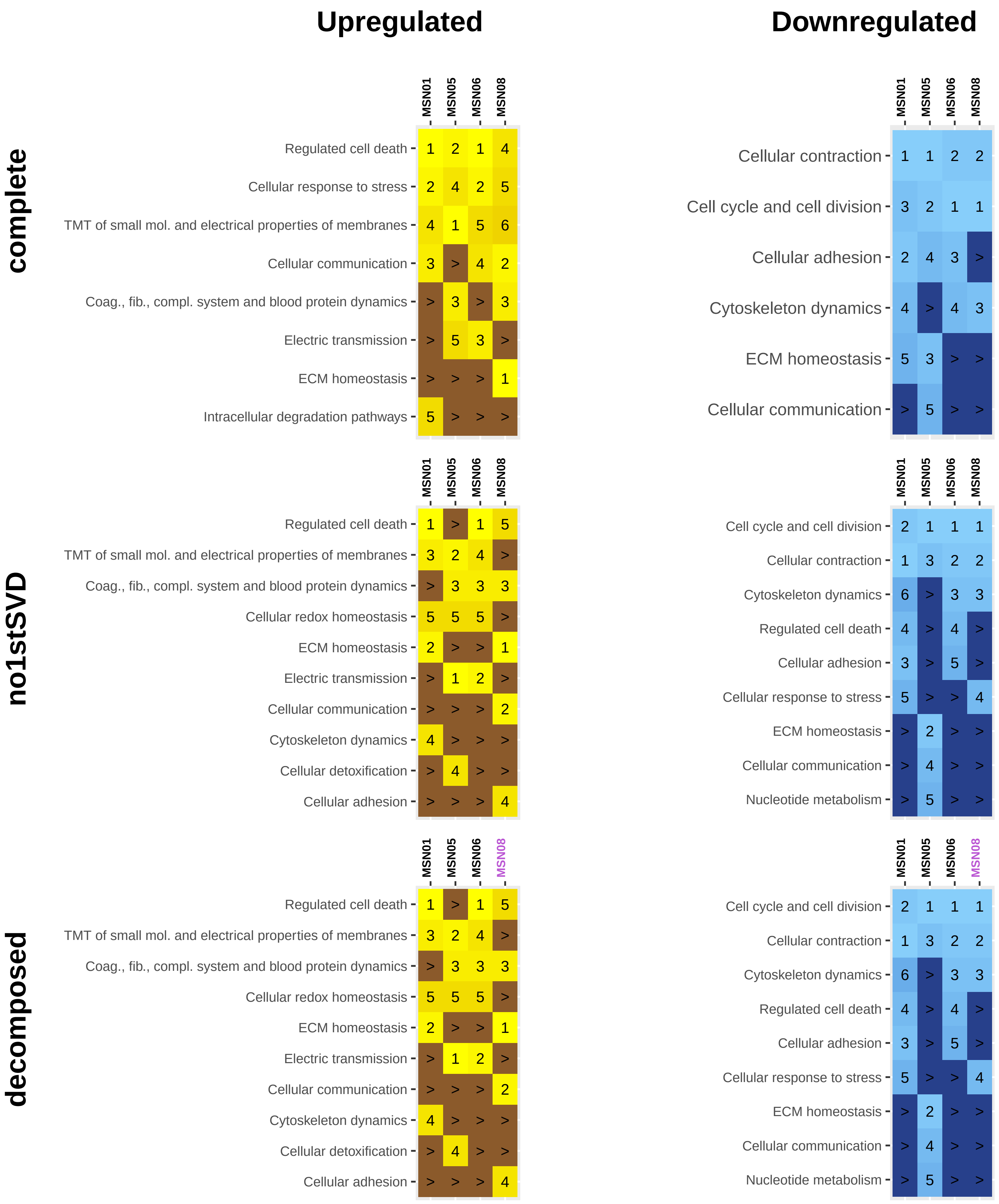

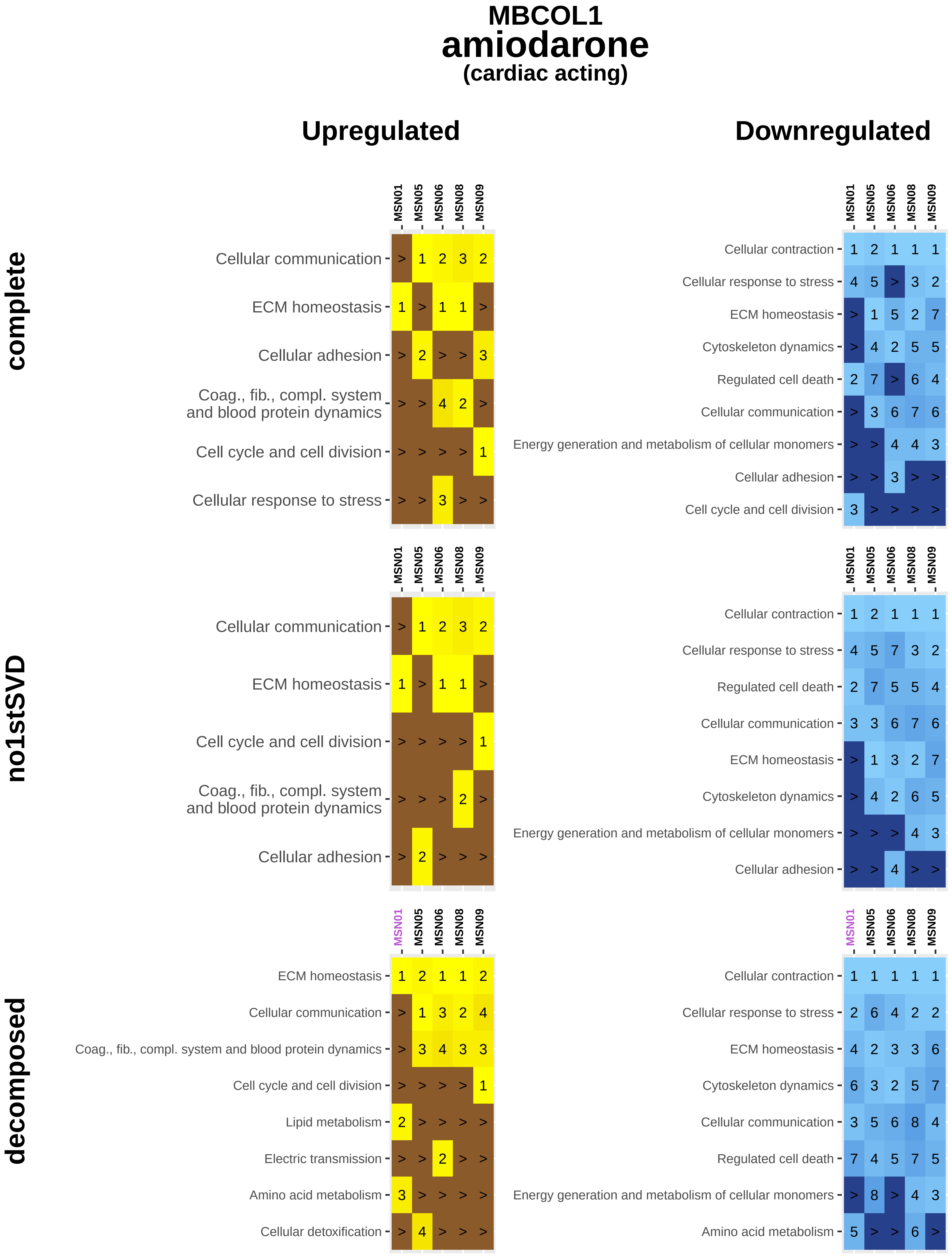
MBCOL1
amiodarone
(cardiac acting)
Upregulated
Downregulated
complete
no1stSVD
decomposed
MSN01
MSN05
MSN06
MSN08
MSN09
Cellular communication
ECM homeostasis
Cellular adhesion
Coag., fib., compl. system and blood protein dynamics
Cell cycle and cell division
Cellular response to stress
Cellular contraction
Cytoskeleton dynamics
Regulated cell death
Energy generation and metabolism of cellular monomers
Lipid metabolism
Electric transmission
Amino acid metabolism
Cellular detoxification

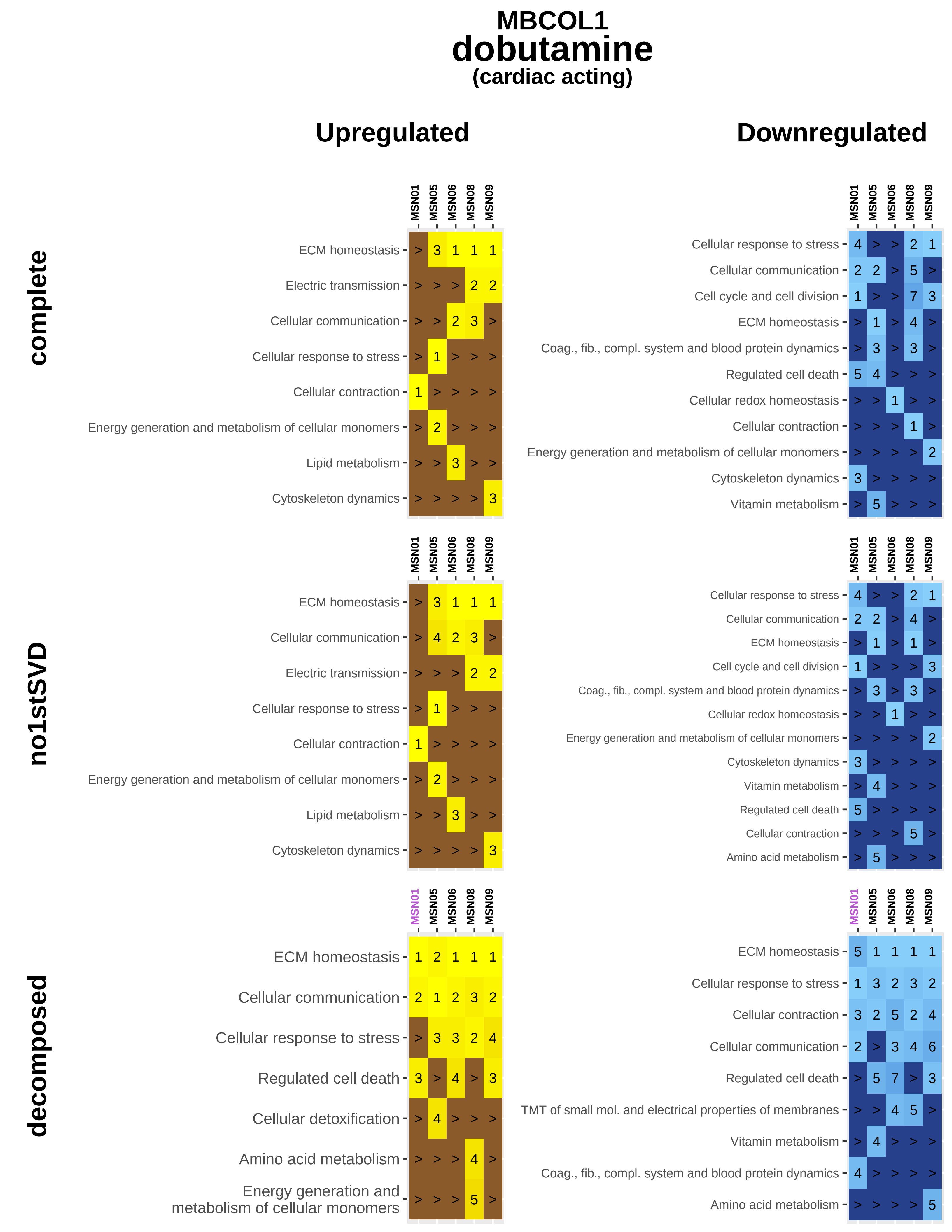

MBCOL1
dobutamine
(cardiac acting)
Upregulated
Downregulated
complete
no1stSVD
decomposed
MSN01
MSN05
MSN06
MSN08
MSN09
ECM homeostasis
Electric transmission
Cellular communication
Cellular response to stress
Cellular contraction
Energy generation and metabolism of cellular monomers
Lipid metabolism
Cytoskeleton dynamics
Cell cycle and cell division
Coag., fib., compl. system and blood protein dynamics
Regulated cell death
Cellular redox homeostasis
Vitamin metabolism
Amino acid metabolism
Cellular detoxification
TMT of small mol. and electrical properties of membranes

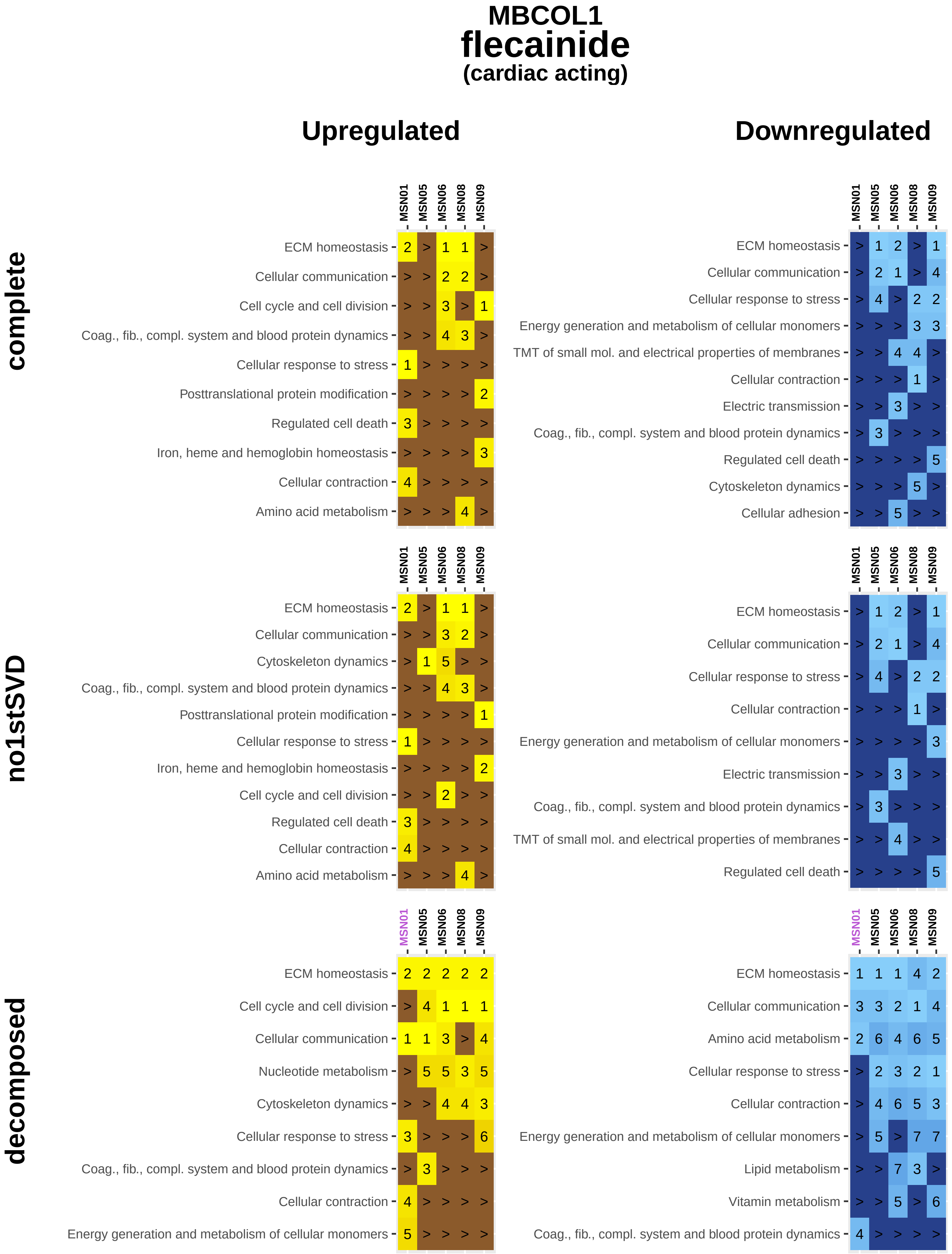
MBCOL1
flecainide
(cardiac acting)
Upregulated
Downregulated
complete
no1stSVD
decomposed
MSN01
MSN05
MSN06
MSN08
MSN09
ECM homeostasis
Cellular communication
Cell cycle and cell division
Coag., fib., compl. system and blood protein dynamics
Cellular response to stress
Posttranslational protein modification
Regulated cell death
Iron, heme and hemoglobin homeostasis
Cellular contraction
Amino acid metabolism
Energy generation and metabolism of cellular monomers
TMT of small mol. and electrical properties of membranes
Electric transmission
Cytoskeleton dynamics
Cellular adhesion
Nucleotide metabolism
Lipid metabolism
Vitamin metabolism

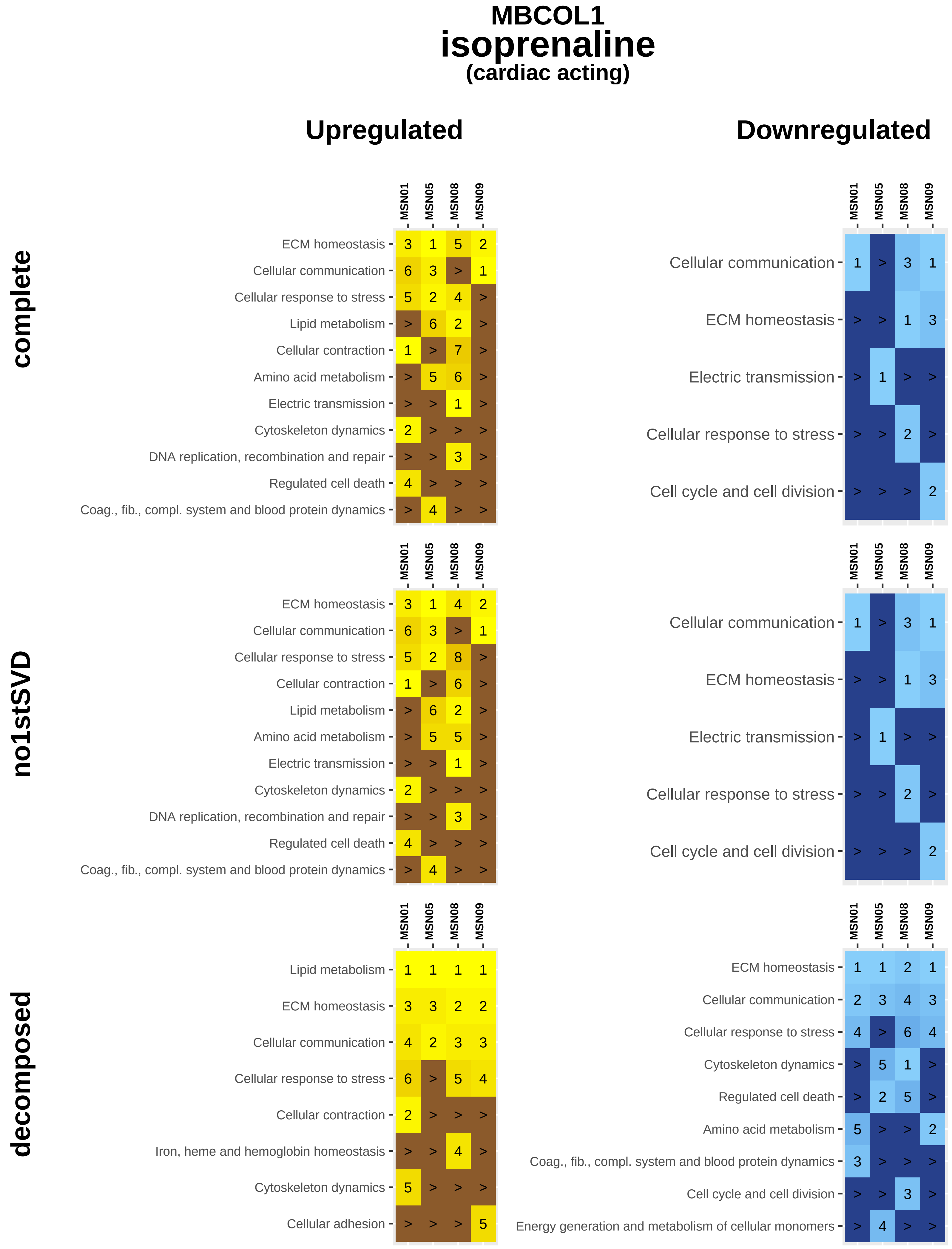
MBCOL1
isoprenaline
(cardiac acting)
Upregulated
Downregulated
complete
no1stSVD
decomposed
MSN01
MSN05
MSN08
MSN09
ECM homeostasis 3 1 5 2
Cellular communication 6 3 > 1
Cellular response to stress 5 2 4 >
Lipid metabolism > 6 2 >
Cellular contraction 1 > 7 >
Amino acid metabolism > 5 6 >
Electric transmission > > 1 >
Cytoskeleton dynamics 2 > > >
DNA replication, recombination and repair > > 3 >
Regulated cell death 4 > > >
Coag., fib., compl. system and blood protein dynamics > 4 > >
Cellular communication 1 > 3 1
ECM homeostasis > > 1 3
Electric transmission > 1 > >
Cellular response to stress > > 2 >
Cell cycle and cell division > > > 2
ECM homeostasis 3 1 4 2
Cellular communication 6 3 > 1
Cellular response to stress 5 2 8 >
Cellular contraction 1 > 6 >
Lipid metabolism > 6 2 >
Amino acid metabolism > 5 5 >
Electric transmission > > 1 >
Cytoskeleton dynamics 2 > > >
DNA replication, recombination and repair > > 3 >
Regulated cell death 4 > > >
Coag., fib., compl. system and blood protein dynamics > 4 > >
Cellular communication 1 > 3 1
ECM homeostasis > > 1 3
Electric transmission > 1 > >
Cellular response to stress > > 2 >
Cell cycle and cell division > > > 2
Lipid metabolism 1 1 1 1
ECM homeostasis 3 3 2 2
Cellular communication 4 2 3 3
Cellular response to stress 6 > 5 4
Cellular contraction 2 > > >
Iron, heme and hemoglobin homeostasis > > 4 >
Cytoskeleton dynamics 5 > > >
Cellular adhesion > > > 5
ECM homeostasis 1 1 2 1
Cellular communication 2 3 4 3
Cellular response to stress 4 > 6 4
Cytoskeleton dynamics > 5 1 >
Regulated cell death > 2 5 >
Amino acid metabolism 5 > > 2
Coag., fib., compl. system and blood protein dynamics 3 > > >
Cell cycle and cell division > > 3 >
Energy generation and metabolism of cellular monomers > 4 > >

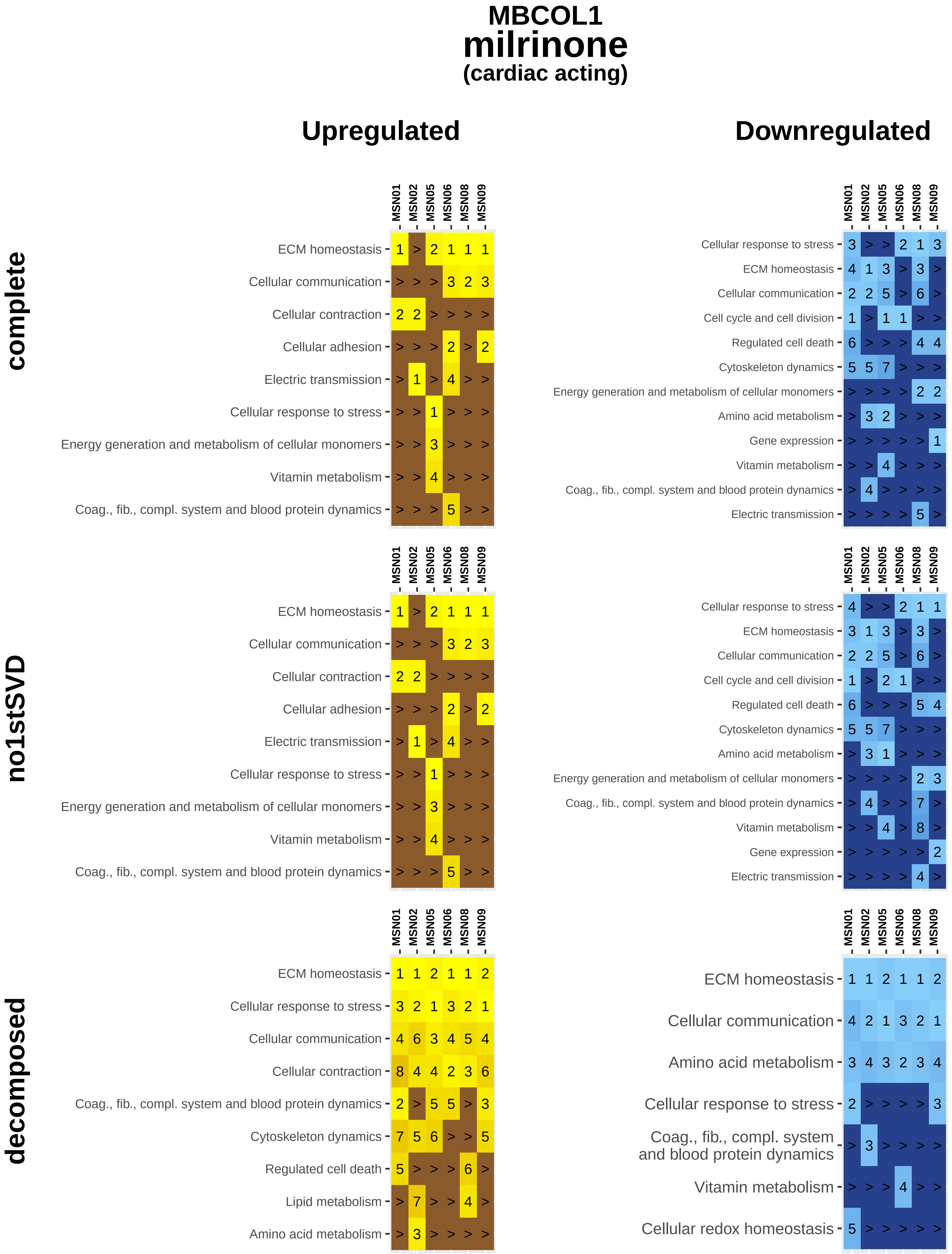
MBCOL1
milrinone
(cardiac acting)
Upregulated
Downregulated
complete
no1stSVD
decomposed
MSN01
MSN02
MSN05
MSN06
MSN08
MSN09
ECM homeostasis
Cellular communication
Cellular contraction
Cellular adhesion
Electric transmission
Cellular response to stress
Energy generation and metabolism of cellular monomers
Vitamin metabolism
Coag., fib., compl. system and blood protein dynamics
Cell cycle and cell division
Regulated cell death
Cytoskeleton dynamics
Amino acid metabolism
Gene expression
Lipid metabolism
Cellular redox homeostasis

### MBCOL1
### phenylephrine
#### (cardiac acting)

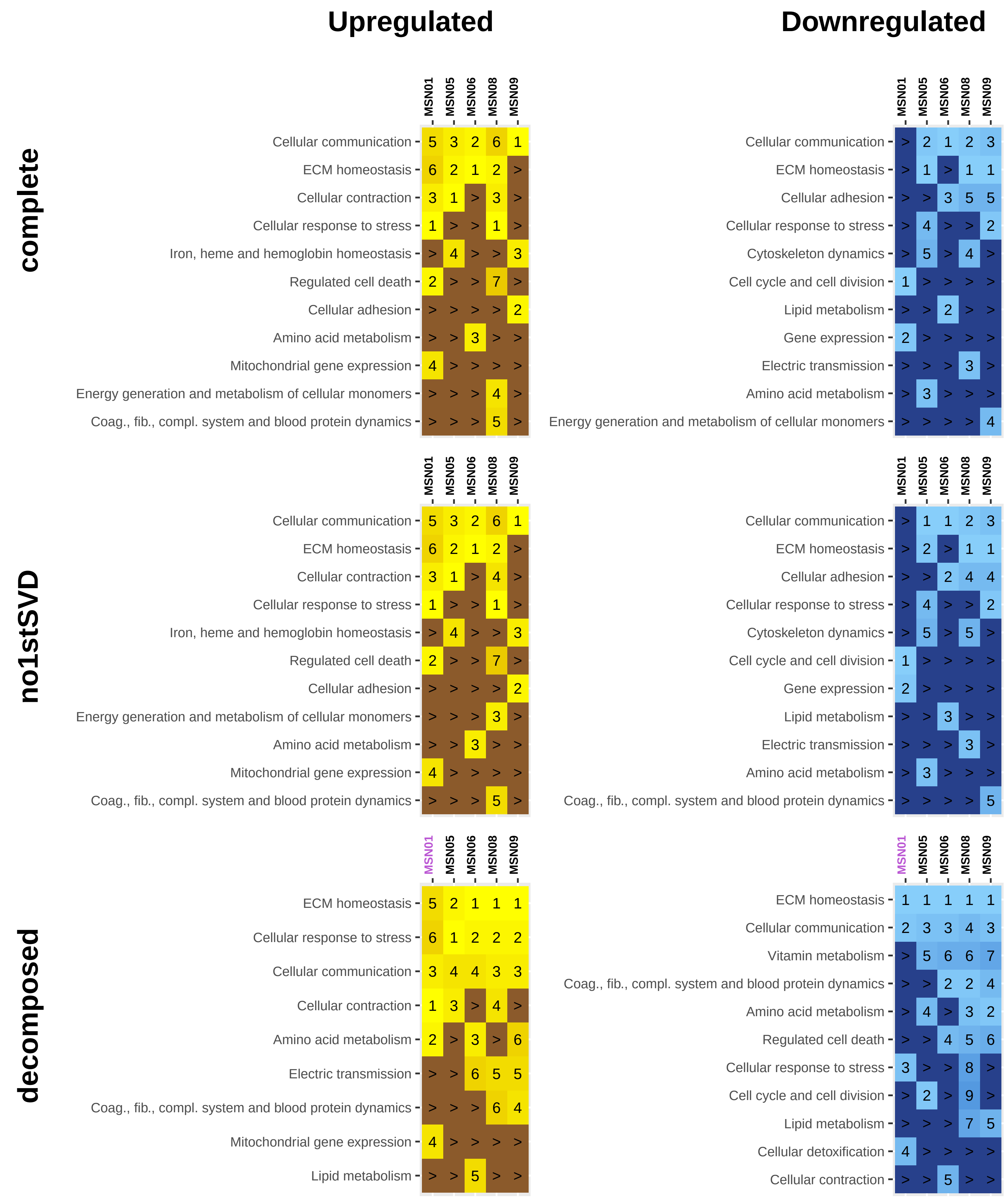

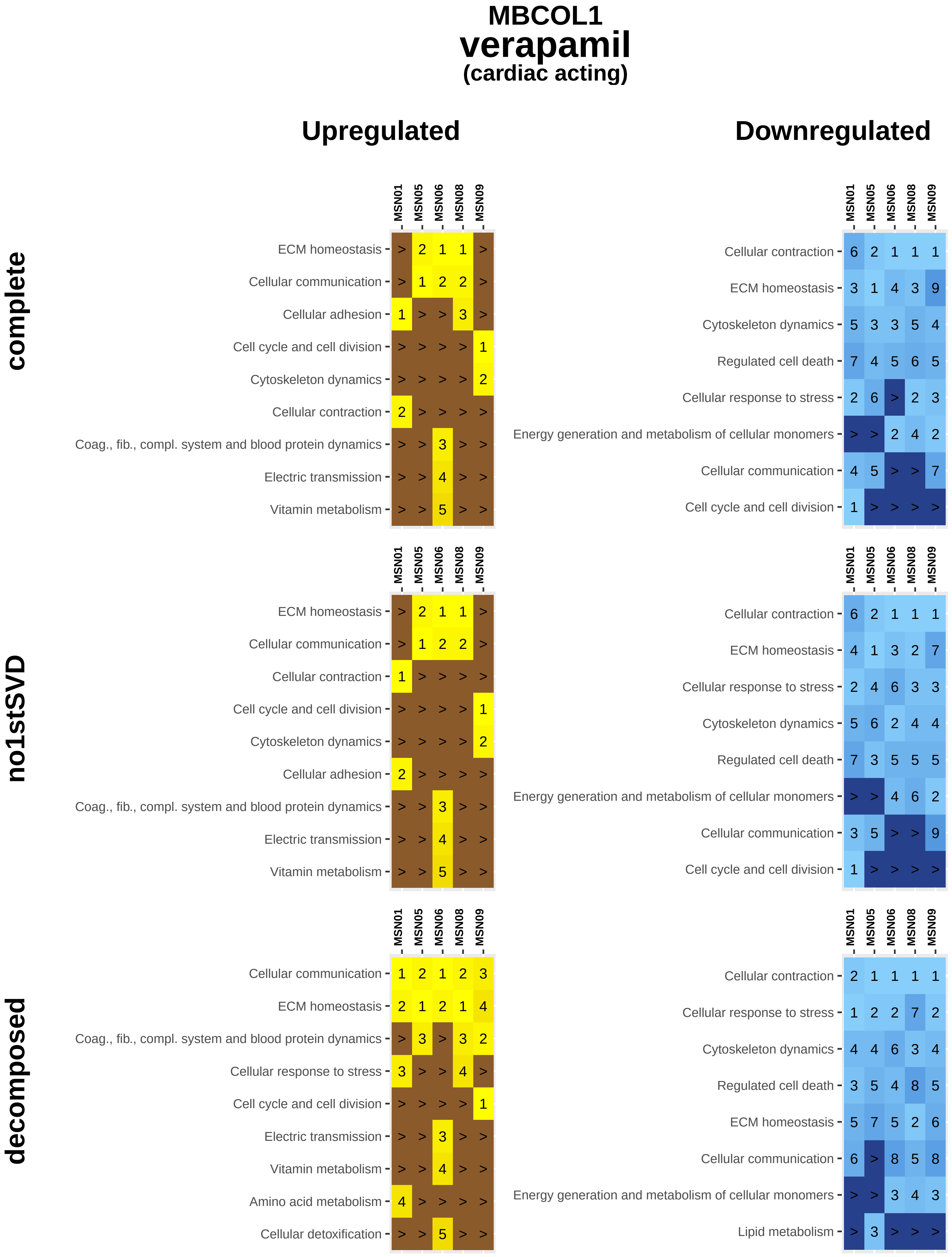
MBCOL1
verapamil
(cardiac acting)
Upregulated
Downregulated
complete
no1stSVD
decomposed
MSN01
MSN05
MSN06
MSN08
MSN09
ECM homeostasis
Cellular communication
Cellular adhesion
Cell cycle and cell division
Cytoskeleton dynamics
Cellular contraction
Coag., fib., compl. system and blood protein dynamics
Electric transmission
Vitamin metabolism
Regulated cell death
Cellular response to stress
Energy generation and metabolism of cellular monomers
Amino acid metabolism
Cellular detoxification
Lipid metabolism

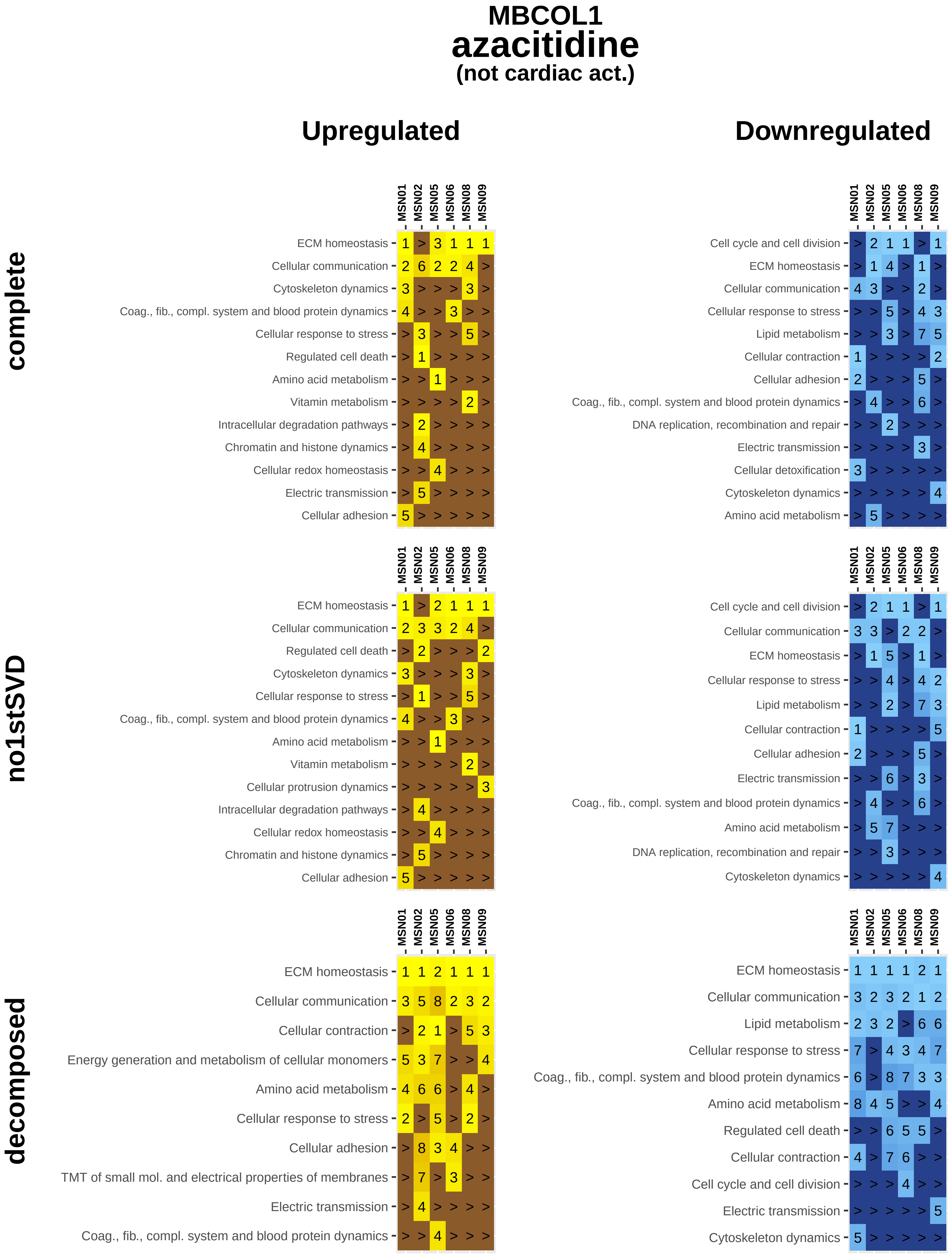
MBCOL1
azacitidine
(not cardiac act.)
Upregulated
Downregulated
complete
no1stSVD
decomposed
MSN01
MSN02
MSN05
MSN06
MSN08
MSN09
ECM homeostasis
Cellular communication
Cytoskeleton dynamics
Coag., fib., compl. system and blood protein dynamics
Cellular response to stress
Regulated cell death
Amino acid metabolism
Vitamin metabolism
Intracellular degradation pathways
Chromatin and histone dynamics
Cellular redox homeostasis
Electric transmission
Cellular adhesion
Cell cycle and cell division
Lipid metabolism
Cellular contraction
DNA replication, recombination and repair
Cellular detoxification
Cellular protrusion dynamics
Energy generation and metabolism of cellular monomers
TMT of small mol. and electrical properties of membranes

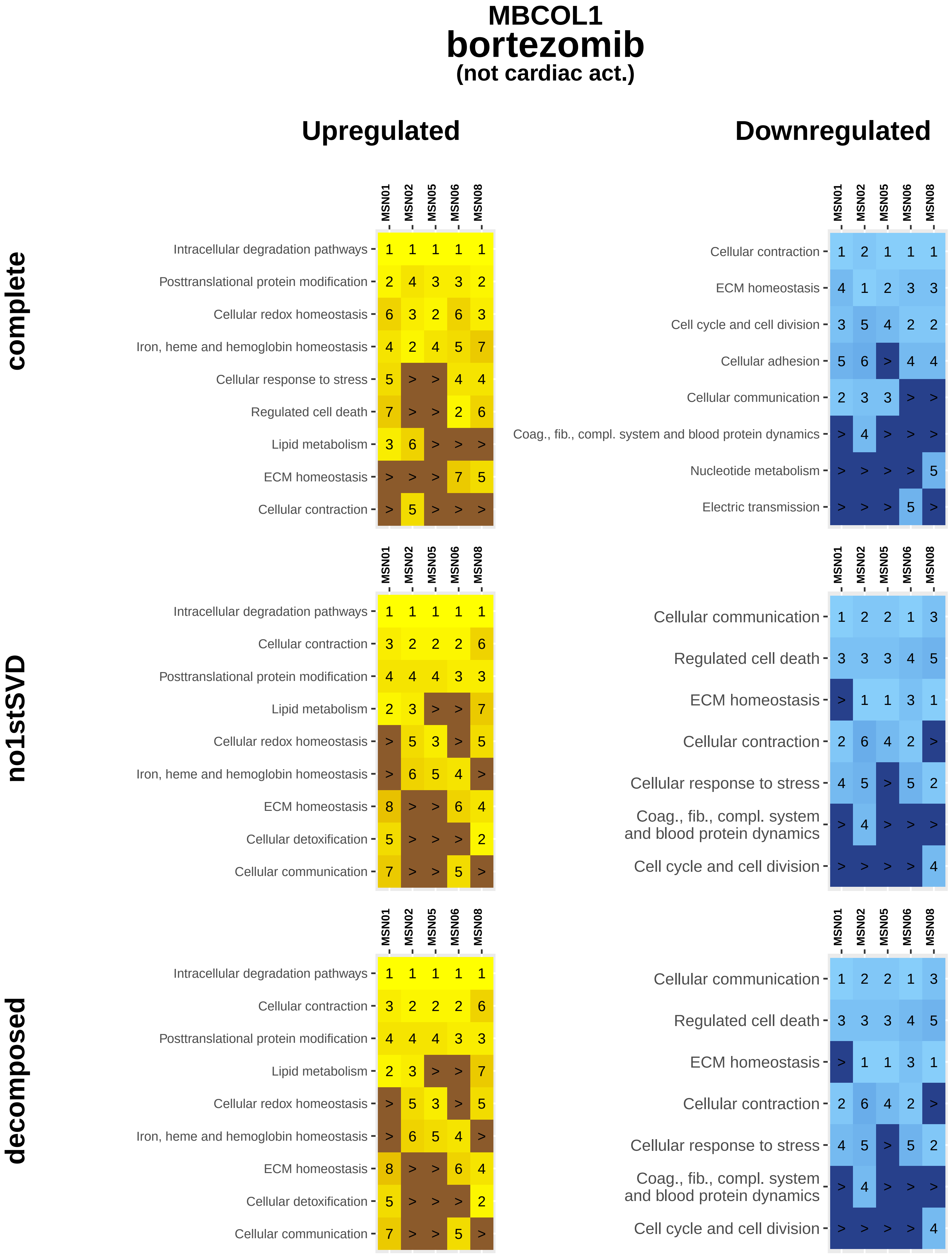

MBCOL1
bortezomib
(not cardiac act.)
Upregulated
Downregulated
complete
no1stSVD
decomposed
MSN01 MSN02 MSN05 MSN06 MSN08
Intracellular degradation pathways 1 1 1 1 1
Posttranslational protein modification 2 4 3 3 2
Cellular redox homeostasis 6 3 2 6 3
Iron, heme and hemoglobin homeostasis 4 2 4 5 7
Cellular response to stress 5 > > 4 4
Regulated cell death 7 > > 2 6
Lipid metabolism 3 6 > > >
ECM homeostasis > > > 7 5
Cellular contraction > 5 > > >
Cellular contraction 1 2 1 1 1
ECM homeostasis 4 1 2 3 3
Cell cycle and cell division 3 5 4 2 2
Cellular adhesion 5 6 > 4 4
Cellular communication 2 3 3 > >
Coag., fib., compl. system and blood protein dynamics > 4 > > >
Nucleotide metabolism > > > > 5
Electric transmission > > > 5 >
Intracellular degradation pathways 1 1 1 1 1
Cellular contraction 3 2 2 2 6
Posttranslational protein modification 4 4 4 3 3
Lipid metabolism 2 3 > > 7
Cellular redox homeostasis > 5 3 > 5
Iron, heme and hemoglobin homeostasis > 6 5 4 >
ECM homeostasis 8 > > 6 4
Cellular detoxification 5 > > > 2
Cellular communication 7 > > 5 >
Cellular communication 1 2 2 1 3
Regulated cell death 3 3 3 4 5
ECM homeostasis > 1 1 3 1
Cellular contraction 2 6 4 2 >
Cellular response to stress 4 5 > 5 2
Coag., fib., compl. system and blood protein dynamics > 4 > > >
Cell cycle and cell division > > > > 4
Intracellular degradation pathways 1 1 1 1 1
Cellular contraction 3 2 2 2 6
Posttranslational protein modification 4 4 4 3 3
Lipid metabolism 2 3 > > 7
Cellular redox homeostasis > 5 3 > 5
Iron, heme and hemoglobin homeostasis > 6 5 4 >
ECM homeostasis 8 > > 6 4
Cellular detoxification 5 > > > 2
Cellular communication 7 > > 5 >
Cellular communication 1 2 2 1 3
Regulated cell death 3 3 3 4 5
ECM homeostasis > 1 1 3 1
Cellular contraction 2 6 4 2 >
Cellular response to stress 4 5 > 5 2
Coag., fib., compl. system and blood protein dynamics > 4 > > >
Cell cycle and cell division > > > > 4

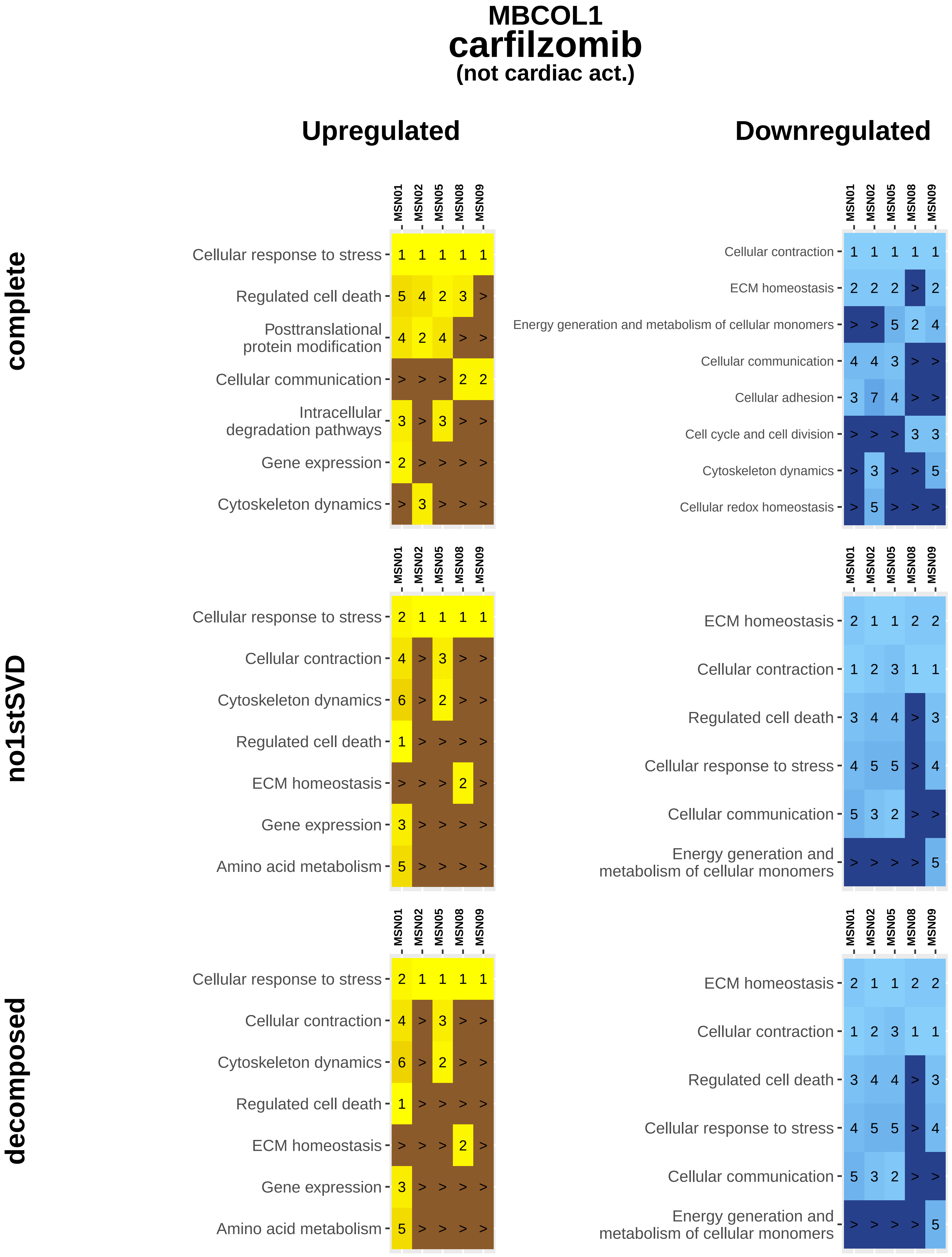
MBCOL1
carfilzomib
(not cardiac act.)
Upregulated
Downregulated
complete
no1stSVD
decomposed
MSN01
MSN02
MSN05
MSN08
MSN09
Cellular response to stress
Regulated cell death
Posttranslational protein modification
Cellular communication
Intracellular degradation pathways
Gene expression
Cytoskeleton dynamics
Cellular contraction
ECM homeostasis
Energy generation and metabolism of cellular monomers
Cellular adhesion
Cell cycle and cell division
Cellular redox homeostasis
Amino acid metabolism

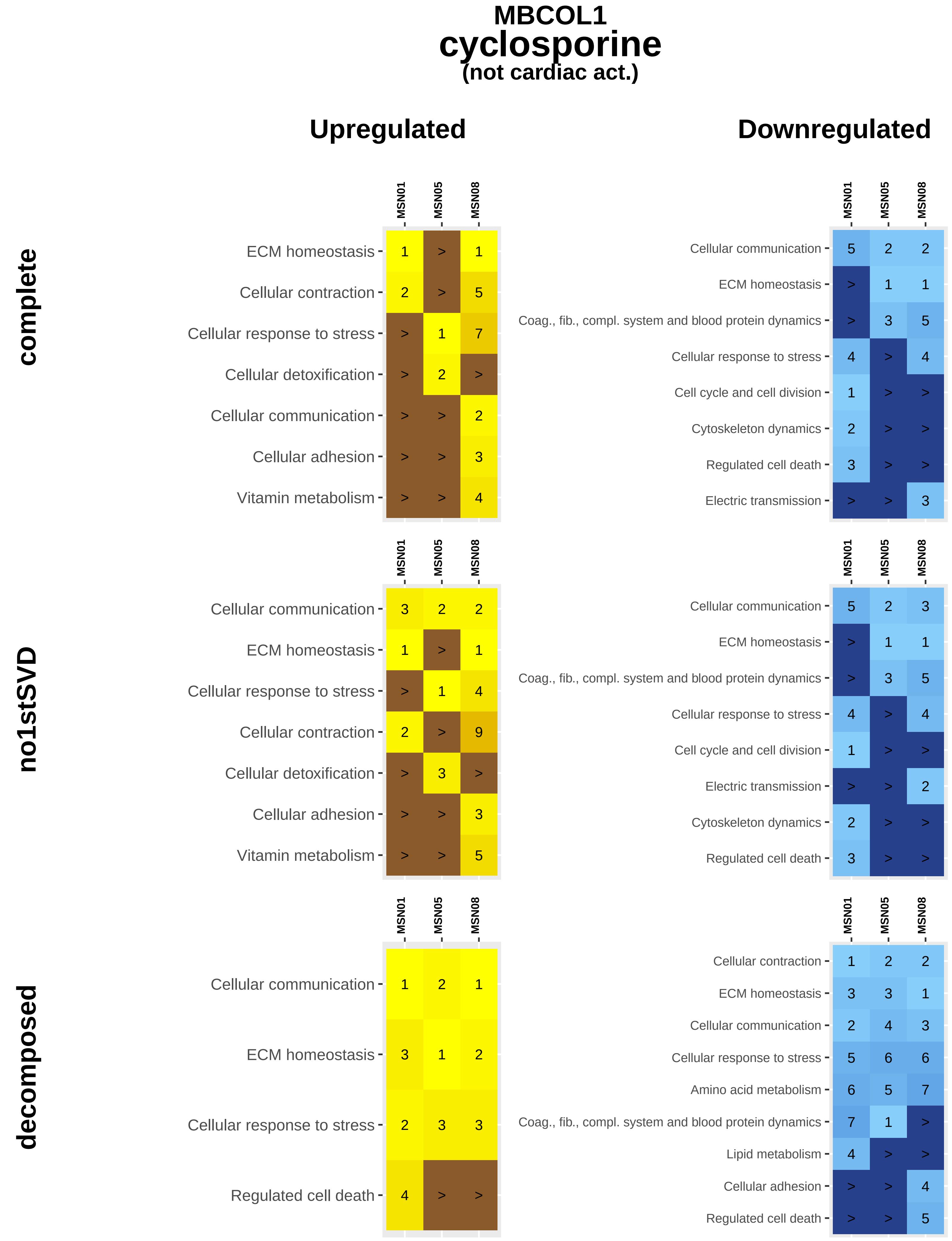
MBCOL1
cyclosporine
(not cardiac act.)
Upregulated
Downregulated
complete
no1stSVD
decomposed
MSN01
MSN05
MSN08
ECM homeostasis 1 > 1
Cellular contraction 2 > 5
Cellular response to stress > 1 7
Cellular detoxification > 2 >
Cellular communication > > 2
Cellular adhesion > > 3
Vitamin metabolism > > 4
Cellular communication 5 2 2
ECM homeostasis > 1 1
Coag., fib., compl. system and blood protein dynamics > 3 5
Cellular response to stress 4 > 4
Cell cycle and cell division 1 > >
Cytoskeleton dynamics 2 > >
Regulated cell death 3 > >
Electric transmission > > 3
Cellular communication 3 2 2
ECM homeostasis 1 > 1
Cellular response to stress > 1 4
Cellular contraction 2 > 9
Cellular detoxification > 3 >
Cellular adhesion > > 3
Vitamin metabolism > > 5
Cellular communication 5 2 3
ECM homeostasis > 1 1
Coag., fib., compl. system and blood protein dynamics > 3 5
Cellular response to stress 4 > 4
Cell cycle and cell division 1 > >
Electric transmission > > 2
Cytoskeleton dynamics 2 > >
Regulated cell death 3 > >
Cellular communication 1 2 1
ECM homeostasis 3 1 2
Cellular response to stress 2 3 3
Regulated cell death 4 > >
Cellular contraction 1 2 2
ECM homeostasis 3 3 1
Cellular communication 2 4 3
Cellular response to stress 5 6 6
Amino acid metabolism 6 5 7
Coag., fib., compl. system and blood protein dynamics 7 1 >
Lipid metabolism 4 > >
Cellular adhesion > > 4
Regulated cell death > > 5

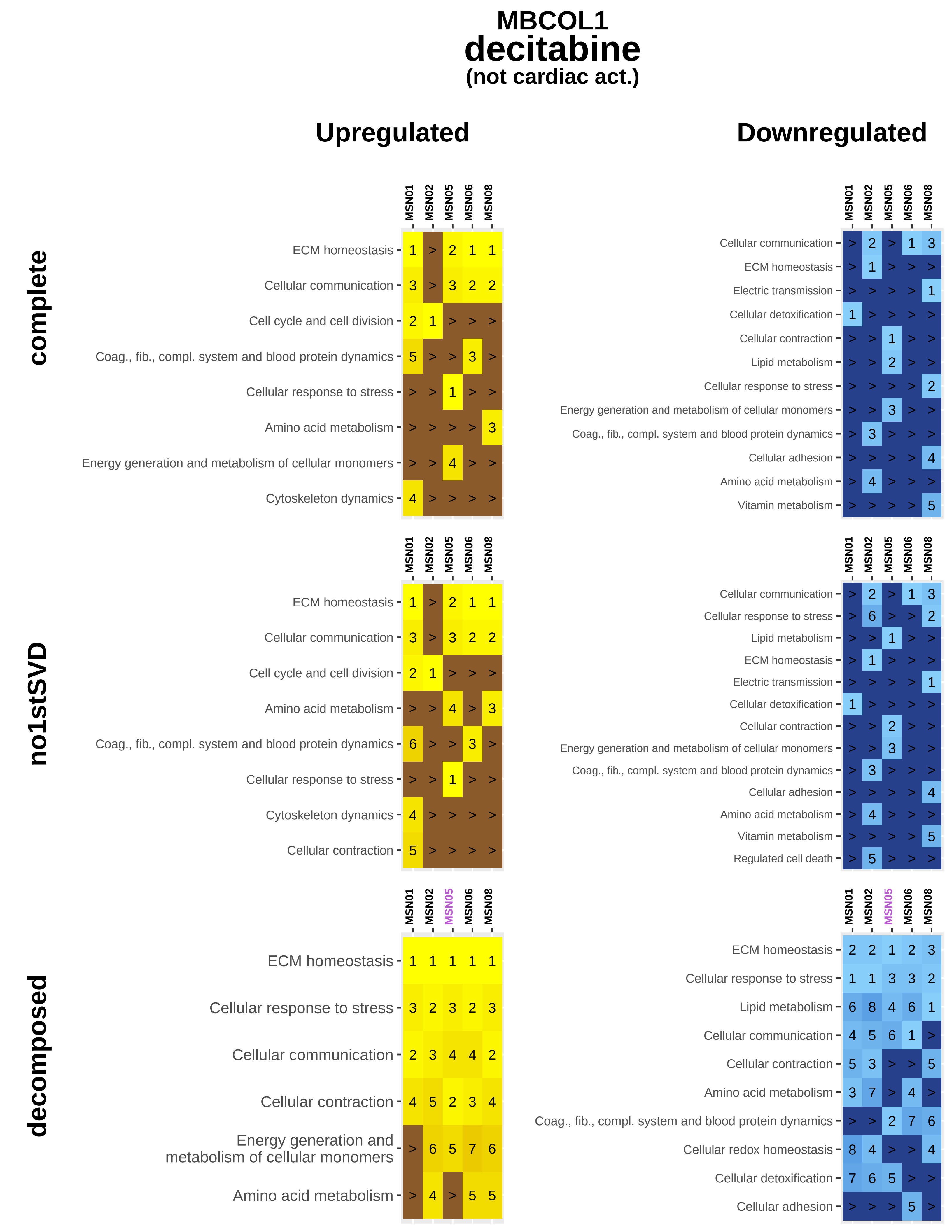

MBCOL1
decitabine
(not cardiac act.)
Upregulated
Downregulated
complete
no1stSVD
decomposed
MSN01
MSN02
MSN05
MSN06
MSN08
ECM homeostasis
Cellular communication
Cell cycle and cell division
Coag., fib., compl. system and blood protein dynamics
Cellular response to stress
Amino acid metabolism
Energy generation and metabolism of cellular monomers
Cytoskeleton dynamics
Electric transmission
Cellular detoxification
Cellular contraction
Lipid metabolism
Cellular adhesion
Vitamin metabolism
Regulated cell death
Cellular redox homeostasis

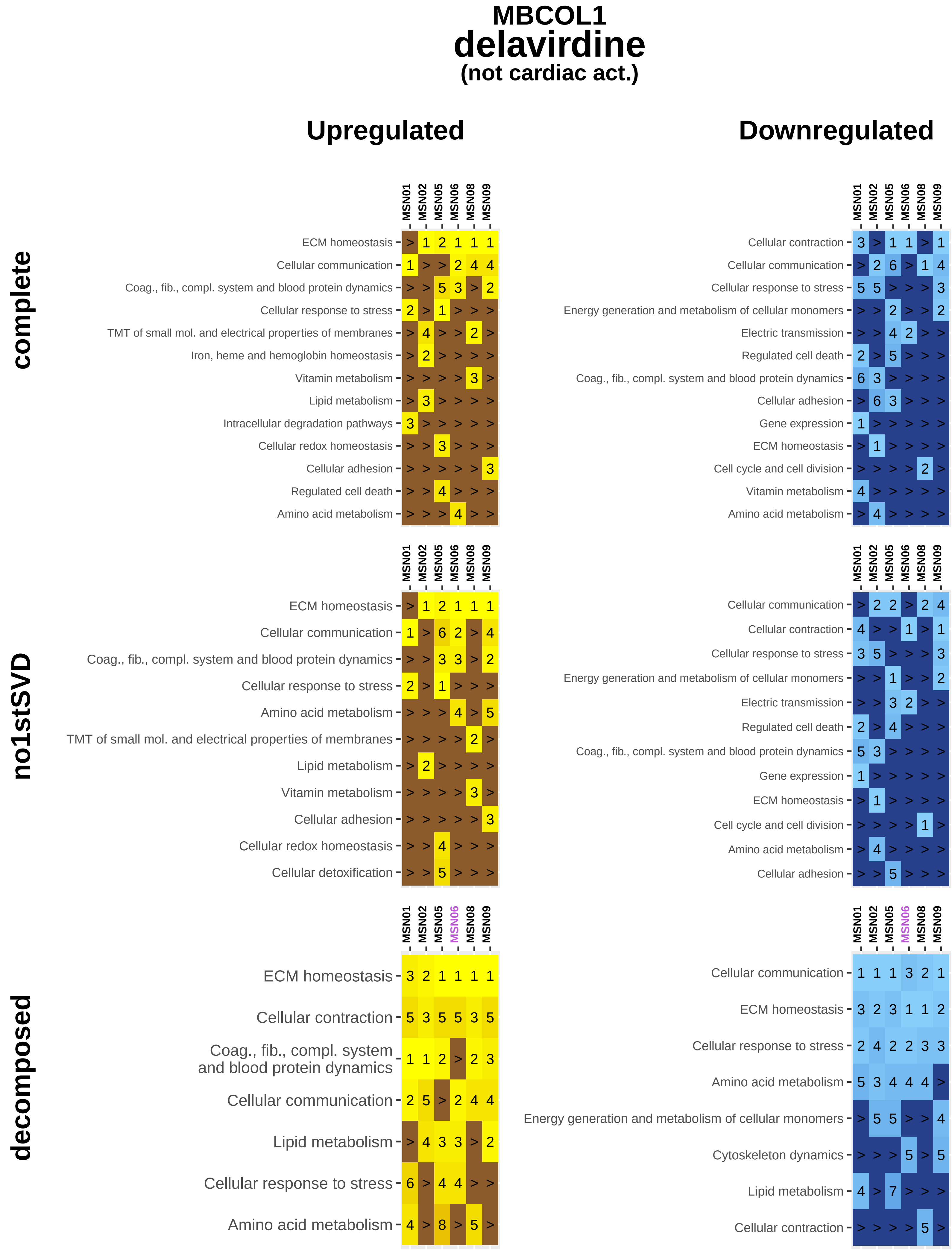
MBCOL1
delavirdine
(not cardiac act.)
Upregulated
Downregulated
complete
no1stSVD
decomposed
MSN01
MSN02
MSN05
MSN06
MSN08
MSN09
ECM homeostasis
Cellular communication
Coag., fib., compl. system and blood protein dynamics
Cellular response to stress
TMT of small mol. and electrical properties of membranes
Iron, heme and hemoglobin homeostasis
Vitamin metabolism
Lipid metabolism
Intracellular degradation pathways
Cellular redox homeostasis
Cellular adhesion
Regulated cell death
Amino acid metabolism
Cellular contraction
Energy generation and metabolism of cellular monomers
Electric transmission
Gene expression
Cell cycle and cell division
Cellular detoxification
Cytoskeleton dynamics

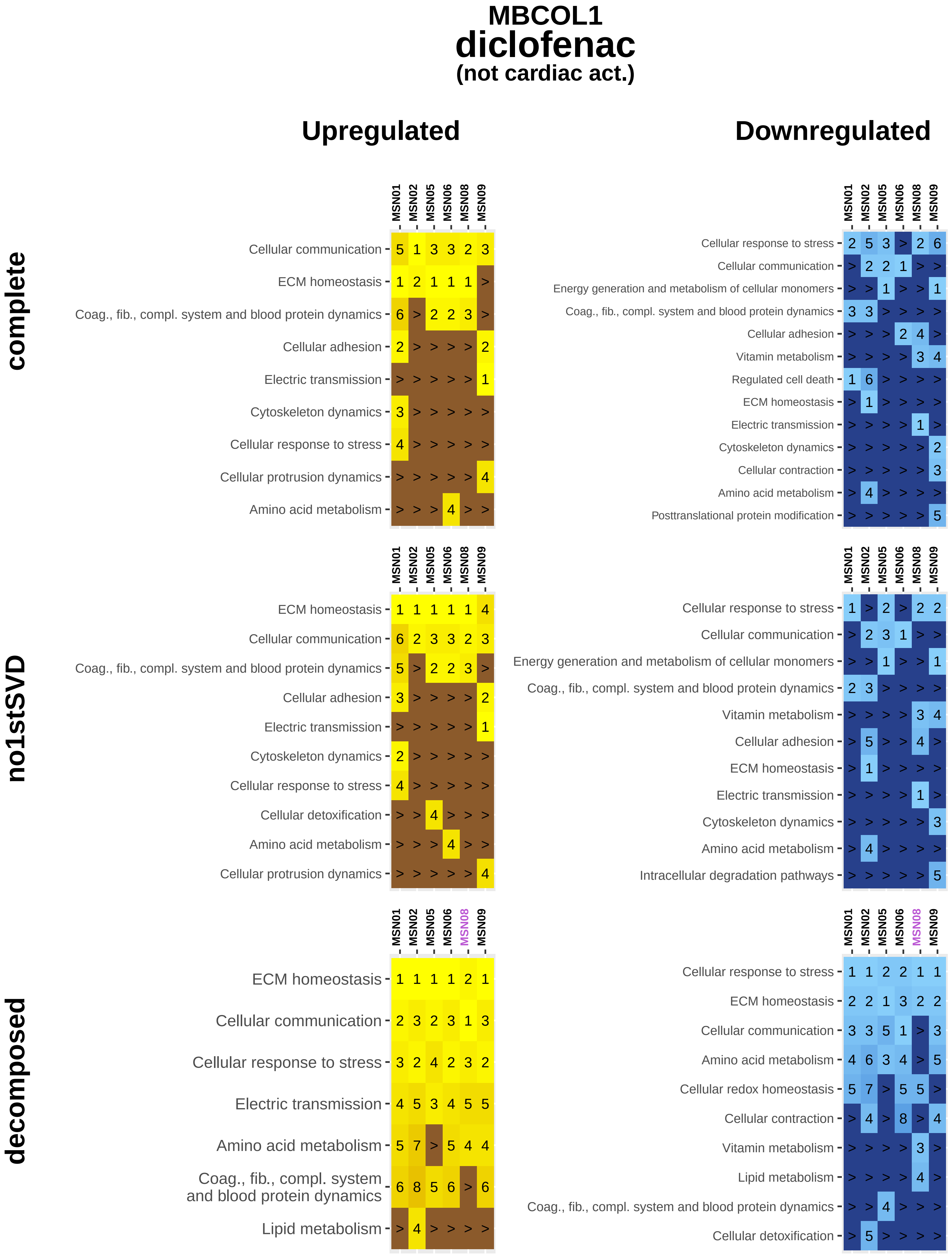
MBCOL1
diclofenac
(not cardiac act.)
Upregulated
Downregulated
complete
no1stSVD
decomposed
MSN01 MSN02 MSN05 MSN06 MSN08 MSN09
Cellular communication 5 1 3 3 2 3
ECM homeostasis 1 2 1 1 1 >
Coag., fib., compl. system and blood protein dynamics 6 > 2 2 3 >
Cellular adhesion 2 > > > > 2
Electric transmission > > > > > 1
Cytoskeleton dynamics 3 > > > > >
Cellular response to stress 4 > > > > >
Cellular protrusion dynamics > > > > > 4
Amino acid metabolism > > > 4 > >
Cellular response to stress 2 5 3 > 2 6
Cellular communication > 2 2 1 > >
Energy generation and metabolism of cellular monomers > > 1 > > 1
Coag., fib., compl. system and blood protein dynamics 3 3 > > > >
Cellular adhesion > > > 2 4 >
Vitamin metabolism > > > > 3 4
Regulated cell death 1 6 > > > >
ECM homeostasis > 1 > > > >
Electric transmission > > > > 1 >
Cytoskeleton dynamics > > > > > 2
Cellular contraction > > > > > 3
Amino acid metabolism > 4 > > > >
Posttranslational protein modification > > > > > 5
ECM homeostasis 1 1 1 1 1 4
Cellular communication 6 2 3 3 2 3
Coag., fib., compl. system and blood protein dynamics 5 > 2 2 3 >
Cellular adhesion 3 > > > > 2
Electric transmission > > > > > 1
Cytoskeleton dynamics 2 > > > > >
Cellular response to stress 4 > > > > >
Cellular detoxification > > 4 > > >
Amino acid metabolism > > > 4 > >
Cellular protrusion dynamics > > > > > 4
Cellular response to stress 1 > 2 > 2 2
Cellular communication > 2 3 1 > >
Energy generation and metabolism of cellular monomers > > 1 > > 1
Coag., fib., compl. system and blood protein dynamics 2 3 > > > >
Vitamin metabolism > > > > 3 4
Cellular adhesion > 5 > > 4 >
ECM homeostasis > 1 > > > >
Electric transmission > > > > 1 >
Cytoskeleton dynamics > > > > > 3
Amino acid metabolism > 4 > > > >
Intracellular degradation pathways > > > > > 5
ECM homeostasis 1 1 1 1 2 1
Cellular communication 2 3 2 3 1 3
Cellular response to stress 3 2 4 2 3 2
Electric transmission 4 5 3 4 5 5
Amino acid metabolism 5 7 > 5 4 4
Coag., fib., compl. system and blood protein dynamics 6 8 5 6 > 6
Lipid metabolism > 4 > > > >
Cellular response to stress 1 1 2 2 1 1
ECM homeostasis 2 2 1 3 2 2
Cellular communication 3 3 5 1 > 3
Amino acid metabolism 4 6 3 4 > 5
Cellular redox homeostasis 5 7 > 5 5 >
Cellular contraction > 4 > 8 > 4
Vitamin metabolism > > > > 3 >
Lipid metabolism > > > > 4 >
Coag., fib., compl. system and blood protein dynamics > > 4 > > >
Cellular detoxification > 5 > > > >

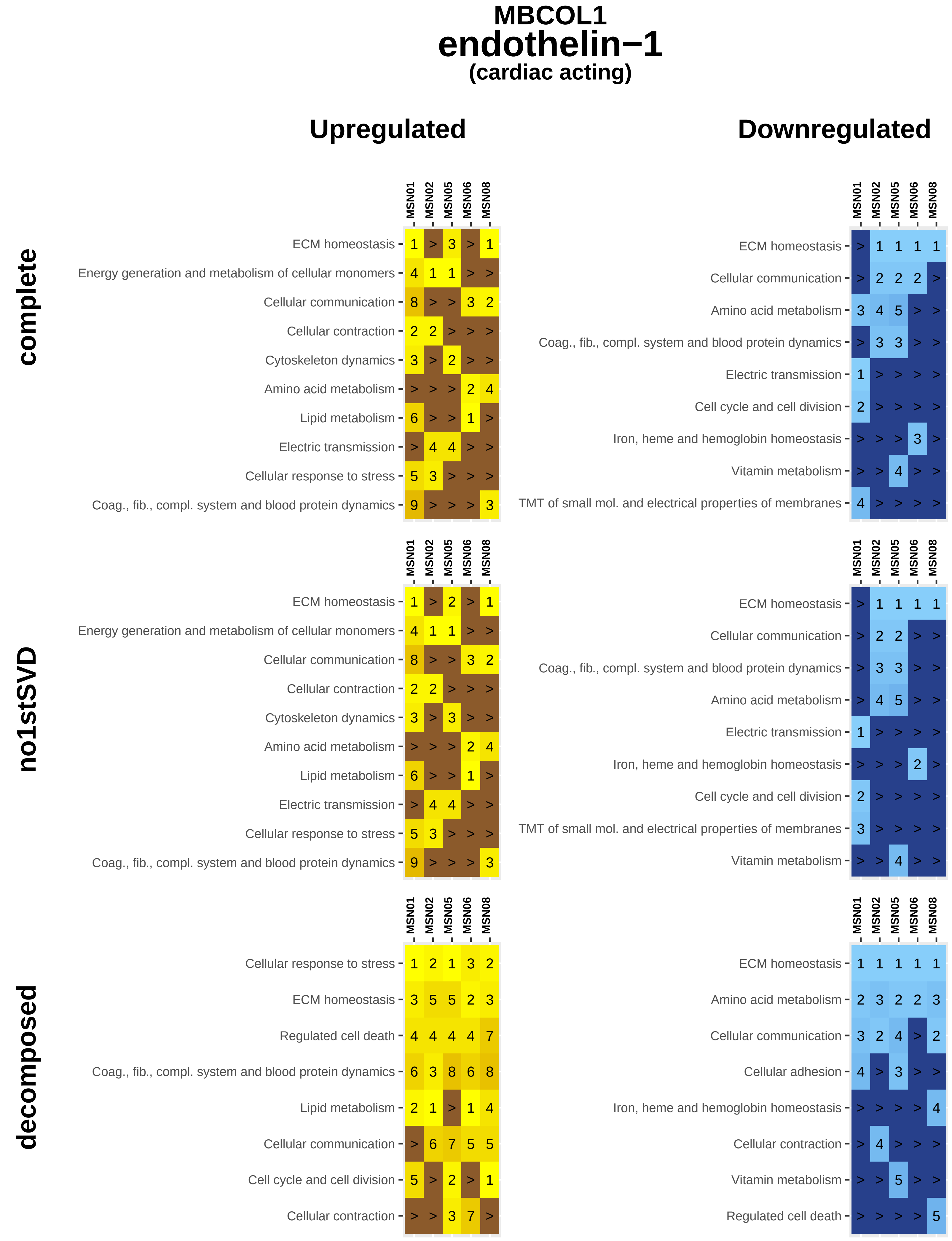
MBCOL1
endothelin−1
(cardiac acting)
Upregulated
Downregulated
complete
no1stSVD
decomposed
MSN01
MSN02
MSN05
MSN06
MSN08
ECM homeostasis
Energy generation and metabolism of cellular monomers
Cellular communication
Cellular contraction
Cytoskeleton dynamics
Amino acid metabolism
Lipid metabolism
Electric transmission
Cellular response to stress
Coag., fib., compl. system and blood protein dynamics
Cell cycle and cell division
Iron, heme and hemoglobin homeostasis
Vitamin metabolism
TMT of small mol. and electrical properties of membranes
Regulated cell death
Cellular adhesion

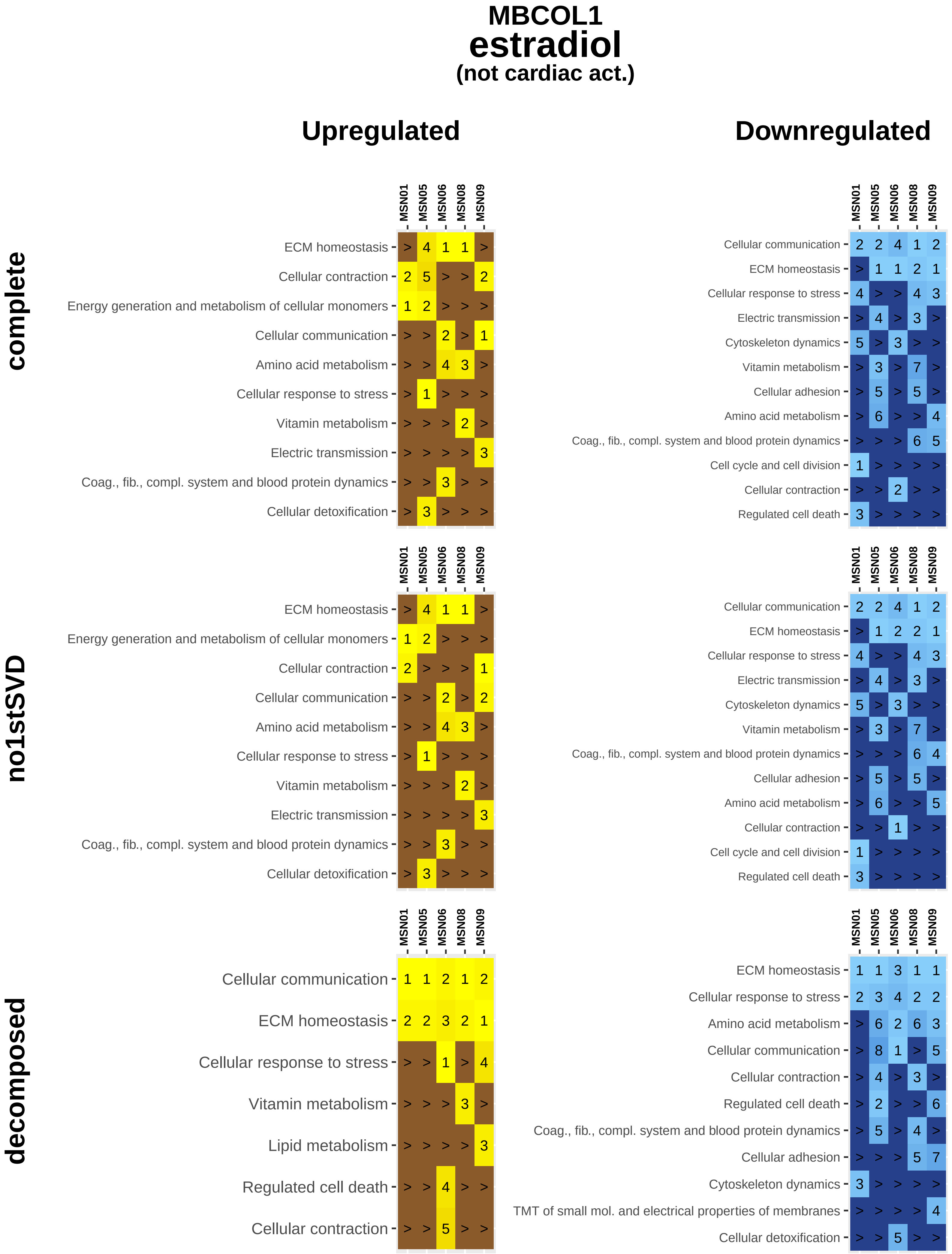

MBCOL1
estradiol
(not cardiac act.)
Upregulated
Downregulated
complete
no1stSVD
decomposed
MSN01
MSN05
MSN06
MSN08
MSN09
ECM homeostasis
Cellular contraction
Energy generation and metabolism of cellular monomers
Cellular communication
Amino acid metabolism
Cellular response to stress
Vitamin metabolism
Electric transmission
Coag., fib., compl. system and blood protein dynamics
Cellular detoxification
Cytoskeleton dynamics
Cellular adhesion
Cell cycle and cell division
Regulated cell death
Lipid metabolism
TMT of small mol. and electrical properties of membranes

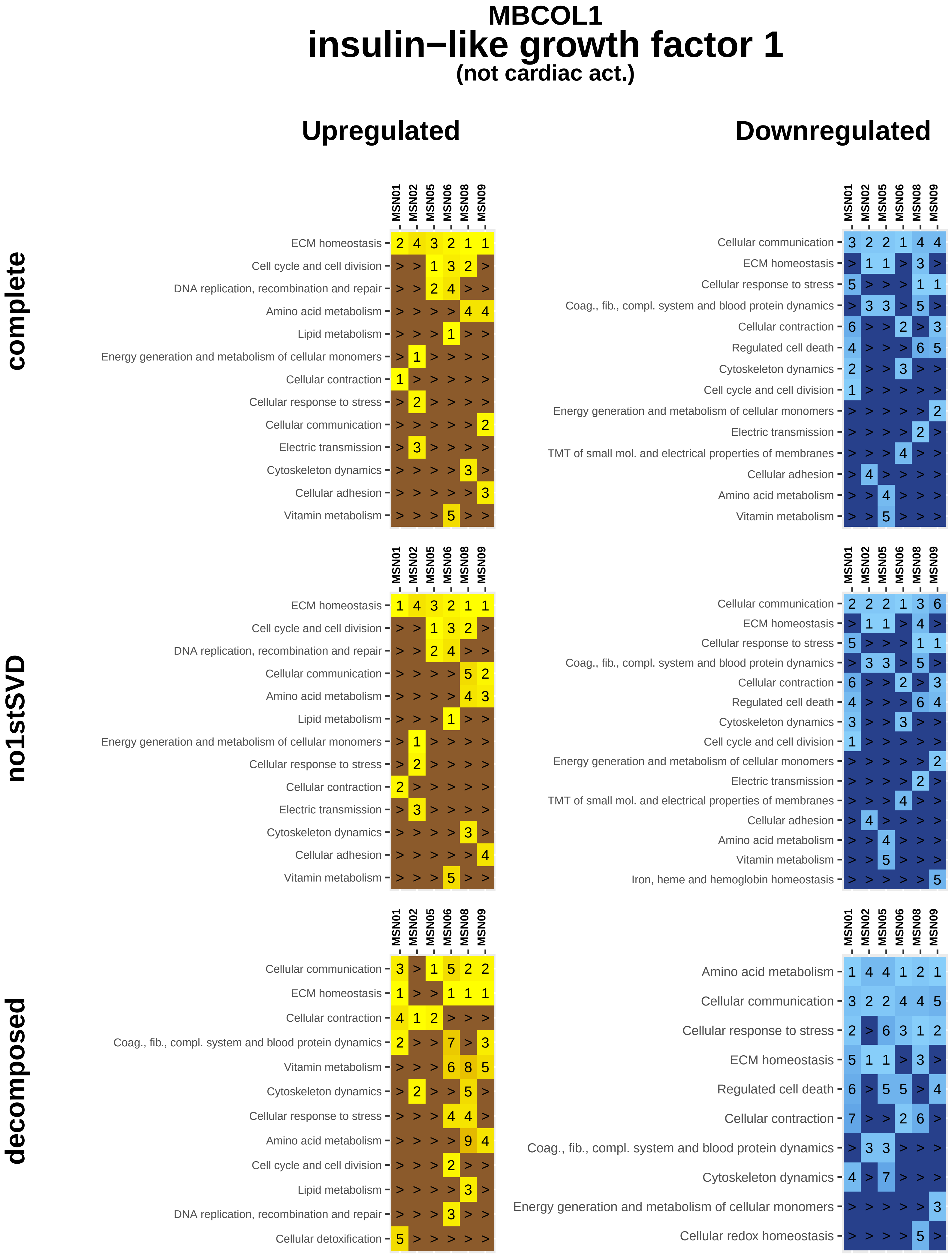

MBCOL1
insulin−like growth factor 1
(not cardiac act.)
Upregulated
Downregulated
complete
no1stSVD
decomposed
MSN01
MSN02
MSN05
MSN06
MSN08
MSN09
ECM homeostasis
Cell cycle and cell division
DNA replication, recombination and repair
Amino acid metabolism
Lipid metabolism
Energy generation and metabolism of cellular monomers
Cellular contraction
Cellular response to stress
Cellular communication
Electric transmission
Cytoskeleton dynamics
Cellular adhesion
Vitamin metabolism
Coag., fib., compl. system and blood protein dynamics
Regulated cell death
TMT of small mol. and electrical properties of membranes
Iron, heme and hemoglobin homeostasis
Cellular detoxification
Cellular redox homeostasis

### MBCOL1
### olmesartan
#### (cardiac acting)

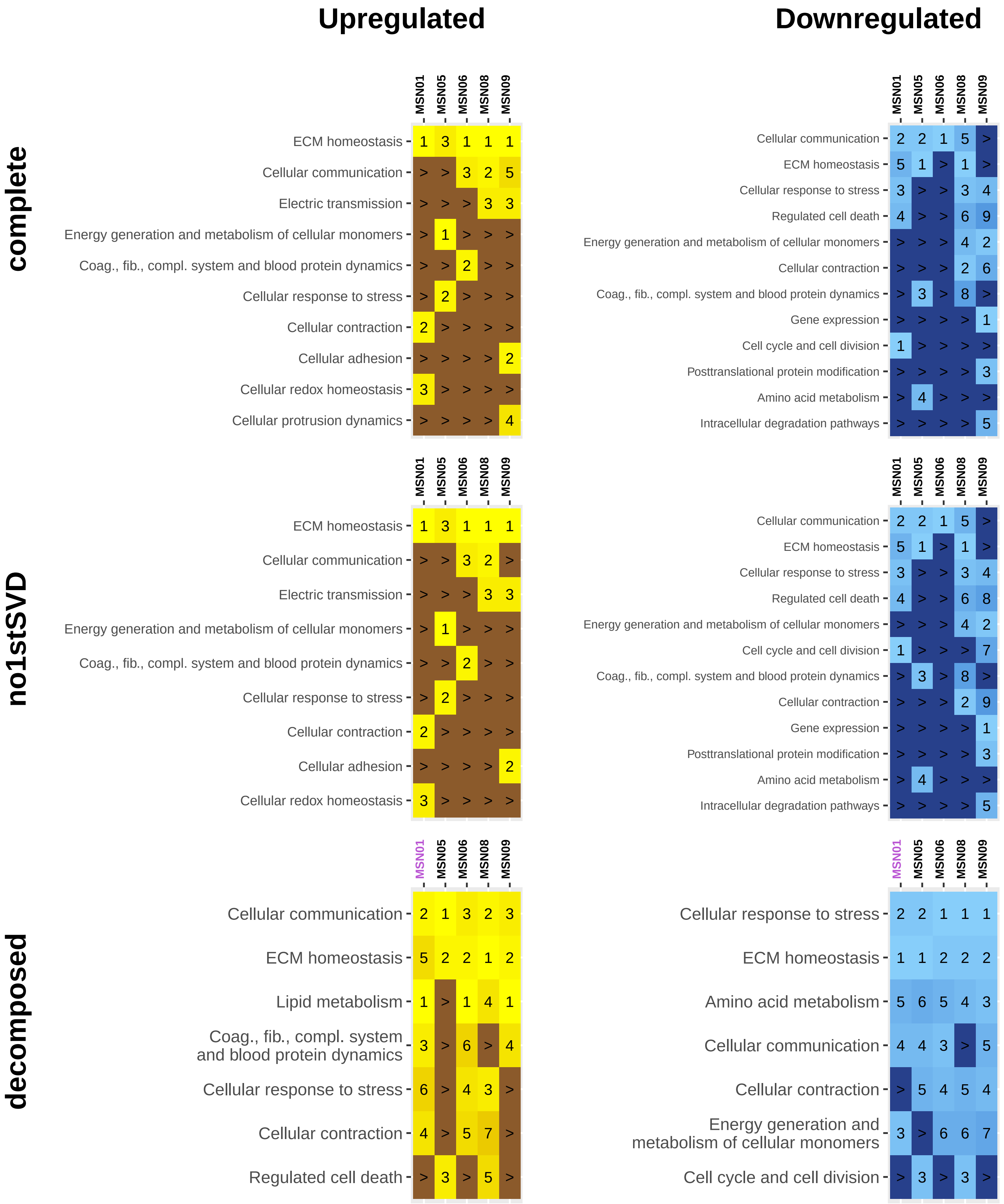

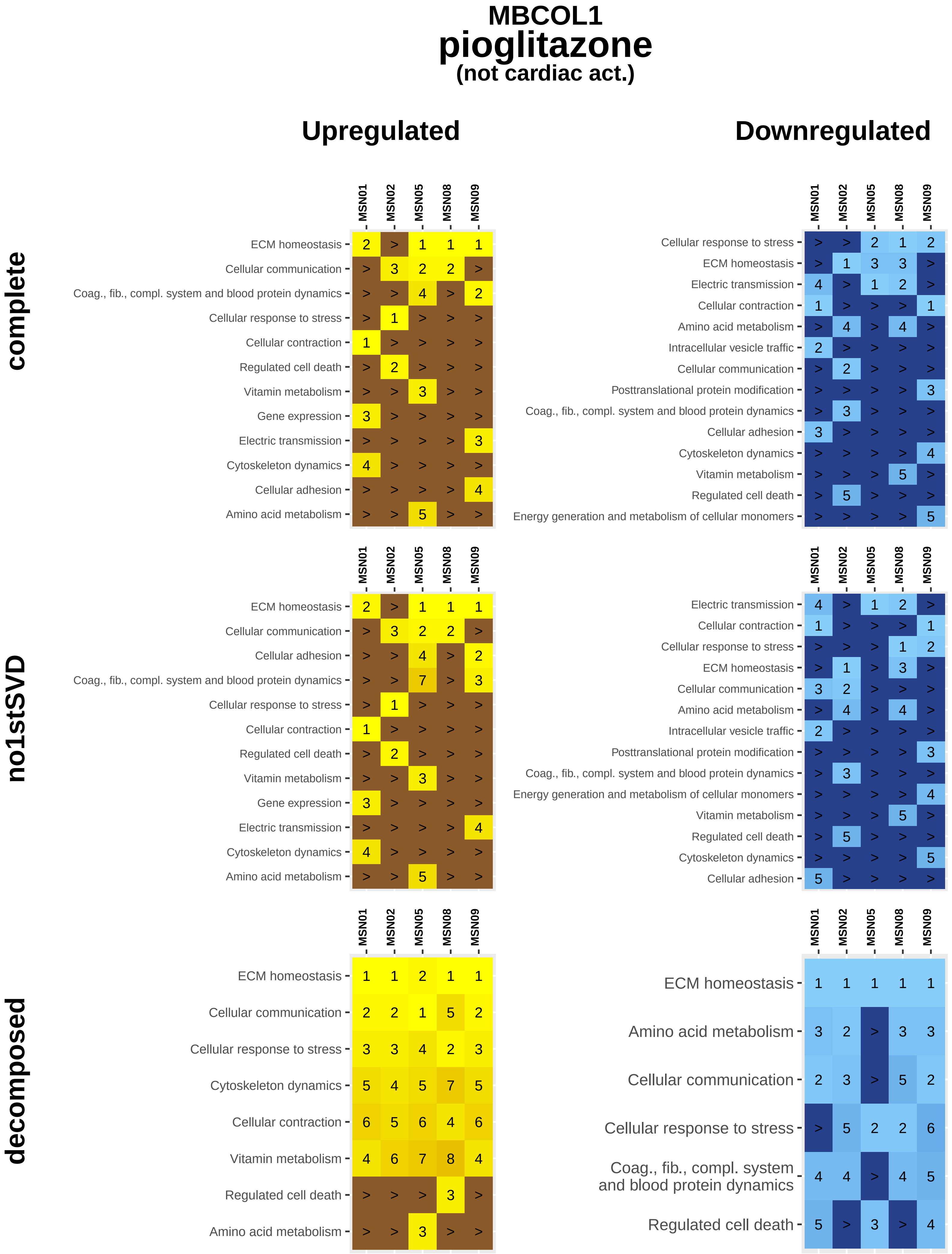
MBCOL1
pioglitazone
(not cardiac act.)
Upregulated
Downregulated
complete
no1stSVD
decomposed
MSN01
MSN02
MSN05
MSN08
MSN09
ECM homeostasis
Cellular communication
Coag., fib., compl. system and blood protein dynamics
Cellular response to stress
Cellular contraction
Regulated cell death
Vitamin metabolism
Gene expression
Electric transmission
Cytoskeleton dynamics
Cellular adhesion
Amino acid metabolism
Intracellular vesicle traffic
Posttranslational protein modification
Energy generation and metabolism of cellular monomers

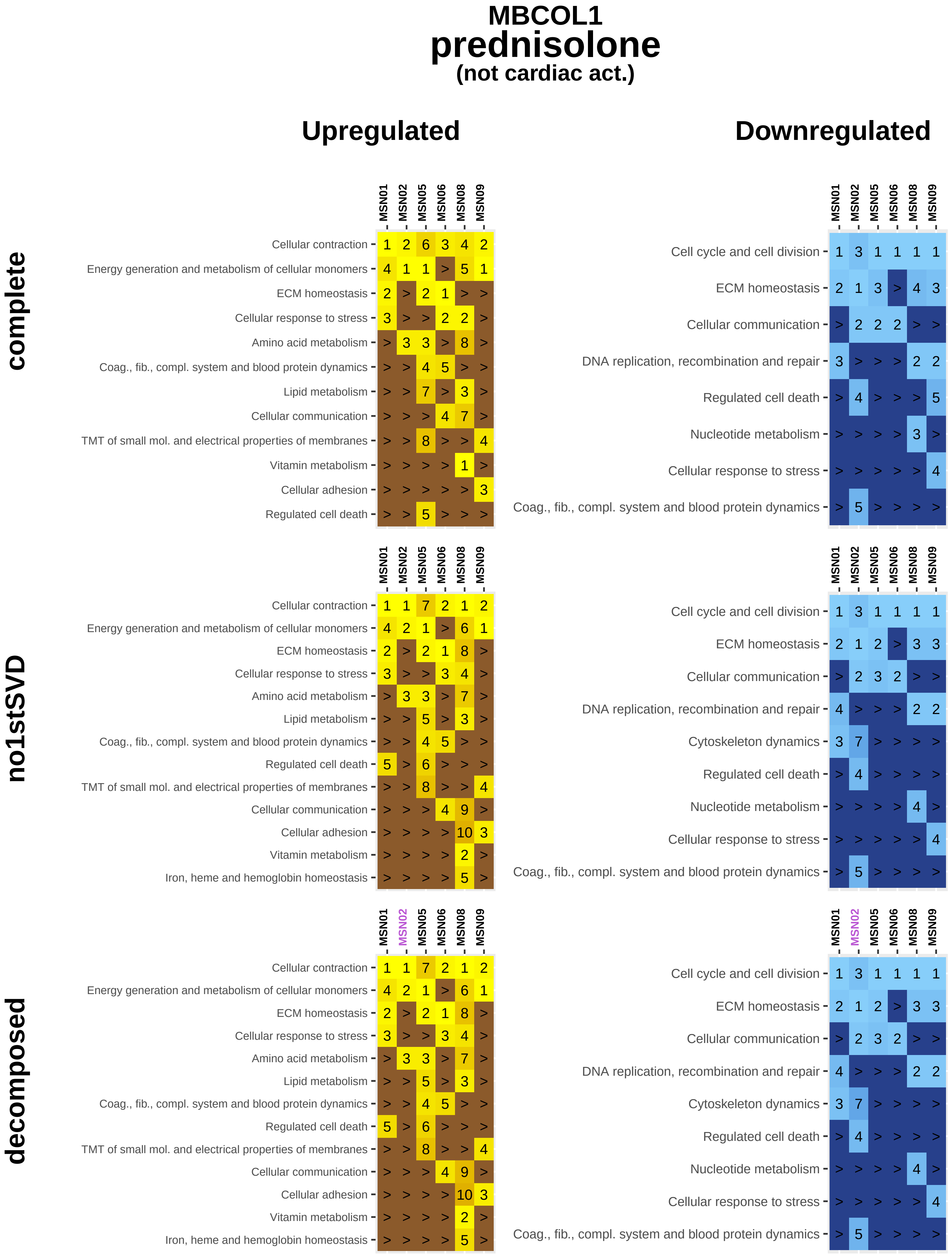
MBCOL1
prednisolone
(not cardiac act.)
Upregulated
Downregulated
complete
no1stSVD
decomposed
MSN01
MSN02
MSN05
MSN06
MSN08
MSN09
Cellular contraction
Energy generation and metabolism of cellular monomers
ECM homeostasis
Cellular response to stress
Amino acid metabolism
Coag., fib., compl. system and blood protein dynamics
Lipid metabolism
Cellular communication
TMT of small mol. and electrical properties of membranes
Vitamin metabolism
Cellular adhesion
Regulated cell death
Iron, heme and hemoglobin homeostasis
Cell cycle and cell division
DNA replication, recombination and repair
Nucleotide metabolism
Cytoskeleton dynamics

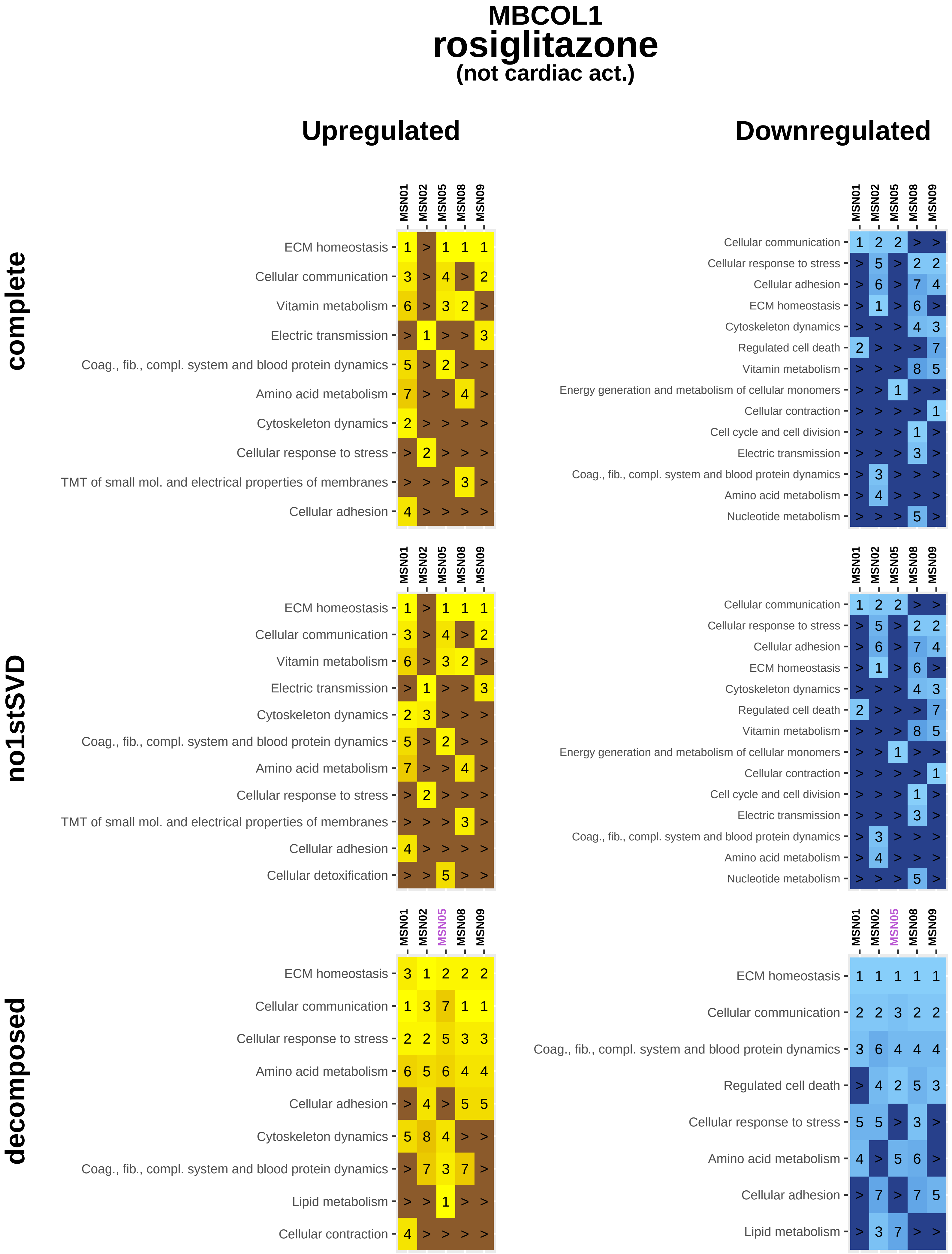

MBCOL1
rosiglitazone
(not cardiac act.)
Upregulated
Downregulated
complete
no1stSVD
decomposed
MSN01
MSN02
MSN05
MSN08
MSN09
ECM homeostasis 1 > 1 1 1
Cellular communication 3 > 4 > 2
Vitamin metabolism 6 > 3 2 >
Electric transmission > 1 > > 3
Coag., fib., compl. system and blood protein dynamics 5 > 2 > >
Amino acid metabolism 7 > > 4 >
Cytoskeleton dynamics 2 > > > >
Cellular response to stress > 2 > > >
TMT of small mol. and electrical properties of membranes > > > 3 >
Cellular adhesion 4 > > > >
Cellular communication 1 2 2 > >
Cellular response to stress > 5 > 2 2
Cellular adhesion > 6 > 7 4
ECM homeostasis > 1 > 6 >
Cytoskeleton dynamics > > > 4 3
Regulated cell death 2 > > > 7
Vitamin metabolism > > > 8 5
Energy generation and metabolism of cellular monomers > > 1 > >
Cellular contraction > > > > 1
Cell cycle and cell division > > > 1 >
Electric transmission > > > 3 >
Coag., fib., compl. system and blood protein dynamics > 3 > > >
Amino acid metabolism > 4 > > >
Nucleotide metabolism > > > 5 >
ECM homeostasis 1 > 1 1 1
Cellular communication 3 > 4 > 2
Vitamin metabolism 6 > 3 2 >
Electric transmission > 1 > > 3
Cytoskeleton dynamics 2 3 > > >
Coag., fib., compl. system and blood protein dynamics 5 > 2 > >
Amino acid metabolism 7 > > 4 >
Cellular response to stress > 2 > > >
TMT of small mol. and electrical properties of membranes > > > 3 >
Cellular adhesion 4 > > > >
Cellular detoxification > > 5 > >
Cellular communication 1 2 2 > >
Cellular response to stress > 5 > 2 2
Cellular adhesion > 6 > 7 4
ECM homeostasis > 1 > 6 >
Cytoskeleton dynamics > > > 4 3
Regulated cell death 2 > > > 7
Vitamin metabolism > > > 8 5
Energy generation and metabolism of cellular monomers > > 1 > >
Cellular contraction > > > > 1
Cell cycle and cell division > > > 1 >
Electric transmission > > > 3 >
Coag., fib., compl. system and blood protein dynamics > 3 > > >
Amino acid metabolism > 4 > > >
Nucleotide metabolism > > > 5 >
ECM homeostasis 3 1 2 2 2
Cellular communication 1 3 7 1 1
Cellular response to stress 2 2 5 3 3
Amino acid metabolism 6 5 6 4 4
Cellular adhesion > 4 > 5 5
Cytoskeleton dynamics 5 8 4 > >
Coag., fib., compl. system and blood protein dynamics > 7 3 7 >
Lipid metabolism > > 1 > >
Cellular contraction 4 > > > >
ECM homeostasis 1 1 1 1 1
Cellular communication 2 2 3 2 2
Coag., fib., compl. system and blood protein dynamics 3 6 4 4 4
Regulated cell death > 4 2 5 3
Cellular response to stress 5 5 > 3 >
Amino acid metabolism 4 > 5 6 >
Cellular adhesion > 7 > 7 5
Lipid metabolism > 3 7 > >

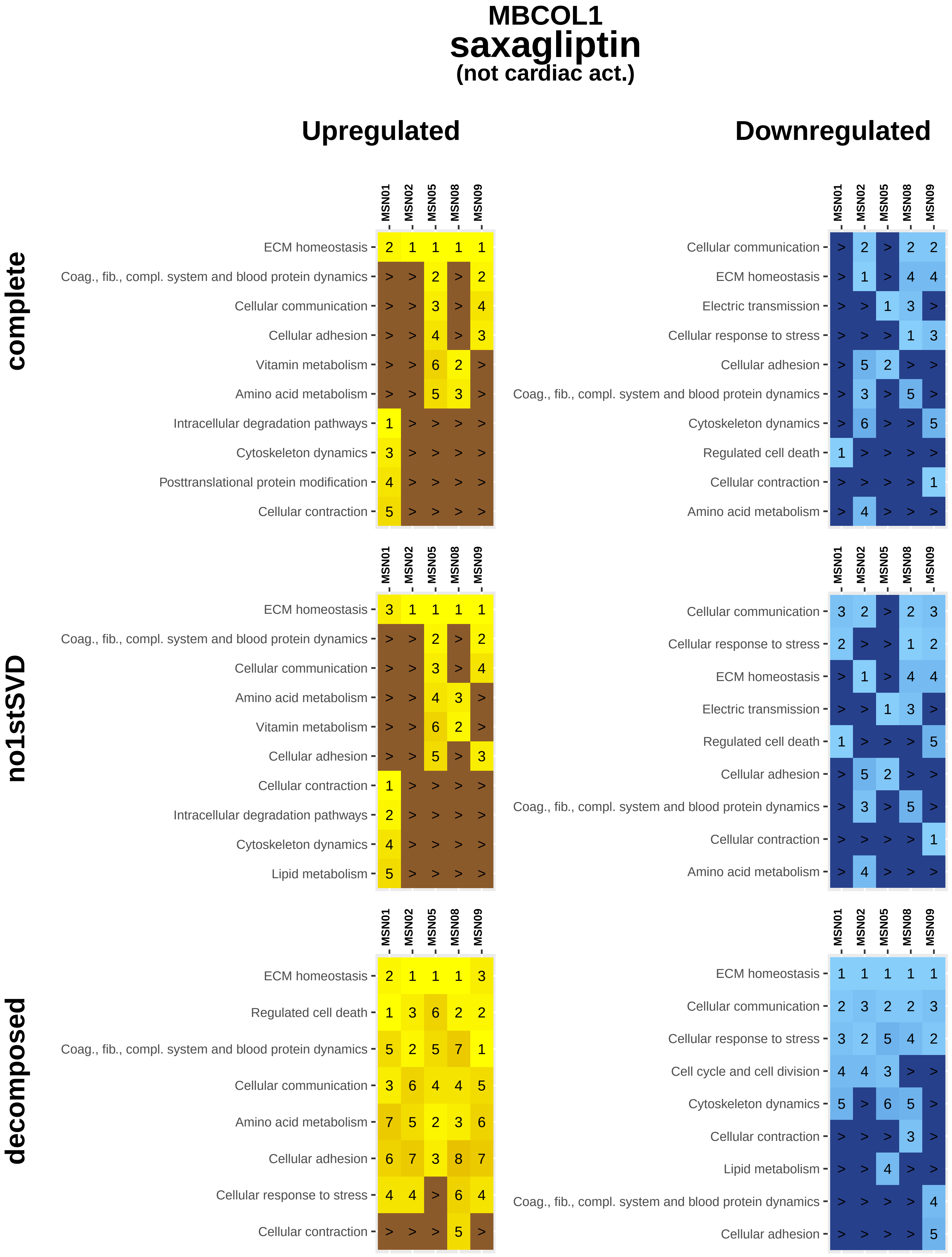
MBCOL1
saxagliptin
(not cardiac act.)
Upregulated
Downregulated
complete
no1stSVD
decomposed
MSN01 MSN02 MSN05 MSN08 MSN09
ECM homeostasis 2 1 1 1 1
Coag., fib., compl. system and blood protein dynamics > > 2 > 2
Cellular communication > > 3 > 4
Cellular adhesion > > 4 > 3
Vitamin metabolism > > 6 2 >
Amino acid metabolism > > 5 3 >
Intracellular degradation pathways 1 > > > >
Cytoskeleton dynamics 3 > > > >
Posttranslational protein modification 4 > > > >
Cellular contraction 5 > > > >
MSN01 MSN02 MSN05 MSN08 MSN09
Cellular communication > 2 > 2 2
ECM homeostasis > 1 > 4 4
Electric transmission > > 1 3 >
Cellular response to stress > > > 1 3
Cellular adhesion > 5 2 > >
Coag., fib., compl. system and blood protein dynamics > 3 > 5 >
Cytoskeleton dynamics > 6 > > 5
Regulated cell death 1 > > > >
Cellular contraction > > > > 1
Amino acid metabolism > 4 > > >
MSN01 MSN02 MSN05 MSN08 MSN09
ECM homeostasis 3 1 1 1 1
Coag., fib., compl. system and blood protein dynamics > > 2 > 2
Cellular communication > > 3 > 4
Amino acid metabolism > > 4 3 >
Vitamin metabolism > > 6 2 >
Cellular adhesion > > 5 > 3
Cellular contraction 1 > > > >
Intracellular degradation pathways 2 > > > >
Cytoskeleton dynamics 4 > > > >
Lipid metabolism 5 > > > >
MSN01 MSN02 MSN05 MSN08 MSN09
Cellular communication 3 2 > 2 3
Cellular response to stress 2 > > 1 2
ECM homeostasis > 1 > 4 4
Electric transmission > > 1 3 >
Regulated cell death 1 > > > 5
Cellular adhesion > 5 2 > >
Coag., fib., compl. system and blood protein dynamics > 3 > 5 >
Cellular contraction > > > > 1
Amino acid metabolism > 4 > > >
MSN01 MSN02 MSN05 MSN08 MSN09
ECM homeostasis 2 1 1 1 3
Regulated cell death 1 3 6 2 2
Coag., fib., compl. system and blood protein dynamics 5 2 5 7 1
Cellular communication 3 6 4 4 5
Amino acid metabolism 7 5 2 3 6
Cellular adhesion 6 7 3 8 7
Cellular response to stress 4 4 > 6 4
Cellular contraction > > > 5 >
MSN01 MSN02 MSN05 MSN08 MSN09
ECM homeostasis 1 1 1 1 1
Cellular communication 2 3 2 2 3
Cellular response to stress 3 2 5 4 2
Cell cycle and cell division 4 4 3 > >
Cytoskeleton dynamics 5 > 6 5 >
Cellular contraction > > > 3 >
Lipid metabolism > > 4 > >
Coag., fib., compl. system and blood protein dynamics > > > > 4
Cellular adhesion > > > > 5

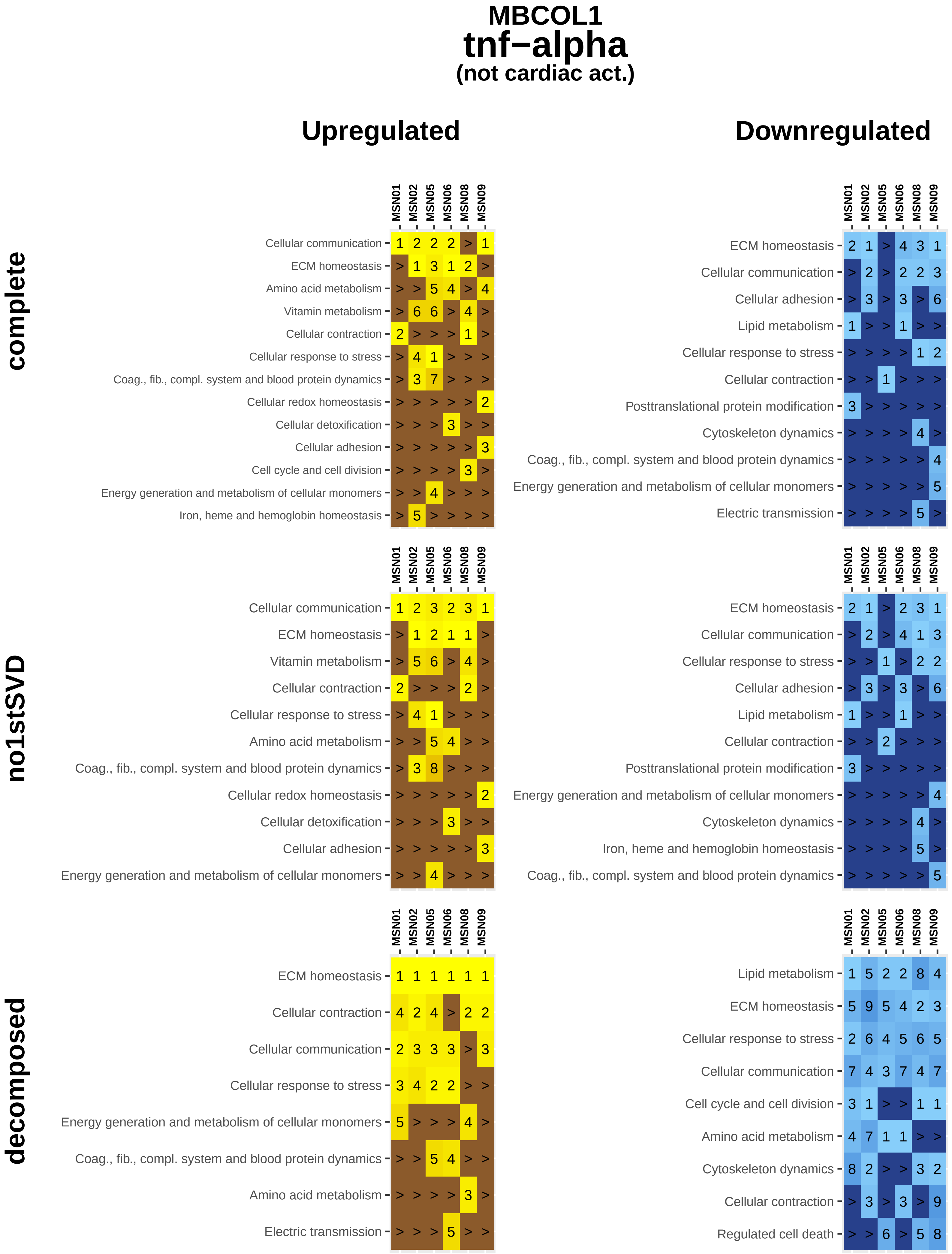
MBCOL1
tnf−alpha
(not cardiac act.)
Upregulated
Downregulated
complete
no1stSVD
decomposed
MSN01
MSN02
MSN05
MSN06
MSN08
MSN09
Cellular communication
ECM homeostasis
Amino acid metabolism
Vitamin metabolism
Cellular contraction
Cellular response to stress
Coag., fib., compl. system and blood protein dynamics
Cellular redox homeostasis
Cellular detoxification
Cellular adhesion
Cell cycle and cell division
Energy generation and metabolism of cellular monomers
Iron, heme and hemoglobin homeostasis
Lipid metabolism
Posttranslational protein modification
Cytoskeleton dynamics
Electric transmission
Regulated cell death
