## Supplemental Figure 14C for "Multiscale mapping of transcriptomic signatures for cardiotoxic drugs"

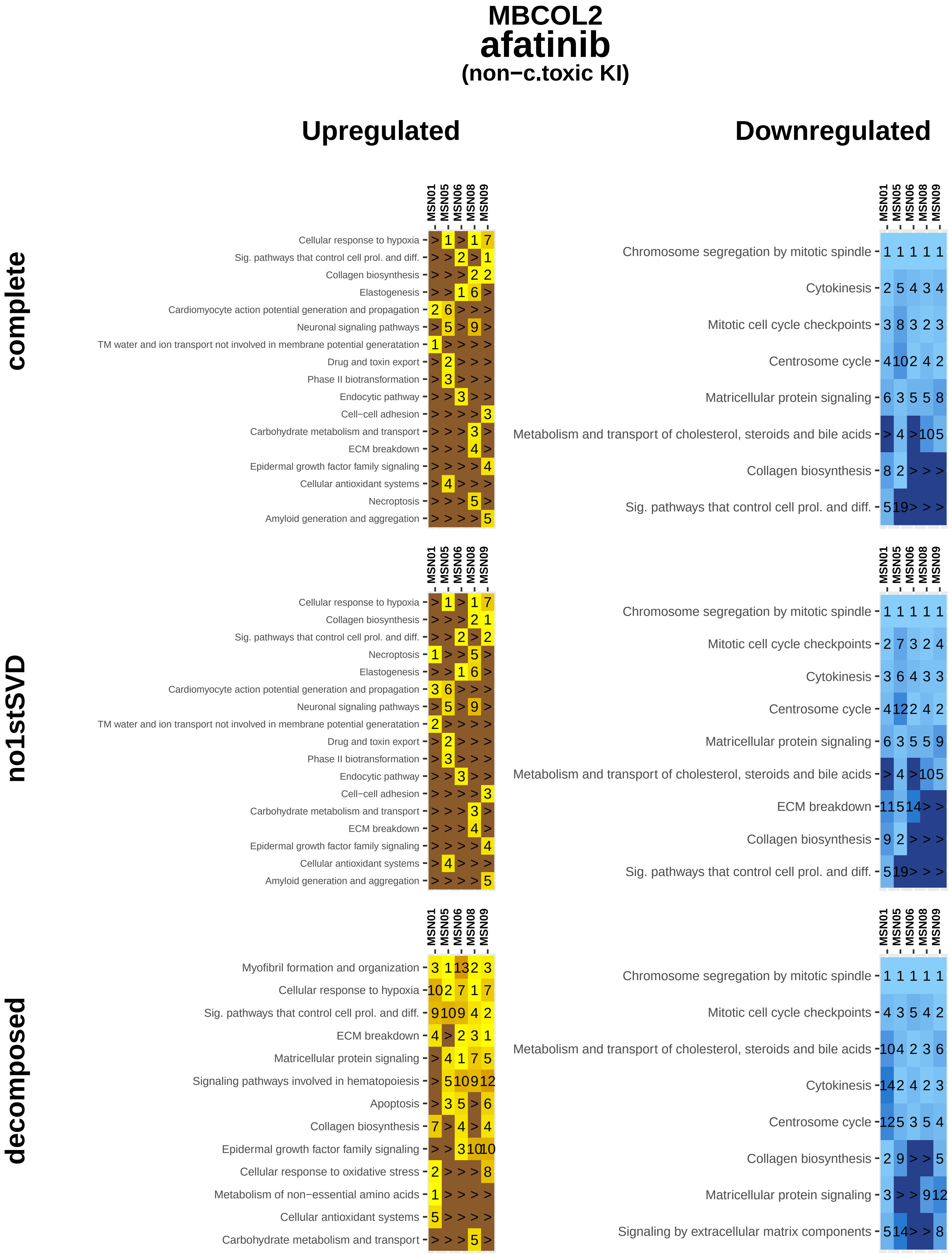

MBCOL2
afatinib
(non−c.toxic KI)
Upregulated
Downregulated
complete
no1stSVD
decomposed
MSN01
MSN05
MSN06
MSN08
MSN09
Cellular response to hypoxia
Sig. pathways that control cell prol. and diff.
Collagen biosynthesis
Elastogenesis
Cardiomyocyte action potential generation and propagation
Neuronal signaling pathways
TM water and ion transport not involved in membrane potential generatation
Drug and toxin export
Phase II biotransformation
Endocytic pathway
Cell−cell adhesion
Carbohydrate metabolism and transport
ECM breakdown
Epidermal growth factor family signaling
Cellular antioxidant systems
Necroptosis
Amyloid generation and aggregation
Chromosome segregation by mitotic spindle
Cytokinesis
Mitotic cell cycle checkpoints
Centrosome cycle
Matricellular protein signaling
Metabolism and transport of cholesterol, steroids and bile acids
Myofibril formation and organization
Signaling pathways involved in hematopoiesis
Apoptosis
Cellular response to oxidative stress
Metabolism of non−essential amino acids
Signaling by extracellular matrix components

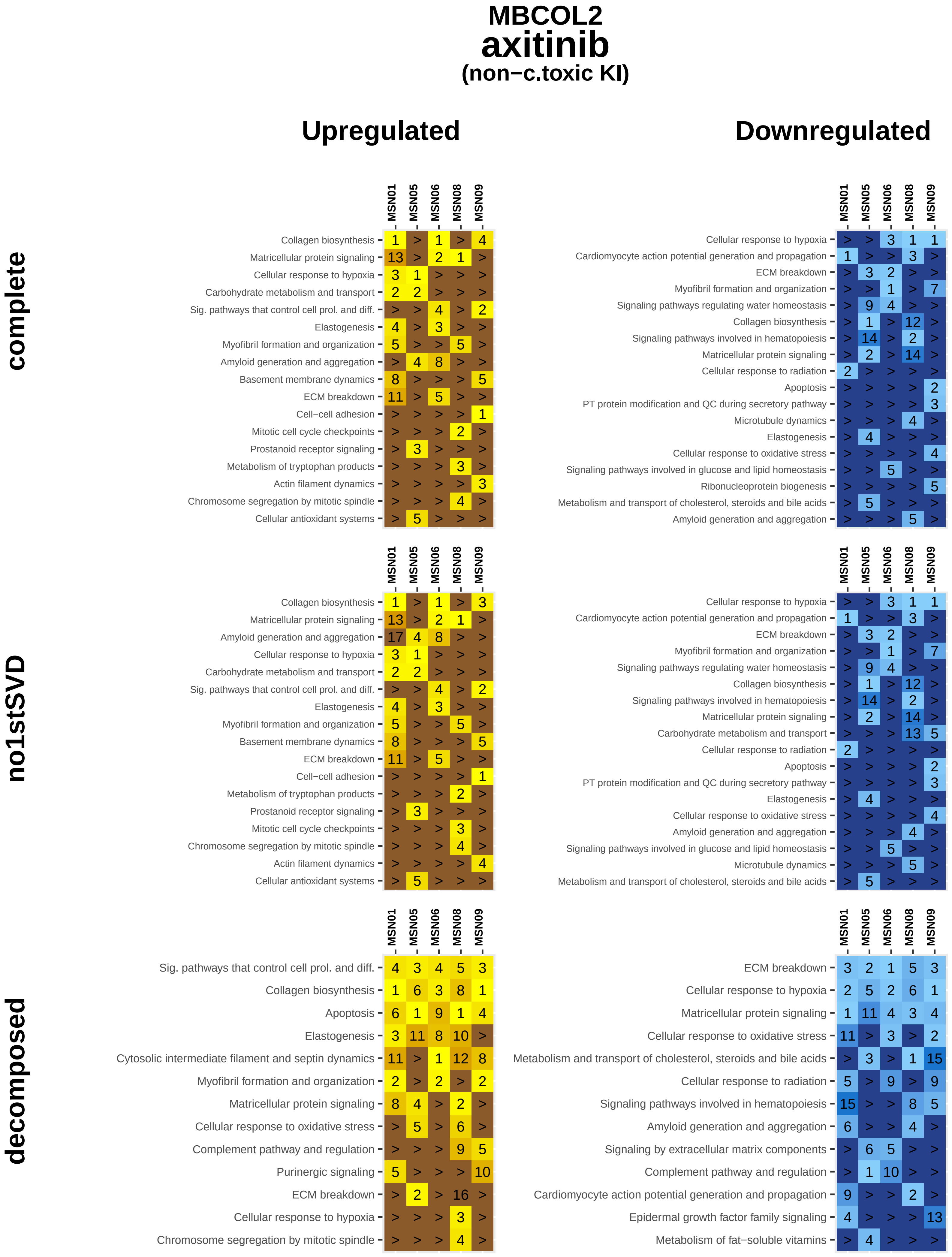

MBCOL2
axitinib
(non−c.toxic KI)
Upregulated
Downregulated
complete
no1stSVD
decomposed
MSN01
MSN05
MSN06
MSN08
MSN09
Collagen biosynthesis
Matricellular protein signaling
Cellular response to hypoxia
Carbohydrate metabolism and transport
Sig. pathways that control cell prol. and diff.
Elastogenesis
Myofibril formation and organization
Amyloid generation and aggregation
Basement membrane dynamics
ECM breakdown
Cell−cell adhesion
Mitotic cell cycle checkpoints
Prostanoid receptor signaling
Metabolism of tryptophan products
Actin filament dynamics
Chromosome segregation by mitotic spindle
Cellular antioxidant systems
Cardiomyocyte action potential generation and propagation
Signaling pathways regulating water homeostasis
Signaling pathways involved in hematopoiesis
Cellular response to radiation
Apoptosis
PT protein modification and QC during secretory pathway
Microtubule dynamics
Cellular response to oxidative stress
Signaling pathways involved in glucose and lipid homeostasis
Ribonucleoprotein biogenesis
Metabolism and transport of cholesterol, steroids and bile acids
Cytosolic intermediate filament and septin dynamics
Complement pathway and regulation
Purinergic signaling
Signaling by extracellular matrix components
Epidermal growth factor family signaling
Metabolism of fat−soluble vitamins

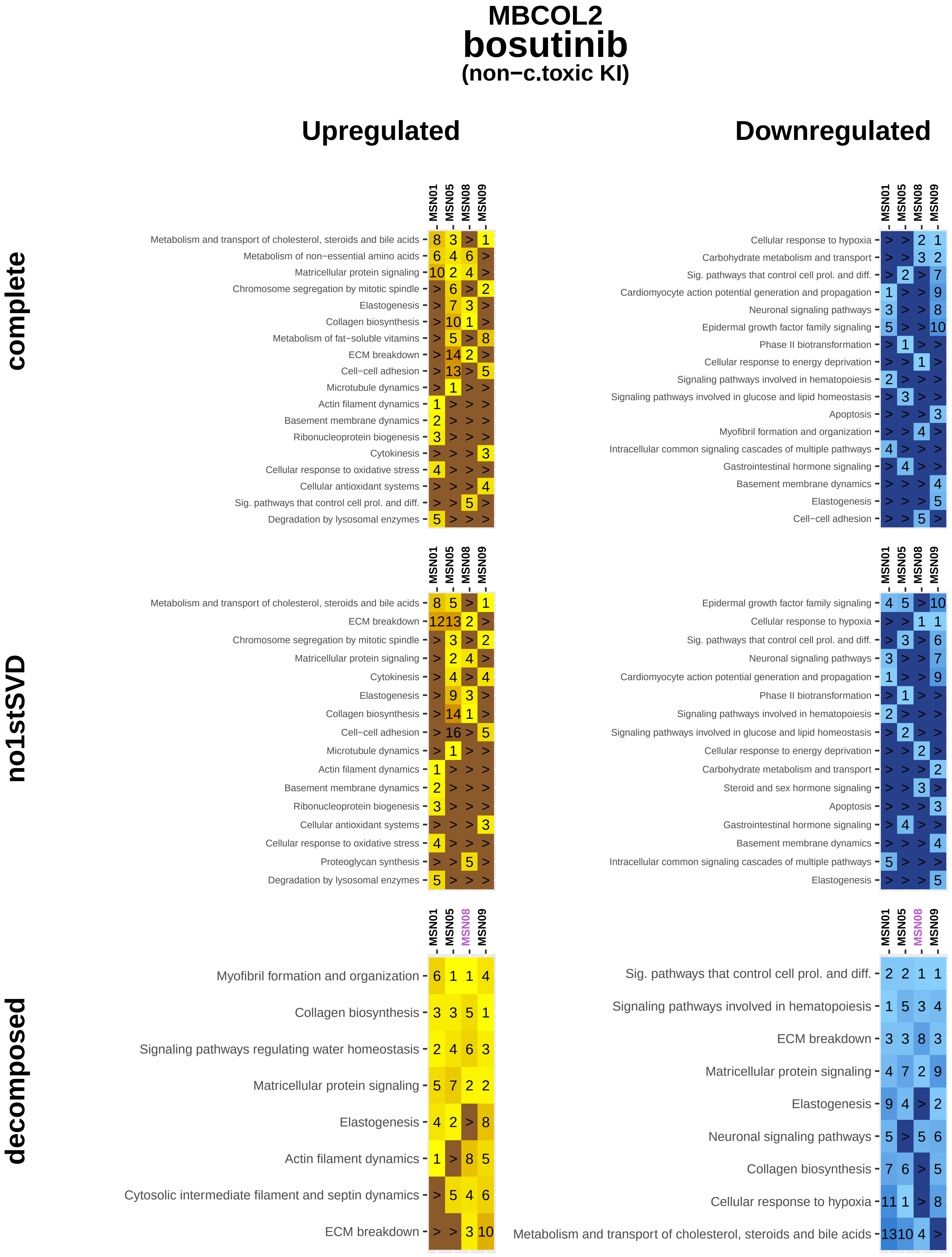
MBCOL2
bosutinib
(non−c.toxic KI)
Upregulated
Downregulated
complete
no1stSVD
decomposed
MSN01
MSN05
MSN08
MSN09
Metabolism and transport of cholesterol, steroids and bile acids 8 3 > 1
Metabolism of non−essential amino acids 6 4 6 >
Matricellular protein signaling 10 2 4 >
Chromosome segregation by mitotic spindle > 6 > 2
Elastogenesis > 7 3 >
Collagen biosynthesis > 10 1 >
Metabolism of fat−soluble vitamins > 5 > 8
ECM breakdown > 14 2 >
Cell−cell adhesion > 13 > 5
Microtubule dynamics > 1 > >
Actin filament dynamics 1 > > >
Basement membrane dynamics 2 > > >
Ribonucleoprotein biogenesis 3 > > >
Cytokinesis > > > 3
Cellular response to oxidative stress 4 > > >
Cellular antioxidant systems > > > 4
Sig. pathways that control cell prol. and diff. > > 5 >
Degradation by lysosomal enzymes 5 > > >
Cellular response to hypoxia > > 2 1
Carbohydrate metabolism and transport > > 3 2
Sig. pathways that control cell prol. and diff. > 2 > 7
Cardiomyocyte action potential generation and propagation 1 > > 9
Neuronal signaling pathways 3 > > 8
Epidermal growth factor family signaling 5 > > 10
Phase II biotransformation > 1 > >
Cellular response to energy deprivation > > 1 >
Signaling pathways involved in hematopoiesis 2 > > >
Signaling pathways involved in glucose and lipid homeostasis > 3 > >
Apoptosis > > > 3
Myofibril formation and organization > > 4 >
Intracellular common signaling cascades of multiple pathways 4 > > >
Gastrointestinal hormone signaling > 4 > >
Basement membrane dynamics > > > 4
Elastogenesis > > > 5
Cell−cell adhesion > > 5 >
Metabolism and transport of cholesterol, steroids and bile acids 8 5 > 1
ECM breakdown 12 13 2 >
Chromosome segregation by mitotic spindle > 3 > 2
Matricellular protein signaling > 2 4 >
Cytokinesis > 4 > 4
Elastogenesis > 9 3 >
Collagen biosynthesis > 14 1 >
Cell−cell adhesion > 16 > 5
Microtubule dynamics > 1 > >
Actin filament dynamics 1 > > >
Basement membrane dynamics 2 > > >
Ribonucleoprotein biogenesis 3 > > >
Cellular antioxidant systems > > > 3
Cellular response to oxidative stress 4 > > >
Proteoglycan synthesis > > 5 >
Degradation by lysosomal enzymes 5 > > >
Epidermal growth factor family signaling 4 5 > 10
Cellular response to hypoxia > > 1 1
Sig. pathways that control cell prol. and diff. > 3 > 6
Neuronal signaling pathways 3 > > 7
Cardiomyocyte action potential generation and propagation 1 > > 9
Phase II biotransformation > 1 > >
Signaling pathways involved in hematopoiesis 2 > > >
Signaling pathways involved in glucose and lipid homeostasis > 2 > >
Cellular response to energy deprivation > > 2 >
Carbohydrate metabolism and transport > > > 2
Steroid and sex hormone signaling > > 3 >
Apoptosis > > > 3
Gastrointestinal hormone signaling > 4 > >
Basement membrane dynamics > > > 4
Intracellular common signaling cascades of multiple pathways 5 > > >
Elastogenesis > > > 5
Myofibril formation and organization 6 1 1 4
Collagen biosynthesis 3 3 5 1
Signaling pathways regulating water homeostasis 2 4 6 3
Matricellular protein signaling 5 7 2 2
Elastogenesis 4 2 > 8
Actin filament dynamics 1 > 8 5
Cytosolic intermediate filament and septin dynamics > 5 4 6
ECM breakdown > > 3 10
Sig. pathways that control cell prol. and diff. 2 2 1 1
Signaling pathways involved in hematopoiesis 1 5 3 4
ECM breakdown 3 3 8 3
Matricellular protein signaling 4 7 2 9
Elastogenesis 9 4 > 2
Neuronal signaling pathways 5 > 5 6
Collagen biosynthesis 7 6 > 5
Cellular response to hypoxia 11 1 > 8
Metabolism and transport of cholesterol, steroids and bile acids 13 10 4 >

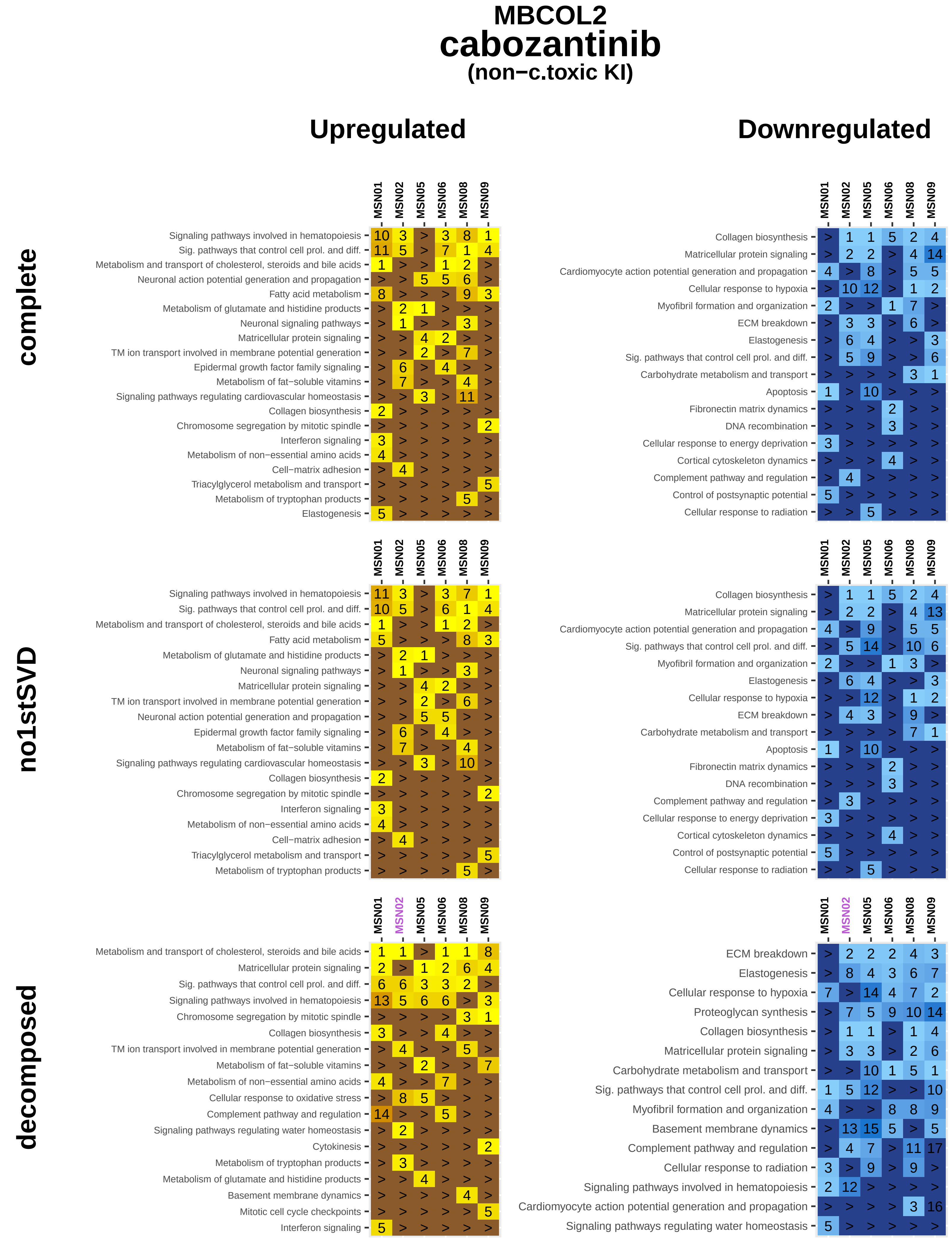
MBCOL2
cabozantinib
(non−c.toxic KI)
Upregulated
Downregulated
complete
no1stSVD
decomposed
MSN01
MSN02
MSN05
MSN06
MSN08
MSN09
Signaling pathways involved in hematopoiesis
Sig. pathways that control cell prol. and diff.
Metabolism and transport of cholesterol, steroids and bile acids
Neuronal action potential generation and propagation
Fatty acid metabolism
Metabolism of glutamate and histidine products
Neuronal signaling pathways
Matricellular protein signaling
TM ion transport involved in membrane potential generation
Epidermal growth factor family signaling
Metabolism of fat−soluble vitamins
Signaling pathways regulating cardiovascular homeostasis
Collagen biosynthesis
Chromosome segregation by mitotic spindle
Interferon signaling
Metabolism of non−essential amino acids
Cell−matrix adhesion
Triacylglycerol metabolism and transport
Metabolism of tryptophan products
Elastogenesis
Cardiomyocyte action potential generation and propagation
Cellular response to hypoxia
Myofibril formation and organization
ECM breakdown
Carbohydrate metabolism and transport
Apoptosis
Fibronectin matrix dynamics
DNA recombination
Cellular response to energy deprivation
Cortical cytoskeleton dynamics
Complement pathway and regulation
Control of postsynaptic potential
Cellular response to radiation
Cellular response to oxidative stress
Signaling pathways regulating water homeostasis
Cytokinesis
Basement membrane dynamics
Mitotic cell cycle checkpoints
Proteoglycan synthesis

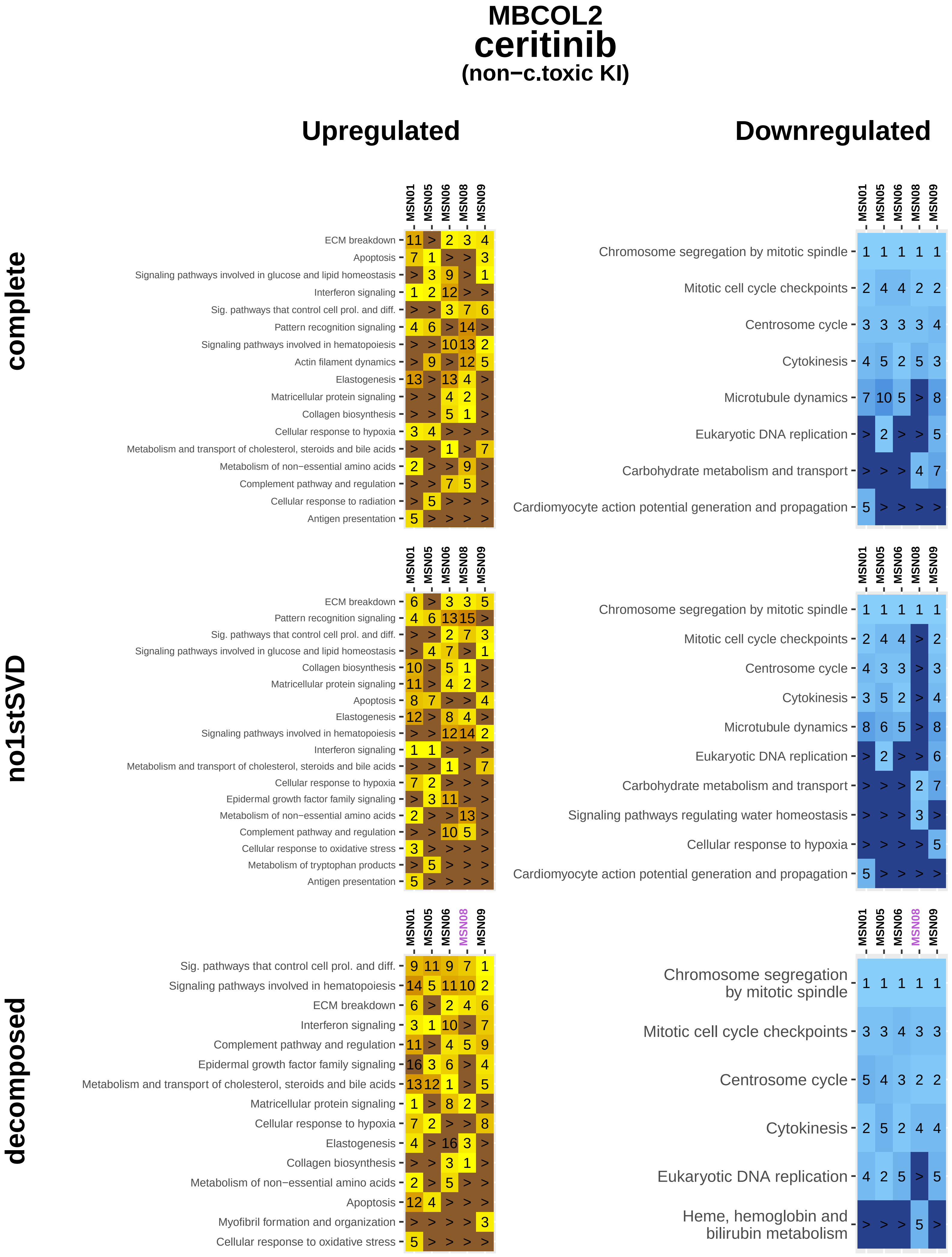

MBCOL2
ceritinib
(non−c.toxic KI)
Upregulated
Downregulated
complete
no1stSVD
decomposed
MSN01
MSN05
MSN06
MSN08
MSN09
ECM breakdown
Apoptosis
Signaling pathways involved in glucose and lipid homeostasis
Interferon signaling
Sig. pathways that control cell prol. and diff.
Pattern recognition signaling
Signaling pathways involved in hematopoiesis
Actin filament dynamics
Elastogenesis
Matricellular protein signaling
Collagen biosynthesis
Cellular response to hypoxia
Metabolism and transport of cholesterol, steroids and bile acids
Metabolism of non−essential amino acids
Complement pathway and regulation
Cellular response to radiation
Antigen presentation
Epidermal growth factor family signaling
Cellular response to oxidative stress
Metabolism of tryptophan products
Myofibril formation and organization
Chromosome segregation by mitotic spindle
Mitotic cell cycle checkpoints
Centrosome cycle
Cytokinesis
Microtubule dynamics
Eukaryotic DNA replication
Carbohydrate metabolism and transport
Cardiomyocyte action potential generation and propagation
Signaling pathways regulating water homeostasis
Heme, hemoglobin and bilirubin metabolism

### MBCOL2
### crizotinib
#### (non−c.toxic KI)

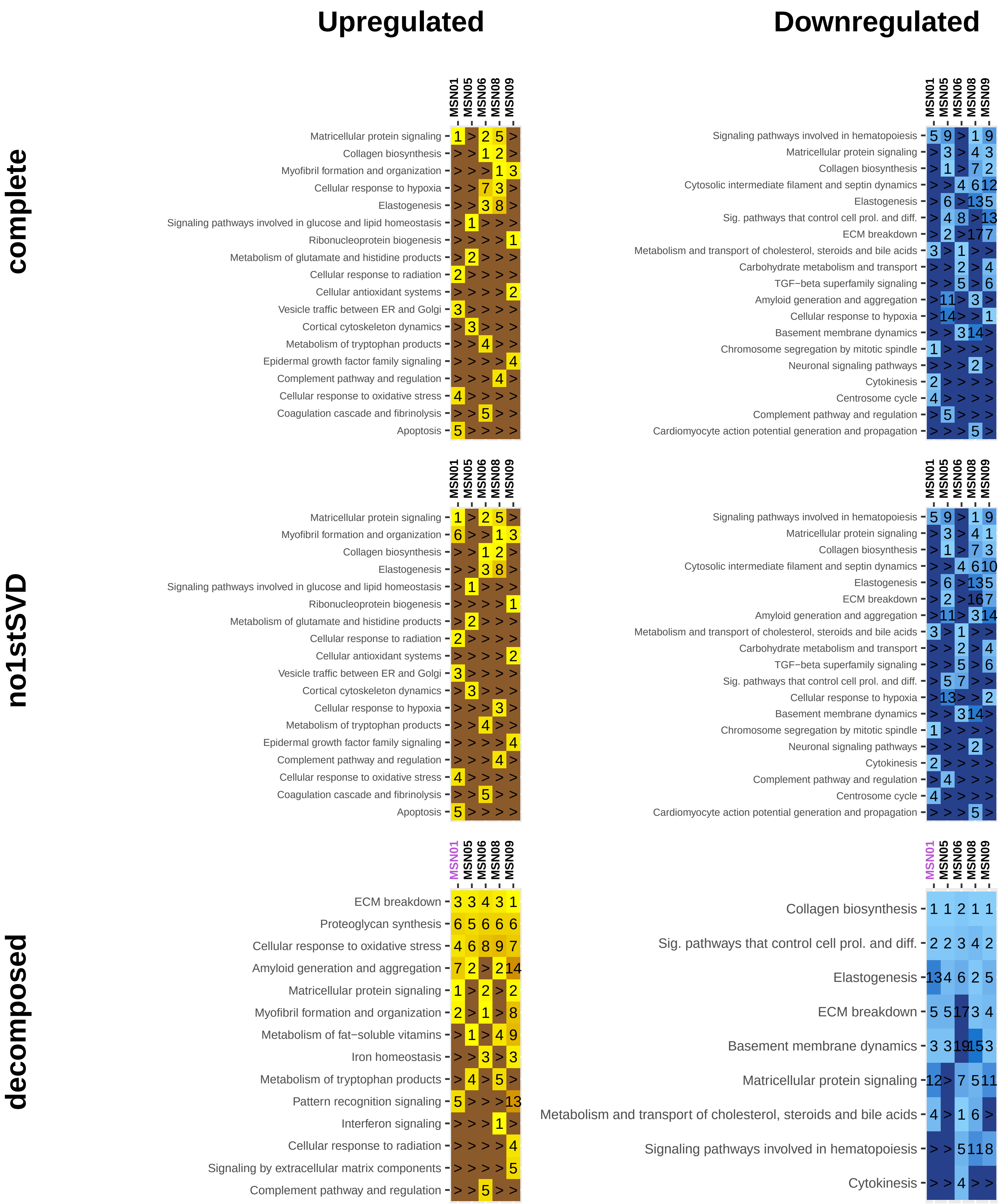

### MBCOL2
### dabrafenib
#### (c.toxic KI)

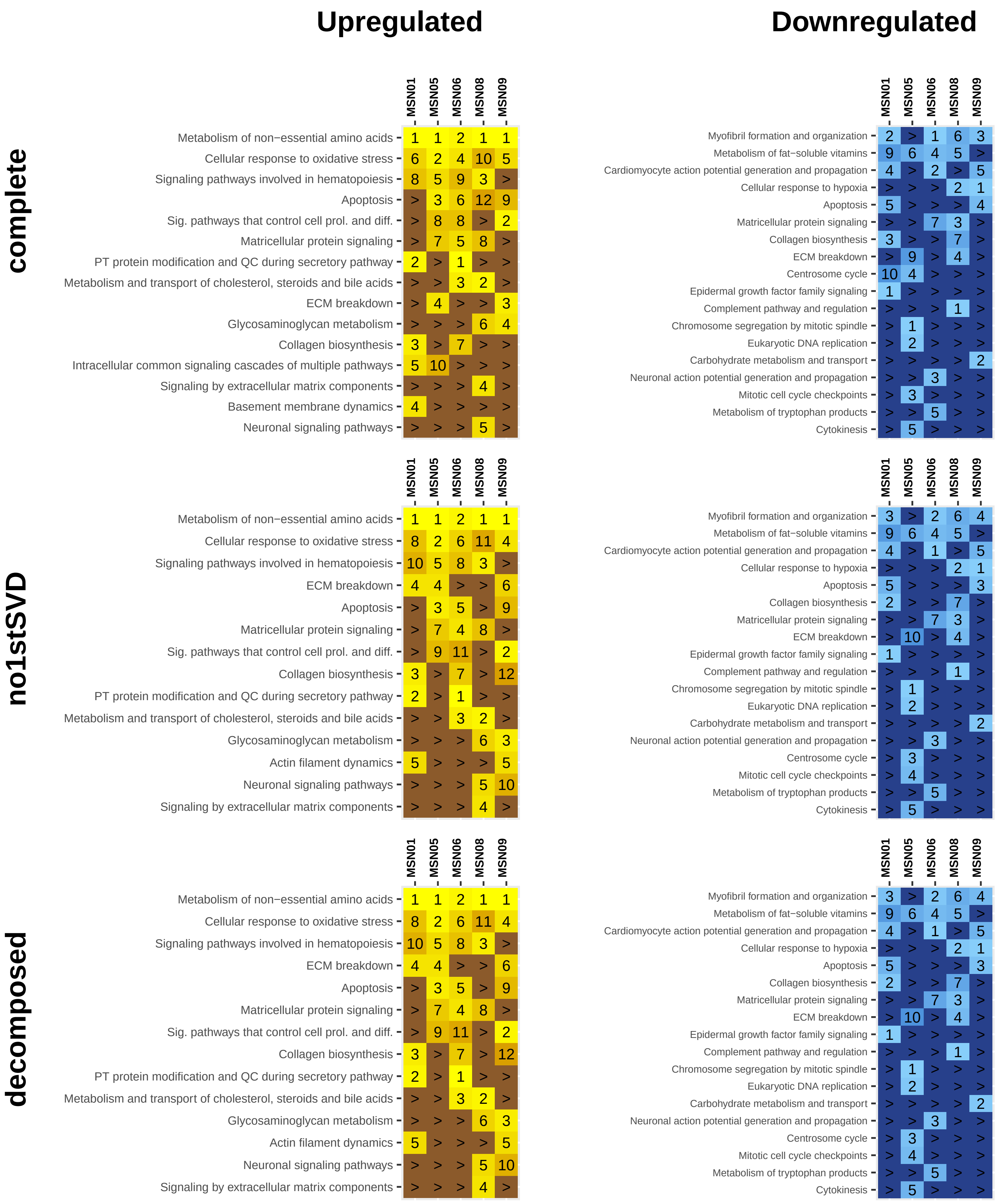

### MBCOL2
### dasatinib
#### (non−c.toxic KI)

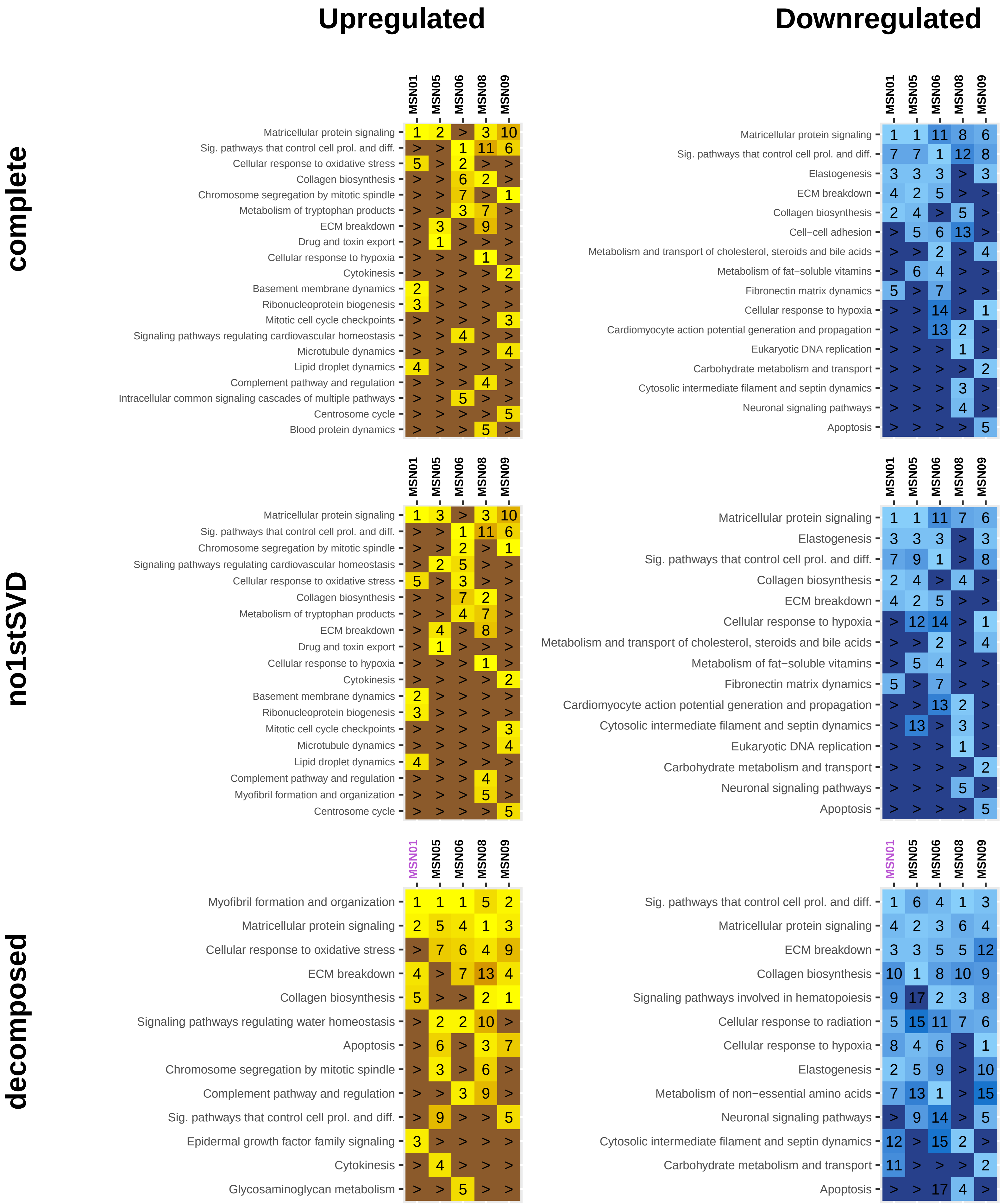

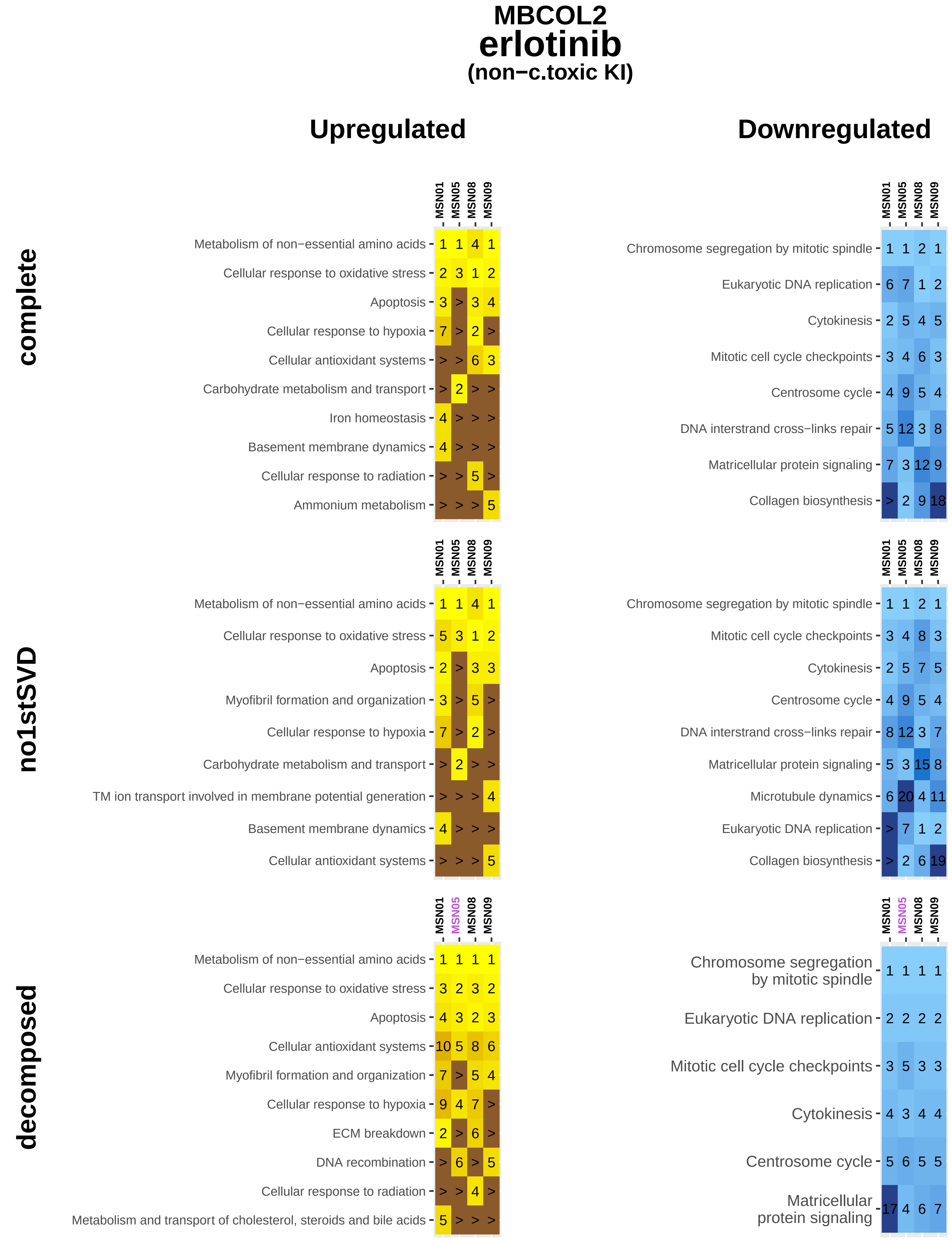
MBCOL2
erlotinib
(non−c.toxic KI)
Upregulated
Downregulated
complete
no1stSVD
decomposed
MSN01
MSN05
MSN08
MSN09
Metabolism of non−essential amino acids
Cellular response to oxidative stress
Apoptosis
Cellular response to hypoxia
Cellular antioxidant systems
Carbohydrate metabolism and transport
Iron homeostasis
Basement membrane dynamics
Cellular response to radiation
Ammonium metabolism
Chromosome segregation by mitotic spindle
Eukaryotic DNA replication
Cytokinesis
Mitotic cell cycle checkpoints
Centrosome cycle
DNA interstrand cross−links repair
Matricellular protein signaling
Collagen biosynthesis
Myofibril formation and organization
TM ion transport involved in membrane potential generation
Microtubule dynamics
ECM breakdown
DNA recombination
Metabolism and transport of cholesterol, steroids and bile acids

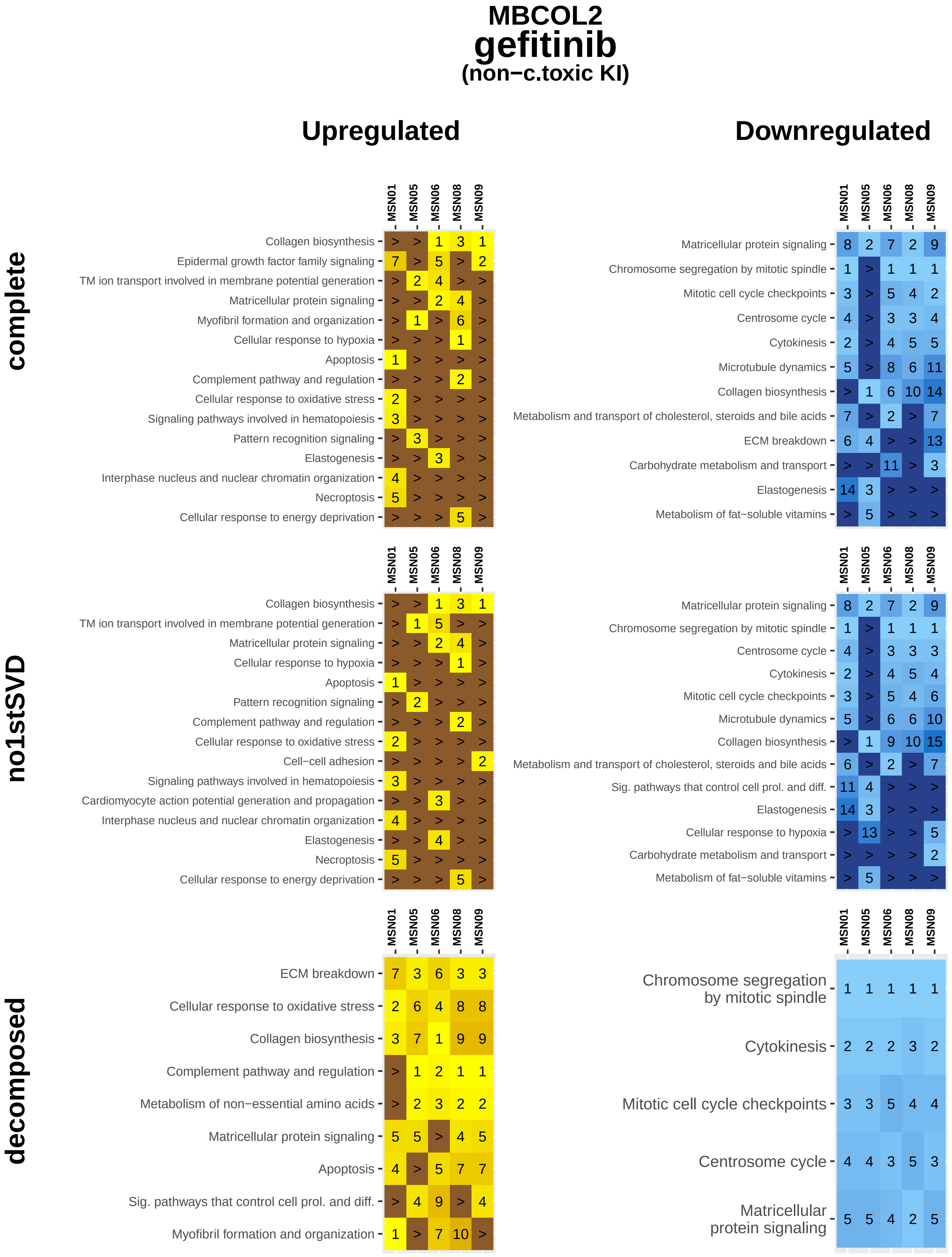
MBCOL2
gefitinib
(non−c.toxic KI)
Upregulated
Downregulated
complete
no1stSVD
decomposed
MSN01
MSN05
MSN06
MSN08
MSN09
Collagen biosynthesis
Epidermal growth factor family signaling
TM ion transport involved in membrane potential generation
Matricellular protein signaling
Myofibril formation and organization
Cellular response to hypoxia
Apoptosis
Complement pathway and regulation
Cellular response to oxidative stress
Signaling pathways involved in hematopoiesis
Pattern recognition signaling
Elastogenesis
Interphase nucleus and nuclear chromatin organization
Necroptosis
Cellular response to energy deprivation
Chromosome segregation by mitotic spindle
Mitotic cell cycle checkpoints
Centrosome cycle
Cytokinesis
Microtubule dynamics
Metabolism and transport of cholesterol, steroids and bile acids
ECM breakdown
Carbohydrate metabolism and transport
Metabolism of fat−soluble vitamins
Cell−cell adhesion
Cardiomyocyte action potential generation and propagation
Sig. pathways that control cell prol. and diff.
Metabolism of non−essential amino acids

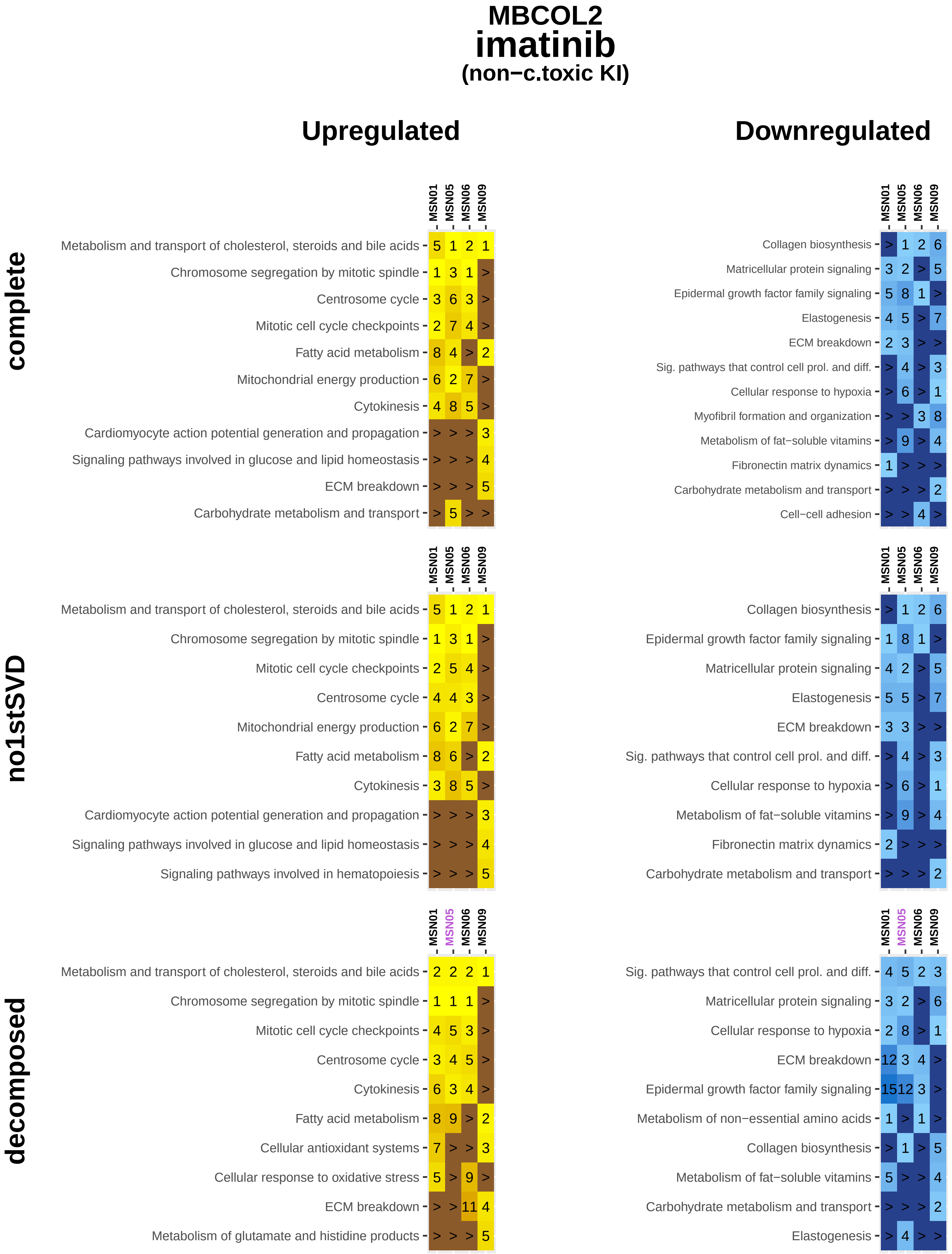

MBCOL2
imatinib
(non−c.toxic KI)
Upregulated
Downregulated
complete
MSN01 MSN05 MSN06 MSN09
Metabolism and transport of cholesterol, steroids and bile acids 5 1 2 1
Chromosome segregation by mitotic spindle 1 3 1 >
Centrosome cycle 3 6 3 >
Mitotic cell cycle checkpoints 2 7 4 >
Fatty acid metabolism 8 4 > 2
Mitochondrial energy production 6 2 7 >
Cytokinesis 4 8 5 >
Cardiomyocyte action potential generation and propagation > > > 3
Signaling pathways involved in glucose and lipid homeostasis > > > 4
ECM breakdown > > > 5
Carbohydrate metabolism and transport > 5 > >
Collagen biosynthesis > 1 2 6
Matricellular protein signaling 3 2 > 5
Epidermal growth factor family signaling 5 8 1 >
Elastogenesis 4 5 > 7
ECM breakdown 2 3 > >
Sig. pathways that control cell prol. and diff. > 4 > 3
Cellular response to hypoxia > 6 > 1
Myofibril formation and organization > > 3 8
Metabolism of fat−soluble vitamins > 9 > 4
Fibronectin matrix dynamics 1 > > >
Carbohydrate metabolism and transport > > > 2
Cell−cell adhesion > > 4 >
no1stSVD
Metabolism and transport of cholesterol, steroids and bile acids 5 1 2 1
Chromosome segregation by mitotic spindle 1 3 1 >
Mitotic cell cycle checkpoints 2 5 4 >
Centrosome cycle 4 4 3 >
Mitochondrial energy production 6 2 7 >
Fatty acid metabolism 8 6 > 2
Cytokinesis 3 8 5 >
Cardiomyocyte action potential generation and propagation > > > 3
Signaling pathways involved in glucose and lipid homeostasis > > > 4
Signaling pathways involved in hematopoiesis > > > 5
Collagen biosynthesis > 1 2 6
Epidermal growth factor family signaling 1 8 1 >
Matricellular protein signaling 4 2 > 5
Elastogenesis 5 5 > 7
ECM breakdown 3 3 > >
Sig. pathways that control cell prol. and diff. > 4 > 3
Cellular response to hypoxia > 6 > 1
Metabolism of fat−soluble vitamins > 9 > 4
Fibronectin matrix dynamics 2 > > >
Carbohydrate metabolism and transport > > > 2
decomposed
Metabolism and transport of cholesterol, steroids and bile acids 2 2 2 1
Chromosome segregation by mitotic spindle 1 1 1 >
Mitotic cell cycle checkpoints 4 5 3 >
Centrosome cycle 3 4 5 >
Cytokinesis 6 3 4 >
Fatty acid metabolism 8 9 > 2
Cellular antioxidant systems 7 > > 3
Cellular response to oxidative stress 5 > 9 >
ECM breakdown > > 11 4
Metabolism of glutamate and histidine products > > > 5
Sig. pathways that control cell prol. and diff. 4 5 2 3
Matricellular protein signaling 3 2 > 6
Cellular response to hypoxia 2 8 > 1
ECM breakdown 12 3 4 >
Epidermal growth factor family signaling 15 12 3 >
Metabolism of non−essential amino acids 1 > 1 >
Collagen biosynthesis > 1 > 5
Metabolism of fat−soluble vitamins 5 > > 4
Carbohydrate metabolism and transport > > > 2
Elastogenesis > 4 > >

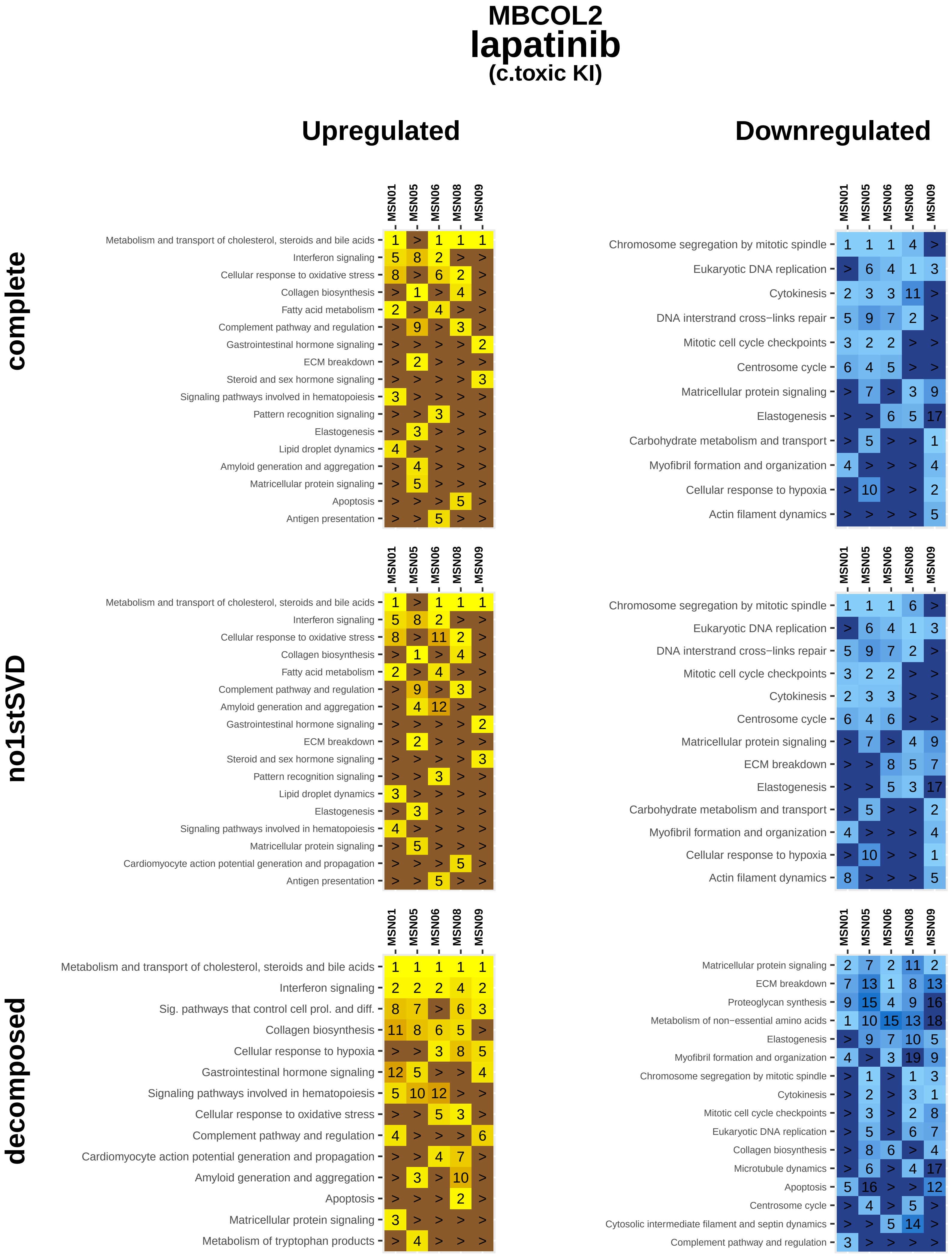

MBCOL2
lapatinib
(c.toxic KI)
Upregulated
Downregulated
complete
no1stSVD
decomposed
MSN01
MSN05
MSN06
MSN08
MSN09
Metabolism and transport of cholesterol, steroids and bile acids
Interferon signaling
Cellular response to oxidative stress
Collagen biosynthesis
Fatty acid metabolism
Complement pathway and regulation
Gastrointestinal hormone signaling
ECM breakdown
Steroid and sex hormone signaling
Signaling pathways involved in hematopoiesis
Pattern recognition signaling
Elastogenesis
Lipid droplet dynamics
Amyloid generation and aggregation
Matricellular protein signaling
Apoptosis
Antigen presentation
Chromosome segregation by mitotic spindle
Eukaryotic DNA replication
Cytokinesis
DNA interstrand cross−links repair
Mitotic cell cycle checkpoints
Centrosome cycle
Carbohydrate metabolism and transport
Myofibril formation and organization
Cellular response to hypoxia
Actin filament dynamics
Cardiomyocyte action potential generation and propagation
Sig. pathways that control cell prol. and diff.
Metabolism of tryptophan products
Proteoglycan synthesis
Metabolism of non−essential amino acids
Collagen biosynthesis
Microtubule dynamics
Cytosolic intermediate filament and septin dynamics

### MBCOL2
### nilotinib
#### (non−c.toxic KI)

##### complete

**Upregulated**

| | MSN01 | MSN05 | MSN06 | MSN08 | MSN09 |
|---|---|---|---|---|---|
| Chromosome segregation by mitotic spindle | 9 | 1 | 1 | 1 | 1 |
| Metabolism of non−essential amino acids | 5 | 2 | 5 | 3 | 5 |
| Centrosome cycle | 14 | 6 | 2 | 2 | 2 |
| Cytokinesis | > | 9 | 3 | 5 | 4 |
| Mitotic cell cycle checkpoints | 13 | > | 4 | 4 | 3 |
| Cellular response to hypoxia | 1 | > | > | > | > |
| Collagen biosynthesis | 2 | > | > | > | > |
| Rolling cell adhesion | > | 3 | > | > | > |
| Myofibril formation and organization | 3 | > | > | > | > |
| Metabolism of glutamate and histidine products | > | 4 | > | > | > |
| Elastogenesis | 4 | > | > | > | > |
| Metabolism and transport of cholesterol, steroids and bile acids | > | 5 | > | > | > |

**Downregulated**

| | MSN01 | MSN05 | MSN06 | MSN08 | MSN09 |
|---|---|---|---|---|---|
| Myofibril formation and organization | > | 2 | 1 | 1 | 1 |
| Neuronal action potential generation and propagation | > | 5 | 3 | 14 | 11 |
| Sig. pathways that control cell prol. and diff. | 2 | 8 | 5 | > | > |
| Actin filament dynamics | > | 9 | > | 2 | 5 |
| Cytosolic intermediate filament and septin dynamics | > | 10 | > | 4 | 4 |
| Amyloid generation and aggregation | 3 | 13 | > | 14 | > |
| Cellular response to hypoxia | > | 3 | > | > | 3 |
| Collagen biosynthesis | > | 1 | > | 6 | > |
| Signaling pathways involved in hematopoiesis | > | > | 7 | 3 | > |
| Cardiomyocyte action potential generation and propagation | 1 | > | > | 9 | > |
| Fibronectin matrix dynamics | > | > | > | 5 | 8 |
| Epidermal growth factor family signaling | > | > | 2 | 11 | > |
| Metabolism of fat−soluble vitamins | > | > | 4 | 12 | > |
| Carbohydrate metabolism and transport | > | > | > | > | 2 |
| Intracellular common signaling cascades of multiple pathways | 4 | > | > | > | > |
| Elastogenesis | > | 4 | > | > | > |

##### no1stSVD

**Upregulated**

| | MSN01 | MSN05 | MSN06 | MSN08 | MSN09 |
|---|---|---|---|---|---|
| Chromosome segregation by mitotic spindle | 9 | 1 | 1 | 1 | 1 |
| Metabolism of non−essential amino acids | 5 | 2 | 5 | 4 | 5 |
| Centrosome cycle | 14 | 6 | 2 | 2 | 2 |
| Mitotic cell cycle checkpoints | 13 | > | 3 | 3 | 3 |
| Cytokinesis | > | 9 | 4 | 5 | 4 |
| Myofibril formation and organization | 1 | > | > | > | > |
| Collagen biosynthesis | 2 | > | > | > | > |
| Rolling cell adhesion | > | 3 | > | > | > |
| Cellular response to hypoxia | 3 | > | > | > | > |
| Metabolism of glutamate and histidine products | > | 4 | > | > | > |
| Elastogenesis | 4 | > | > | > | > |
| Metabolism and transport of cholesterol, steroids and bile acids | > | 5 | > | > | > |

**Downregulated**

| | MSN01 | MSN05 | MSN06 | MSN08 | MSN09 |
|---|---|---|---|---|---|
| Myofibril formation and organization | > | 3 | 1 | 1 | 1 |
| Neuronal action potential generation and propagation | > | 5 | 2 | 10 | 11 |
| Sig. pathways that control cell prol. and diff. | 2 | 8 | 5 | > | > |
| Cytosolic intermediate filament and septin dynamics | > | 10 | > | 2 | 4 |
| Actin filament dynamics | > | 9 | > | 3 | 6 |
| Amyloid generation and aggregation | 3 | 13 | > | 6 | > |
| Collagen biosynthesis | > | 1 | > | 4 | > |
| Cellular response to hypoxia | > | 2 | > | > | 3 |
| Cardiomyocyte action potential generation and propagation | 1 | > | > | 9 | > |
| Fibronectin matrix dynamics | > | > | > | 5 | 7 |
| Metabolism of fat−soluble vitamins | > | > | 4 | 11 | > |
| Apoptosis | > | 11 | > | > | 5 |
| Carbohydrate metabolism and transport | > | > | > | > | 2 |
| Epidermal growth factor family signaling | > | > | 3 | > | > |
| Elastogenesis | > | 4 | > | > | > |
| Autophagy | 4 | > | > | > | > |
| Intracellular common signaling cascades of multiple pathways | 5 | > | > | > | > |

##### decomposed

**Upregulated**

| | MSN01 | MSN05 | MSN06 | MSN08 | MSN09 |
|---|---|---|---|---|---|
| Chromosome segregation by mitotic spindle | 1 | 1 | 1 | 1 | 1 |
| Cytokinesis | 2 | 2 | 3 | 4 | 2 |
| Mitotic cell cycle checkpoints | 3 | 3 | 4 | 3 | 3 |
| Centrosome cycle | 4 | 4 | 2 | 2 | 4 |
| Metabolism of non−essential amino acids | 7 | 5 | 5 | 5 | 5 |
| Cellular response to oxidative stress | 5 | 6 | 6 | 6 | 6 |

**Downregulated**

| | MSN01 | MSN05 | MSN06 | MSN08 | MSN09 |
|---|---|---|---|---|---|
| Collagen biosynthesis | 7 | 1 | 2 | 6 | 6 |
| Sig. pathways that control cell prol. and diff. | 3 | 9 | 11 | 7 | 4 |
| Myofibril formation and organization | > | 2 | 1 | 1 | 1 |
| ECM breakdown | 2 | 11 | > | 4 | 3 |
| Cardiomyocyte action potential generation and propagation | > | 6 | 5 | 8 | 2 |
| Signaling pathways involved in hematopoiesis | 6 | 12 | 3 | 2 | > |
| Amyloid generation and aggregation | 8 | 4 | 8 | > | 12 |
| Phase II biotransformation | > | 7 | 4 | > | 8 |
| Matricellular protein signaling | 1 | > | > | 3 | > |
| Metabolism of fat−soluble vitamins | > | 3 | > | 5 | > |
| Elastogenesis | 4 | 10 | > | > | > |
| Steroid hormone metabolism | > | 5 | > | > | > |
| Signaling by extracellular matrix components | 5 | > | > | > | > |
| Cellular response to hypoxia | > | > | > | > | 5 |

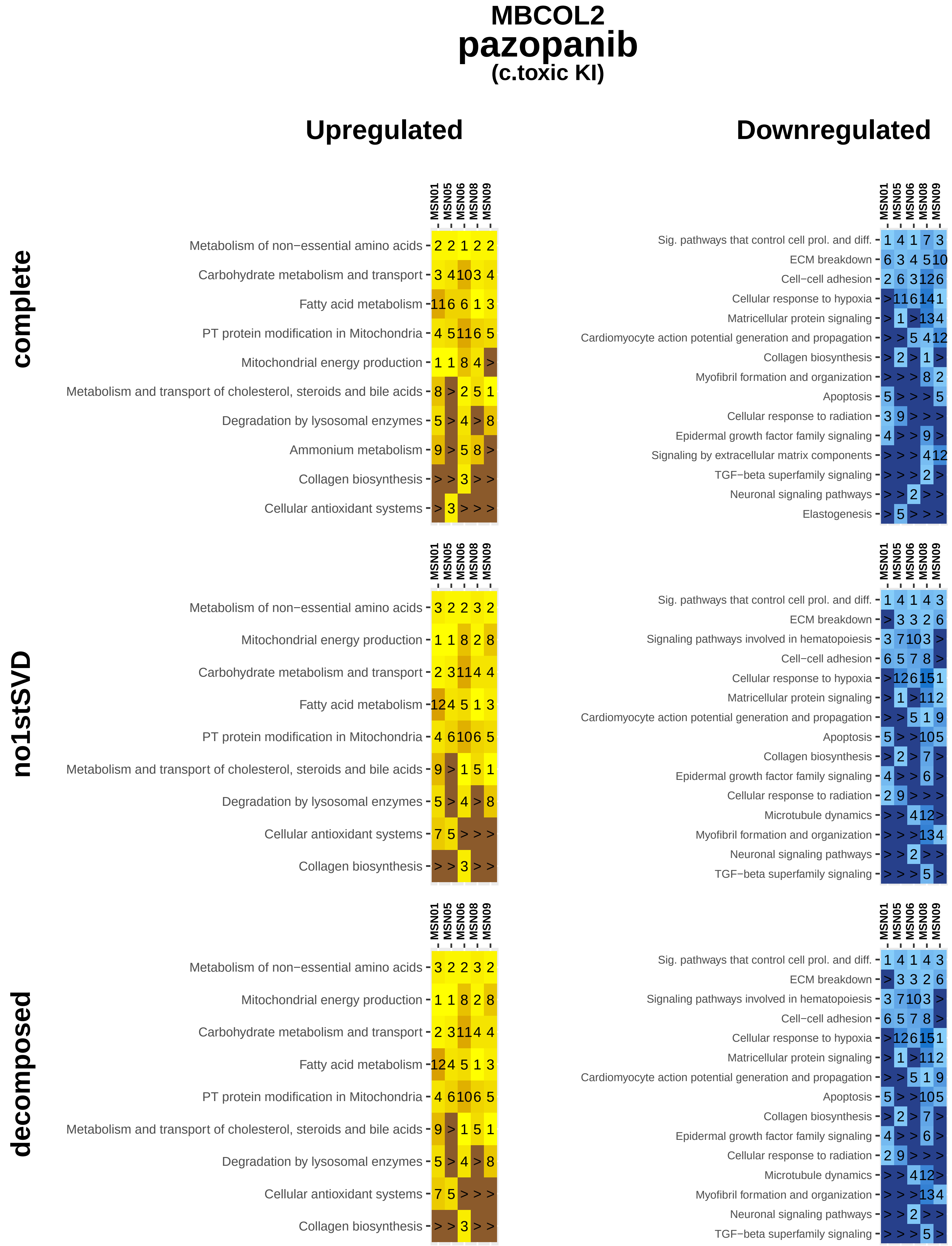
MBCOL2
pazopanib
(c.toxic KI)
Upregulated
Downregulated
complete
no1stSVD
decomposed
MSN01
MSN05
MSN06
MSN08
MSN09
Metabolism of non−essential amino acids
Carbohydrate metabolism and transport
Fatty acid metabolism
PT protein modification in Mitochondria
Mitochondrial energy production
Metabolism and transport of cholesterol, steroids and bile acids
Degradation by lysosomal enzymes
Ammonium metabolism
Collagen biosynthesis
Cellular antioxidant systems
Sig. pathways that control cell prol. and diff.
ECM breakdown
Cell−cell adhesion
Cellular response to hypoxia
Matricellular protein signaling
Cardiomyocyte action potential generation and propagation
Collagen biosynthesis
Myofibril formation and organization
Apoptosis
Cellular response to radiation
Epidermal growth factor family signaling
Signaling by extracellular matrix components
TGF−beta superfamily signaling
Neuronal signaling pathways
Elastogenesis
Signaling pathways involved in hematopoiesis
Microtubule dynamics

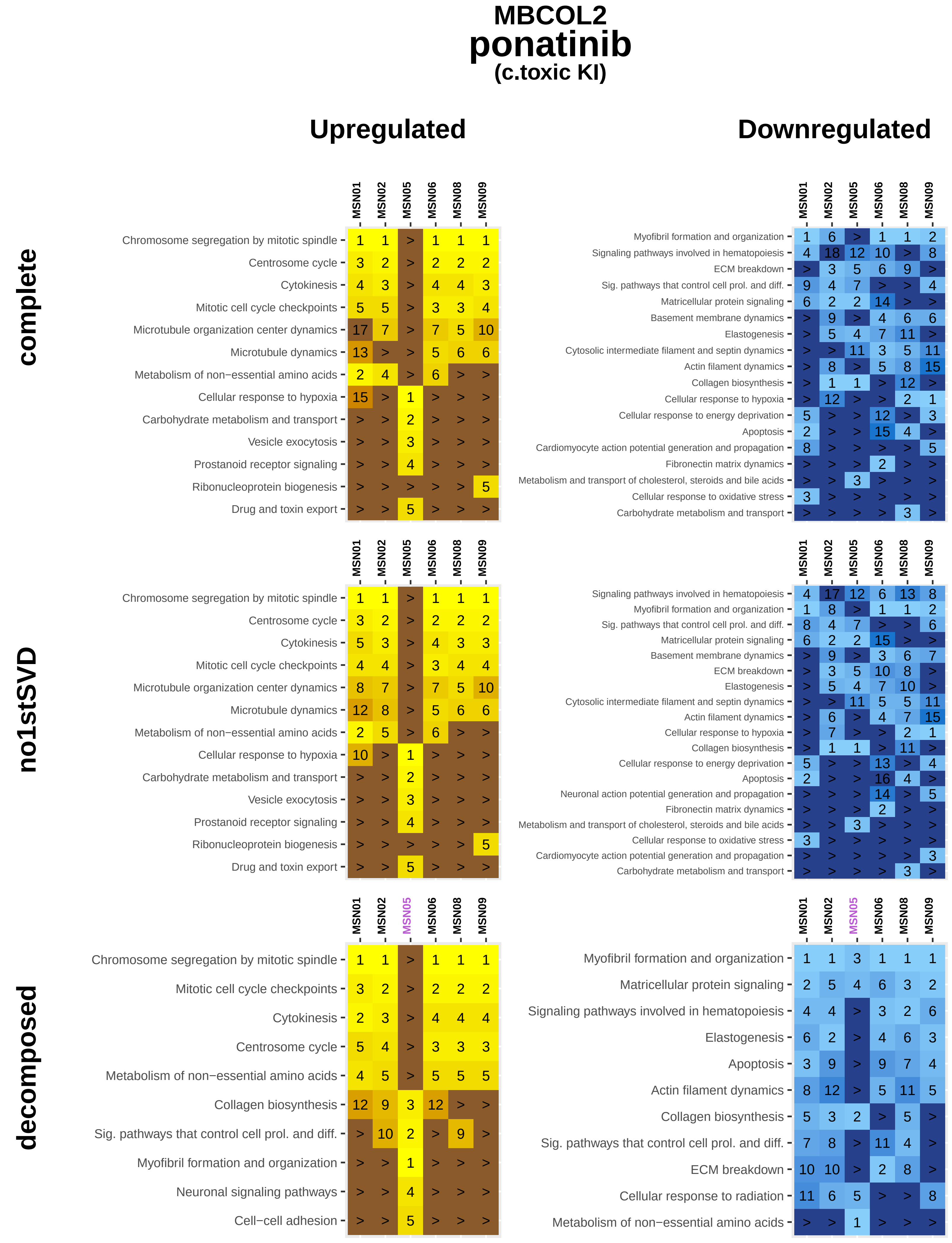
MBCOL2
ponatinib
(c.toxic KI)
Upregulated
Downregulated
complete
no1stSVD
decomposed
MSN01 MSN02 MSN05 MSN06 MSN08 MSN09
Chromosome segregation by mitotic spindle 1 1 > 1 1 1
Centrosome cycle 3 2 > 2 2 2
Cytokinesis 4 3 > 4 4 3
Mitotic cell cycle checkpoints 5 5 > 3 3 4
Microtubule organization center dynamics 17 7 > 7 5 10
Microtubule dynamics 13 > > 5 6 6
Metabolism of non−essential amino acids 2 4 > 6 > >
Cellular response to hypoxia 15 > 1 > > >
Carbohydrate metabolism and transport > > 2 > > >
Vesicle exocytosis > > 3 > > >
Prostanoid receptor signaling > > 4 > > >
Ribonucleoprotein biogenesis > > > > > 5
Drug and toxin export > > 5 > > >
MSN01 MSN02 MSN05 MSN06 MSN08 MSN09
Myofibril formation and organization 1 6 > 1 1 2
Signaling pathways involved in hematopoiesis 4 18 12 10 > 8
ECM breakdown > 3 5 6 9 >
Sig. pathways that control cell prol. and diff. 9 4 7 > > 4
Matricellular protein signaling 6 2 2 14 > >
Basement membrane dynamics > 9 > 4 6 6
Elastogenesis > 5 4 7 11 >
Cytosolic intermediate filament and septin dynamics > > 11 3 5 11
Actin filament dynamics > 8 > 5 8 15
Collagen biosynthesis > 1 1 > 12 >
Cellular response to hypoxia > 12 > > 2 1
Cellular response to energy deprivation 5 > > 12 > 3
Apoptosis 2 > > 15 4 >
Cardiomyocyte action potential generation and propagation 8 > > > > 5
Fibronectin matrix dynamics > > > 2 > >
Metabolism and transport of cholesterol, steroids and bile acids > > 3 > > >
Cellular response to oxidative stress 3 > > > > >
Carbohydrate metabolism and transport > > > > 3 >
MSN01 MSN02 MSN05 MSN06 MSN08 MSN09
Chromosome segregation by mitotic spindle 1 1 > 1 1 1
Centrosome cycle 3 2 > 2 2 2
Cytokinesis 5 3 > 4 3 3
Mitotic cell cycle checkpoints 4 4 > 3 4 4
Microtubule organization center dynamics 8 7 > 7 5 10
Microtubule dynamics 12 8 > 5 6 6
Metabolism of non−essential amino acids 2 5 > 6 > >
Cellular response to hypoxia 10 > 1 > > >
Carbohydrate metabolism and transport > > 2 > > >
Vesicle exocytosis > > 3 > > >
Prostanoid receptor signaling > > 4 > > >
Ribonucleoprotein biogenesis > > > > > 5
Drug and toxin export > > 5 > > >
MSN01 MSN02 MSN05 MSN06 MSN08 MSN09
Signaling pathways involved in hematopoiesis 4 17 12 6 13 8
Myofibril formation and organization 1 8 > 1 1 2
Sig. pathways that control cell prol. and diff. 8 4 7 > > 6
Matricellular protein signaling 6 2 2 15 > >
Basement membrane dynamics > 9 > 3 6 7
ECM breakdown > 3 5 10 8 >
Elastogenesis > 5 4 7 10 >
Cytosolic intermediate filament and septin dynamics > > 11 5 5 11
Actin filament dynamics > 6 > 4 7 15
Cellular response to hypoxia > 7 > > 2 1
Collagen biosynthesis > 1 1 > 11 >
Cellular response to energy deprivation 5 > > 13 > 4
Apoptosis 2 > > 16 4 >
Neuronal action potential generation and propagation > > > 14 > 5
Fibronectin matrix dynamics > > > 2 > >
Metabolism and transport of cholesterol, steroids and bile acids > > 3 > > >
Cellular response to oxidative stress 3 > > > > >
Cardiomyocyte action potential generation and propagation > > > > > 3
Carbohydrate metabolism and transport > > > > 3 >
MSN01 MSN02 MSN05 MSN06 MSN08 MSN09
Chromosome segregation by mitotic spindle 1 1 > 1 1 1
Mitotic cell cycle checkpoints 3 2 > 2 2 2
Cytokinesis 2 3 > 4 4 4
Centrosome cycle 5 4 > 3 3 3
Metabolism of non−essential amino acids 4 5 > 5 5 5
Collagen biosynthesis 12 9 3 12 > >
Sig. pathways that control cell prol. and diff. > 10 2 > 9 >
Myofibril formation and organization > > 1 > > >
Neuronal signaling pathways > > 4 > > >
Cell−cell adhesion > > 5 > > >
MSN01 MSN02 MSN05 MSN06 MSN08 MSN09
Myofibril formation and organization 1 1 3 1 1 1
Matricellular protein signaling 2 5 4 6 3 2
Signaling pathways involved in hematopoiesis 4 4 > 3 2 6
Elastogenesis 6 2 > 4 6 3
Apoptosis 3 9 > 9 7 4
Actin filament dynamics 8 12 > 5 11 5
Collagen biosynthesis 5 3 2 > 5 >
Sig. pathways that control cell prol. and diff. 7 8 > 11 4 >
ECM breakdown 10 10 > 2 8 >
Cellular response to radiation 11 6 5 > > 8
Metabolism of non−essential amino acids > > 1 > > >

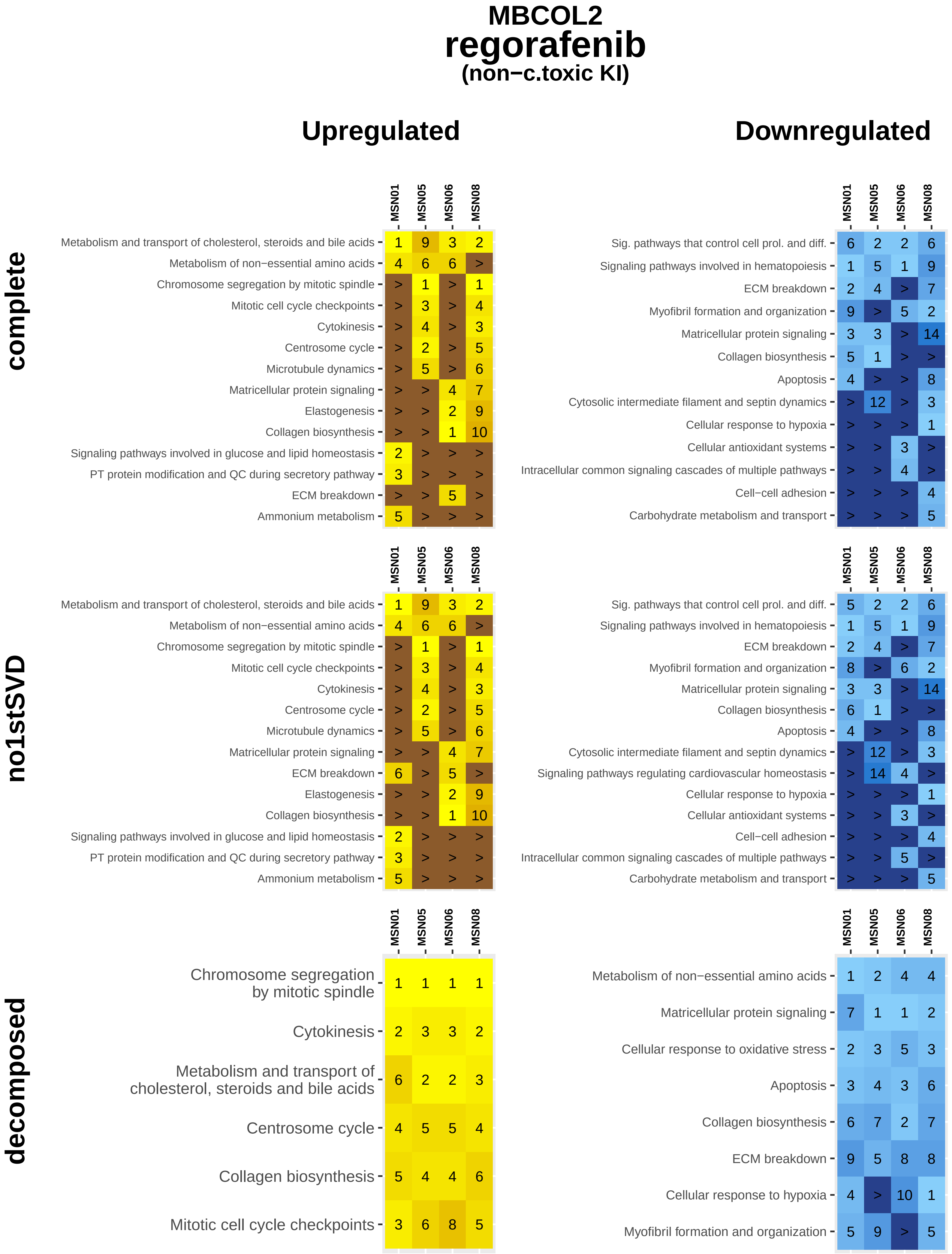
MBCOL2
regorafenib
(non−c.toxic KI)
Upregulated
Downregulated
complete
MSN01 MSN05 MSN06 MSN08
Metabolism and transport of cholesterol, steroids and bile acids 1 9 3 2
Metabolism of non−essential amino acids 4 6 6 >
Chromosome segregation by mitotic spindle > 1 > 1
Mitotic cell cycle checkpoints > 3 > 4
Cytokinesis > 4 > 3
Centrosome cycle > 2 > 5
Microtubule dynamics > 5 > 6
Matricellular protein signaling > > 4 7
Elastogenesis > > 2 9
Collagen biosynthesis > > 1 10
Signaling pathways involved in glucose and lipid homeostasis 2 > > >
PT protein modification and QC during secretory pathway 3 > > >
ECM breakdown > > 5 >
Ammonium metabolism 5 > > >
MSN01 MSN05 MSN06 MSN08
Sig. pathways that control cell prol. and diff. 6 2 2 6
Signaling pathways involved in hematopoiesis 1 5 1 9
ECM breakdown 2 4 > 7
Myofibril formation and organization 9 > 5 2
Matricellular protein signaling 3 3 > 14
Collagen biosynthesis 5 1 > >
Apoptosis 4 > > 8
Cytosolic intermediate filament and septin dynamics > 12 > 3
Cellular response to hypoxia > > > 1
Cellular antioxidant systems > > 3 >
Intracellular common signaling cascades of multiple pathways > > 4 >
Cell−cell adhesion > > > 4
Carbohydrate metabolism and transport > > > 5
no1stSVD
MSN01 MSN05 MSN06 MSN08
Metabolism and transport of cholesterol, steroids and bile acids 1 9 3 2
Metabolism of non−essential amino acids 4 6 6 >
Chromosome segregation by mitotic spindle > 1 > 1
Mitotic cell cycle checkpoints > 3 > 4
Cytokinesis > 4 > 3
Centrosome cycle > 2 > 5
Microtubule dynamics > 5 > 6
Matricellular protein signaling > > 4 7
ECM breakdown 6 > 5 >
Elastogenesis > > 2 9
Collagen biosynthesis > > 1 10
Signaling pathways involved in glucose and lipid homeostasis 2 > > >
PT protein modification and QC during secretory pathway 3 > > >
Ammonium metabolism 5 > > >
MSN01 MSN05 MSN06 MSN08
Sig. pathways that control cell prol. and diff. 5 2 2 6
Signaling pathways involved in hematopoiesis 1 5 1 9
ECM breakdown 2 4 > 7
Myofibril formation and organization 8 > 6 2
Matricellular protein signaling 3 3 > 14
Collagen biosynthesis 6 1 > >
Apoptosis 4 > > 8
Cytosolic intermediate filament and septin dynamics > 12 > 3
Signaling pathways regulating cardiovascular homeostasis > 14 4 >
Cellular response to hypoxia > > > 1
Cellular antioxidant systems > > 3 >
Cell−cell adhesion > > > 4
Intracellular common signaling cascades of multiple pathways > > 5 >
Carbohydrate metabolism and transport > > > 5
decomposed
MSN01 MSN05 MSN06 MSN08
Chromosome segregation by mitotic spindle 1 1 1 1
Cytokinesis 2 3 3 2
Metabolism and transport of cholesterol, steroids and bile acids 6 2 2 3
Centrosome cycle 4 5 5 4
Collagen biosynthesis 5 4 4 6
Mitotic cell cycle checkpoints 3 6 8 5
MSN01 MSN05 MSN06 MSN08
Metabolism of non−essential amino acids 1 2 4 4
Matricellular protein signaling 7 1 1 2
Cellular response to oxidative stress 2 3 5 3
Apoptosis 3 4 3 6
Collagen biosynthesis 6 7 2 7
ECM breakdown 9 5 8 8
Cellular response to hypoxia 4 > 10 1
Myofibril formation and organization 5 9 > 5

### MBCOL2
### ruxolitinib
#### (non−c.toxic KI)

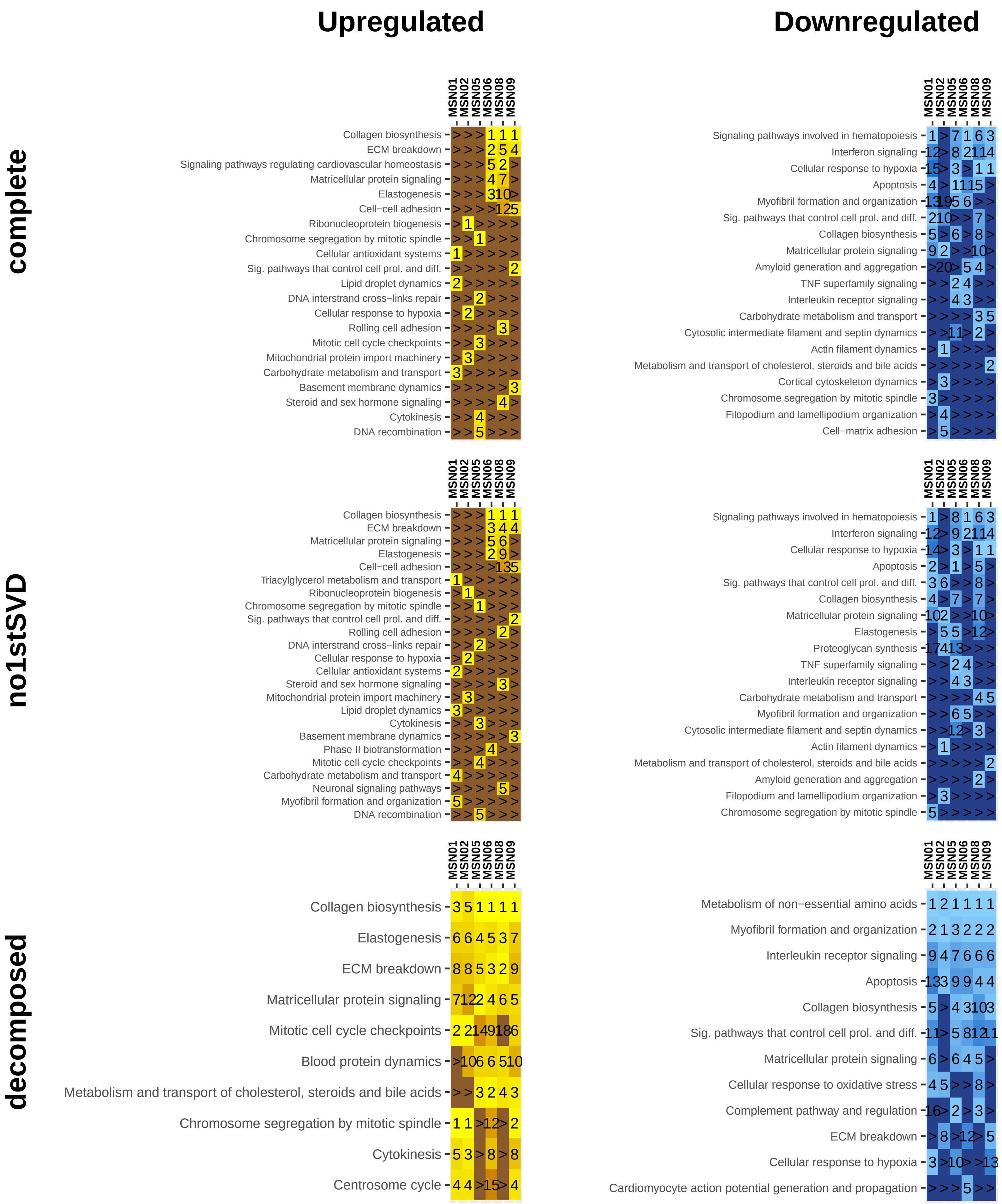

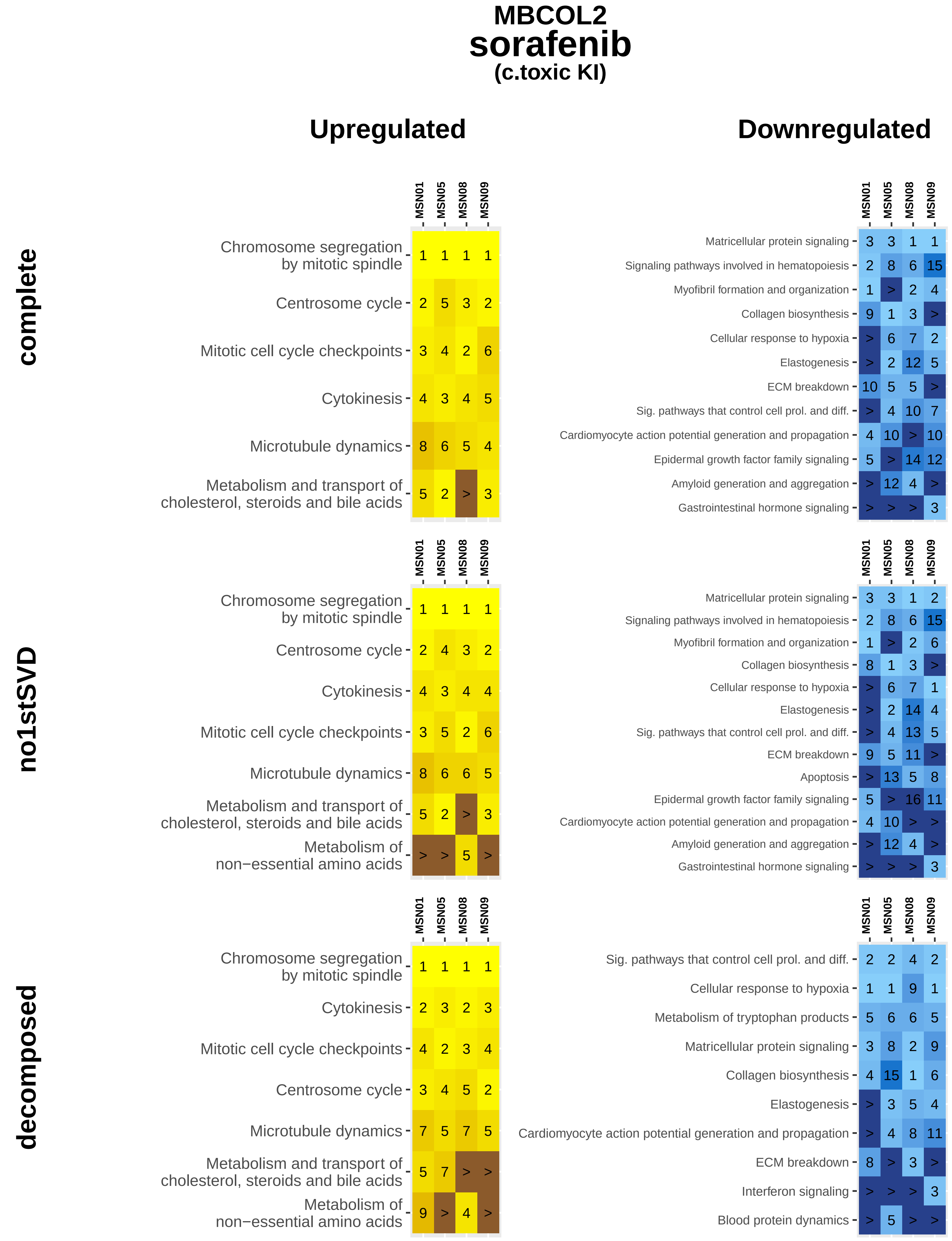
MBCOL2
sorafenib
(c.toxic KI)
Upregulated
Downregulated
complete
no1stSVD
decomposed
MSN01
MSN05
MSN08
MSN09
Chromosome segregation by mitotic spindle 1 1 1 1
Centrosome cycle 2 5 3 2
Mitotic cell cycle checkpoints 3 4 2 6
Cytokinesis 4 3 4 5
Microtubule dynamics 8 6 5 4
Metabolism and transport of cholesterol, steroids and bile acids 5 2 > 3
Matricellular protein signaling 3 3 1 1
Signaling pathways involved in hematopoiesis 2 8 6 15
Myofibril formation and organization 1 > 2 4
Collagen biosynthesis 9 1 3 >
Cellular response to hypoxia > 6 7 2
Elastogenesis > 2 12 5
ECM breakdown 10 5 5 >
Sig. pathways that control cell prol. and diff. > 4 10 7
Cardiomyocyte action potential generation and propagation 4 10 > 10
Epidermal growth factor family signaling 5 > 14 12
Amyloid generation and aggregation > 12 4 >
Gastrointestinal hormone signaling > > > 3
Chromosome segregation by mitotic spindle 1 1 1 1
Centrosome cycle 2 4 3 2
Cytokinesis 4 3 4 4
Mitotic cell cycle checkpoints 3 5 2 6
Microtubule dynamics 8 6 6 5
Metabolism and transport of cholesterol, steroids and bile acids 5 2 > 3
Metabolism of non−essential amino acids > > 5 >
Matricellular protein signaling 3 3 1 2
Signaling pathways involved in hematopoiesis 2 8 6 15
Myofibril formation and organization 1 > 2 6
Collagen biosynthesis 8 1 3 >
Cellular response to hypoxia > 6 7 1
Elastogenesis > 2 14 4
Sig. pathways that control cell prol. and diff. > 4 13 5
ECM breakdown 9 5 11 >
Apoptosis > 13 5 8
Epidermal growth factor family signaling 5 > 16 11
Cardiomyocyte action potential generation and propagation 4 10 > >
Amyloid generation and aggregation > 12 4 >
Gastrointestinal hormone signaling > > > 3
Chromosome segregation by mitotic spindle 1 1 1 1
Cytokinesis 2 3 2 3
Mitotic cell cycle checkpoints 4 2 3 4
Centrosome cycle 3 4 5 2
Microtubule dynamics 7 5 7 5
Metabolism and transport of cholesterol, steroids and bile acids 5 7 > >
Metabolism of non−essential amino acids 9 > 4 >
Sig. pathways that control cell prol. and diff. 2 2 4 2
Cellular response to hypoxia 1 1 9 1
Metabolism of tryptophan products 5 6 6 5
Matricellular protein signaling 3 8 2 9
Collagen biosynthesis 4 15 1 6
Elastogenesis > 3 5 4
Cardiomyocyte action potential generation and propagation > 4 8 11
ECM breakdown 8 > 3 >
Interferon signaling > > > 3
Blood protein dynamics > 5 > >

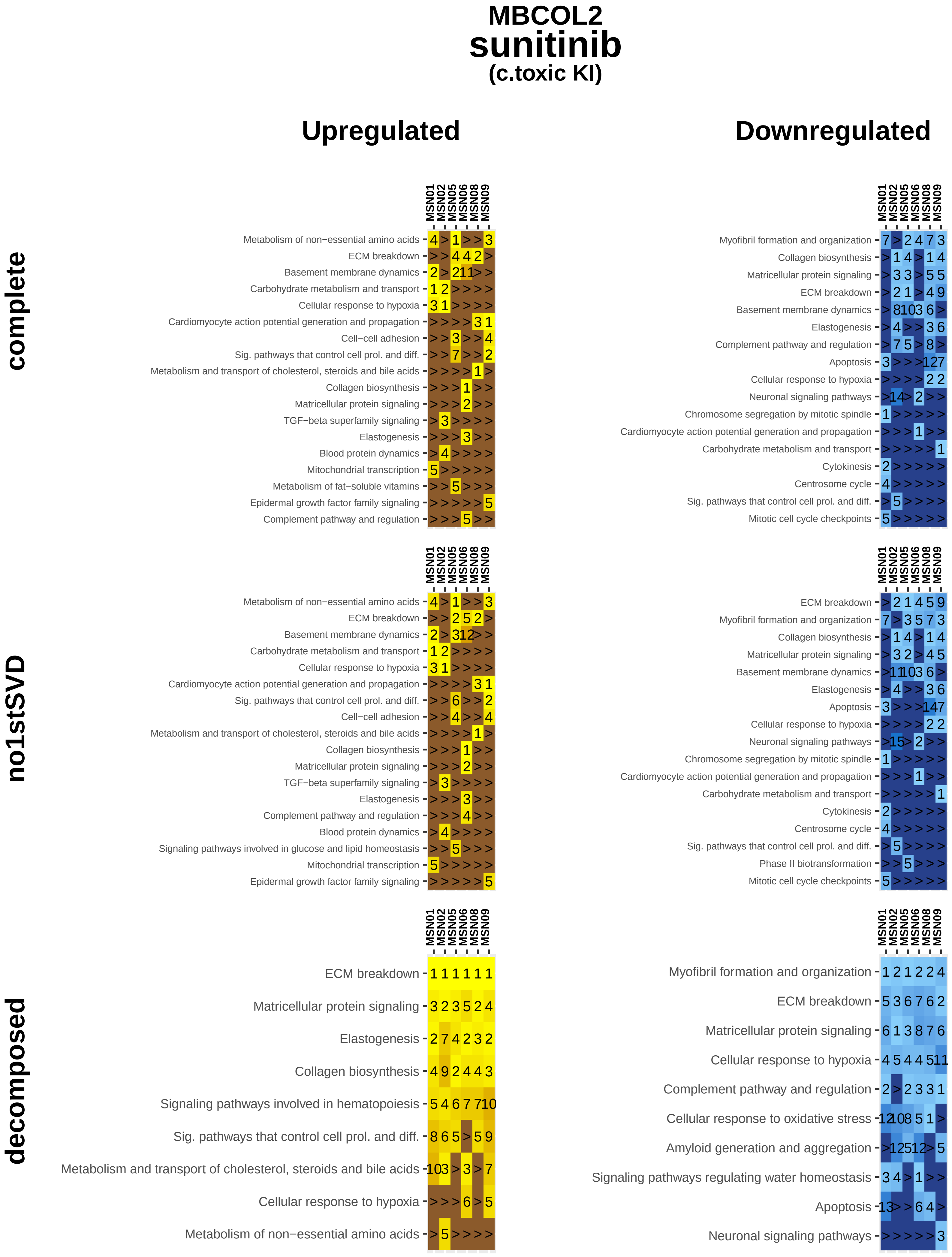
MBCOL2
sunitinib
(c.toxic KI)
Upregulated
Downregulated
complete
no1stSVD
decomposed
MSN01
MSN02
MSN05
MSN06
MSN08
MSN09
Metabolism of non−essential amino acids
ECM breakdown
Basement membrane dynamics
Carbohydrate metabolism and transport
Cellular response to hypoxia
Cardiomyocyte action potential generation and propagation
Cell−cell adhesion
Sig. pathways that control cell prol. and diff.
Metabolism and transport of cholesterol, steroids and bile acids
Collagen biosynthesis
Matricellular protein signaling
TGF−beta superfamily signaling
Elastogenesis
Blood protein dynamics
Mitochondrial transcription
Metabolism of fat−soluble vitamins
Epidermal growth factor family signaling
Complement pathway and regulation
Myofibril formation and organization
Apoptosis
Neuronal signaling pathways
Chromosome segregation by mitotic spindle
Cytokinesis
Centrosome cycle
Mitotic cell cycle checkpoints
Signaling pathways involved in glucose and lipid homeostasis
Phase II biotransformation
Signaling pathways involved in hematopoiesis
Cellular response to oxidative stress
Amyloid generation and aggregation
Signaling pathways regulating water homeostasis

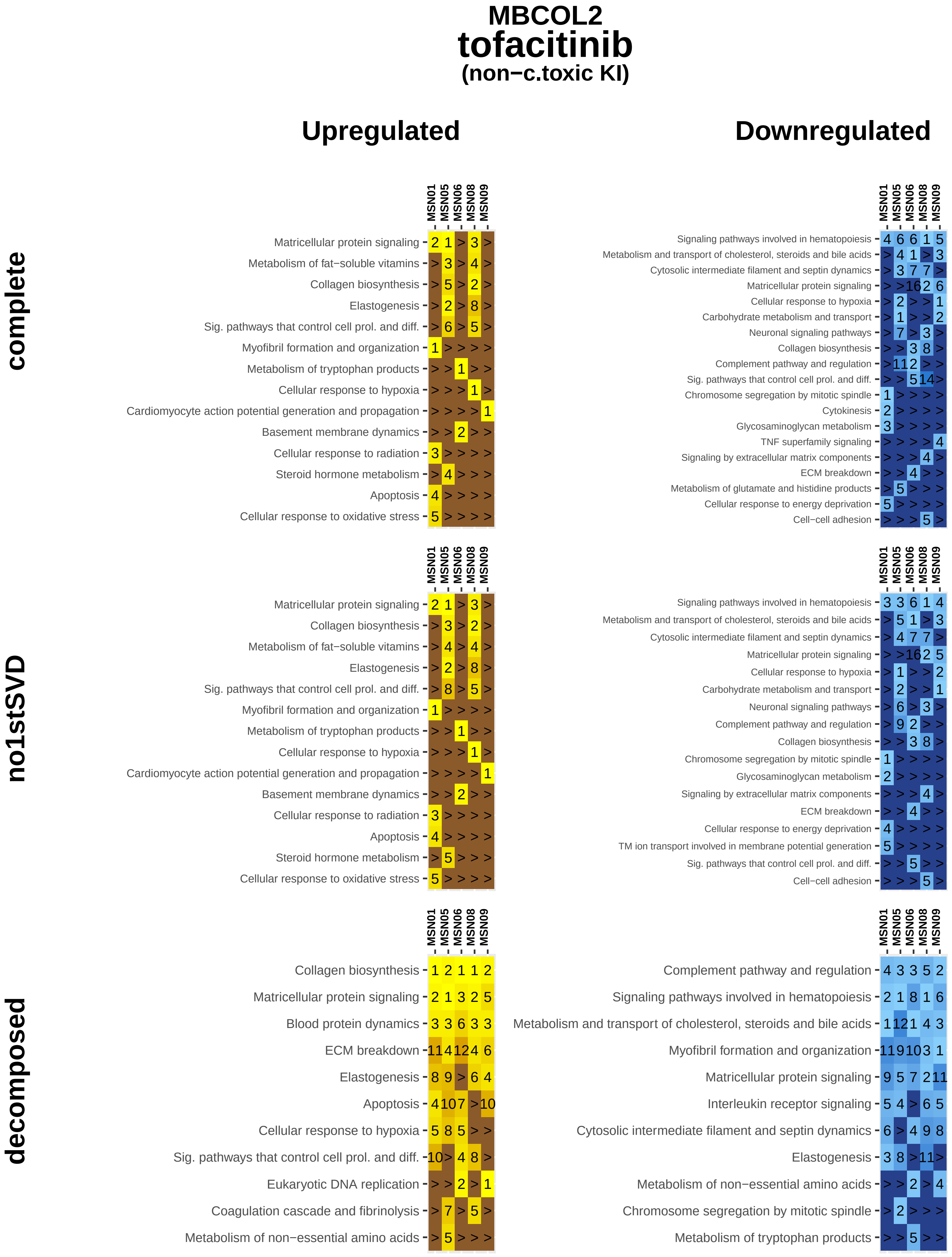
MBCOL2
tofacitinib
(non−c.toxic KI)
Upregulated
Downregulated
complete
no1stSVD
decomposed
MSN01
MSN05
MSN06
MSN08
MSN09
Matricellular protein signaling
Metabolism of fat−soluble vitamins
Collagen biosynthesis
Elastogenesis
Sig. pathways that control cell prol. and diff.
Myofibril formation and organization
Metabolism of tryptophan products
Cellular response to hypoxia
Cardiomyocyte action potential generation and propagation
Basement membrane dynamics
Cellular response to radiation
Steroid hormone metabolism
Apoptosis
Cellular response to oxidative stress
Signaling pathways involved in hematopoiesis
Metabolism and transport of cholesterol, steroids and bile acids
Cytosolic intermediate filament and septin dynamics
Carbohydrate metabolism and transport
Neuronal signaling pathways
Complement pathway and regulation
Chromosome segregation by mitotic spindle
Cytokinesis
Glycosaminoglycan metabolism
TNF superfamily signaling
Signaling by extracellular matrix components
ECM breakdown
Metabolism of glutamate and histidine products
Cellular response to energy deprivation
Cell−cell adhesion
TM ion transport involved in membrane potential generation
Blood protein dynamics
Eukaryotic DNA replication
Coagulation cascade and fibrinolysis
Metabolism of non−essential amino acids
Interleukin receptor signaling

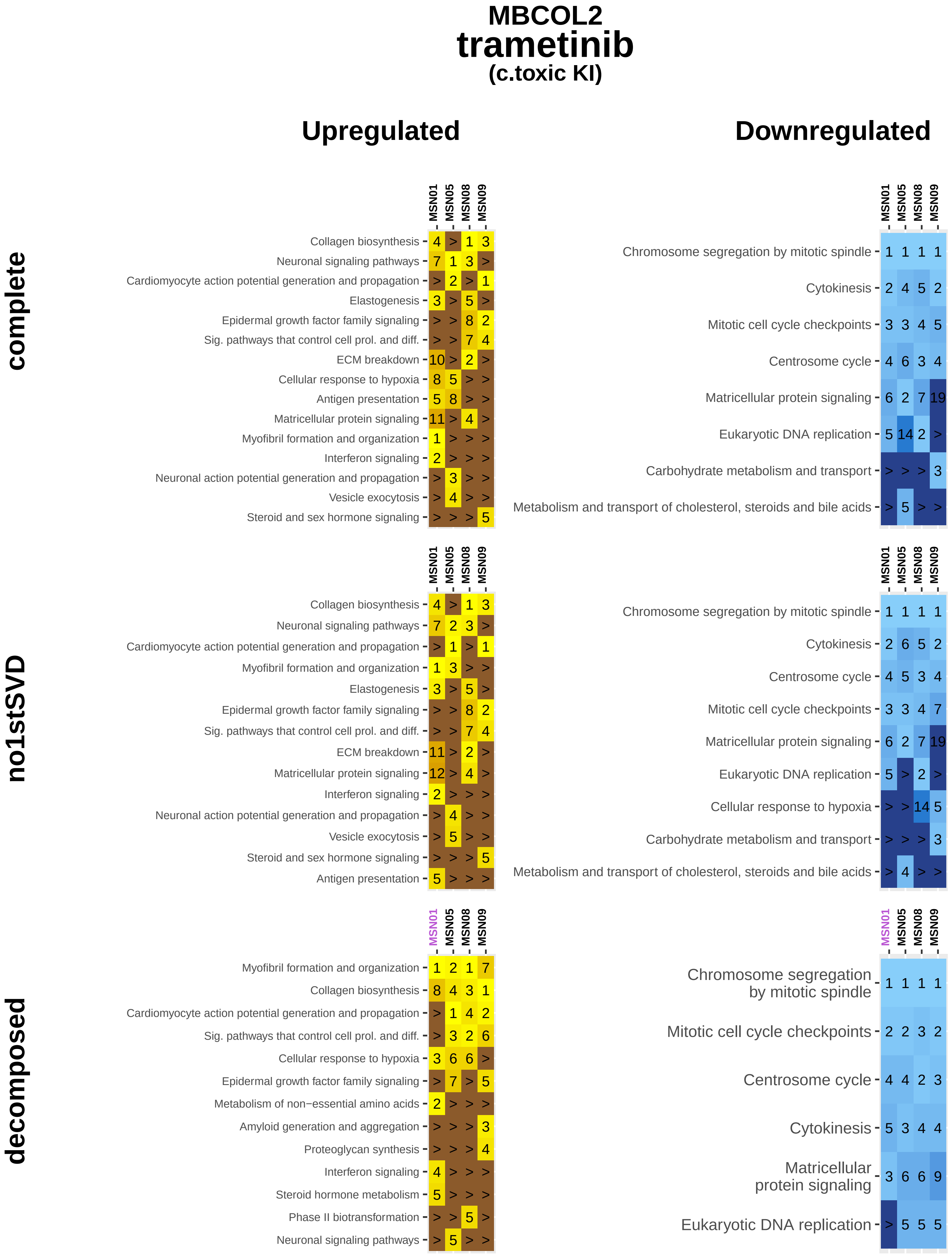
MBCOL2
trametinib
(c.toxic KI)
Upregulated
Downregulated
complete
no1stSVD
decomposed
MSN01 MSN05 MSN08 MSN09
Collagen biosynthesis 4 > 1 3
Neuronal signaling pathways 7 1 3 >
Cardiomyocyte action potential generation and propagation > 2 > 1
Elastogenesis 3 > 5 >
Epidermal growth factor family signaling > > 8 2
Sig. pathways that control cell prol. and diff. > > 7 4
ECM breakdown 10 > 2 >
Cellular response to hypoxia 8 5 > >
Antigen presentation 5 8 > >
Matricellular protein signaling 11 > 4 >
Myofibril formation and organization 1 > > >
Interferon signaling 2 > > >
Neuronal action potential generation and propagation > 3 > >
Vesicle exocytosis > 4 > >
Steroid and sex hormone signaling > > > 5
Chromosome segregation by mitotic spindle 1 1 1 1
Cytokinesis 2 4 5 2
Mitotic cell cycle checkpoints 3 3 4 5
Centrosome cycle 4 6 3 4
Matricellular protein signaling 6 2 7 19
Eukaryotic DNA replication 5 14 2 >
Carbohydrate metabolism and transport > > > 3
Metabolism and transport of cholesterol, steroids and bile acids > 5 > >
Collagen biosynthesis 4 > 1 3
Neuronal signaling pathways 7 2 3 >
Cardiomyocyte action potential generation and propagation > 1 > 1
Myofibril formation and organization 1 3 > >
Elastogenesis 3 > 5 >
Epidermal growth factor family signaling > > 8 2
Sig. pathways that control cell prol. and diff. > > 7 4
ECM breakdown 11 > 2 >
Matricellular protein signaling 12 > 4 >
Interferon signaling 2 > > >
Neuronal action potential generation and propagation > 4 > >
Vesicle exocytosis > 5 > >
Steroid and sex hormone signaling > > > 5
Antigen presentation 5 > > >
Chromosome segregation by mitotic spindle 1 1 1 1
Cytokinesis 2 6 5 2
Centrosome cycle 4 5 3 4
Mitotic cell cycle checkpoints 3 3 4 7
Matricellular protein signaling 6 2 7 19
Eukaryotic DNA replication 5 > 2 >
Cellular response to hypoxia > > 14 5
Carbohydrate metabolism and transport > > > 3
Metabolism and transport of cholesterol, steroids and bile acids > 4 > >
Myofibril formation and organization 1 2 1 7
Collagen biosynthesis 8 4 3 1
Cardiomyocyte action potential generation and propagation > 1 4 2
Sig. pathways that control cell prol. and diff. > 3 2 6
Cellular response to hypoxia 3 6 6 >
Epidermal growth factor family signaling > 7 > 5
Metabolism of non−essential amino acids 2 > > >
Amyloid generation and aggregation > > > 3
Proteoglycan synthesis > > > 4
Interferon signaling 4 > > >
Steroid hormone metabolism 5 > > >
Phase II biotransformation > > 5 >
Neuronal signaling pathways > 5 > >
Chromosome segregation by mitotic spindle 1 1 1 1
Mitotic cell cycle checkpoints 2 2 3 2
Centrosome cycle 4 4 2 3
Cytokinesis 5 3 4 4
Matricellular protein signaling 3 6 6 9
Eukaryotic DNA replication > 5 5 5

### MBCOL2
### vandetanib
#### (c.toxic KI)

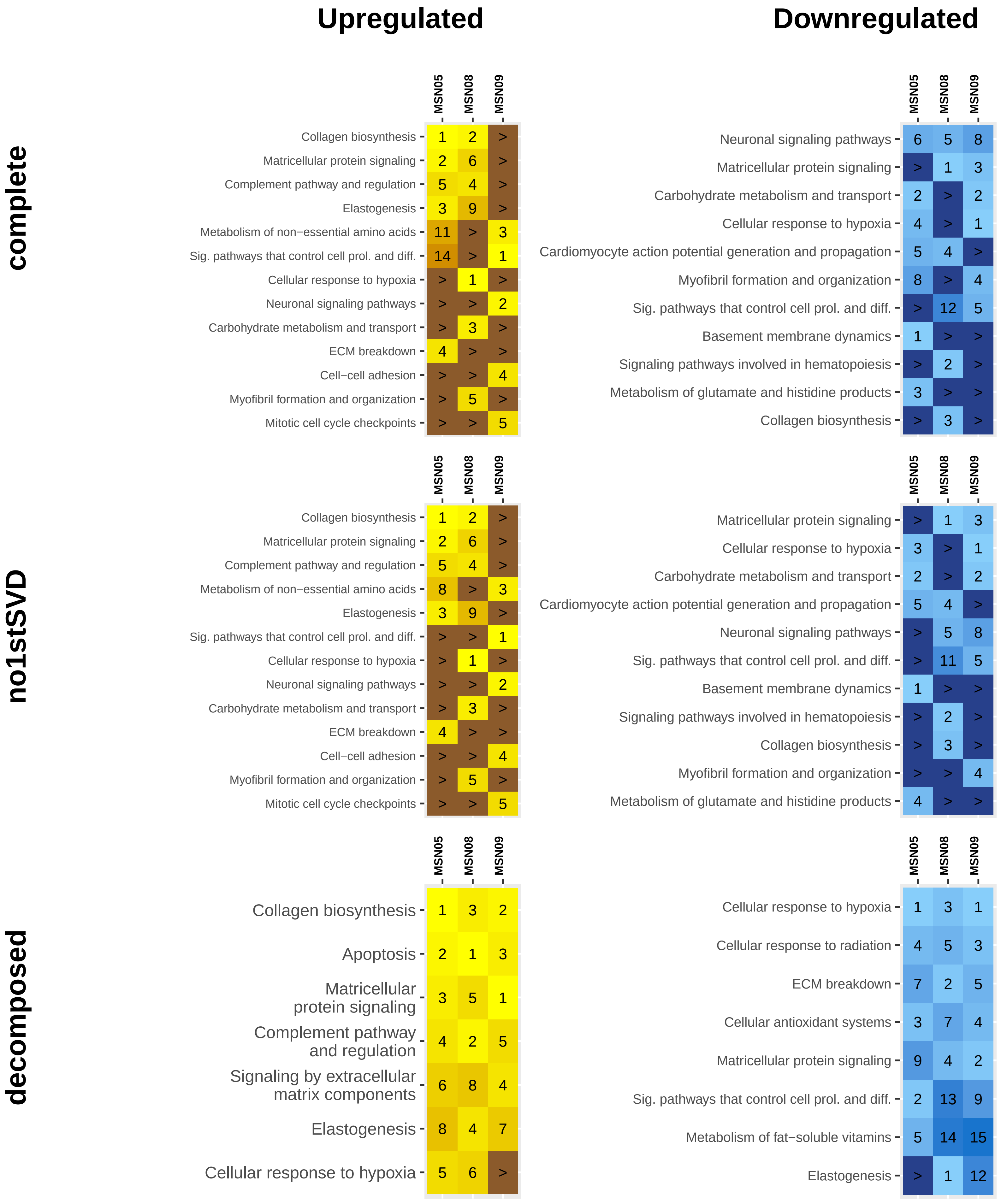

### MBCOL2
### vemurafenib
#### (non−c.toxic KI)

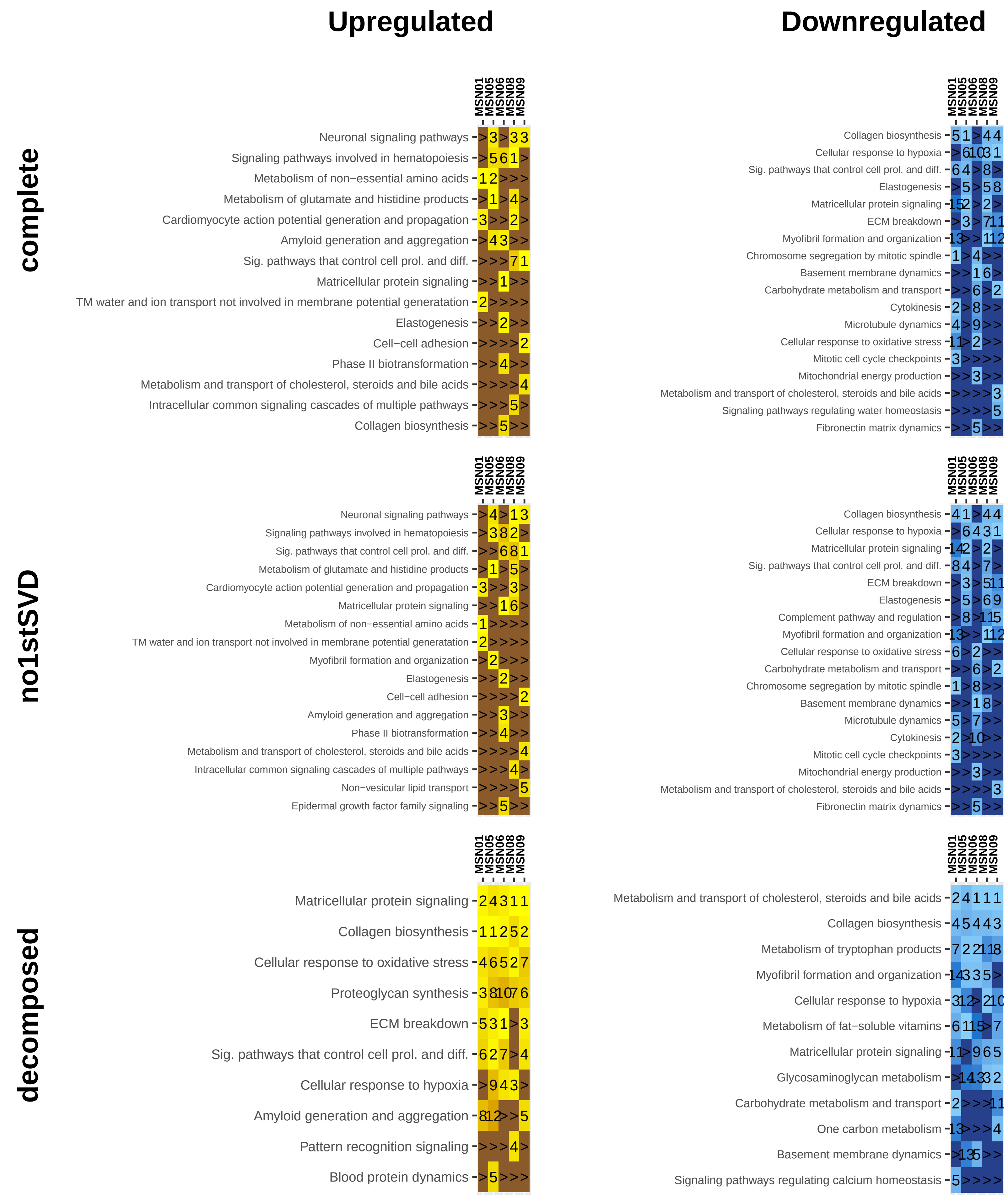

### MBCOL2
### bevacizumab
#### (c.toxic mAb)

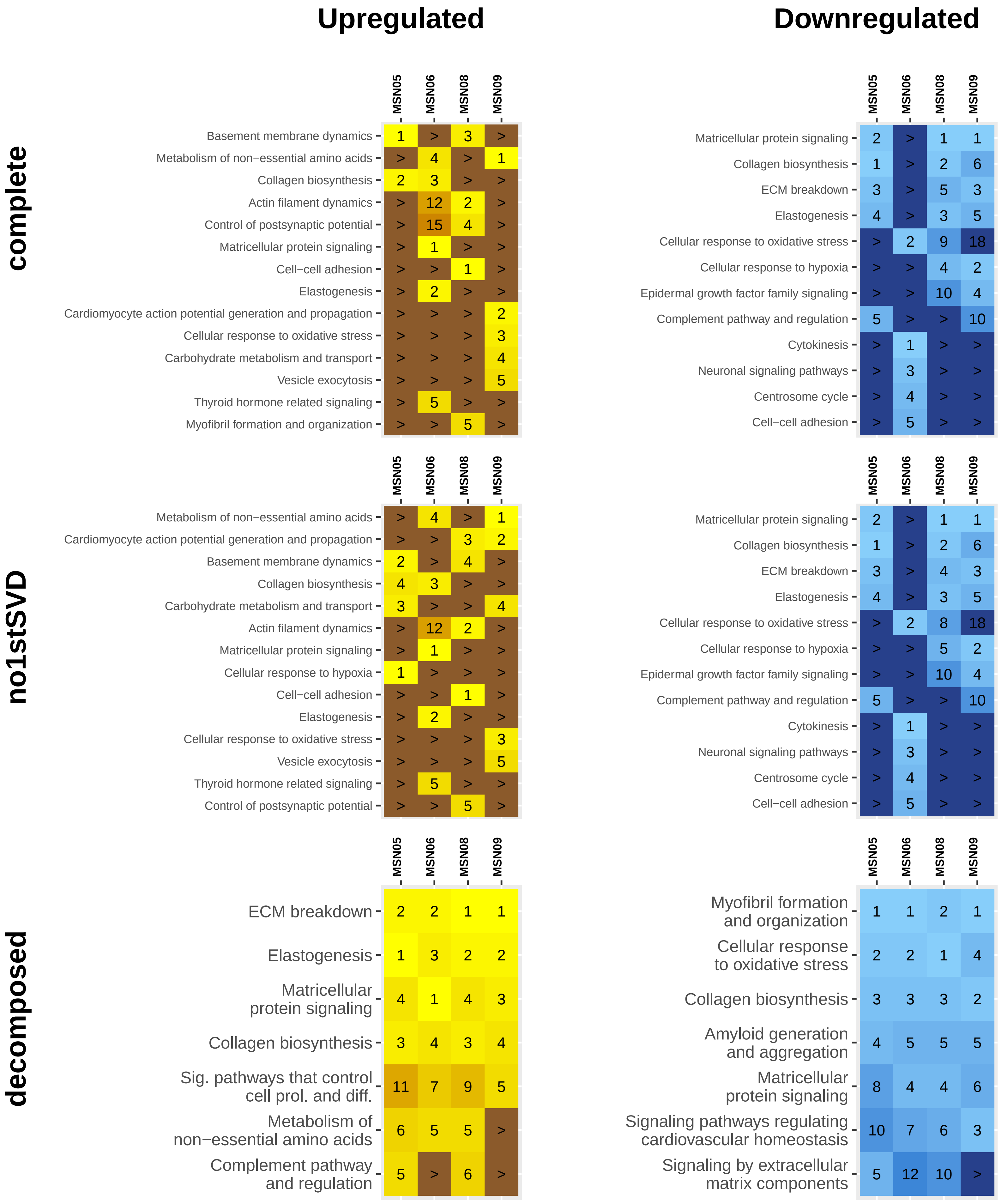

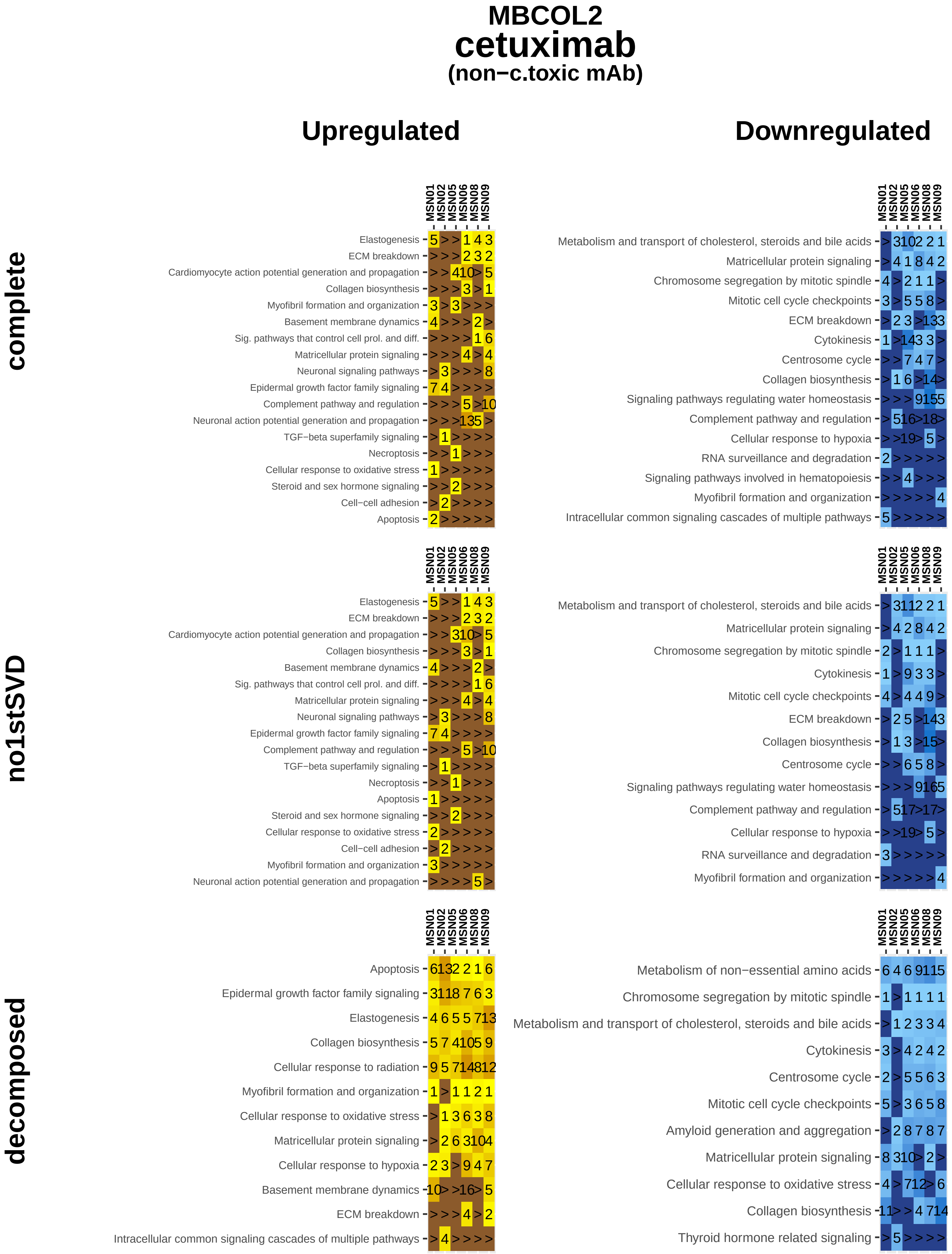

MBCOL2
cetuximab
(non−c.toxic mAb)
Upregulated
Downregulated
complete
no1stSVD
decomposed
MSN01
MSN02
MSN05
MSN06
MSN08
MSN09
Elastogenesis
ECM breakdown
Cardiomyocyte action potential generation and propagation
Collagen biosynthesis
Myofibril formation and organization
Basement membrane dynamics
Sig. pathways that control cell prol. and diff.
Matricellular protein signaling
Neuronal signaling pathways
Epidermal growth factor family signaling
Complement pathway and regulation
Neuronal action potential generation and propagation
TGF−beta superfamily signaling
Necroptosis
Cellular response to oxidative stress
Steroid and sex hormone signaling
Cell−cell adhesion
Apoptosis
Metabolism and transport of cholesterol, steroids and bile acids
Chromosome segregation by mitotic spindle
Mitotic cell cycle checkpoints
Cytokinesis
Centrosome cycle
Signaling pathways regulating water homeostasis
Cellular response to hypoxia
RNA surveillance and degradation
Signaling pathways involved in hematopoiesis
Intracellular common signaling cascades of multiple pathways
Cellular response to radiation
Metabolism of non−essential amino acids
Amyloid generation and aggregation
Thyroid hormone related signaling

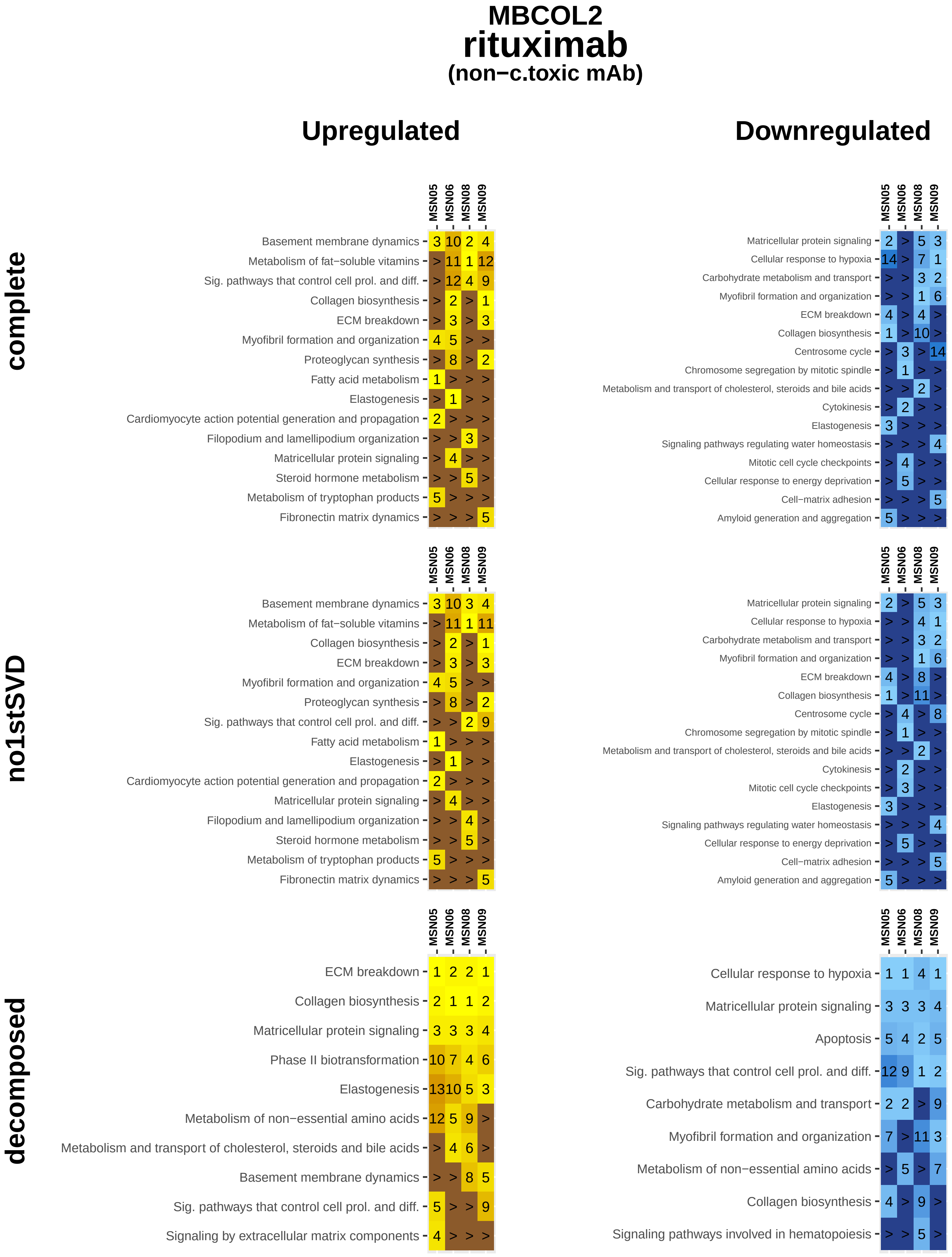

MBCOL2
rituximab
(non−c.toxic mAb)
Upregulated
Downregulated
complete
MSN05 MSN06 MSN08 MSN09
Basement membrane dynamics 3 10 2 4
Metabolism of fat−soluble vitamins > 11 1 12
Sig. pathways that control cell prol. and diff. > 12 4 9
Collagen biosynthesis > 2 > 1
ECM breakdown > 3 > 3
Myofibril formation and organization 4 5 > >
Proteoglycan synthesis > 8 > 2
Fatty acid metabolism 1 > > >
Elastogenesis > 1 > >
Cardiomyocyte action potential generation and propagation 2 > > >
Filopodium and lamellipodium organization > > 3 >
Matricellular protein signaling > 4 > >
Steroid hormone metabolism > > 5 >
Metabolism of tryptophan products 5 > > >
Fibronectin matrix dynamics > > > 5
MSN05 MSN06 MSN08 MSN09
Matricellular protein signaling 2 > 5 3
Cellular response to hypoxia 14 > 7 1
Carbohydrate metabolism and transport > > 3 2
Myofibril formation and organization > > 1 6
ECM breakdown 4 > 4 >
Collagen biosynthesis 1 > 10 >
Centrosome cycle > 3 > 14
Chromosome segregation by mitotic spindle > 1 > >
Metabolism and transport of cholesterol, steroids and bile acids > > 2 >
Cytokinesis > 2 > >
Elastogenesis 3 > > >
Signaling pathways regulating water homeostasis > > > 4
Mitotic cell cycle checkpoints > 4 > >
Cellular response to energy deprivation > 5 > >
Cell−matrix adhesion > > > 5
Amyloid generation and aggregation 5 > > >
no1stSVD
MSN05 MSN06 MSN08 MSN09
Basement membrane dynamics 3 10 3 4
Metabolism of fat−soluble vitamins > 11 1 11
Collagen biosynthesis > 2 > 1
ECM breakdown > 3 > 3
Myofibril formation and organization 4 5 > >
Proteoglycan synthesis > 8 > 2
Sig. pathways that control cell prol. and diff. > > 2 9
Fatty acid metabolism 1 > > >
Elastogenesis > 1 > >
Cardiomyocyte action potential generation and propagation 2 > > >
Matricellular protein signaling > 4 > >
Filopodium and lamellipodium organization > > 4 >
Steroid hormone metabolism > > 5 >
Metabolism of tryptophan products 5 > > >
Fibronectin matrix dynamics > > > 5
MSN05 MSN06 MSN08 MSN09
Matricellular protein signaling 2 > 5 3
Cellular response to hypoxia > > 4 1
Carbohydrate metabolism and transport > > 3 2
Myofibril formation and organization > > 1 6
ECM breakdown 4 > 8 >
Collagen biosynthesis 1 > 11 >
Centrosome cycle > 4 > 8
Chromosome segregation by mitotic spindle > 1 > >
Metabolism and transport of cholesterol, steroids and bile acids > > 2 >
Cytokinesis > 2 > >
Mitotic cell cycle checkpoints > 3 > >
Elastogenesis 3 > > >
Signaling pathways regulating water homeostasis > > > 4
Cellular response to energy deprivation > 5 > >
Cell−matrix adhesion > > > 5
Amyloid generation and aggregation 5 > > >
decomposed
MSN05 MSN06 MSN08 MSN09
ECM breakdown 1 2 2 1
Collagen biosynthesis 2 1 1 2
Matricellular protein signaling 3 3 3 4
Phase II biotransformation 10 7 4 6
Elastogenesis 13 10 5 3
Metabolism of non−essential amino acids 12 5 9 >
Metabolism and transport of cholesterol, steroids and bile acids > 4 6 >
Basement membrane dynamics > > 8 5
Sig. pathways that control cell prol. and diff. 5 > > 9
Signaling by extracellular matrix components 4 > > >
MSN05 MSN06 MSN08 MSN09
Cellular response to hypoxia 1 1 4 1
Matricellular protein signaling 3 3 3 4
Apoptosis 5 4 2 5
Sig. pathways that control cell prol. and diff. 12 9 1 2
Carbohydrate metabolism and transport 2 2 > 9
Myofibril formation and organization 7 > 11 3
Metabolism of non−essential amino acids > 5 > 7
Collagen biosynthesis 4 > 9 >
Signaling pathways involved in hematopoiesis > > 5 >

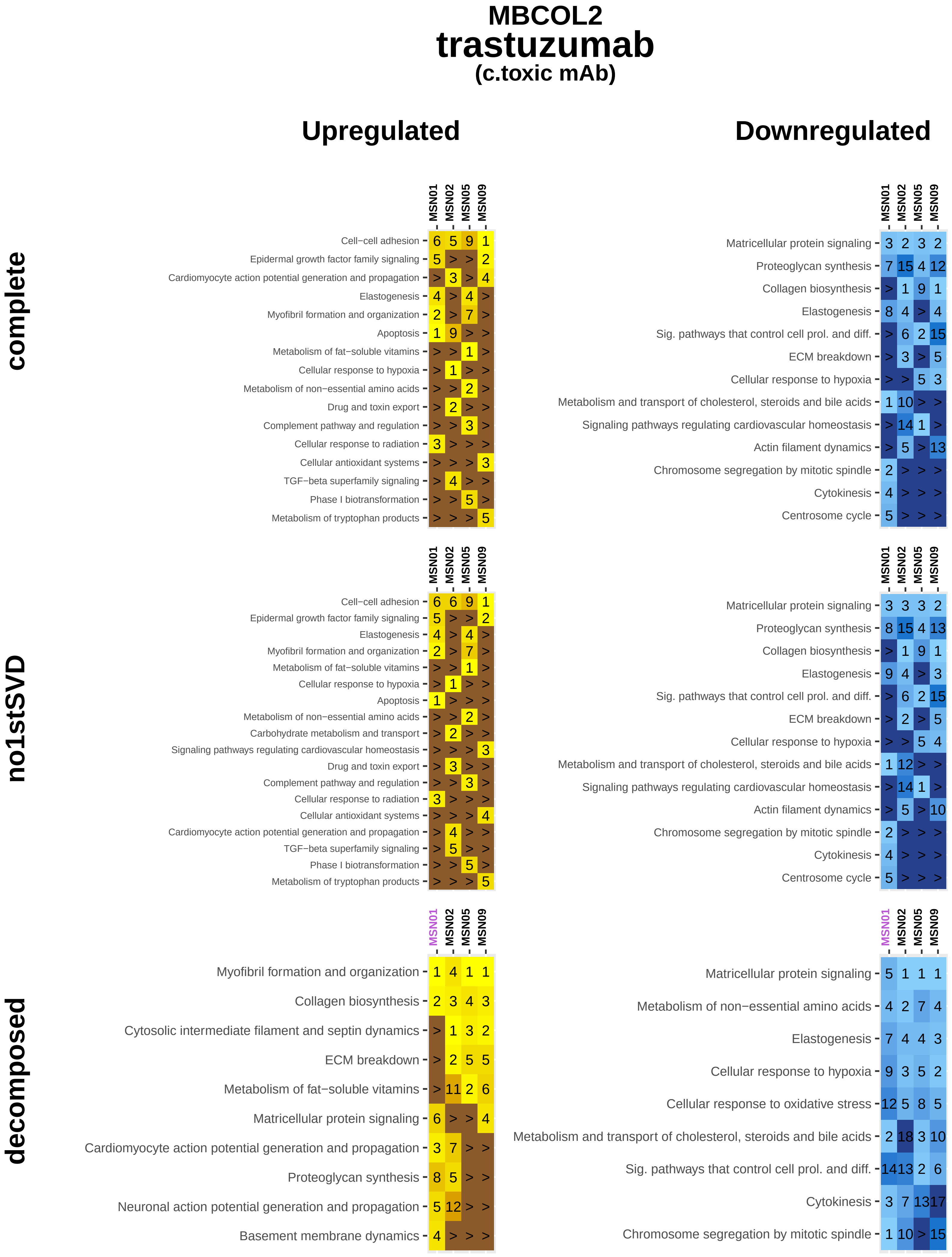
MBCOL2
trastuzumab
(c.toxic mAb)
Upregulated
Downregulated
complete
no1stSVD
decomposed
MSN01
MSN02
MSN05
MSN09
Cell−cell adhesion
Epidermal growth factor family signaling
Cardiomyocyte action potential generation and propagation
Elastogenesis
Myofibril formation and organization
Apoptosis
Metabolism of fat−soluble vitamins
Cellular response to hypoxia
Metabolism of non−essential amino acids
Drug and toxin export
Complement pathway and regulation
Cellular response to radiation
Cellular antioxidant systems
TGF−beta superfamily signaling
Phase I biotransformation
Metabolism of tryptophan products
Matricellular protein signaling
Proteoglycan synthesis
Collagen biosynthesis
Sig. pathways that control cell prol. and diff.
ECM breakdown
Metabolism and transport of cholesterol, steroids and bile acids
Signaling pathways regulating cardiovascular homeostasis
Actin filament dynamics
Chromosome segregation by mitotic spindle
Cytokinesis
Centrosome cycle
Carbohydrate metabolism and transport
Cytosolic intermediate filament and septin dynamics
Neuronal action potential generation and propagation
Basement membrane dynamics
Cellular response to oxidative stress

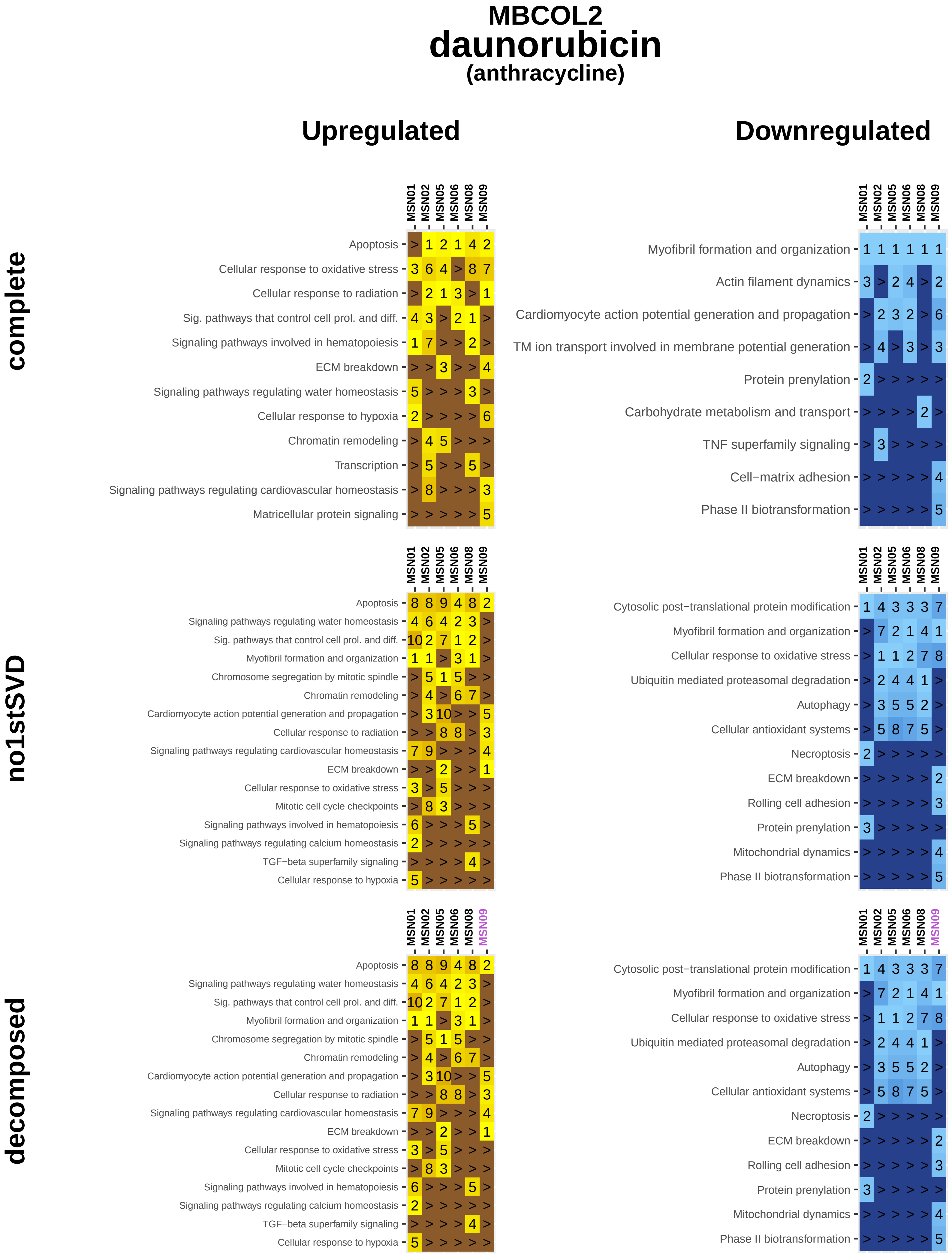
MBCOL2
daunorubicin
(anthracycline)
Upregulated
Downregulated
complete
no1stSVD
decomposed
MSN01
MSN02
MSN05
MSN06
MSN08
MSN09
Apoptosis
Cellular response to oxidative stress
Cellular response to radiation
Sig. pathways that control cell prol. and diff.
Signaling pathways involved in hematopoiesis
ECM breakdown
Signaling pathways regulating water homeostasis
Cellular response to hypoxia
Chromatin remodeling
Transcription
Signaling pathways regulating cardiovascular homeostasis
Matricellular protein signaling
Myofibril formation and organization
Actin filament dynamics
Cardiomyocyte action potential generation and propagation
TM ion transport involved in membrane potential generation
Protein prenylation
Carbohydrate metabolism and transport
TNF superfamily signaling
Cell−matrix adhesion
Phase II biotransformation
Chromosome segregation by mitotic spindle
Mitotic cell cycle checkpoints
Signaling pathways regulating calcium homeostasis
TGF−beta superfamily signaling
Cytosolic post−translational protein modification
Ubiquitin mediated proteasomal degradation
Autophagy
Cellular antioxidant systems
Necroptosis
Rolling cell adhesion
Mitochondrial dynamics

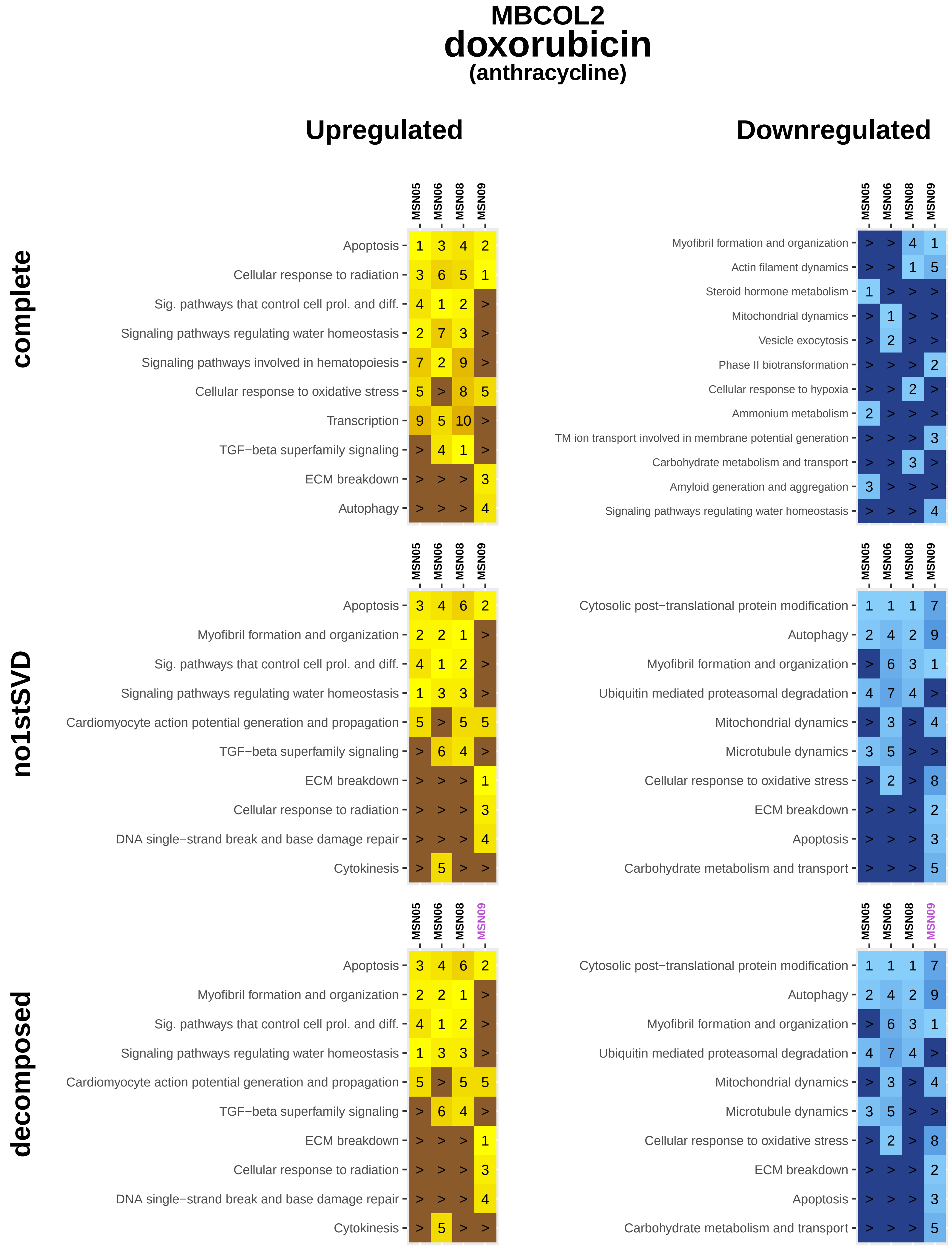
MBCOL2
doxorubicin
(anthracycline)
Upregulated
Downregulated
complete
no1stSVD
decomposed
MSN05
MSN06
MSN08
MSN09
Apoptosis
Cellular response to radiation
Sig. pathways that control cell prol. and diff.
Signaling pathways regulating water homeostasis
Signaling pathways involved in hematopoiesis
Cellular response to oxidative stress
Transcription
TGF−beta superfamily signaling
ECM breakdown
Autophagy
Myofibril formation and organization
Actin filament dynamics
Steroid hormone metabolism
Mitochondrial dynamics
Vesicle exocytosis
Phase II biotransformation
Cellular response to hypoxia
Ammonium metabolism
TM ion transport involved in membrane potential generation
Carbohydrate metabolism and transport
Amyloid generation and aggregation
Cardiomyocyte action potential generation and propagation
DNA single−strand break and base damage repair
Cytokinesis
Cytosolic post−translational protein modification
Ubiquitin mediated proteasomal degradation
Microtubule dynamics

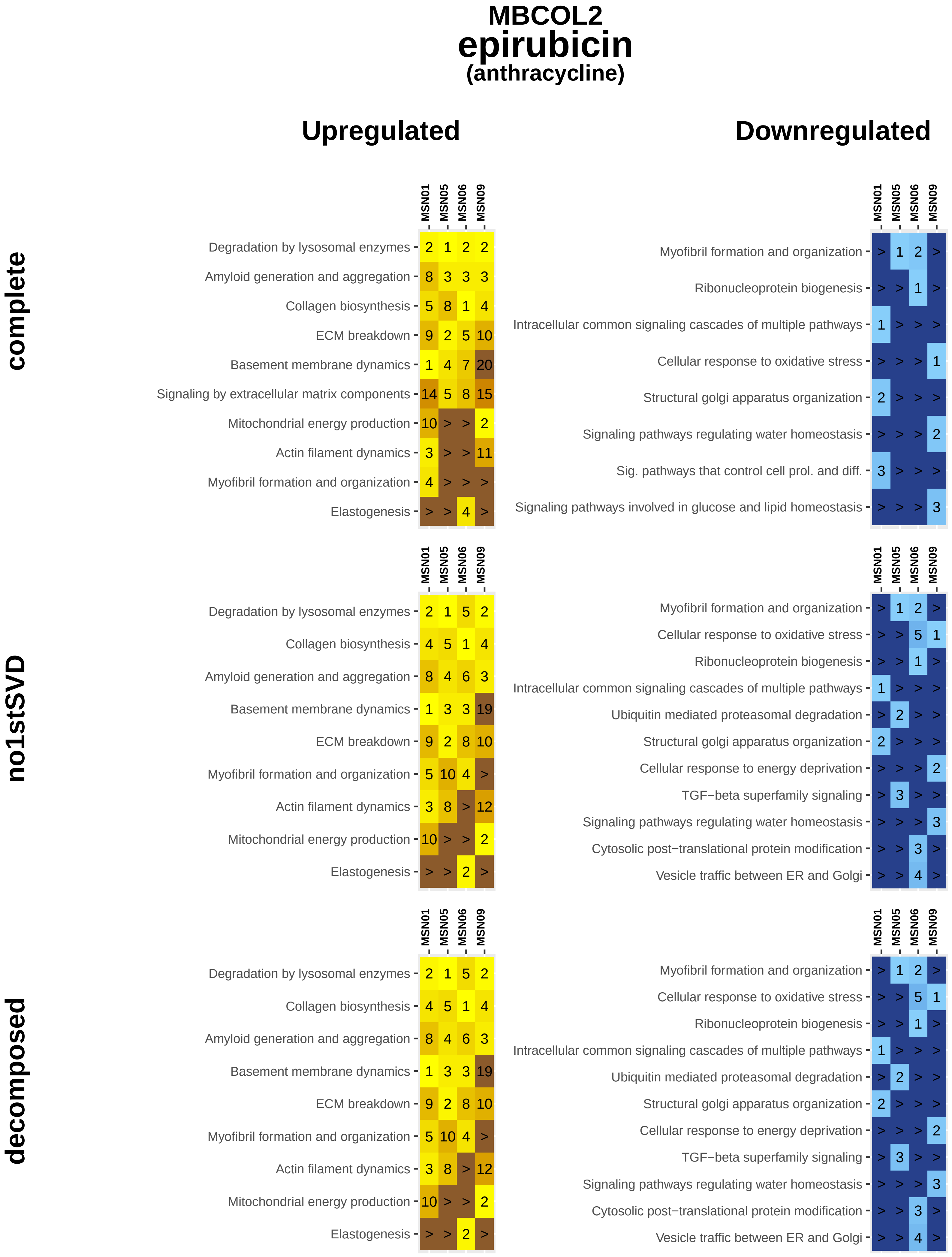
MBCOL2
epirubicin
(anthracycline)
Upregulated
Downregulated
complete
no1stSVD
decomposed
MSN01 MSN05 MSN06 MSN09
Degradation by lysosomal enzymes 2 1 2 2
Amyloid generation and aggregation 8 3 3 3
Collagen biosynthesis 5 8 1 4
ECM breakdown 9 2 5 10
Basement membrane dynamics 1 4 7 20
Signaling by extracellular matrix components 14 5 8 15
Mitochondrial energy production 10 > > 2
Actin filament dynamics 3 > > 11
Myofibril formation and organization 4 > > >
Elastogenesis > > 4 >
MSN01 MSN05 MSN06 MSN09
Myofibril formation and organization > 1 2 >
Ribonucleoprotein biogenesis > > 1 >
Intracellular common signaling cascades of multiple pathways 1 > > >
Cellular response to oxidative stress > > > 1
Structural golgi apparatus organization 2 > > >
Signaling pathways regulating water homeostasis > > > 2
Sig. pathways that control cell prol. and diff. 3 > > >
Signaling pathways involved in glucose and lipid homeostasis > > > 3
MSN01 MSN05 MSN06 MSN09
Degradation by lysosomal enzymes 2 1 5 2
Collagen biosynthesis 4 5 1 4
Amyloid generation and aggregation 8 4 6 3
Basement membrane dynamics 1 3 3 19
ECM breakdown 9 2 8 10
Myofibril formation and organization 5 10 4 >
Actin filament dynamics 3 8 > 12
Mitochondrial energy production 10 > > 2
Elastogenesis > > 2 >
MSN01 MSN05 MSN06 MSN09
Myofibril formation and organization > 1 2 >
Cellular response to oxidative stress > > 5 1
Ribonucleoprotein biogenesis > > 1 >
Intracellular common signaling cascades of multiple pathways 1 > > >
Ubiquitin mediated proteasomal degradation > 2 > >
Structural golgi apparatus organization 2 > > >
Cellular response to energy deprivation > > > 2
TGF−beta superfamily signaling > 3 > >
Signaling pathways regulating water homeostasis > > > 3
Cytosolic post−translational protein modification > > 3 >
Vesicle traffic between ER and Golgi > > 4 >
MSN01 MSN05 MSN06 MSN09
Degradation by lysosomal enzymes 2 1 5 2
Collagen biosynthesis 4 5 1 4
Amyloid generation and aggregation 8 4 6 3
Basement membrane dynamics 1 3 3 19
ECM breakdown 9 2 8 10
Myofibril formation and organization 5 10 4 >
Actin filament dynamics 3 8 > 12
Mitochondrial energy production 10 > > 2
Elastogenesis > > 2 >
MSN01 MSN05 MSN06 MSN09
Myofibril formation and organization > 1 2 >
Cellular response to oxidative stress > > 5 1
Ribonucleoprotein biogenesis > > 1 >
Intracellular common signaling cascades of multiple pathways 1 > > >
Ubiquitin mediated proteasomal degradation > 2 > >
Structural golgi apparatus organization 2 > > >
Cellular response to energy deprivation > > > 2
TGF−beta superfamily signaling > 3 > >
Signaling pathways regulating water homeostasis > > > 3
Cytosolic post−translational protein modification > > 3 >
Vesicle traffic between ER and Golgi > > 4 >

**MBCOL2**

### idarubicin

**(anthracycline)**

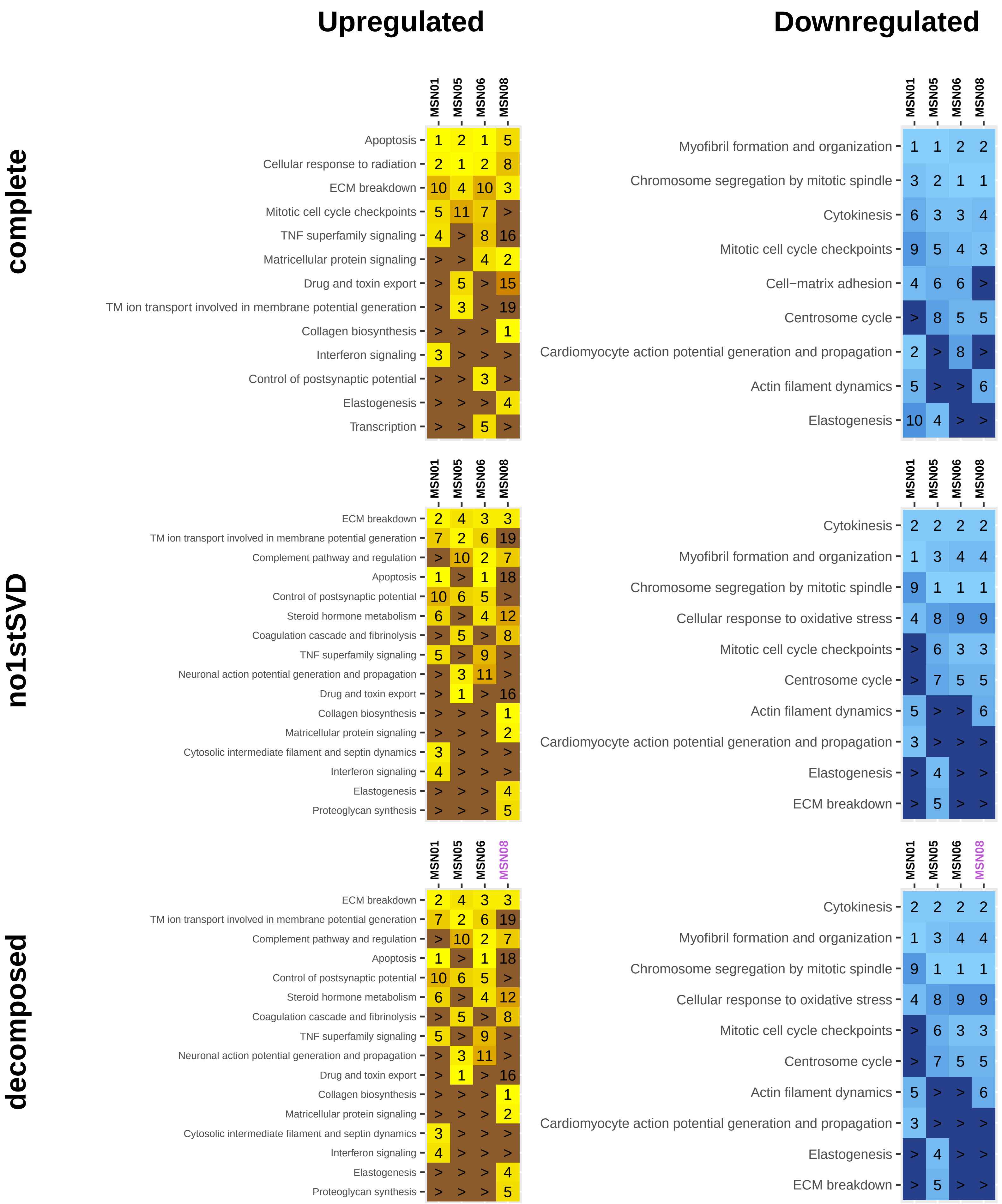

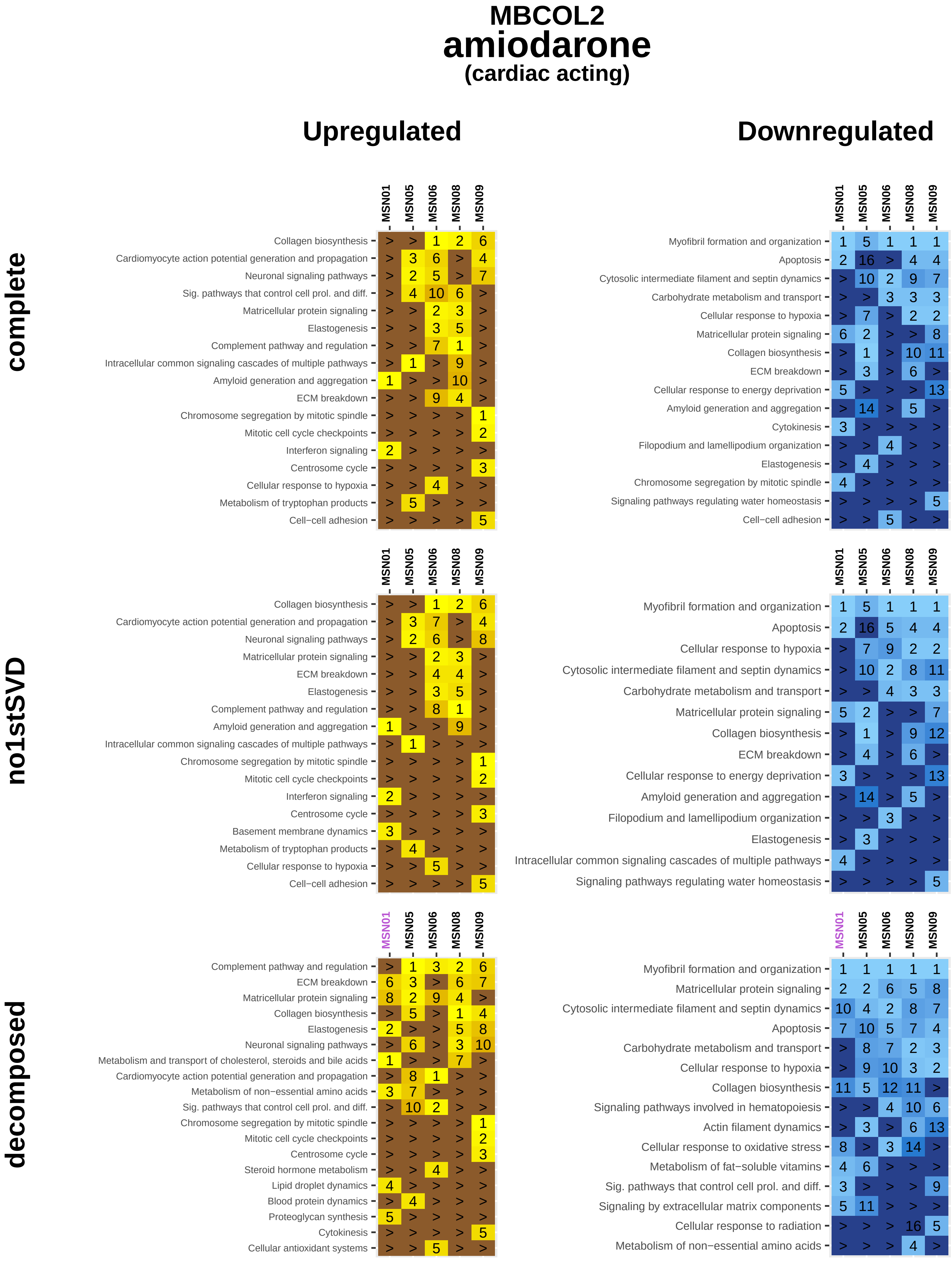
MBCOL2
amiodarone
(cardiac acting)
Upregulated
Downregulated
complete
no1stSVD
decomposed
MSN01
MSN05
MSN06
MSN08
MSN09
Collagen biosynthesis
Cardiomyocyte action potential generation and propagation
Neuronal signaling pathways
Sig. pathways that control cell prol. and diff.
Matricellular protein signaling
Elastogenesis
Complement pathway and regulation
Intracellular common signaling cascades of multiple pathways
Amyloid generation and aggregation
ECM breakdown
Chromosome segregation by mitotic spindle
Mitotic cell cycle checkpoints
Interferon signaling
Centrosome cycle
Cellular response to hypoxia
Metabolism of tryptophan products
Cell−cell adhesion
Myofibril formation and organization
Apoptosis
Cytosolic intermediate filament and septin dynamics
Carbohydrate metabolism and transport
Cellular response to energy deprivation
Cytokinesis
Filopodium and lamellipodium organization
Signaling pathways regulating water homeostasis
Basement membrane dynamics
Metabolism and transport of cholesterol, steroids and bile acids
Metabolism of non−essential amino acids
Steroid hormone metabolism
Lipid droplet dynamics
Blood protein dynamics
Proteoglycan synthesis
Cellular antioxidant systems
Signaling pathways involved in hematopoiesis
Actin filament dynamics
Cellular response to oxidative stress
Metabolism of fat−soluble vitamins
Signaling by extracellular matrix components
Cellular response to radiation

### MBCOL2
### dobutamine
#### (cardiac acting)

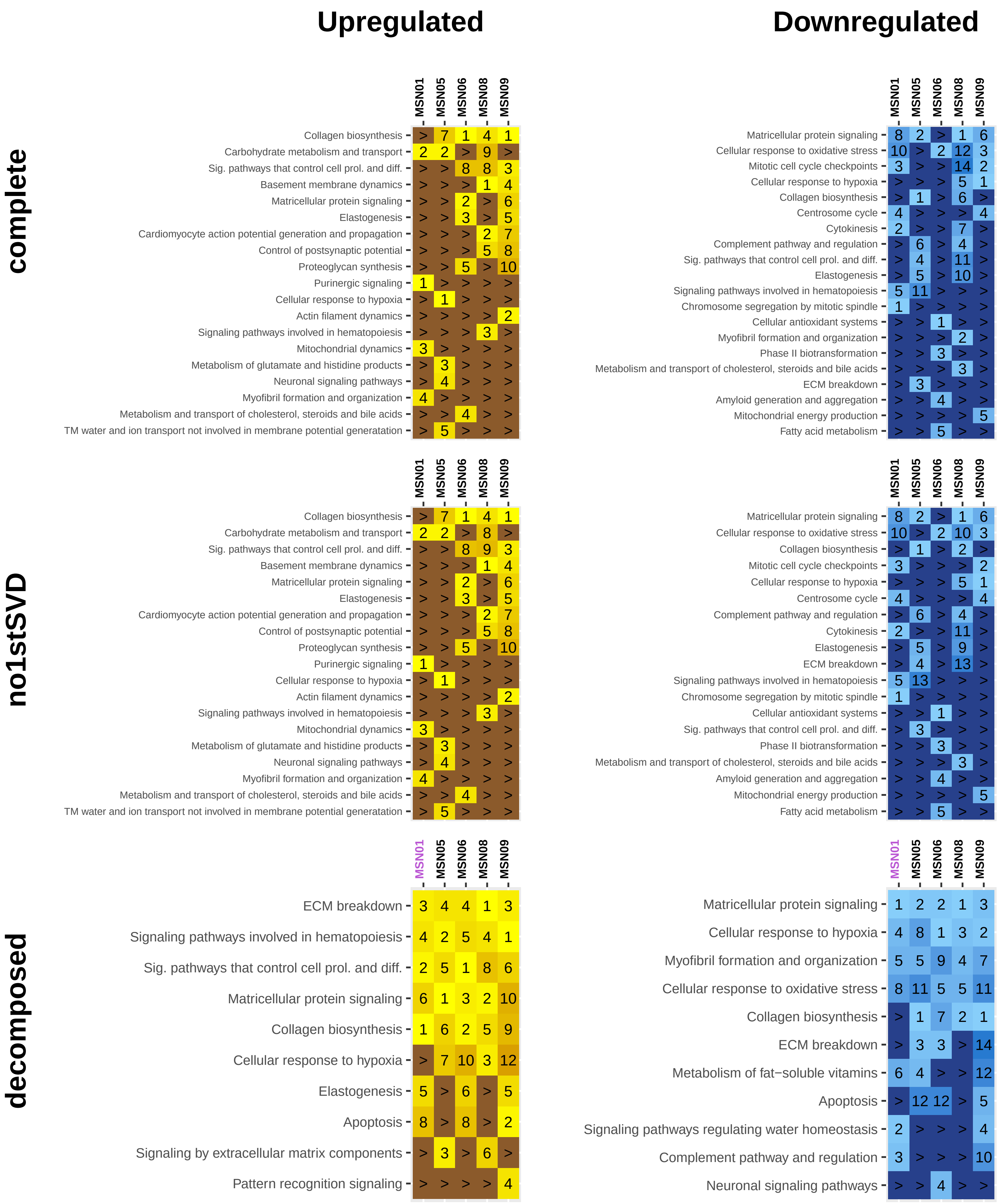

### MBCOL2
### flecainide
#### (cardiac acting)

#### Upregulated | Downregulated

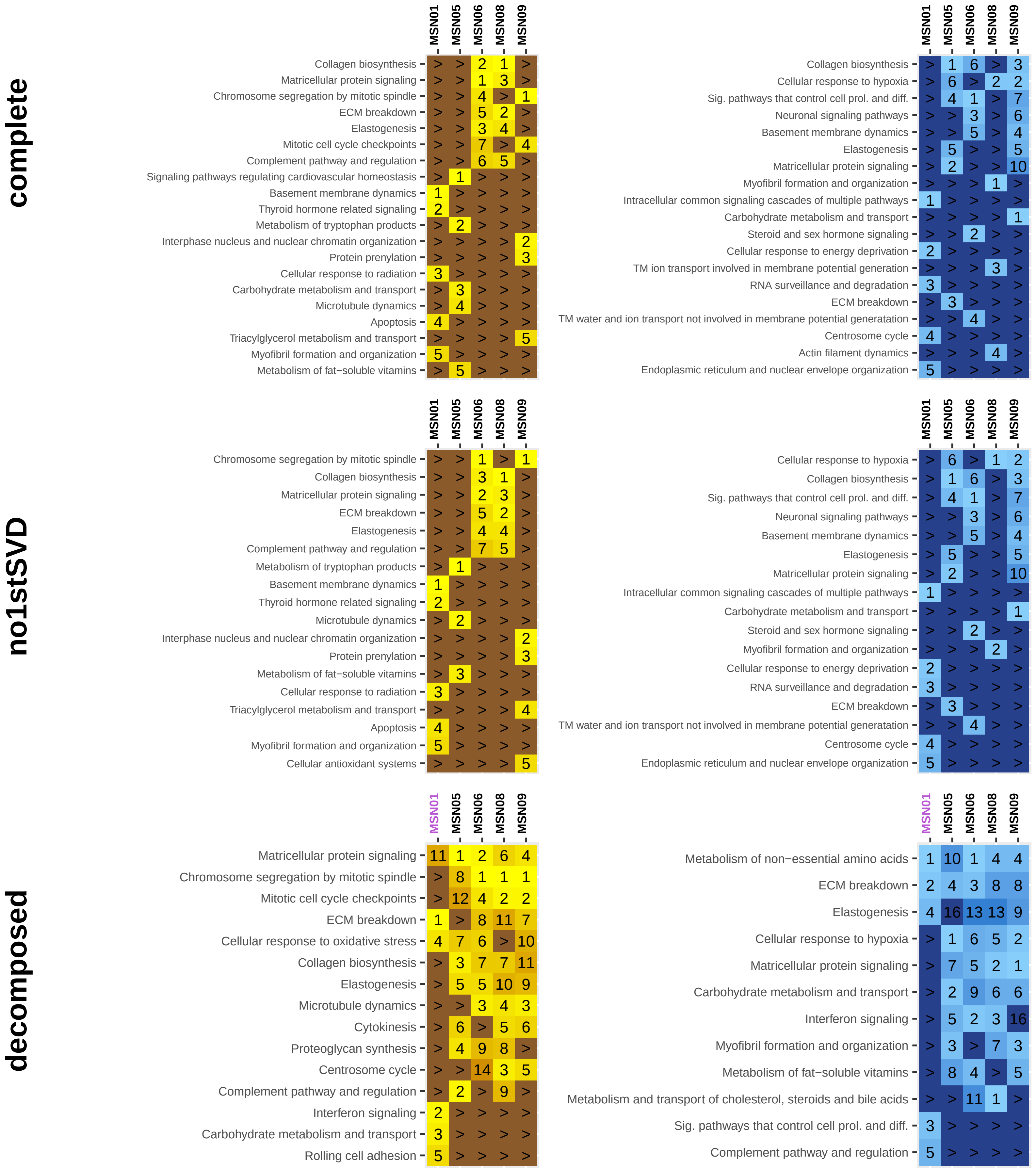

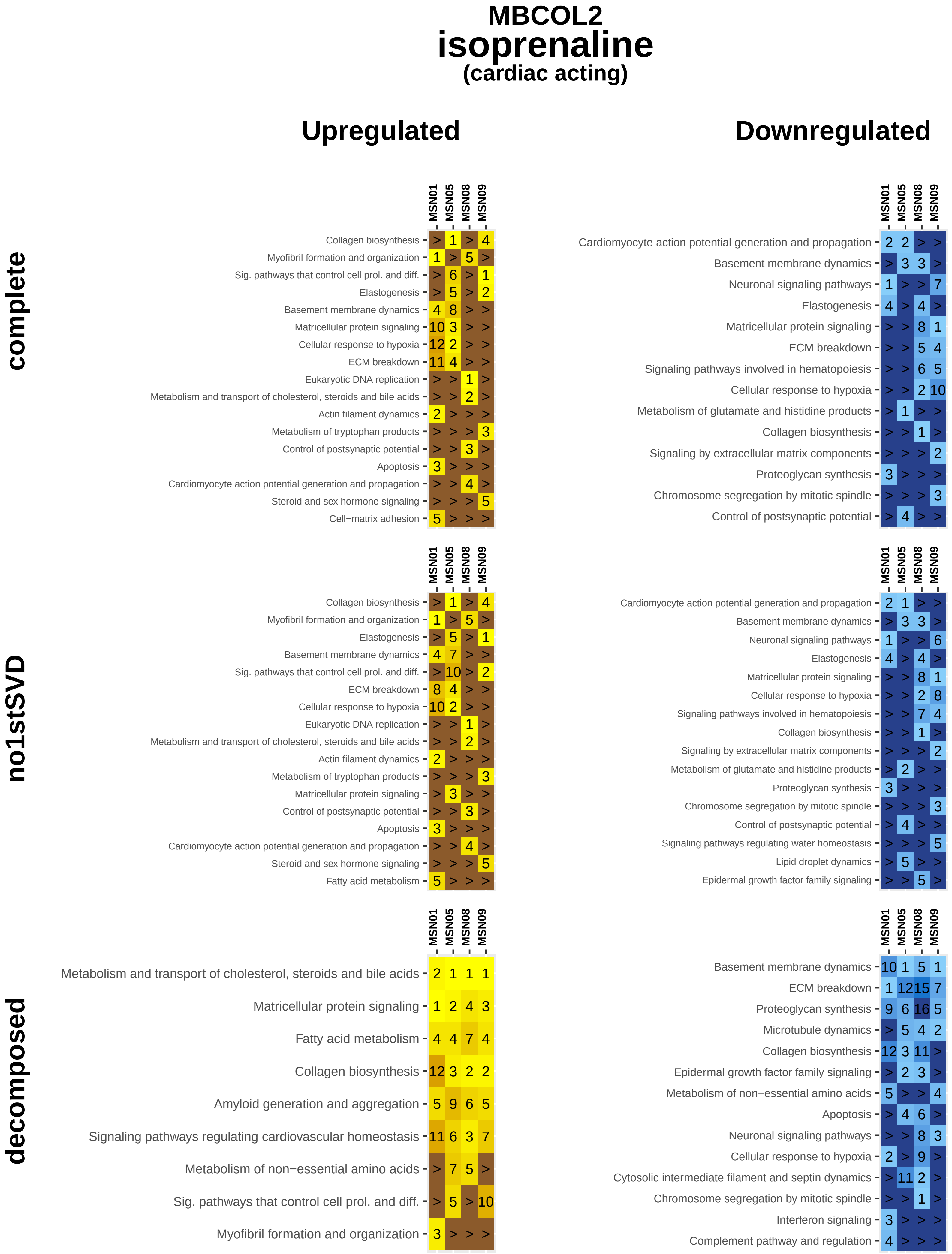

MBCOL2
isoprenaline
(cardiac acting)
Upregulated
Downregulated
complete
no1stSVD
decomposed
MSN01
MSN05
MSN08
MSN09
Collagen biosynthesis > 1 > 4
Myofibril formation and organization 1 > 5 >
Sig. pathways that control cell prol. and diff. > 6 > 1
Elastogenesis > 5 > 2
Basement membrane dynamics 4 8 > >
Matricellular protein signaling 10 3 > >
Cellular response to hypoxia 12 2 > >
ECM breakdown 11 4 > >
Eukaryotic DNA replication > > 1 >
Metabolism and transport of cholesterol, steroids and bile acids > > 2 >
Actin filament dynamics 2 > > >
Metabolism of tryptophan products > > > 3
Control of postsynaptic potential > > 3 >
Apoptosis 3 > > >
Cardiomyocyte action potential generation and propagation > > 4 >
Steroid and sex hormone signaling > > > 5
Cell−matrix adhesion 5 > > >
Cardiomyocyte action potential generation and propagation 2 2 > >
Basement membrane dynamics > 3 3 >
Neuronal signaling pathways 1 > > 7
Elastogenesis 4 > 4 >
Matricellular protein signaling > > 8 1
ECM breakdown > > 5 4
Signaling pathways involved in hematopoiesis > > 6 5
Cellular response to hypoxia > > 2 10
Metabolism of glutamate and histidine products > 1 > >
Collagen biosynthesis > > 1 >
Signaling by extracellular matrix components > > > 2
Proteoglycan synthesis 3 > > >
Chromosome segregation by mitotic spindle > > > 3
Control of postsynaptic potential > 4 > >
Collagen biosynthesis > 1 > 4
Myofibril formation and organization 1 > 5 >
Elastogenesis > 5 > 1
Basement membrane dynamics 4 7 > >
Sig. pathways that control cell prol. and diff. > 10 > 2
ECM breakdown 8 4 > >
Cellular response to hypoxia 10 2 > >
Eukaryotic DNA replication > > 1 >
Metabolism and transport of cholesterol, steroids and bile acids > > 2 >
Actin filament dynamics 2 > > >
Metabolism of tryptophan products > > > 3
Matricellular protein signaling > 3 > >
Control of postsynaptic potential > > 3 >
Apoptosis 3 > > >
Cardiomyocyte action potential generation and propagation > > 4 >
Steroid and sex hormone signaling > > > 5
Fatty acid metabolism 5 > > >
Cardiomyocyte action potential generation and propagation 2 1 > >
Basement membrane dynamics > 3 3 >
Neuronal signaling pathways 1 > > 6
Elastogenesis 4 > 4 >
Matricellular protein signaling > > 8 1
Cellular response to hypoxia > > 2 8
Signaling pathways involved in hematopoiesis > > 7 4
Collagen biosynthesis > > 1 >
Signaling by extracellular matrix components > > > 2
Metabolism of glutamate and histidine products > 2 > >
Proteoglycan synthesis 3 > > >
Chromosome segregation by mitotic spindle > > > 3
Control of postsynaptic potential > 4 > >
Signaling pathways regulating water homeostasis > > > 5
Lipid droplet dynamics > 5 > >
Epidermal growth factor family signaling > > 5 >
Metabolism and transport of cholesterol, steroids and bile acids 2 1 1 1
Matricellular protein signaling 1 2 4 3
Fatty acid metabolism 4 4 7 4
Collagen biosynthesis 12 3 2 2
Amyloid generation and aggregation 5 9 6 5
Signaling pathways regulating cardiovascular homeostasis 11 6 3 7
Metabolism of non−essential amino acids > 7 5 >
Sig. pathways that control cell prol. and diff. > 5 > 10
Myofibril formation and organization 3 > > >
Basement membrane dynamics 10 1 5 1
ECM breakdown 1 12 15 7
Proteoglycan synthesis 9 6 16 5
Microtubule dynamics > 5 4 2
Collagen biosynthesis 12 3 11 >
Epidermal growth factor family signaling > 2 3 >
Metabolism of non−essential amino acids 5 > > 4
Apoptosis > 4 6 >
Neuronal signaling pathways > > 8 3
Cellular response to hypoxia 2 > 9 >
Cytosolic intermediate filament and septin dynamics > 11 2 >
Chromosome segregation by mitotic spindle > > 1 >
Interferon signaling 3 > > >
Complement pathway and regulation 4 > > >

### MBCOL2
### milrinone
#### (cardiac acting)

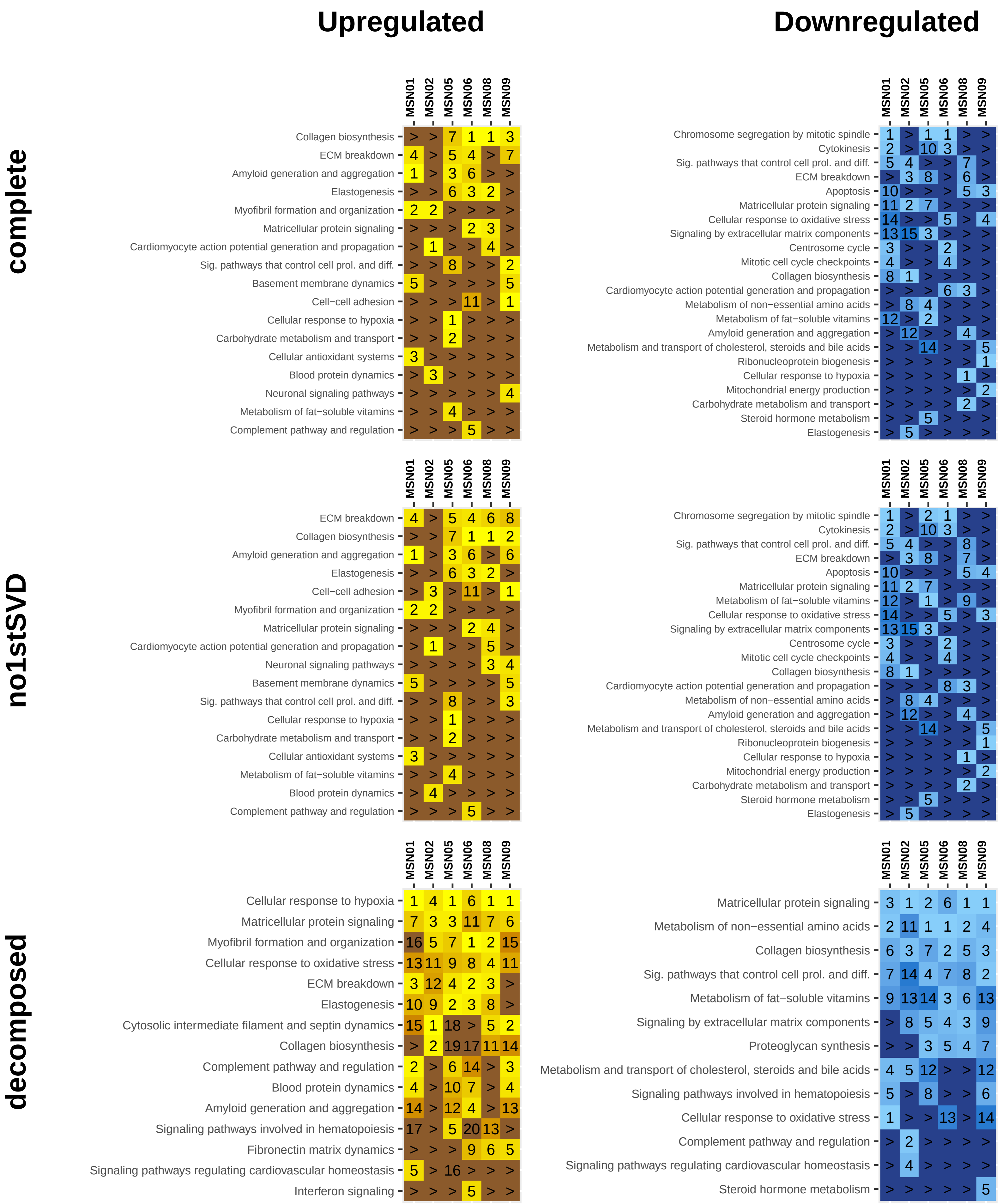

### MBCOL2
### phenylephrine
#### (cardiac acting)

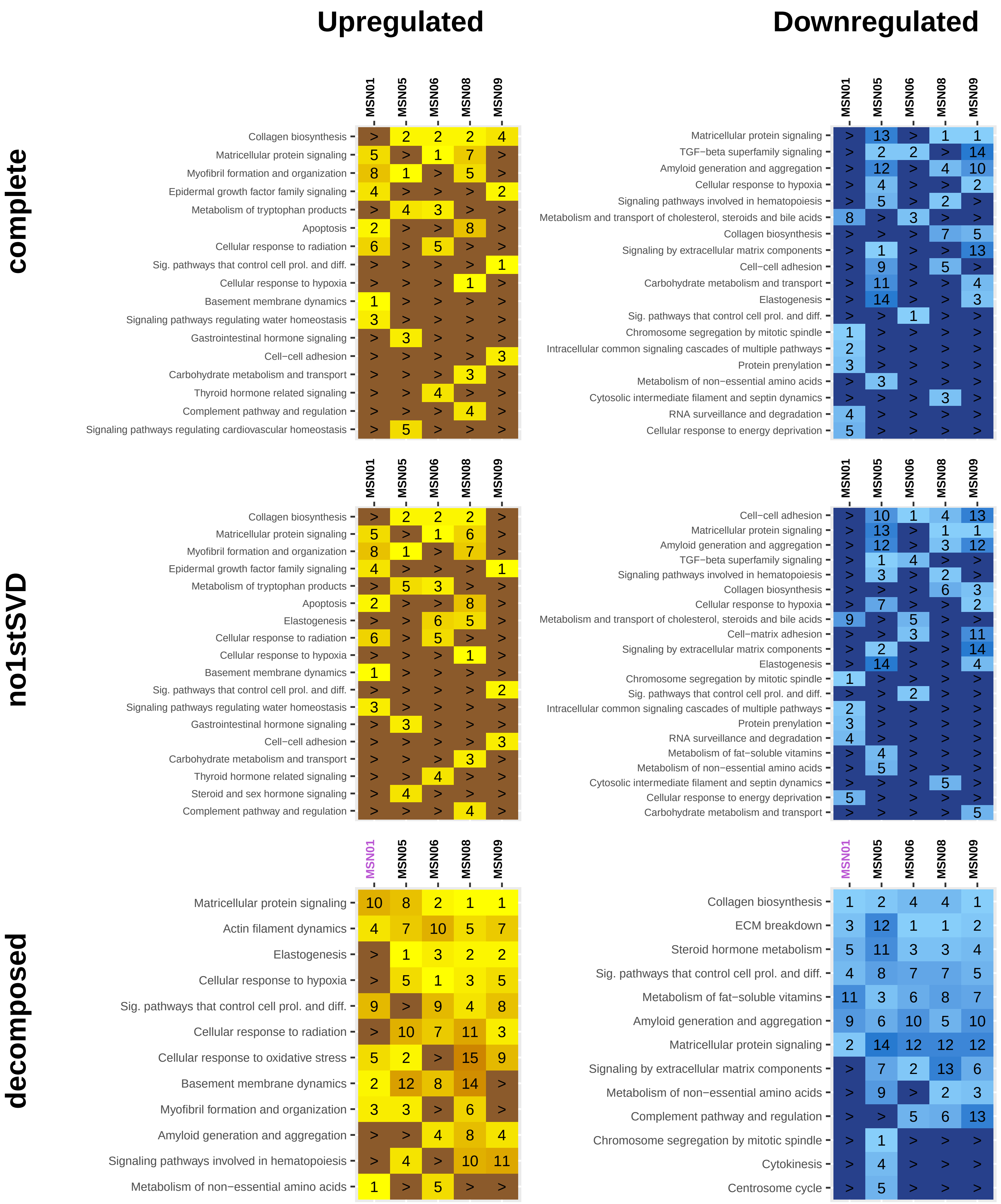

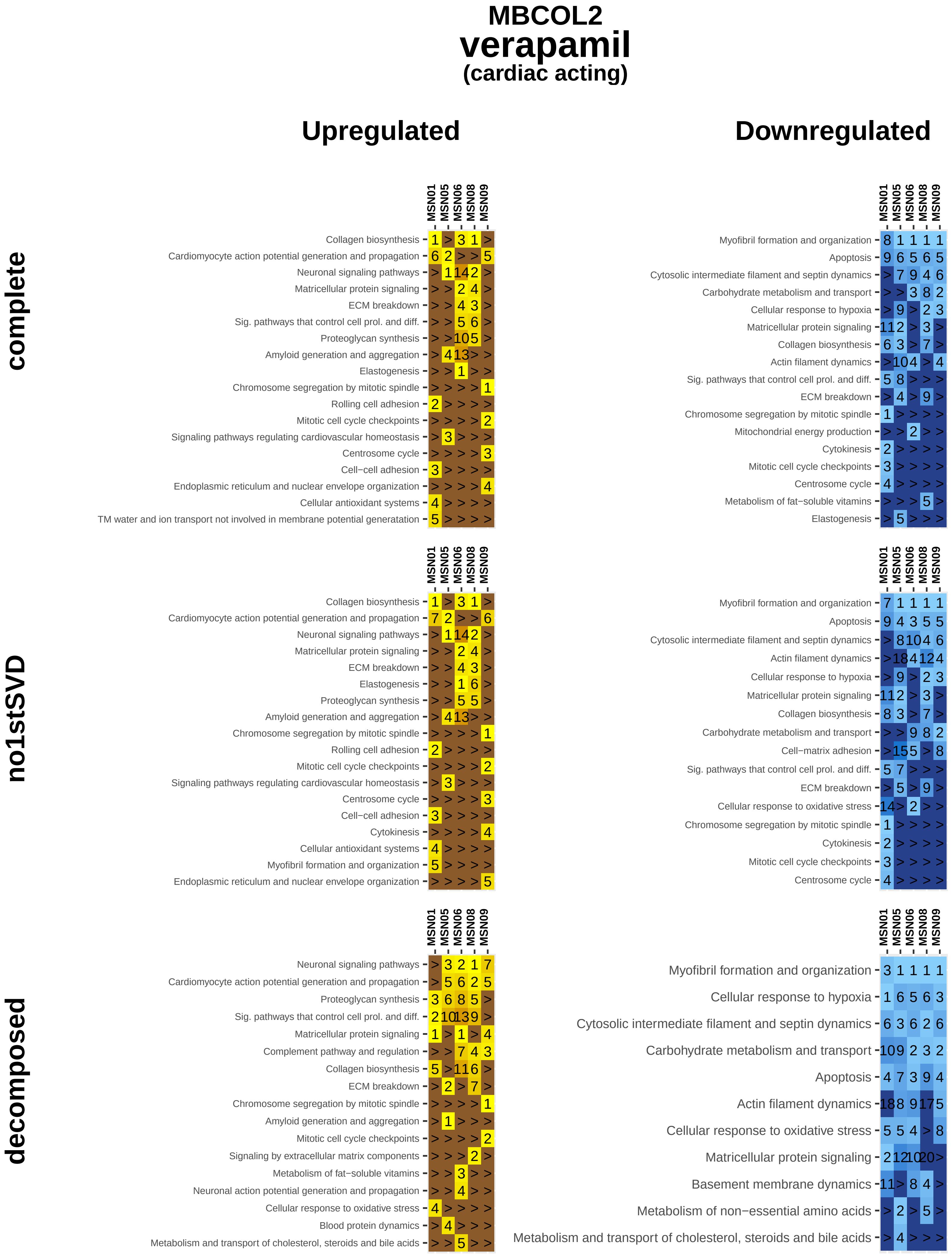
MBCOL2
verapamil
(cardiac acting)
Upregulated
Downregulated
complete
no1stSVD
decomposed
MSN01
MSN05
MSN06
MSN08
MSN09
Collagen biosynthesis
Cardiomyocyte action potential generation and propagation
Neuronal signaling pathways
Matricellular protein signaling
ECM breakdown
Sig. pathways that control cell prol. and diff.
Proteoglycan synthesis
Amyloid generation and aggregation
Elastogenesis
Chromosome segregation by mitotic spindle
Rolling cell adhesion
Mitotic cell cycle checkpoints
Signaling pathways regulating cardiovascular homeostasis
Centrosome cycle
Cell−cell adhesion
Endoplasmic reticulum and nuclear envelope organization
Cellular antioxidant systems
TM water and ion transport not involved in membrane potential generatation
Myofibril formation and organization
Apoptosis
Cytosolic intermediate filament and septin dynamics
Carbohydrate metabolism and transport
Cellular response to hypoxia
Actin filament dynamics
Mitochondrial energy production
Cytokinesis
Metabolism of fat−soluble vitamins
Cell−matrix adhesion
Cellular response to oxidative stress
Complement pathway and regulation
Signaling by extracellular matrix components
Neuronal action potential generation and propagation
Blood protein dynamics
Metabolism and transport of cholesterol, steroids and bile acids
Basement membrane dynamics
Metabolism of non−essential amino acids

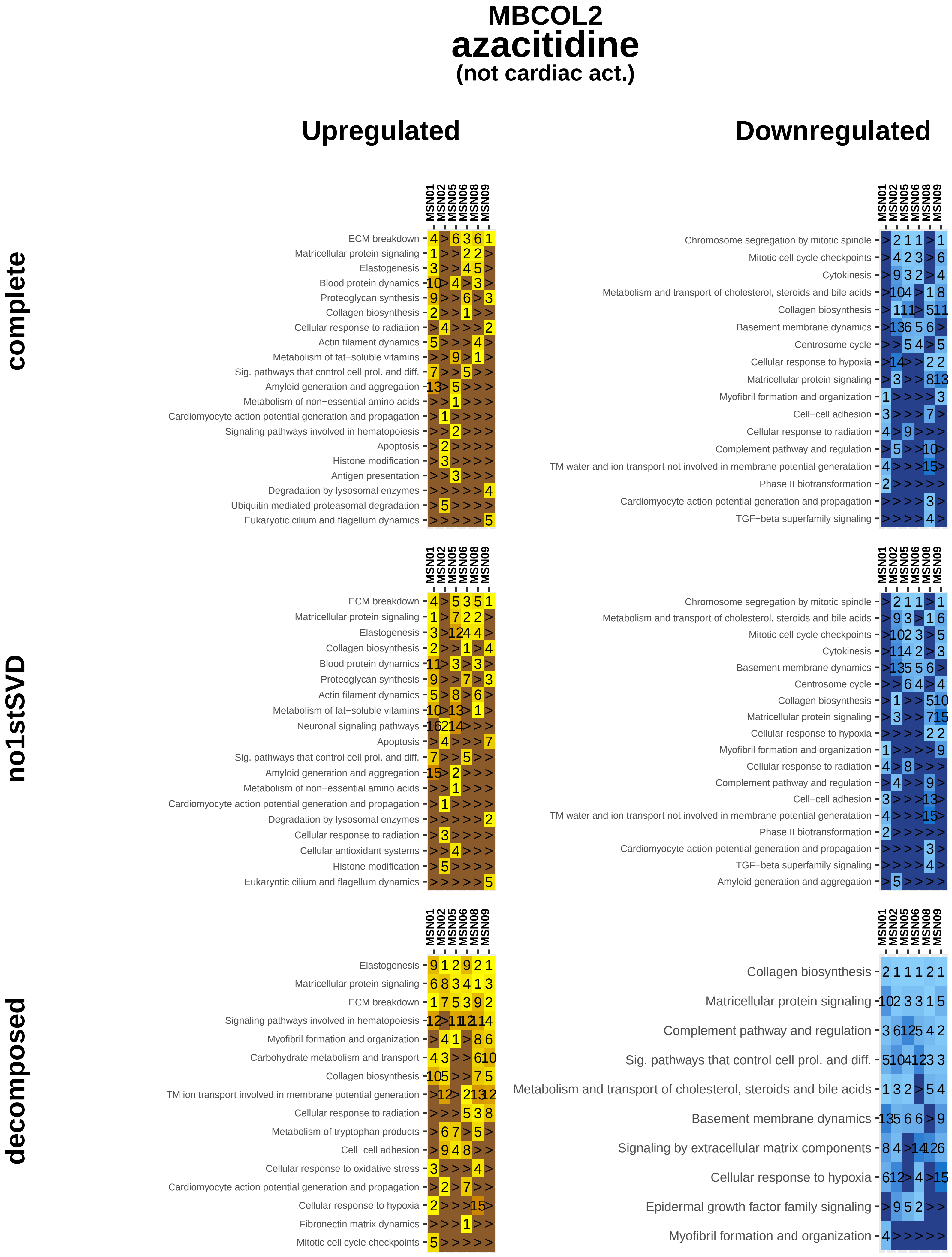

MBCOL2
azacitidine
(not cardiac act.)
Upregulated
Downregulated
complete
no1stSVD
decomposed
MSN01
MSN02
MSN05
MSN06
MSN08
MSN09
ECM breakdown
Matricellular protein signaling
Elastogenesis
Blood protein dynamics
Proteoglycan synthesis
Collagen biosynthesis
Cellular response to radiation
Actin filament dynamics
Metabolism of fat−soluble vitamins
Sig. pathways that control cell prol. and diff.
Amyloid generation and aggregation
Metabolism of non−essential amino acids
Cardiomyocyte action potential generation and propagation
Signaling pathways involved in hematopoiesis
Apoptosis
Histone modification
Antigen presentation
Degradation by lysosomal enzymes
Ubiquitin mediated proteasomal degradation
Eukaryotic cilium and flagellum dynamics
Chromosome segregation by mitotic spindle
Mitotic cell cycle checkpoints
Cytokinesis
Metabolism and transport of cholesterol, steroids and bile acids
Basement membrane dynamics
Centrosome cycle
Cellular response to hypoxia
Myofibril formation and organization
Cell−cell adhesion
Complement pathway and regulation
TM water and ion transport not involved in membrane potential generatation
Phase II biotransformation
TGF−beta superfamily signaling
Neuronal signaling pathways
Cellular antioxidant systems
Carbohydrate metabolism and transport
TM ion transport involved in membrane potential generation
Metabolism of tryptophan products
Cellular response to oxidative stress
Fibronectin matrix dynamics
Signaling by extracellular matrix components
Epidermal growth factor family signaling

#### MBCOL2
### bortezomib
##### (not cardiac act.)

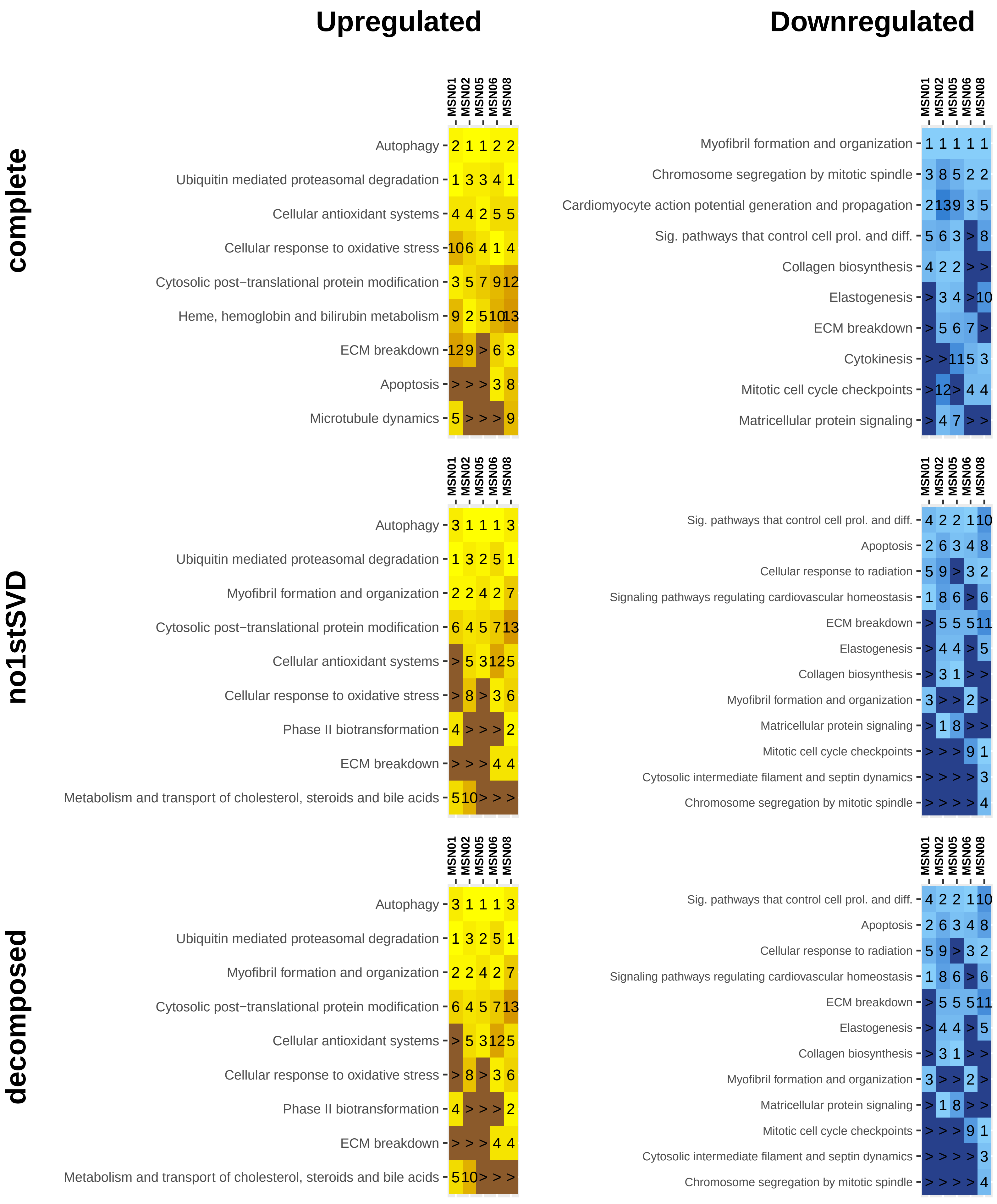

### MBCOL2
### carfilzomib
#### (not cardiac act.)

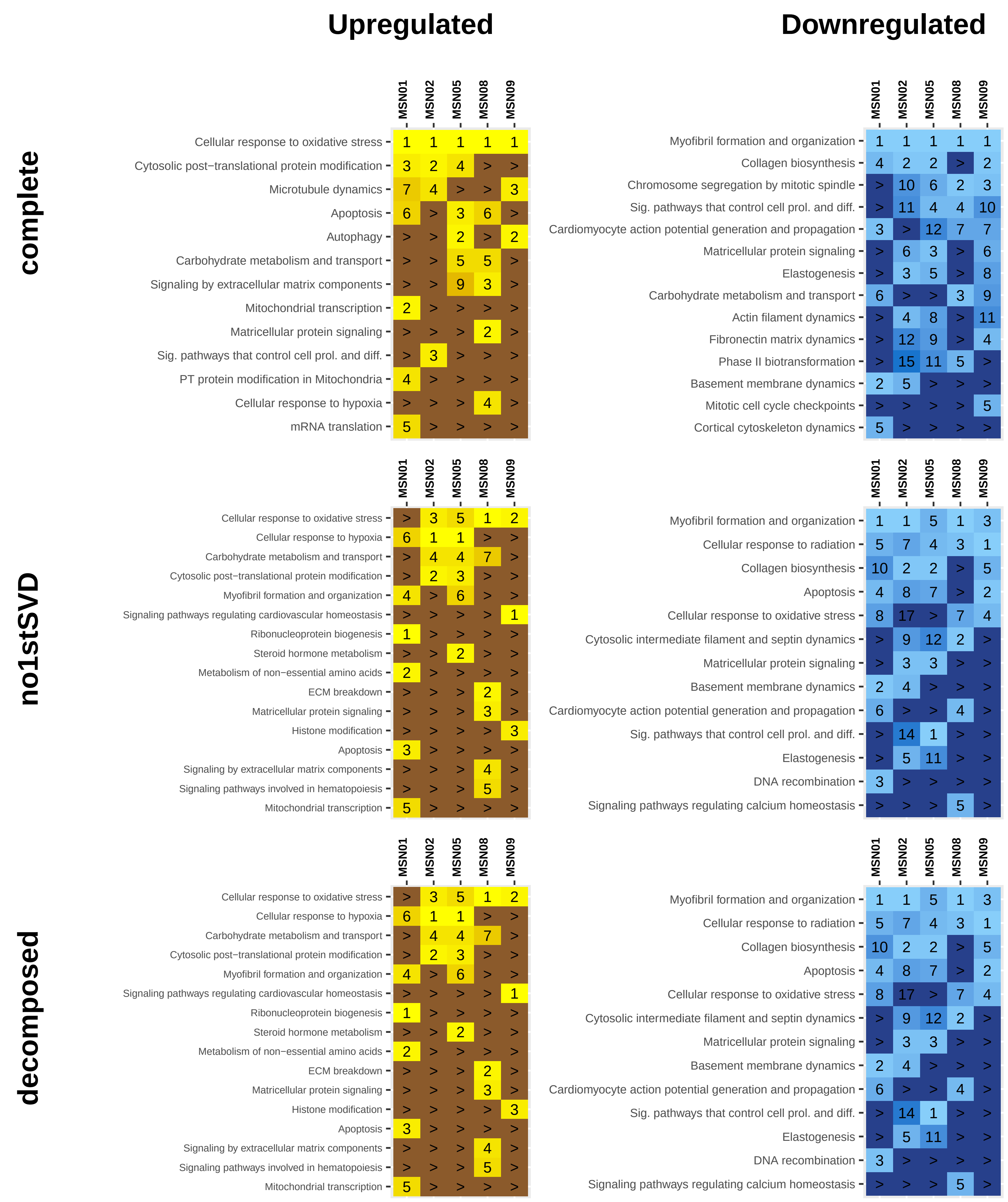

### MBCOL2
### cyclosporine
(not cardiac act.)

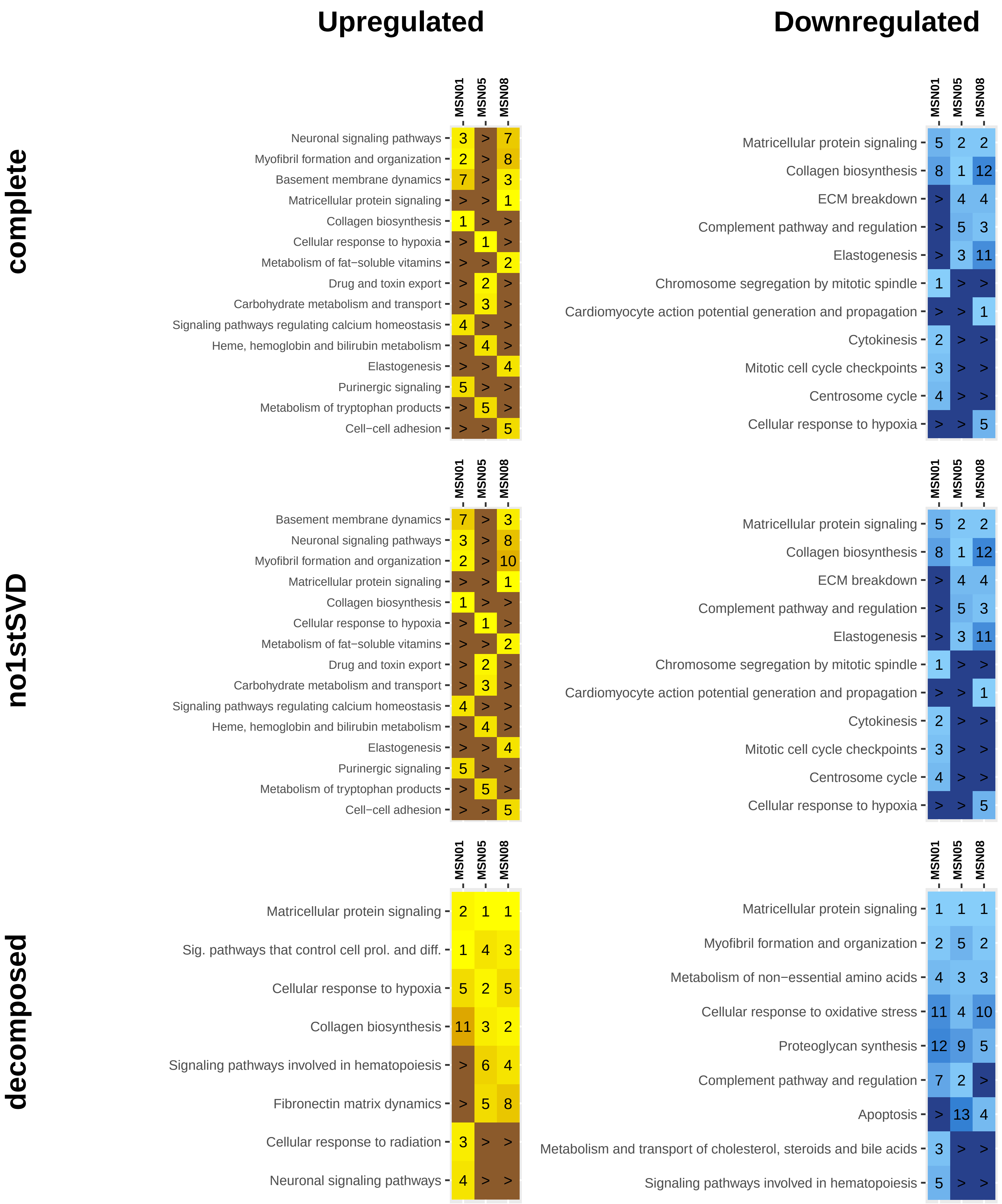

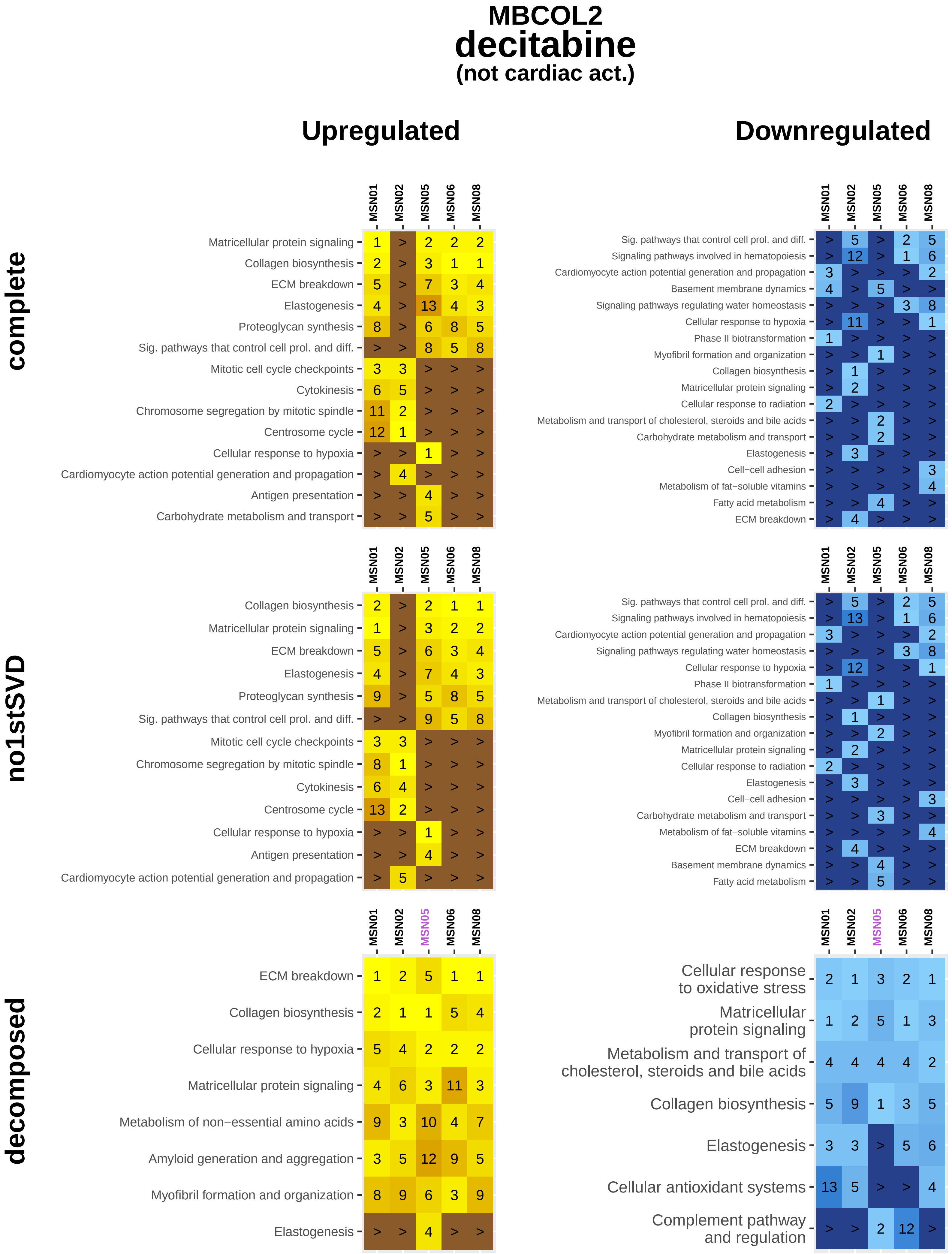
MBCOL2
decitabine
(not cardiac act.)
Upregulated
Downregulated
complete
MSN01 MSN02 MSN05 MSN06 MSN08
Matricellular protein signaling 1 > 2 2 2
Collagen biosynthesis 2 > 3 1 1
ECM breakdown 5 > 7 3 4
Elastogenesis 4 > 13 4 3
Proteoglycan synthesis 8 > 6 8 5
Sig. pathways that control cell prol. and diff. > > 8 5 8
Mitotic cell cycle checkpoints 3 3 > > >
Cytokinesis 6 5 > > >
Chromosome segregation by mitotic spindle 11 2 > > >
Centrosome cycle 12 1 > > >
Cellular response to hypoxia > > 1 > >
Cardiomyocyte action potential generation and propagation > 4 > > >
Antigen presentation > > 4 > >
Carbohydrate metabolism and transport > > 5 > >
MSN01 MSN02 MSN05 MSN06 MSN08
Sig. pathways that control cell prol. and diff. > 5 > 2 5
Signaling pathways involved in hematopoiesis > 12 > 1 6
Cardiomyocyte action potential generation and propagation 3 > > > 2
Basement membrane dynamics 4 > 5 > >
Signaling pathways regulating water homeostasis > > > 3 8
Cellular response to hypoxia > 11 > > 1
Phase II biotransformation 1 > > > >
Myofibril formation and organization > > 1 > >
Collagen biosynthesis > 1 > > >
Matricellular protein signaling > 2 > > >
Cellular response to radiation 2 > > > >
Metabolism and transport of cholesterol, steroids and bile acids > > 2 > >
Carbohydrate metabolism and transport > > 2 > >
Elastogenesis > 3 > > >
Cell−cell adhesion > > > > 3
Metabolism of fat−soluble vitamins > > > > 4
Fatty acid metabolism > > 4 > >
ECM breakdown > 4 > > >
no1stSVD
MSN01 MSN02 MSN05 MSN06 MSN08
Collagen biosynthesis 2 > 2 1 1
Matricellular protein signaling 1 > 3 2 2
ECM breakdown 5 > 6 3 4
Elastogenesis 4 > 7 4 3
Proteoglycan synthesis 9 > 5 8 5
Sig. pathways that control cell prol. and diff. > > 9 5 8
Mitotic cell cycle checkpoints 3 3 > > >
Chromosome segregation by mitotic spindle 8 1 > > >
Cytokinesis 6 4 > > >
Centrosome cycle 13 2 > > >
Cellular response to hypoxia > > 1 > >
Antigen presentation > > 4 > >
Cardiomyocyte action potential generation and propagation > 5 > > >
MSN01 MSN02 MSN05 MSN06 MSN08
Sig. pathways that control cell prol. and diff. > 5 > 2 5
Signaling pathways involved in hematopoiesis > 13 > 1 6
Cardiomyocyte action potential generation and propagation 3 > > > 2
Signaling pathways regulating water homeostasis > > > 3 8
Cellular response to hypoxia > 12 > > 1
Phase II biotransformation 1 > > > >
Metabolism and transport of cholesterol, steroids and bile acids > > 1 > >
Collagen biosynthesis > 1 > > >
Myofibril formation and organization > > 2 > >
Matricellular protein signaling > 2 > > >
Cellular response to radiation 2 > > > >
Elastogenesis > 3 > > >
Cell−cell adhesion > > > > 3
Carbohydrate metabolism and transport > > 3 > >
Metabolism of fat−soluble vitamins > > > > 4
ECM breakdown > 4 > > >
Basement membrane dynamics > > 4 > >
Fatty acid metabolism > > 5 > >
decomposed
MSN01 MSN02 MSN05 MSN06 MSN08
ECM breakdown 1 2 5 1 1
Collagen biosynthesis 2 1 1 5 4
Cellular response to hypoxia 5 4 2 2 2
Matricellular protein signaling 4 6 3 11 3
Metabolism of non−essential amino acids 9 3 10 4 7
Amyloid generation and aggregation 3 5 12 9 5
Myofibril formation and organization 8 9 6 3 9
Elastogenesis > > 4 > >
MSN01 MSN02 MSN05 MSN06 MSN08
Cellular response to oxidative stress 2 1 3 2 1
Matricellular protein signaling 1 2 5 1 3
Metabolism and transport of cholesterol, steroids and bile acids 4 4 4 4 2
Collagen biosynthesis 5 9 1 3 5
Elastogenesis 3 3 > 5 6
Cellular antioxidant systems 13 5 > > 4
Complement pathway and regulation > > 2 12 >

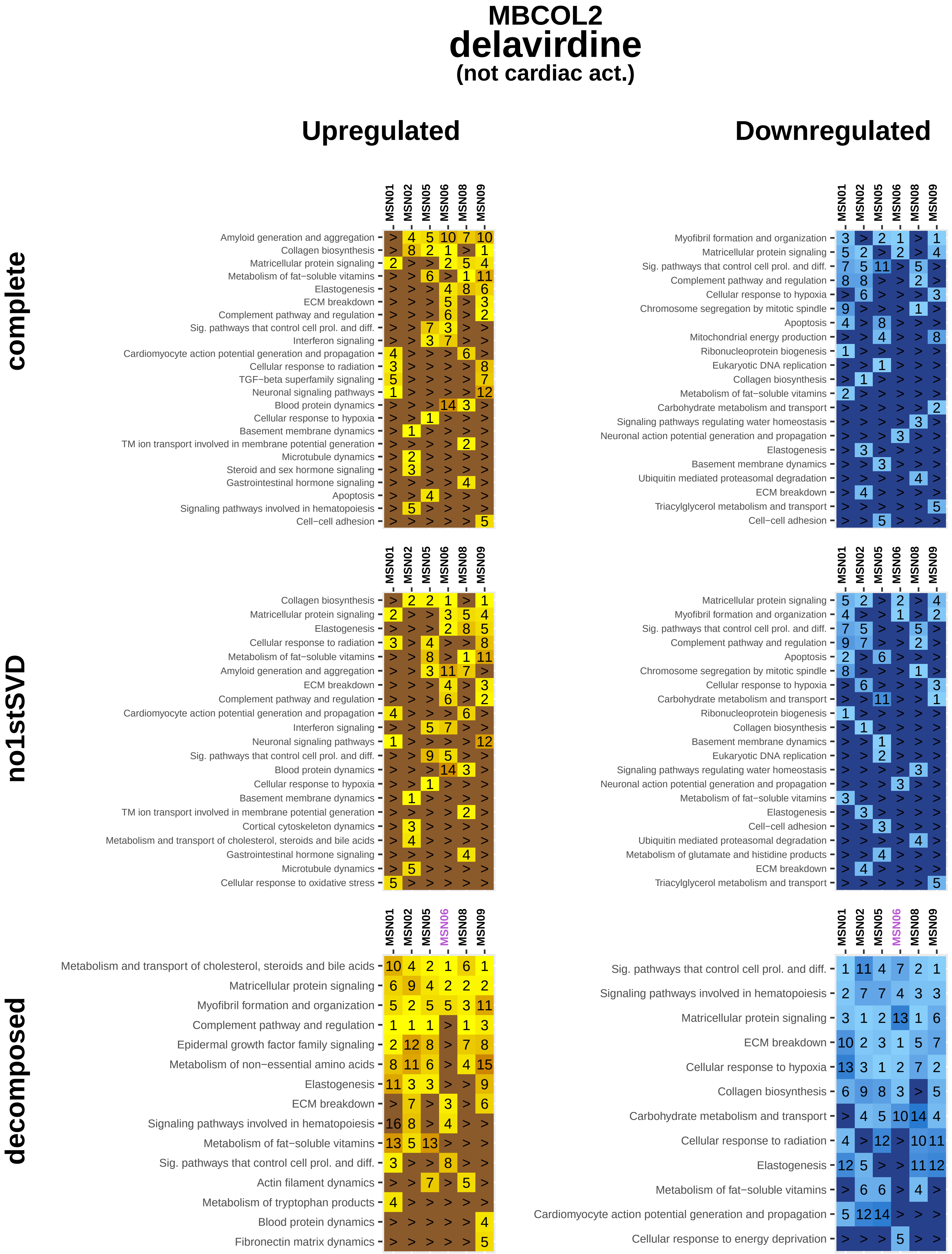
MBCOL2
delavirdine
(not cardiac act.)
Upregulated
Downregulated
complete
no1stSVD
decomposed
MSN01
MSN02
MSN05
MSN06
MSN08
MSN09
Amyloid generation and aggregation
Collagen biosynthesis
Matricellular protein signaling
Metabolism of fat−soluble vitamins
Elastogenesis
ECM breakdown
Complement pathway and regulation
Sig. pathways that control cell prol. and diff.
Interferon signaling
Cardiomyocyte action potential generation and propagation
Cellular response to radiation
TGF−beta superfamily signaling
Neuronal signaling pathways
Blood protein dynamics
Cellular response to hypoxia
Basement membrane dynamics
TM ion transport involved in membrane potential generation
Microtubule dynamics
Steroid and sex hormone signaling
Gastrointestinal hormone signaling
Apoptosis
Signaling pathways involved in hematopoiesis
Cell−cell adhesion
Myofibril formation and organization
Chromosome segregation by mitotic spindle
Mitochondrial energy production
Ribonucleoprotein biogenesis
Eukaryotic DNA replication
Carbohydrate metabolism and transport
Signaling pathways regulating water homeostasis
Neuronal action potential generation and propagation
Ubiquitin mediated proteasomal degradation
Triacylglycerol metabolism and transport
Cortical cytoskeleton dynamics
Metabolism and transport of cholesterol, steroids and bile acids
Cellular response to oxidative stress
Metabolism of glutamate and histidine products
Epidermal growth factor family signaling
Metabolism of non−essential amino acids
Actin filament dynamics
Metabolism of tryptophan products
Fibronectin matrix dynamics
Cellular response to energy deprivation

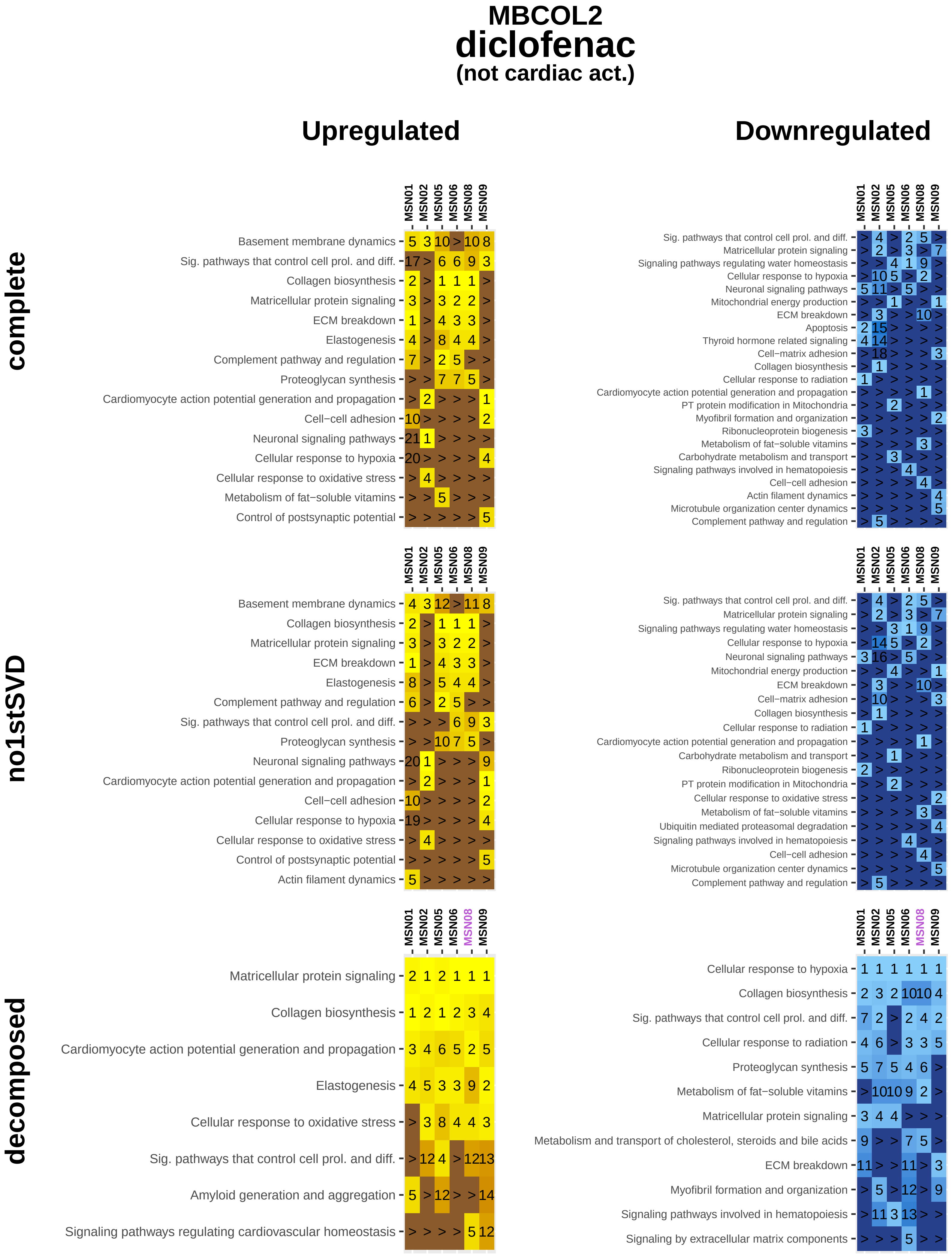
MBCOL2
diclofenac
(not cardiac act.)
Upregulated
Downregulated
complete
no1stSVD
decomposed
MSN01
MSN02
MSN05
MSN06
MSN08
MSN09
Basement membrane dynamics
Sig. pathways that control cell prol. and diff.
Collagen biosynthesis
Matricellular protein signaling
ECM breakdown
Elastogenesis
Complement pathway and regulation
Proteoglycan synthesis
Cardiomyocyte action potential generation and propagation
Cell−cell adhesion
Neuronal signaling pathways
Cellular response to hypoxia
Cellular response to oxidative stress
Metabolism of fat−soluble vitamins
Control of postsynaptic potential
Actin filament dynamics
Amyloid generation and aggregation
Signaling pathways regulating cardiovascular homeostasis
Signaling pathways regulating water homeostasis
Mitochondrial energy production
Apoptosis
Thyroid hormone related signaling
Cell−matrix adhesion
Cellular response to radiation
PT protein modification in Mitochondria
Myofibril formation and organization
Ribonucleoprotein biogenesis
Carbohydrate metabolism and transport
Signaling pathways involved in hematopoiesis
Microtubule organization center dynamics
Ubiquitin mediated proteasomal degradation
Metabolism and transport of cholesterol, steroids and bile acids
Signaling by extracellular matrix components

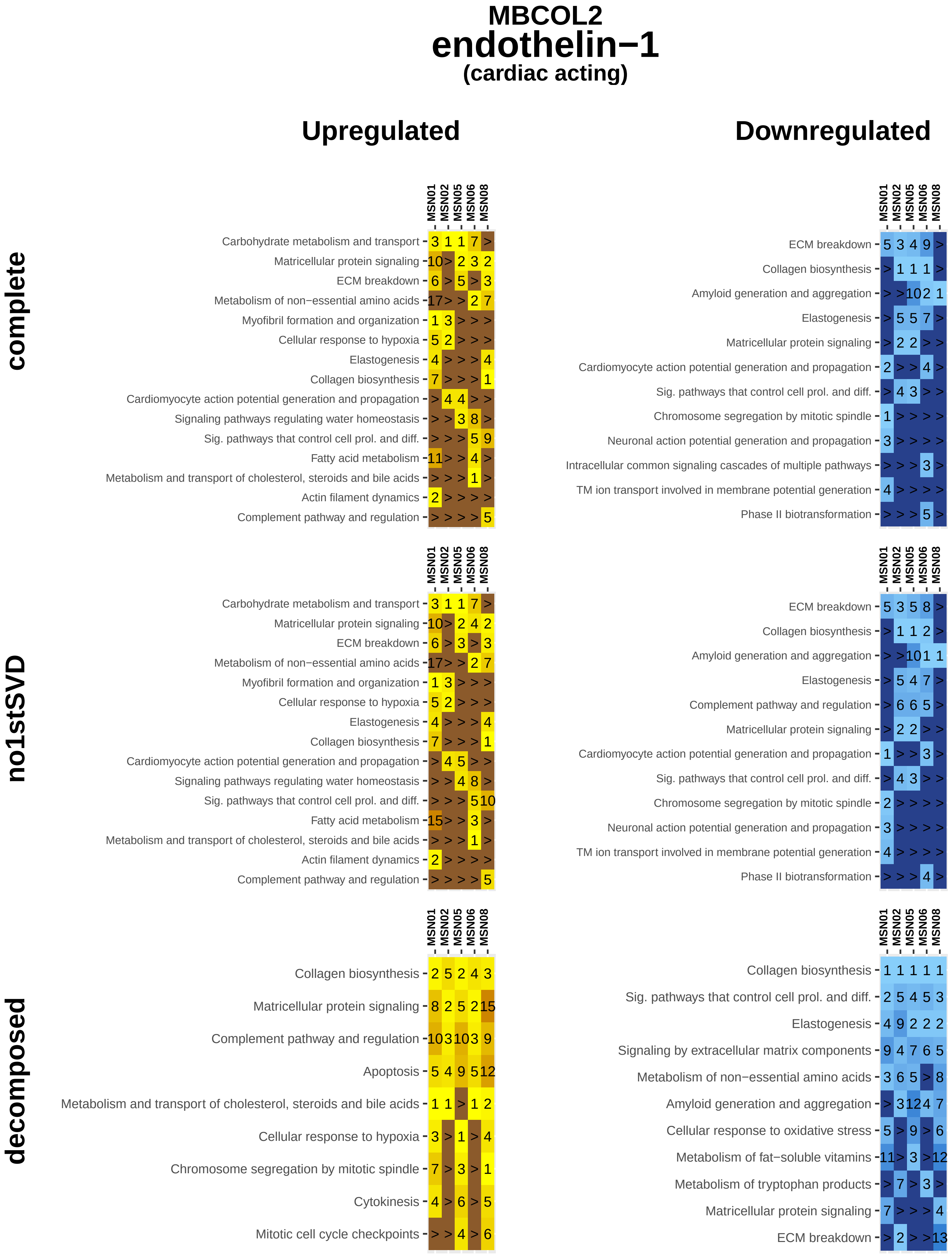
MBCOL2
endothelin−1
(cardiac acting)
Upregulated
Downregulated
complete
no1stSVD
decomposed
MSN01
MSN02
MSN05
MSN06
MSN08
Carbohydrate metabolism and transport
Matricellular protein signaling
ECM breakdown
Metabolism of non−essential amino acids
Myofibril formation and organization
Cellular response to hypoxia
Elastogenesis
Collagen biosynthesis
Cardiomyocyte action potential generation and propagation
Signaling pathways regulating water homeostasis
Sig. pathways that control cell prol. and diff.
Fatty acid metabolism
Metabolism and transport of cholesterol, steroids and bile acids
Actin filament dynamics
Complement pathway and regulation
Amyloid generation and aggregation
Chromosome segregation by mitotic spindle
Neuronal action potential generation and propagation
Intracellular common signaling cascades of multiple pathways
TM ion transport involved in membrane potential generation
Phase II biotransformation
Apoptosis
Cytokinesis
Mitotic cell cycle checkpoints
Signaling by extracellular matrix components
Cellular response to oxidative stress
Metabolism of fat−soluble vitamins
Metabolism of tryptophan products

**MBCOL2**

### estradiol

**(not cardiac act.)**

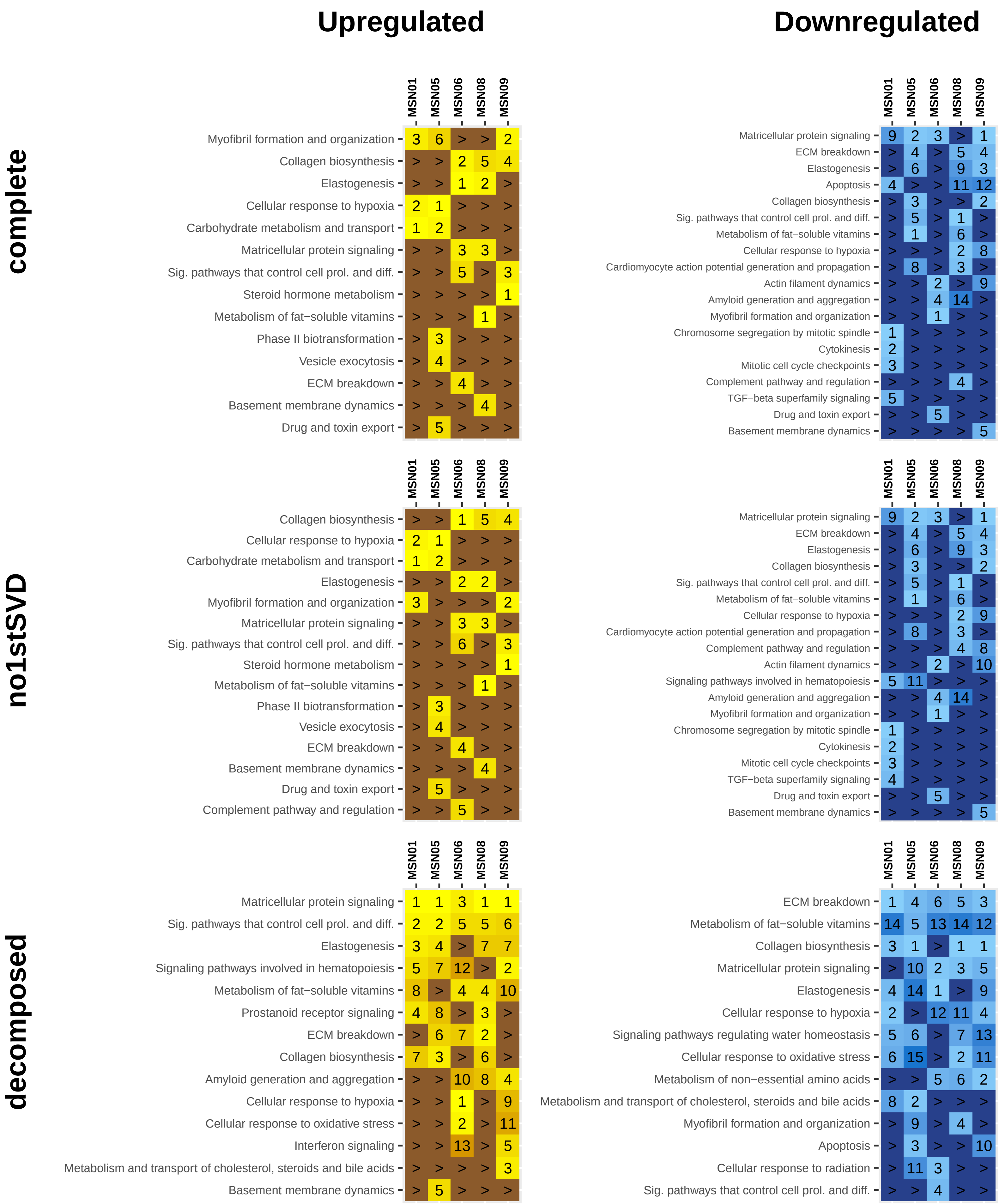

### MBCOL2
### insulin−like growth factor 1
#### (not cardiac act.)

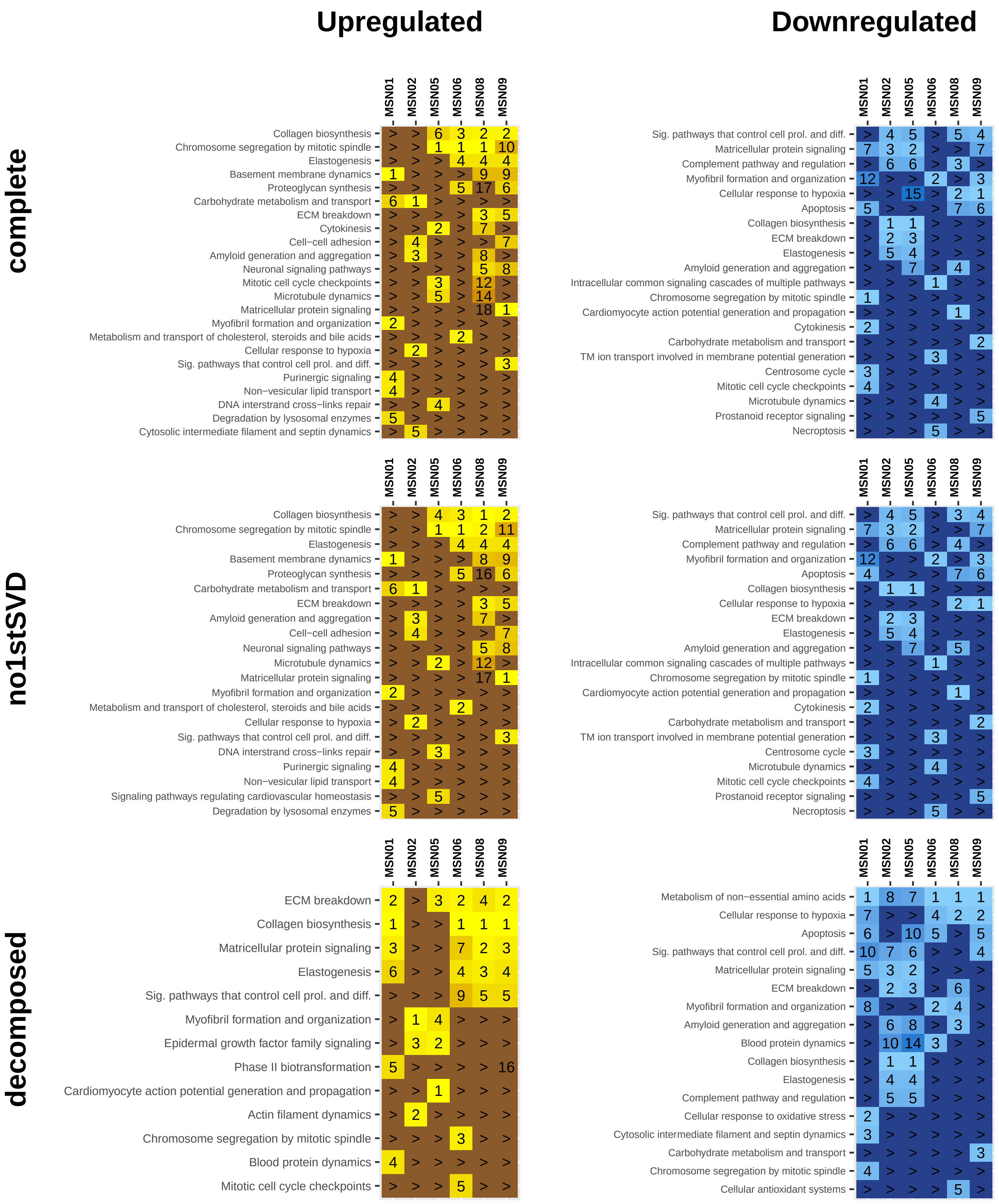

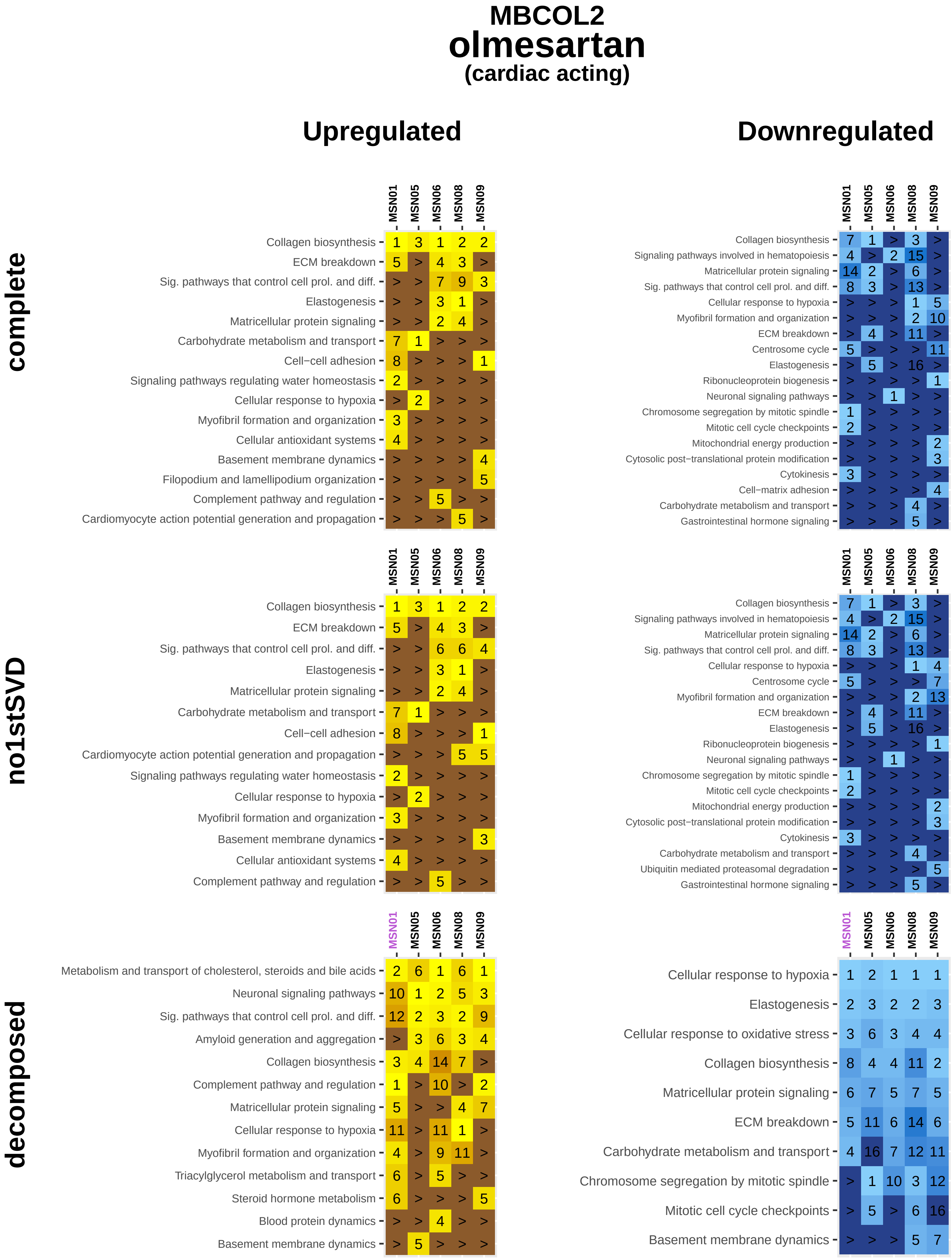
MBCOL2
olmesartan
(cardiac acting)
Upregulated
Downregulated
complete
no1stSVD
decomposed
MSN01 MSN05 MSN06 MSN08 MSN09
Collagen biosynthesis 1 3 1 2 2
ECM breakdown 5 > 4 3 >
Sig. pathways that control cell prol. and diff. > > 7 9 3
Elastogenesis > > 3 1 >
Matricellular protein signaling > > 2 4 >
Carbohydrate metabolism and transport 7 1 > > >
Cell−cell adhesion 8 > > > 1
Signaling pathways regulating water homeostasis 2 > > > >
Cellular response to hypoxia > 2 > > >
Myofibril formation and organization 3 > > > >
Cellular antioxidant systems 4 > > > >
Basement membrane dynamics > > > > 4
Filopodium and lamellipodium organization > > > > 5
Complement pathway and regulation > > 5 > >
Cardiomyocyte action potential generation and propagation > > > 5 >
Collagen biosynthesis 7 1 > 3 >
Signaling pathways involved in hematopoiesis 4 > 2 15 >
Matricellular protein signaling 14 2 > 6 >
Sig. pathways that control cell prol. and diff. 8 3 > 13 >
Cellular response to hypoxia > > > 1 5
Myofibril formation and organization > > > 2 10
ECM breakdown > 4 > 11 >
Centrosome cycle 5 > > > 11
Elastogenesis > 5 > 16 >
Ribonucleoprotein biogenesis > > > > 1
Neuronal signaling pathways > > 1 > >
Chromosome segregation by mitotic spindle 1 > > > >
Mitotic cell cycle checkpoints 2 > > > >
Mitochondrial energy production > > > > 2
Cytosolic post−translational protein modification > > > > 3
Cytokinesis 3 > > > >
Cell−matrix adhesion > > > > 4
Carbohydrate metabolism and transport > > > 4 >
Gastrointestinal hormone signaling > > > 5 >
Collagen biosynthesis 1 3 1 2 2
ECM breakdown 5 > 4 3 >
Sig. pathways that control cell prol. and diff. > > 6 6 4
Elastogenesis > > 3 1 >
Matricellular protein signaling > > 2 4 >
Carbohydrate metabolism and transport 7 1 > > >
Cell−cell adhesion 8 > > > 1
Cardiomyocyte action potential generation and propagation > > > 5 5
Signaling pathways regulating water homeostasis 2 > > > >
Cellular response to hypoxia > 2 > > >
Myofibril formation and organization 3 > > > >
Basement membrane dynamics > > > > 3
Cellular antioxidant systems 4 > > > >
Complement pathway and regulation > > 5 > >
Collagen biosynthesis 7 1 > 3 >
Signaling pathways involved in hematopoiesis 4 > 2 15 >
Matricellular protein signaling 14 2 > 6 >
Sig. pathways that control cell prol. and diff. 8 3 > 13 >
Cellular response to hypoxia > > > 1 4
Centrosome cycle 5 > > > 7
Myofibril formation and organization > > > 2 13
ECM breakdown > 4 > 11 >
Elastogenesis > 5 > 16 >
Ribonucleoprotein biogenesis > > > > 1
Neuronal signaling pathways > > 1 > >
Chromosome segregation by mitotic spindle 1 > > > >
Mitotic cell cycle checkpoints 2 > > > >
Mitochondrial energy production > > > > 2
Cytosolic post−translational protein modification > > > > 3
Cytokinesis 3 > > > >
Carbohydrate metabolism and transport > > > 4 >
Ubiquitin mediated proteasomal degradation > > > > 5
Gastrointestinal hormone signaling > > > 5 >
Metabolism and transport of cholesterol, steroids and bile acids 2 6 1 6 1
Neuronal signaling pathways 10 1 2 5 3
Sig. pathways that control cell prol. and diff. 12 2 3 2 9
Amyloid generation and aggregation > 3 6 3 4
Collagen biosynthesis 3 4 14 7 >
Complement pathway and regulation 1 > 10 > 2
Matricellular protein signaling 5 > > 4 7
Cellular response to hypoxia 11 > 11 1 >
Myofibril formation and organization 4 > 9 11 >
Triacylglycerol metabolism and transport 6 > 5 > >
Steroid hormone metabolism 6 > > > 5
Blood protein dynamics > > 4 > >
Basement membrane dynamics > 5 > > >
Cellular response to hypoxia 1 2 1 1 1
Elastogenesis 2 3 2 2 3
Cellular response to oxidative stress 3 6 3 4 4
Collagen biosynthesis 8 4 4 11 2
Matricellular protein signaling 6 7 5 7 5
ECM breakdown 5 11 6 14 6
Carbohydrate metabolism and transport 4 16 7 12 11
Chromosome segregation by mitotic spindle > 1 10 3 12
Mitotic cell cycle checkpoints > 5 > 6 16
Basement membrane dynamics > > > 5 7

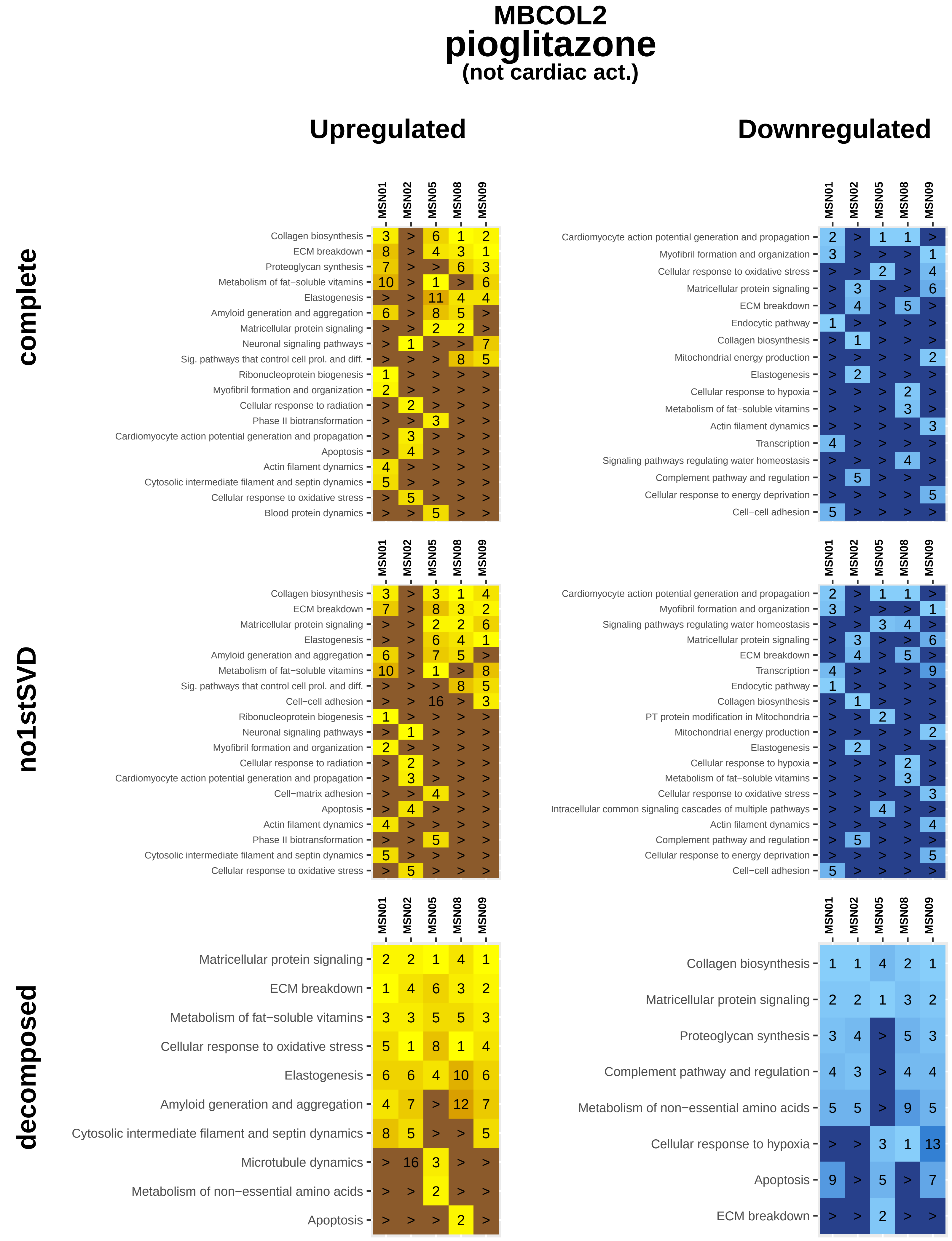
MBCOL2
pioglitazone
(not cardiac act.)
Upregulated
Downregulated
complete
no1stSVD
decomposed
MSN01
MSN02
MSN05
MSN08
MSN09
Collagen biosynthesis
ECM breakdown
Proteoglycan synthesis
Metabolism of fat−soluble vitamins
Elastogenesis
Amyloid generation and aggregation
Matricellular protein signaling
Neuronal signaling pathways
Sig. pathways that control cell prol. and diff.
Ribonucleoprotein biogenesis
Myofibril formation and organization
Cellular response to radiation
Phase II biotransformation
Cardiomyocyte action potential generation and propagation
Apoptosis
Actin filament dynamics
Cytosolic intermediate filament and septin dynamics
Cellular response to oxidative stress
Blood protein dynamics
Endocytic pathway
Mitochondrial energy production
Cellular response to hypoxia
Transcription
Signaling pathways regulating water homeostasis
Complement pathway and regulation
Cellular response to energy deprivation
Cell−cell adhesion
Cell−matrix adhesion
PT protein modification in Mitochondria
Intracellular common signaling cascades of multiple pathways
Microtubule dynamics
Metabolism of non−essential amino acids

**MBCOL2**

### prednisolone

**(not cardiac act.)**

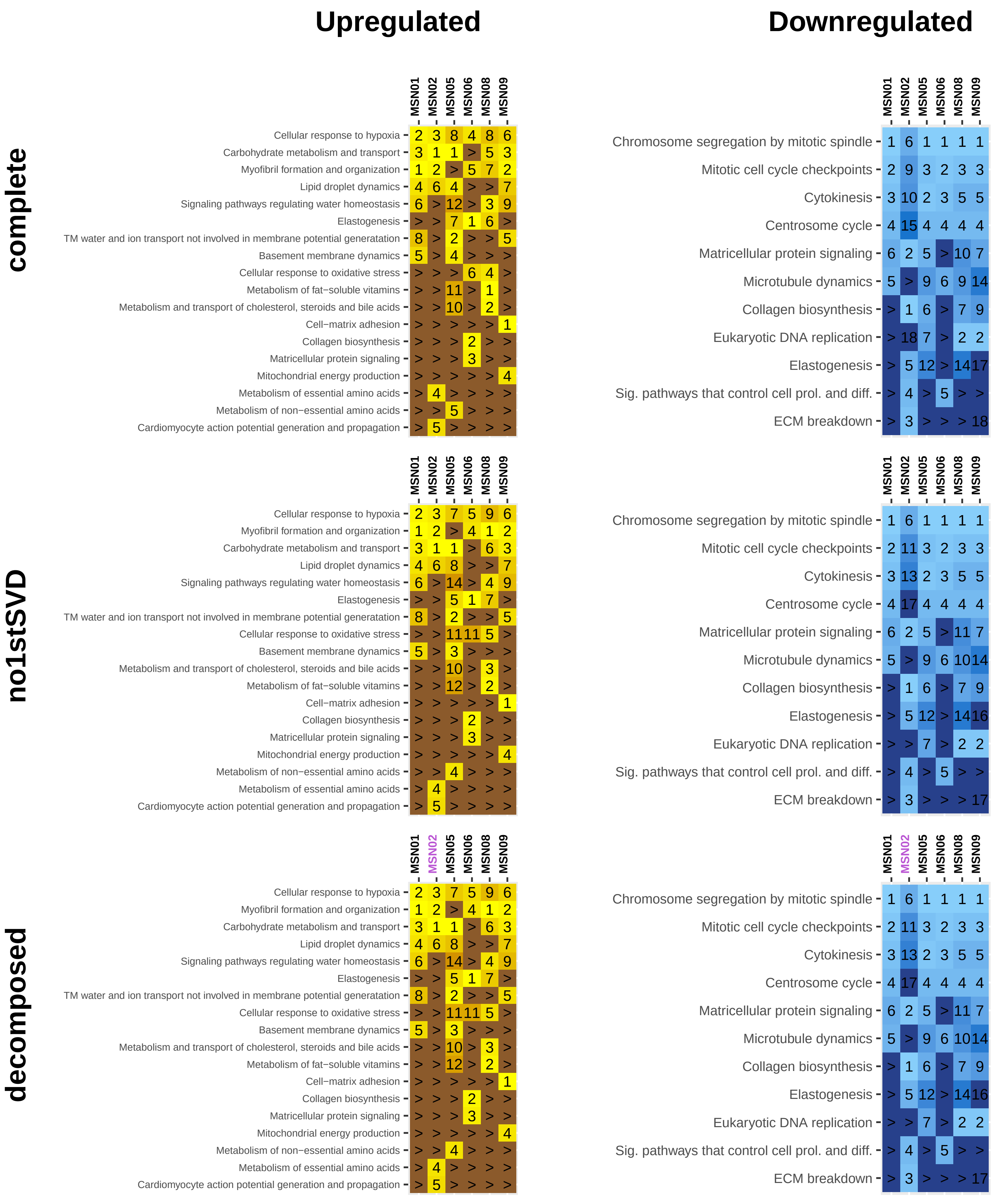

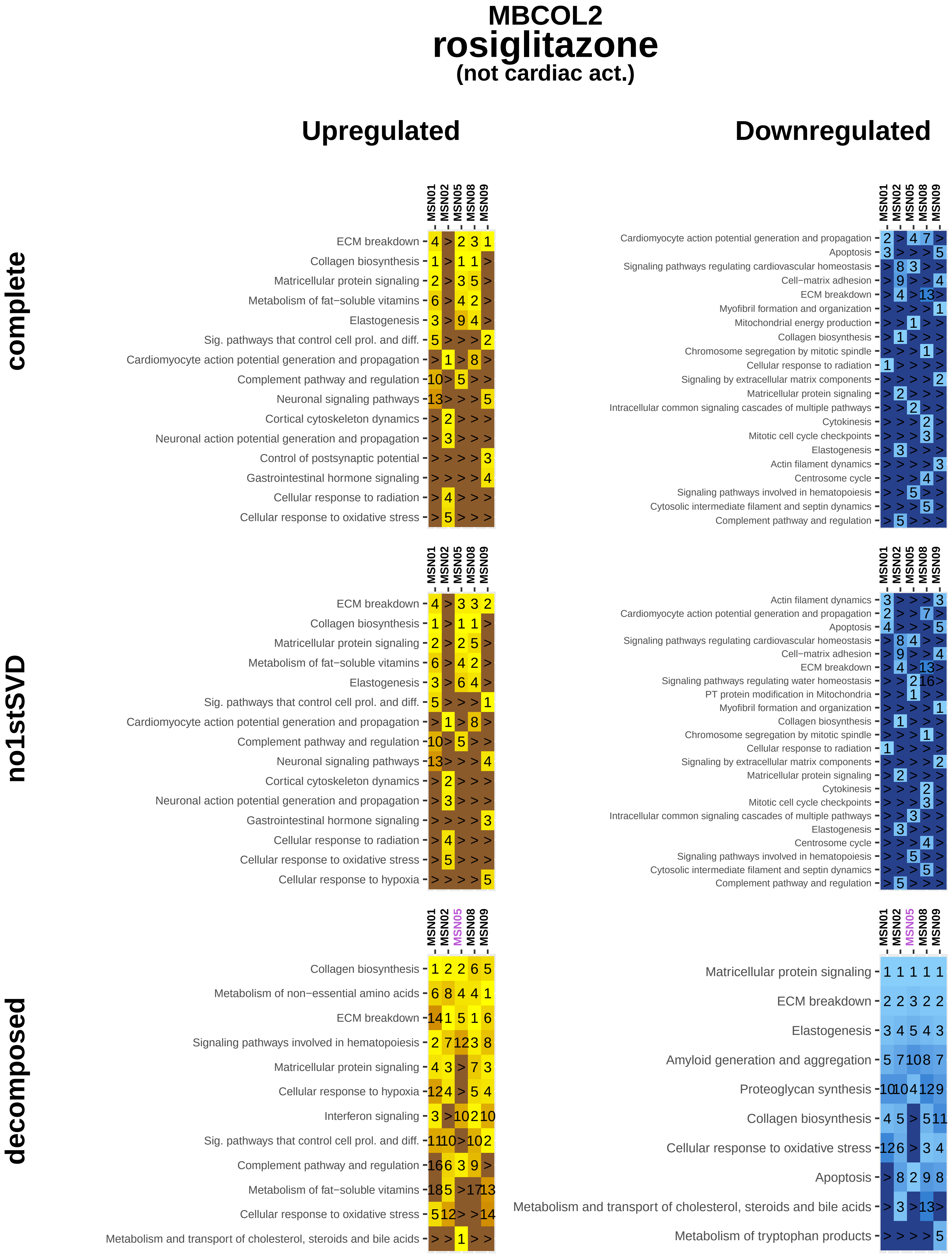
MBCOL2
rosiglitazone
(not cardiac act.)
Upregulated
Downregulated
complete
no1stSVD
decomposed
MSN01
MSN02
MSN05
MSN08
MSN09
ECM breakdown
Collagen biosynthesis
Matricellular protein signaling
Metabolism of fat−soluble vitamins
Elastogenesis
Sig. pathways that control cell prol. and diff.
Cardiomyocyte action potential generation and propagation
Complement pathway and regulation
Neuronal signaling pathways
Cortical cytoskeleton dynamics
Neuronal action potential generation and propagation
Control of postsynaptic potential
Gastrointestinal hormone signaling
Cellular response to radiation
Cellular response to oxidative stress
Apoptosis
Signaling pathways regulating cardiovascular homeostasis
Cell−matrix adhesion
Myofibril formation and organization
Mitochondrial energy production
Chromosome segregation by mitotic spindle
Signaling by extracellular matrix components
Intracellular common signaling cascades of multiple pathways
Cytokinesis
Mitotic cell cycle checkpoints
Actin filament dynamics
Centrosome cycle
Signaling pathways involved in hematopoiesis
Cytosolic intermediate filament and septin dynamics
Cellular response to hypoxia
Signaling pathways regulating water homeostasis
PT protein modification in Mitochondria
Metabolism of non−essential amino acids
Interferon signaling
Metabolism and transport of cholesterol, steroids and bile acids
Amyloid generation and aggregation
Proteoglycan synthesis
Metabolism of tryptophan products

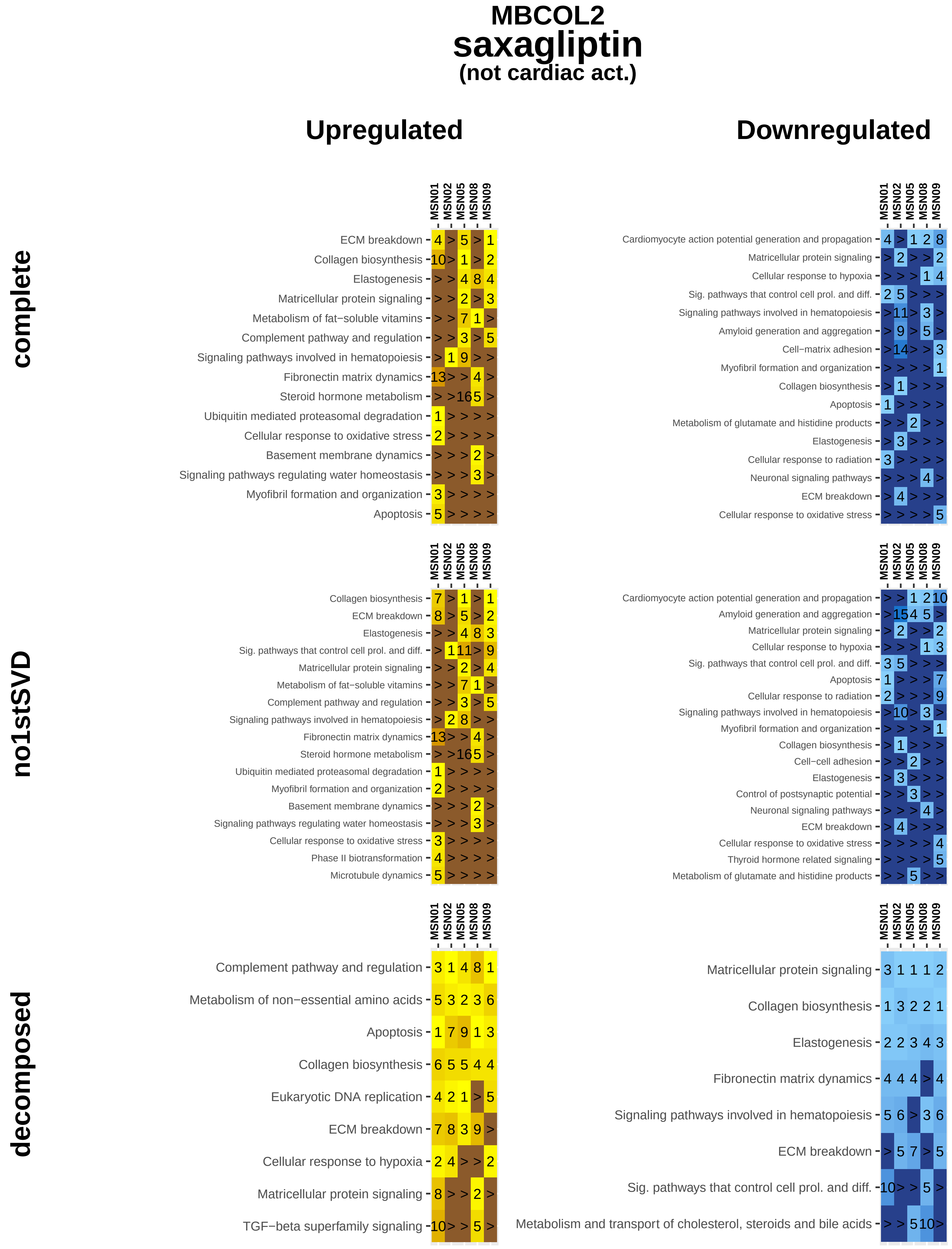
MBCOL2
saxagliptin
(not cardiac act.)
Upregulated
Downregulated
complete
no1stSVD
decomposed
MSN01
MSN02
MSN05
MSN08
MSN09
ECM breakdown
Collagen biosynthesis
Elastogenesis
Matricellular protein signaling
Metabolism of fat−soluble vitamins
Complement pathway and regulation
Signaling pathways involved in hematopoiesis
Fibronectin matrix dynamics
Steroid hormone metabolism
Ubiquitin mediated proteasomal degradation
Cellular response to oxidative stress
Basement membrane dynamics
Signaling pathways regulating water homeostasis
Myofibril formation and organization
Apoptosis
Cardiomyocyte action potential generation and propagation
Cellular response to hypoxia
Sig. pathways that control cell prol. and diff.
Amyloid generation and aggregation
Cell−matrix adhesion
Metabolism of glutamate and histidine products
Cellular response to radiation
Neuronal signaling pathways
Phase II biotransformation
Microtubule dynamics
Cell−cell adhesion
Control of postsynaptic potential
Thyroid hormone related signaling
Metabolism of non−essential amino acids
Eukaryotic DNA replication
TGF−beta superfamily signaling
Metabolism and transport of cholesterol, steroids and bile acids

### MBCOL2
### tnf−alpha
#### (not cardiac act.)

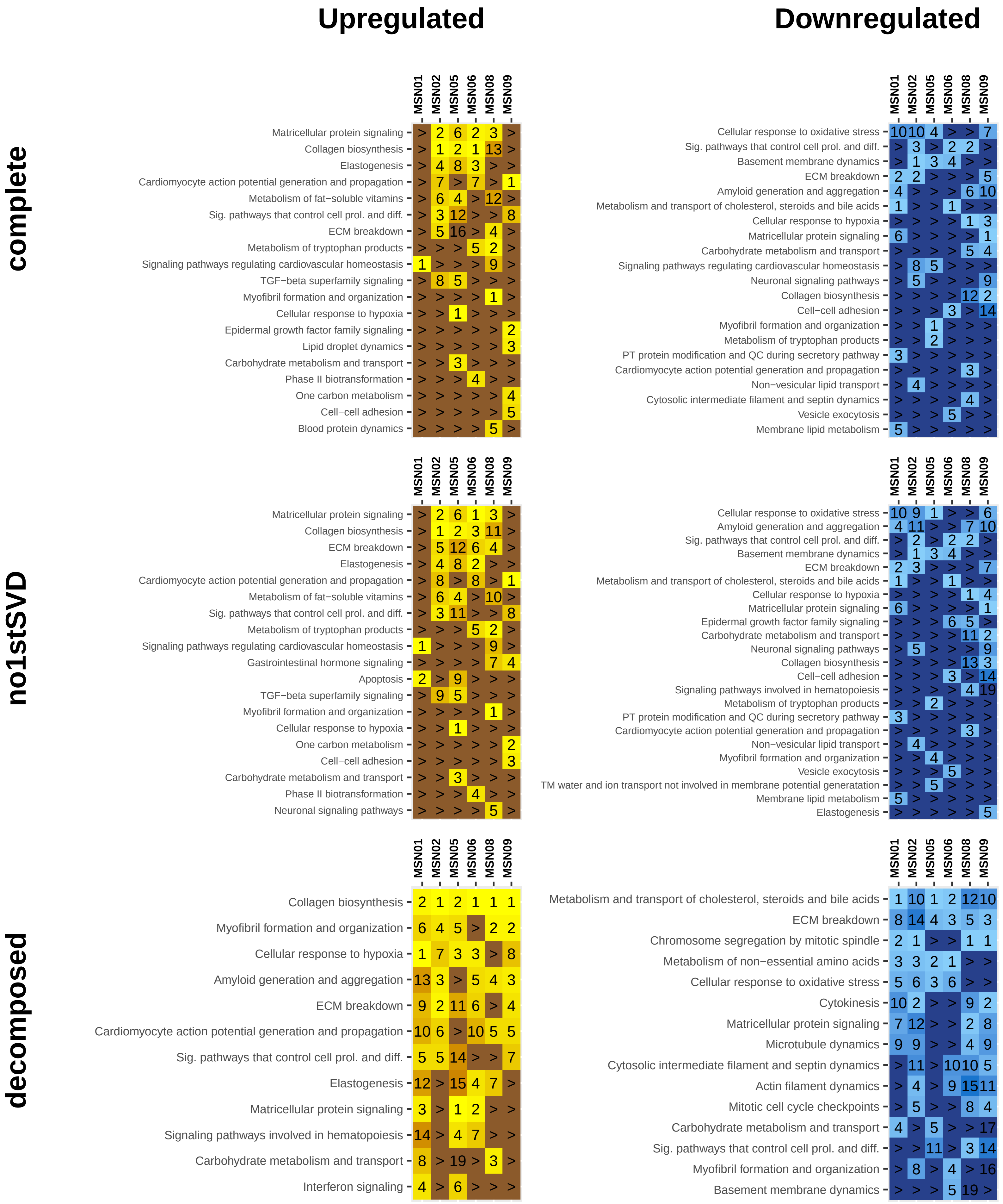
