## Supplemental Figure 14D for "Multiscale mapping of transcriptomic signatures for cardiotoxic drugs"

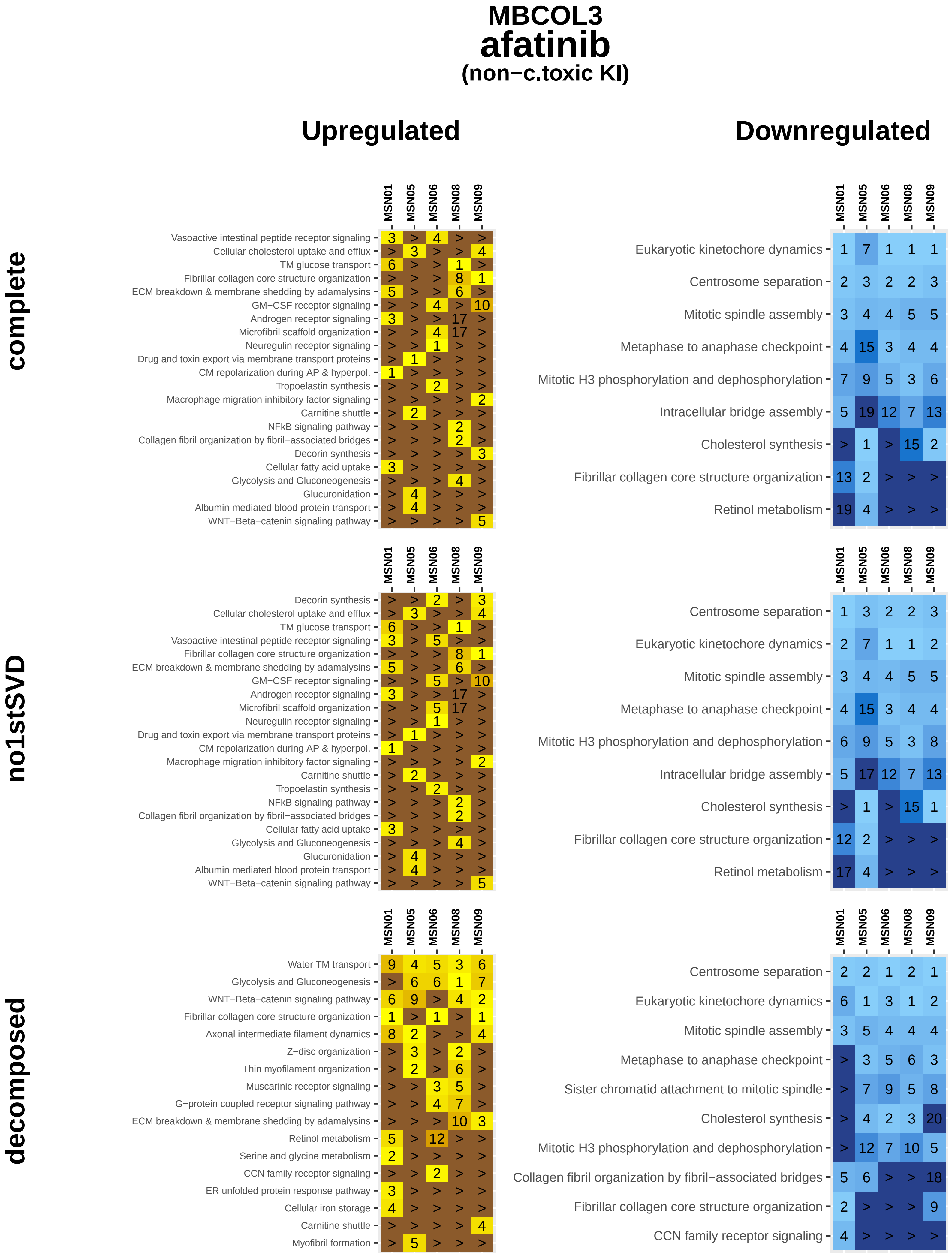
MBCOL3
afatinib
(non−c.toxic KI)
Upregulated
Downregulated
complete
no1stSVD
decomposed
MSN01
MSN05
MSN06
MSN08
MSN09
Vasoactive intestinal peptide receptor signaling
Cellular cholesterol uptake and efflux
TM glucose transport
Fibrillar collagen core structure organization
ECM breakdown & membrane shedding by adamalysins
GM−CSF receptor signaling
Androgen receptor signaling
Microfibril scaffold organization
Neuregulin receptor signaling
Drug and toxin export via membrane transport proteins
CM repolarization during AP & hyperpol.
Tropoelastin synthesis
Macrophage migration inhibitory factor signaling
Carnitine shuttle
NFkB signaling pathway
Collagen fibril organization by fibril−associated bridges
Decorin synthesis
Cellular fatty acid uptake
Glycolysis and Gluconeogenesis
Glucuronidation
Albumin mediated blood protein transport
WNT−Beta−catenin signaling pathway
Eukaryotic kinetochore dynamics
Centrosome separation
Mitotic spindle assembly
Metaphase to anaphase checkpoint
Mitotic H3 phosphorylation and dephosphorylation
Intracellular bridge assembly
Cholesterol synthesis
Retinol metabolism
Water TM transport
Axonal intermediate filament dynamics
Z−disc organization
Thin myofilament organization
Muscarinic receptor signaling
G−protein coupled receptor signaling pathway
Serine and glycine metabolism
CCN family receptor signaling
ER unfolded protein response pathway
Cellular iron storage
Myofibril formation
Sister chromatid attachment to mitotic spindle

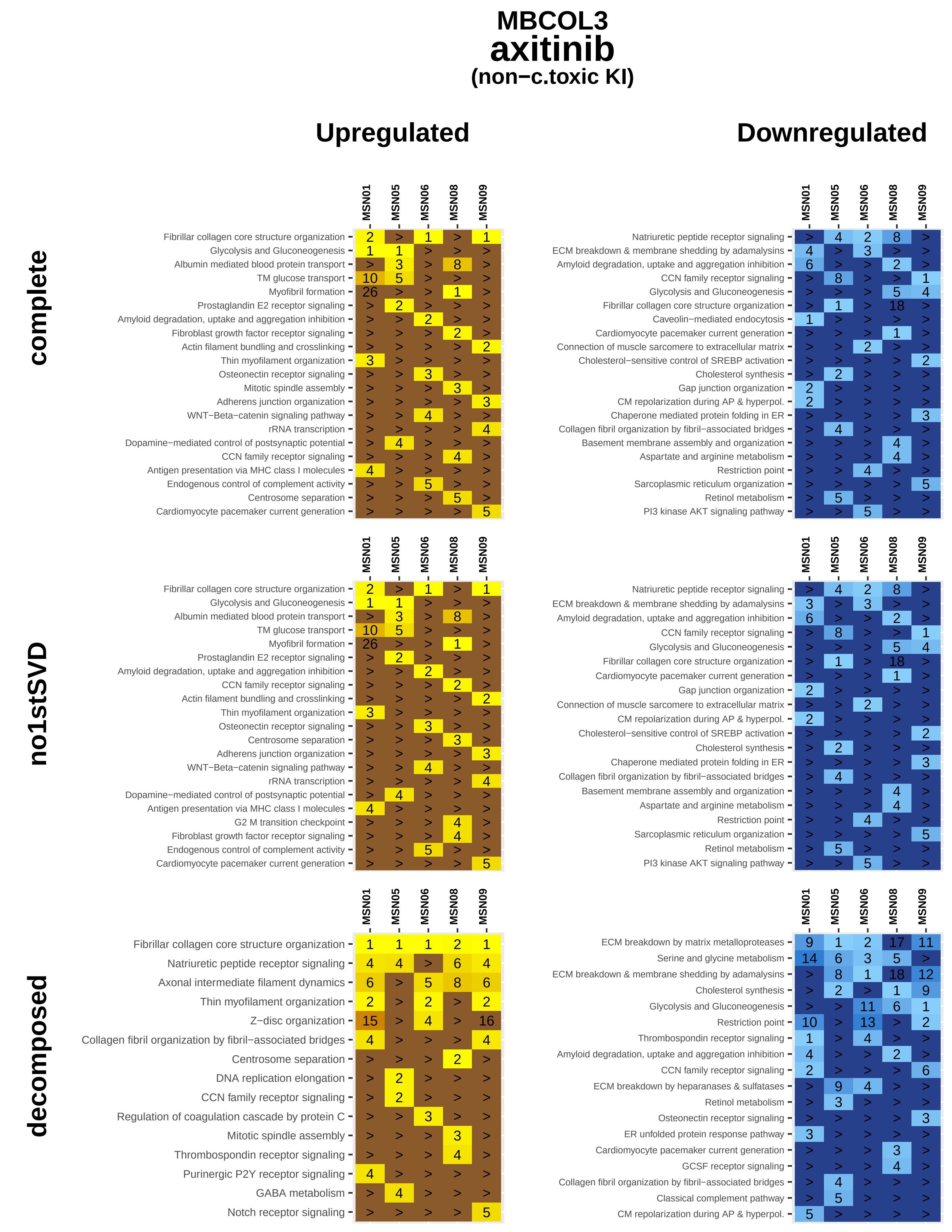

MBCOL3
axitinib
(non−c.toxic KI)
Upregulated
Downregulated
complete
no1stSVD
decomposed
MSN01
MSN05
MSN06
MSN08
MSN09
Fibrillar collagen core structure organization
Glycolysis and Gluconeogenesis
Albumin mediated blood protein transport
TM glucose transport
Myofibril formation
Prostaglandin E2 receptor signaling
Amyloid degradation, uptake and aggregation inhibition
Fibroblast growth factor receptor signaling
Actin filament bundling and crosslinking
Thin myofilament organization
Osteonectin receptor signaling
Mitotic spindle assembly
Adherens junction organization
WNT−Beta−catenin signaling pathway
rRNA transcription
Dopamine−mediated control of postsynaptic potential
CCN family receptor signaling
Antigen presentation via MHC class I molecules
Endogenous control of complement activity
Centrosome separation
Cardiomyocyte pacemaker current generation
Natriuretic peptide receptor signaling
ECM breakdown & membrane shedding by adamalysins
Amyloid degradation, uptake and aggregation inhibition
CCN family receptor signaling
Glycolysis and Gluconeogenesis
Fibrillar collagen core structure organization
Caveolin−mediated endocytosis
Cardiomyocyte pacemaker current generation
Connection of muscle sarcomere to extracellular matrix
Cholesterol−sensitive control of SREBP activation
Cholesterol synthesis
Gap junction organization
CM repolarization during AP & hyperpol.
Chaperone mediated protein folding in ER
Collagen fibril organization by fibril−associated bridges
Basement membrane assembly and organization
Aspartate and arginine metabolism
Restriction point
Sarcoplasmic reticulum organization
Retinol metabolism
PI3 kinase AKT signaling pathway
G2 M transition checkpoint
Axonal intermediate filament dynamics
Z−disc organization
DNA replication elongation
Regulation of coagulation cascade by protein C
Thrombospondin receptor signaling
Purinergic P2Y receptor signaling
GABA metabolism
Notch receptor signaling
ECM breakdown by matrix metalloproteases
Serine and glycine metabolism
ECM breakdown by heparanases & sulfatases
Osteonectin receptor signaling
ER unfolded protein response pathway
GCSF receptor signaling
Classical complement pathway

### MBCOL3
### bosutinib
#### (non−c.toxic KI)

#### complete

##### Upregulated

| | MSN01 | MSN05 | MSN08 | MSN09 |
|---|---|---|---|---|
| Cholesterol synthesis | 1 | 2 | > | 1 |
| Serine and glycine metabolism | 3 | 5 | 6 | > |
| Retinol metabolism | > | 3 | > | 12 |
| Fibrillar collagen core structure organization | > | > | 1 | > |
| Centrosome separation | > | 1 | > | > |
| Intrinsic apoptosis pathway | 2 | > | > | > |
| Eukaryotic kinetochore dynamics | > | > | > | 2 |
| Procollagen processing in the ER | > | > | 2 | > |
| Amyloid degradation, uptake and aggregation inhibition | > | > | 2 | > |
| Protein palmitoylation | > | > | > | 3 |
| Microtubule crosslinking and bundling | > | 4 | > | > |
| Large ribosomal subunit organization | 4 | > | > | > |
| ECM breakdown & membrane shedding by adamalysins | > | > | 4 | > |
| Small ribosomal subunit organization | 5 | > | > | > |
| Collagen fiber crosslinking | > | > | 5 | > |

##### Downregulated

| | MSN01 | MSN05 | MSN08 | MSN09 |
|---|---|---|---|---|
| Cellular cholesterol uptake and efflux | 4 | > | > | 4 |
| Vasoactive intestinal peptide receptor signaling | > | 1 | > | > |
| Progesterone receptor signaling | > | > | 1 | > |
| Glycolysis and Gluconeogenesis | > | > | > | 1 |
| ECM breakdown & membrane shedding by adamalysins | 1 | > | > | > |
| Histone methylation and demethylation | 2 | > | > | > |
| GM−CSF receptor signaling | > | > | 2 | > |
| Basement membrane attachment to cell surface | > | > | > | 2 |
| Fatty acid omega−hydroxylation | > | 2 | > | > |
| Decorin synthesis | > | 2 | > | > |
| Thrombopoietin receptor signaling | > | > | 3 | > |
| Semaphorin signaling | 3 | > | > | > |
| Cardiomyocyte pacemaker current generation | > | > | > | 3 |
| Thin myofilament organization | > | > | 4 | > |
| Intrinsic apoptosis pathway | > | 4 | > | > |
| Centriolar satelites organization | 4 | > | > | > |
| Sarcoplasmic reticulum organization | > | > | 5 | > |
| Inhibin receptor signaling | > | > | > | 5 |

#### no1stSVD

##### Upregulated

| | MSN01 | MSN05 | MSN08 | MSN09 |
|---|---|---|---|---|
| Cholesterol synthesis | 1 | 2 | > | 1 |
| Serine and glycine metabolism | 3 | 6 | 6 | > |
| Retinol metabolism | > | 4 | > | 12 |
| ECM breakdown & membrane shedding by adamalysins | > | 22 | 4 | > |
| Fibrillar collagen core structure organization | > | > | 1 | > |
| Centrosome separation | > | 1 | > | > |
| Intrinsic apoptosis pathway | 2 | > | > | > |
| Eukaryotic kinetochore dynamics | > | > | > | 2 |
| Procollagen processing in the ER | > | > | 2 | > |
| Amyloid degradation, uptake and aggregation inhibition | > | > | 2 | > |
| Protein palmitoylation | > | > | > | 3 |
| Inhibition of apoptosis | > | 3 | > | > |
| Large ribosomal subunit organization | 4 | > | > | > |
| Small ribosomal subunit organization | 5 | > | > | > |
| Microtubule crosslinking and bundling | > | 5 | > | > |
| Collagen fiber crosslinking | > | > | 5 | > |

##### Downregulated

| | MSN01 | MSN05 | MSN08 | MSN09 |
|---|---|---|---|---|
| Cellular cholesterol uptake and efflux | 4 | > | > | 5 |
| Hepatocyte growth factor receptor signaling | 8 | 4 | > | > |
| Vasoactive intestinal peptide receptor signaling | > | 1 | > | > |
| Progesterone receptor signaling | > | > | 1 | > |
| Glycolysis and Gluconeogenesis | > | > | > | 1 |
| ECM breakdown & membrane shedding by adamalysins | 1 | > | > | > |
| Histone methylation and demethylation | 2 | > | > | > |
| HIF−1 receptor signaling pathway | > | > | 2 | > |
| Fatty acid omega−hydroxylation | > | 2 | > | > |
| Basement membrane attachment to cell surface | > | > | > | 2 |
| Semaphorin signaling | 3 | > | > | > |
| Intrinsic apoptosis pathway | > | 3 | > | > |
| Electron transport chain | > | > | 3 | > |
| Cardiomyocyte pacemaker current generation | > | > | > | 3 |
| TM glucose transport | > | > | > | 4 |
| Insulin receptor signaling | > | > | 4 | > |
| Serotonin inactivation | > | 4 | > | > |
| Centriolar satelites organization | 4 | > | > | > |
| Glycogen synthesis and glycogenolysis | > | > | 5 | > |

#### decomposed

##### Upregulated

| | MSN01 | MSN05 | MSN08 | MSN09 |
|---|---|---|---|---|
| Natriuretic peptide receptor signaling | 1 | 1 | 2 | 1 |
| Thin myofilament organization | 2 | 2 | 1 | 5 |
| Osteonectin receptor signaling | 4 | > | 4 | 2 |
| Contractile ring constriction | 6 | 4 | > | 3 |
| Collagen fibril organization by fibril−associated bridges | > | 8 | 6 | 4 |
| Potassium TM transport | > | > | 3 | > |
| Elastin cross−linking and assembly | > | 3 | > | > |
| Actin filament bundling and crosslinking | 3 | > | > | > |
| Collagen fiber crosslinking | > | 4 | > | > |
| Z−disc organization | > | > | 5 | > |
| Sodium TM transport | 5 | > | > | > |

##### Downregulated

| | MSN01 | MSN05 | MSN08 | MSN09 |
|---|---|---|---|---|
| Fibrillar collagen core structure organization | 10 | 1 | 8 | 1 |
| Retinol metabolism | 12 | 12 | 4 | 14 |
| Cholesterol synthesis | 7 | 4 | 1 | > |
| Semaphorin signaling | 2 | > | 6 | 6 |
| WNT−Beta−catenin signaling pathway | > | 9 | 3 | 3 |
| VEGF receptor signaling | 5 | 2 | > | 8 |
| ECM breakdown by matrix metalloproteases | 8 | 3 | > | 10 |
| Alternative complement pathway | 1 | > | > | 4 |
| Amyloid degradation, uptake and aggregation inhibition | > | 5 | 2 | > |
| Non−vesicular phospholipid transport | 5 | > | > | 8 |
| Glutamate and glutamine metabolism | > | 12 | > | 2 |
| ECM breakdown by cathepsins | 3 | > | > | > |
| Thrombospondin receptor signaling | > | > | 5 | > |
| Collagen fibril organization by fibril−associated bridges | > | > | > | 5 |
| Classical complement pathway | 5 | > | > | > |

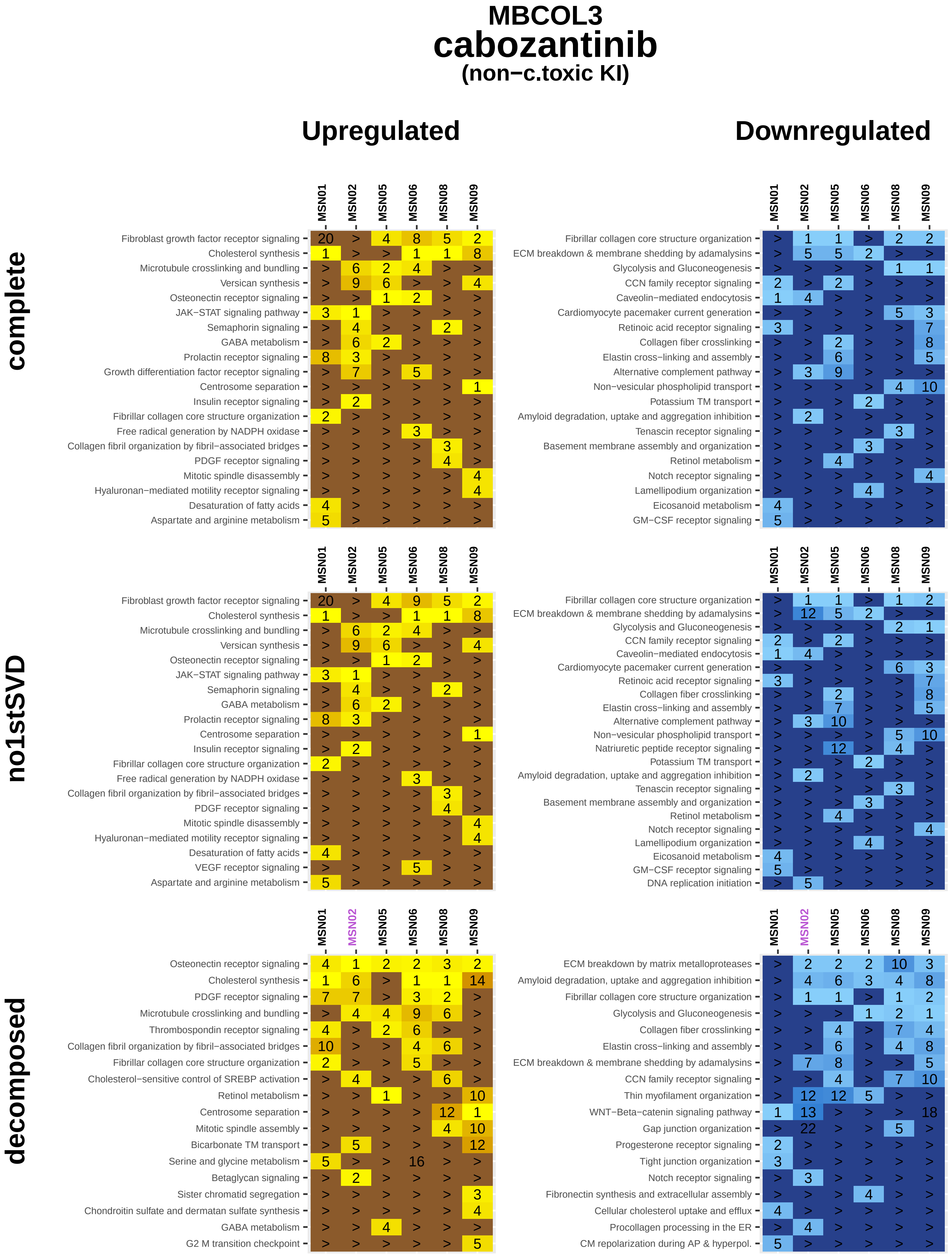
MBCOL3
cabozantinib
(non−c.toxic KI)
Upregulated
Downregulated
complete
no1stSVD
decomposed
MSN01
MSN02
MSN05
MSN06
MSN08
MSN09
Fibroblast growth factor receptor signaling
Cholesterol synthesis
Microtubule crosslinking and bundling
Versican synthesis
Osteonectin receptor signaling
JAK−STAT signaling pathway
Semaphorin signaling
GABA metabolism
Prolactin receptor signaling
Growth differentiation factor receptor signaling
Centrosome separation
Insulin receptor signaling
Fibrillar collagen core structure organization
Free radical generation by NADPH oxidase
Collagen fibril organization by fibril−associated bridges
PDGF receptor signaling
Mitotic spindle disassembly
Hyaluronan−mediated motility receptor signaling
Desaturation of fatty acids
Aspartate and arginine metabolism
Fibrillar collagen core structure organization
ECM breakdown & membrane shedding by adamalysins
Glycolysis and Gluconeogenesis
CCN family receptor signaling
Caveolin−mediated endocytosis
Cardiomyocyte pacemaker current generation
Retinoic acid receptor signaling
Collagen fiber crosslinking
Elastin cross−linking and assembly
Alternative complement pathway
Non−vesicular phospholipid transport
Potassium TM transport
Amyloid degradation, uptake and aggregation inhibition
Tenascin receptor signaling
Basement membrane assembly and organization
Retinol metabolism
Notch receptor signaling
Lamellipodium organization
Eicosanoid metabolism
GM−CSF receptor signaling
VEGF receptor signaling
Natriuretic peptide receptor signaling
DNA replication initiation
Thrombospondin receptor signaling
Cholesterol−sensitive control of SREBP activation
Mitotic spindle assembly
Bicarbonate TM transport
Serine and glycine metabolism
Betaglycan signaling
Sister chromatid segregation
Chondroitin sulfate and dermatan sulfate synthesis
G2 M transition checkpoint
ECM breakdown by matrix metalloproteases
Thin myofilament organization
WNT−Beta−catenin signaling pathway
Gap junction organization
Progesterone receptor signaling
Tight junction organization
Fibronectin synthesis and extracellular assembly
Cellular cholesterol uptake and efflux
Procollagen processing in the ER
CM repolarization during AP & hyperpol.

### MBCOL3
### ceritinib
#### (non−c.toxic KI)

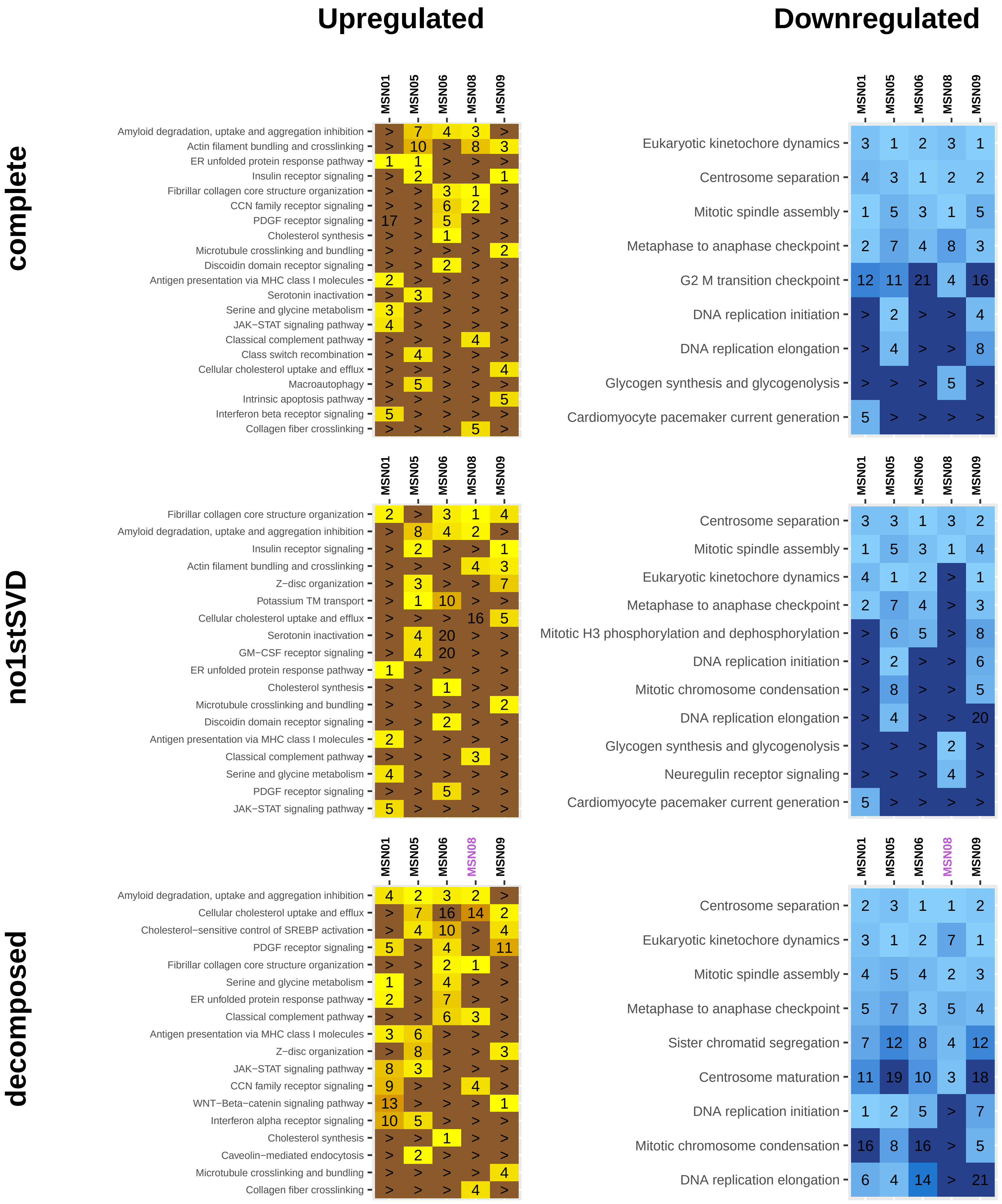

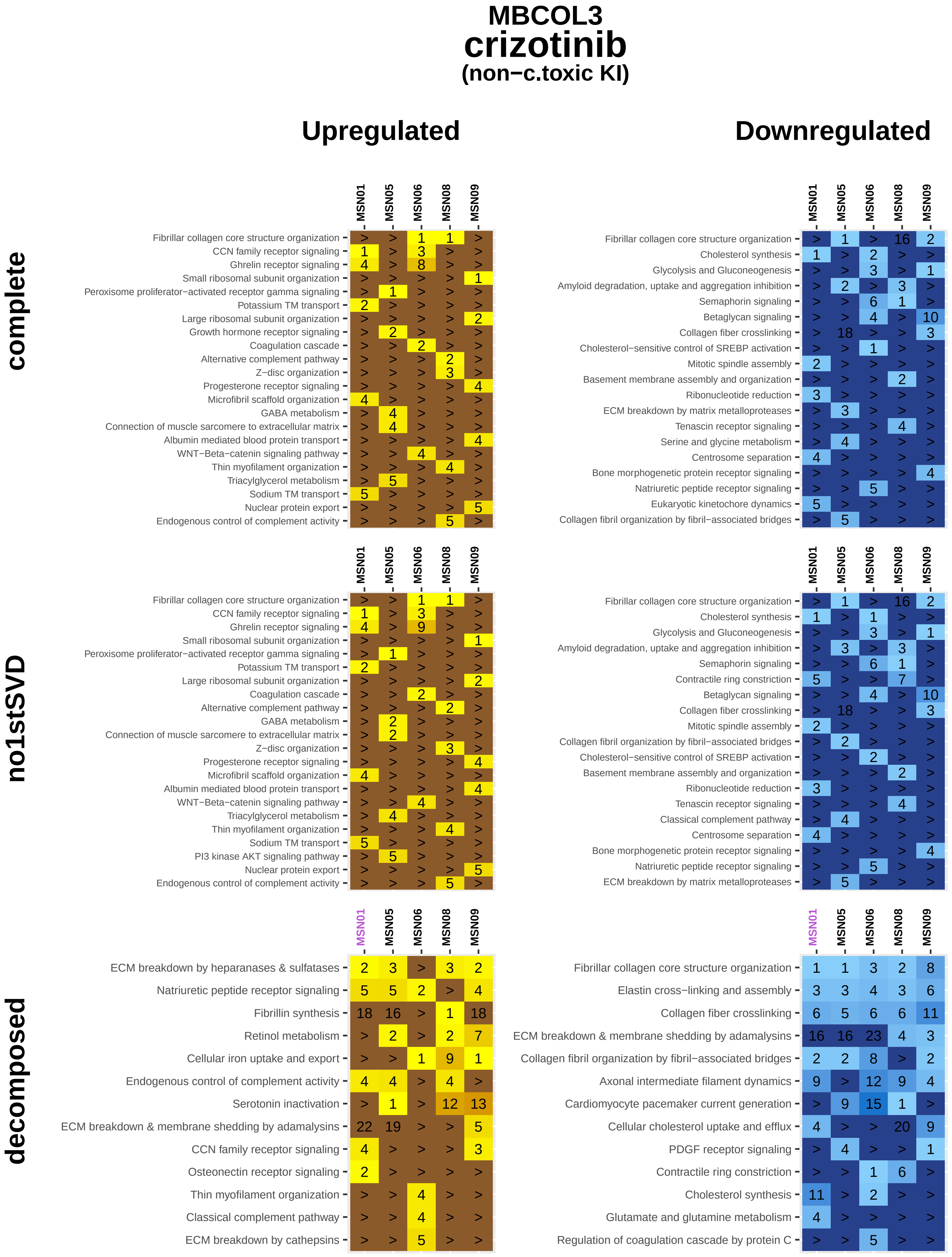

MBCOL3
crizotinib
(non−c.toxic KI)
Upregulated
Downregulated
complete
MSN01 MSN05 MSN06 MSN08 MSN09
Fibrillar collagen core structure organization > > 1 1 >
CCN family receptor signaling 1 > 3 > >
Ghrelin receptor signaling 4 > 8 > >
Small ribosomal subunit organization > > > > 1
Peroxisome proliferator−activated receptor gamma signaling > 1 > > >
Potassium TM transport 2 > > > >
Large ribosomal subunit organization > > > > 2
Growth hormone receptor signaling > 2 > > >
Coagulation cascade > > 2 > >
Alternative complement pathway > > > 2 >
Z−disc organization > > > 3 >
Progesterone receptor signaling > > > > 4
Microfibril scaffold organization 4 > > > >
GABA metabolism > 4 > > >
Connection of muscle sarcomere to extracellular matrix > 4 > > >
Albumin mediated blood protein transport > > > > 4
WNT−Beta−catenin signaling pathway > > 4 > >
Thin myofilament organization > > > 4 >
Triacylglycerol metabolism > 5 > > >
Sodium TM transport 5 > > > >
Nuclear protein export > > > > 5
Endogenous control of complement activity > > > 5 >
MSN01 MSN05 MSN06 MSN08 MSN09
Fibrillar collagen core structure organization > 1 > 16 2
Cholesterol synthesis 1 > 2 > >
Glycolysis and Gluconeogenesis > > 3 > 1
Amyloid degradation, uptake and aggregation inhibition > 2 > 3 >
Semaphorin signaling > > 6 1 >
Betaglycan signaling > > 4 > 10
Collagen fiber crosslinking > 18 > > 3
Cholesterol−sensitive control of SREBP activation > > 1 > >
Mitotic spindle assembly 2 > > > >
Basement membrane assembly and organization > > > 2 >
Ribonucleotide reduction 3 > > > >
ECM breakdown by matrix metalloproteases > 3 > > >
Tenascin receptor signaling > > > 4 >
Serine and glycine metabolism > 4 > > >
Centrosome separation 4 > > > >
Bone morphogenetic protein receptor signaling > > > > 4
Natriuretic peptide receptor signaling > > 5 > >
Eukaryotic kinetochore dynamics 5 > > > >
Collagen fibril organization by fibril−associated bridges > 5 > > >
no1stSVD
MSN01 MSN05 MSN06 MSN08 MSN09
Fibrillar collagen core structure organization > > 1 1 >
CCN family receptor signaling 1 > 3 > >
Ghrelin receptor signaling 4 > 9 > >
Small ribosomal subunit organization > > > > 1
Peroxisome proliferator−activated receptor gamma signaling > 1 > > >
Potassium TM transport 2 > > > >
Large ribosomal subunit organization > > > > 2
Coagulation cascade > > 2 > >
Alternative complement pathway > > > 2 >
GABA metabolism > 2 > > >
Connection of muscle sarcomere to extracellular matrix > 2 > > >
Z−disc organization > > > 3 >
Progesterone receptor signaling > > > > 4
Microfibril scaffold organization 4 > > > >
Albumin mediated blood protein transport > > > > 4
WNT−Beta−catenin signaling pathway > > 4 > >
Triacylglycerol metabolism > 4 > > >
Thin myofilament organization > > > 4 >
Sodium TM transport 5 > > > >
PI3 kinase AKT signaling pathway > 5 > > >
Nuclear protein export > > > > 5
Endogenous control of complement activity > > > 5 >
MSN01 MSN05 MSN06 MSN08 MSN09
Fibrillar collagen core structure organization > 1 > 16 2
Cholesterol synthesis 1 > 1 > >
Glycolysis and Gluconeogenesis > > 3 > 1
Amyloid degradation, uptake and aggregation inhibition > 3 > 3 >
Semaphorin signaling > > 6 1 >
Contractile ring constriction 5 > > 7 >
Betaglycan signaling > > 4 > 10
Collagen fiber crosslinking > 18 > > 3
Mitotic spindle assembly 2 > > > >
Collagen fibril organization by fibril−associated bridges > 2 > > >
Cholesterol−sensitive control of SREBP activation > > 2 > >
Basement membrane assembly and organization > > > 2 >
Ribonucleotide reduction 3 > > > >
Tenascin receptor signaling > > > 4 >
Classical complement pathway > 4 > > >
Centrosome separation 4 > > > >
Bone morphogenetic protein receptor signaling > > > > 4
Natriuretic peptide receptor signaling > > 5 > >
ECM breakdown by matrix metalloproteases > 5 > > >
decomposed
MSN01 MSN05 MSN06 MSN08 MSN09
ECM breakdown by heparanases & sulfatases 2 3 > 3 2
Natriuretic peptide receptor signaling 5 5 2 > 4
Fibrillin synthesis 18 16 > 1 18
Retinol metabolism > 2 > 2 7
Cellular iron uptake and export > > 1 9 1
Endogenous control of complement activity 4 4 > 4 >
Serotonin inactivation > 1 > 12 13
ECM breakdown & membrane shedding by adamalysins 22 19 > > 5
CCN family receptor signaling 4 > > > 3
Osteonectin receptor signaling 2 > > > >
Thin myofilament organization > > 4 > >
Classical complement pathway > > 4 > >
ECM breakdown by cathepsins > > 5 > >
MSN01 MSN05 MSN06 MSN08 MSN09
Fibrillar collagen core structure organization 1 1 3 2 8
Elastin cross−linking and assembly 3 3 4 3 6
Collagen fiber crosslinking 6 5 6 6 11
ECM breakdown & membrane shedding by adamalysins 16 16 23 4 3
Collagen fibril organization by fibril−associated bridges 2 2 8 > 2
Axonal intermediate filament dynamics 9 > 12 9 4
Cardiomyocyte pacemaker current generation > 9 15 1 >
Cellular cholesterol uptake and efflux 4 > > 20 9
PDGF receptor signaling > 4 > > 1
Contractile ring constriction > > 1 6 >
Cholesterol synthesis 11 > 2 > >
Glutamate and glutamine metabolism 4 > > > >
Regulation of coagulation cascade by protein C > > 5 > >

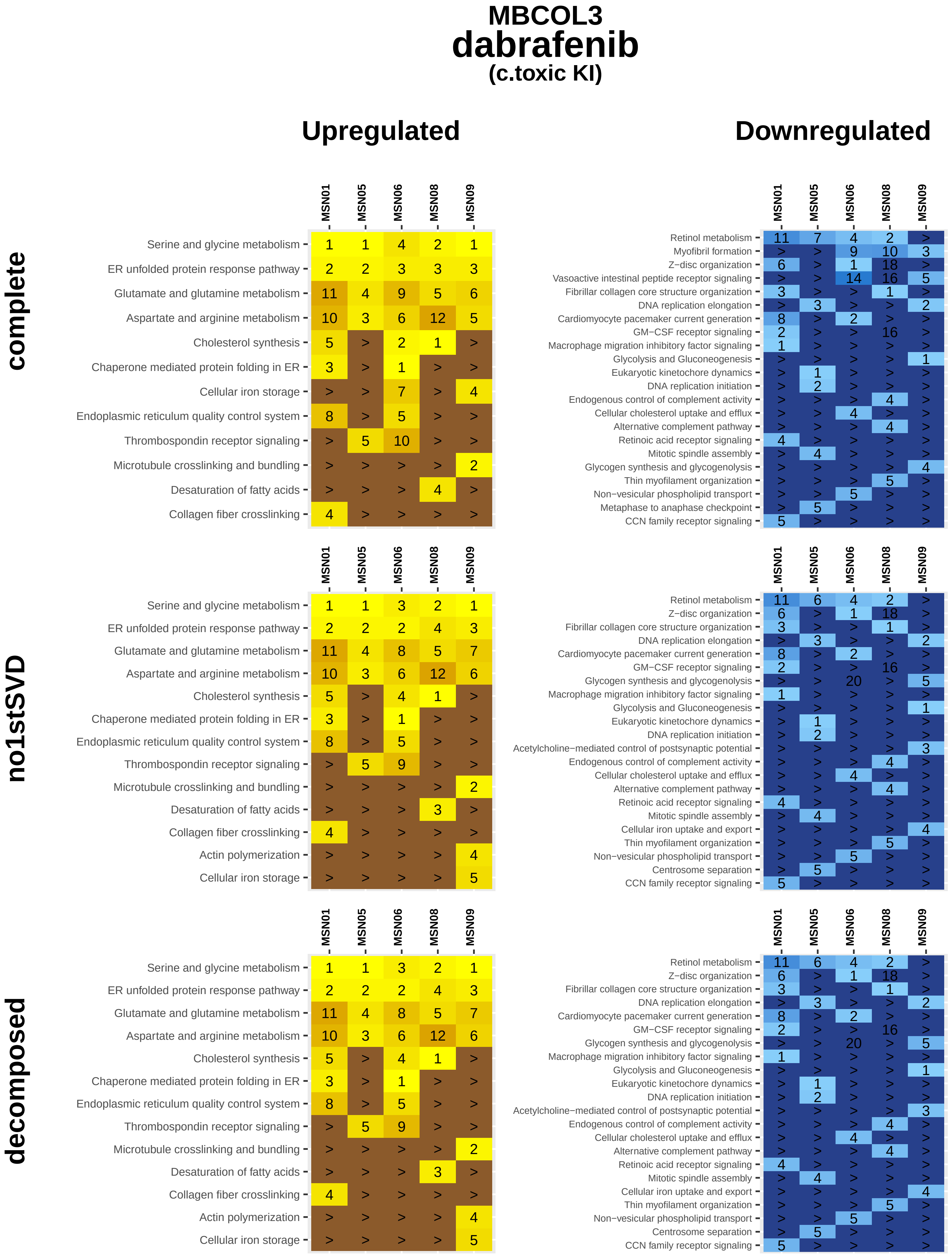
MBCOL3
dabrafenib
(c.toxic KI)
Upregulated
Downregulated
complete
MSN01 MSN05 MSN06 MSN08 MSN09
Serine and glycine metabolism 1 1 4 2 1
ER unfolded protein response pathway 2 2 3 3 3
Glutamate and glutamine metabolism 11 4 9 5 6
Aspartate and arginine metabolism 10 3 6 12 5
Cholesterol synthesis 5 > 2 1 >
Chaperone mediated protein folding in ER 3 > 1 > >
Cellular iron storage > > 7 > 4
Endoplasmic reticulum quality control system 8 > 5 > >
Thrombospondin receptor signaling > 5 10 > >
Microtubule crosslinking and bundling > > > > 2
Desaturation of fatty acids > > > 4 >
Collagen fiber crosslinking 4 > > > >
MSN01 MSN05 MSN06 MSN08 MSN09
Retinol metabolism 11 7 4 2 >
Myofibril formation > > 9 10 3
Z−disc organization 6 > 1 18 >
Vasoactive intestinal peptide receptor signaling > > 14 16 5
Fibrillar collagen core structure organization 3 > > 1 >
DNA replication elongation > 3 > > 2
Cardiomyocyte pacemaker current generation 8 > 2 > >
GM−CSF receptor signaling 2 > > 16 >
Macrophage migration inhibitory factor signaling 1 > > > >
Glycolysis and Gluconeogenesis > > > > 1
Eukaryotic kinetochore dynamics > 1 > > >
DNA replication initiation > 2 > > >
Endogenous control of complement activity > > > 4 >
Cellular cholesterol uptake and efflux > > 4 > >
Alternative complement pathway > > > 4 >
Retinoic acid receptor signaling 4 > > > >
Mitotic spindle assembly > 4 > > >
Glycogen synthesis and glycogenolysis > > > > 4
Thin myofilament organization > > > 5 >
Non−vesicular phospholipid transport > > 5 > >
Metaphase to anaphase checkpoint > 5 > > >
CCN family receptor signaling 5 > > > >
no1stSVD
MSN01 MSN05 MSN06 MSN08 MSN09
Serine and glycine metabolism 1 1 3 2 1
ER unfolded protein response pathway 2 2 2 4 3
Glutamate and glutamine metabolism 11 4 8 5 7
Aspartate and arginine metabolism 10 3 6 12 6
Cholesterol synthesis 5 > 4 1 >
Chaperone mediated protein folding in ER 3 > 1 > >
Endoplasmic reticulum quality control system 8 > 5 > >
Thrombospondin receptor signaling > 5 9 > >
Microtubule crosslinking and bundling > > > > 2
Desaturation of fatty acids > > > 3 >
Collagen fiber crosslinking 4 > > > >
Actin polymerization > > > > 4
Cellular iron storage > > > > 5
MSN01 MSN05 MSN06 MSN08 MSN09
Retinol metabolism 11 6 4 2 >
Z−disc organization 6 > 1 18 >
Fibrillar collagen core structure organization 3 > > 1 >
DNA replication elongation > 3 > > 2
Cardiomyocyte pacemaker current generation 8 > 2 > >
GM−CSF receptor signaling 2 > > 16 >
Glycogen synthesis and glycogenolysis > > 20 > 5
Macrophage migration inhibitory factor signaling 1 > > > >
Glycolysis and Gluconeogenesis > > > > 1
Eukaryotic kinetochore dynamics > 1 > > >
DNA replication initiation > 2 > > >
Acetylcholine−mediated control of postsynaptic potential > > > > 3
Endogenous control of complement activity > > > 4 >
Cellular cholesterol uptake and efflux > > 4 > >
Alternative complement pathway > > > 4 >
Retinoic acid receptor signaling 4 > > > >
Mitotic spindle assembly > 4 > > >
Cellular iron uptake and export > > > > 4
Thin myofilament organization > > > 5 >
Non−vesicular phospholipid transport > > 5 > >
Centrosome separation > 5 > > >
CCN family receptor signaling 5 > > > >
decomposed
MSN01 MSN05 MSN06 MSN08 MSN09
Serine and glycine metabolism 1 1 3 2 1
ER unfolded protein response pathway 2 2 2 4 3
Glutamate and glutamine metabolism 11 4 8 5 7
Aspartate and arginine metabolism 10 3 6 12 6
Cholesterol synthesis 5 > 4 1 >
Chaperone mediated protein folding in ER 3 > 1 > >
Endoplasmic reticulum quality control system 8 > 5 > >
Thrombospondin receptor signaling > 5 9 > >
Microtubule crosslinking and bundling > > > > 2
Desaturation of fatty acids > > > 3 >
Collagen fiber crosslinking 4 > > > >
Actin polymerization > > > > 4
Cellular iron storage > > > > 5
MSN01 MSN05 MSN06 MSN08 MSN09
Retinol metabolism 11 6 4 2 >
Z−disc organization 6 > 1 18 >
Fibrillar collagen core structure organization 3 > > 1 >
DNA replication elongation > 3 > > 2
Cardiomyocyte pacemaker current generation 8 > 2 > >
GM−CSF receptor signaling 2 > > 16 >
Glycogen synthesis and glycogenolysis > > 20 > 5
Macrophage migration inhibitory factor signaling 1 > > > >
Glycolysis and Gluconeogenesis > > > > 1
Eukaryotic kinetochore dynamics > 1 > > >
DNA replication initiation > 2 > > >
Acetylcholine−mediated control of postsynaptic potential > > > > 3
Endogenous control of complement activity > > > 4 >
Cellular cholesterol uptake and efflux > > 4 > >
Alternative complement pathway > > > 4 >
Retinoic acid receptor signaling 4 > > > >
Mitotic spindle assembly > 4 > > >
Cellular iron uptake and export > > > > 4
Thin myofilament organization > > > 5 >
Non−vesicular phospholipid transport > > 5 > >
Centrosome separation > 5 > > >
CCN family receptor signaling 5 > > > >

### MBCOL3
### dasatinib
#### (non−c.toxic KI)

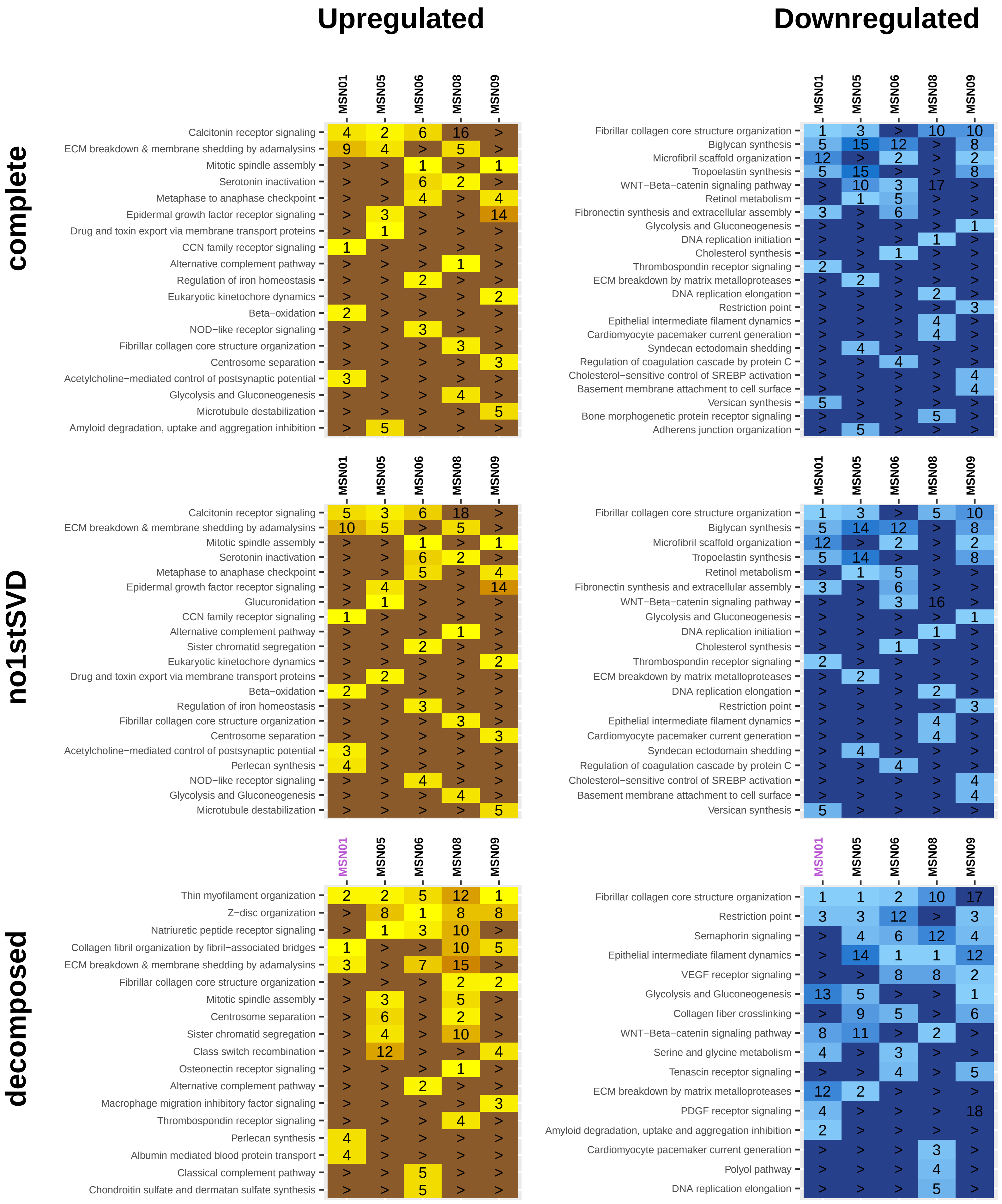

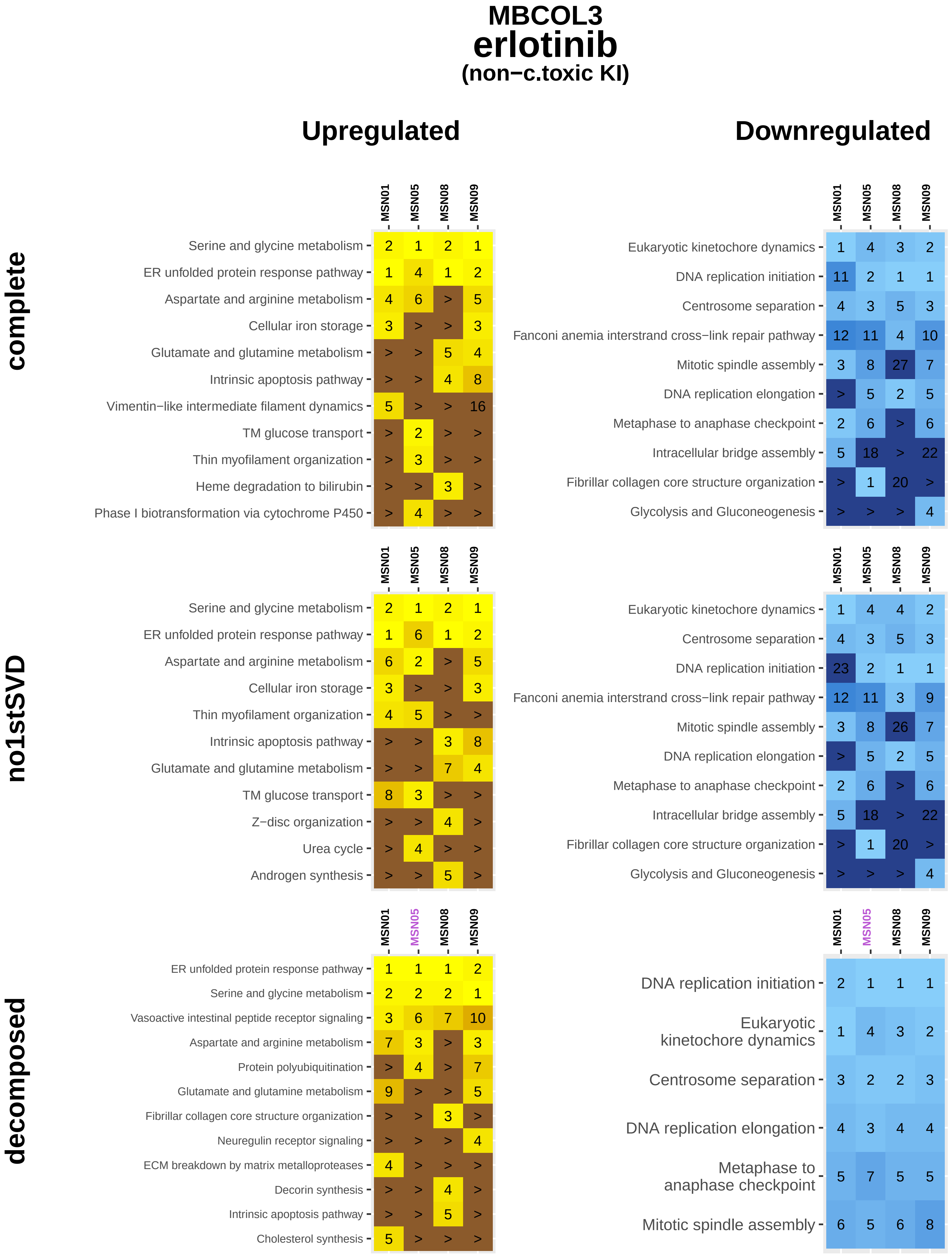
MBCOL3
erlotinib
(non−c.toxic KI)
Upregulated
Downregulated
complete
no1stSVD
decomposed
MSN01
MSN05
MSN08
MSN09
Serine and glycine metabolism 2 1 2 1
ER unfolded protein response pathway 1 4 1 2
Aspartate and arginine metabolism 4 6 > 5
Cellular iron storage 3 > > 3
Glutamate and glutamine metabolism > > 5 4
Intrinsic apoptosis pathway > > 4 8
Vimentin−like intermediate filament dynamics 5 > > 16
TM glucose transport > 2 > >
Thin myofilament organization > 3 > >
Heme degradation to bilirubin > > 3 >
Phase I biotransformation via cytochrome P450 > 4 > >
Eukaryotic kinetochore dynamics 1 4 3 2
DNA replication initiation 11 2 1 1
Centrosome separation 4 3 5 3
Fanconi anemia interstrand cross−link repair pathway 12 11 4 10
Mitotic spindle assembly 3 8 27 7
DNA replication elongation > 5 2 5
Metaphase to anaphase checkpoint 2 6 > 6
Intracellular bridge assembly 5 18 > 22
Fibrillar collagen core structure organization > 1 20 >
Glycolysis and Gluconeogenesis > > > 4
Serine and glycine metabolism 2 1 2 1
ER unfolded protein response pathway 1 6 1 2
Aspartate and arginine metabolism 6 2 > 5
Cellular iron storage 3 > > 3
Thin myofilament organization 4 5 > >
Intrinsic apoptosis pathway > > 3 8
Glutamate and glutamine metabolism > > 7 4
TM glucose transport 8 3 > >
Z−disc organization > > 4 >
Urea cycle > 4 > >
Androgen synthesis > > 5 >
Eukaryotic kinetochore dynamics 1 4 4 2
Centrosome separation 4 3 5 3
DNA replication initiation 23 2 1 1
Fanconi anemia interstrand cross−link repair pathway 12 11 3 9
Mitotic spindle assembly 3 8 26 7
DNA replication elongation > 5 2 5
Metaphase to anaphase checkpoint 2 6 > 6
Intracellular bridge assembly 5 18 > 22
Fibrillar collagen core structure organization > 1 20 >
Glycolysis and Gluconeogenesis > > > 4
ER unfolded protein response pathway 1 1 1 2
Serine and glycine metabolism 2 2 2 1
Vasoactive intestinal peptide receptor signaling 3 6 7 10
Aspartate and arginine metabolism 7 3 > 3
Protein polyubiquitination > 4 > 7
Glutamate and glutamine metabolism 9 > > 5
Fibrillar collagen core structure organization > > 3 >
Neuregulin receptor signaling > > > 4
ECM breakdown by matrix metalloproteases 4 > > >
Decorin synthesis > > 4 >
Intrinsic apoptosis pathway > > 5 >
Cholesterol synthesis 5 > > >
DNA replication initiation 2 1 1 1
Eukaryotic kinetochore dynamics 1 4 3 2
Centrosome separation 3 2 2 3
DNA replication elongation 4 3 4 4
Metaphase to anaphase checkpoint 5 7 5 5
Mitotic spindle assembly 6 5 6 8

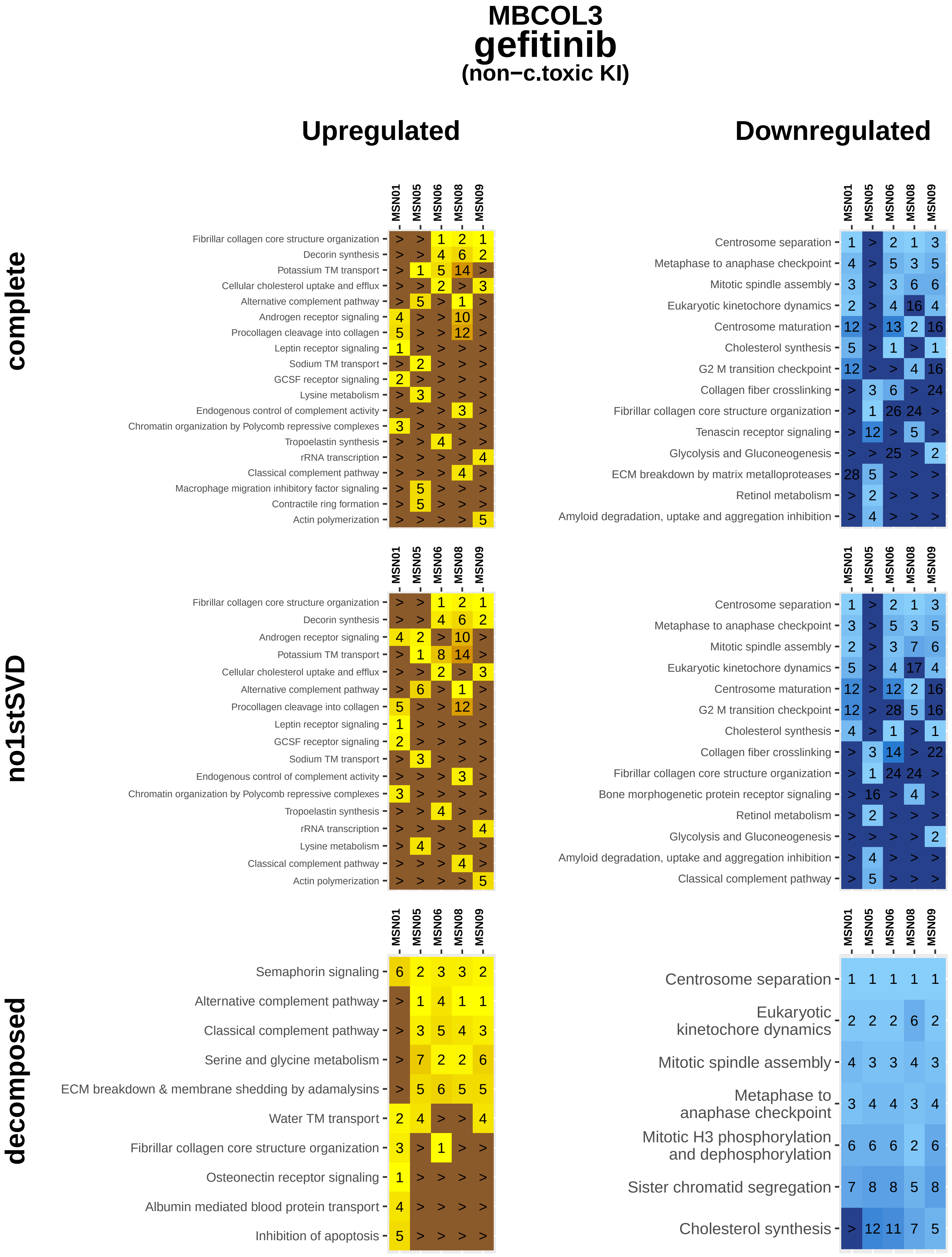

MBCOL3
gefitinib
(non−c.toxic KI)
Upregulated
Downregulated
complete
no1stSVD
decomposed
MSN01 MSN05 MSN06 MSN08 MSN09
Fibrillar collagen core structure organization > > 1 2 1
Decorin synthesis > > 4 6 2
Potassium TM transport > 1 5 14 >
Cellular cholesterol uptake and efflux > > 2 > 3
Alternative complement pathway > 5 > 1 >
Androgen receptor signaling 4 > > 10 >
Procollagen cleavage into collagen 5 > > 12 >
Leptin receptor signaling 1 > > > >
Sodium TM transport > 2 > > >
GCSF receptor signaling 2 > > > >
Lysine metabolism > 3 > > >
Endogenous control of complement activity > > > 3 >
Chromatin organization by Polycomb repressive complexes 3 > > > >
Tropoelastin synthesis > > 4 > >
rRNA transcription > > > > 4
Classical complement pathway > > > 4 >
Macrophage migration inhibitory factor signaling > 5 > > >
Contractile ring formation > 5 > > >
Actin polymerization > > > > 5
MSN01 MSN05 MSN06 MSN08 MSN09
Centrosome separation 1 > 2 1 3
Metaphase to anaphase checkpoint 4 > 5 3 5
Mitotic spindle assembly 3 > 3 6 6
Eukaryotic kinetochore dynamics 2 > 4 16 4
Centrosome maturation 12 > 13 2 16
Cholesterol synthesis 5 > 1 > 1
G2 M transition checkpoint 12 > > 4 16
Collagen fiber crosslinking > 3 6 > 24
Fibrillar collagen core structure organization > 1 26 24 >
Tenascin receptor signaling > 12 > 5 >
Glycolysis and Gluconeogenesis > > 25 > 2
ECM breakdown by matrix metalloproteases 28 5 > > >
Retinol metabolism > 2 > > >
Amyloid degradation, uptake and aggregation inhibition > 4 > > >
MSN01 MSN05 MSN06 MSN08 MSN09
Fibrillar collagen core structure organization > > 1 2 1
Decorin synthesis > > 4 6 2
Androgen receptor signaling 4 2 > 10 >
Potassium TM transport > 1 8 14 >
Cellular cholesterol uptake and efflux > > 2 > 3
Alternative complement pathway > 6 > 1 >
Procollagen cleavage into collagen 5 > > 12 >
Leptin receptor signaling 1 > > > >
GCSF receptor signaling 2 > > > >
Sodium TM transport > 3 > > >
Endogenous control of complement activity > > > 3 >
Chromatin organization by Polycomb repressive complexes 3 > > > >
Tropoelastin synthesis > > 4 > >
rRNA transcription > > > > 4
Lysine metabolism > 4 > > >
Classical complement pathway > > > 4 >
Actin polymerization > > > > 5
MSN01 MSN05 MSN06 MSN08 MSN09
Centrosome separation 1 > 2 1 3
Metaphase to anaphase checkpoint 3 > 5 3 5
Mitotic spindle assembly 2 > 3 7 6
Eukaryotic kinetochore dynamics 5 > 4 17 4
Centrosome maturation 12 > 12 2 16
G2 M transition checkpoint 12 > 28 5 16
Cholesterol synthesis 4 > 1 > 1
Collagen fiber crosslinking > 3 14 > 22
Fibrillar collagen core structure organization > 1 24 24 >
Bone morphogenetic protein receptor signaling > 16 > 4 >
Retinol metabolism > 2 > > >
Glycolysis and Gluconeogenesis > > > > 2
Amyloid degradation, uptake and aggregation inhibition > 4 > > >
Classical complement pathway > 5 > > >
MSN01 MSN05 MSN06 MSN08 MSN09
Semaphorin signaling 6 2 3 3 2
Alternative complement pathway > 1 4 1 1
Classical complement pathway > 3 5 4 3
Serine and glycine metabolism > 7 2 2 6
ECM breakdown & membrane shedding by adamalysins > 5 6 5 5
Water TM transport 2 4 > > 4
Fibrillar collagen core structure organization 3 > 1 > >
Osteonectin receptor signaling 1 > > > >
Albumin mediated blood protein transport 4 > > > >
Inhibition of apoptosis 5 > > > >
MSN01 MSN05 MSN06 MSN08 MSN09
Centrosome separation 1 1 1 1 1
Eukaryotic kinetochore dynamics 2 2 2 6 2
Mitotic spindle assembly 4 3 3 4 3
Metaphase to anaphase checkpoint 3 4 4 3 4
Mitotic H3 phosphorylation and dephosphorylation 6 6 6 2 6
Sister chromatid segregation 7 8 8 5 8
Cholesterol synthesis > 12 11 7 5

### MBCOL3
### imatinib
#### (non−c.toxic KI)

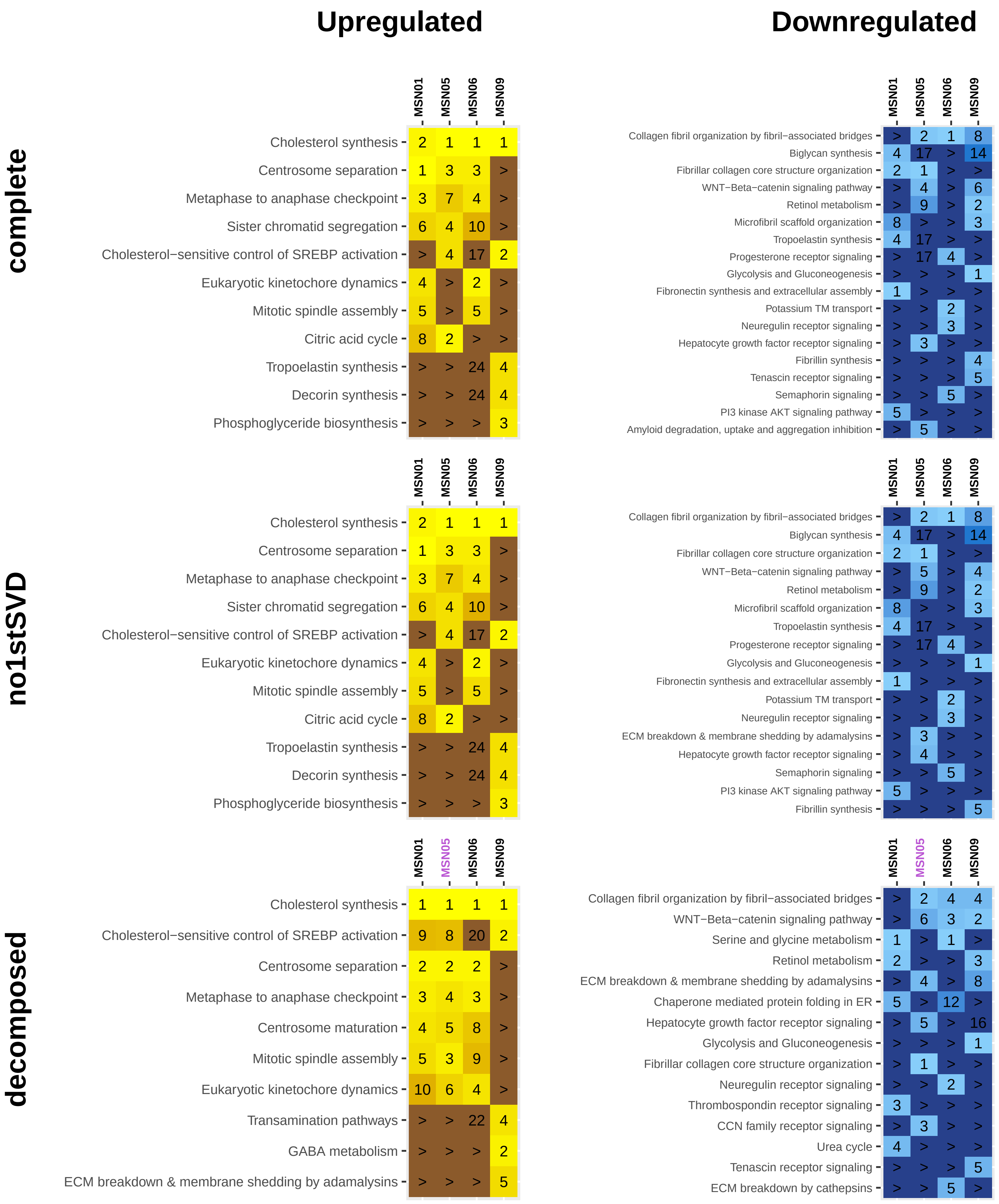

### MBCOL3
### lapatinib
#### (c.toxic KI)

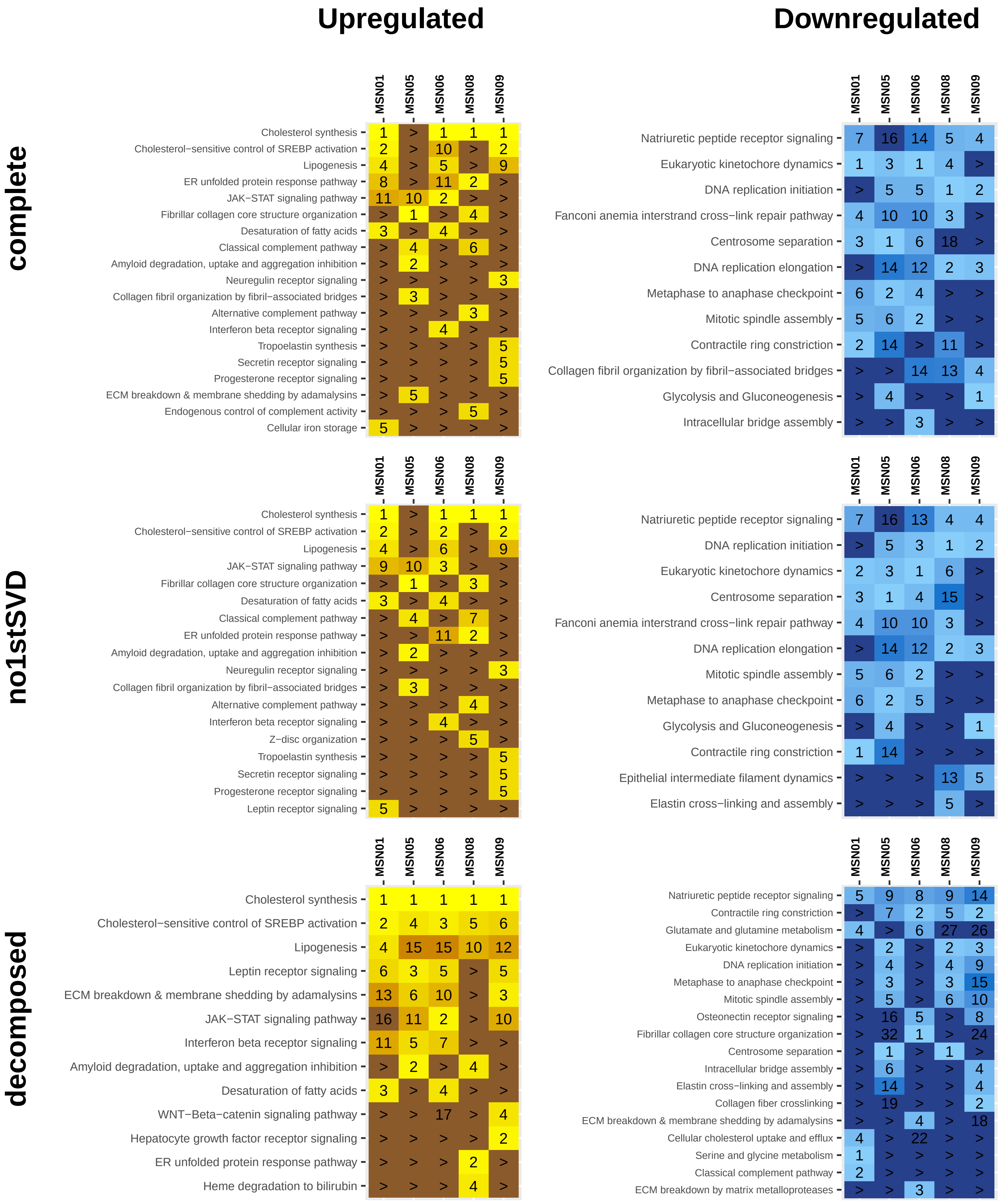

### MBCOL3
### nilotinib
#### (non−c.toxic KI)

##### complete

**Upregulated**

| | MSN01 | MSN05 | MSN06 | MSN08 | MSN09 |
|---|---|---|---|---|---|
| Centrosome separation | 1 | 8 | 2 | 1 | 2 |
| Serine and glycine metabolism | 4 | 1 | 5 | 4 | 8 |
| Eukaryotic kinetochore dynamics | > | 19 | 1 | 2 | 1 |
| Metaphase to anaphase checkpoint | > | 16 | 4 | 3 | 4 |
| Mitotic spindle assembly | > | 17 | 3 | 5 | 3 |
| Sister chromatid segregation | > | 3 | 9 | 16 | 12 |
| Centrosome maturation | > | > | 14 | 11 | 5 |
| Cholesterol synthesis | 18 | 5 | > | > | > |
| Glycolysis and Gluconeogenesis | 2 | > | > | > | > |
| Selectin−mediated Leukocyte rolling | > | 3 | > | > | > |
| GABA metabolism | > | 3 | > | > | > |
| Fibrillar collagen core structure organization | 3 | > | > | > | > |
| Fibronectin synthesis and extracellular assembly | 5 | > | > | > | > |

**Downregulated**

| | MSN01 | MSN05 | MSN06 | MSN08 | MSN09 |
|---|---|---|---|---|---|
| Z−disc organization | > | 5 | 1 | 1 | 2 |
| Thin myofilament organization | > | 3 | 5 | 2 | 3 |
| Thick myofilament organization | > | > | 2 | 5 | 5 |
| Epithelial intermediate filament dynamics | > | 6 | > | 4 | 4 |
| Fibrillar collagen core structure organization | > | 1 | > | 13 | 12 |
| GM−CSF receptor signaling | > | > | 11 | 3 | 17 |
| Collagen fibril organization by fibril−associated bridges | > | 2 | > | 10 | > |
| Potassium TM transport | > | > | 4 | > | 10 |
| Gap junction organization | 2 | > | > | 20 | > |
| Glycolysis and Gluconeogenesis | > | > | > | > | 1 |
| CM repolarization during AP & hyperpol. | 2 | > | > | > | > |
| Connection of muscle sarcomere to plasma membrane | 3 | > | > | > | > |
| ECM breakdown & membrane shedding by adamalysins | > | > | 4 | > | > |
| Hepatocyte growth factor receptor signaling | > | 4 | > | > | > |
| Amyloid plaque organization | 4 | > | > | > | > |
| Fibroblast growth factor receptor signaling | 5 | > | > | > | > |

##### no1stSVD

**Upregulated**

| | MSN01 | MSN05 | MSN06 | MSN08 | MSN09 |
|---|---|---|---|---|---|
| Centrosome separation | 1 | 8 | 2 | 1 | 2 |
| Serine and glycine metabolism | 9 | 1 | 5 | 4 | 8 |
| Eukaryotic kinetochore dynamics | > | 19 | 1 | 2 | 1 |
| Metaphase to anaphase checkpoint | > | 16 | 3 | 3 | 4 |
| Mitotic spindle assembly | > | 17 | 4 | 5 | 3 |
| Sister chromatid segregation | > | 3 | 11 | 20 | 12 |
| Centrosome maturation | > | > | 10 | 12 | 5 |
| Cholesterol synthesis | 18 | 5 | > | > | > |
| Glycolysis and Gluconeogenesis | 2 | > | > | > | > |
| Selectin−mediated Leukocyte rolling | > | 3 | > | > | > |
| GABA metabolism | > | 3 | > | > | > |
| Fibrillar collagen core structure organization | 3 | > | > | > | > |
| Fibronectin synthesis and extracellular assembly | 4 | > | > | > | > |
| Thin myofilament organization | 5 | > | > | > | > |

**Downregulated**

| | MSN01 | MSN05 | MSN06 | MSN08 | MSN09 |
|---|---|---|---|---|---|
| Z−disc organization | > | 5 | 1 | 1 | 2 |
| Thin myofilament organization | > | 3 | 4 | 2 | 3 |
| Epithelial intermediate filament dynamics | > | 6 | > | 3 | 4 |
| Fibrillar collagen core structure organization | > | 1 | > | 11 | 12 |
| Collagen fibril organization by fibril−associated bridges | > | 2 | > | 7 | > |
| Potassium TM transport | > | > | 5 | > | 10 |
| Gap junction organization | 2 | > | > | 18 | > |
| Glycolysis and Gluconeogenesis | > | > | > | > | 1 |
| CM repolarization during AP & hyperpol. | 2 | > | > | > | > |
| ECM breakdown & membrane shedding by adamalysins | > | > | 2 | > | > |
| Microtubule crosslinking and bundling | > | > | 3 | > | > |
| Connection of muscle sarcomere to plasma membrane | 3 | > | > | > | > |
| Tenascin receptor signaling | > | > | > | 4 | > |
| Hepatocyte growth factor receptor signaling | > | 4 | > | > | > |
| Amyloid plaque organization | 4 | > | > | > | > |
| Thick myofilament organization | > | > | > | > | 5 |
| Glycogen synthesis and glycogenolysis | > | > | > | 5 | > |
| Fibroblast growth factor receptor signaling | 5 | > | > | > | > |

##### decomposed

**Upregulated**

| | MSN01 | MSN05 | MSN06 | MSN08 | MSN09 |
|---|---|---|---|---|---|
| Centrosome separation | 1 | 1 | 1 | 1 | 1 |
| Metaphase to anaphase checkpoint | 3 | 3 | 3 | 3 | 3 |
| Eukaryotic kinetochore dynamics | 2 | 7 | 2 | 2 | 2 |
| Mitotic spindle assembly | 4 | 6 | 4 | 4 | 4 |
| Serine and glycine metabolism | 10 | 4 | 5 | 5 | 5 |
| Mitotic H3 phosphorylation and dephosphorylation | 5 | 8 | 6 | 6 | 6 |
| Contractile ring constriction | 13 | 2 | 6 | 18 | 6 |
| Cholesterol synthesis | > | 5 | > | > | > |

**Downregulated**

| | MSN01 | MSN05 | MSN06 | MSN08 | MSN09 |
|---|---|---|---|---|---|
| Fibrillar collagen core structure organization | 10 | 1 | 3 | 10 | 3 |
| Z−disc organization | > | 4 | 1 | 1 | 1 |
| Thin myofilament organization | > | > | 2 | 6 | 2 |
| Epithelial intermediate filament dynamics | > | 5 | 6 | > | 4 |
| ECM breakdown & membrane shedding by adamalysins | 8 | > | > | 2 | 6 |
| Cardiomyocyte pacemaker current generation | > | > | 6 | 8 | 4 |
| Thick myofilament organization | > | > | 4 | 3 | > |
| Retinol metabolism | > | 3 | > | 4 | > |
| Collagen fibril organization by fibril−associated bridges | > | 2 | 5 | > | > |
| Amyloid degradation, uptake and aggregation inhibition | 1 | > | > | > | > |
| Thrombospondin receptor signaling | 2 | > | > | > | > |
| Osteonectin receptor signaling | 2 | > | > | > | > |
| Muscarinic receptor signaling | 4 | > | > | > | > |
| CCN family receptor signaling | > | > | > | 5 | > |

### MBCOL3
### pazopanib
#### (c.toxic KI)

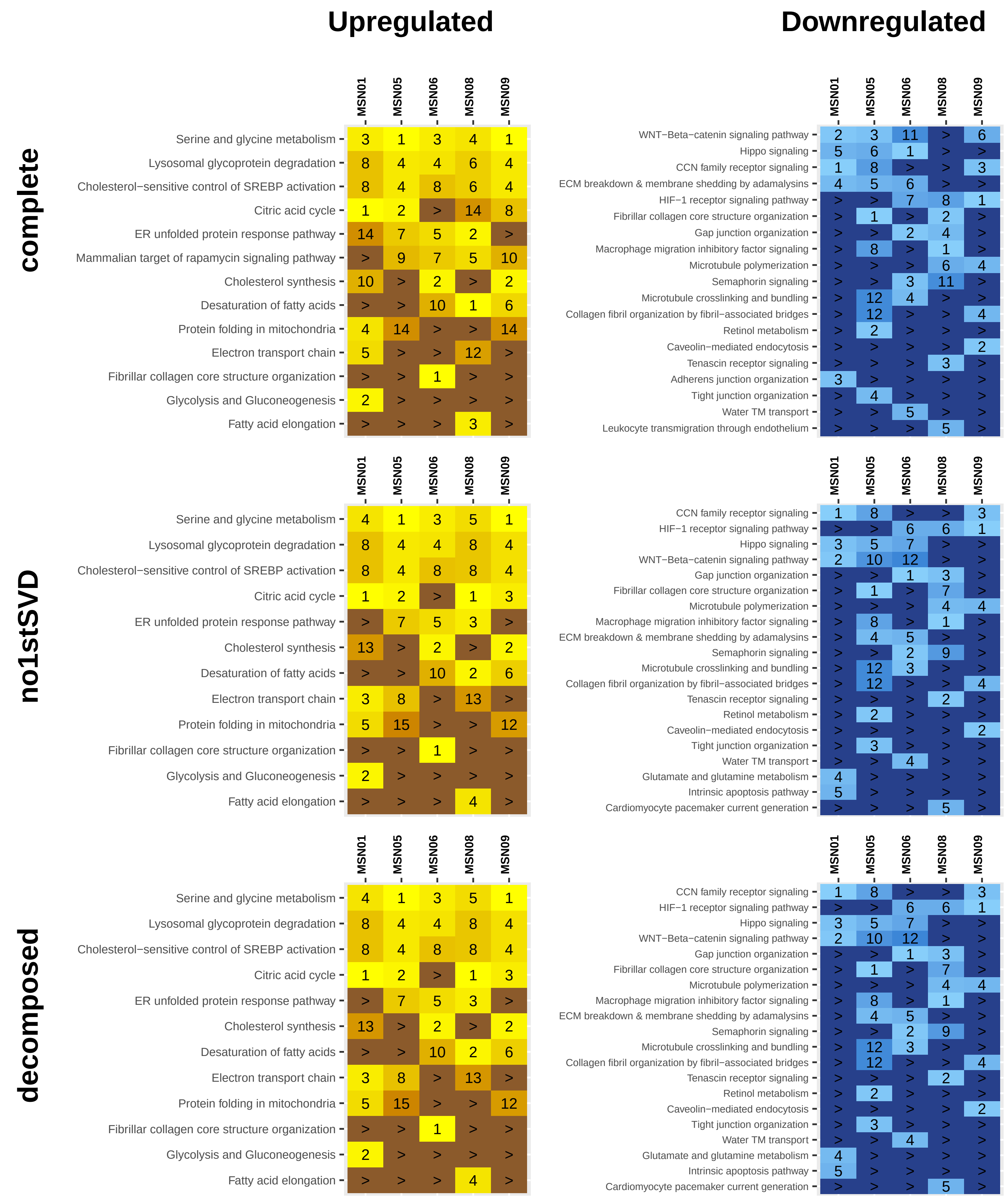

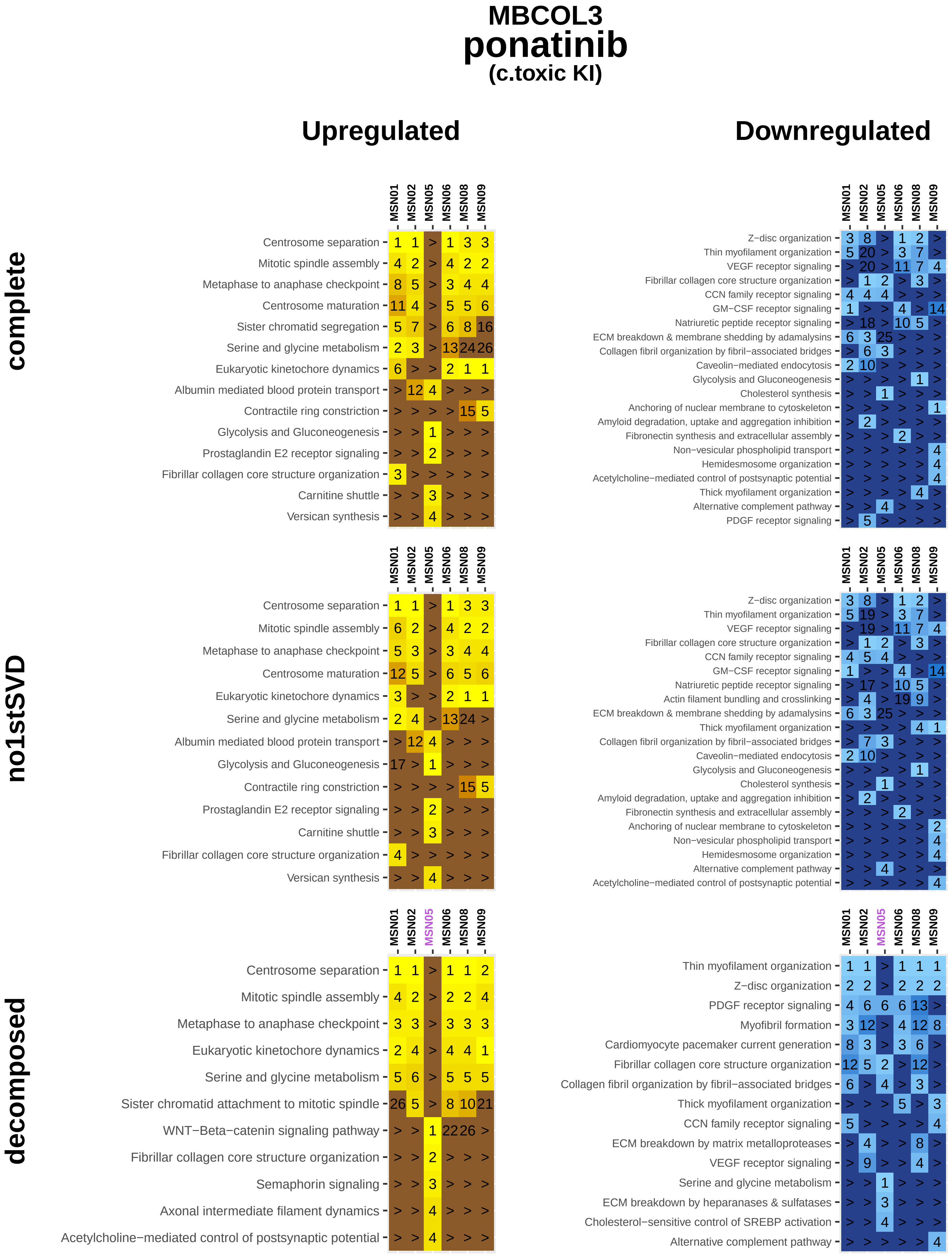
MBCOL3
ponatinib
(c.toxic KI)
Upregulated
Downregulated
complete
MSN01 MSN02 MSN05 MSN06 MSN08 MSN09
Centrosome separation 1 1 > 1 3 3
Mitotic spindle assembly 4 2 > 4 2 2
Metaphase to anaphase checkpoint 8 5 > 3 4 4
Centrosome maturation 11 4 > 5 5 6
Sister chromatid segregation 5 7 > 6 8 16
Serine and glycine metabolism 2 3 > 13 24 26
Eukaryotic kinetochore dynamics 6 > > 2 1 1
Albumin mediated blood protein transport > 12 4 > > >
Contractile ring constriction > > > > 15 5
Glycolysis and Gluconeogenesis > > 1 > > >
Prostaglandin E2 receptor signaling > > 2 > > >
Fibrillar collagen core structure organization 3 > > > > >
Carnitine shuttle > > 3 > > >
Versican synthesis > > 4 > > >
MSN01 MSN02 MSN05 MSN06 MSN08 MSN09
Z−disc organization 3 8 > 1 2 >
Thin myofilament organization 5 20 > 3 7 >
VEGF receptor signaling > 20 > 11 7 4
Fibrillar collagen core structure organization > 1 2 > 3 >
CCN family receptor signaling 4 4 4 > > >
GM−CSF receptor signaling 1 > > 4 > 14
Natriuretic peptide receptor signaling > 18 > 10 5 >
ECM breakdown & membrane shedding by adamalysins 6 3 25 > > >
Collagen fibril organization by fibril−associated bridges > 6 3 > > >
Caveolin−mediated endocytosis 2 10 > > > >
Glycolysis and Gluconeogenesis > > > > 1 >
Cholesterol synthesis > > 1 > > >
Anchoring of nuclear membrane to cytoskeleton > > > > > 1
Amyloid degradation, uptake and aggregation inhibition > 2 > > > >
Fibronectin synthesis and extracellular assembly > > > 2 > >
Non−vesicular phospholipid transport > > > > > 4
Hemidesmosome organization > > > > > 4
Acetylcholine−mediated control of postsynaptic potential > > > > > 4
Thick myofilament organization > > > > 4 >
Alternative complement pathway > > 4 > > >
PDGF receptor signaling > 5 > > > >
no1stSVD
MSN01 MSN02 MSN05 MSN06 MSN08 MSN09
Centrosome separation 1 1 > 1 3 3
Mitotic spindle assembly 6 2 > 4 2 2
Metaphase to anaphase checkpoint 5 3 > 3 4 4
Centrosome maturation 12 5 > 6 5 6
Eukaryotic kinetochore dynamics 3 > > 2 1 1
Serine and glycine metabolism 2 4 > 13 24 >
Albumin mediated blood protein transport > 12 4 > > >
Glycolysis and Gluconeogenesis 17 > 1 > > >
Contractile ring constriction > > > > 15 5
Prostaglandin E2 receptor signaling > > 2 > > >
Carnitine shuttle > > 3 > > >
Fibrillar collagen core structure organization 4 > > > > >
Versican synthesis > > 4 > > >
MSN01 MSN02 MSN05 MSN06 MSN08 MSN09
Z−disc organization 3 8 > 1 2 >
Thin myofilament organization 5 19 > 3 7 >
VEGF receptor signaling > 19 > 11 7 4
Fibrillar collagen core structure organization > 1 2 > 3 >
CCN family receptor signaling 4 5 4 > > >
GM−CSF receptor signaling 1 > > 4 > 14
Natriuretic peptide receptor signaling > 17 > 10 5 >
Actin filament bundling and crosslinking > 4 > 19 9 >
ECM breakdown & membrane shedding by adamalysins 6 3 25 > > >
Thick myofilament organization > > > > 4 1
Collagen fibril organization by fibril−associated bridges > 7 3 > > >
Caveolin−mediated endocytosis 2 10 > > > >
Glycolysis and Gluconeogenesis > > > > 1 >
Cholesterol synthesis > > 1 > > >
Amyloid degradation, uptake and aggregation inhibition > 2 > > > >
Fibronectin synthesis and extracellular assembly > > > 2 > >
Anchoring of nuclear membrane to cytoskeleton > > > > > 2
Non−vesicular phospholipid transport > > > > > 4
Hemidesmosome organization > > > > > 4
Alternative complement pathway > > 4 > > >
Acetylcholine−mediated control of postsynaptic potential > > > > > 4
decomposed
MSN01 MSN02 MSN05 MSN06 MSN08 MSN09
Centrosome separation 1 1 > 1 1 2
Mitotic spindle assembly 4 2 > 2 2 4
Metaphase to anaphase checkpoint 3 3 > 3 3 3
Eukaryotic kinetochore dynamics 2 4 > 4 4 1
Serine and glycine metabolism 5 6 > 5 5 5
Sister chromatid attachment to mitotic spindle 26 5 > 8 10 21
WNT−Beta−catenin signaling pathway > > 1 22 26 >
Fibrillar collagen core structure organization > > 2 > > >
Semaphorin signaling > > 3 > > >
Axonal intermediate filament dynamics > > 4 > > >
Acetylcholine−mediated control of postsynaptic potential > > 4 > > >
MSN01 MSN02 MSN05 MSN06 MSN08 MSN09
Thin myofilament organization 1 1 > 1 1 1
Z−disc organization 2 2 > 2 2 2
PDGF receptor signaling 4 6 6 6 13 >
Myofibril formation 3 12 > 4 12 8
Cardiomyocyte pacemaker current generation 8 3 > 3 6 >
Fibrillar collagen core structure organization 12 5 2 > 12 >
Collagen fibril organization by fibril−associated bridges 6 > 4 > 3 >
Thick myofilament organization > > > 5 > 3
CCN family receptor signaling 5 > > > > 4
ECM breakdown by matrix metalloproteases > 4 > > 8 >
VEGF receptor signaling > 9 > > 4 >
Serine and glycine metabolism > > 1 > > >
ECM breakdown by heparanases & sulfatases > > 3 > > >
Cholesterol−sensitive control of SREBP activation > > 4 > > >
Alternative complement pathway > > > > > 4

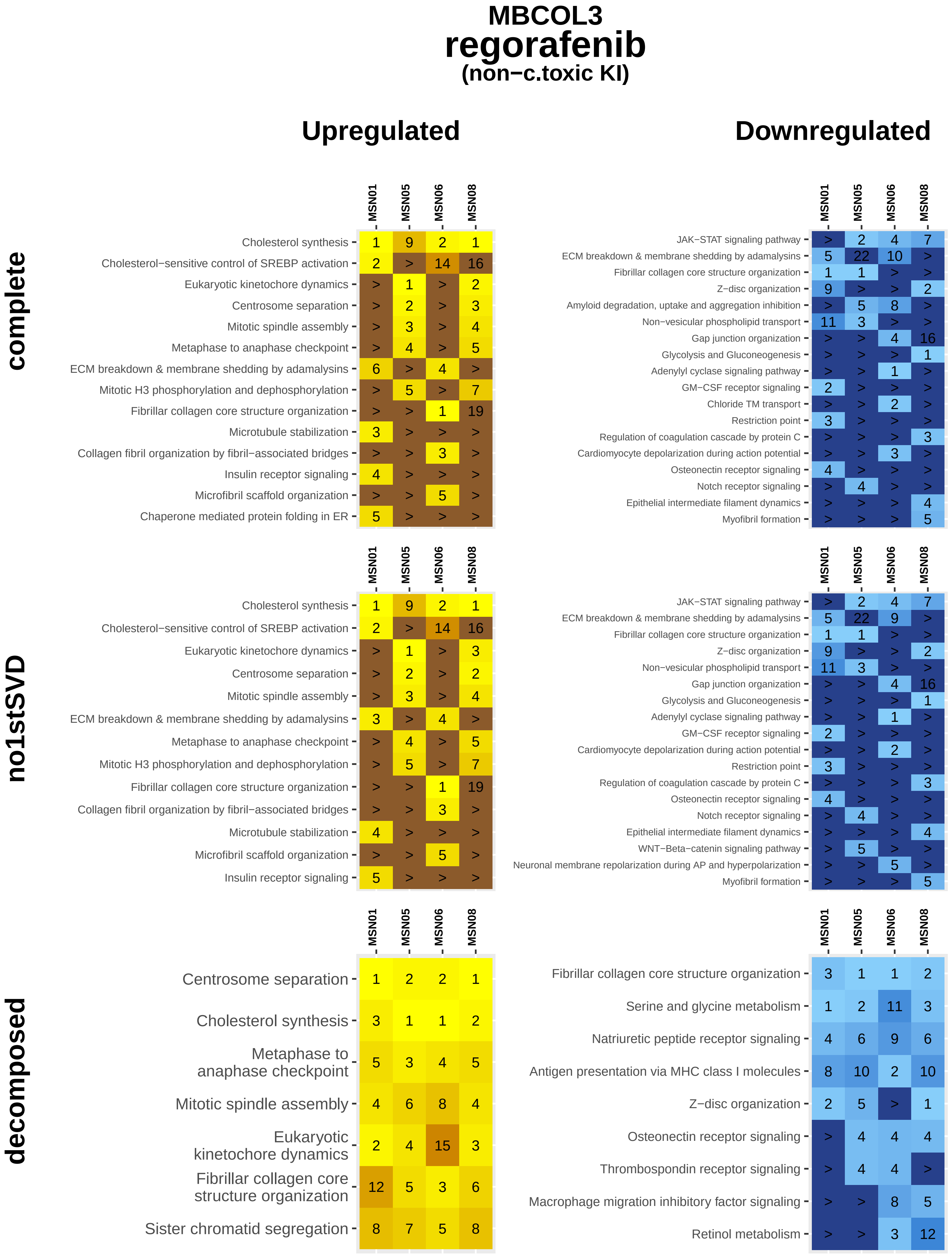
MBCOL3
regorafenib
(non−c.toxic KI)
Upregulated
Downregulated
complete
no1stSVD
decomposed
MSN01 MSN05 MSN06 MSN08
Cholesterol synthesis 1 9 2 1
Cholesterol−sensitive control of SREBP activation 2 > 14 16
Eukaryotic kinetochore dynamics > 1 > 2
Centrosome separation > 2 > 3
Mitotic spindle assembly > 3 > 4
Metaphase to anaphase checkpoint > 4 > 5
ECM breakdown & membrane shedding by adamalysins 6 > 4 >
Mitotic H3 phosphorylation and dephosphorylation > 5 > 7
Fibrillar collagen core structure organization > > 1 19
Microtubule stabilization 3 > > >
Collagen fibril organization by fibril−associated bridges > > 3 >
Insulin receptor signaling 4 > > >
Microfibril scaffold organization > > 5 >
Chaperone mediated protein folding in ER 5 > > >
JAK−STAT signaling pathway > 2 4 7
ECM breakdown & membrane shedding by adamalysins 5 22 10 >
Fibrillar collagen core structure organization 1 1 > >
Z−disc organization 9 > > 2
Amyloid degradation, uptake and aggregation inhibition > 5 8 >
Non−vesicular phospholipid transport 11 3 > >
Gap junction organization > > 4 16
Glycolysis and Gluconeogenesis > > > 1
Adenylyl cyclase signaling pathway > > 1 >
GM−CSF receptor signaling 2 > > >
Chloride TM transport > > 2 >
Restriction point 3 > > >
Regulation of coagulation cascade by protein C > > > 3
Cardiomyocyte depolarization during action potential > > 3 >
Osteonectin receptor signaling 4 > > >
Notch receptor signaling > 4 > >
Epithelial intermediate filament dynamics > > > 4
Myofibril formation > > > 5
Cholesterol synthesis 1 9 2 1
Cholesterol−sensitive control of SREBP activation 2 > 14 16
Eukaryotic kinetochore dynamics > 1 > 3
Centrosome separation > 2 > 2
Mitotic spindle assembly > 3 > 4
ECM breakdown & membrane shedding by adamalysins 3 > 4 >
Metaphase to anaphase checkpoint > 4 > 5
Mitotic H3 phosphorylation and dephosphorylation > 5 > 7
Fibrillar collagen core structure organization > > 1 19
Collagen fibril organization by fibril−associated bridges > > 3 >
Microtubule stabilization 4 > > >
Microfibril scaffold organization > > 5 >
Insulin receptor signaling 5 > > >
JAK−STAT signaling pathway > 2 4 7
ECM breakdown & membrane shedding by adamalysins 5 22 9 >
Fibrillar collagen core structure organization 1 1 > >
Z−disc organization 9 > > 2
Non−vesicular phospholipid transport 11 3 > >
Gap junction organization > > 4 16
Glycolysis and Gluconeogenesis > > > 1
Adenylyl cyclase signaling pathway > > 1 >
GM−CSF receptor signaling 2 > > >
Cardiomyocyte depolarization during action potential > > 2 >
Restriction point 3 > > >
Regulation of coagulation cascade by protein C > > > 3
Osteonectin receptor signaling 4 > > >
Notch receptor signaling > 4 > >
Epithelial intermediate filament dynamics > > > 4
WNT−Beta−catenin signaling pathway > 5 > >
Neuronal membrane repolarization during AP and hyperpolarization > > 5 >
Myofibril formation > > > 5
Centrosome separation 1 2 2 1
Cholesterol synthesis 3 1 1 2
Metaphase to anaphase checkpoint 5 3 4 5
Mitotic spindle assembly 4 6 8 4
Eukaryotic kinetochore dynamics 2 4 15 3
Fibrillar collagen core structure organization 12 5 3 6
Sister chromatid segregation 8 7 5 8
Fibrillar collagen core structure organization 3 1 1 2
Serine and glycine metabolism 1 2 11 3
Natriuretic peptide receptor signaling 4 6 9 6
Antigen presentation via MHC class I molecules 8 10 2 10
Z−disc organization 2 5 > 1
Osteonectin receptor signaling > 4 4 4
Thrombospondin receptor signaling > 4 4 >
Macrophage migration inhibitory factor signaling > > 8 5
Retinol metabolism > > 3 12

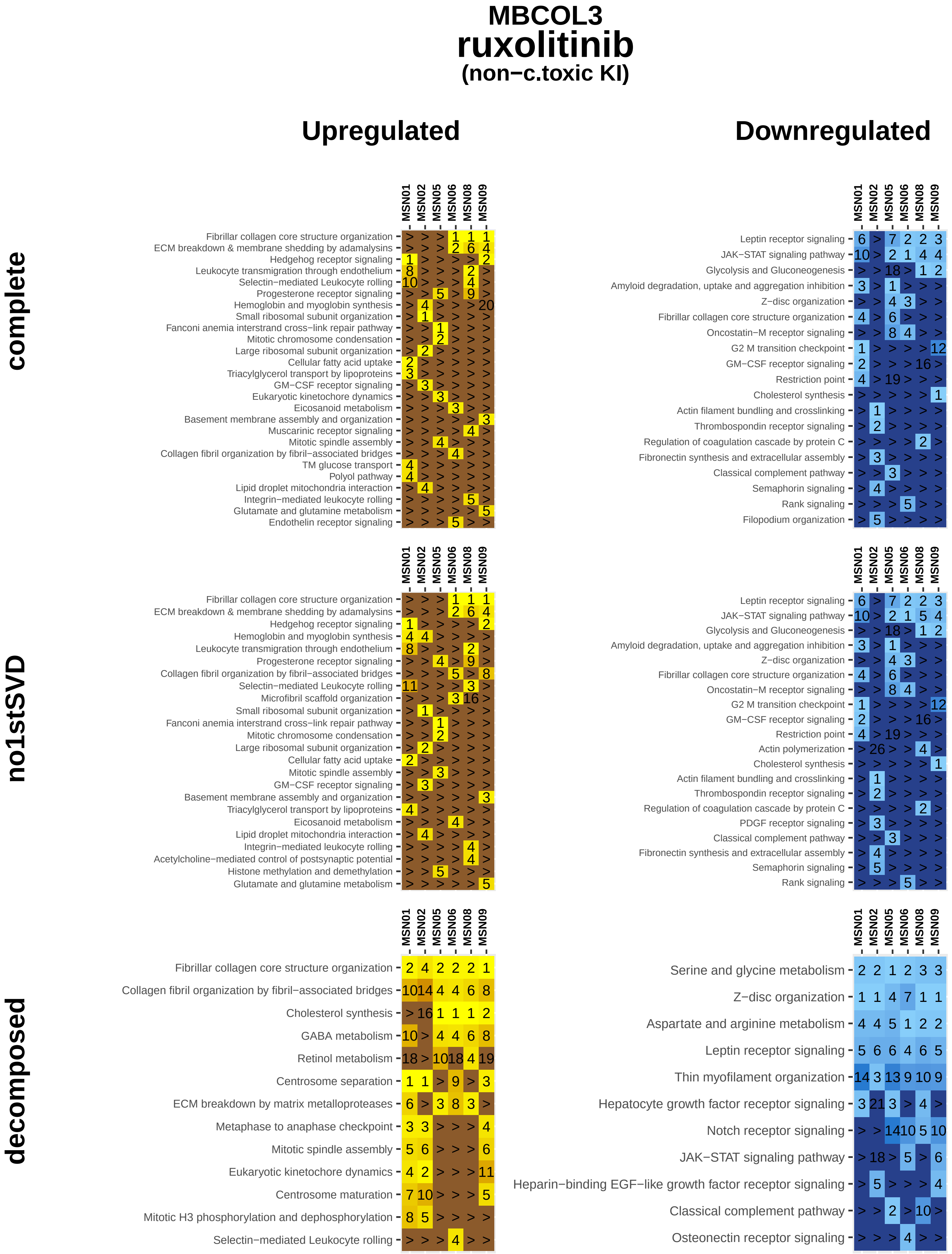

MBCOL3
ruxolitinib
(non−c.toxic KI)
Upregulated
Downregulated
complete
no1stSVD
decomposed
MSN01
MSN02
MSN05
MSN06
MSN08
MSN09
Fibrillar collagen core structure organization
ECM breakdown & membrane shedding by adamalysins
Hedgehog receptor signaling
Leukocyte transmigration through endothelium
Selectin−mediated Leukocyte rolling
Progesterone receptor signaling
Hemoglobin and myoglobin synthesis
Small ribosomal subunit organization
Fanconi anemia interstrand cross−link repair pathway
Mitotic chromosome condensation
Large ribosomal subunit organization
Cellular fatty acid uptake
Triacylglycerol transport by lipoproteins
GM−CSF receptor signaling
Eukaryotic kinetochore dynamics
Eicosanoid metabolism
Basement membrane assembly and organization
Muscarinic receptor signaling
Mitotic spindle assembly
Collagen fibril organization by fibril−associated bridges
TM glucose transport
Polyol pathway
Lipid droplet mitochondria interaction
Integrin−mediated leukocyte rolling
Glutamate and glutamine metabolism
Endothelin receptor signaling
Leptin receptor signaling
JAK−STAT signaling pathway
Glycolysis and Gluconeogenesis
Amyloid degradation, uptake and aggregation inhibition
Z−disc organization
Fibrillar collagen core structure organization
Oncostatin−M receptor signaling
G2 M transition checkpoint
GM−CSF receptor signaling
Restriction point
Cholesterol synthesis
Actin filament bundling and crosslinking
Thrombospondin receptor signaling
Regulation of coagulation cascade by protein C
Fibronectin synthesis and extracellular assembly
Classical complement pathway
Semaphorin signaling
Rank signaling
Filopodium organization
Fibrillar collagen core structure organization
ECM breakdown & membrane shedding by adamalysins
Hedgehog receptor signaling
Hemoglobin and myoglobin synthesis
Leukocyte transmigration through endothelium
Progesterone receptor signaling
Collagen fibril organization by fibril−associated bridges
Selectin−mediated Leukocyte rolling
Microfibril scaffold organization
Small ribosomal subunit organization
Fanconi anemia interstrand cross−link repair pathway
Mitotic chromosome condensation
Large ribosomal subunit organization
Cellular fatty acid uptake
Mitotic spindle assembly
GM−CSF receptor signaling
Basement membrane assembly and organization
Triacylglycerol transport by lipoproteins
Eicosanoid metabolism
Lipid droplet mitochondria interaction
Integrin−mediated leukocyte rolling
Acetylcholine−mediated control of postsynaptic potential
Histone methylation and demethylation
Glutamate and glutamine metabolism
Leptin receptor signaling
JAK−STAT signaling pathway
Glycolysis and Gluconeogenesis
Amyloid degradation, uptake and aggregation inhibition
Z−disc organization
Fibrillar collagen core structure organization
Oncostatin−M receptor signaling
G2 M transition checkpoint
GM−CSF receptor signaling
Restriction point
Actin polymerization
Cholesterol synthesis
Actin filament bundling and crosslinking
Thrombospondin receptor signaling
Regulation of coagulation cascade by protein C
PDGF receptor signaling
Classical complement pathway
Fibronectin synthesis and extracellular assembly
Semaphorin signaling
Rank signaling
Fibrillar collagen core structure organization
Collagen fibril organization by fibril−associated bridges
Cholesterol synthesis
GABA metabolism
Retinol metabolism
Centrosome separation
ECM breakdown by matrix metalloproteases
Metaphase to anaphase checkpoint
Mitotic spindle assembly
Eukaryotic kinetochore dynamics
Centrosome maturation
Mitotic H3 phosphorylation and dephosphorylation
Selectin−mediated Leukocyte rolling
Serine and glycine metabolism
Z−disc organization
Aspartate and arginine metabolism
Leptin receptor signaling
Thin myofilament organization
Hepatocyte growth factor receptor signaling
Notch receptor signaling
JAK−STAT signaling pathway
Heparin−binding EGF−like growth factor receptor signaling
Classical complement pathway
Osteonectin receptor signaling

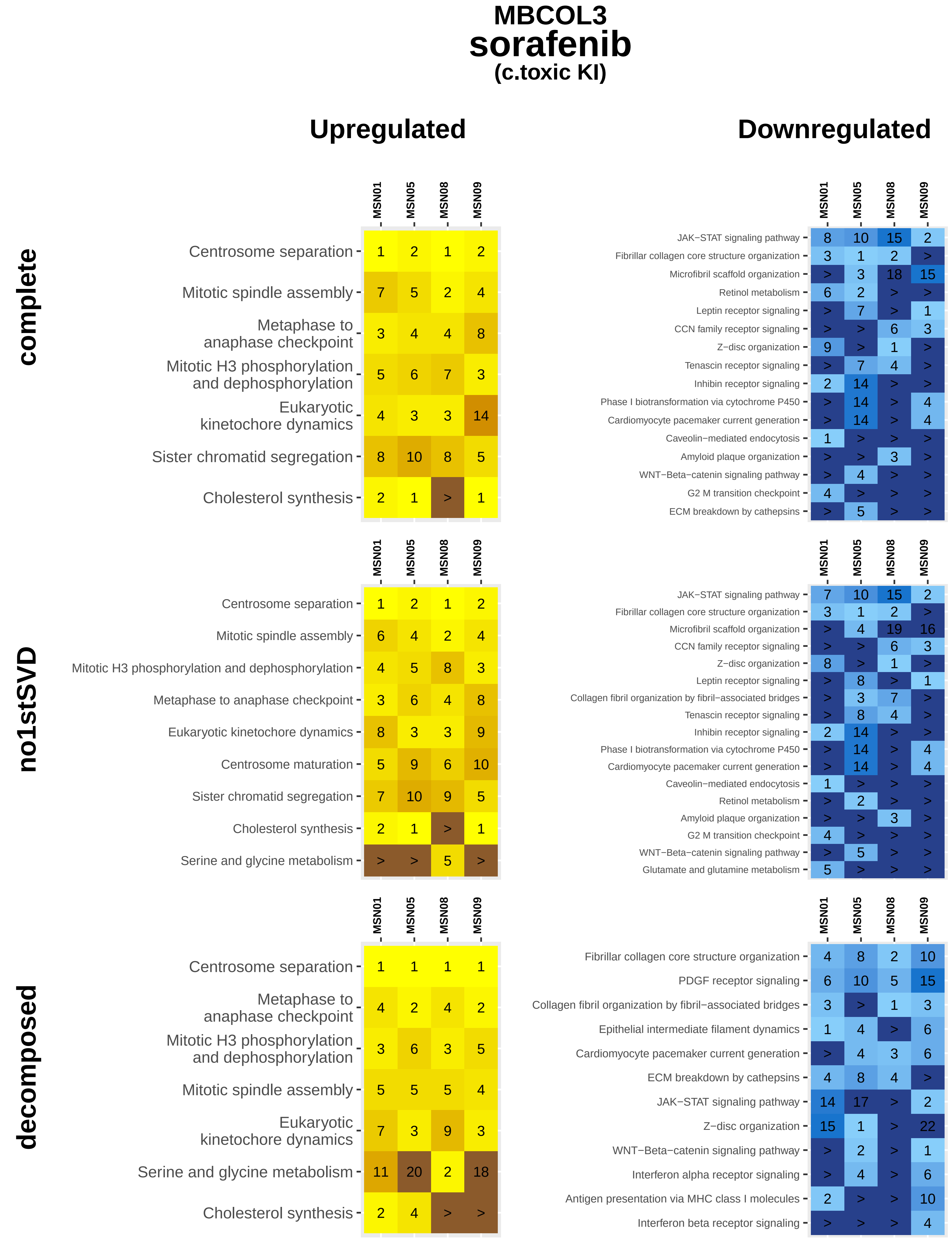

MBCOL3
sorafenib
(c.toxic KI)
Upregulated
Downregulated
complete
no1stSVD
decomposed
MSN01
MSN05
MSN08
MSN09
Centrosome separation
Mitotic spindle assembly
Metaphase to anaphase checkpoint
Mitotic H3 phosphorylation and dephosphorylation
Eukaryotic kinetochore dynamics
Sister chromatid segregation
Cholesterol synthesis
Centrosome maturation
Serine and glycine metabolism
JAK−STAT signaling pathway
Fibrillar collagen core structure organization
Microfibril scaffold organization
Retinol metabolism
Leptin receptor signaling
CCN family receptor signaling
Z−disc organization
Tenascin receptor signaling
Inhibin receptor signaling
Phase I biotransformation via cytochrome P450
Cardiomyocyte pacemaker current generation
Caveolin−mediated endocytosis
Amyloid plaque organization
WNT−Beta−catenin signaling pathway
G2 M transition checkpoint
ECM breakdown by cathepsins
Collagen fibril organization by fibril−associated bridges
Glutamate and glutamine metabolism
PDGF receptor signaling
Epithelial intermediate filament dynamics
Interferon alpha receptor signaling
Antigen presentation via MHC class I molecules
Interferon beta receptor signaling

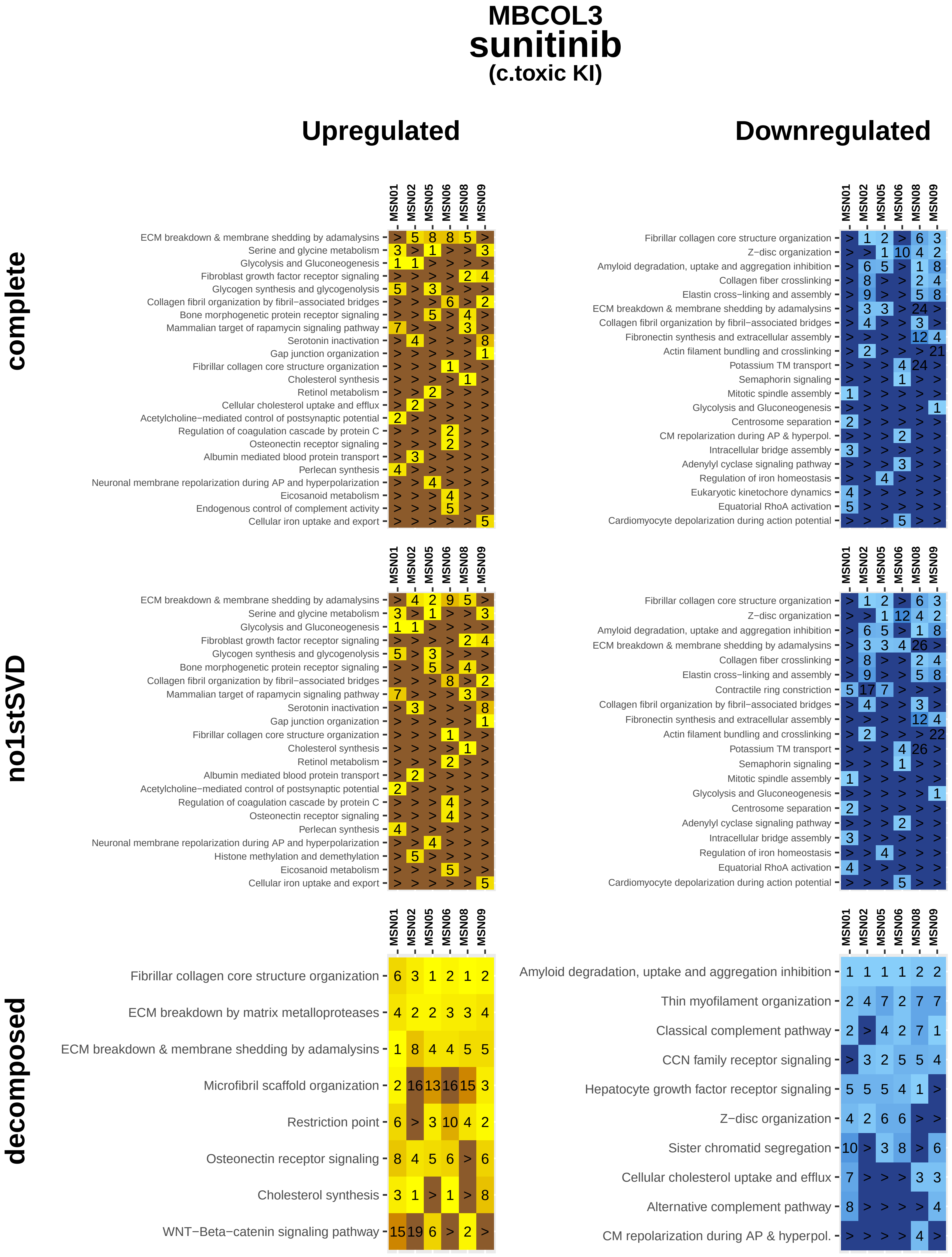

MBCOL3
sunitinib
(c.toxic KI)
Upregulated
Downregulated
complete
no1stSVD
decomposed
MSN01
MSN02
MSN05
MSN06
MSN08
MSN09
ECM breakdown & membrane shedding by adamalysins
Serine and glycine metabolism
Glycolysis and Gluconeogenesis
Fibroblast growth factor receptor signaling
Glycogen synthesis and glycogenolysis
Collagen fibril organization by fibril−associated bridges
Bone morphogenetic protein receptor signaling
Mammalian target of rapamycin signaling pathway
Serotonin inactivation
Gap junction organization
Fibrillar collagen core structure organization
Cholesterol synthesis
Retinol metabolism
Cellular cholesterol uptake and efflux
Acetylcholine−mediated control of postsynaptic potential
Regulation of coagulation cascade by protein C
Osteonectin receptor signaling
Albumin mediated blood protein transport
Perlecan synthesis
Neuronal membrane repolarization during AP and hyperpolarization
Eicosanoid metabolism
Endogenous control of complement activity
Cellular iron uptake and export
Fibrillar collagen core structure organization
Z−disc organization
Amyloid degradation, uptake and aggregation inhibition
Collagen fiber crosslinking
Elastin cross−linking and assembly
ECM breakdown & membrane shedding by adamalysins
Collagen fibril organization by fibril−associated bridges
Fibronectin synthesis and extracellular assembly
Actin filament bundling and crosslinking
Potassium TM transport
Semaphorin signaling
Mitotic spindle assembly
Glycolysis and Gluconeogenesis
Centrosome separation
CM repolarization during AP & hyperpol.
Intracellular bridge assembly
Adenylyl cyclase signaling pathway
Regulation of iron homeostasis
Eukaryotic kinetochore dynamics
Equatorial RhoA activation
Cardiomyocyte depolarization during action potential
ECM breakdown & membrane shedding by adamalysins
Serine and glycine metabolism
Glycolysis and Gluconeogenesis
Fibroblast growth factor receptor signaling
Glycogen synthesis and glycogenolysis
Bone morphogenetic protein receptor signaling
Collagen fibril organization by fibril−associated bridges
Mammalian target of rapamycin signaling pathway
Serotonin inactivation
Gap junction organization
Fibrillar collagen core structure organization
Cholesterol synthesis
Retinol metabolism
Albumin mediated blood protein transport
Acetylcholine−mediated control of postsynaptic potential
Regulation of coagulation cascade by protein C
Osteonectin receptor signaling
Perlecan synthesis
Neuronal membrane repolarization during AP and hyperpolarization
Histone methylation and demethylation
Eicosanoid metabolism
Cellular iron uptake and export
Fibrillar collagen core structure organization
Z−disc organization
Amyloid degradation, uptake and aggregation inhibition
ECM breakdown & membrane shedding by adamalysins
Collagen fiber crosslinking
Elastin cross−linking and assembly
Contractile ring constriction
Collagen fibril organization by fibril−associated bridges
Fibronectin synthesis and extracellular assembly
Actin filament bundling and crosslinking
Potassium TM transport
Semaphorin signaling
Mitotic spindle assembly
Glycolysis and Gluconeogenesis
Centrosome separation
Adenylyl cyclase signaling pathway
Intracellular bridge assembly
Regulation of iron homeostasis
Equatorial RhoA activation
Cardiomyocyte depolarization during action potential
Fibrillar collagen core structure organization
ECM breakdown by matrix metalloproteases
ECM breakdown & membrane shedding by adamalysins
Microfibril scaffold organization
Restriction point
Osteonectin receptor signaling
Cholesterol synthesis
WNT−Beta−catenin signaling pathway
Amyloid degradation, uptake and aggregation inhibition
Thin myofilament organization
Classical complement pathway
CCN family receptor signaling
Hepatocyte growth factor receptor signaling
Z−disc organization
Sister chromatid segregation
Cellular cholesterol uptake and efflux
Alternative complement pathway
CM repolarization during AP & hyperpol.

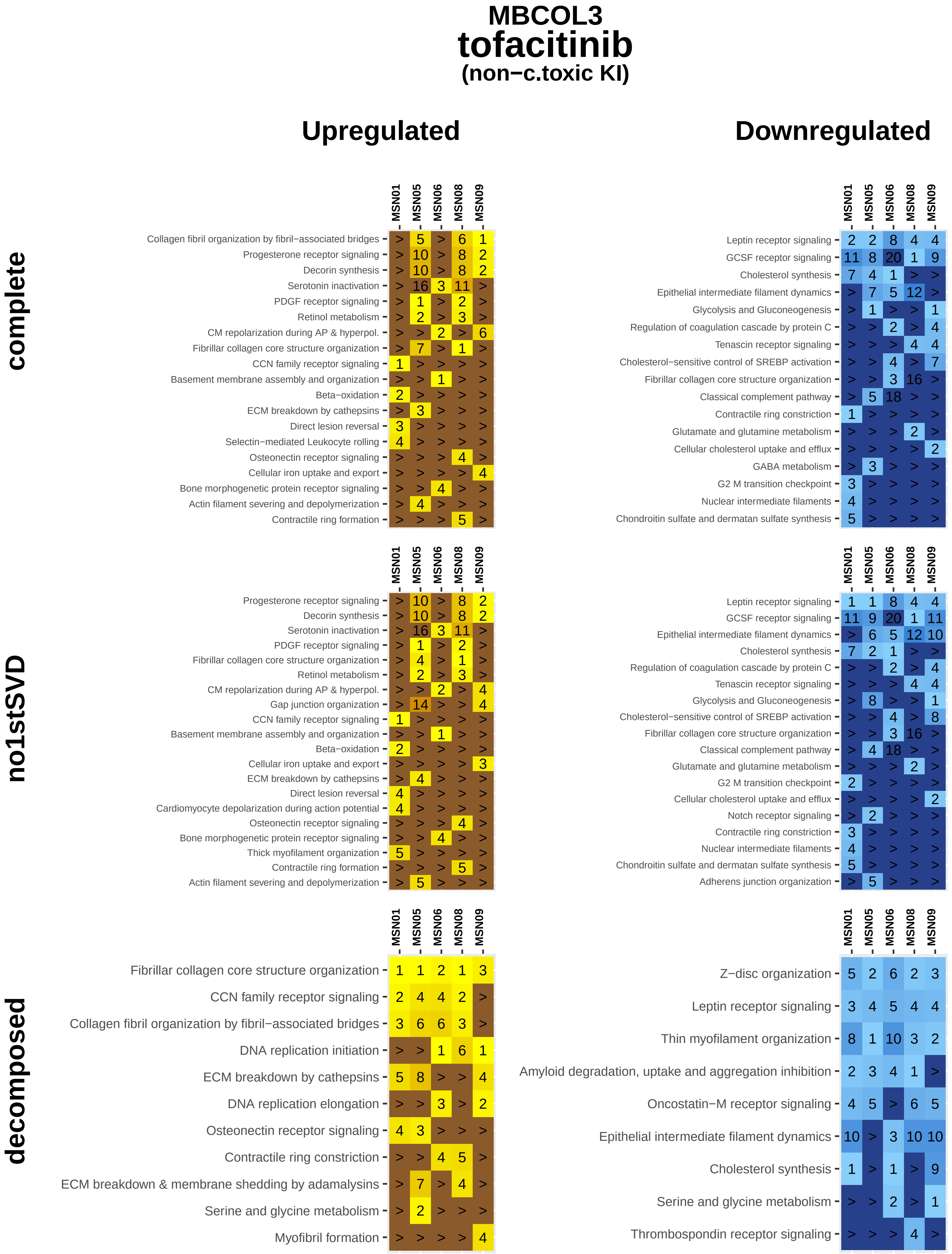

MBCOL3
tofacitinib
(non−c.toxic KI)
Upregulated
Downregulated
complete
no1stSVD
decomposed
MSN01 MSN05 MSN06 MSN08 MSN09
Collagen fibril organization by fibril−associated bridges > 5 > 6 1
Progesterone receptor signaling > 10 > 8 2
Decorin synthesis > 10 > 8 2
Serotonin inactivation > 16 3 11 >
PDGF receptor signaling > 1 > 2 >
Retinol metabolism > 2 > 3 >
CM repolarization during AP & hyperpol. > > 2 > 6
Fibrillar collagen core structure organization > 7 > 1 >
CCN family receptor signaling 1 > > > >
Basement membrane assembly and organization > > 1 > >
Beta−oxidation 2 > > > >
ECM breakdown by cathepsins > 3 > > >
Direct lesion reversal 3 > > > >
Selectin−mediated Leukocyte rolling 4 > > > >
Osteonectin receptor signaling > > > 4 >
Cellular iron uptake and export > > > > 4
Bone morphogenetic protein receptor signaling > > 4 > >
Actin filament severing and depolymerization > 4 > > >
Contractile ring formation > > > 5 >
Leptin receptor signaling 2 2 8 4 4
GCSF receptor signaling 11 8 20 1 9
Cholesterol synthesis 7 4 1 > >
Epithelial intermediate filament dynamics > 7 5 12 >
Glycolysis and Gluconeogenesis > 1 > > 1
Regulation of coagulation cascade by protein C > > 2 > 4
Tenascin receptor signaling > > > 4 4
Cholesterol−sensitive control of SREBP activation > > 4 > 7
Fibrillar collagen core structure organization > > 3 16 >
Classical complement pathway > 5 18 > >
Contractile ring constriction 1 > > > >
Glutamate and glutamine metabolism > > > 2 >
Cellular cholesterol uptake and efflux > > > > 2
GABA metabolism > 3 > > >
G2 M transition checkpoint 3 > > > >
Nuclear intermediate filaments 4 > > > >
Chondroitin sulfate and dermatan sulfate synthesis 5 > > > >
Progesterone receptor signaling > 10 > 8 2
Decorin synthesis > 10 > 8 2
Serotonin inactivation > 16 3 11 >
PDGF receptor signaling > 1 > 2 >
Fibrillar collagen core structure organization > 4 > 1 >
Retinol metabolism > 2 > 3 >
CM repolarization during AP & hyperpol. > > 2 > 4
Gap junction organization > 14 > > 4
CCN family receptor signaling 1 > > > >
Basement membrane assembly and organization > > 1 > >
Beta−oxidation 2 > > > >
Cellular iron uptake and export > > > > 3
ECM breakdown by cathepsins > 4 > > >
Direct lesion reversal 4 > > > >
Cardiomyocyte depolarization during action potential 4 > > > >
Osteonectin receptor signaling > > > 4 >
Bone morphogenetic protein receptor signaling > > 4 > >
Thick myofilament organization 5 > > > >
Contractile ring formation > > > 5 >
Actin filament severing and depolymerization > 5 > > >
Leptin receptor signaling 1 1 8 4 4
GCSF receptor signaling 11 9 20 1 11
Epithelial intermediate filament dynamics > 6 5 12 10
Cholesterol synthesis 7 2 1 > >
Regulation of coagulation cascade by protein C > > 2 > 4
Tenascin receptor signaling > > > 4 4
Glycolysis and Gluconeogenesis > 8 > > 1
Cholesterol−sensitive control of SREBP activation > > 4 > 8
Fibrillar collagen core structure organization > > 3 16 >
Classical complement pathway > 4 18 > >
Glutamate and glutamine metabolism > > > 2 >
G2 M transition checkpoint 2 > > > >
Cellular cholesterol uptake and efflux > > > > 2
Notch receptor signaling > 2 > > >
Contractile ring constriction 3 > > > >
Nuclear intermediate filaments 4 > > > >
Chondroitin sulfate and dermatan sulfate synthesis 5 > > > >
Adherens junction organization > 5 > > >
Fibrillar collagen core structure organization 1 1 2 1 3
CCN family receptor signaling 2 4 4 2 >
Collagen fibril organization by fibril−associated bridges 3 6 6 3 >
DNA replication initiation > > 1 6 1
ECM breakdown by cathepsins 5 8 > > 4
DNA replication elongation > > 3 > 2
Osteonectin receptor signaling 4 3 > > >
Contractile ring constriction > > 4 5 >
ECM breakdown & membrane shedding by adamalysins > 7 > 4 >
Serine and glycine metabolism > 2 > > >
Myofibril formation > > > > 4
Z−disc organization 5 2 6 2 3
Leptin receptor signaling 3 4 5 4 4
Thin myofilament organization 8 1 10 3 2
Amyloid degradation, uptake and aggregation inhibition 2 3 4 1 >
Oncostatin−M receptor signaling 4 5 > 6 5
Epithelial intermediate filament dynamics 10 > 3 10 10
Cholesterol synthesis 1 > 1 > 9
Serine and glycine metabolism > > 2 > 1
Thrombospondin receptor signaling > > > 4 >

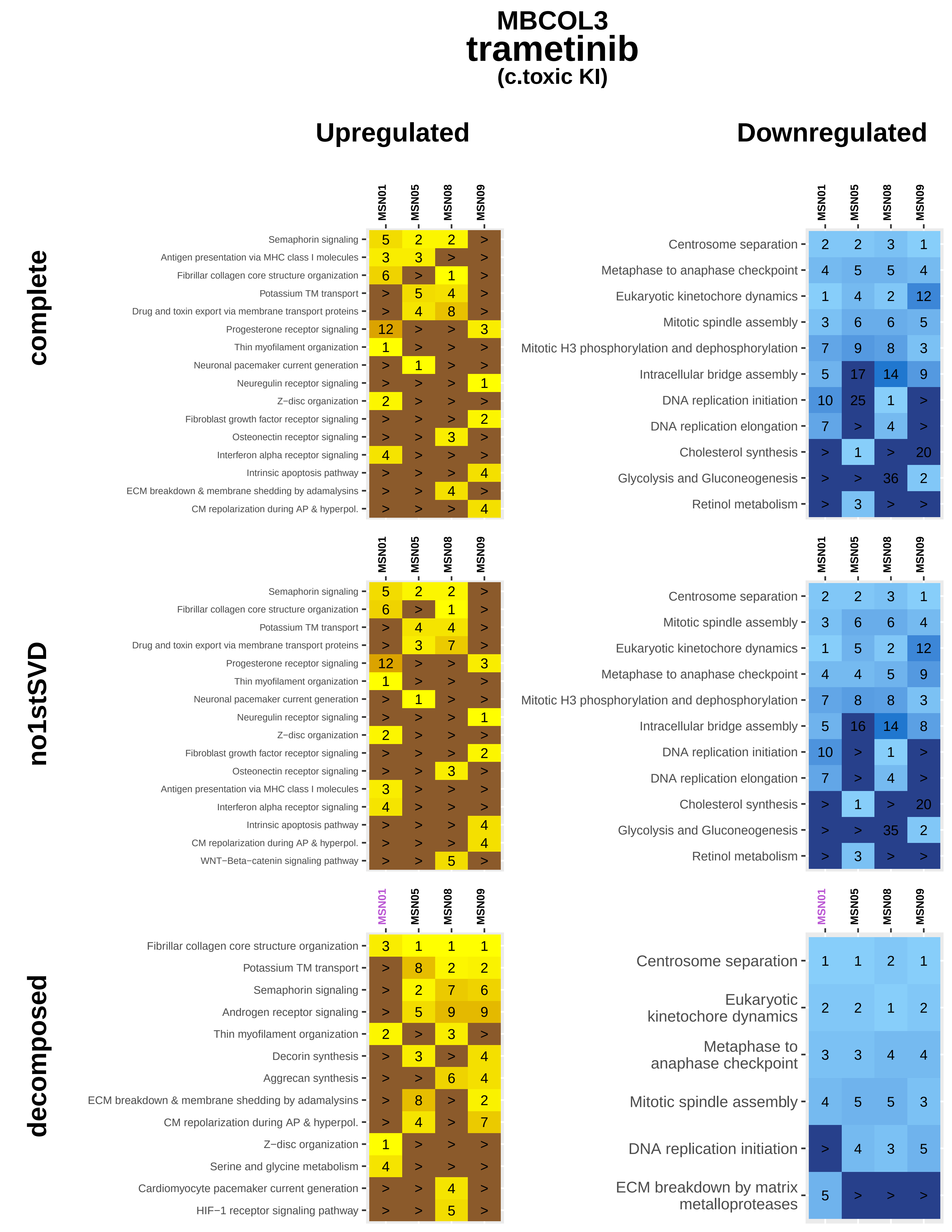
MBCOL3
trametinib
(c.toxic KI)
Upregulated
Downregulated
complete
no1stSVD
decomposed
MSN01
MSN05
MSN08
MSN09
Semaphorin signaling
Antigen presentation via MHC class I molecules
Fibrillar collagen core structure organization
Potassium TM transport
Drug and toxin export via membrane transport proteins
Progesterone receptor signaling
Thin myofilament organization
Neuronal pacemaker current generation
Neuregulin receptor signaling
Z−disc organization
Fibroblast growth factor receptor signaling
Osteonectin receptor signaling
Interferon alpha receptor signaling
Intrinsic apoptosis pathway
ECM breakdown & membrane shedding by adamalysins
CM repolarization during AP & hyperpol.
WNT−Beta−catenin signaling pathway
Androgen receptor signaling
Decorin synthesis
Aggrecan synthesis
Serine and glycine metabolism
Cardiomyocyte pacemaker current generation
HIF−1 receptor signaling pathway
Centrosome separation
Metaphase to anaphase checkpoint
Eukaryotic kinetochore dynamics
Mitotic spindle assembly
Mitotic H3 phosphorylation and dephosphorylation
Intracellular bridge assembly
DNA replication initiation
DNA replication elongation
Cholesterol synthesis
Glycolysis and Gluconeogenesis
Retinol metabolism
ECM breakdown by matrix metalloproteases

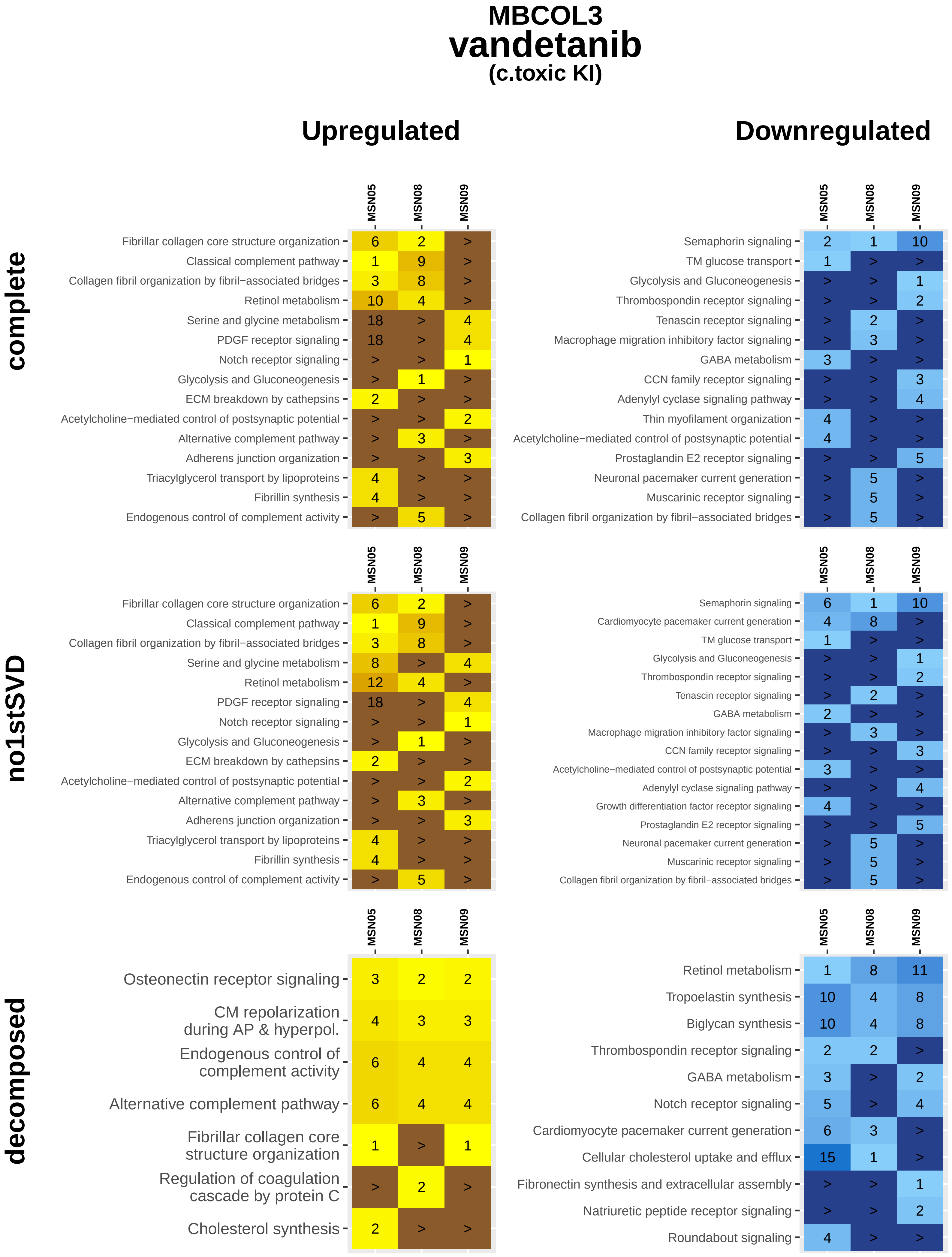
MBCOL3
vandetanib
(c.toxic KI)
Upregulated
Downregulated
complete
no1stSVD
decomposed
MSN05
MSN08
MSN09
Fibrillar collagen core structure organization 6 2 >
Classical complement pathway 1 9 >
Collagen fibril organization by fibril−associated bridges 3 8 >
Retinol metabolism 10 4 >
Serine and glycine metabolism 18 > 4
PDGF receptor signaling 18 > 4
Notch receptor signaling > > 1
Glycolysis and Gluconeogenesis > 1 >
ECM breakdown by cathepsins 2 > >
Acetylcholine−mediated control of postsynaptic potential > > 2
Alternative complement pathway > 3 >
Adherens junction organization > > 3
Triacylglycerol transport by lipoproteins 4 > >
Fibrillin synthesis 4 > >
Endogenous control of complement activity > 5 >
Semaphorin signaling 2 1 10
TM glucose transport 1 > >
Glycolysis and Gluconeogenesis > > 1
Thrombospondin receptor signaling > > 2
Tenascin receptor signaling > 2 >
Macrophage migration inhibitory factor signaling > 3 >
GABA metabolism 3 > >
CCN family receptor signaling > > 3
Adenylyl cyclase signaling pathway > > 4
Thin myofilament organization 4 > >
Acetylcholine−mediated control of postsynaptic potential 4 > >
Prostaglandin E2 receptor signaling > > 5
Neuronal pacemaker current generation > 5 >
Muscarinic receptor signaling > 5 >
Collagen fibril organization by fibril−associated bridges > 5 >
Fibrillar collagen core structure organization 6 2 >
Classical complement pathway 1 9 >
Collagen fibril organization by fibril−associated bridges 3 8 >
Serine and glycine metabolism 8 > 4
Retinol metabolism 12 4 >
PDGF receptor signaling 18 > 4
Notch receptor signaling > > 1
Glycolysis and Gluconeogenesis > 1 >
ECM breakdown by cathepsins 2 > >
Acetylcholine−mediated control of postsynaptic potential > > 2
Alternative complement pathway > 3 >
Adherens junction organization > > 3
Triacylglycerol transport by lipoproteins 4 > >
Fibrillin synthesis 4 > >
Endogenous control of complement activity > 5 >
Semaphorin signaling 6 1 10
Cardiomyocyte pacemaker current generation 4 8 >
TM glucose transport 1 > >
Glycolysis and Gluconeogenesis > > 1
Thrombospondin receptor signaling > > 2
Tenascin receptor signaling > 2 >
GABA metabolism 2 > >
Macrophage migration inhibitory factor signaling > 3 >
CCN family receptor signaling > > 3
Acetylcholine−mediated control of postsynaptic potential 3 > >
Adenylyl cyclase signaling pathway > > 4
Growth differentiation factor receptor signaling 4 > >
Prostaglandin E2 receptor signaling > > 5
Neuronal pacemaker current generation > 5 >
Muscarinic receptor signaling > 5 >
Collagen fibril organization by fibril−associated bridges > 5 >
Osteonectin receptor signaling 3 2 2
CM repolarization during AP & hyperpol. 4 3 3
Endogenous control of complement activity 6 4 4
Alternative complement pathway 6 4 4
Fibrillar collagen core structure organization 1 > 1
Regulation of coagulation cascade by protein C > 2 >
Cholesterol synthesis 2 > >
Retinol metabolism 1 8 11
Tropoelastin synthesis 10 4 8
Biglycan synthesis 10 4 8
Thrombospondin receptor signaling 2 2 >
GABA metabolism 3 > 2
Notch receptor signaling 5 > 4
Cardiomyocyte pacemaker current generation 6 3 >
Cellular cholesterol uptake and efflux 15 1 >
Fibronectin synthesis and extracellular assembly > > 1
Natriuretic peptide receptor signaling > > 2
Roundabout signaling 4 > >

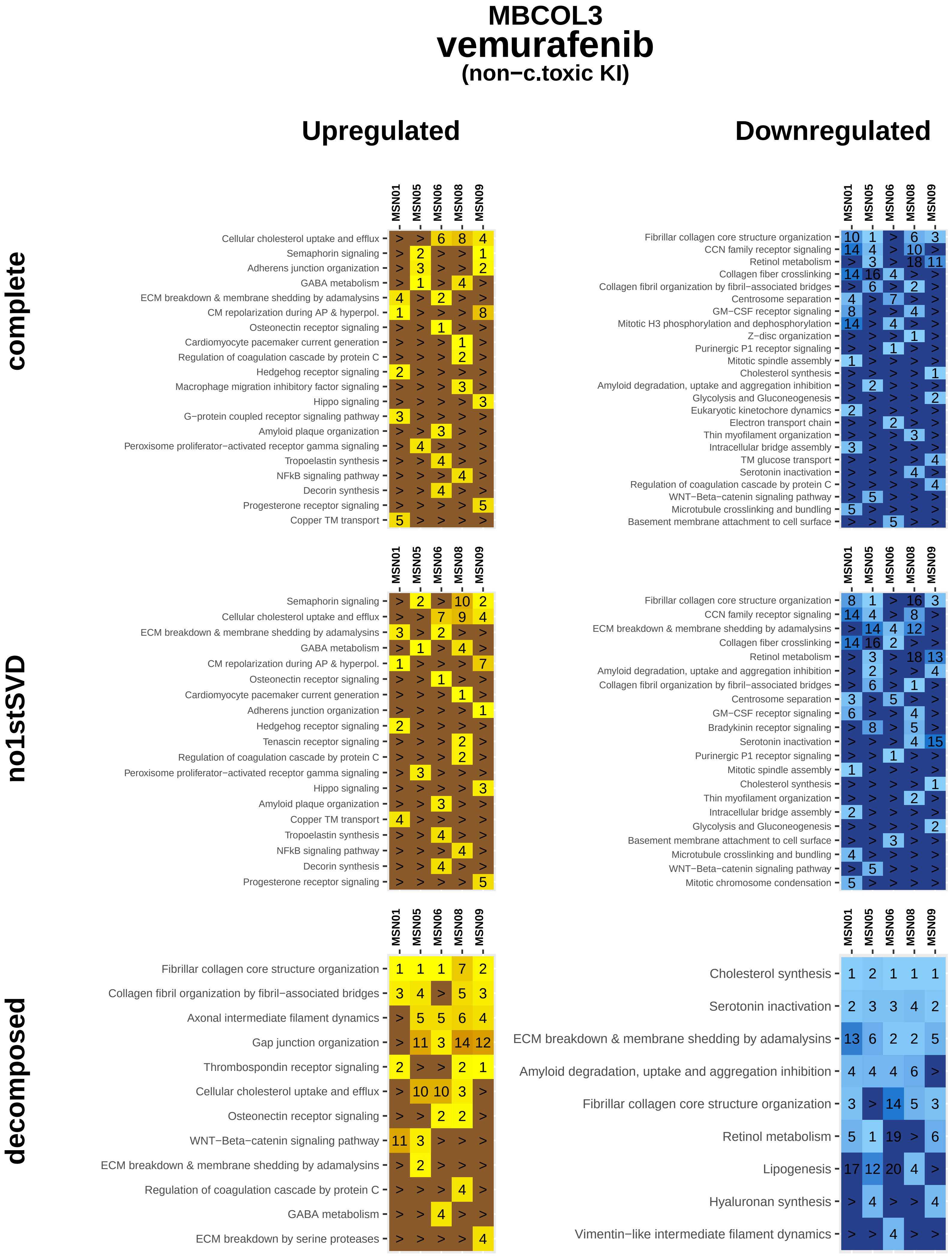
MBCOL3
vemurafenib
(non−c.toxic KI)
Upregulated
Downregulated
complete
no1stSVD
decomposed
MSN01
MSN05
MSN06
MSN08
MSN09
Cellular cholesterol uptake and efflux
Semaphorin signaling
Adherens junction organization
GABA metabolism
ECM breakdown & membrane shedding by adamalysins
CM repolarization during AP & hyperpol.
Osteonectin receptor signaling
Cardiomyocyte pacemaker current generation
Regulation of coagulation cascade by protein C
Hedgehog receptor signaling
Macrophage migration inhibitory factor signaling
Hippo signaling
G−protein coupled receptor signaling pathway
Amyloid plaque organization
Peroxisome proliferator−activated receptor gamma signaling
Tropoelastin synthesis
NFkB signaling pathway
Decorin synthesis
Progesterone receptor signaling
Copper TM transport
Fibrillar collagen core structure organization
CCN family receptor signaling
Retinol metabolism
Collagen fiber crosslinking
Collagen fibril organization by fibril−associated bridges
Centrosome separation
GM−CSF receptor signaling
Mitotic H3 phosphorylation and dephosphorylation
Z−disc organization
Purinergic P1 receptor signaling
Mitotic spindle assembly
Cholesterol synthesis
Amyloid degradation, uptake and aggregation inhibition
Glycolysis and Gluconeogenesis
Eukaryotic kinetochore dynamics
Electron transport chain
Thin myofilament organization
Intracellular bridge assembly
TM glucose transport
Serotonin inactivation
WNT−Beta−catenin signaling pathway
Microtubule crosslinking and bundling
Basement membrane attachment to cell surface
Tenascin receptor signaling
Bradykinin receptor signaling
Mitotic chromosome condensation
Axonal intermediate filament dynamics
Gap junction organization
Thrombospondin receptor signaling
ECM breakdown by serine proteases
Lipogenesis
Hyaluronan synthesis
Vimentin−like intermediate filament dynamics

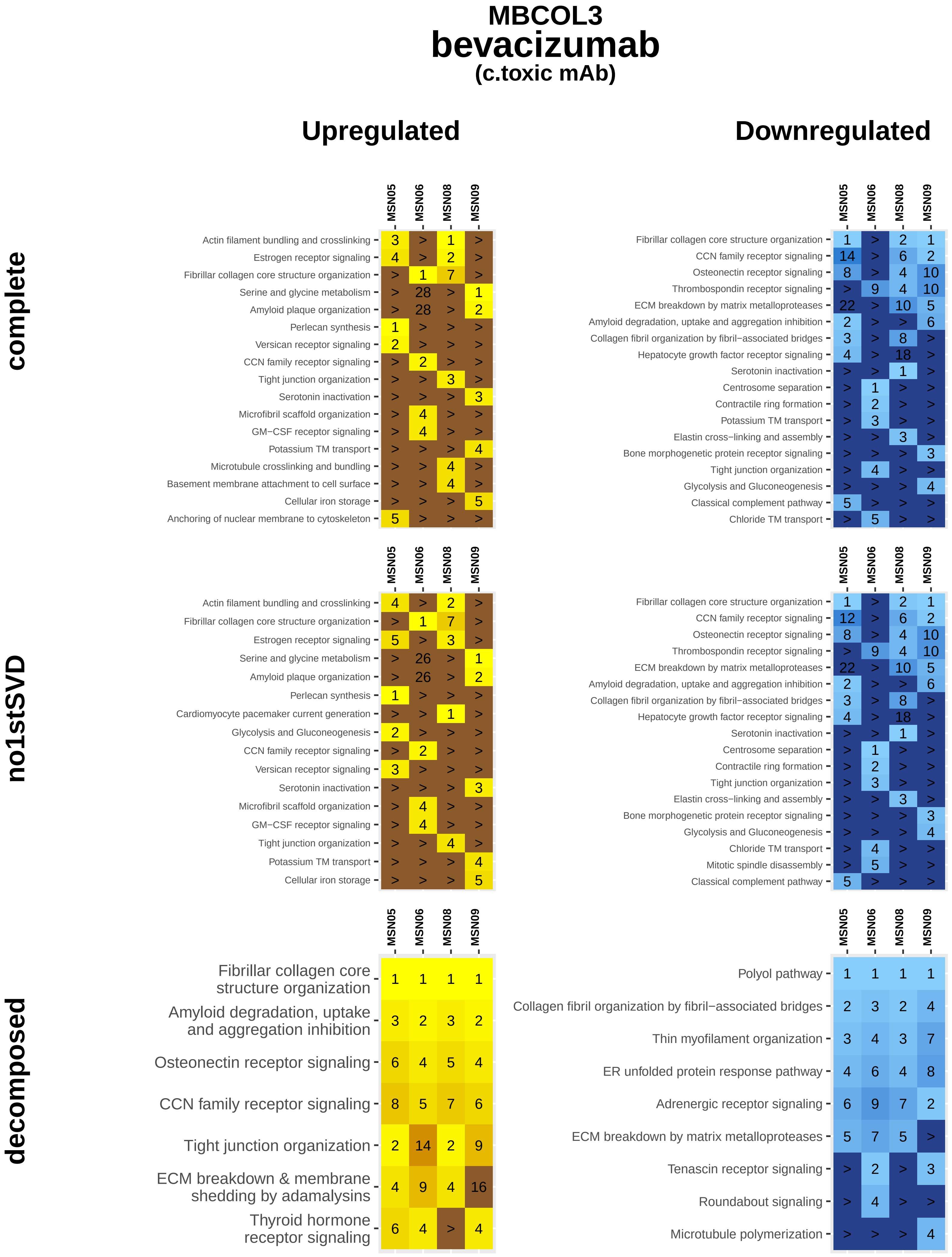
MBCOL3
bevacizumab
(c.toxic mAb)
Upregulated
Downregulated
complete
no1stSVD
decomposed
MSN05
MSN06
MSN08
MSN09
Actin filament bundling and crosslinking
Estrogen receptor signaling
Fibrillar collagen core structure organization
Serine and glycine metabolism
Amyloid plaque organization
Perlecan synthesis
Versican receptor signaling
CCN family receptor signaling
Tight junction organization
Serotonin inactivation
Microfibril scaffold organization
GM−CSF receptor signaling
Potassium TM transport
Microtubule crosslinking and bundling
Basement membrane attachment to cell surface
Cellular iron storage
Anchoring of nuclear membrane to cytoskeleton
Osteonectin receptor signaling
Thrombospondin receptor signaling
ECM breakdown by matrix metalloproteases
Amyloid degradation, uptake and aggregation inhibition
Collagen fibril organization by fibril−associated bridges
Hepatocyte growth factor receptor signaling
Centrosome separation
Contractile ring formation
Elastin cross−linking and assembly
Bone morphogenetic protein receptor signaling
Glycolysis and Gluconeogenesis
Classical complement pathway
Chloride TM transport
Cardiomyocyte pacemaker current generation
Mitotic spindle disassembly
ECM breakdown & membrane shedding by adamalysins
Thyroid hormone receptor signaling
Polyol pathway
Thin myofilament organization
ER unfolded protein response pathway
Adrenergic receptor signaling
Tenascin receptor signaling
Roundabout signaling
Microtubule polymerization

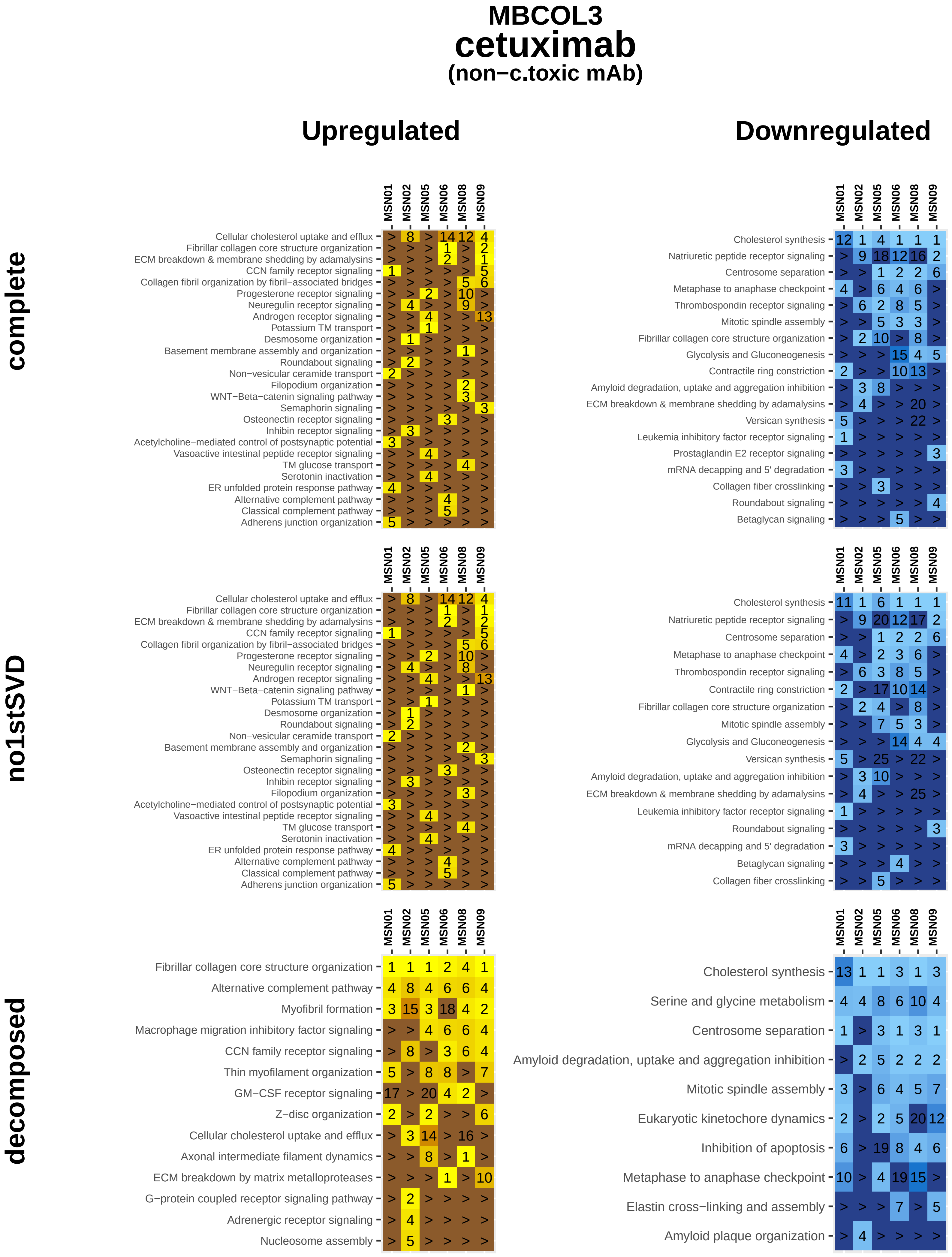

MBCOL3
cetuximab
(non−c.toxic mAb)
Upregulated
Downregulated
complete
no1stSVD
decomposed
MSN01
MSN02
MSN05
MSN06
MSN08
MSN09
Cellular cholesterol uptake and efflux
Fibrillar collagen core structure organization
ECM breakdown & membrane shedding by adamalysins
CCN family receptor signaling
Collagen fibril organization by fibril−associated bridges
Progesterone receptor signaling
Neuregulin receptor signaling
Androgen receptor signaling
Potassium TM transport
Desmosome organization
Basement membrane assembly and organization
Roundabout signaling
Non−vesicular ceramide transport
Filopodium organization
WNT−Beta−catenin signaling pathway
Semaphorin signaling
Osteonectin receptor signaling
Inhibin receptor signaling
Acetylcholine−mediated control of postsynaptic potential
Vasoactive intestinal peptide receptor signaling
TM glucose transport
Serotonin inactivation
ER unfolded protein response pathway
Alternative complement pathway
Classical complement pathway
Adherens junction organization
Cholesterol synthesis
Natriuretic peptide receptor signaling
Centrosome separation
Metaphase to anaphase checkpoint
Thrombospondin receptor signaling
Mitotic spindle assembly
Glycolysis and Gluconeogenesis
Contractile ring constriction
Amyloid degradation, uptake and aggregation inhibition
Versican synthesis
Leukemia inhibitory factor receptor signaling
Prostaglandin E2 receptor signaling
mRNA decapping and 5' degradation
Collagen fiber crosslinking
Betaglycan signaling
Myofibril formation
Macrophage migration inhibitory factor signaling
Thin myofilament organization
GM−CSF receptor signaling
Z−disc organization
Axonal intermediate filament dynamics
ECM breakdown by matrix metalloproteases
G−protein coupled receptor signaling pathway
Adrenergic receptor signaling
Nucleosome assembly
Serine and glycine metabolism
Eukaryotic kinetochore dynamics
Inhibition of apoptosis
Elastin cross−linking and assembly
Amyloid plaque organization

### MBCOL3
### rituximab
#### (non−c.toxic mAb)

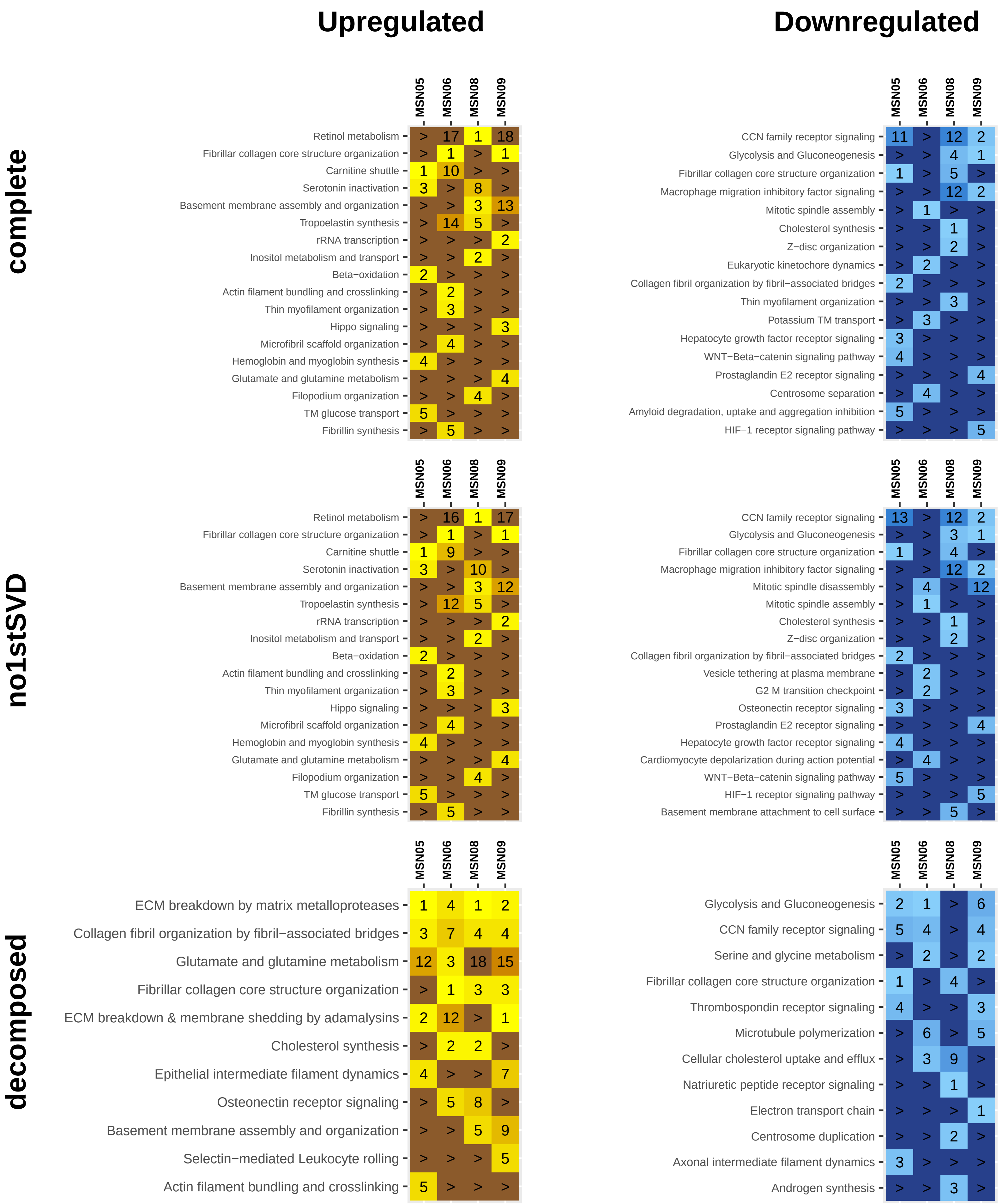

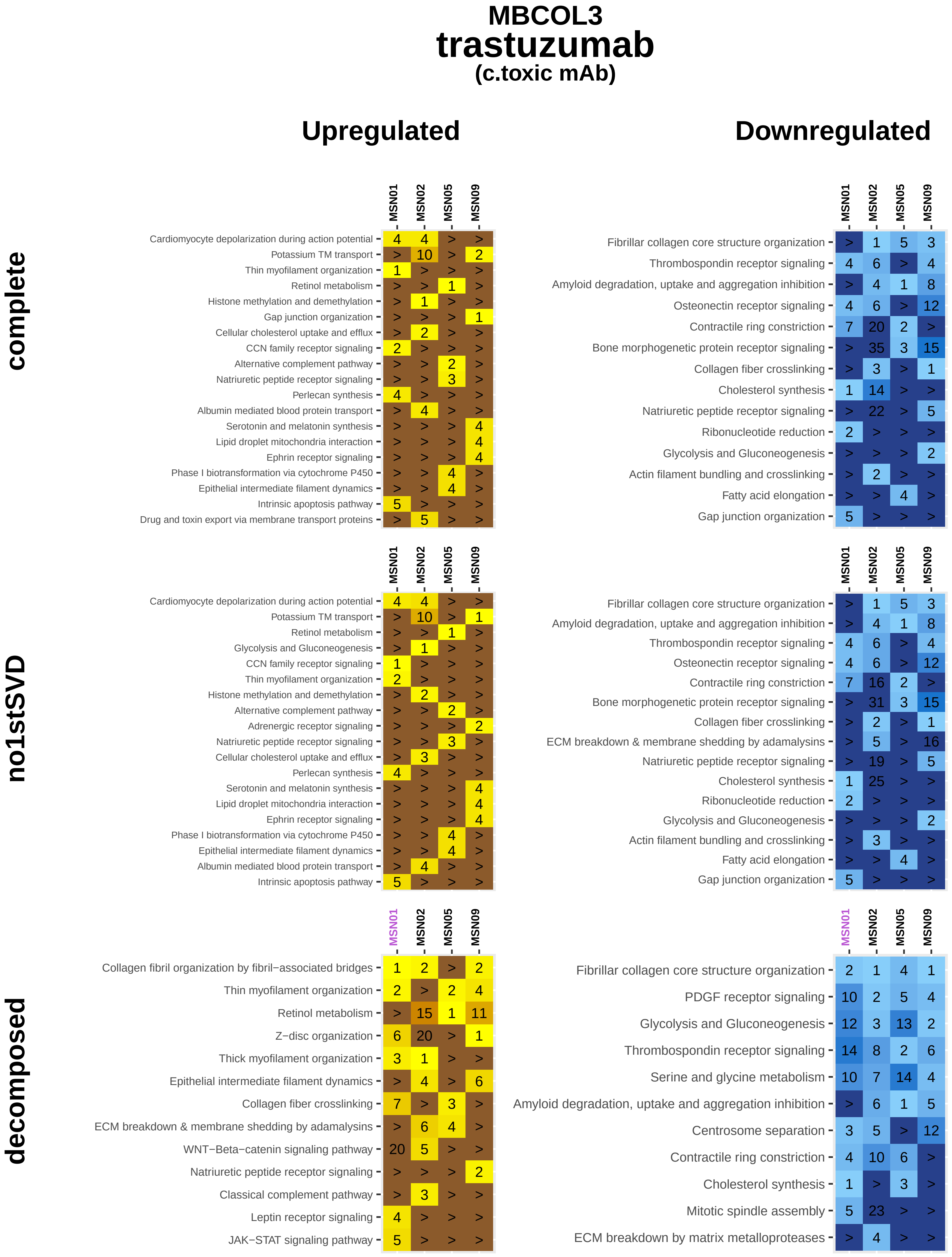
MBCOL3
trastuzumab
(c.toxic mAb)
Upregulated
Downregulated
complete
no1stSVD
decomposed
MSN01
MSN02
MSN05
MSN09
Cardiomyocyte depolarization during action potential
Potassium TM transport
Thin myofilament organization
Retinol metabolism
Histone methylation and demethylation
Gap junction organization
Cellular cholesterol uptake and efflux
CCN family receptor signaling
Alternative complement pathway
Natriuretic peptide receptor signaling
Perlecan synthesis
Albumin mediated blood protein transport
Serotonin and melatonin synthesis
Lipid droplet mitochondria interaction
Ephrin receptor signaling
Phase I biotransformation via cytochrome P450
Epithelial intermediate filament dynamics
Intrinsic apoptosis pathway
Drug and toxin export via membrane transport proteins
Fibrillar collagen core structure organization
Thrombospondin receptor signaling
Amyloid degradation, uptake and aggregation inhibition
Osteonectin receptor signaling
Contractile ring constriction
Bone morphogenetic protein receptor signaling
Collagen fiber crosslinking
Cholesterol synthesis
Ribonucleotide reduction
Glycolysis and Gluconeogenesis
Actin filament bundling and crosslinking
Fatty acid elongation
Adrenergic receptor signaling
ECM breakdown & membrane shedding by adamalysins
Collagen fibril organization by fibril−associated bridges
Z−disc organization
Thick myofilament organization
WNT−Beta−catenin signaling pathway
Classical complement pathway
Leptin receptor signaling
JAK−STAT signaling pathway
PDGF receptor signaling
Serine and glycine metabolism
Centrosome separation
Mitotic spindle assembly
ECM breakdown by matrix metalloproteases

### MBCOL3
### daunorubicin
##### (anthracycline)

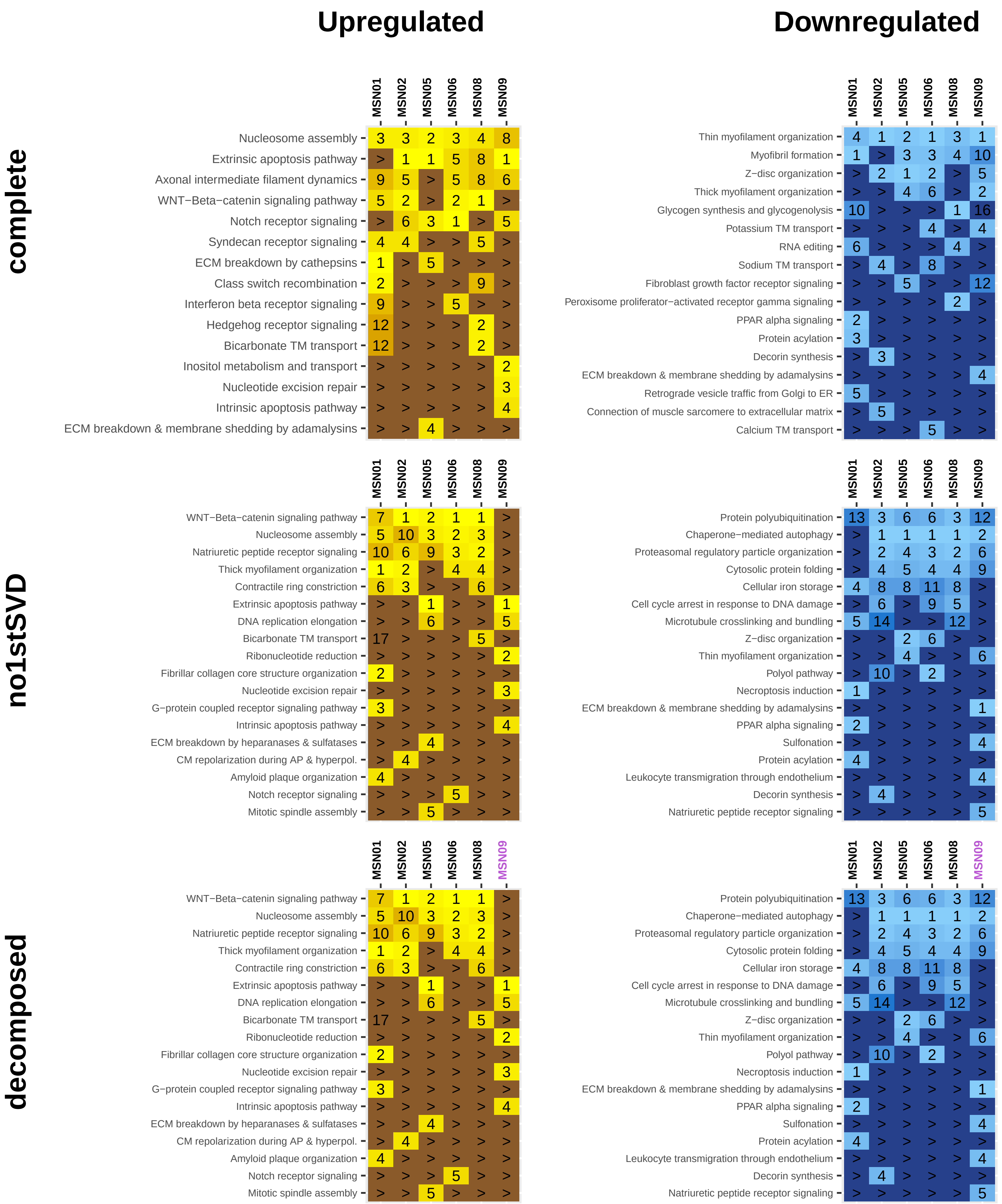

### MBCOL3
### doxorubicin
#### (anthracycline)

#### complete

##### Upregulated

| | MSN05 | MSN06 | MSN08 | MSN09 |
|---|---|---|---|---|
| Nucleosome assembly | 2 | 3 | 1 | 2 |
| Extrinsic apoptosis pathway | 6 | 6 | 8 | 1 |
| Syndecan receptor signaling | 4 | 2 | 4 | > |
| WNT−Beta−catenin signaling pathway | 7 | 1 | 3 | > |
| Axonal intermediate filament dynamics | 6 | 6 | 2 | > |
| Prostaglandin E2 receptor signaling | > | 5 | 6 | > |
| Motilin receptor signaling | 1 | > | > | > |
| Intrinsic apoptosis pathway | > | > | > | 3 |
| Cannabinoid receptor signaling | 4 | > | > | > |
| Hedgehog receptor signaling | > | 4 | > | > |
| Acetylcholine−mediated control of postsynaptic potential | > | > | > | 4 |
| Notch receptor signaling | > | > | > | 5 |
| Nodal growth differentiation factor receptor signaling | > | > | 5 | > |

##### Downregulated

| | MSN05 | MSN06 | MSN08 | MSN09 |
|---|---|---|---|---|
| Thin myofilament organization | 2 | > | 1 | 1 |
| Actin polymerization | 5 | > | 3 | > |
| Myofibril formation | > | 2 | > | 7 |
| Progestagen synthesis | 1 | > | > | > |
| Mitochondrial transport | > | 2 | > | > |
| Potassium TM transport | > | > | > | 2 |
| Glycolysis and Gluconeogenesis | > | > | 2 | > |
| Transamination pathways | 2 | > | > | > |
| Sulfonation | > | > | > | 4 |
| Leukocyte transmigration through endothelium | > | > | > | 4 |
| Androgen synthesis | 4 | > | > | > |
| Natriuretic peptide receptor signaling | > | > | > | 5 |

#### no1stSVD

##### Upregulated

| | MSN05 | MSN06 | MSN08 | MSN09 |
|---|---|---|---|---|
| WNT−Beta−catenin signaling pathway | 3 | 1 | 1 | > |
| Natriuretic peptide receptor signaling | 1 | 3 | 2 | > |
| Nucleosome assembly | 2 | 4 | 5 | > |
| Thick myofilament organization | 4 | 5 | 4 | > |
| Extrinsic apoptosis pathway | > | 10 | > | 2 |
| Nucleotide excision repair | > | > | > | 1 |
| Syndecan receptor signaling | > | 2 | > | > |
| Nodal growth differentiation factor receptor signaling | > | > | 3 | > |
| Inositol metabolism and transport | > | > | > | 3 |
| Ribonucleotide reduction | > | > | > | 4 |
| Intrinsic apoptosis pathway | > | > | > | 5 |

##### Downregulated

| | MSN05 | MSN06 | MSN08 | MSN09 |
|---|---|---|---|---|
| Chaperone−mediated autophagy | 1 | 2 | 2 | 1 |
| Cellular iron storage | 6 | 4 | 4 | > |
| Protein polyubiquitination | 2 | > | 1 | 12 |
| Hepatocyte growth factor receptor signaling | 4 | > | 2 | 14 |
| Cytosolic protein folding | 3 | > | > | 10 |
| CD44−Mediated leukocyte rolling | 4 | > | > | 14 |
| Necroptosis induction | > | 1 | > | > |
| ECM breakdown & membrane shedding by adamalysins | > | > | > | 2 |
| Z−disc organization | > | > | > | 3 |
| Heme degradation to bilirubin | > | 4 | > | > |
| Collagen fiber crosslinking | > | > | > | 4 |
| Natriuretic peptide receptor signaling | > | > | > | 5 |

#### decomposed

##### Upregulated

| | MSN05 | MSN06 | MSN08 | MSN09 |
|---|---|---|---|---|
| WNT−Beta−catenin signaling pathway | 3 | 1 | 1 | > |
| Natriuretic peptide receptor signaling | 1 | 3 | 2 | > |
| Nucleosome assembly | 2 | 4 | 5 | > |
| Thick myofilament organization | 4 | 5 | 4 | > |
| Extrinsic apoptosis pathway | > | 10 | > | 2 |
| Nucleotide excision repair | > | > | > | 1 |
| Syndecan receptor signaling | > | 2 | > | > |
| Nodal growth differentiation factor receptor signaling | > | > | 3 | > |
| Inositol metabolism and transport | > | > | > | 3 |
| Ribonucleotide reduction | > | > | > | 4 |
| Intrinsic apoptosis pathway | > | > | > | 5 |

##### Downregulated

| | MSN05 | MSN06 | MSN08 | MSN09 |
|---|---|---|---|---|
| Chaperone−mediated autophagy | 1 | 2 | 2 | 1 |
| Cellular iron storage | 6 | 4 | 4 | > |
| Protein polyubiquitination | 2 | > | 1 | 12 |
| Hepatocyte growth factor receptor signaling | 4 | > | 2 | 14 |
| Cytosolic protein folding | 3 | > | > | 10 |
| CD44−Mediated leukocyte rolling | 4 | > | > | 14 |
| Necroptosis induction | > | 1 | > | > |
| ECM breakdown & membrane shedding by adamalysins | > | > | > | 2 |
| Z−disc organization | > | > | > | 3 |
| Heme degradation to bilirubin | > | 4 | > | > |
| Collagen fiber crosslinking | > | > | > | 4 |
| Natriuretic peptide receptor signaling | > | > | > | 5 |

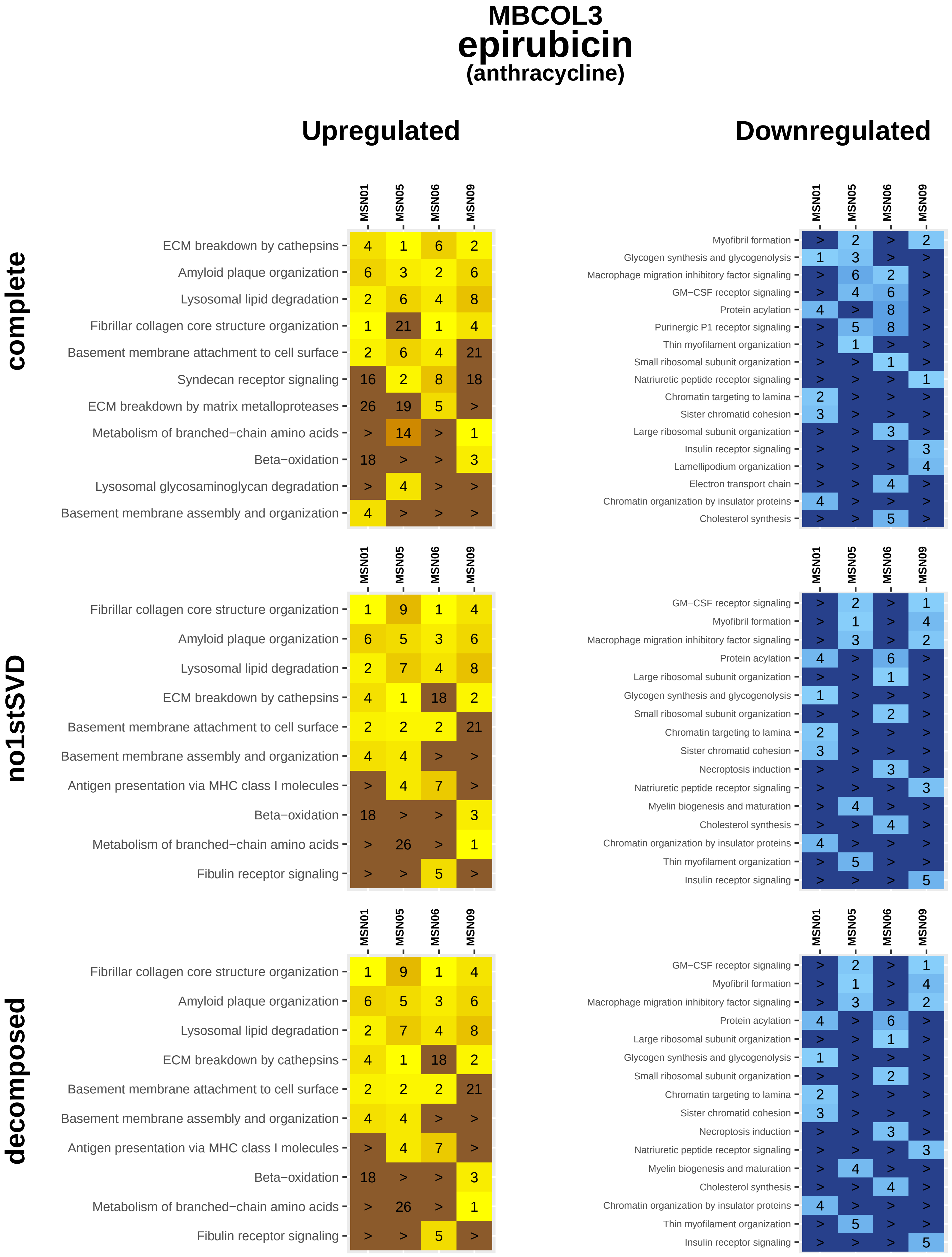
MBCOL3
epirubicin
(anthracycline)
Upregulated
Downregulated
complete
no1stSVD
decomposed
MSN01 MSN05 MSN06 MSN09
ECM breakdown by cathepsins 4 1 6 2
Amyloid plaque organization 6 3 2 6
Lysosomal lipid degradation 2 6 4 8
Fibrillar collagen core structure organization 1 21 1 4
Basement membrane attachment to cell surface 2 6 4 21
Syndecan receptor signaling 16 2 8 18
ECM breakdown by matrix metalloproteases 26 19 5 >
Metabolism of branched−chain amino acids > 14 > 1
Beta−oxidation 18 > > 3
Lysosomal glycosaminoglycan degradation > 4 > >
Basement membrane assembly and organization 4 > > >
MSN01 MSN05 MSN06 MSN09
Myofibril formation > 2 > 2
Glycogen synthesis and glycogenolysis 1 3 > >
Macrophage migration inhibitory factor signaling > 6 2 >
GM−CSF receptor signaling > 4 6 >
Protein acylation 4 > 8 >
Purinergic P1 receptor signaling > 5 8 >
Thin myofilament organization > 1 > >
Small ribosomal subunit organization > > 1 >
Natriuretic peptide receptor signaling > > > 1
Chromatin targeting to lamina 2 > > >
Sister chromatid cohesion 3 > > >
Large ribosomal subunit organization > > 3 >
Insulin receptor signaling > > > 3
Lamellipodium organization > > > 4
Electron transport chain > > 4 >
Chromatin organization by insulator proteins 4 > > >
Cholesterol synthesis > > 5 >
MSN01 MSN05 MSN06 MSN09
Fibrillar collagen core structure organization 1 9 1 4
Amyloid plaque organization 6 5 3 6
Lysosomal lipid degradation 2 7 4 8
ECM breakdown by cathepsins 4 1 18 2
Basement membrane attachment to cell surface 2 2 2 21
Basement membrane assembly and organization 4 4 > >
Antigen presentation via MHC class I molecules > 4 7 >
Beta−oxidation 18 > > 3
Metabolism of branched−chain amino acids > 26 > 1
Fibulin receptor signaling > > 5 >
MSN01 MSN05 MSN06 MSN09
GM−CSF receptor signaling > 2 > 1
Myofibril formation > 1 > 4
Macrophage migration inhibitory factor signaling > 3 > 2
Protein acylation 4 > 6 >
Large ribosomal subunit organization > > 1 >
Glycogen synthesis and glycogenolysis 1 > > >
Small ribosomal subunit organization > > 2 >
Chromatin targeting to lamina 2 > > >
Sister chromatid cohesion 3 > > >
Necroptosis induction > > 3 >
Natriuretic peptide receptor signaling > > > 3
Myelin biogenesis and maturation > 4 > >
Cholesterol synthesis > > 4 >
Chromatin organization by insulator proteins 4 > > >
Thin myofilament organization > 5 > >
Insulin receptor signaling > > > 5
MSN01 MSN05 MSN06 MSN09
Fibrillar collagen core structure organization 1 9 1 4
Amyloid plaque organization 6 5 3 6
Lysosomal lipid degradation 2 7 4 8
ECM breakdown by cathepsins 4 1 18 2
Basement membrane attachment to cell surface 2 2 2 21
Basement membrane assembly and organization 4 4 > >
Antigen presentation via MHC class I molecules > 4 7 >
Beta−oxidation 18 > > 3
Metabolism of branched−chain amino acids > 26 > 1
Fibulin receptor signaling > > 5 >
MSN01 MSN05 MSN06 MSN09
GM−CSF receptor signaling > 2 > 1
Myofibril formation > 1 > 4
Macrophage migration inhibitory factor signaling > 3 > 2
Protein acylation 4 > 6 >
Large ribosomal subunit organization > > 1 >
Glycogen synthesis and glycogenolysis 1 > > >
Small ribosomal subunit organization > > 2 >
Chromatin targeting to lamina 2 > > >
Sister chromatid cohesion 3 > > >
Necroptosis induction > > 3 >
Natriuretic peptide receptor signaling > > > 3
Myelin biogenesis and maturation > 4 > >
Cholesterol synthesis > > 4 >
Chromatin organization by insulator proteins 4 > > >
Thin myofilament organization > 5 > >
Insulin receptor signaling > > > 5

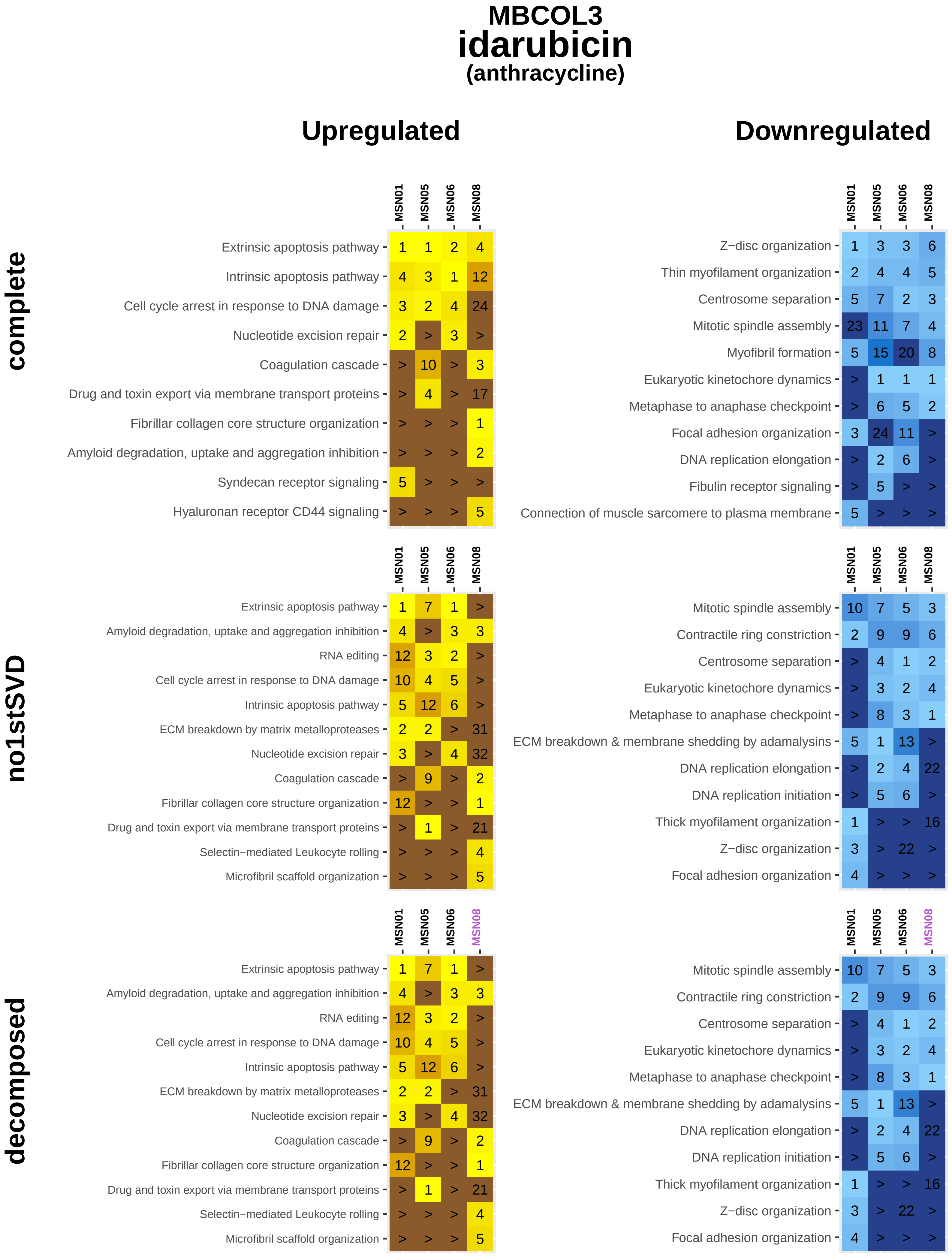

MBCOL3
idarubicin
(anthracycline)
Upregulated
Downregulated
complete
MSN01 MSN05 MSN06 MSN08
Extrinsic apoptosis pathway 1 1 2 4
Intrinsic apoptosis pathway 4 3 1 12
Cell cycle arrest in response to DNA damage 3 2 4 24
Nucleotide excision repair 2 > 3 >
Coagulation cascade > 10 > 3
Drug and toxin export via membrane transport proteins > 4 > 17
Fibrillar collagen core structure organization > > > 1
Amyloid degradation, uptake and aggregation inhibition > > > 2
Syndecan receptor signaling 5 > > >
Hyaluronan receptor CD44 signaling > > > 5
MSN01 MSN05 MSN06 MSN08
Z−disc organization 1 3 3 6
Thin myofilament organization 2 4 4 5
Centrosome separation 5 7 2 3
Mitotic spindle assembly 23 11 7 4
Myofibril formation 5 15 20 8
Eukaryotic kinetochore dynamics > 1 1 1
Metaphase to anaphase checkpoint > 6 5 2
Focal adhesion organization 3 24 11 >
DNA replication elongation > 2 6 >
Fibulin receptor signaling > 5 > >
Connection of muscle sarcomere to plasma membrane 5 > > >
no1stSVD
MSN01 MSN05 MSN06 MSN08
Extrinsic apoptosis pathway 1 7 1 >
Amyloid degradation, uptake and aggregation inhibition 4 > 3 3
RNA editing 12 3 2 >
Cell cycle arrest in response to DNA damage 10 4 5 >
Intrinsic apoptosis pathway 5 12 6 >
ECM breakdown by matrix metalloproteases 2 2 > 31
Nucleotide excision repair 3 > 4 32
Coagulation cascade > 9 > 2
Fibrillar collagen core structure organization 12 > > 1
Drug and toxin export via membrane transport proteins > 1 > 21
Selectin−mediated Leukocyte rolling > > > 4
Microfibril scaffold organization > > > 5
MSN01 MSN05 MSN06 MSN08
Mitotic spindle assembly 10 7 5 3
Contractile ring constriction 2 9 9 6
Centrosome separation > 4 1 2
Eukaryotic kinetochore dynamics > 3 2 4
Metaphase to anaphase checkpoint > 8 3 1
ECM breakdown & membrane shedding by adamalysins 5 1 13 >
DNA replication elongation > 2 4 22
DNA replication initiation > 5 6 >
Thick myofilament organization 1 > > 16
Z−disc organization 3 > 22 >
Focal adhesion organization 4 > > >
decomposed
MSN01 MSN05 MSN06 MSN08
Extrinsic apoptosis pathway 1 7 1 >
Amyloid degradation, uptake and aggregation inhibition 4 > 3 3
RNA editing 12 3 2 >
Cell cycle arrest in response to DNA damage 10 4 5 >
Intrinsic apoptosis pathway 5 12 6 >
ECM breakdown by matrix metalloproteases 2 2 > 31
Nucleotide excision repair 3 > 4 32
Coagulation cascade > 9 > 2
Fibrillar collagen core structure organization 12 > > 1
Drug and toxin export via membrane transport proteins > 1 > 21
Selectin−mediated Leukocyte rolling > > > 4
Microfibril scaffold organization > > > 5
MSN01 MSN05 MSN06 MSN08
Mitotic spindle assembly 10 7 5 3
Contractile ring constriction 2 9 9 6
Centrosome separation > 4 1 2
Eukaryotic kinetochore dynamics > 3 2 4
Metaphase to anaphase checkpoint > 8 3 1
ECM breakdown & membrane shedding by adamalysins 5 1 13 >
DNA replication elongation > 2 4 22
DNA replication initiation > 5 6 >
Thick myofilament organization 1 > > 16
Z−disc organization 3 > 22 >
Focal adhesion organization 4 > > >

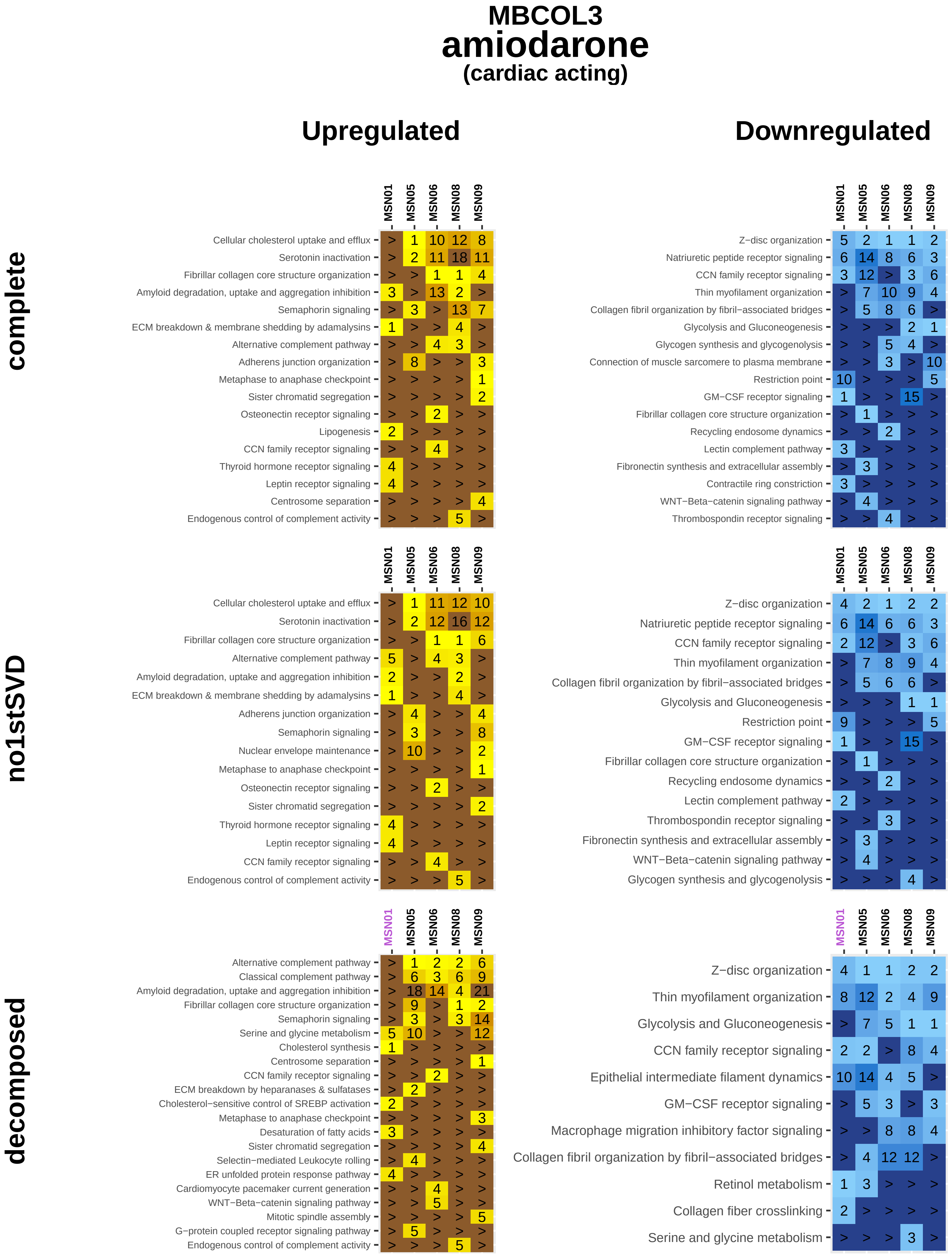
MBCOL3
amiodarone
(cardiac acting)
Upregulated
Downregulated
complete
no1stSVD
decomposed
MSN01
MSN05
MSN06
MSN08
MSN09
Cellular cholesterol uptake and efflux
Serotonin inactivation
Fibrillar collagen core structure organization
Amyloid degradation, uptake and aggregation inhibition
Semaphorin signaling
ECM breakdown & membrane shedding by adamalysins
Alternative complement pathway
Adherens junction organization
Metaphase to anaphase checkpoint
Sister chromatid segregation
Osteonectin receptor signaling
Lipogenesis
CCN family receptor signaling
Thyroid hormone receptor signaling
Leptin receptor signaling
Centrosome separation
Endogenous control of complement activity
Z−disc organization
Natriuretic peptide receptor signaling
Thin myofilament organization
Collagen fibril organization by fibril−associated bridges
Glycolysis and Gluconeogenesis
Glycogen synthesis and glycogenolysis
Connection of muscle sarcomere to plasma membrane
Restriction point
GM−CSF receptor signaling
Recycling endosome dynamics
Lectin complement pathway
Fibronectin synthesis and extracellular assembly
Contractile ring constriction
WNT−Beta−catenin signaling pathway
Thrombospondin receptor signaling
Nuclear envelope maintenance
Classical complement pathway
Serine and glycine metabolism
Cholesterol synthesis
ECM breakdown by heparanases & sulfatases
Cholesterol−sensitive control of SREBP activation
Desaturation of fatty acids
Selectin−mediated Leukocyte rolling
ER unfolded protein response pathway
Cardiomyocyte pacemaker current generation
Mitotic spindle assembly
G−protein coupled receptor signaling pathway
Epithelial intermediate filament dynamics
Macrophage migration inhibitory factor signaling
Retinol metabolism
Collagen fiber crosslinking

### MBCOL3
### dobutamine
#### (cardiac acting)

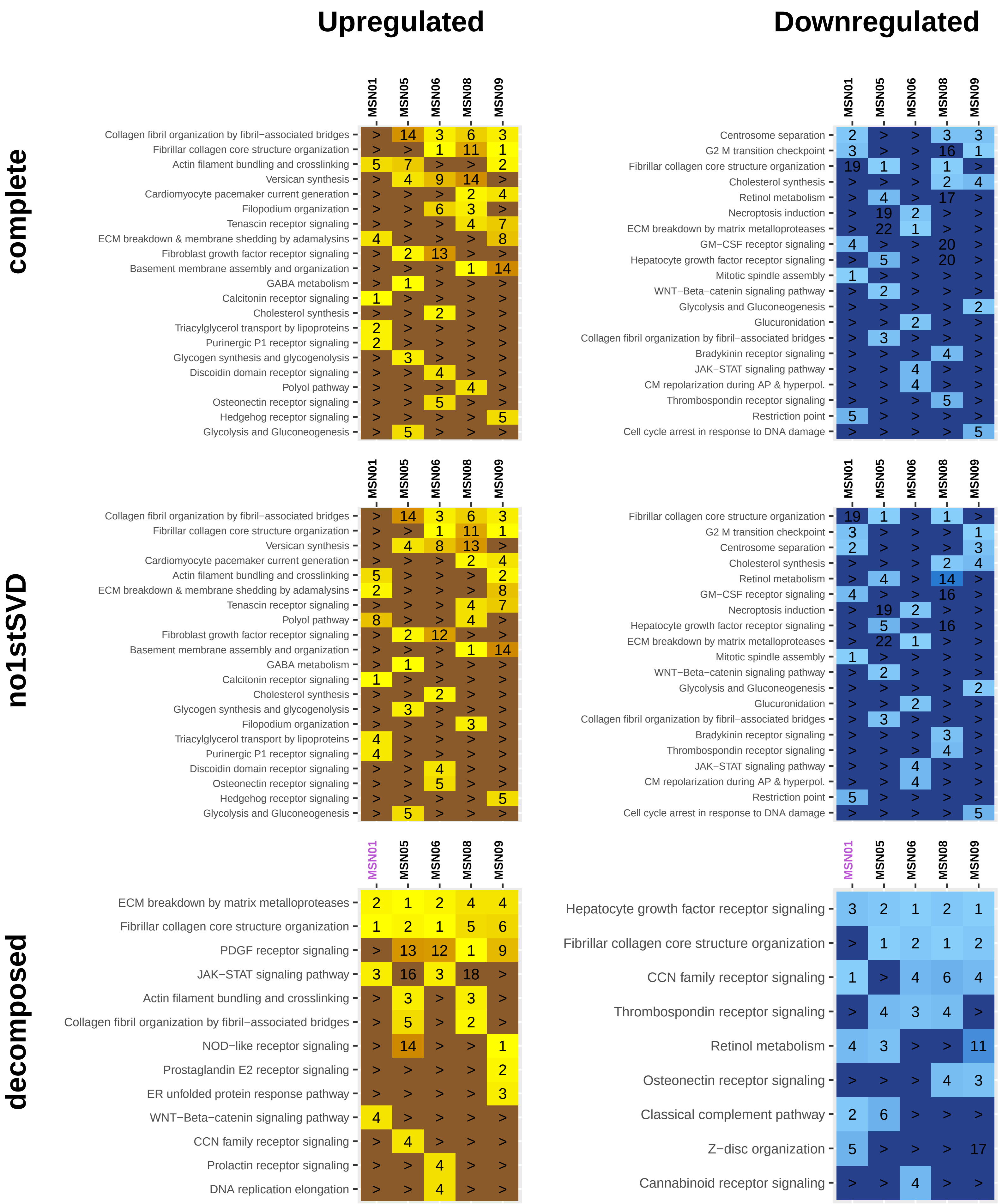

### MBCOL3
### flecainide
#### (cardiac acting)

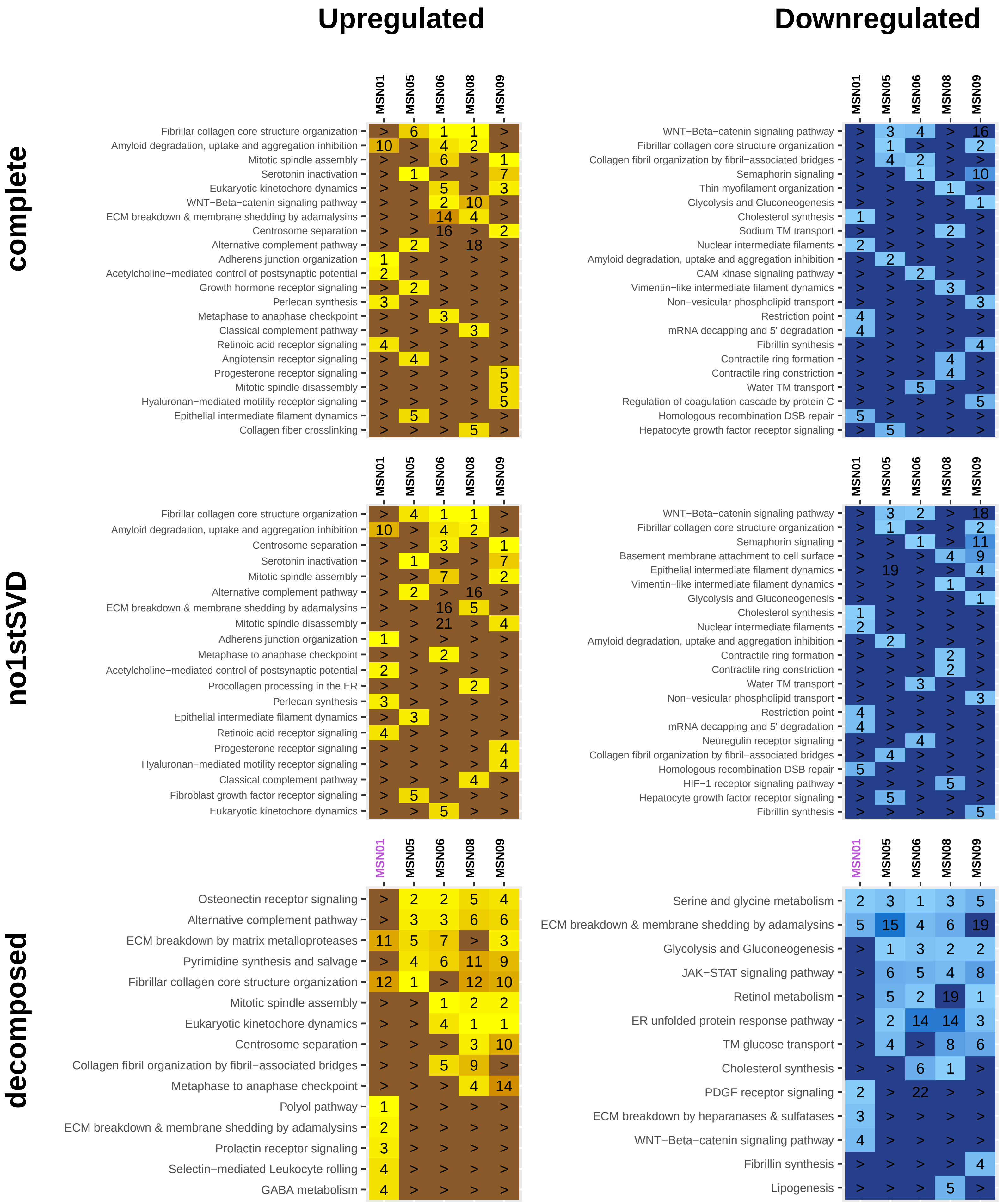

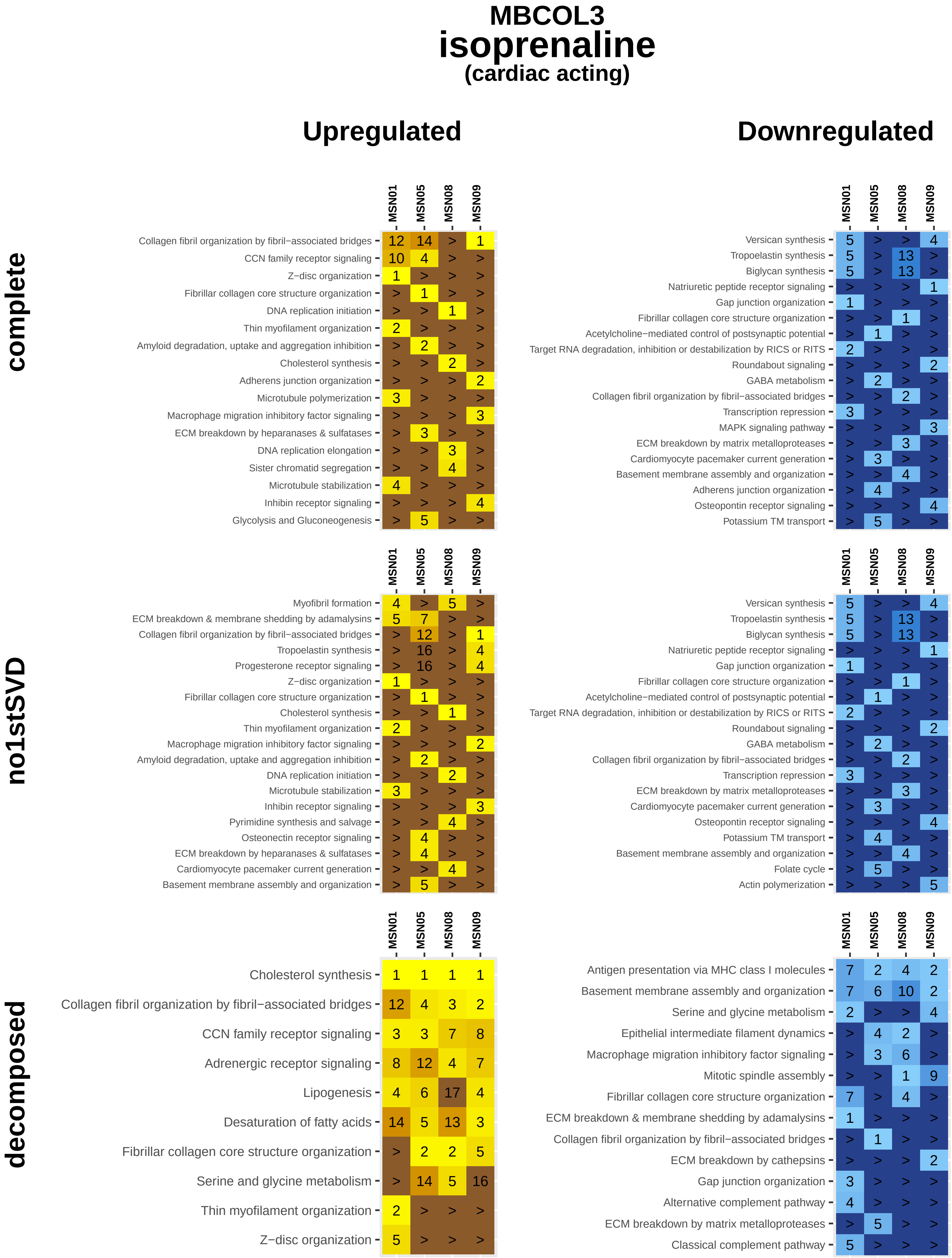
MBCOL3
isoprenaline
(cardiac acting)
Upregulated
Downregulated
complete
MSN01 MSN05 MSN08 MSN09
Collagen fibril organization by fibril−associated bridges 12 14 > 1
CCN family receptor signaling 10 4 > >
Z−disc organization 1 > > >
Fibrillar collagen core structure organization > 1 > >
DNA replication initiation > > 1 >
Thin myofilament organization 2 > > >
Amyloid degradation, uptake and aggregation inhibition > 2 > >
Cholesterol synthesis > > 2 >
Adherens junction organization > > > 2
Microtubule polymerization 3 > > >
Macrophage migration inhibitory factor signaling > > > 3
ECM breakdown by heparanases & sulfatases > 3 > >
DNA replication elongation > > 3 >
Sister chromatid segregation > > 4 >
Microtubule stabilization 4 > > >
Inhibin receptor signaling > > > 4
Glycolysis and Gluconeogenesis > 5 > >
MSN01 MSN05 MSN08 MSN09
Versican synthesis 5 > > 4
Tropoelastin synthesis 5 > 13 >
Biglycan synthesis 5 > 13 >
Natriuretic peptide receptor signaling > > > 1
Gap junction organization 1 > > >
Fibrillar collagen core structure organization > > 1 >
Acetylcholine−mediated control of postsynaptic potential > 1 > >
Target RNA degradation, inhibition or destabilization by RICS or RITS 2 > > >
Roundabout signaling > > > 2
GABA metabolism > 2 > >
Collagen fibril organization by fibril−associated bridges > > 2 >
Transcription repression 3 > > >
MAPK signaling pathway > > > 3
ECM breakdown by matrix metalloproteases > > 3 >
Cardiomyocyte pacemaker current generation > 3 > >
Basement membrane assembly and organization > > 4 >
Adherens junction organization > 4 > >
Osteopontin receptor signaling > > > 4
Potassium TM transport > 5 > >
no1stSVD
MSN01 MSN05 MSN08 MSN09
Myofibril formation 4 > 5 >
ECM breakdown & membrane shedding by adamalysins 5 7 > >
Collagen fibril organization by fibril−associated bridges > 12 > 1
Tropoelastin synthesis > 16 > 4
Progesterone receptor signaling > 16 > 4
Z−disc organization 1 > > >
Fibrillar collagen core structure organization > 1 > >
Cholesterol synthesis > > 1 >
Thin myofilament organization 2 > > >
Macrophage migration inhibitory factor signaling > > > 2
Amyloid degradation, uptake and aggregation inhibition > 2 > >
DNA replication initiation > > 2 >
Microtubule stabilization 3 > > >
Inhibin receptor signaling > > > 3
Pyrimidine synthesis and salvage > > 4 >
Osteonectin receptor signaling > 4 > >
ECM breakdown by heparanases & sulfatases > 4 > >
Cardiomyocyte pacemaker current generation > > 4 >
Basement membrane assembly and organization > 5 > >
MSN01 MSN05 MSN08 MSN09
Versican synthesis 5 > > 4
Tropoelastin synthesis 5 > 13 >
Biglycan synthesis 5 > 13 >
Natriuretic peptide receptor signaling > > > 1
Gap junction organization 1 > > >
Fibrillar collagen core structure organization > > 1 >
Acetylcholine−mediated control of postsynaptic potential > 1 > >
Target RNA degradation, inhibition or destabilization by RICS or RITS 2 > > >
Roundabout signaling > > > 2
GABA metabolism > 2 > >
Collagen fibril organization by fibril−associated bridges > > 2 >
Transcription repression 3 > > >
ECM breakdown by matrix metalloproteases > > 3 >
Cardiomyocyte pacemaker current generation > 3 > >
Osteopontin receptor signaling > > > 4
Potassium TM transport > 4 > >
Basement membrane assembly and organization > > 4 >
Folate cycle > 5 > >
Actin polymerization > > > 5
decomposed
MSN01 MSN05 MSN08 MSN09
Cholesterol synthesis 1 1 1 1
Collagen fibril organization by fibril−associated bridges 12 4 3 2
CCN family receptor signaling 3 3 7 8
Adrenergic receptor signaling 8 12 4 7
Lipogenesis 4 6 17 4
Desaturation of fatty acids 14 5 13 3
Fibrillar collagen core structure organization > 2 2 5
Serine and glycine metabolism > 14 5 16
Thin myofilament organization 2 > > >
Z−disc organization 5 > > >
MSN01 MSN05 MSN08 MSN09
Antigen presentation via MHC class I molecules 7 2 4 2
Basement membrane assembly and organization 7 6 10 2
Serine and glycine metabolism 2 > > 4
Epithelial intermediate filament dynamics > 4 2 >
Macrophage migration inhibitory factor signaling > 3 6 >
Mitotic spindle assembly > > 1 9
Fibrillar collagen core structure organization 7 > 4 >
ECM breakdown & membrane shedding by adamalysins 1 > > >
Collagen fibril organization by fibril−associated bridges > 1 > >
ECM breakdown by cathepsins > > > 2
Gap junction organization 3 > > >
Alternative complement pathway 4 > > >
ECM breakdown by matrix metalloproteases > 5 > >
Classical complement pathway 5 > > >

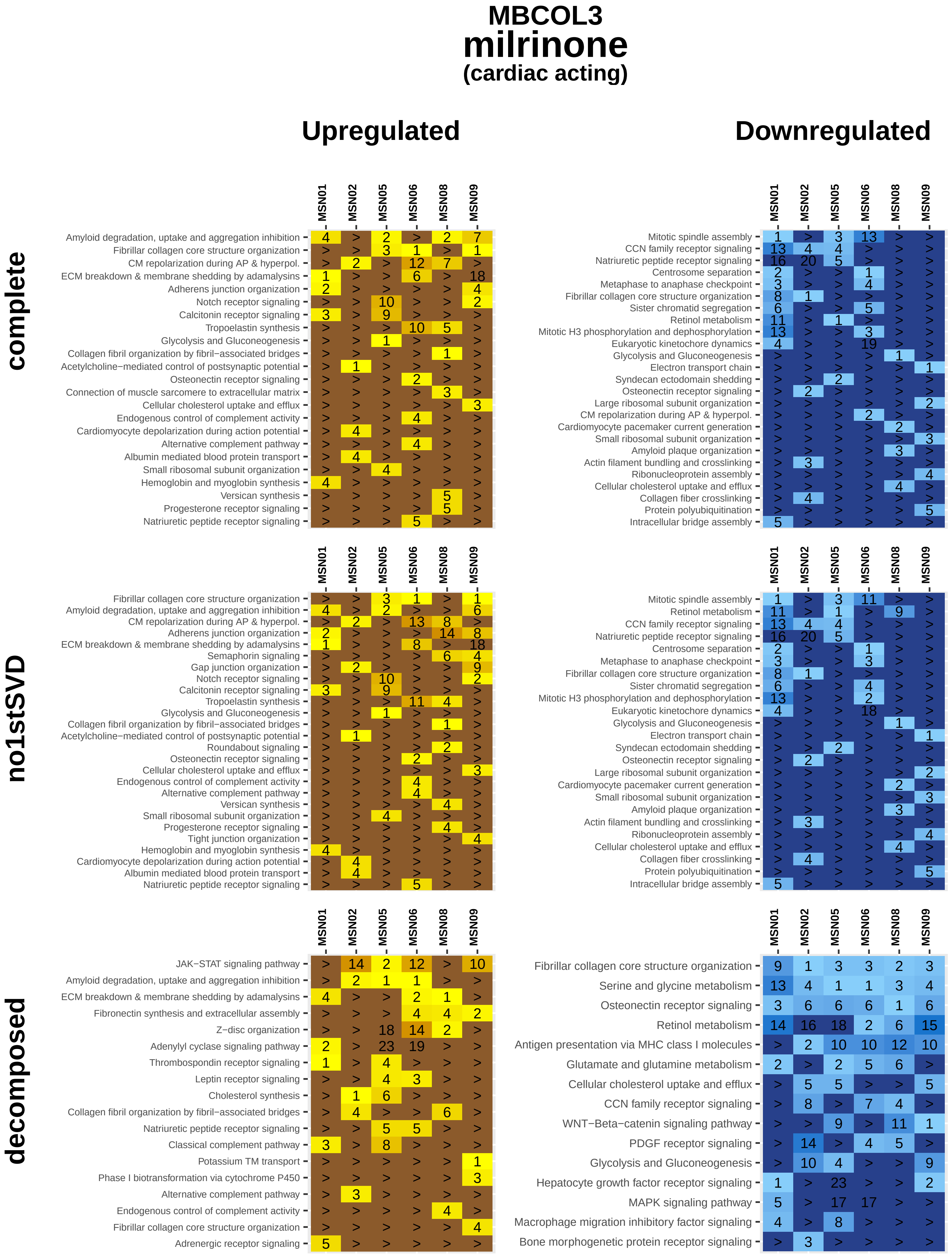
MBCOL3
milrinone
(cardiac acting)
Upregulated
Downregulated
complete
no1stSVD
decomposed
MSN01
MSN02
MSN05
MSN06
MSN08
MSN09
Amyloid degradation, uptake and aggregation inhibition
Fibrillar collagen core structure organization
CM repolarization during AP & hyperpol.
ECM breakdown & membrane shedding by adamalysins
Adherens junction organization
Notch receptor signaling
Calcitonin receptor signaling
Tropoelastin synthesis
Glycolysis and Gluconeogenesis
Collagen fibril organization by fibril−associated bridges
Acetylcholine−mediated control of postsynaptic potential
Osteonectin receptor signaling
Connection of muscle sarcomere to extracellular matrix
Cellular cholesterol uptake and efflux
Endogenous control of complement activity
Cardiomyocyte depolarization during action potential
Alternative complement pathway
Albumin mediated blood protein transport
Small ribosomal subunit organization
Hemoglobin and myoglobin synthesis
Versican synthesis
Progesterone receptor signaling
Natriuretic peptide receptor signaling
Mitotic spindle assembly
CCN family receptor signaling
Centrosome separation
Metaphase to anaphase checkpoint
Sister chromatid segregation
Retinol metabolism
Mitotic H3 phosphorylation and dephosphorylation
Eukaryotic kinetochore dynamics
Electron transport chain
Syndecan ectodomain shedding
Large ribosomal subunit organization
Cardiomyocyte pacemaker current generation
Amyloid plaque organization
Actin filament bundling and crosslinking
Ribonucleoprotein assembly
Collagen fiber crosslinking
Protein polyubiquitination
Intracellular bridge assembly
Semaphorin signaling
Gap junction organization
Roundabout signaling
Tight junction organization
JAK−STAT signaling pathway
Fibronectin synthesis and extracellular assembly
Z−disc organization
Adenylyl cyclase signaling pathway
Thrombospondin receptor signaling
Leptin receptor signaling
Cholesterol synthesis
Classical complement pathway
Potassium TM transport
Phase I biotransformation via cytochrome P450
Adrenergic receptor signaling
Serine and glycine metabolism
Antigen presentation via MHC class I molecules
Glutamate and glutamine metabolism
WNT−Beta−catenin signaling pathway
PDGF receptor signaling
Hepatocyte growth factor receptor signaling
MAPK signaling pathway
Macrophage migration inhibitory factor signaling
Bone morphogenetic protein receptor signaling

**MBCOL3**

### phenylephrine

**(cardiac acting)**

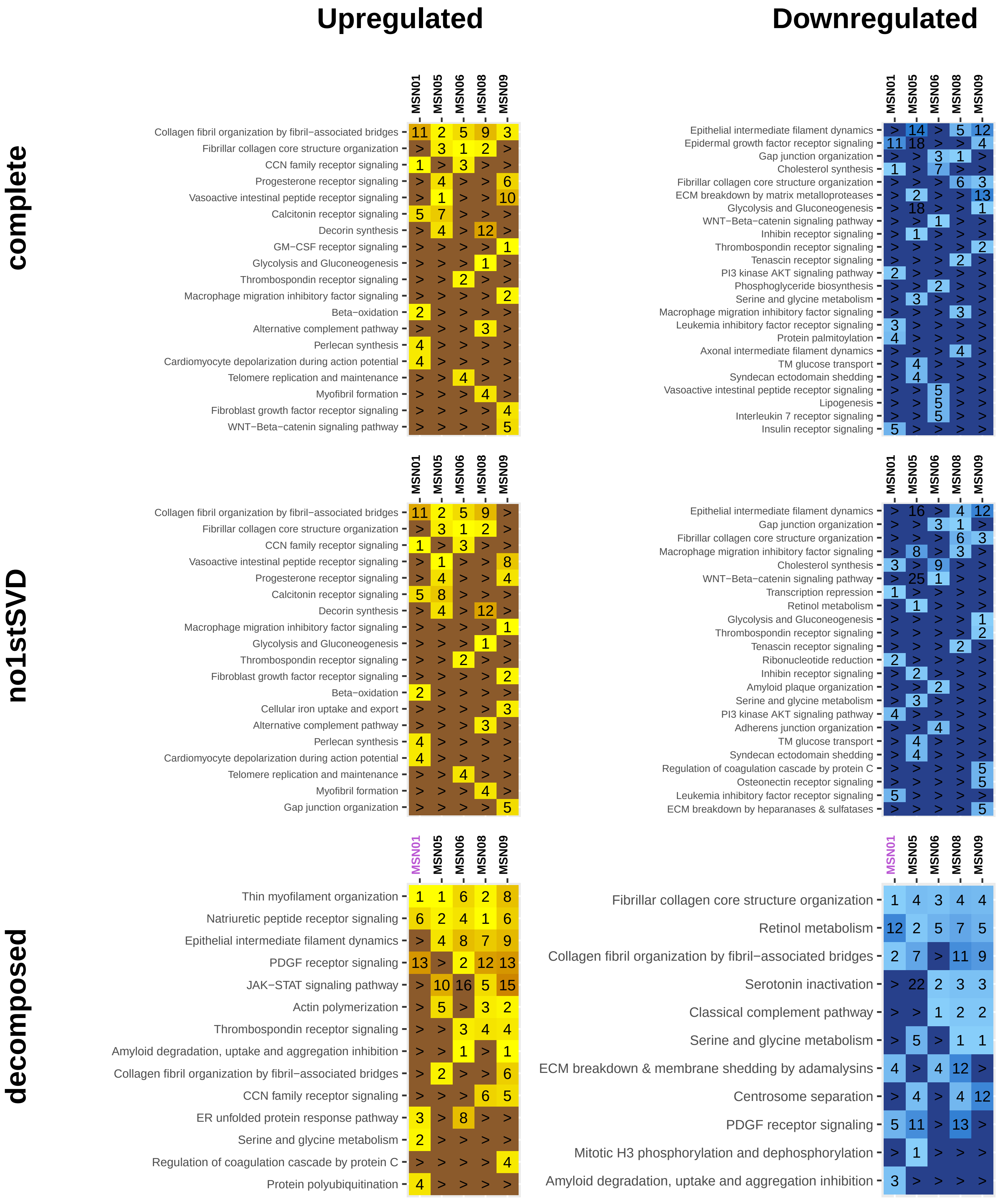

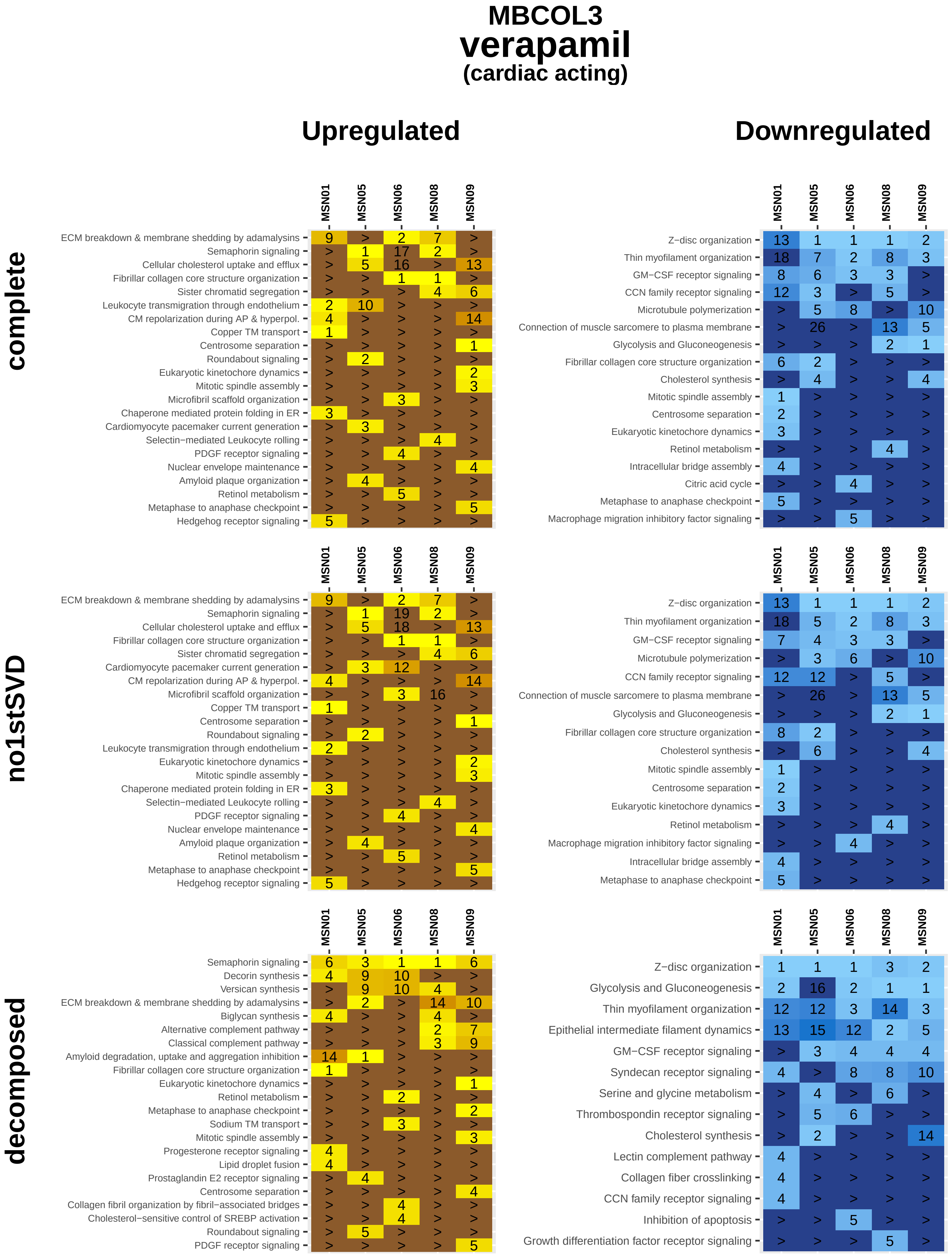

MBCOL3
verapamil
(cardiac acting)
Upregulated
Downregulated
complete
no1stSVD
decomposed
MSN01
MSN05
MSN06
MSN08
MSN09
ECM breakdown & membrane shedding by adamalysins
Semaphorin signaling
Cellular cholesterol uptake and efflux
Fibrillar collagen core structure organization
Sister chromatid segregation
Leukocyte transmigration through endothelium
CM repolarization during AP & hyperpol.
Copper TM transport
Centrosome separation
Roundabout signaling
Eukaryotic kinetochore dynamics
Mitotic spindle assembly
Microfibril scaffold organization
Chaperone mediated protein folding in ER
Cardiomyocyte pacemaker current generation
Selectin−mediated Leukocyte rolling
PDGF receptor signaling
Nuclear envelope maintenance
Amyloid plaque organization
Retinol metabolism
Metaphase to anaphase checkpoint
Hedgehog receptor signaling
Z−disc organization
Thin myofilament organization
GM−CSF receptor signaling
CCN family receptor signaling
Microtubule polymerization
Connection of muscle sarcomere to plasma membrane
Glycolysis and Gluconeogenesis
Cholesterol synthesis
Intracellular bridge assembly
Citric acid cycle
Macrophage migration inhibitory factor signaling
Decorin synthesis
Versican synthesis
Biglycan synthesis
Alternative complement pathway
Classical complement pathway
Amyloid degradation, uptake and aggregation inhibition
Sodium TM transport
Progesterone receptor signaling
Lipid droplet fusion
Prostaglandin E2 receptor signaling
Collagen fibril organization by fibril−associated bridges
Cholesterol−sensitive control of SREBP activation
Epithelial intermediate filament dynamics
Syndecan receptor signaling
Serine and glycine metabolism
Thrombospondin receptor signaling
Lectin complement pathway
Collagen fiber crosslinking
Inhibition of apoptosis
Growth differentiation factor receptor signaling

### MBCOL3
### azacitidine
#### (not cardiac act.)

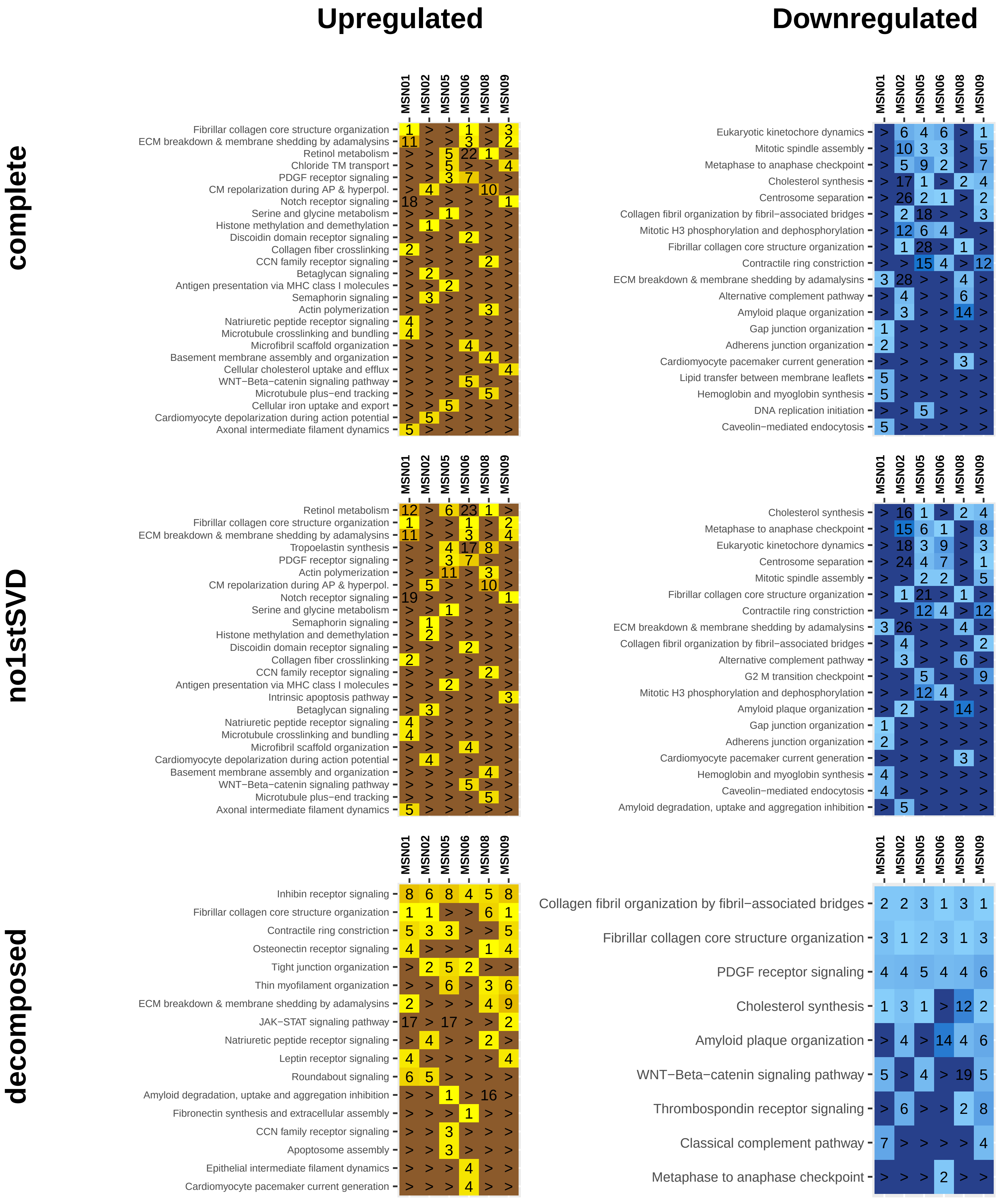

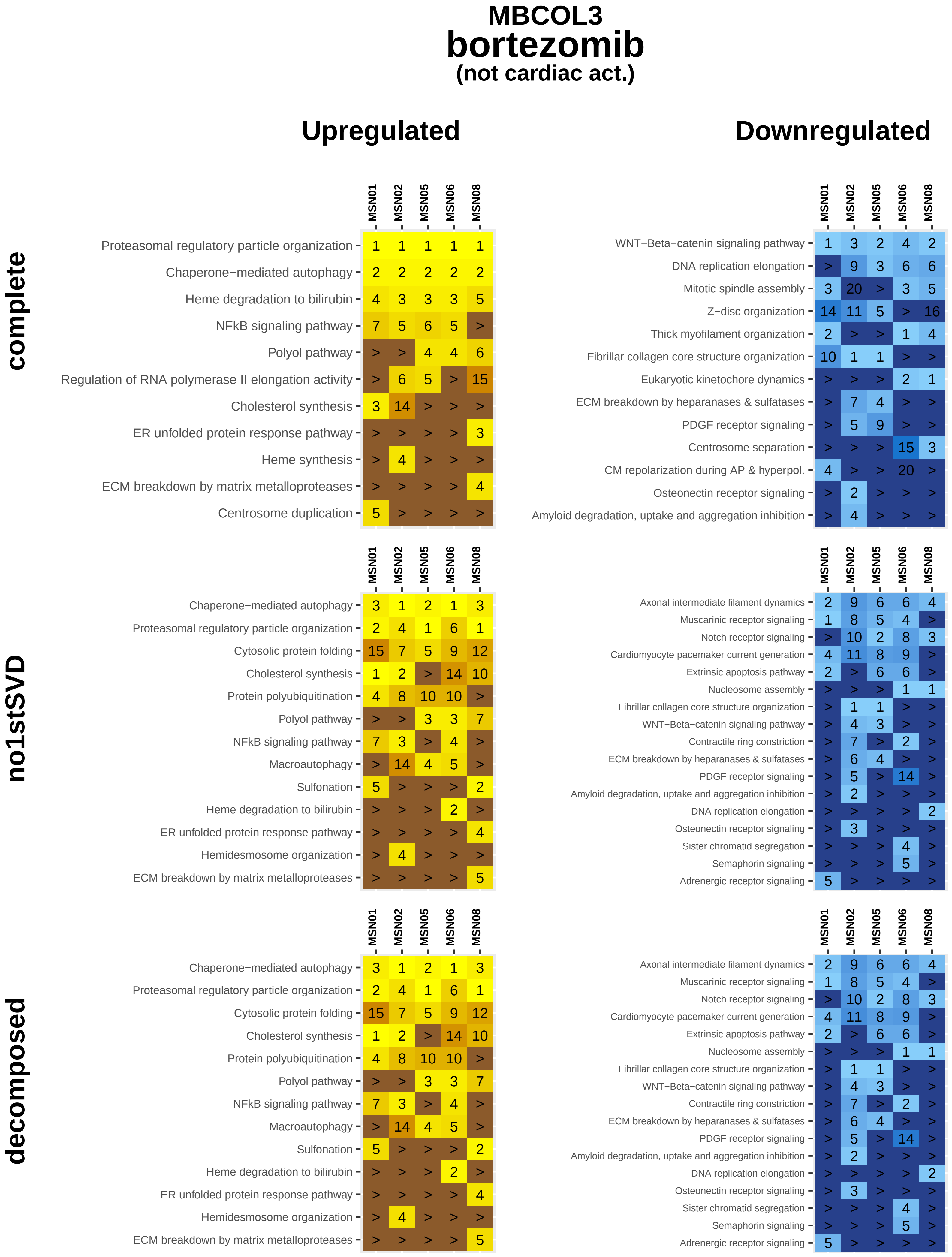
MBCOL3
bortezomib
(not cardiac act.)
Upregulated
Downregulated
complete
MSN01 MSN02 MSN05 MSN06 MSN08
Proteasomal regulatory particle organization 1 1 1 1 1
Chaperone−mediated autophagy 2 2 2 2 2
Heme degradation to bilirubin 4 3 3 3 5
NFkB signaling pathway 7 5 6 5 >
Polyol pathway > > 4 4 6
Regulation of RNA polymerase II elongation activity > 6 5 > 15
Cholesterol synthesis 3 14 > > >
ER unfolded protein response pathway > > > > 3
Heme synthesis > 4 > > >
ECM breakdown by matrix metalloproteases > > > > 4
Centrosome duplication 5 > > > >
MSN01 MSN02 MSN05 MSN06 MSN08
WNT−Beta−catenin signaling pathway 1 3 2 4 2
DNA replication elongation > 9 3 6 6
Mitotic spindle assembly 3 20 > 3 5
Z−disc organization 14 11 5 > 16
Thick myofilament organization 2 > > 1 4
Fibrillar collagen core structure organization 10 1 1 > >
Eukaryotic kinetochore dynamics > > > 2 1
ECM breakdown by heparanases & sulfatases > 7 4 > >
PDGF receptor signaling > 5 9 > >
Centrosome separation > > > 15 3
CM repolarization during AP & hyperpol. 4 > > 20 >
Osteonectin receptor signaling > 2 > > >
Amyloid degradation, uptake and aggregation inhibition > 4 > > >
no1stSVD
MSN01 MSN02 MSN05 MSN06 MSN08
Chaperone−mediated autophagy 3 1 2 1 3
Proteasomal regulatory particle organization 2 4 1 6 1
Cytosolic protein folding 15 7 5 9 12
Cholesterol synthesis 1 2 > 14 10
Protein polyubiquitination 4 8 10 10 >
Polyol pathway > > 3 3 7
NFkB signaling pathway 7 3 > 4 >
Macroautophagy > 14 4 5 >
Sulfonation 5 > > > 2
Heme degradation to bilirubin > > > 2 >
ER unfolded protein response pathway > > > > 4
Hemidesmosome organization > 4 > > >
ECM breakdown by matrix metalloproteases > > > > 5
MSN01 MSN02 MSN05 MSN06 MSN08
Axonal intermediate filament dynamics 2 9 6 6 4
Muscarinic receptor signaling 1 8 5 4 >
Notch receptor signaling > 10 2 8 3
Cardiomyocyte pacemaker current generation 4 11 8 9 >
Extrinsic apoptosis pathway 2 > 6 6 >
Nucleosome assembly > > > 1 1
Fibrillar collagen core structure organization > 1 1 > >
WNT−Beta−catenin signaling pathway > 4 3 > >
Contractile ring constriction > 7 > 2 >
ECM breakdown by heparanases & sulfatases > 6 4 > >
PDGF receptor signaling > 5 > 14 >
Amyloid degradation, uptake and aggregation inhibition > 2 > > >
DNA replication elongation > > > > 2
Osteonectin receptor signaling > 3 > > >
Sister chromatid segregation > > > 4 >
Semaphorin signaling > > > 5 >
Adrenergic receptor signaling 5 > > > >
decomposed
MSN01 MSN02 MSN05 MSN06 MSN08
Chaperone−mediated autophagy 3 1 2 1 3
Proteasomal regulatory particle organization 2 4 1 6 1
Cytosolic protein folding 15 7 5 9 12
Cholesterol synthesis 1 2 > 14 10
Protein polyubiquitination 4 8 10 10 >
Polyol pathway > > 3 3 7
NFkB signaling pathway 7 3 > 4 >
Macroautophagy > 14 4 5 >
Sulfonation 5 > > > 2
Heme degradation to bilirubin > > > 2 >
ER unfolded protein response pathway > > > > 4
Hemidesmosome organization > 4 > > >
ECM breakdown by matrix metalloproteases > > > > 5
MSN01 MSN02 MSN05 MSN06 MSN08
Axonal intermediate filament dynamics 2 9 6 6 4
Muscarinic receptor signaling 1 8 5 4 >
Notch receptor signaling > 10 2 8 3
Cardiomyocyte pacemaker current generation 4 11 8 9 >
Extrinsic apoptosis pathway 2 > 6 6 >
Nucleosome assembly > > > 1 1
Fibrillar collagen core structure organization > 1 1 > >
WNT−Beta−catenin signaling pathway > 4 3 > >
Contractile ring constriction > 7 > 2 >
ECM breakdown by heparanases & sulfatases > 6 4 > >
PDGF receptor signaling > 5 > 14 >
Amyloid degradation, uptake and aggregation inhibition > 2 > > >
DNA replication elongation > > > > 2
Osteonectin receptor signaling > 3 > > >
Sister chromatid segregation > > > 4 >
Semaphorin signaling > > > 5 >
Adrenergic receptor signaling 5 > > > >

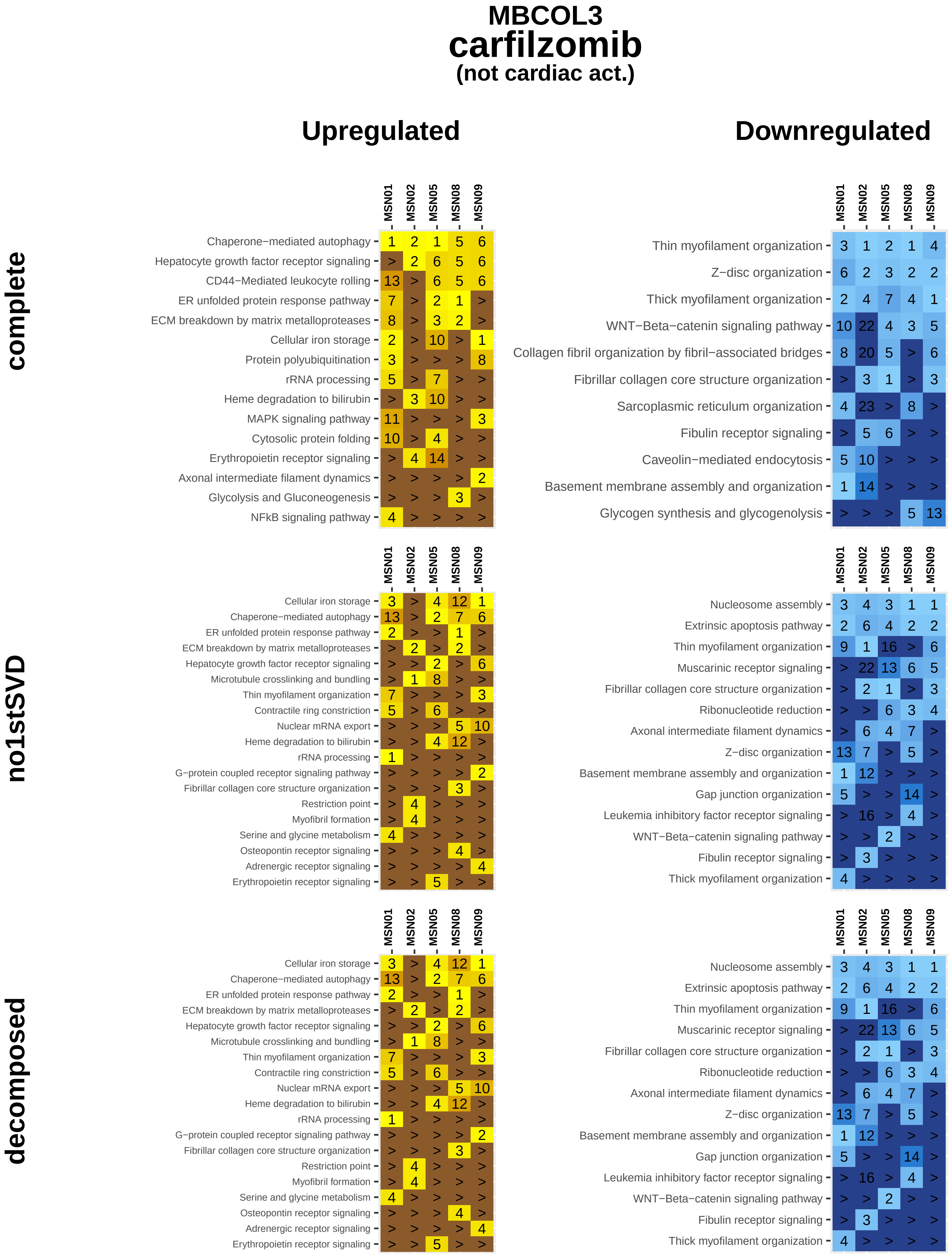

MBCOL3
carfilzomib
(not cardiac act.)
Upregulated
Downregulated
complete
MSN01 MSN02 MSN05 MSN08 MSN09
Chaperone−mediated autophagy 1 2 1 5 6
Hepatocyte growth factor receptor signaling > 2 6 5 6
CD44−Mediated leukocyte rolling 13 > 6 5 6
ER unfolded protein response pathway 7 > 2 1 >
ECM breakdown by matrix metalloproteases 8 > 3 2 >
Cellular iron storage 2 > 10 > 1
Protein polyubiquitination 3 > > > 8
rRNA processing 5 > 7 > >
Heme degradation to bilirubin > 3 10 > >
MAPK signaling pathway 11 > > > 3
Cytosolic protein folding 10 > 4 > >
Erythropoietin receptor signaling > 4 14 > >
Axonal intermediate filament dynamics > > > > 2
Glycolysis and Gluconeogenesis > > > 3 >
NFkB signaling pathway 4 > > > >
Thin myofilament organization 3 1 2 1 4
Z−disc organization 6 2 3 2 2
Thick myofilament organization 2 4 7 4 1
WNT−Beta−catenin signaling pathway 10 22 4 3 5
Collagen fibril organization by fibril−associated bridges 8 20 5 > 6
Fibrillar collagen core structure organization > 3 1 > 3
Sarcoplasmic reticulum organization 4 23 > 8 >
Fibulin receptor signaling > 5 6 > >
Caveolin−mediated endocytosis 5 10 > > >
Basement membrane assembly and organization 1 14 > > >
Glycogen synthesis and glycogenolysis > > > 5 13
no1stSVD
Cellular iron storage 3 > 4 12 1
Chaperone−mediated autophagy 13 > 2 7 6
ER unfolded protein response pathway 2 > > 1 >
ECM breakdown by matrix metalloproteases > 2 > 2 >
Hepatocyte growth factor receptor signaling > > 2 > 6
Microtubule crosslinking and bundling > 1 8 > >
Thin myofilament organization 7 > > > 3
Contractile ring constriction 5 > 6 > >
Nuclear mRNA export > > > 5 10
Heme degradation to bilirubin > > 4 12 >
rRNA processing 1 > > > >
G−protein coupled receptor signaling pathway > > > > 2
Fibrillar collagen core structure organization > > > 3 >
Restriction point > 4 > > >
Myofibril formation > 4 > > >
Serine and glycine metabolism 4 > > > >
Osteopontin receptor signaling > > > 4 >
Adrenergic receptor signaling > > > > 4
Erythropoietin receptor signaling > > 5 > >
Nucleosome assembly 3 4 3 1 1
Extrinsic apoptosis pathway 2 6 4 2 2
Thin myofilament organization 9 1 16 > 6
Muscarinic receptor signaling > 22 13 6 5
Fibrillar collagen core structure organization > 2 1 > 3
Ribonucleotide reduction > > 6 3 4
Axonal intermediate filament dynamics > 6 4 7 >
Z−disc organization 13 7 > 5 >
Basement membrane assembly and organization 1 12 > > >
Gap junction organization 5 > > 14 >
Leukemia inhibitory factor receptor signaling > 16 > 4 >
WNT−Beta−catenin signaling pathway > > 2 > >
Fibulin receptor signaling > 3 > > >
Thick myofilament organization 4 > > > >
decomposed
Cellular iron storage 3 > 4 12 1
Chaperone−mediated autophagy 13 > 2 7 6
ER unfolded protein response pathway 2 > > 1 >
ECM breakdown by matrix metalloproteases > 2 > 2 >
Hepatocyte growth factor receptor signaling > > 2 > 6
Microtubule crosslinking and bundling > 1 8 > >
Thin myofilament organization 7 > > > 3
Contractile ring constriction 5 > 6 > >
Nuclear mRNA export > > > 5 10
Heme degradation to bilirubin > > 4 12 >
rRNA processing 1 > > > >
G−protein coupled receptor signaling pathway > > > > 2
Fibrillar collagen core structure organization > > > 3 >
Restriction point > 4 > > >
Myofibril formation > 4 > > >
Serine and glycine metabolism 4 > > > >
Osteopontin receptor signaling > > > 4 >
Adrenergic receptor signaling > > > > 4
Erythropoietin receptor signaling > > 5 > >
Nucleosome assembly 3 4 3 1 1
Extrinsic apoptosis pathway 2 6 4 2 2
Thin myofilament organization 9 1 16 > 6
Muscarinic receptor signaling > 22 13 6 5
Fibrillar collagen core structure organization > 2 1 > 3
Ribonucleotide reduction > > 6 3 4
Axonal intermediate filament dynamics > 6 4 7 >
Z−disc organization 13 7 > 5 >
Basement membrane assembly and organization 1 12 > > >
Gap junction organization 5 > > 14 >
Leukemia inhibitory factor receptor signaling > 16 > 4 >
WNT−Beta−catenin signaling pathway > > 2 > >
Fibulin receptor signaling > 3 > > >
Thick myofilament organization 4 > > > >

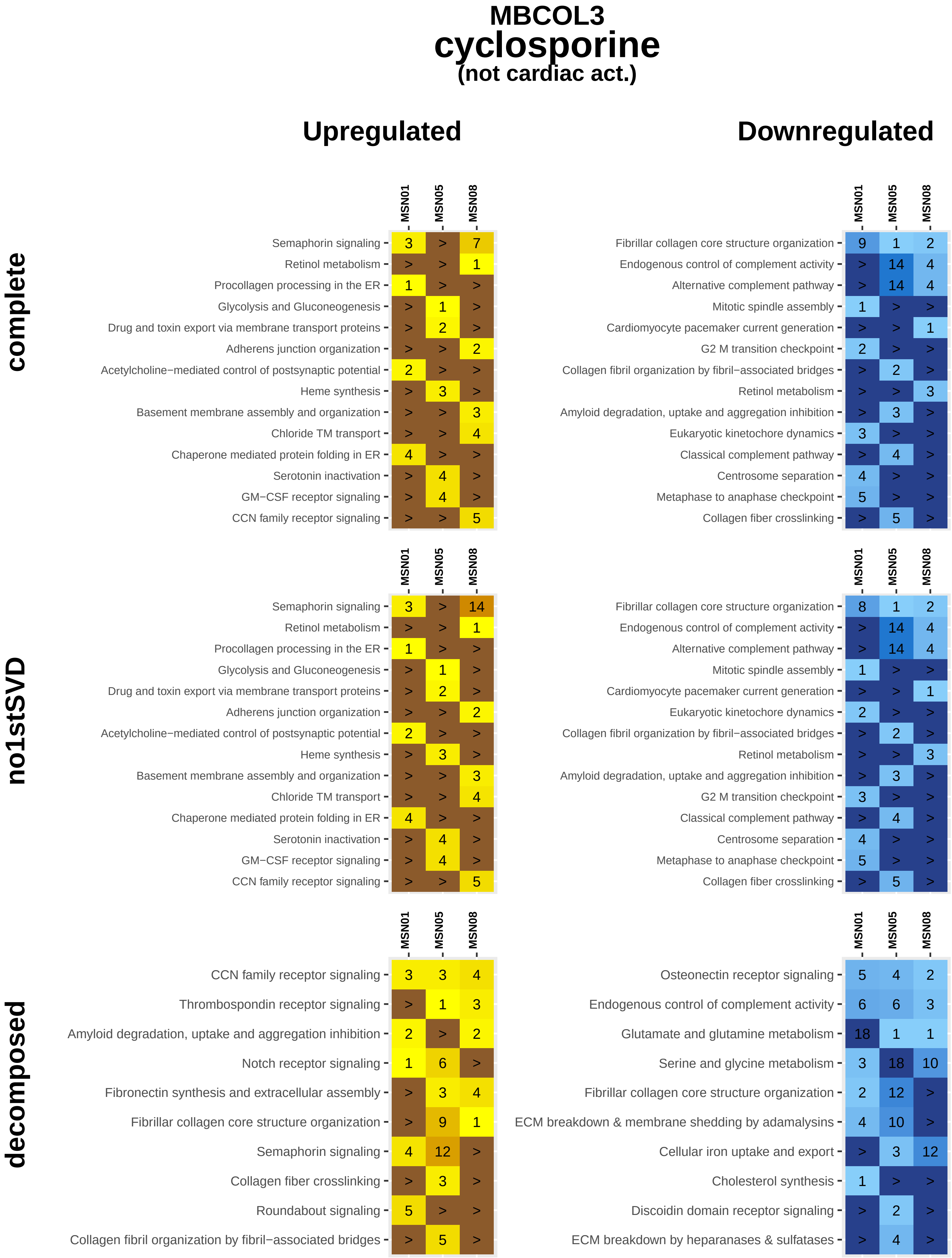

MBCOL3
cyclosporine
(not cardiac act.)
Upregulated
Downregulated
complete
MSN01 MSN05 MSN08
Semaphorin signaling 3 > 7
Retinol metabolism > > 1
Procollagen processing in the ER 1 > >
Glycolysis and Gluconeogenesis > 1 >
Drug and toxin export via membrane transport proteins > 2 >
Adherens junction organization > > 2
Acetylcholine−mediated control of postsynaptic potential 2 > >
Heme synthesis > 3 >
Basement membrane assembly and organization > > 3
Chloride TM transport > > 4
Chaperone mediated protein folding in ER 4 > >
Serotonin inactivation > 4 >
GM−CSF receptor signaling > 4 >
CCN family receptor signaling > > 5
MSN01 MSN05 MSN08
Fibrillar collagen core structure organization 9 1 2
Endogenous control of complement activity > 14 4
Alternative complement pathway > 14 4
Mitotic spindle assembly 1 > >
Cardiomyocyte pacemaker current generation > > 1
G2 M transition checkpoint 2 > >
Collagen fibril organization by fibril−associated bridges > 2 >
Retinol metabolism > > 3
Amyloid degradation, uptake and aggregation inhibition > 3 >
Eukaryotic kinetochore dynamics 3 > >
Classical complement pathway > 4 >
Centrosome separation 4 > >
Metaphase to anaphase checkpoint 5 > >
Collagen fiber crosslinking > 5 >
no1stSVD
MSN01 MSN05 MSN08
Semaphorin signaling 3 > 14
Retinol metabolism > > 1
Procollagen processing in the ER 1 > >
Glycolysis and Gluconeogenesis > 1 >
Drug and toxin export via membrane transport proteins > 2 >
Adherens junction organization > > 2
Acetylcholine−mediated control of postsynaptic potential 2 > >
Heme synthesis > 3 >
Basement membrane assembly and organization > > 3
Chloride TM transport > > 4
Chaperone mediated protein folding in ER 4 > >
Serotonin inactivation > 4 >
GM−CSF receptor signaling > 4 >
CCN family receptor signaling > > 5
MSN01 MSN05 MSN08
Fibrillar collagen core structure organization 8 1 2
Endogenous control of complement activity > 14 4
Alternative complement pathway > 14 4
Mitotic spindle assembly 1 > >
Cardiomyocyte pacemaker current generation > > 1
Eukaryotic kinetochore dynamics 2 > >
Collagen fibril organization by fibril−associated bridges > 2 >
Retinol metabolism > > 3
Amyloid degradation, uptake and aggregation inhibition > 3 >
G2 M transition checkpoint 3 > >
Classical complement pathway > 4 >
Centrosome separation 4 > >
Metaphase to anaphase checkpoint 5 > >
Collagen fiber crosslinking > 5 >
decomposed
MSN01 MSN05 MSN08
CCN family receptor signaling 3 3 4
Thrombospondin receptor signaling > 1 3
Amyloid degradation, uptake and aggregation inhibition 2 > 2
Notch receptor signaling 1 6 >
Fibronectin synthesis and extracellular assembly > 3 4
Fibrillar collagen core structure organization > 9 1
Semaphorin signaling 4 12 >
Collagen fiber crosslinking > 3 >
Roundabout signaling 5 > >
Collagen fibril organization by fibril−associated bridges > 5 >
MSN01 MSN05 MSN08
Osteonectin receptor signaling 5 4 2
Endogenous control of complement activity 6 6 3
Glutamate and glutamine metabolism 18 1 1
Serine and glycine metabolism 3 18 10
Fibrillar collagen core structure organization 2 12 >
ECM breakdown & membrane shedding by adamalysins 4 10 >
Cellular iron uptake and export > 3 12
Cholesterol synthesis 1 > >
Discoidin domain receptor signaling > 2 >
ECM breakdown by heparanases & sulfatases > 4 >

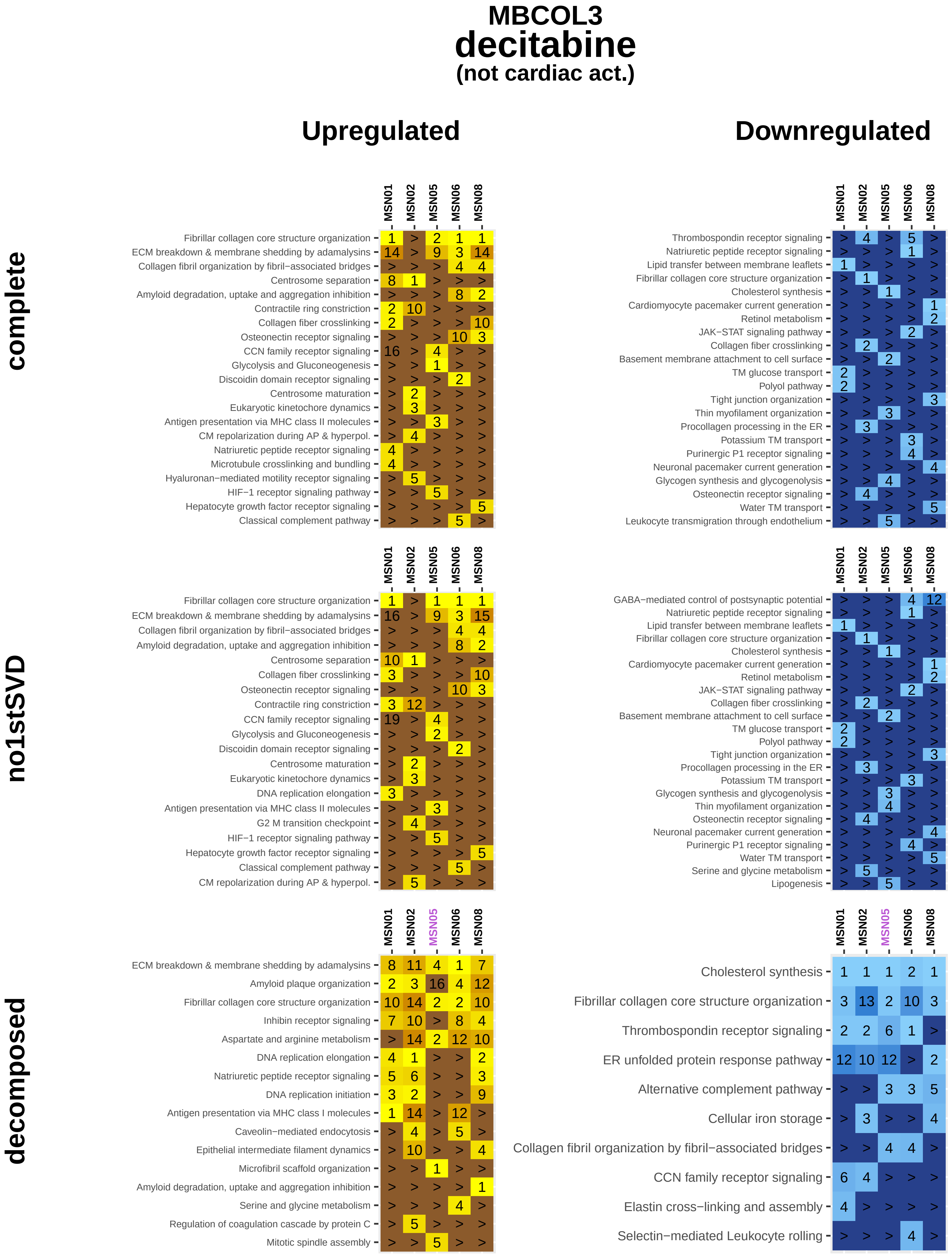

MBCOL3
decitabine
(not cardiac act.)
Upregulated
Downregulated
complete
no1stSVD
decomposed
MSN01
MSN02
MSN05
MSN06
MSN08
Fibrillar collagen core structure organization
ECM breakdown & membrane shedding by adamalysins
Collagen fibril organization by fibril−associated bridges
Centrosome separation
Amyloid degradation, uptake and aggregation inhibition
Contractile ring constriction
Collagen fiber crosslinking
Osteonectin receptor signaling
CCN family receptor signaling
Glycolysis and Gluconeogenesis
Discoidin domain receptor signaling
Centrosome maturation
Eukaryotic kinetochore dynamics
Antigen presentation via MHC class II molecules
CM repolarization during AP & hyperpol.
Natriuretic peptide receptor signaling
Microtubule crosslinking and bundling
Hyaluronan−mediated motility receptor signaling
HIF−1 receptor signaling pathway
Hepatocyte growth factor receptor signaling
Classical complement pathway
Thrombospondin receptor signaling
Lipid transfer between membrane leaflets
Cholesterol synthesis
Cardiomyocyte pacemaker current generation
Retinol metabolism
JAK−STAT signaling pathway
Basement membrane attachment to cell surface
TM glucose transport
Polyol pathway
Tight junction organization
Thin myofilament organization
Procollagen processing in the ER
Potassium TM transport
Purinergic P1 receptor signaling
Neuronal pacemaker current generation
Glycogen synthesis and glycogenolysis
Water TM transport
Leukocyte transmigration through endothelium
DNA replication elongation
G2 M transition checkpoint
GABA−mediated control of postsynaptic potential
Serine and glycine metabolism
Lipogenesis
Amyloid plaque organization
Inhibin receptor signaling
Aspartate and arginine metabolism
DNA replication initiation
Antigen presentation via MHC class I molecules
Caveolin−mediated endocytosis
Epithelial intermediate filament dynamics
Microfibril scaffold organization
Regulation of coagulation cascade by protein C
Mitotic spindle assembly
ER unfolded protein response pathway
Alternative complement pathway
Cellular iron storage
Elastin cross−linking and assembly
Selectin−mediated Leukocyte rolling

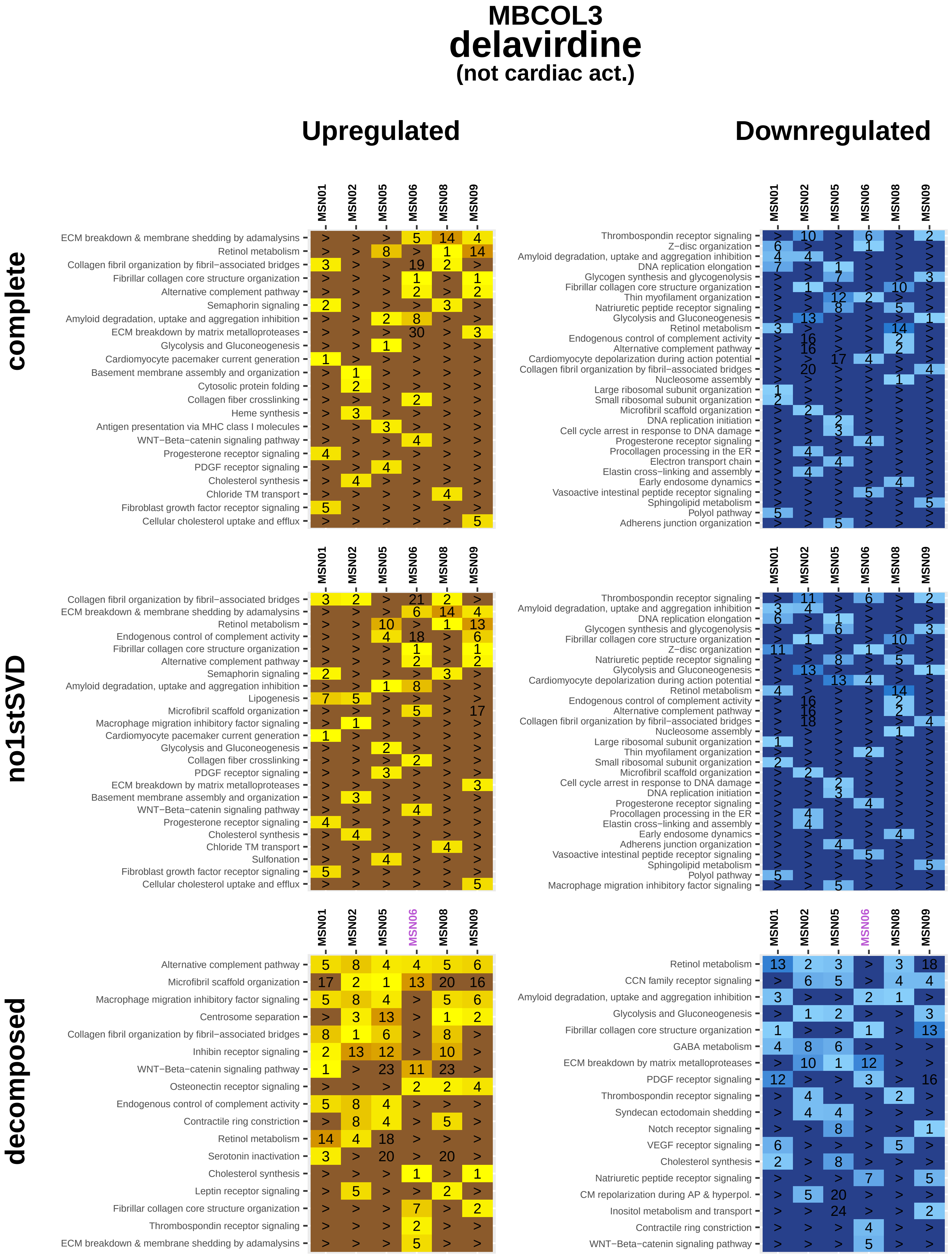
MBCOL3
delavirdine
(not cardiac act.)
Upregulated
Downregulated
complete
no1stSVD
decomposed
MSN01
MSN02
MSN05
MSN06
MSN08
MSN09
ECM breakdown & membrane shedding by adamalysins
Retinol metabolism
Collagen fibril organization by fibril−associated bridges
Fibrillar collagen core structure organization
Alternative complement pathway
Semaphorin signaling
Amyloid degradation, uptake and aggregation inhibition
ECM breakdown by matrix metalloproteases
Glycolysis and Gluconeogenesis
Cardiomyocyte pacemaker current generation
Basement membrane assembly and organization
Cytosolic protein folding
Collagen fiber crosslinking
Heme synthesis
Antigen presentation via MHC class I molecules
WNT−Beta−catenin signaling pathway
Progesterone receptor signaling
PDGF receptor signaling
Cholesterol synthesis
Chloride TM transport
Fibroblast growth factor receptor signaling
Cellular cholesterol uptake and efflux
Thrombospondin receptor signaling
Z−disc organization
DNA replication elongation
Glycogen synthesis and glycogenolysis
Thin myofilament organization
Natriuretic peptide receptor signaling
Endogenous control of complement activity
Cardiomyocyte depolarization during action potential
Nucleosome assembly
Large ribosomal subunit organization
Small ribosomal subunit organization
Microfibril scaffold organization
DNA replication initiation
Cell cycle arrest in response to DNA damage
Procollagen processing in the ER
Electron transport chain
Elastin cross−linking and assembly
Early endosome dynamics
Vasoactive intestinal peptide receptor signaling
Sphingolipid metabolism
Polyol pathway
Adherens junction organization
Lipogenesis
Macrophage migration inhibitory factor signaling
Sulfonation
Centrosome separation
Inhibin receptor signaling
Osteonectin receptor signaling
Contractile ring constriction
Serotonin inactivation
Leptin receptor signaling
CCN family receptor signaling
GABA metabolism
Syndecan ectodomain shedding
Notch receptor signaling
VEGF receptor signaling
CM repolarization during AP & hyperpol.
Inositol metabolism and transport

### MBCOL3
### diclofenac
#### (not cardiac act.)

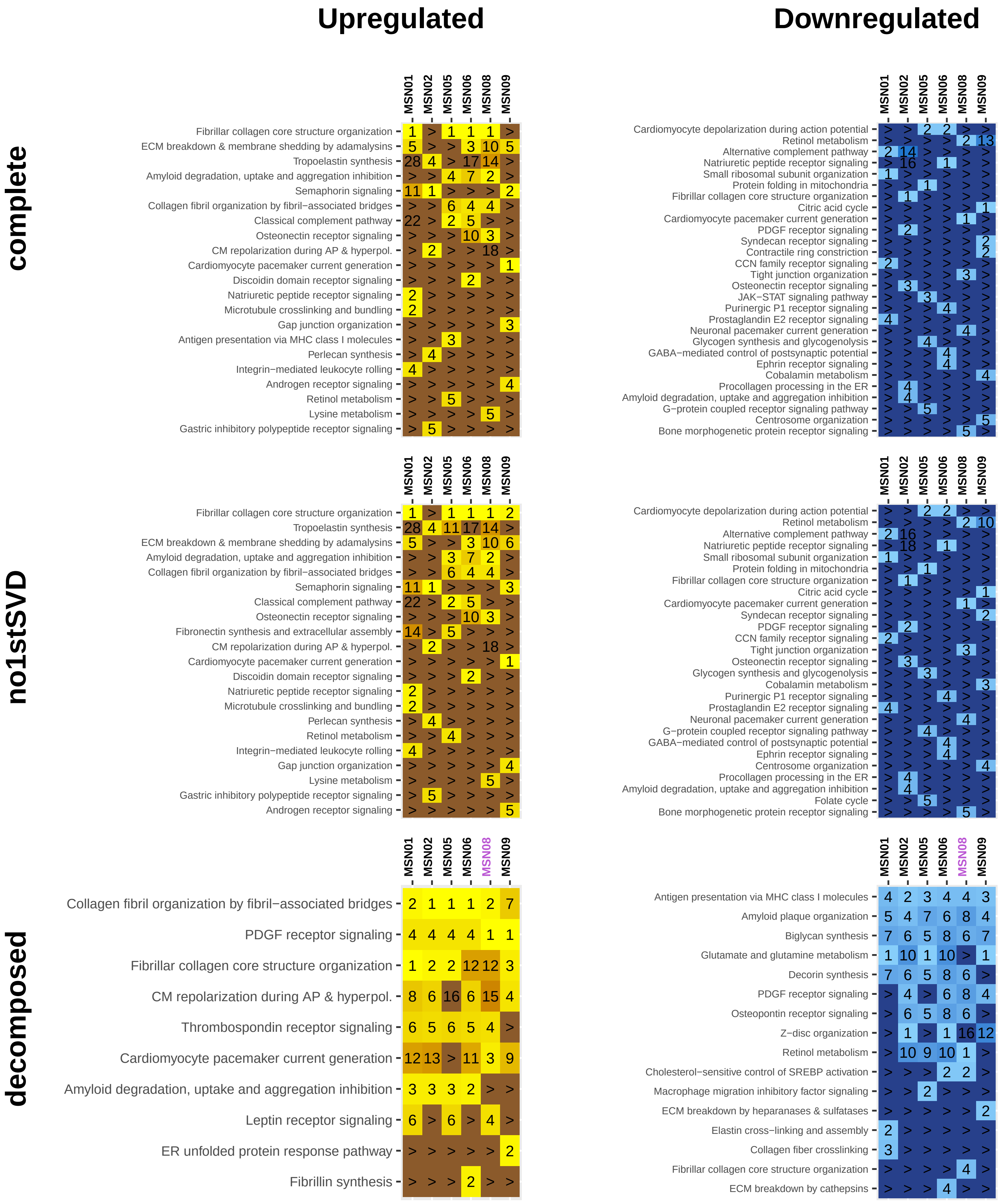

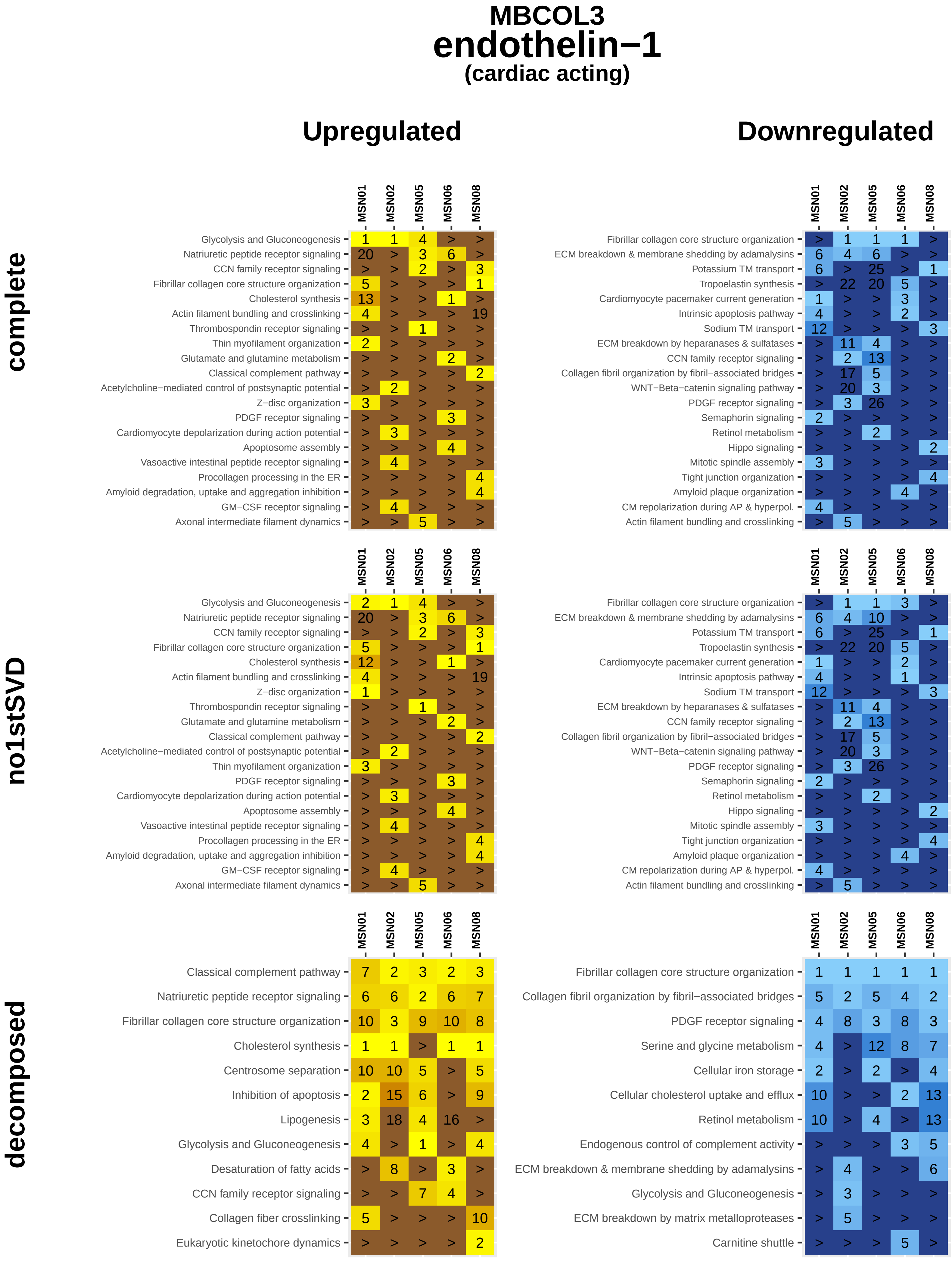

MBCOL3
endothelin−1
(cardiac acting)
Upregulated
Downregulated
complete
no1stSVD
decomposed
MSN01
MSN02
MSN05
MSN06
MSN08
Glycolysis and Gluconeogenesis
Natriuretic peptide receptor signaling
CCN family receptor signaling
Fibrillar collagen core structure organization
Cholesterol synthesis
Actin filament bundling and crosslinking
Thrombospondin receptor signaling
Thin myofilament organization
Glutamate and glutamine metabolism
Classical complement pathway
Acetylcholine−mediated control of postsynaptic potential
Z−disc organization
PDGF receptor signaling
Cardiomyocyte depolarization during action potential
Apoptosome assembly
Vasoactive intestinal peptide receptor signaling
Procollagen processing in the ER
Amyloid degradation, uptake and aggregation inhibition
GM−CSF receptor signaling
Axonal intermediate filament dynamics
ECM breakdown & membrane shedding by adamalysins
Potassium TM transport
Tropoelastin synthesis
Cardiomyocyte pacemaker current generation
Intrinsic apoptosis pathway
Sodium TM transport
ECM breakdown by heparanases & sulfatases
Collagen fibril organization by fibril−associated bridges
WNT−Beta−catenin signaling pathway
Semaphorin signaling
Retinol metabolism
Hippo signaling
Mitotic spindle assembly
Tight junction organization
Amyloid plaque organization
CM repolarization during AP & hyperpol.
Centrosome separation
Inhibition of apoptosis
Lipogenesis
Desaturation of fatty acids
Collagen fiber crosslinking
Eukaryotic kinetochore dynamics
Serine and glycine metabolism
Cellular iron storage
Cellular cholesterol uptake and efflux
Endogenous control of complement activity
ECM breakdown by matrix metalloproteases
Carnitine shuttle

### MBCOL3
### estradiol
#### (not cardiac act.)

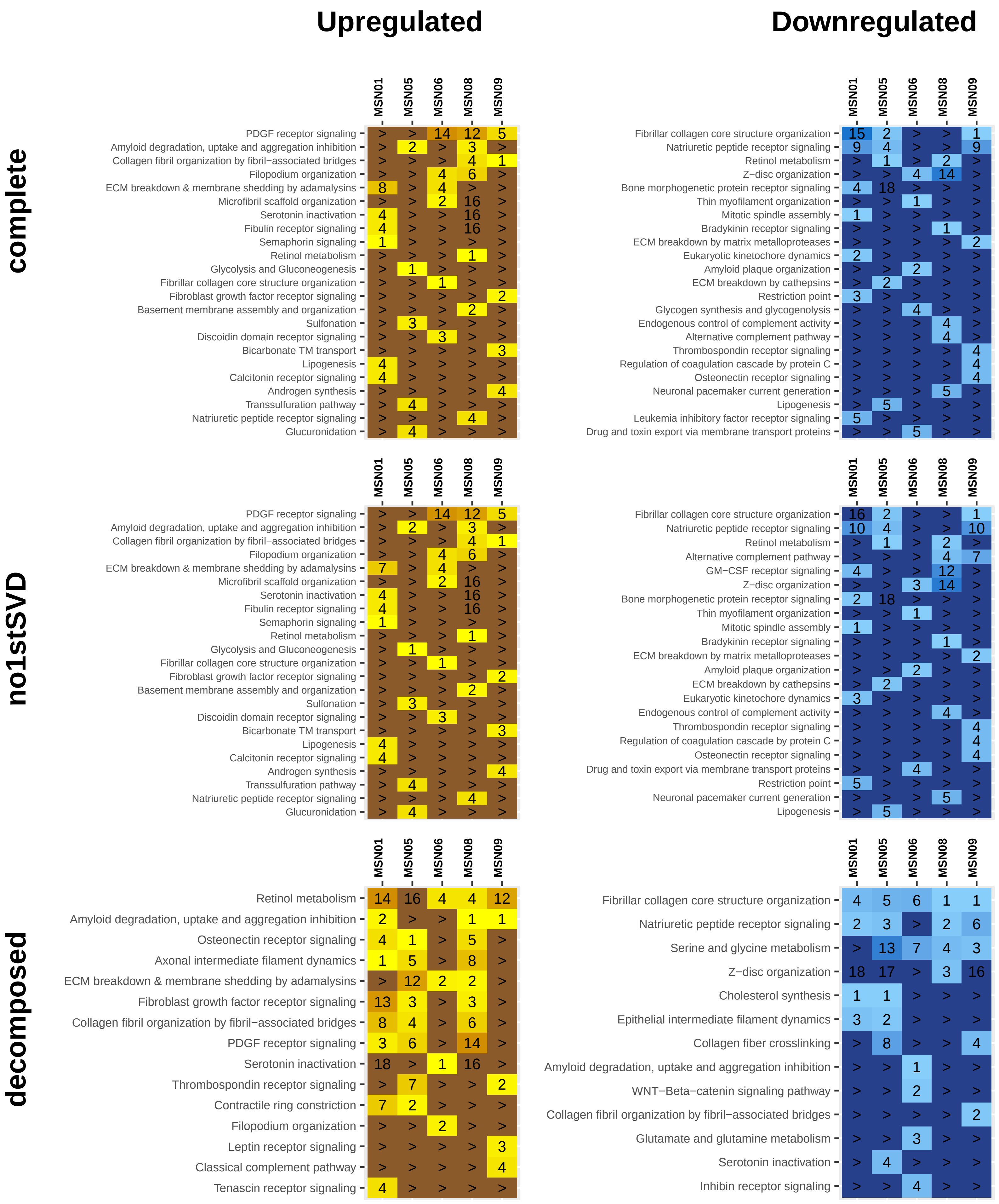

### MBCOL3
### insulin−like growth factor 1
#### (not cardiac act.)

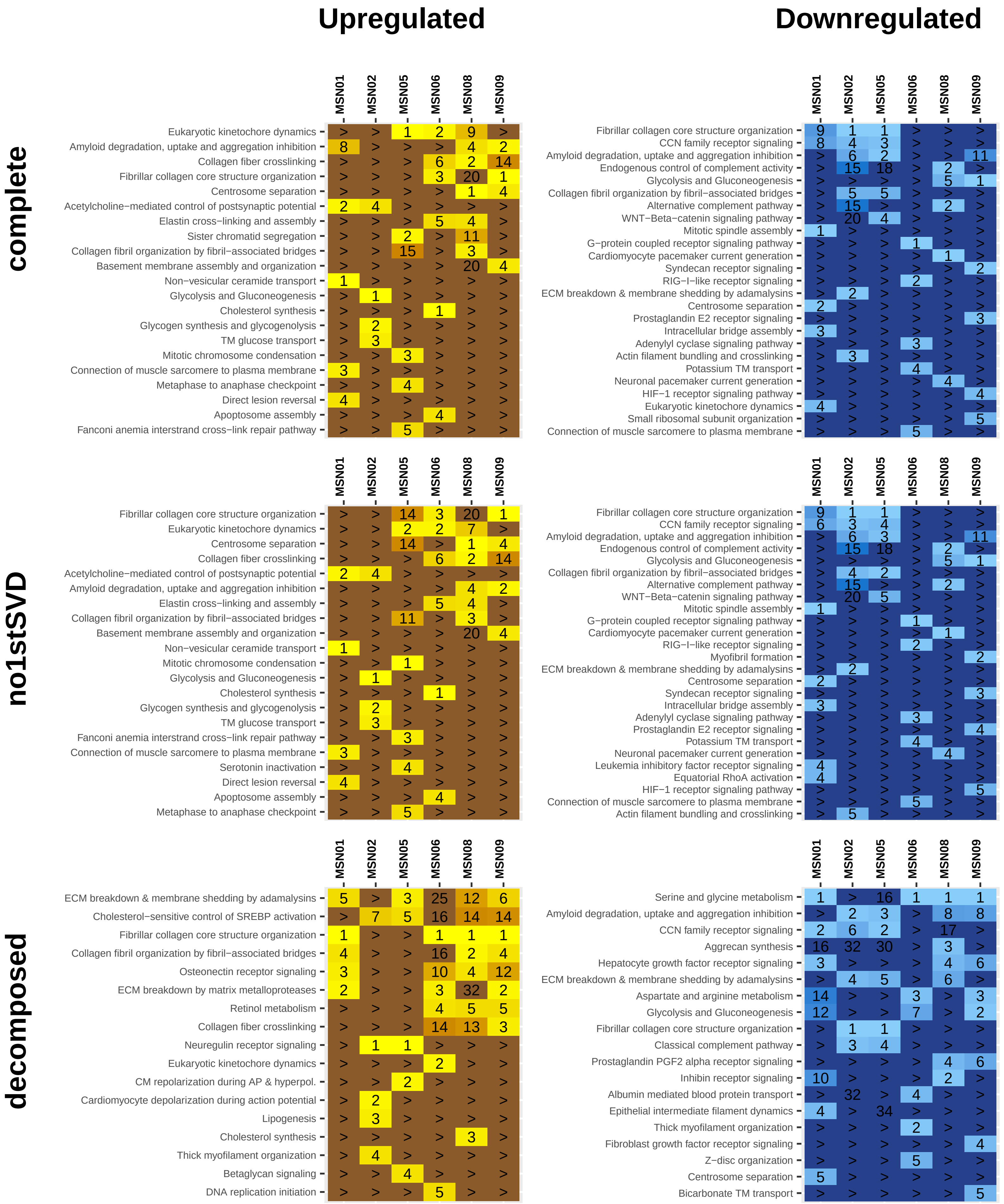

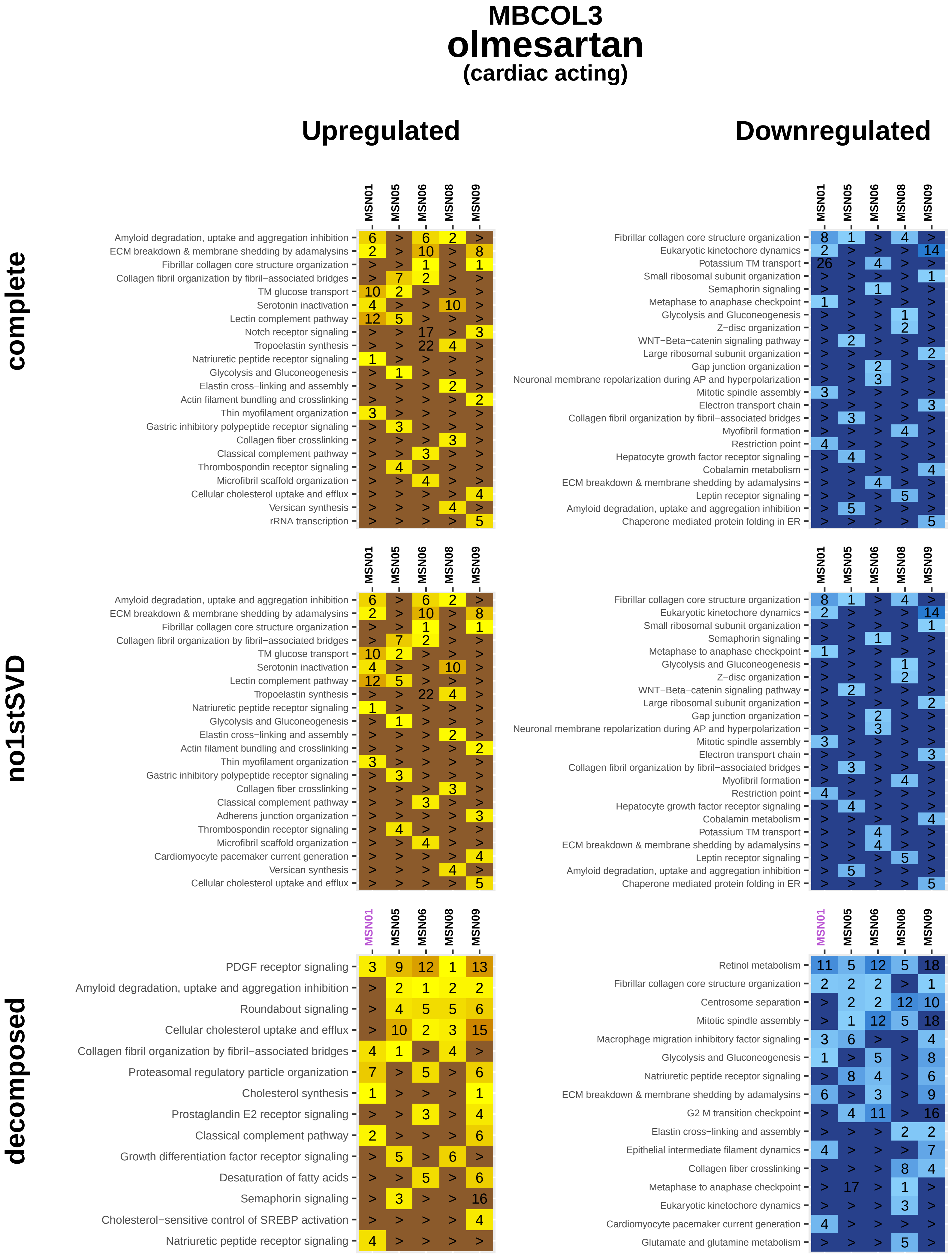

MBCOL3
olmesartan
(cardiac acting)
Upregulated
Downregulated
complete
no1stSVD
decomposed
MSN01
MSN05
MSN06
MSN08
MSN09
Amyloid degradation, uptake and aggregation inhibition
ECM breakdown & membrane shedding by adamalysins
Fibrillar collagen core structure organization
Collagen fibril organization by fibril−associated bridges
TM glucose transport
Serotonin inactivation
Lectin complement pathway
Notch receptor signaling
Tropoelastin synthesis
Natriuretic peptide receptor signaling
Glycolysis and Gluconeogenesis
Elastin cross−linking and assembly
Actin filament bundling and crosslinking
Thin myofilament organization
Gastric inhibitory polypeptide receptor signaling
Collagen fiber crosslinking
Classical complement pathway
Thrombospondin receptor signaling
Microfibril scaffold organization
Cellular cholesterol uptake and efflux
Versican synthesis
rRNA transcription
Eukaryotic kinetochore dynamics
Potassium TM transport
Small ribosomal subunit organization
Semaphorin signaling
Metaphase to anaphase checkpoint
Z−disc organization
WNT−Beta−catenin signaling pathway
Large ribosomal subunit organization
Gap junction organization
Neuronal membrane repolarization during AP and hyperpolarization
Mitotic spindle assembly
Electron transport chain
Myofibril formation
Restriction point
Hepatocyte growth factor receptor signaling
Cobalamin metabolism
Leptin receptor signaling
Chaperone mediated protein folding in ER
Adherens junction organization
Cardiomyocyte pacemaker current generation
PDGF receptor signaling
Roundabout signaling
Proteasomal regulatory particle organization
Cholesterol synthesis
Prostaglandin E2 receptor signaling
Growth differentiation factor receptor signaling
Desaturation of fatty acids
Cholesterol−sensitive control of SREBP activation
Retinol metabolism
Centrosome separation
Macrophage migration inhibitory factor signaling
G2 M transition checkpoint
Epithelial intermediate filament dynamics
Glutamate and glutamine metabolism

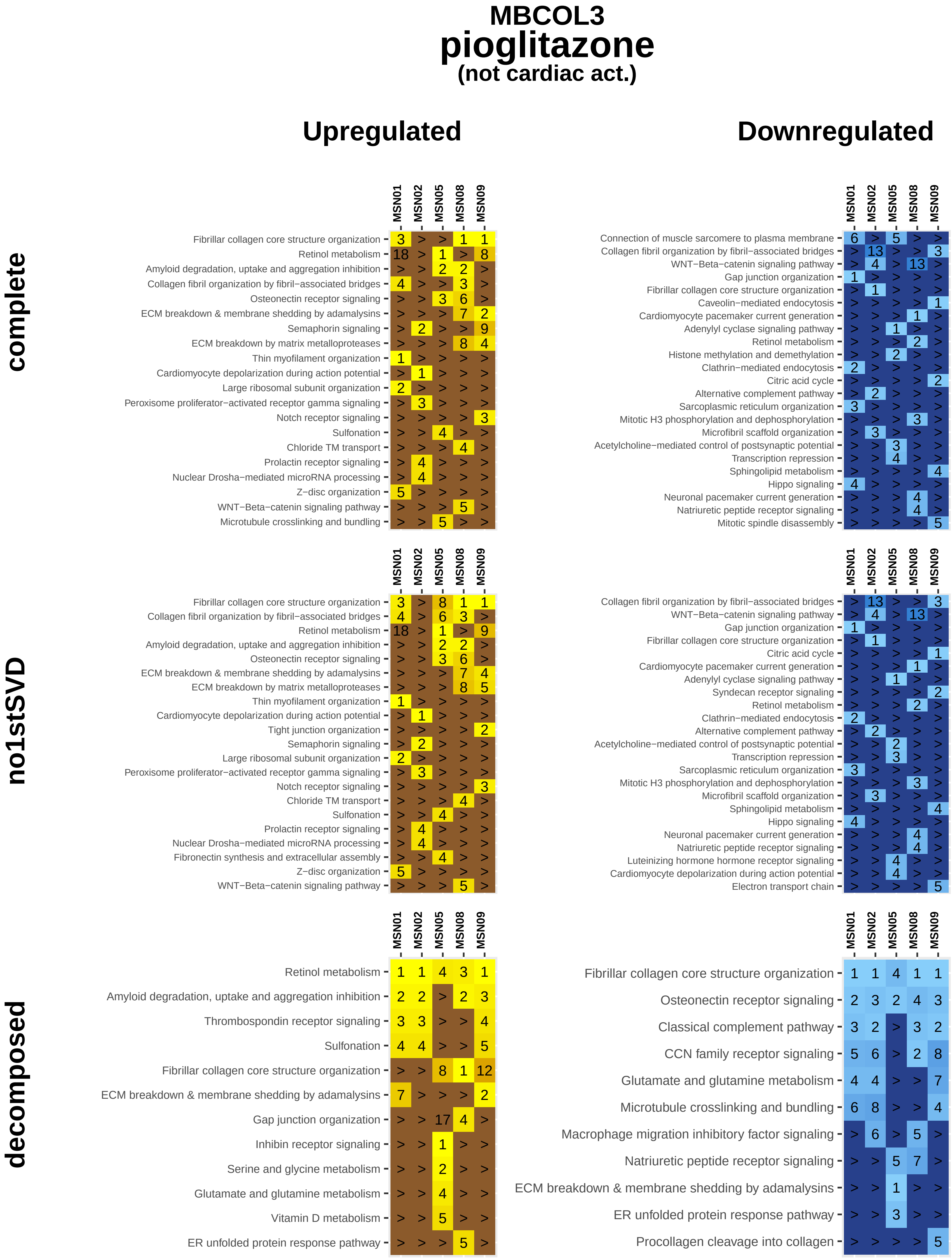
MBCOL3
pioglitazone
(not cardiac act.)
Upregulated
Downregulated
complete
no1stSVD
decomposed
MSN01
MSN02
MSN05
MSN08
MSN09
Fibrillar collagen core structure organization
Retinol metabolism
Amyloid degradation, uptake and aggregation inhibition
Collagen fibril organization by fibril−associated bridges
Osteonectin receptor signaling
ECM breakdown & membrane shedding by adamalysins
Semaphorin signaling
ECM breakdown by matrix metalloproteases
Thin myofilament organization
Cardiomyocyte depolarization during action potential
Large ribosomal subunit organization
Peroxisome proliferator−activated receptor gamma signaling
Notch receptor signaling
Sulfonation
Chloride TM transport
Prolactin receptor signaling
Nuclear Drosha−mediated microRNA processing
Z−disc organization
WNT−Beta−catenin signaling pathway
Microtubule crosslinking and bundling
Connection of muscle sarcomere to plasma membrane
Gap junction organization
Caveolin−mediated endocytosis
Cardiomyocyte pacemaker current generation
Adenylyl cyclase signaling pathway
Histone methylation and demethylation
Clathrin−mediated endocytosis
Citric acid cycle
Alternative complement pathway
Sarcoplasmic reticulum organization
Mitotic H3 phosphorylation and dephosphorylation
Microfibril scaffold organization
Acetylcholine−mediated control of postsynaptic potential
Transcription repression
Sphingolipid metabolism
Hippo signaling
Neuronal pacemaker current generation
Natriuretic peptide receptor signaling
Mitotic spindle disassembly
Tight junction organization
Fibronectin synthesis and extracellular assembly
Syndecan receptor signaling
Luteinizing hormone hormone receptor signaling
Electron transport chain
Thrombospondin receptor signaling
Inhibin receptor signaling
Serine and glycine metabolism
Glutamate and glutamine metabolism
Vitamin D metabolism
ER unfolded protein response pathway
Classical complement pathway
CCN family receptor signaling
Macrophage migration inhibitory factor signaling
Procollagen cleavage into collagen

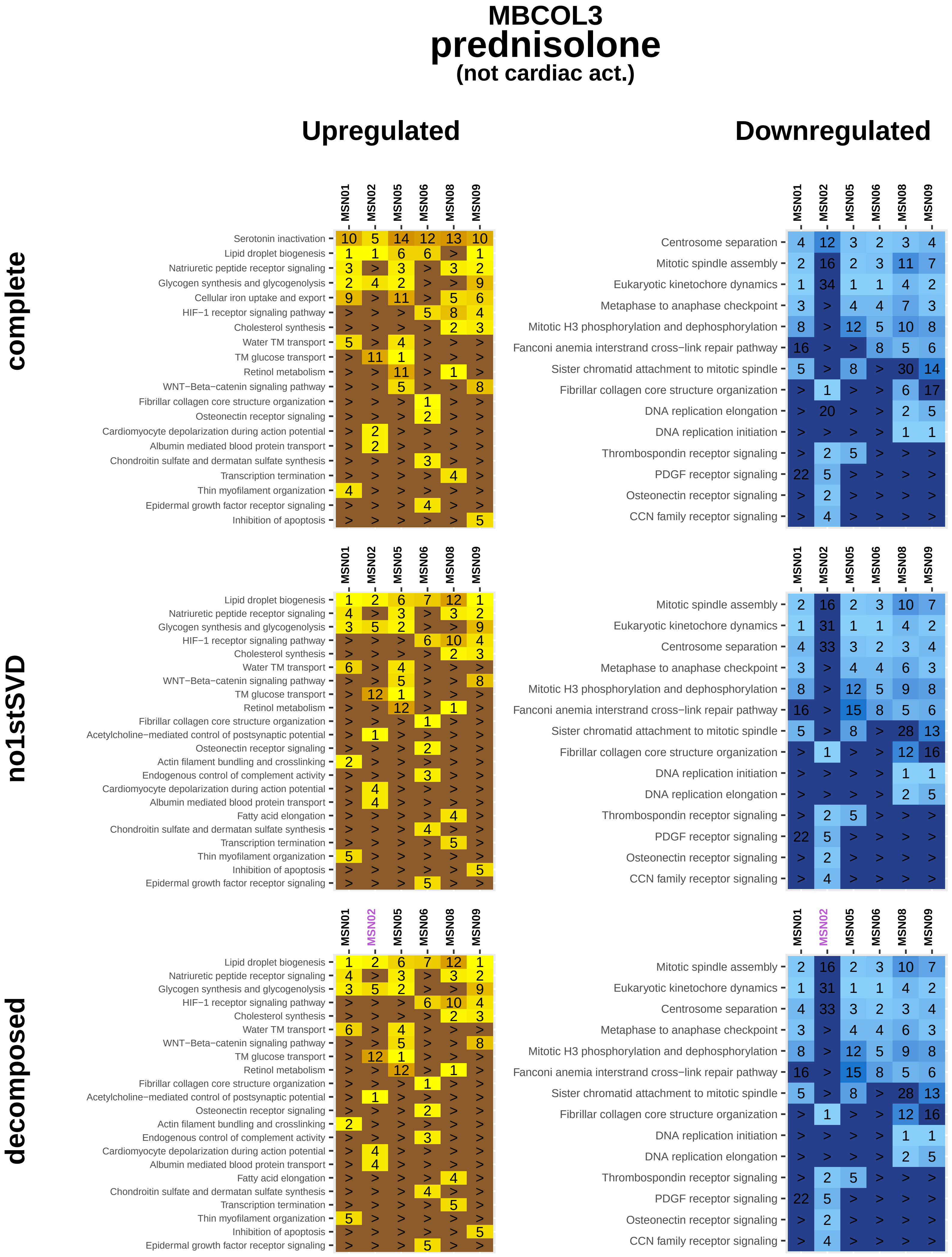
MBCOL3
prednisolone
(not cardiac act.)
Upregulated
Downregulated
complete
no1stSVD
decomposed
MSN01
MSN02
MSN05
MSN06
MSN08
MSN09
Serotonin inactivation
Lipid droplet biogenesis
Natriuretic peptide receptor signaling
Glycogen synthesis and glycogenolysis
Cellular iron uptake and export
HIF−1 receptor signaling pathway
Cholesterol synthesis
Water TM transport
TM glucose transport
Retinol metabolism
WNT−Beta−catenin signaling pathway
Fibrillar collagen core structure organization
Osteonectin receptor signaling
Cardiomyocyte depolarization during action potential
Albumin mediated blood protein transport
Chondroitin sulfate and dermatan sulfate synthesis
Transcription termination
Thin myofilament organization
Epidermal growth factor receptor signaling
Inhibition of apoptosis
Acetylcholine−mediated control of postsynaptic potential
Actin filament bundling and crosslinking
Endogenous control of complement activity
Fatty acid elongation
Centrosome separation
Mitotic spindle assembly
Eukaryotic kinetochore dynamics
Metaphase to anaphase checkpoint
Mitotic H3 phosphorylation and dephosphorylation
Fanconi anemia interstrand cross−link repair pathway
Sister chromatid attachment to mitotic spindle
DNA replication elongation
DNA replication initiation
Thrombospondin receptor signaling
PDGF receptor signaling
CCN family receptor signaling

### MBCOL3
### rosiglitazone
#### (not cardiac act.)

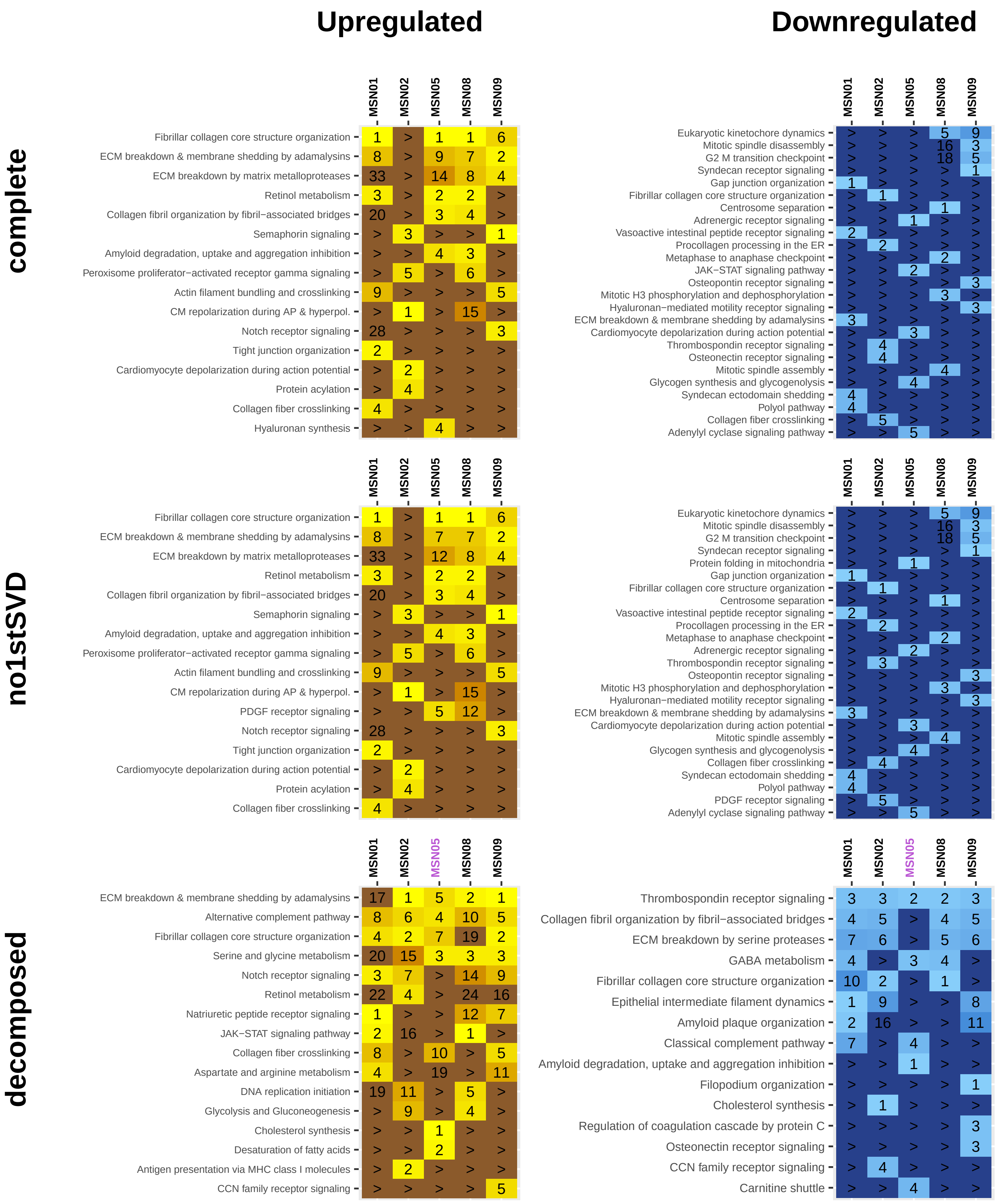

### MBCOL3
### saxagliptin
#### (not cardiac act.)

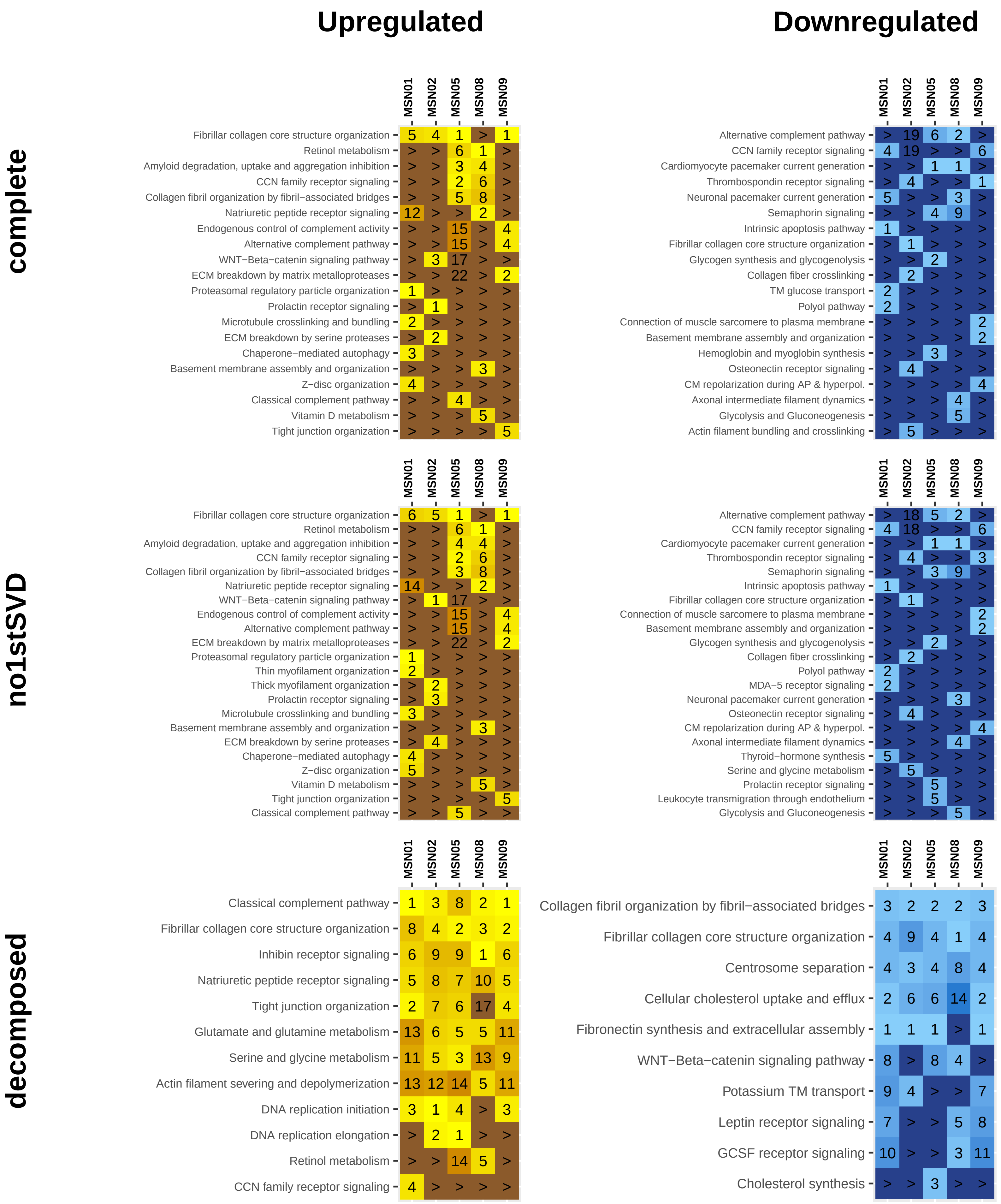

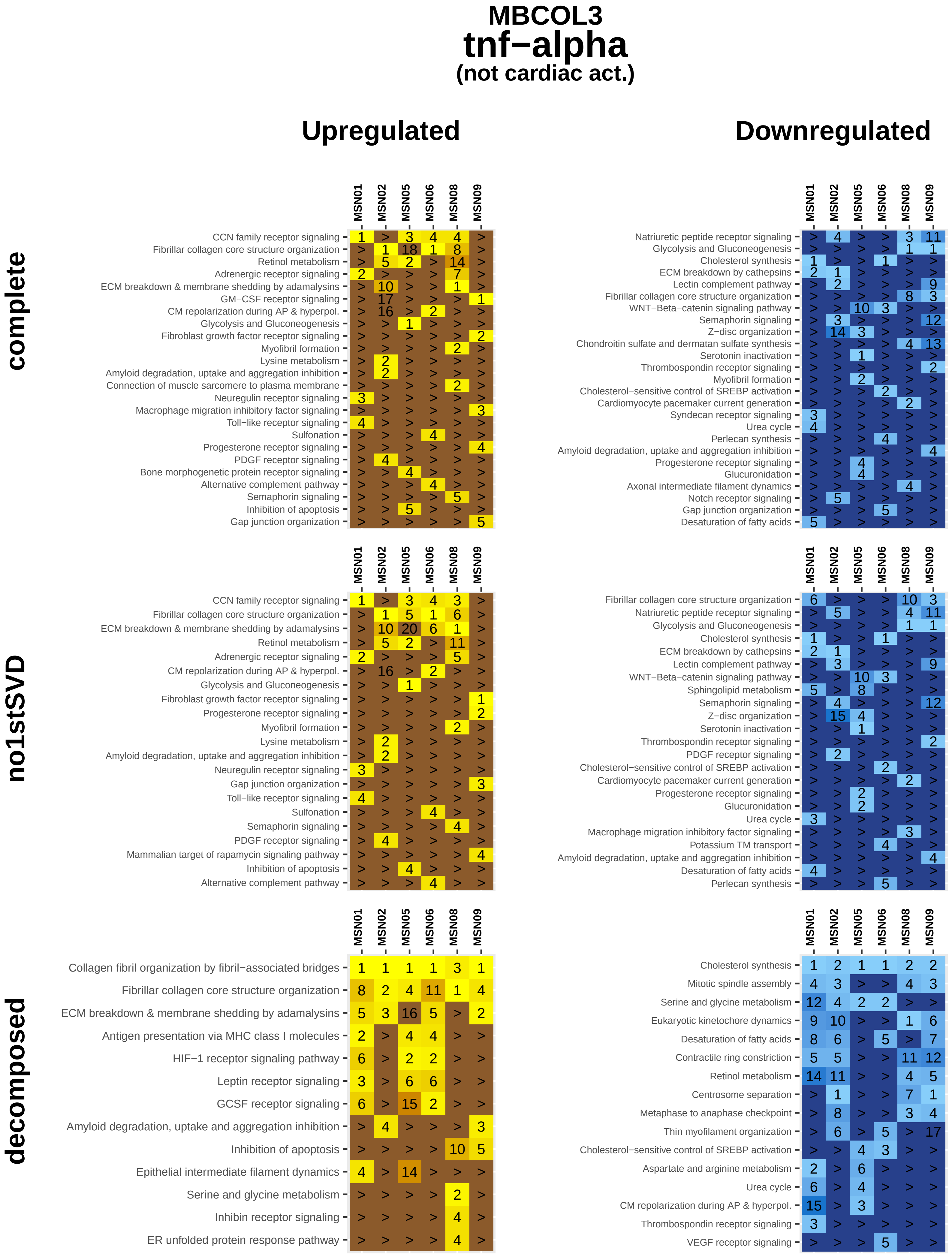

MBCOL3
tnf−alpha
(not cardiac act.)
Upregulated
Downregulated
complete
no1stSVD
decomposed
MSN01
MSN02
MSN05
MSN06
MSN08
MSN09
CCN family receptor signaling
Fibrillar collagen core structure organization
Retinol metabolism
Adrenergic receptor signaling
ECM breakdown & membrane shedding by adamalysins
GM−CSF receptor signaling
CM repolarization during AP & hyperpol.
Glycolysis and Gluconeogenesis
Fibroblast growth factor receptor signaling
Myofibril formation
Lysine metabolism
Amyloid degradation, uptake and aggregation inhibition
Connection of muscle sarcomere to plasma membrane
Neuregulin receptor signaling
Macrophage migration inhibitory factor signaling
Toll−like receptor signaling
Sulfonation
Progesterone receptor signaling
PDGF receptor signaling
Bone morphogenetic protein receptor signaling
Alternative complement pathway
Semaphorin signaling
Inhibition of apoptosis
Gap junction organization
Natriuretic peptide receptor signaling
Cholesterol synthesis
ECM breakdown by cathepsins
Lectin complement pathway
WNT−Beta−catenin signaling pathway
Z−disc organization
Chondroitin sulfate and dermatan sulfate synthesis
Serotonin inactivation
Thrombospondin receptor signaling
Cholesterol−sensitive control of SREBP activation
Cardiomyocyte pacemaker current generation
Syndecan receptor signaling
Urea cycle
Perlecan synthesis
Glucuronidation
Axonal intermediate filament dynamics
Notch receptor signaling
Desaturation of fatty acids
Sphingolipid metabolism
Potassium TM transport
Mammalian target of rapamycin signaling pathway
Collagen fibril organization by fibril−associated bridges
Antigen presentation via MHC class I molecules
HIF−1 receptor signaling pathway
Leptin receptor signaling
GCSF receptor signaling
Epithelial intermediate filament dynamics
Serine and glycine metabolism
Inhibin receptor signaling
ER unfolded protein response pathway
Mitotic spindle assembly
Eukaryotic kinetochore dynamics
Contractile ring constriction
Centrosome separation
Metaphase to anaphase checkpoint
Thin myofilament organization
Aspartate and arginine metabolism
VEGF receptor signaling
