## Supplemental Figure 14E for "Multiscale mapping of transcriptomic signatures for cardiotoxic drugs"

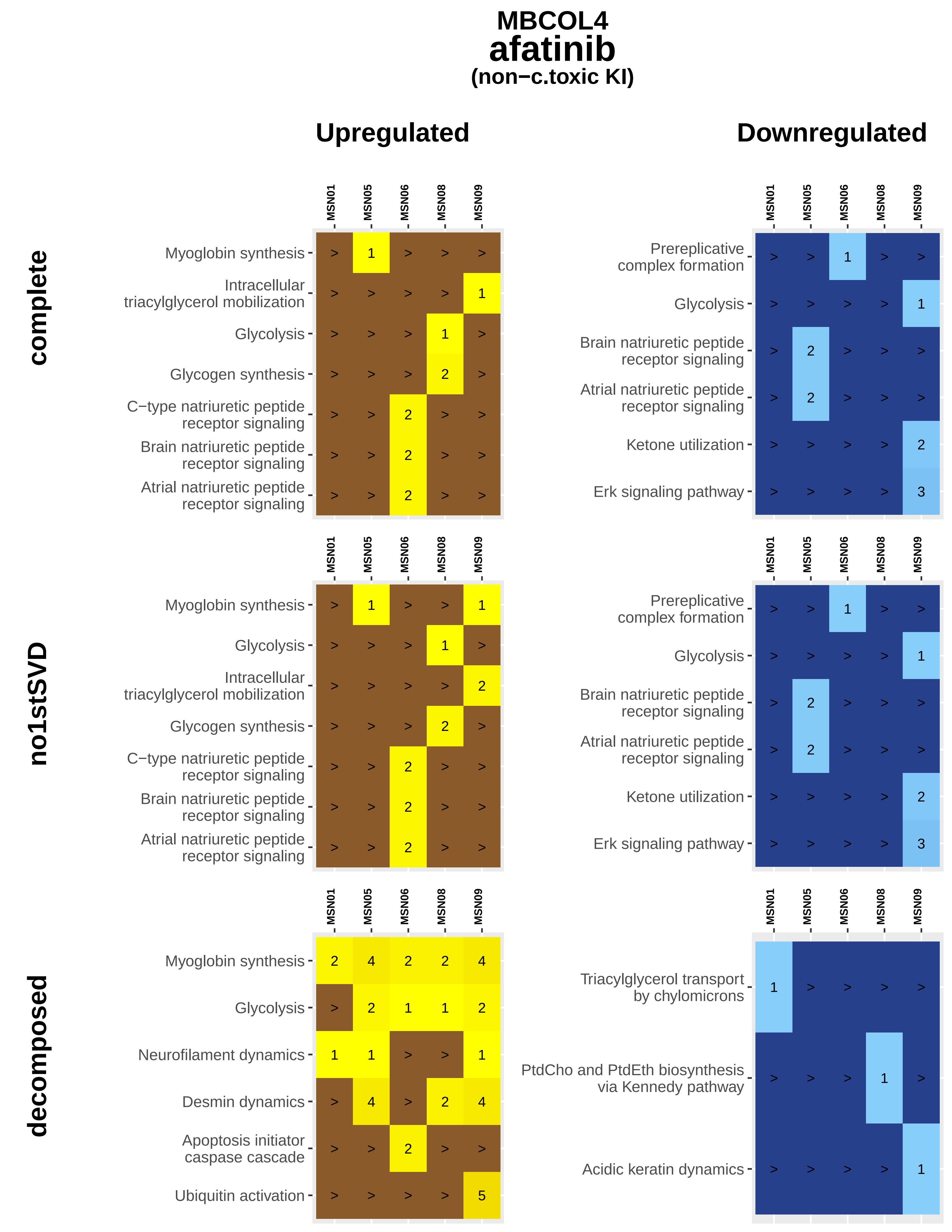
MBCOL4
afatinib
(non−c.toxic KI)
Upregulated
Downregulated
complete
no1stSVD
decomposed
MSN01
MSN05
MSN06
MSN08
MSN09
Myoglobin synthesis
Intracellular triacylglycerol mobilization
Glycolysis
Glycogen synthesis
C−type natriuretic peptide receptor signaling
Brain natriuretic peptide receptor signaling
Atrial natriuretic peptide receptor signaling
Prereplicative complex formation
Ketone utilization
Erk signaling pathway
Neurofilament dynamics
Desmin dynamics
Apoptosis initiator caspase cascade
Ubiquitin activation
Triacylglycerol transport by chylomicrons
PtdCho and PtdEth biosynthesis via Kennedy pathway
Acidic keratin dynamics

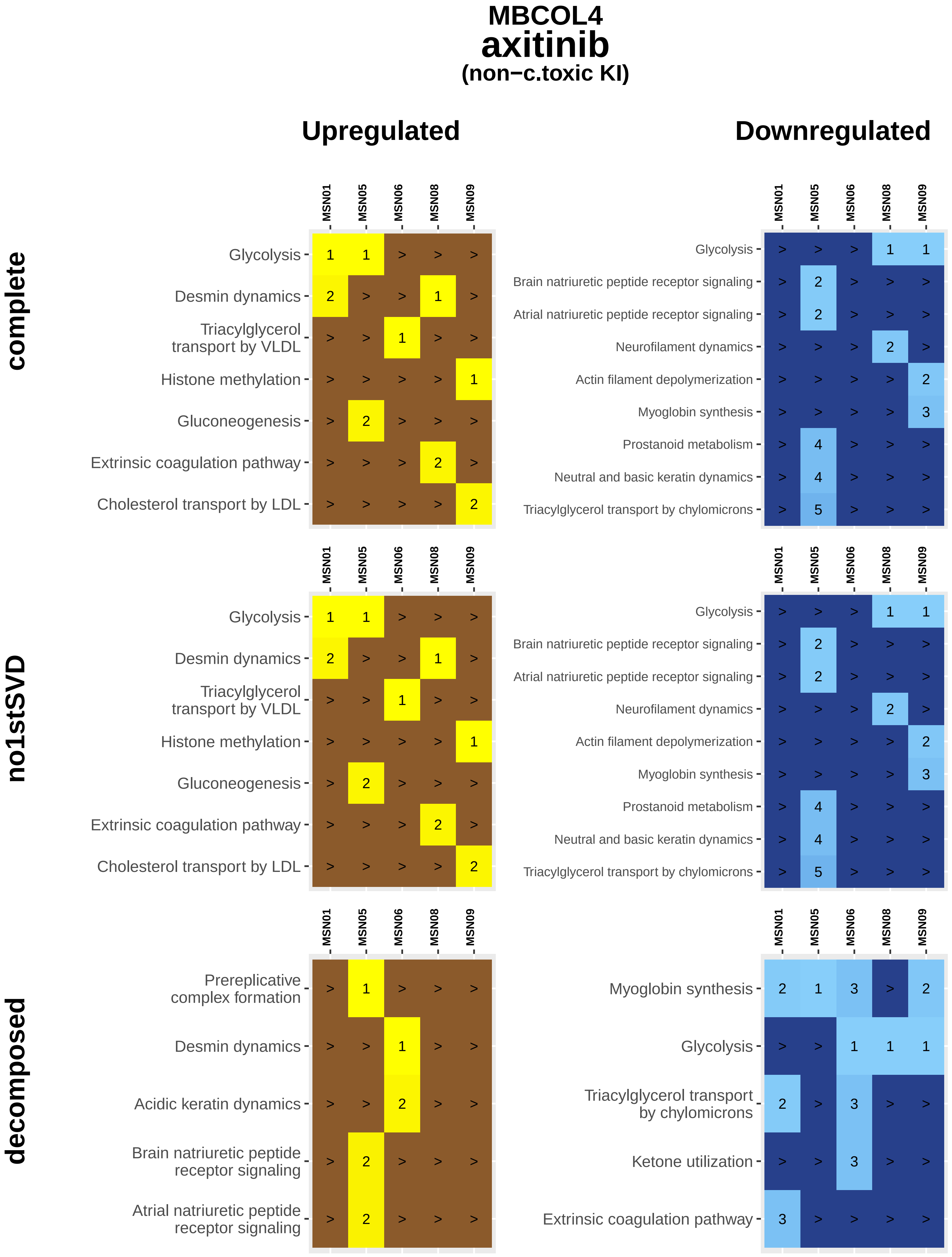
MBCOL4
axitinib
(non−c.toxic KI)
Upregulated
Downregulated
complete
no1stSVD
decomposed
MSN01
MSN05
MSN06
MSN08
MSN09
Glycolysis
Desmin dynamics
Triacylglycerol transport by VLDL
Histone methylation
Gluconeogenesis
Extrinsic coagulation pathway
Cholesterol transport by LDL
Brain natriuretic peptide receptor signaling
Atrial natriuretic peptide receptor signaling
Neurofilament dynamics
Actin filament depolymerization
Myoglobin synthesis
Prostanoid metabolism
Neutral and basic keratin dynamics
Triacylglycerol transport by chylomicrons
Prereplicative complex formation
Acidic keratin dynamics
Ketone utilization

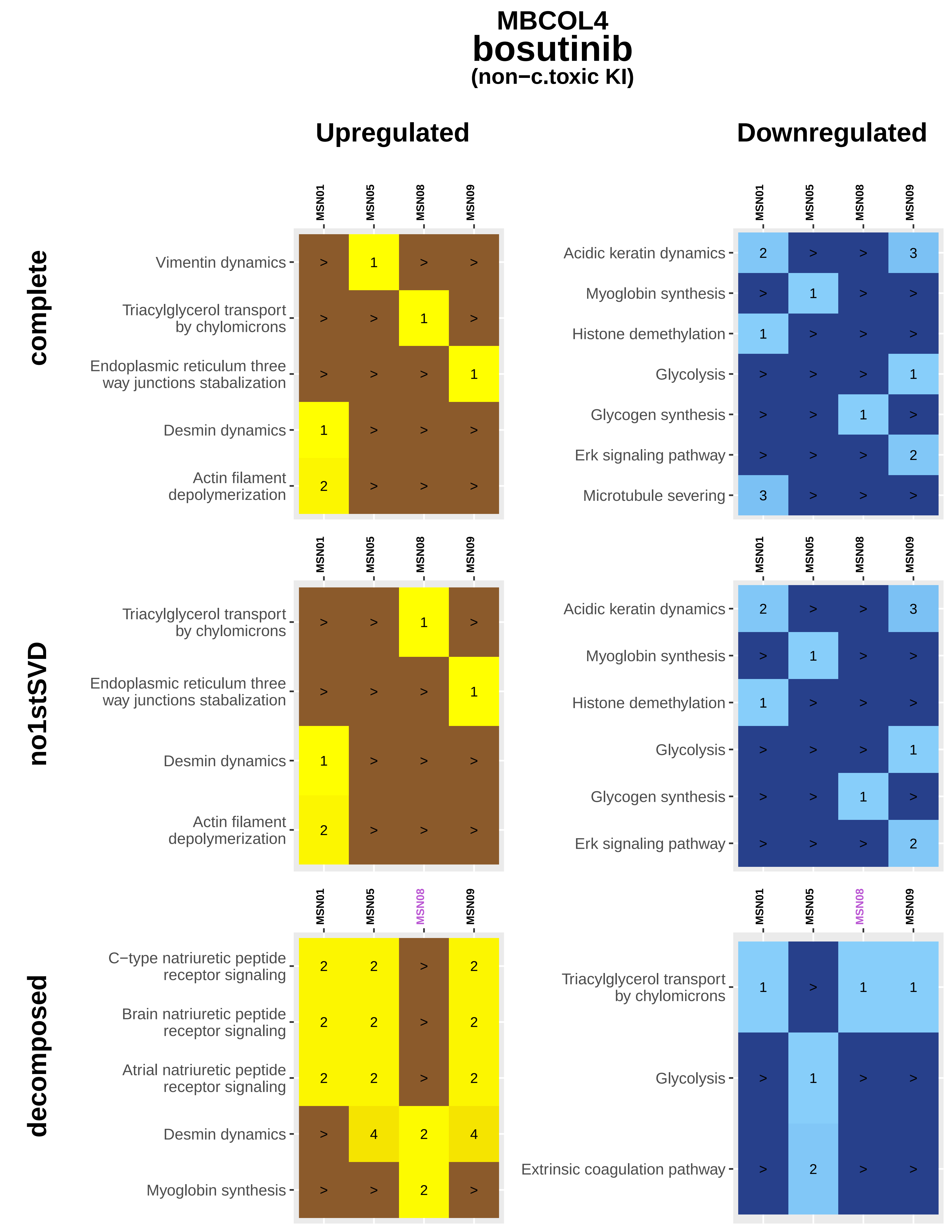
MBCOL4
bosutinib
(non−c.toxic KI)
Upregulated
Downregulated
complete
no1stSVD
decomposed
MSN01
MSN05
MSN08
MSN09
Vimentin dynamics
Triacylglycerol transport by chylomicrons
Endoplasmic reticulum three way junctions stabalization
Desmin dynamics
Actin filament depolymerization
Acidic keratin dynamics
Myoglobin synthesis
Histone demethylation
Glycolysis
Glycogen synthesis
Erk signaling pathway
Microtubule severing
C−type natriuretic peptide receptor signaling
Brain natriuretic peptide receptor signaling
Atrial natriuretic peptide receptor signaling
Extrinsic coagulation pathway

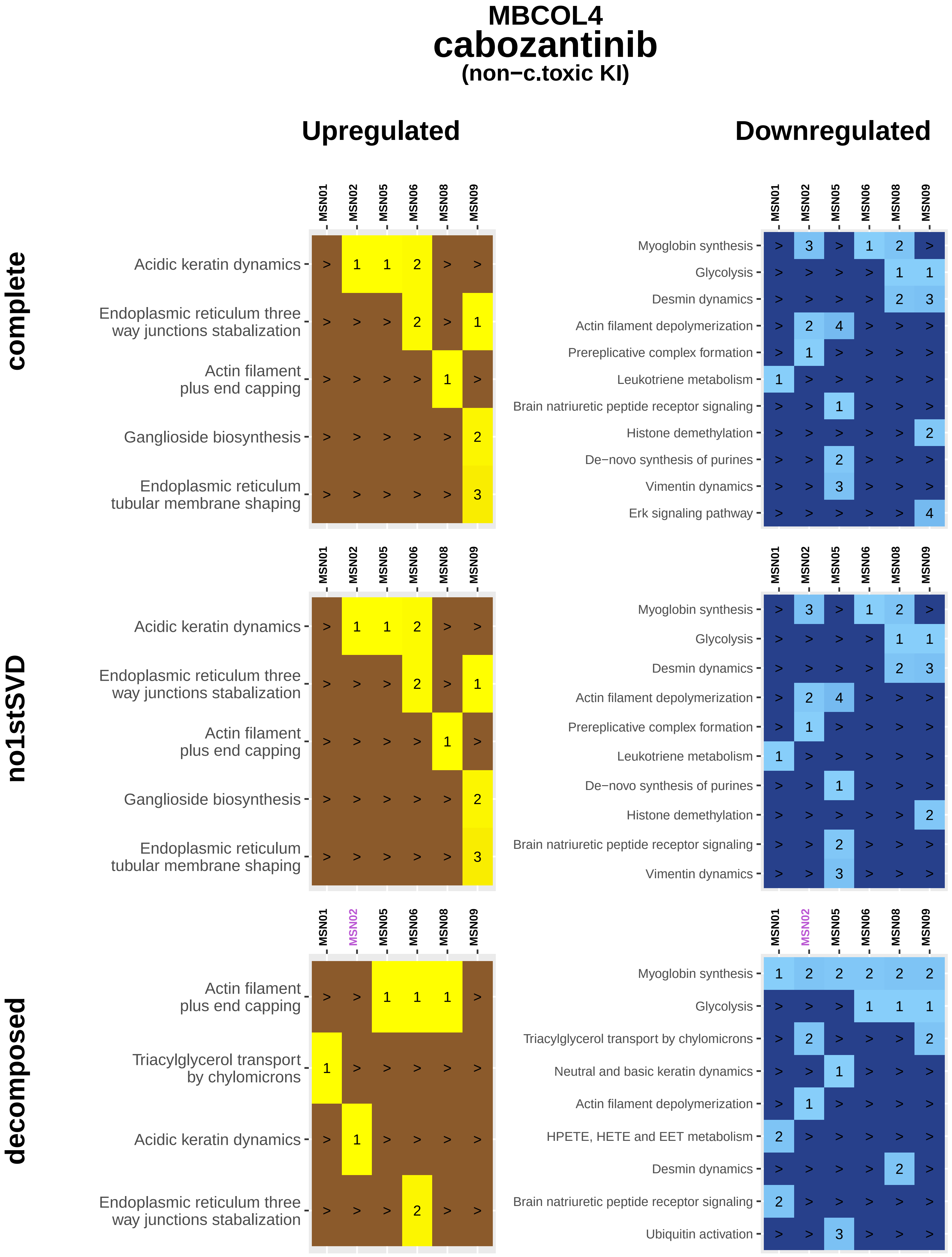
MBCOL4
cabozantinib
(non−c.toxic KI)
Upregulated
Downregulated
complete
no1stSVD
decomposed
MSN01
MSN02
MSN05
MSN06
MSN08
MSN09
Acidic keratin dynamics
Endoplasmic reticulum three way junctions stabalization
Actin filament plus end capping
Ganglioside biosynthesis
Endoplasmic reticulum tubular membrane shaping
Triacylglycerol transport by chylomicrons
Myoglobin synthesis
Glycolysis
Desmin dynamics
Actin filament depolymerization
Prereplicative complex formation
Leukotriene metabolism
Brain natriuretic peptide receptor signaling
Histone demethylation
De−novo synthesis of purines
Vimentin dynamics
Erk signaling pathway
Neutral and basic keratin dynamics
HPETE, HETE and EET metabolism
Ubiquitin activation

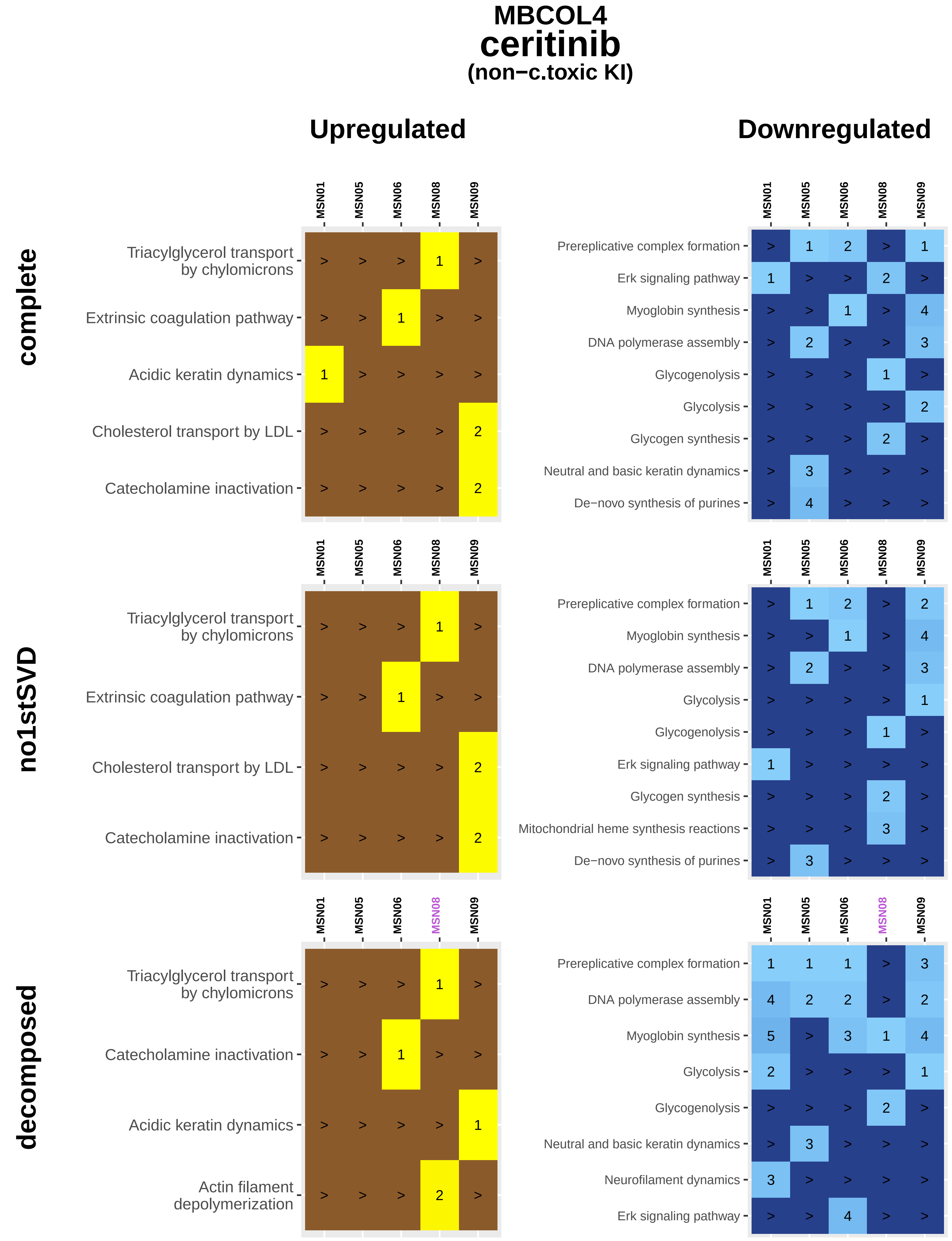
MBCOL4
ceritinib
(non−c.toxic KI)
Upregulated
Downregulated
complete
no1stSVD
decomposed
MSN01
MSN05
MSN06
MSN08
MSN09
Triacylglycerol transport by chylomicrons
Extrinsic coagulation pathway
Acidic keratin dynamics
Cholesterol transport by LDL
Catecholamine inactivation
Actin filament depolymerization
Prereplicative complex formation
Erk signaling pathway
Myoglobin synthesis
DNA polymerase assembly
Glycogenolysis
Glycolysis
Glycogen synthesis
Neutral and basic keratin dynamics
De−novo synthesis of purines
Mitochondrial heme synthesis reactions
Neurofilament dynamics

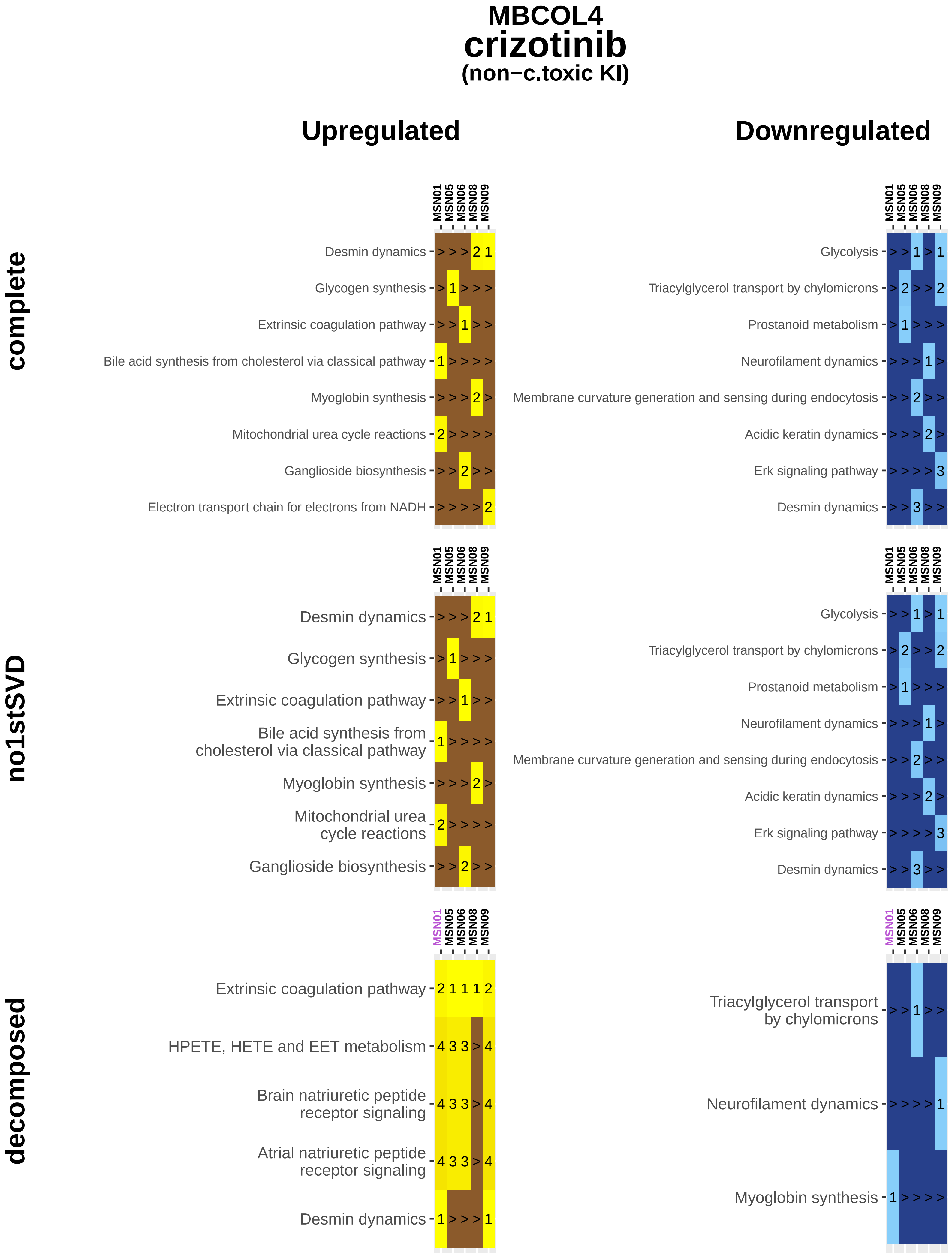
MBCOL4
crizotinib
(non−c.toxic KI)
Upregulated
Downregulated
complete
no1stSVD
decomposed
MSN01
MSN05
MSN06
MSN08
MSN09
Desmin dynamics
Glycogen synthesis
Extrinsic coagulation pathway
Bile acid synthesis from cholesterol via classical pathway
Myoglobin synthesis
Mitochondrial urea cycle reactions
Ganglioside biosynthesis
Electron transport chain for electrons from NADH
Glycolysis
Triacylglycerol transport by chylomicrons
Prostanoid metabolism
Neurofilament dynamics
Membrane curvature generation and sensing during endocytosis
Acidic keratin dynamics
Erk signaling pathway
HPETE, HETE and EET metabolism
Brain natriuretic peptide receptor signaling
Atrial natriuretic peptide receptor signaling

### MBCOL4
### dabrafenib
#### (c.toxic KI)

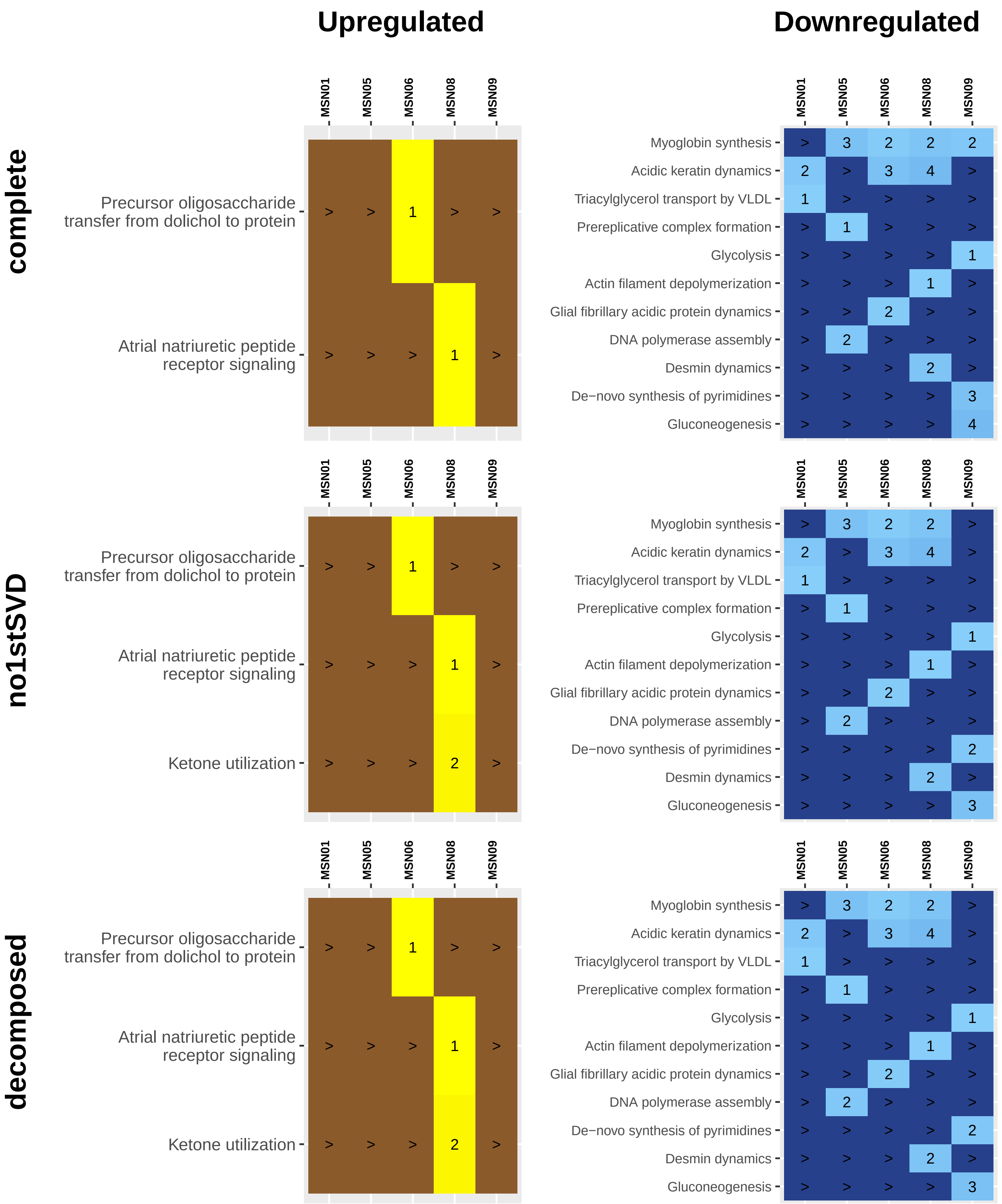

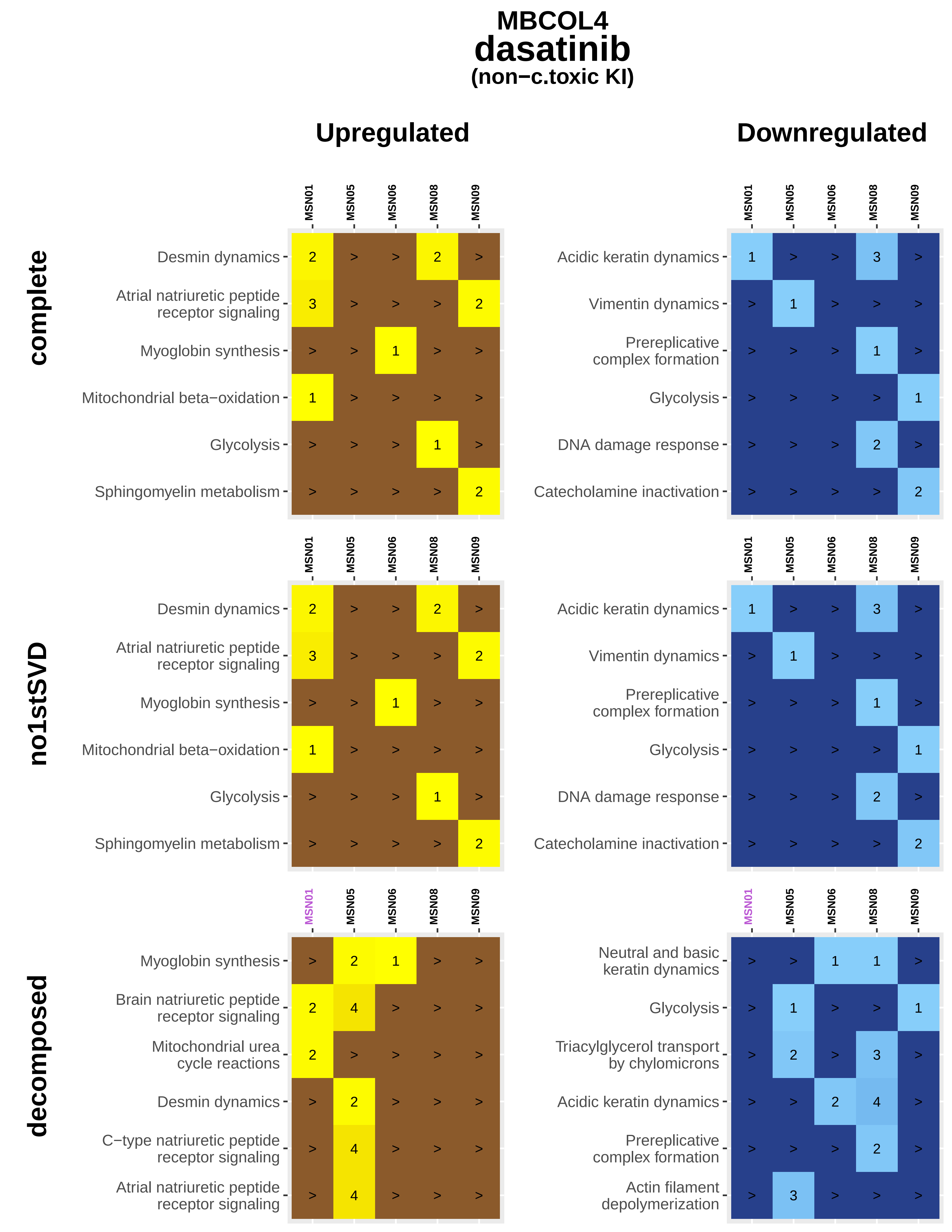
MBCOL4
dasatinib
(non−c.toxic KI)
Upregulated
Downregulated
complete
no1stSVD
decomposed
MSN01
MSN05
MSN06
MSN08
MSN09
Desmin dynamics 2 > > 2 >
Atrial natriuretic peptide receptor signaling 3 > > > 2
Myoglobin synthesis > > 1 > >
Mitochondrial beta−oxidation 1 > > > >
Glycolysis > > > 1 >
Sphingomyelin metabolism > > > > 2
Acidic keratin dynamics 1 > > 3 >
Vimentin dynamics > 1 > > >
Prereplicative complex formation > > > 1 >
Glycolysis > > > > 1
DNA damage response > > > 2 >
Catecholamine inactivation > > > > 2
Desmin dynamics 2 > > 2 >
Atrial natriuretic peptide receptor signaling 3 > > > 2
Myoglobin synthesis > > 1 > >
Mitochondrial beta−oxidation 1 > > > >
Glycolysis > > > 1 >
Sphingomyelin metabolism > > > > 2
Acidic keratin dynamics 1 > > 3 >
Vimentin dynamics > 1 > > >
Prereplicative complex formation > > > 1 >
Glycolysis > > > > 1
DNA damage response > > > 2 >
Catecholamine inactivation > > > > 2
Myoglobin synthesis > 2 1 > >
Brain natriuretic peptide receptor signaling 2 4 > > >
Mitochondrial urea cycle reactions 2 > > > >
Desmin dynamics > 2 > > >
C−type natriuretic peptide receptor signaling > 4 > > >
Atrial natriuretic peptide receptor signaling > 4 > > >
Neutral and basic keratin dynamics > > 1 1 >
Glycolysis > 1 > > 1
Triacylglycerol transport by chylomicrons > 2 > 3 >
Acidic keratin dynamics > > 2 4 >
Prereplicative complex formation > > > 2 >
Actin filament depolymerization > 3 > > >

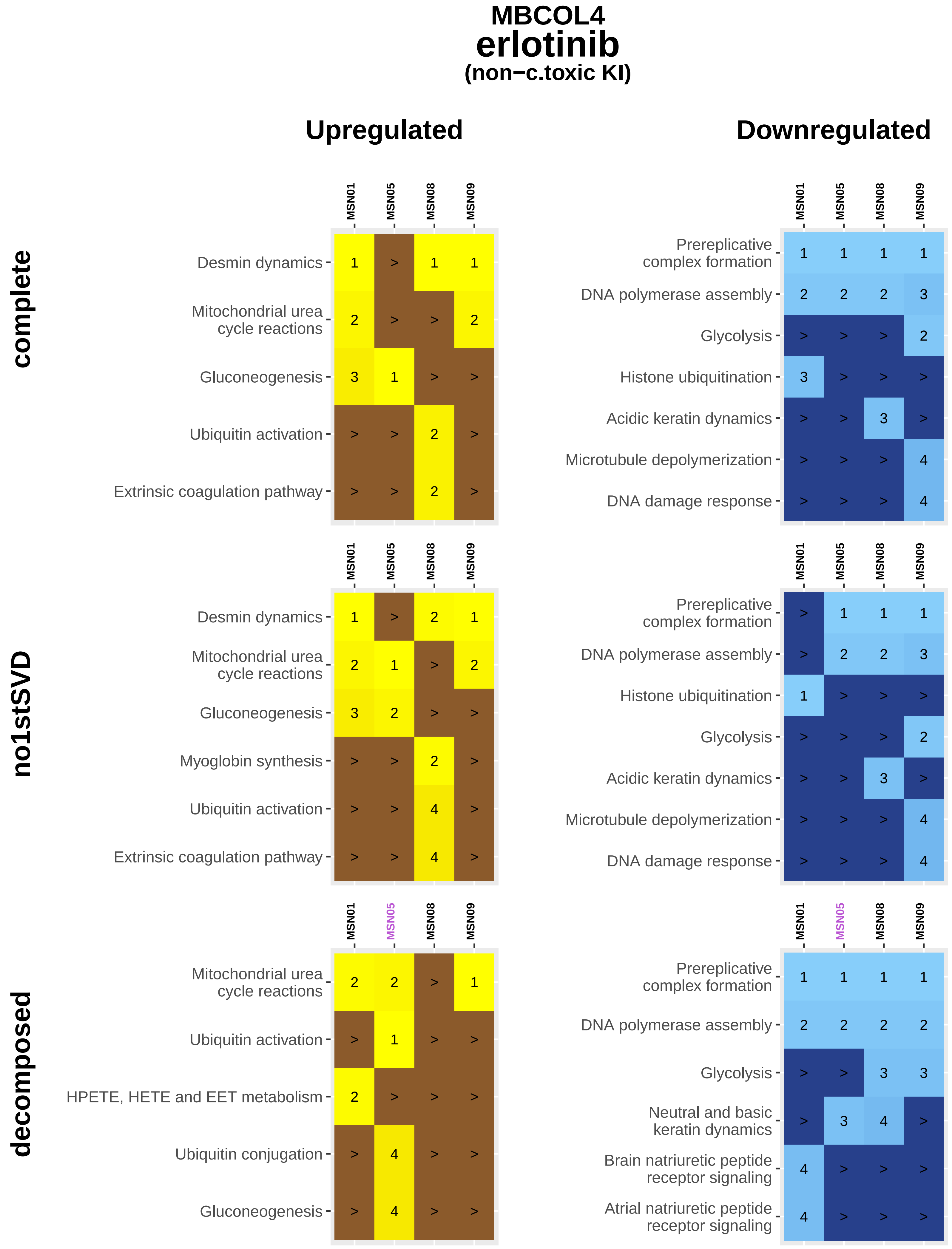
MBCOL4
erlotinib
(non−c.toxic KI)
Upregulated
Downregulated
complete
no1stSVD
decomposed
MSN01
MSN05
MSN08
MSN09
Desmin dynamics
Mitochondrial urea cycle reactions
Gluconeogenesis
Ubiquitin activation
Extrinsic coagulation pathway
Myoglobin synthesis
HPETE, HETE and EET metabolism
Ubiquitin conjugation
Prereplicative complex formation
DNA polymerase assembly
Glycolysis
Histone ubiquitination
Acidic keratin dynamics
Microtubule depolymerization
DNA damage response
Neutral and basic keratin dynamics
Brain natriuretic peptide receptor signaling
Atrial natriuretic peptide receptor signaling

### MBCOL4
### gefitinib
#### (non−c.toxic KI)

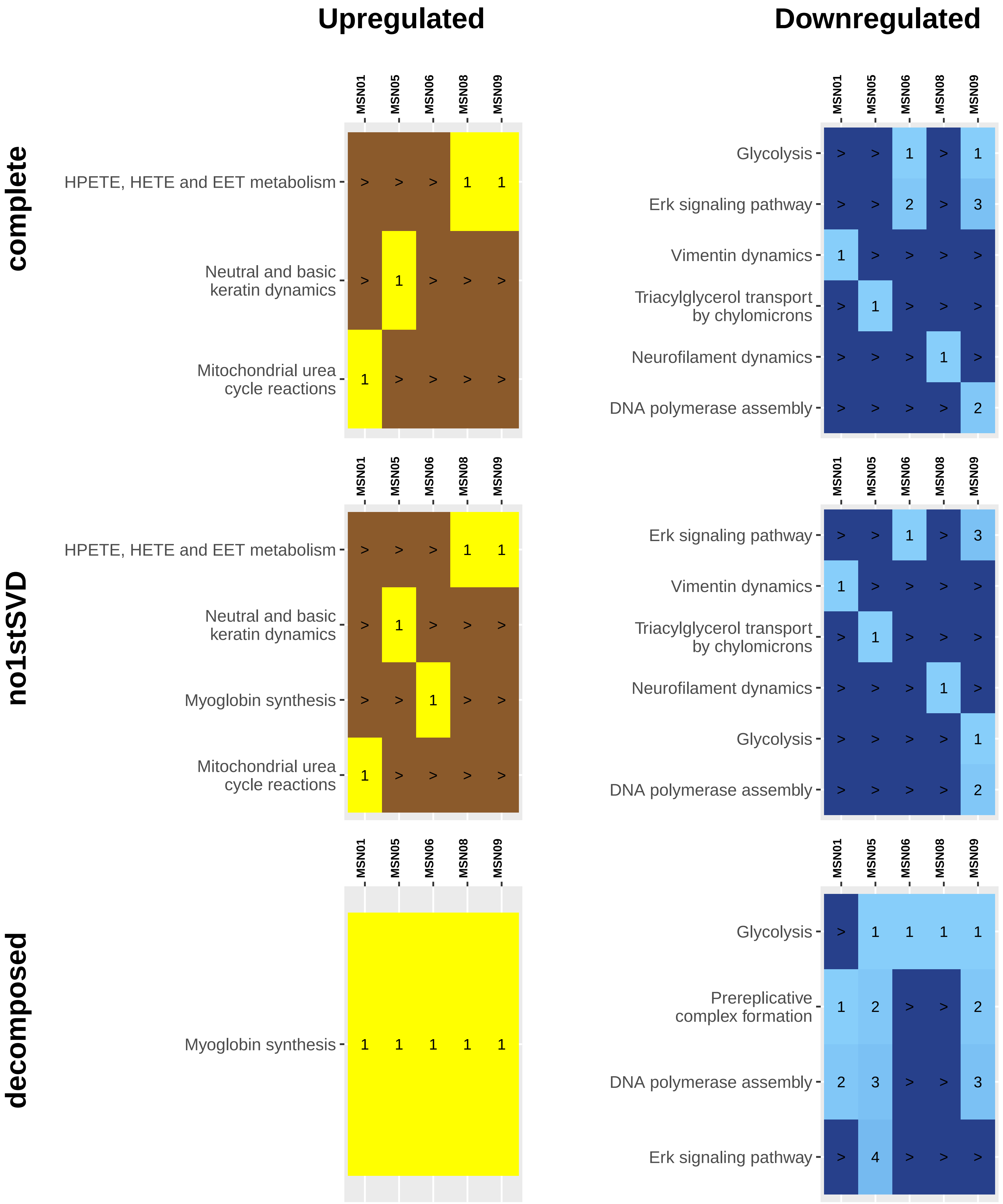

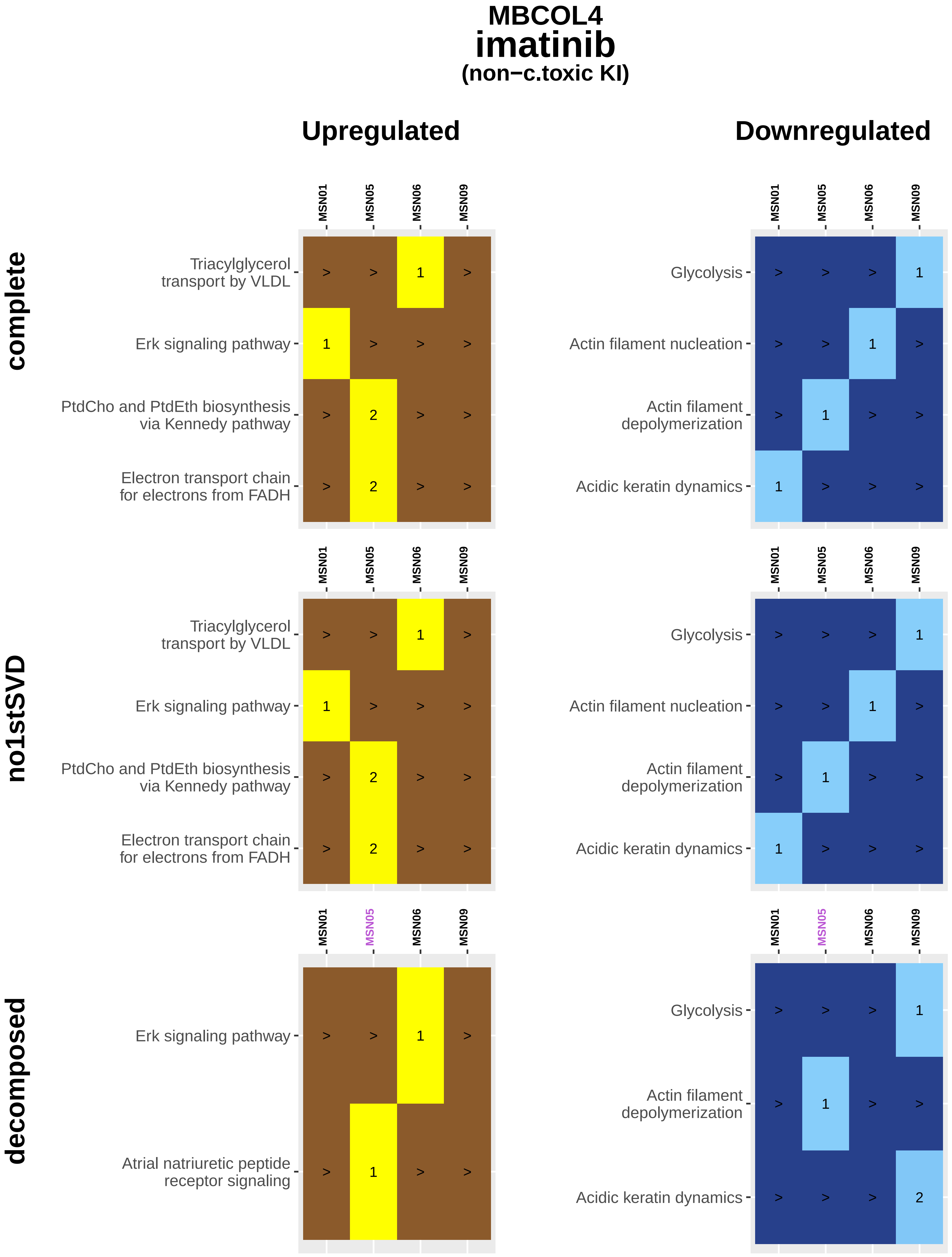

MBCOL4
imatinib
(non−c.toxic KI)
Upregulated
Downregulated
complete
no1stSVD
decomposed
MSN01
MSN05
MSN06
MSN09
Triacylglycerol transport by VLDL
Erk signaling pathway
PtdCho and PtdEth biosynthesis via Kennedy pathway
Electron transport chain for electrons from FADH
Atrial natriuretic peptide receptor signaling
Glycolysis
Actin filament nucleation
Actin filament depolymerization
Acidic keratin dynamics

### MBCOL4
### lapatinib
#### (c.toxic KI)

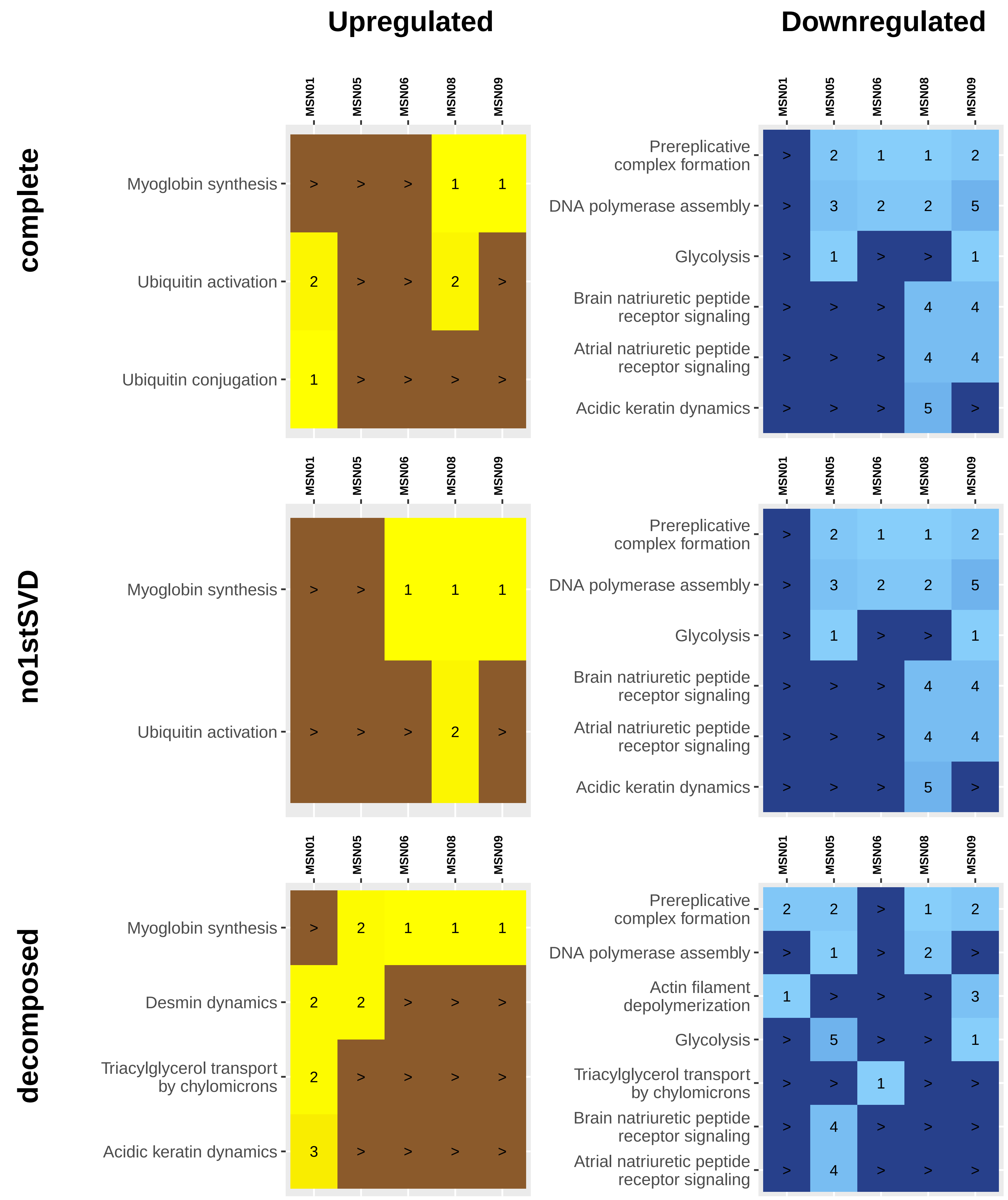

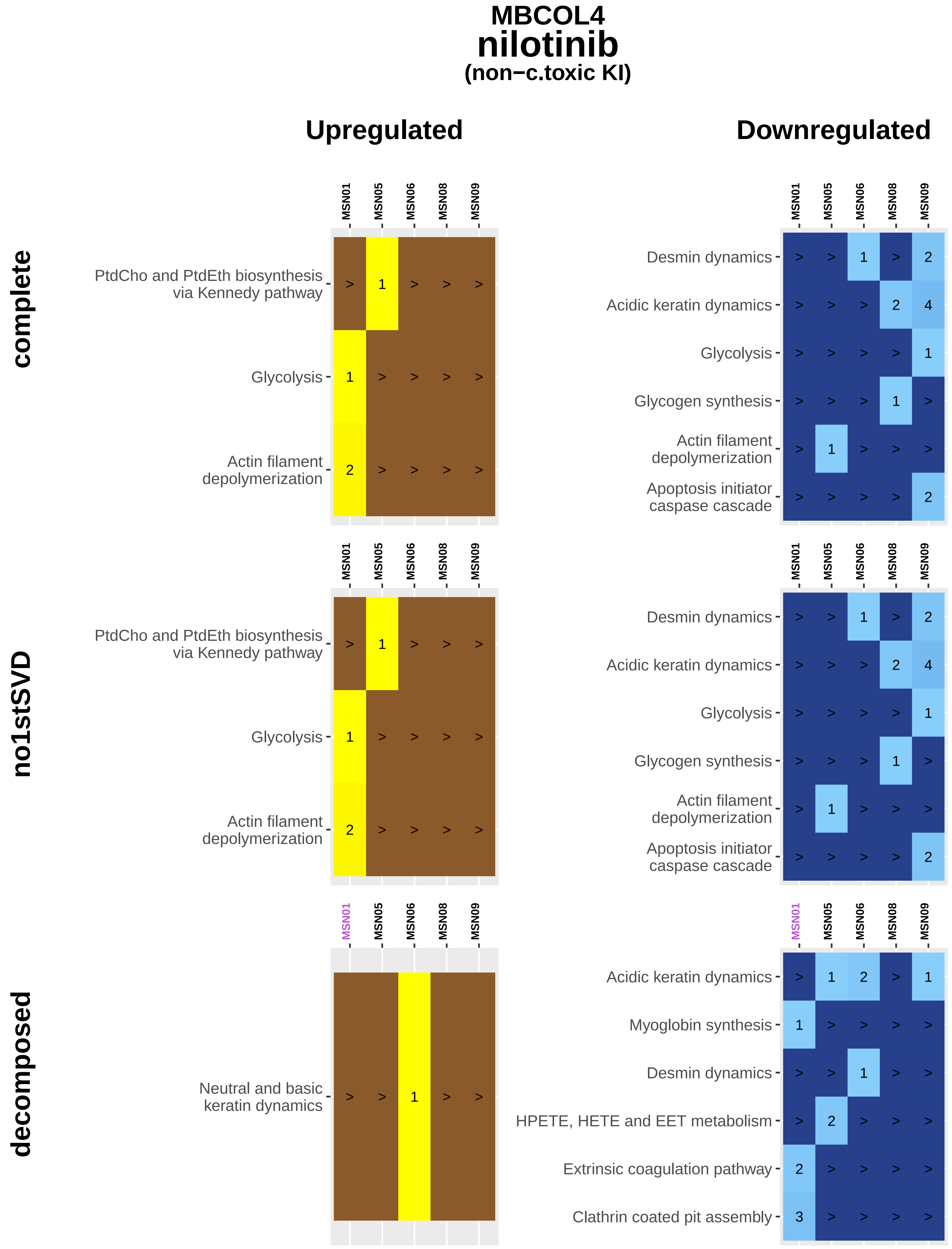
MBCOL4
nilotinib
(non−c.toxic KI)
Upregulated
Downregulated
complete
no1stSVD
decomposed
MSN01
MSN05
MSN06
MSN08
MSN09
PtdCho and PtdEth biosynthesis via Kennedy pathway
Glycolysis
Actin filament depolymerization
Desmin dynamics
Acidic keratin dynamics
Glycogen synthesis
Apoptosis initiator caspase cascade
Neutral and basic keratin dynamics
Myoglobin synthesis
HPETE, HETE and EET metabolism
Extrinsic coagulation pathway
Clathrin coated pit assembly

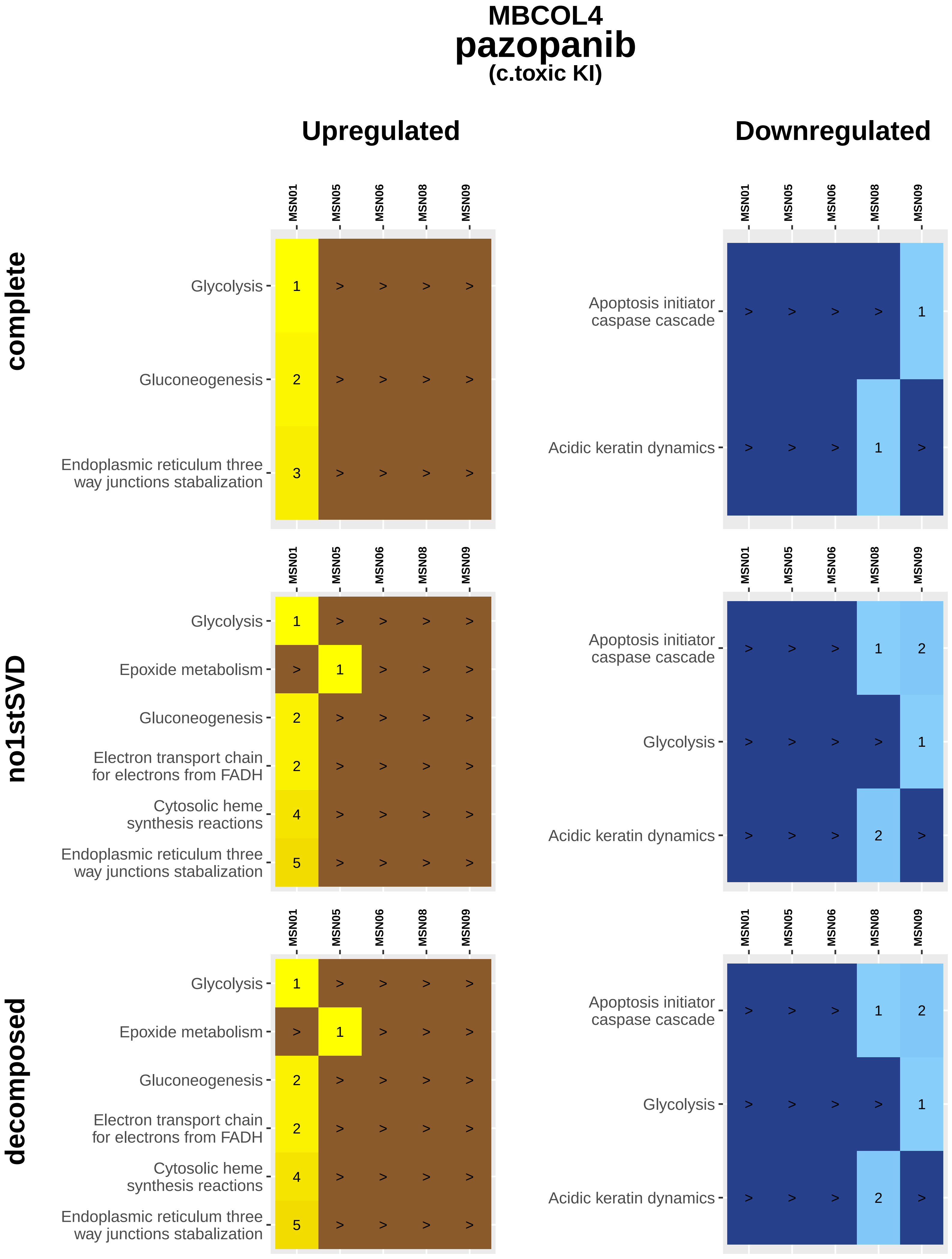

MBCOL4
pazopanib
(c.toxic KI)
Upregulated
Downregulated
complete
no1stSVD
decomposed
MSN01
MSN05
MSN06
MSN08
MSN09
Glycolysis 1 > > > >
Gluconeogenesis 2 > > > >
Endoplasmic reticulum three way junctions stabalization 3 > > > >
Apoptosis initiator caspase cascade > > > > 1
Acidic keratin dynamics > > > 1 >
Glycolysis 1 > > > >
Epoxide metabolism > 1 > > >
Gluconeogenesis 2 > > > >
Electron transport chain for electrons from FADH 2 > > > >
Cytosolic heme synthesis reactions 4 > > > >
Endoplasmic reticulum three way junctions stabalization 5 > > > >
Apoptosis initiator caspase cascade > > > 1 2
Glycolysis > > > > 1
Acidic keratin dynamics > > > 2 >
Glycolysis 1 > > > >
Epoxide metabolism > 1 > > >
Gluconeogenesis 2 > > > >
Electron transport chain for electrons from FADH 2 > > > >
Cytosolic heme synthesis reactions 4 > > > >
Endoplasmic reticulum three way junctions stabalization 5 > > > >
Apoptosis initiator caspase cascade > > > 1 2
Glycolysis > > > > 1
Acidic keratin dynamics > > > 2 >

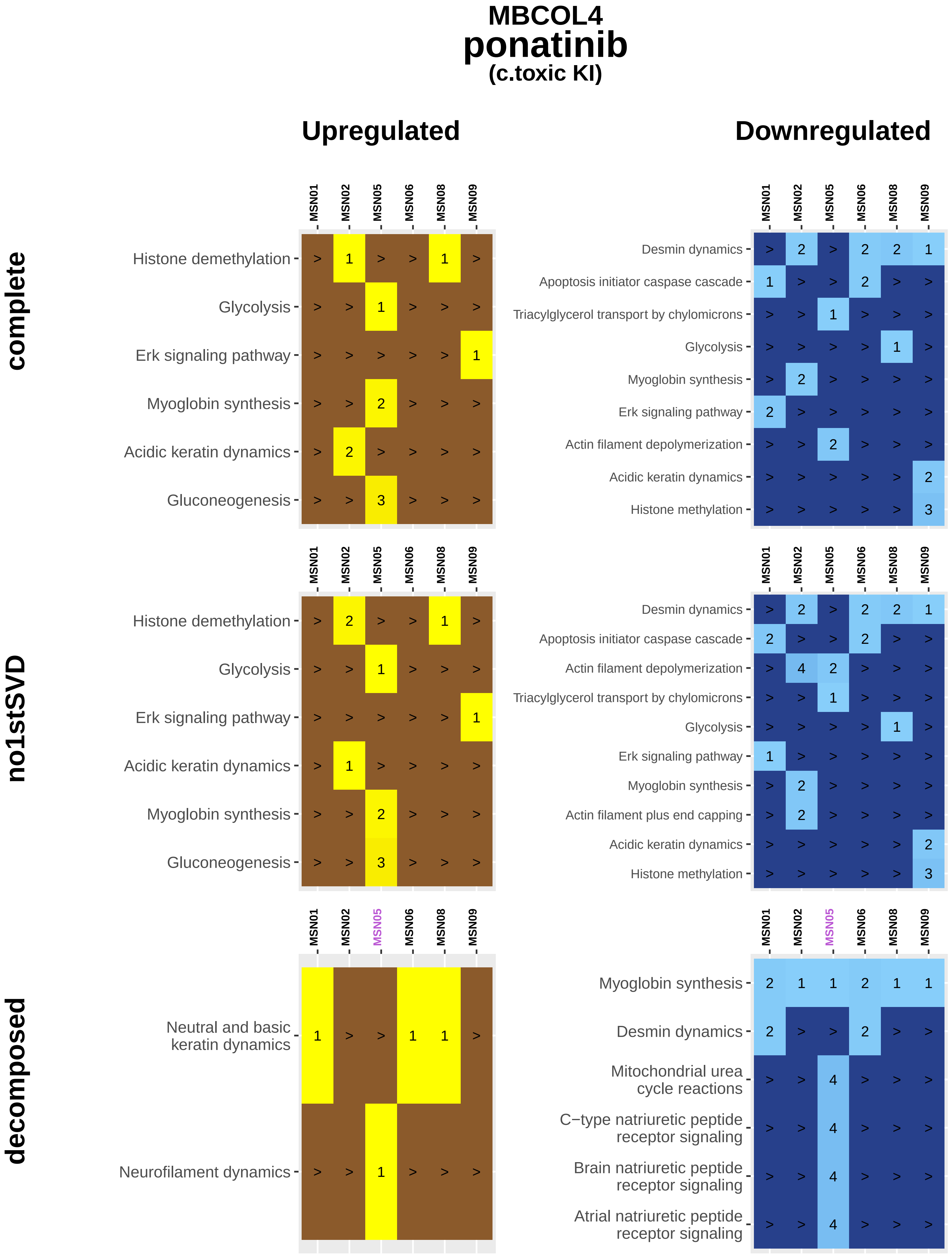
MBCOL4
ponatinib
(c.toxic KI)
Upregulated
Downregulated
complete
no1stSVD
decomposed
MSN01
MSN02
MSN05
MSN06
MSN08
MSN09
Histone demethylation
Glycolysis
Erk signaling pathway
Myoglobin synthesis
Acidic keratin dynamics
Gluconeogenesis
Desmin dynamics
Apoptosis initiator caspase cascade
Triacylglycerol transport by chylomicrons
Actin filament depolymerization
Histone methylation
Actin filament plus end capping
Neutral and basic keratin dynamics
Neurofilament dynamics
Mitochondrial urea cycle reactions
C−type natriuretic peptide receptor signaling
Brain natriuretic peptide receptor signaling
Atrial natriuretic peptide receptor signaling

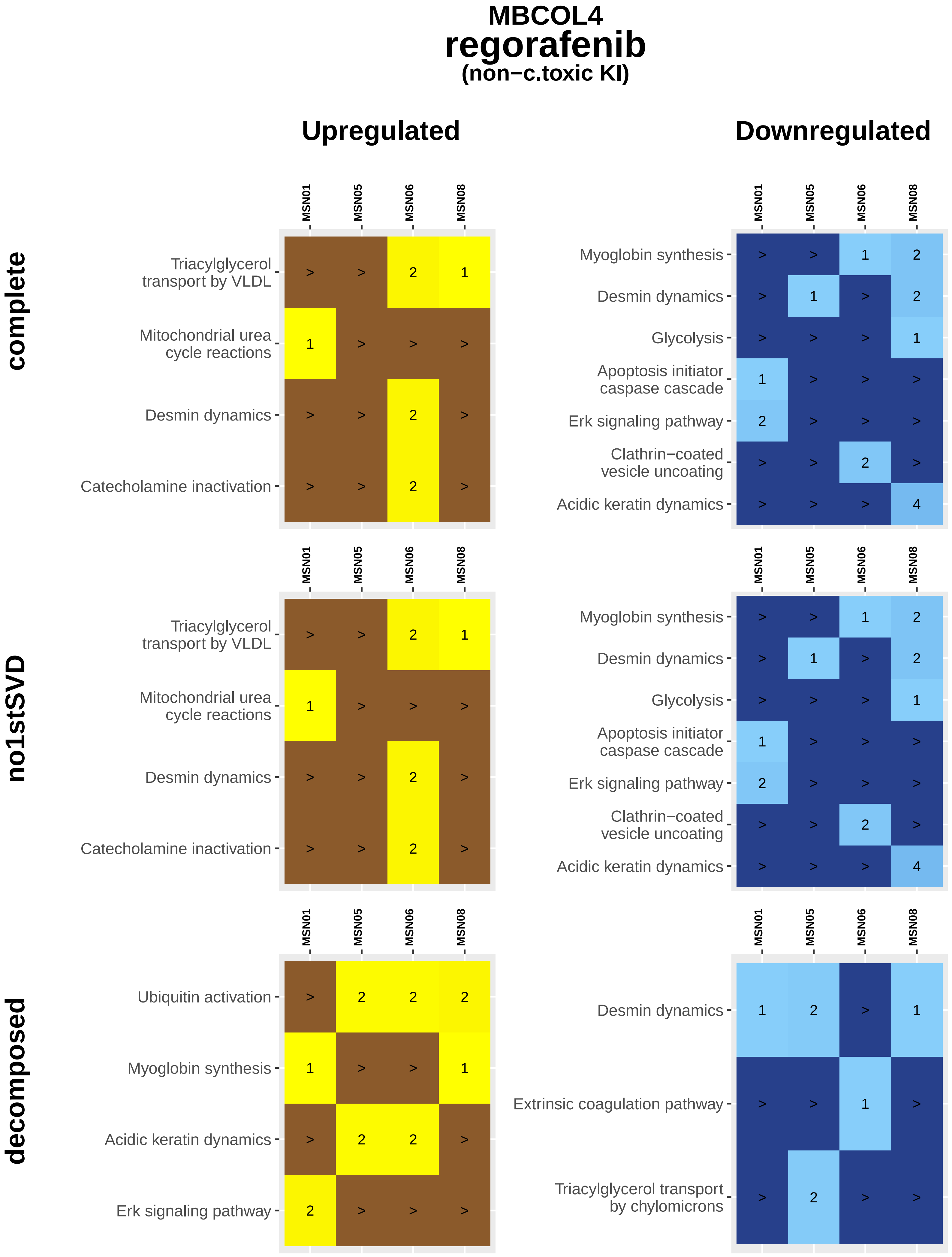

MBCOL4
regorafenib
(non−c.toxic KI)
Upregulated
Downregulated
complete
no1stSVD
decomposed
MSN01
MSN05
MSN06
MSN08
Triacylglycerol transport by VLDL
Mitochondrial urea cycle reactions
Desmin dynamics
Catecholamine inactivation
Myoglobin synthesis
Glycolysis
Apoptosis initiator caspase cascade
Erk signaling pathway
Clathrin−coated vesicle uncoating
Acidic keratin dynamics
Ubiquitin activation
Extrinsic coagulation pathway
Triacylglycerol transport by chylomicrons

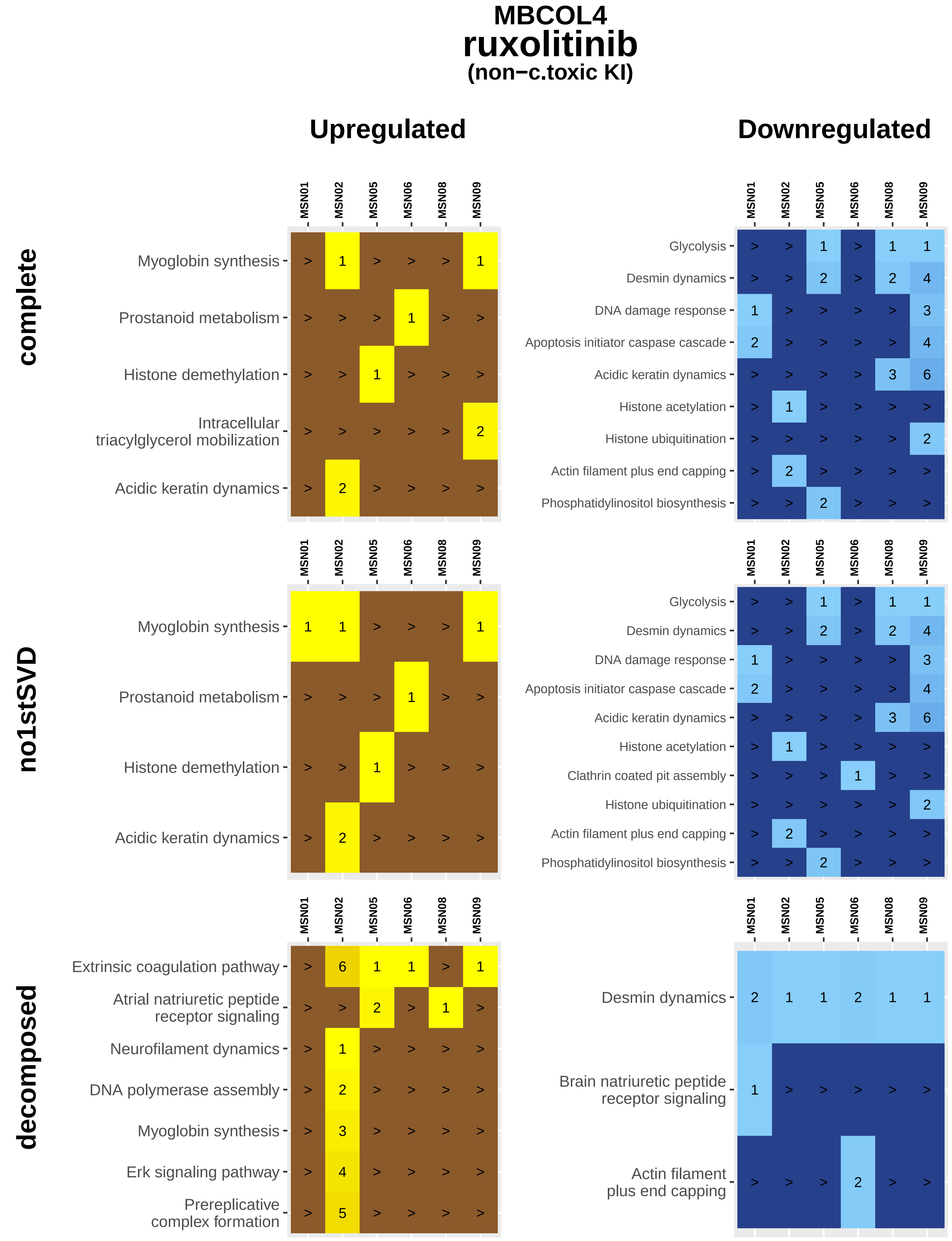

MBCOL4
ruxolitinib
(non−c.toxic KI)
Upregulated
Downregulated
complete
no1stSVD
decomposed
MSN01
MSN02
MSN05
MSN06
MSN08
MSN09
Myoglobin synthesis
Prostanoid metabolism
Histone demethylation
Intracellular triacylglycerol mobilization
Acidic keratin dynamics
Glycolysis
Desmin dynamics
DNA damage response
Apoptosis initiator caspase cascade
Histone acetylation
Histone ubiquitination
Actin filament plus end capping
Phosphatidylinositol biosynthesis
Clathrin coated pit assembly
Extrinsic coagulation pathway
Atrial natriuretic peptide receptor signaling
Neurofilament dynamics
DNA polymerase assembly
Erk signaling pathway
Prereplicative complex formation
Brain natriuretic peptide receptor signaling
Actin filament plus end capping

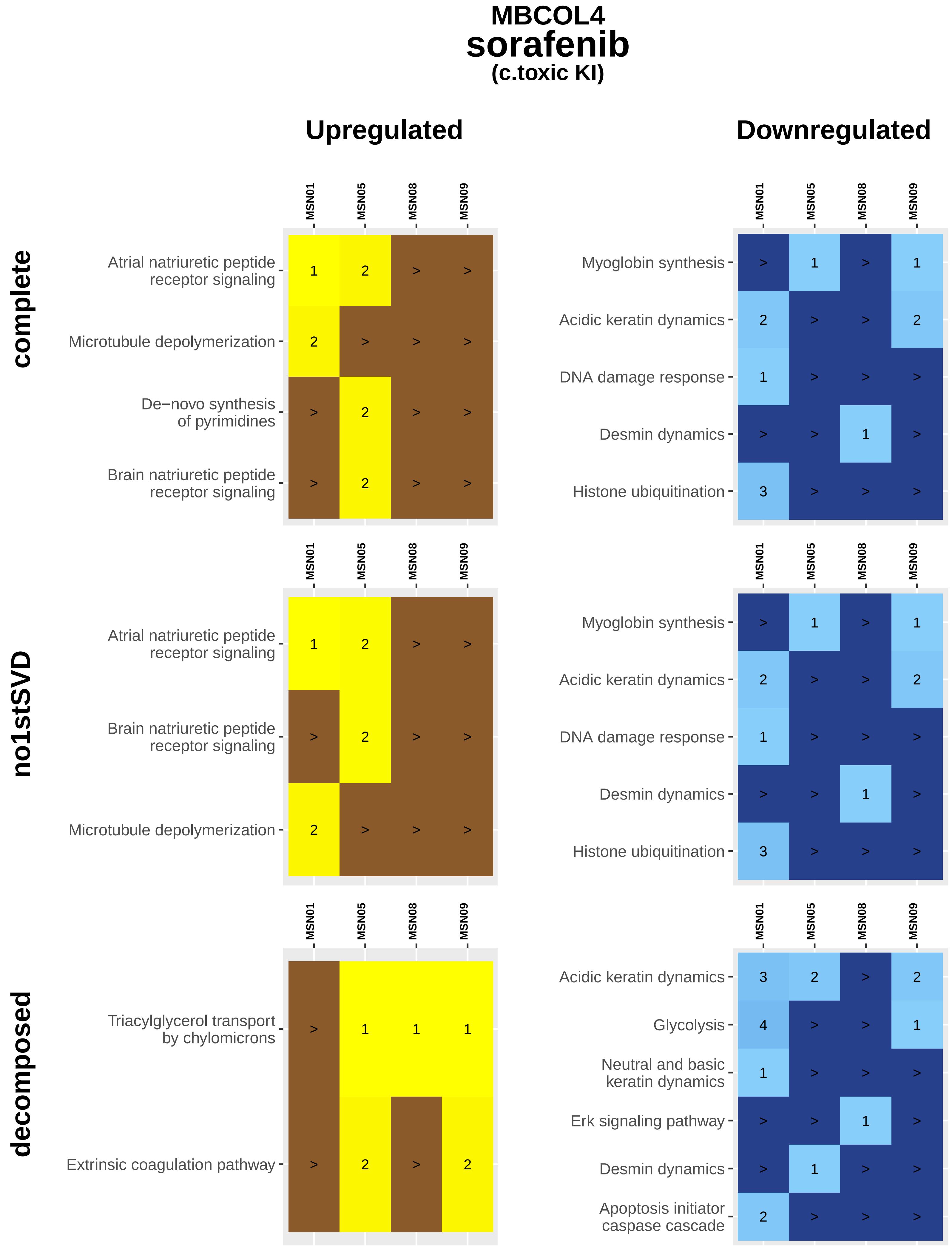
MBCOL4
sorafenib
(c.toxic KI)
Upregulated
Downregulated
complete
no1stSVD
decomposed
MSN01
MSN05
MSN08
MSN09
Atrial natriuretic peptide receptor signaling
Microtubule depolymerization
De−novo synthesis of pyrimidines
Brain natriuretic peptide receptor signaling
Myoglobin synthesis
Acidic keratin dynamics
DNA damage response
Desmin dynamics
Histone ubiquitination
Triacylglycerol transport by chylomicrons
Extrinsic coagulation pathway
Glycolysis
Neutral and basic keratin dynamics
Erk signaling pathway
Apoptosis initiator caspase cascade

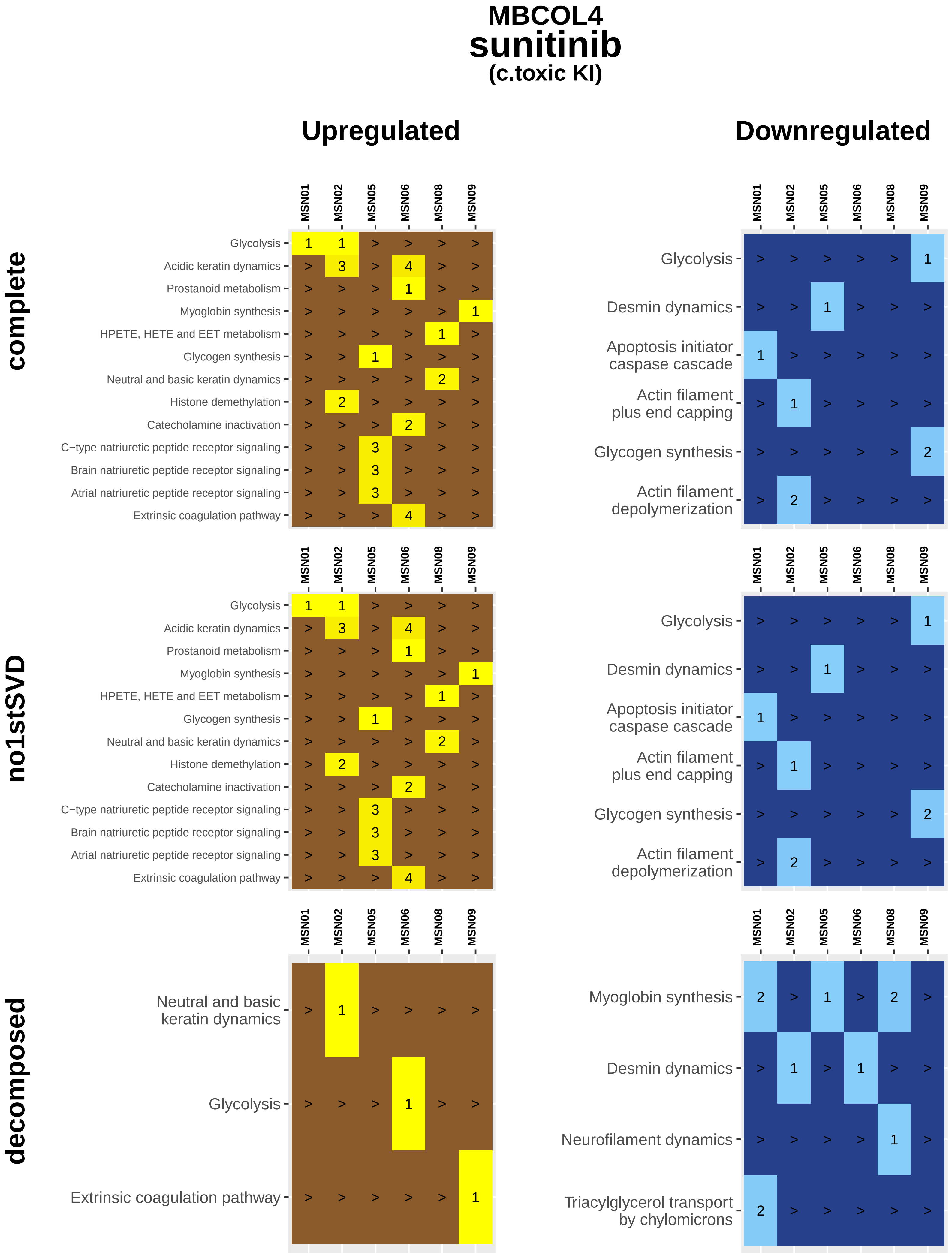
MBCOL4
sunitinib
(c.toxic KI)
Upregulated
Downregulated
complete
no1stSVD
decomposed
MSN01
MSN02
MSN05
MSN06
MSN08
MSN09
Glycolysis
Acidic keratin dynamics
Prostanoid metabolism
Myoglobin synthesis
HPETE, HETE and EET metabolism
Glycogen synthesis
Neutral and basic keratin dynamics
Histone demethylation
Catecholamine inactivation
C−type natriuretic peptide receptor signaling
Brain natriuretic peptide receptor signaling
Atrial natriuretic peptide receptor signaling
Extrinsic coagulation pathway
Desmin dynamics
Apoptosis initiator caspase cascade
Actin filament plus end capping
Actin filament depolymerization
Neutral and basic keratin dynamics
Neurofilament dynamics
Triacylglycerol transport by chylomicrons

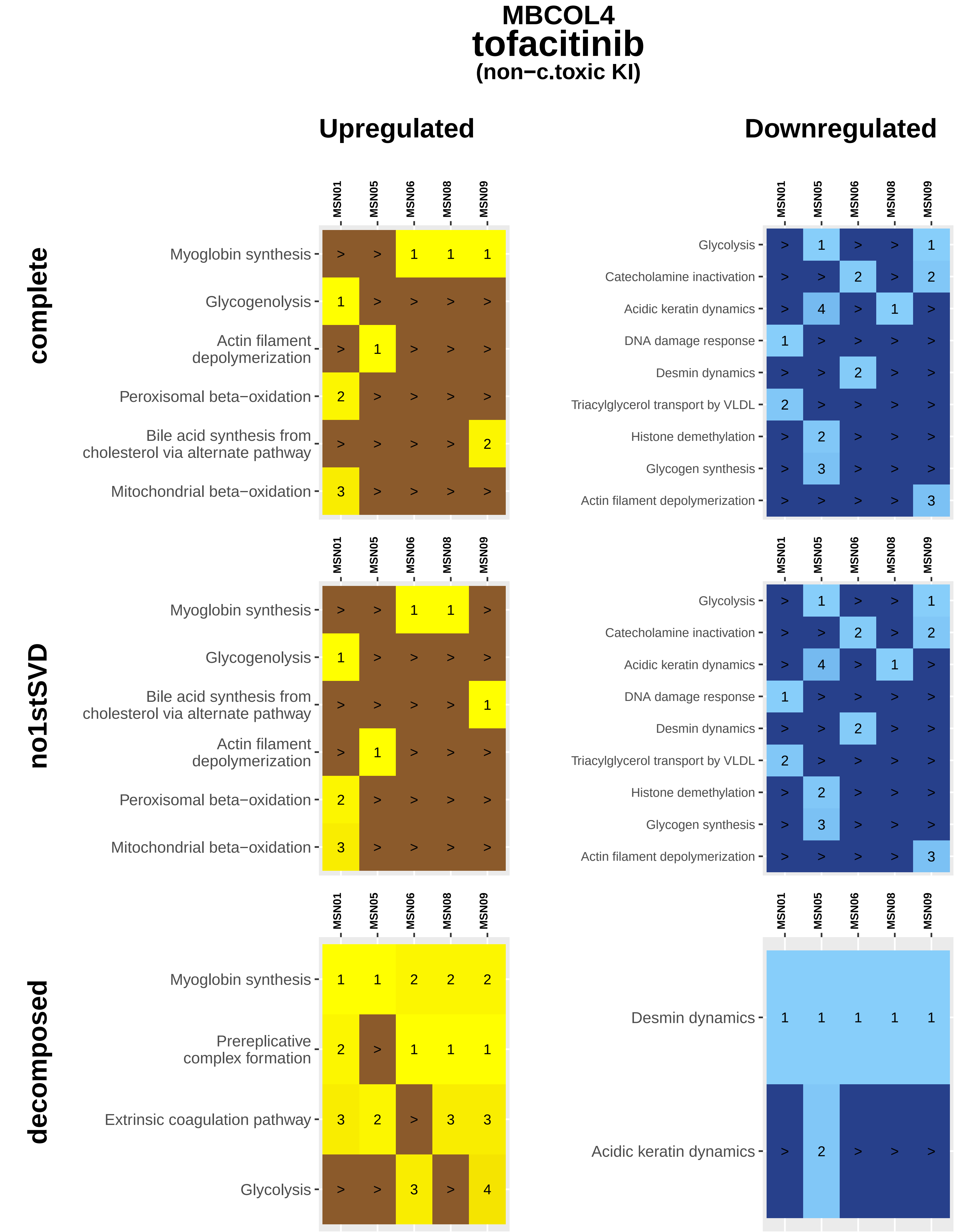
MBCOL4
tofacitinib
(non−c.toxic KI)
Upregulated
Downregulated
complete
no1stSVD
decomposed
MSN01
MSN05
MSN06
MSN08
MSN09
Myoglobin synthesis
Glycogenolysis
Actin filament depolymerization
Peroxisomal beta−oxidation
Bile acid synthesis from cholesterol via alternate pathway
Mitochondrial beta−oxidation
Glycolysis
Catecholamine inactivation
Acidic keratin dynamics
DNA damage response
Desmin dynamics
Triacylglycerol transport by VLDL
Histone demethylation
Glycogen synthesis
Actin filament depolymerization
Prereplicative complex formation
Extrinsic coagulation pathway

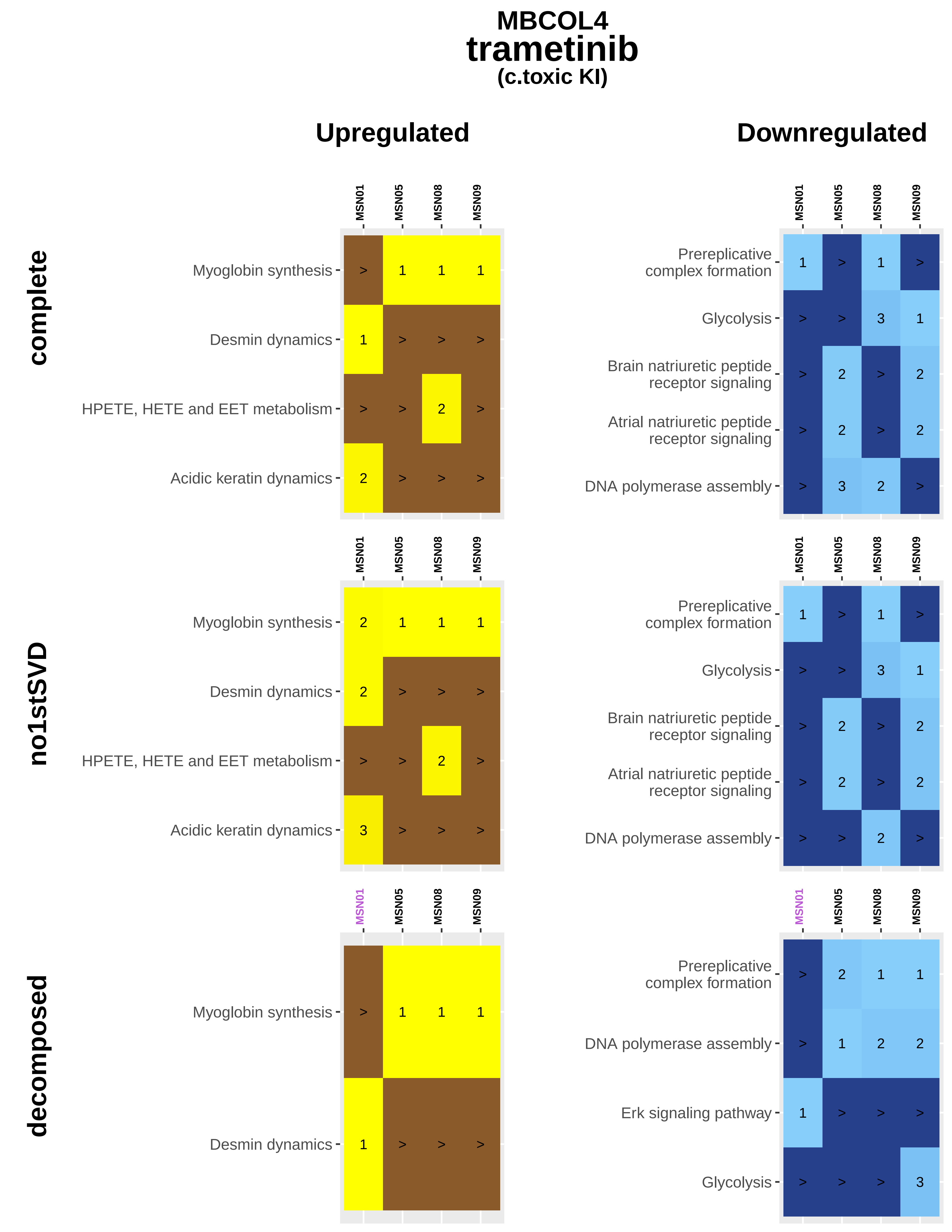
MBCOL4
trametinib
(c.toxic KI)
Upregulated
Downregulated
complete
no1stSVD
decomposed
MSN01
MSN05
MSN08
MSN09
Myoglobin synthesis
Desmin dynamics
HPETE, HETE and EET metabolism
Acidic keratin dynamics
Prereplicative complex formation
Glycolysis
Brain natriuretic peptide receptor signaling
Atrial natriuretic peptide receptor signaling
DNA polymerase assembly
Erk signaling pathway

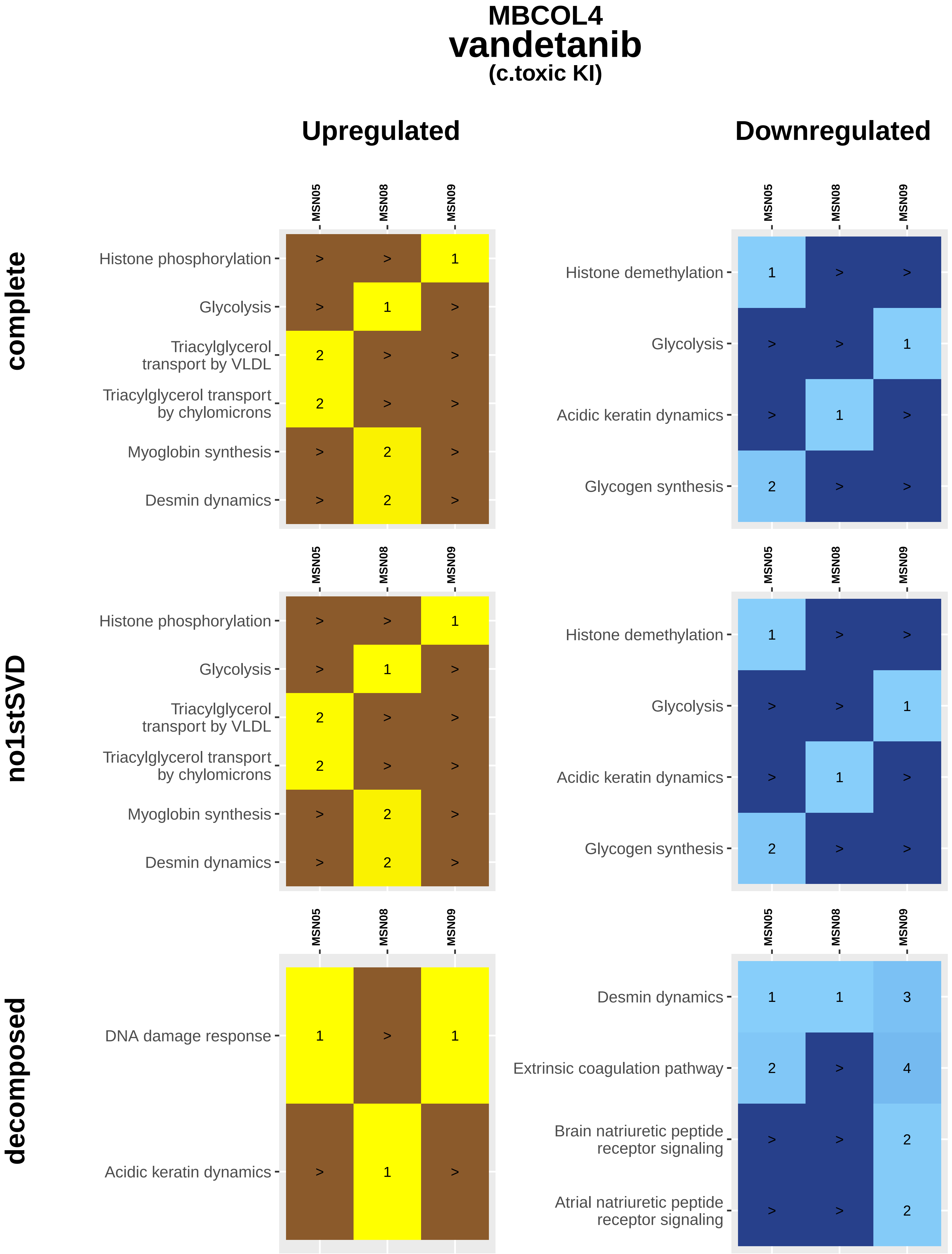

MBCOL4
vandetanib
(c.toxic KI)
Upregulated
Downregulated
complete
no1stSVD
decomposed
MSN05
MSN08
MSN09
Histone phosphorylation
Glycolysis
Triacylglycerol transport by VLDL
Triacylglycerol transport by chylomicrons
Myoglobin synthesis
Desmin dynamics
Histone demethylation
Acidic keratin dynamics
Glycogen synthesis
DNA damage response
Extrinsic coagulation pathway
Brain natriuretic peptide receptor signaling
Atrial natriuretic peptide receptor signaling

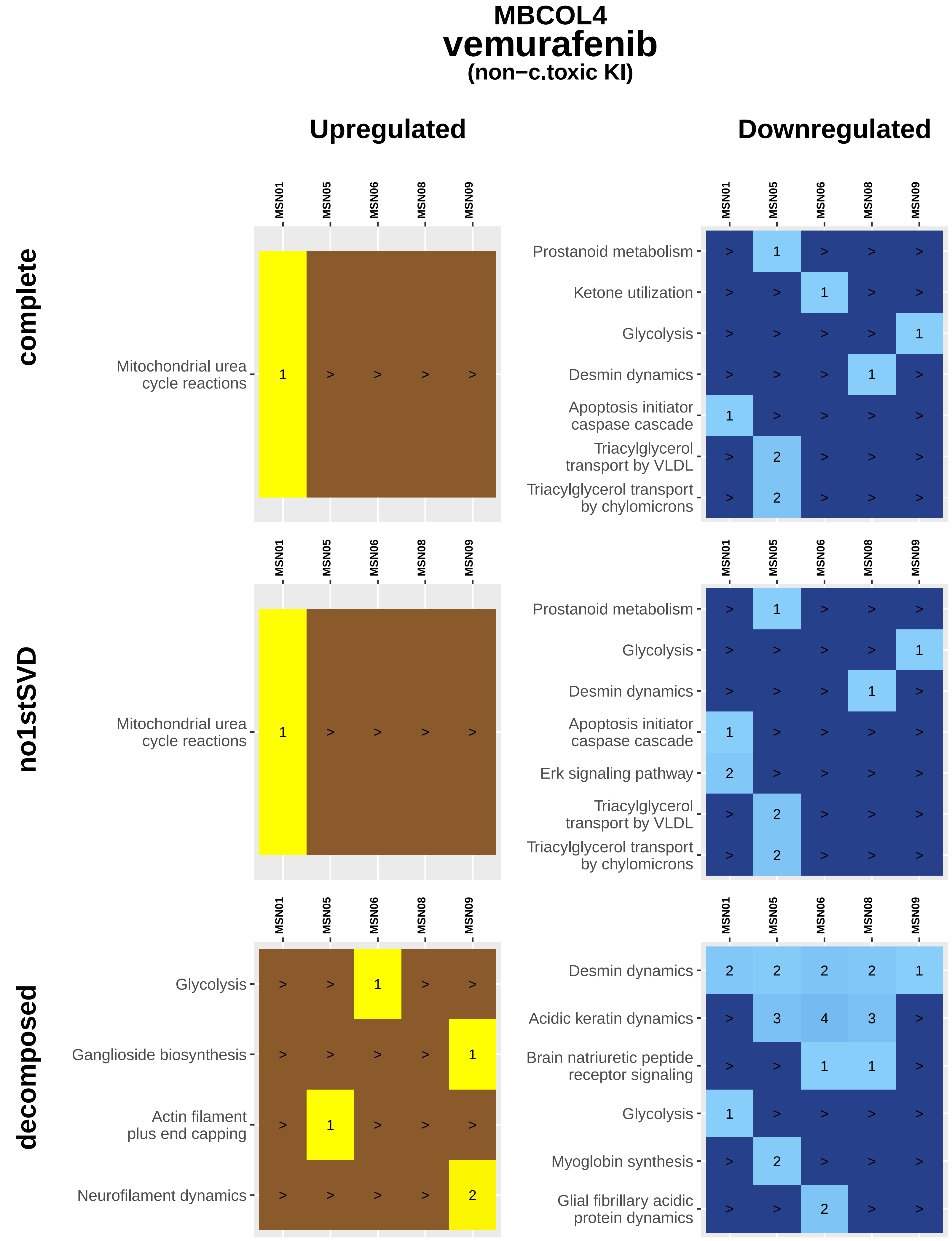

MBCOL4
vemurafenib
(non−c.toxic KI)
Upregulated
Downregulated
complete
no1stSVD
decomposed
MSN01
MSN05
MSN06
MSN08
MSN09
Mitochondrial urea cycle reactions
Prostanoid metabolism
Ketone utilization
Glycolysis
Desmin dynamics
Apoptosis initiator caspase cascade
Triacylglycerol transport by VLDL
Triacylglycerol transport by chylomicrons
Erk signaling pathway
Ganglioside biosynthesis
Actin filament plus end capping
Neurofilament dynamics
Acidic keratin dynamics
Brain natriuretic peptide receptor signaling
Myoglobin synthesis
Glial fibrillary acidic protein dynamics

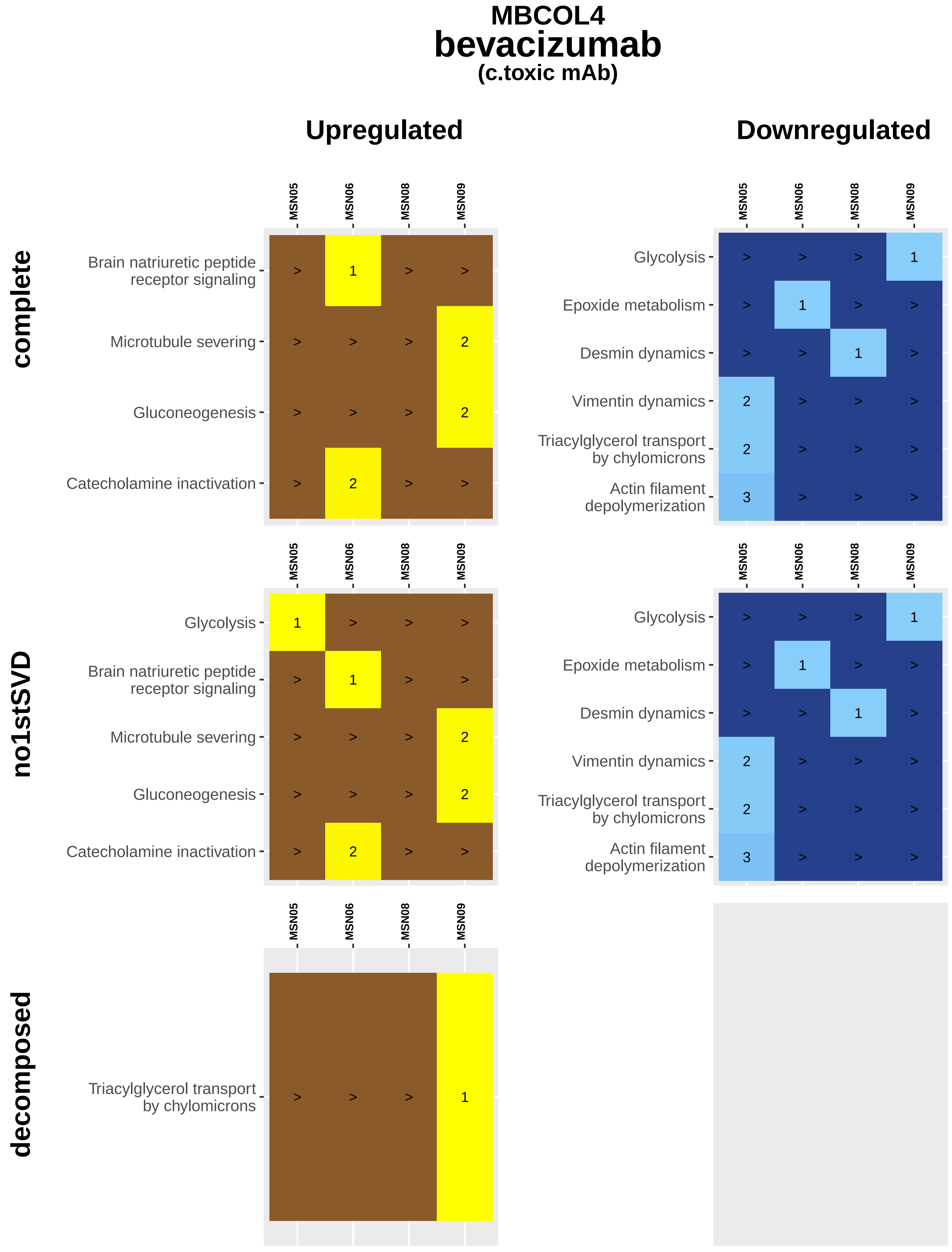

MBCOL4
bevacizumab
(c.toxic mAb)
Upregulated
Downregulated
complete
no1stSVD
decomposed
MSN05
MSN06
MSN08
MSN09
Brain natriuretic peptide receptor signaling
Microtubule severing
Gluconeogenesis
Catecholamine inactivation
Glycolysis
Epoxide metabolism
Desmin dynamics
Vimentin dynamics
Triacylglycerol transport by chylomicrons
Actin filament depolymerization

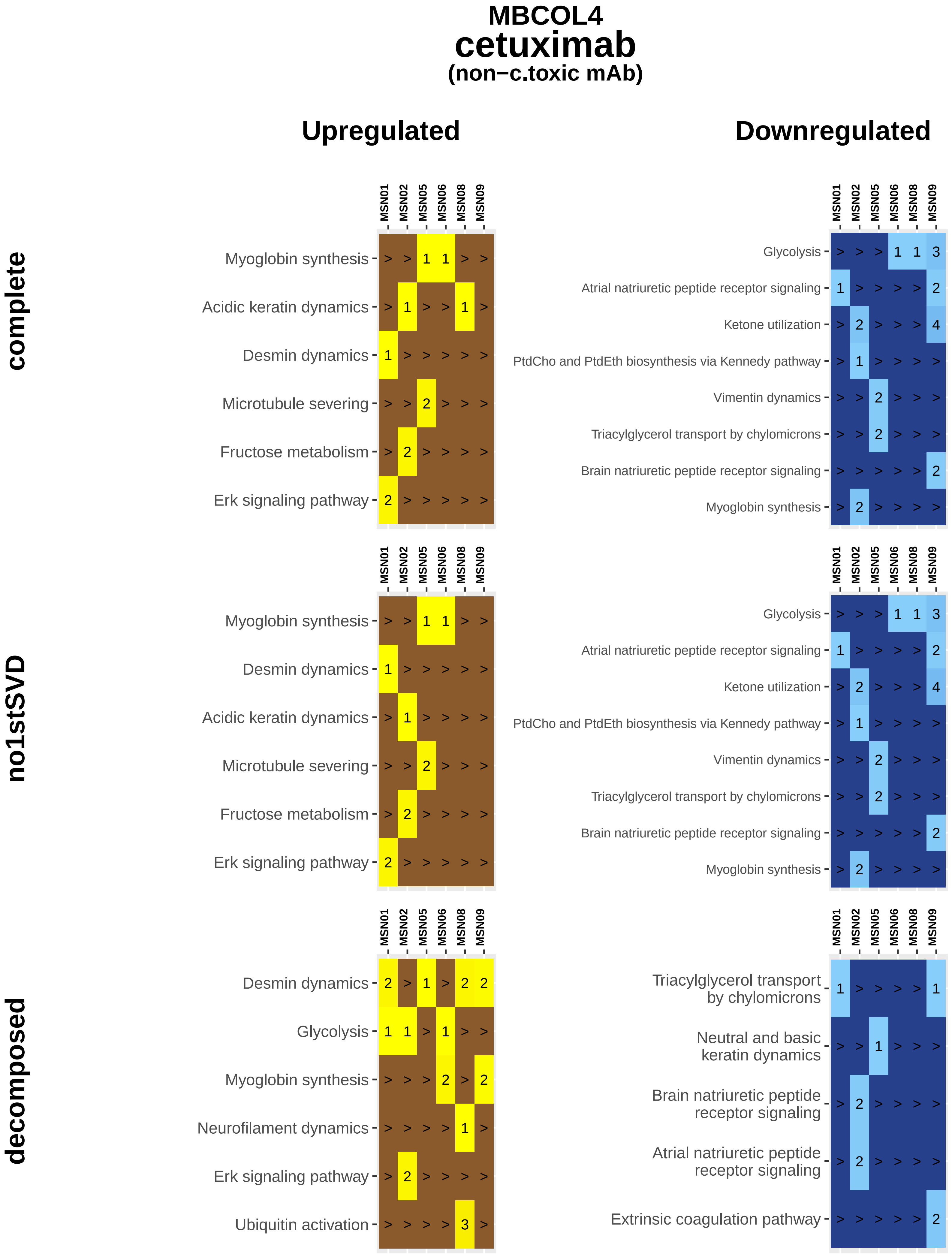

MBCOL4
cetuximab
(non−c.toxic mAb)
Upregulated
Downregulated
complete
no1stSVD
decomposed
MSN01
MSN02
MSN05
MSN06
MSN08
MSN09
Myoglobin synthesis
Acidic keratin dynamics
Desmin dynamics
Microtubule severing
Fructose metabolism
Erk signaling pathway
Glycolysis
Atrial natriuretic peptide receptor signaling
Ketone utilization
PtdCho and PtdEth biosynthesis via Kennedy pathway
Vimentin dynamics
Triacylglycerol transport by chylomicrons
Brain natriuretic peptide receptor signaling
Neurofilament dynamics
Ubiquitin activation
Neutral and basic keratin dynamics
Extrinsic coagulation pathway

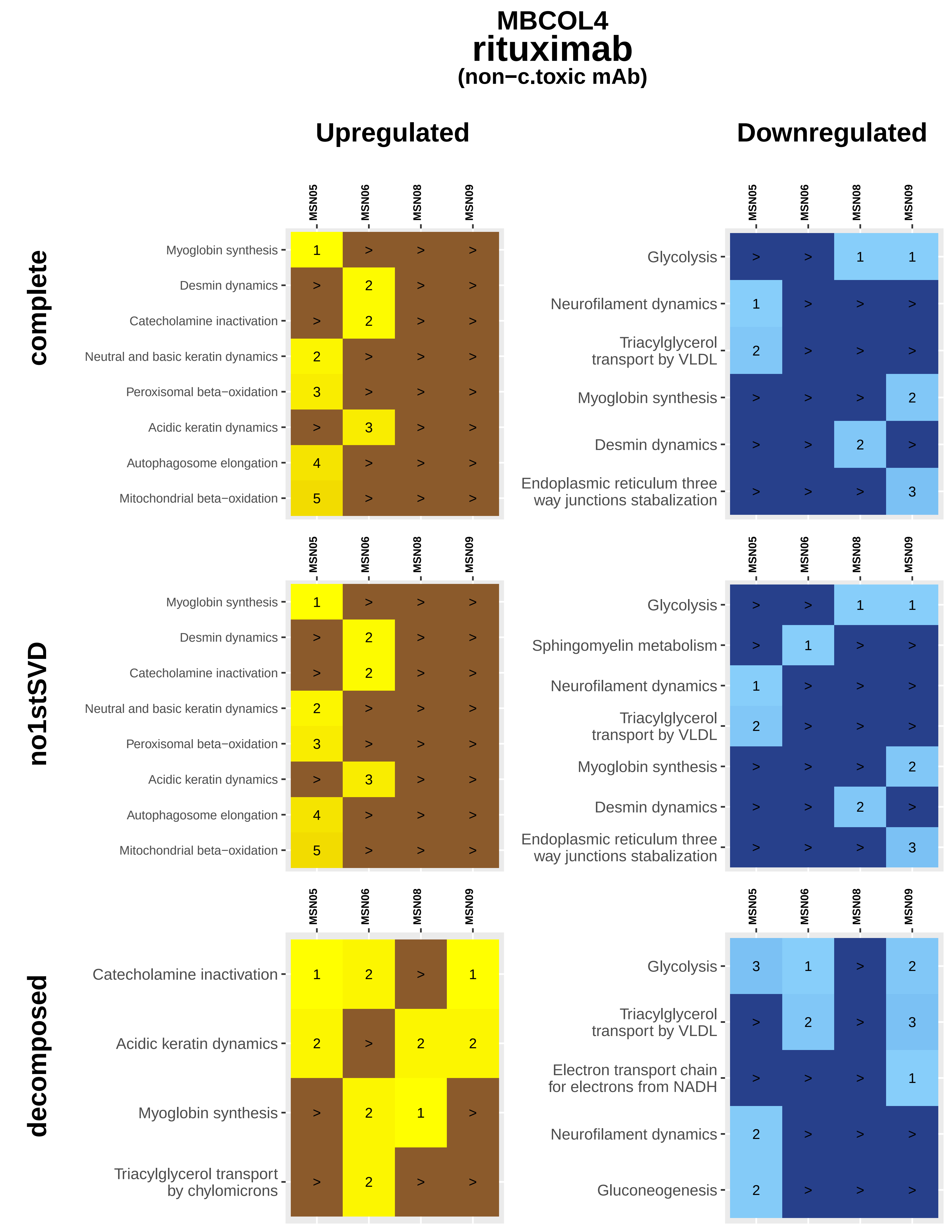
MBCOL4
rituximab
(non−c.toxic mAb)
Upregulated
Downregulated
complete
no1stSVD
decomposed
MSN05
MSN06
MSN08
MSN09
Myoglobin synthesis
Desmin dynamics
Catecholamine inactivation
Neutral and basic keratin dynamics
Peroxisomal beta−oxidation
Acidic keratin dynamics
Autophagosome elongation
Mitochondrial beta−oxidation
Glycolysis
Neurofilament dynamics
Triacylglycerol transport by VLDL
Endoplasmic reticulum three way junctions stabalization
Sphingomyelin metabolism
Triacylglycerol transport by chylomicrons
Electron transport chain for electrons from NADH
Gluconeogenesis

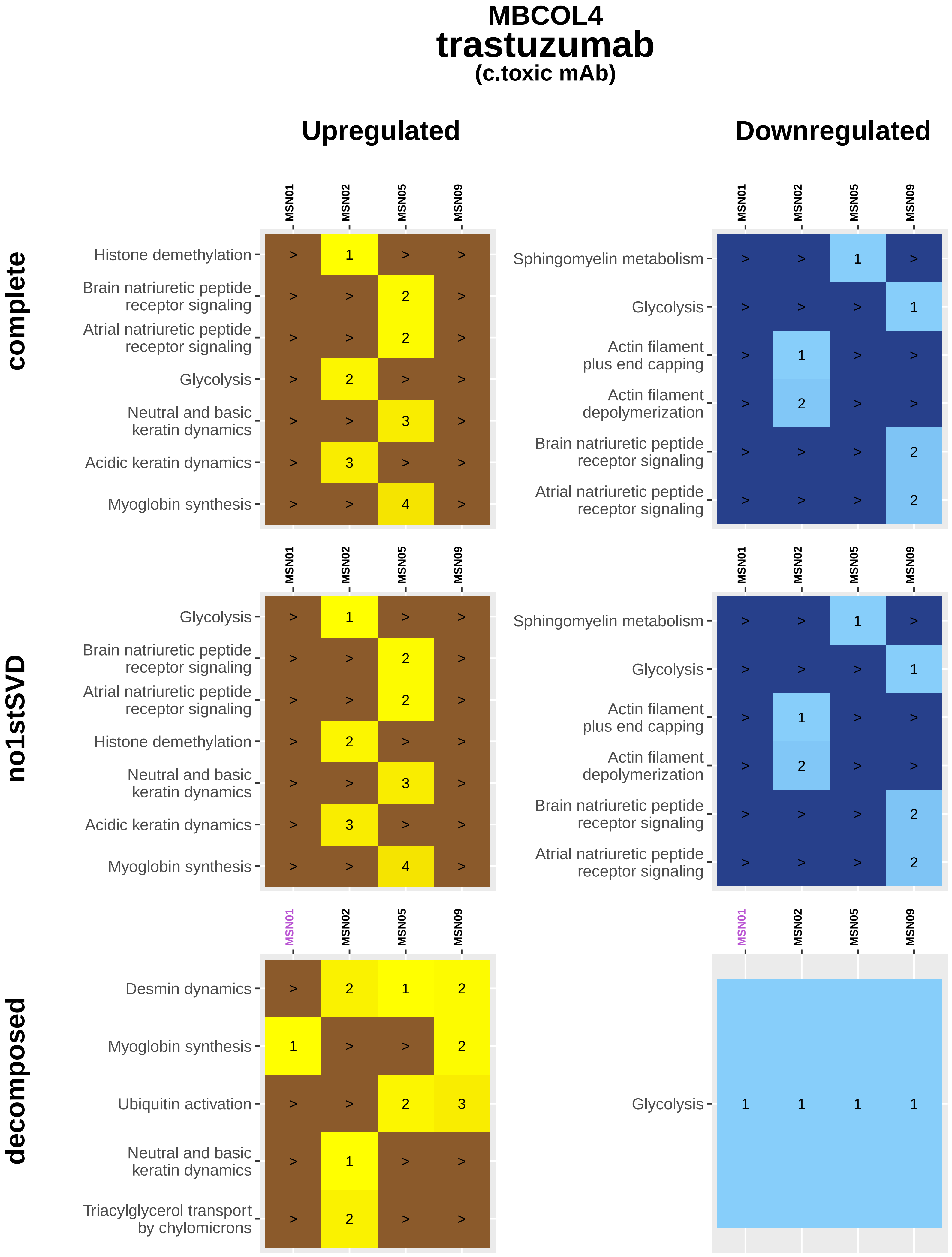
MBCOL4
trastuzumab
(c.toxic mAb)
Upregulated
Downregulated
complete
no1stSVD
decomposed
MSN01
MSN02
MSN05
MSN09
Histone demethylation
Brain natriuretic peptide receptor signaling
Atrial natriuretic peptide receptor signaling
Glycolysis
Neutral and basic keratin dynamics
Acidic keratin dynamics
Myoglobin synthesis
Sphingomyelin metabolism
Actin filament plus end capping
Actin filament depolymerization
Desmin dynamics
Ubiquitin activation
Triacylglycerol transport by chylomicrons

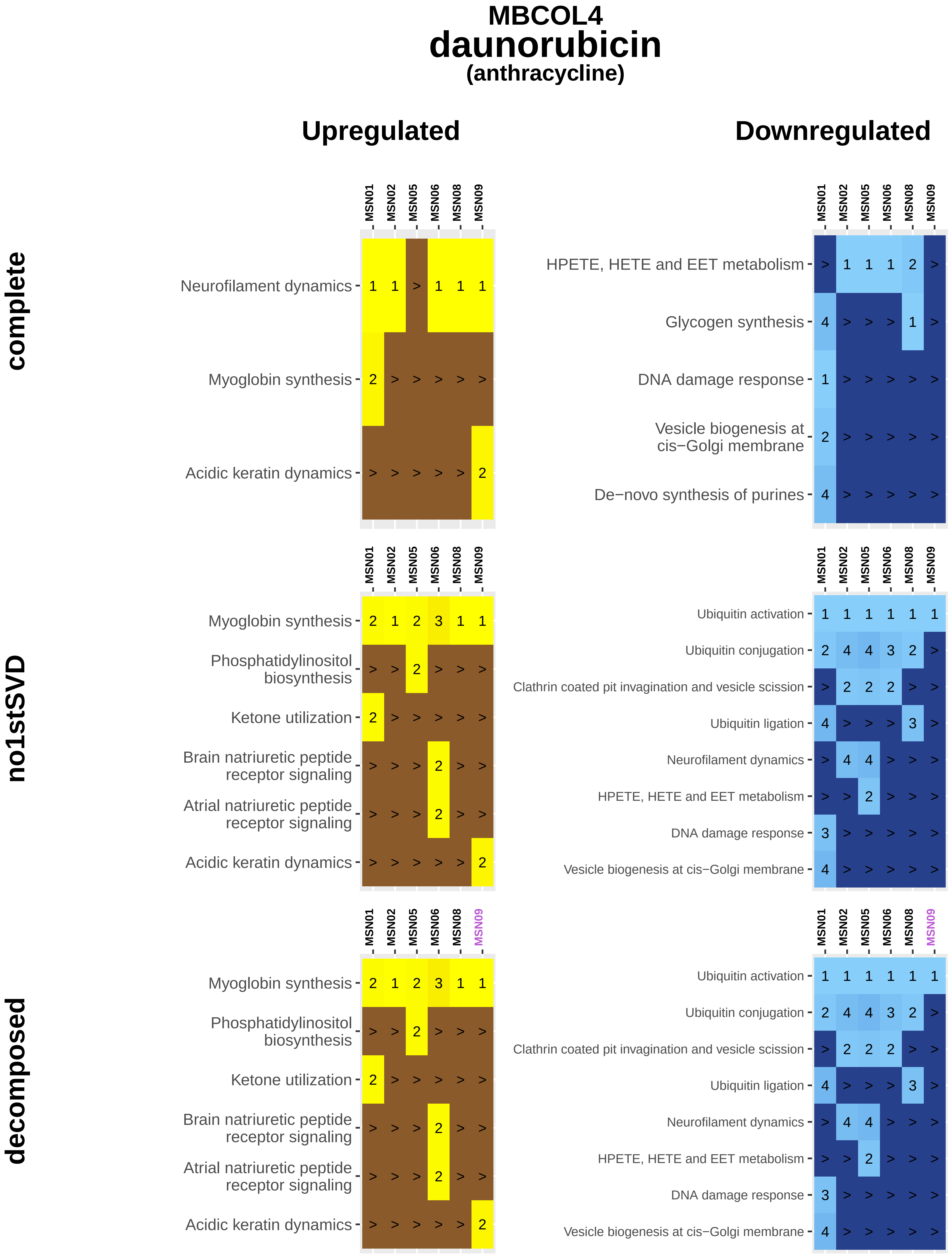
MBCOL4
daunorubicin
(anthracycline)
Upregulated
Downregulated
complete
no1stSVD
decomposed
MSN01
MSN02
MSN05
MSN06
MSN08
MSN09
Neurofilament dynamics
Myoglobin synthesis
Acidic keratin dynamics
HPETE, HETE and EET metabolism
Glycogen synthesis
DNA damage response
Vesicle biogenesis at cis−Golgi membrane
De−novo synthesis of purines
Phosphatidylinositol biosynthesis
Ketone utilization
Brain natriuretic peptide receptor signaling
Atrial natriuretic peptide receptor signaling
Ubiquitin activation
Ubiquitin conjugation
Clathrin coated pit invagination and vesicle scission
Ubiquitin ligation
Neurofilament dynamics

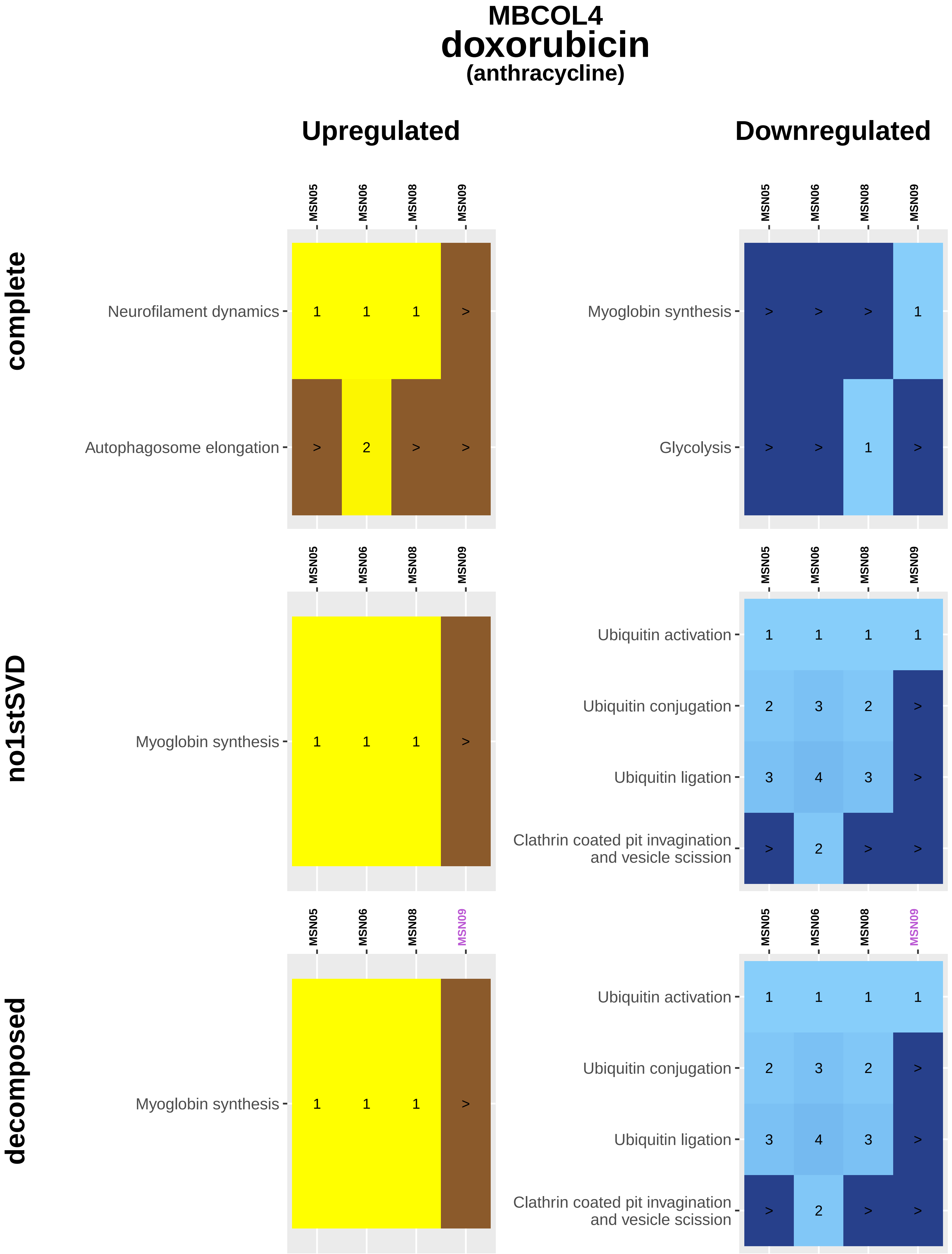

MBCOL4
doxorubicin
(anthracycline)
Upregulated
Downregulated
complete
no1stSVD
decomposed
MSN05
MSN06
MSN08
MSN09
Neurofilament dynamics
Autophagosome elongation
Myoglobin synthesis
Glycolysis
Ubiquitin activation
Ubiquitin conjugation
Ubiquitin ligation
Clathrin coated pit invagination and vesicle scission

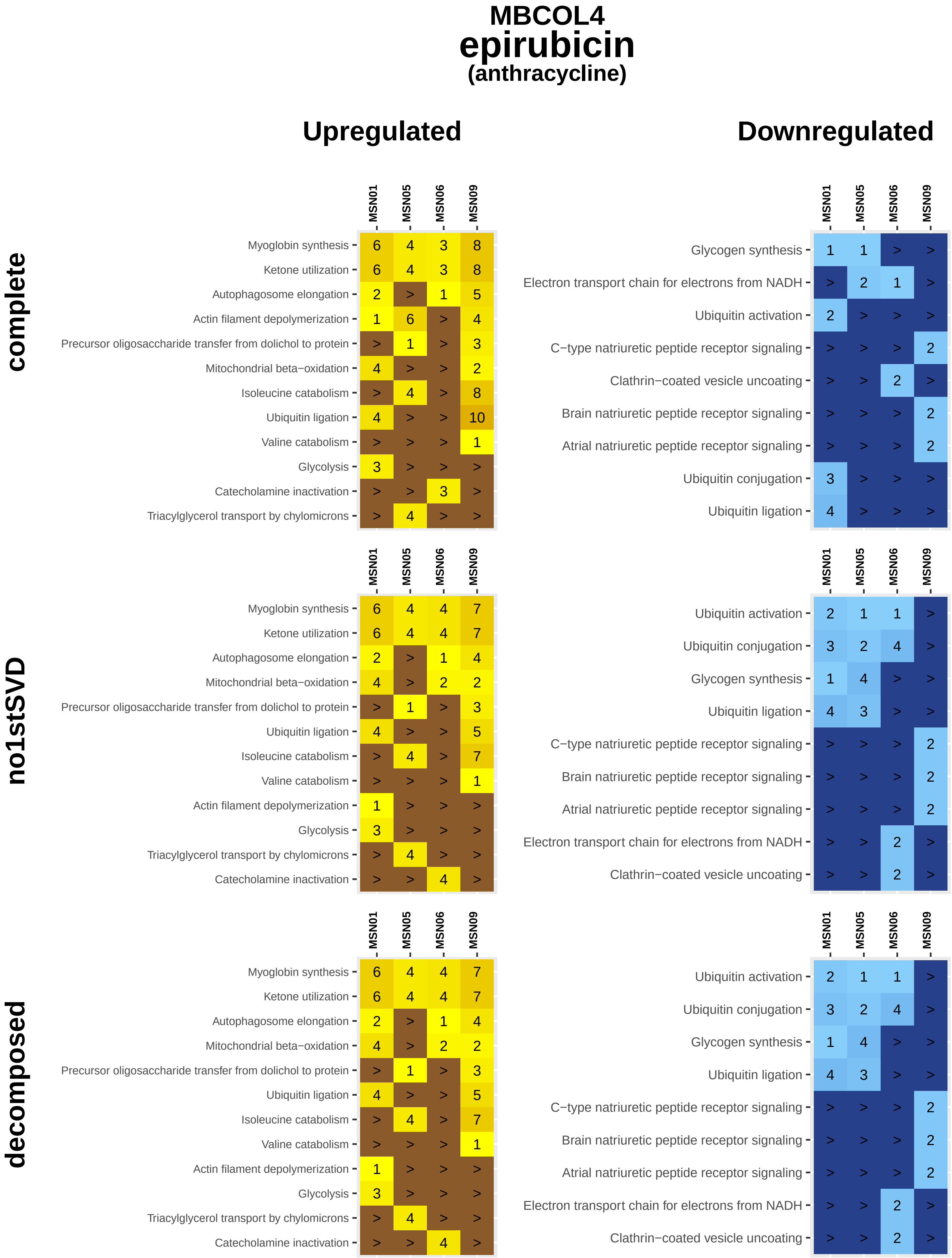
MBCOL4
epirubicin
(anthracycline)
Upregulated
Downregulated
complete
MSN01 MSN05 MSN06 MSN09
Myoglobin synthesis 6 4 3 8
Ketone utilization 6 4 3 8
Autophagosome elongation 2 > 1 5
Actin filament depolymerization 1 6 > 4
Precursor oligosaccharide transfer from dolichol to protein > 1 > 3
Mitochondrial beta−oxidation 4 > > 2
Isoleucine catabolism > 4 > 8
Ubiquitin ligation 4 > > 10
Valine catabolism > > > 1
Glycolysis 3 > > >
Catecholamine inactivation > > 3 >
Triacylglycerol transport by chylomicrons > 4 > >
MSN01 MSN05 MSN06 MSN09
Glycogen synthesis 1 1 > >
Electron transport chain for electrons from NADH > 2 1 >
Ubiquitin activation 2 > > >
C−type natriuretic peptide receptor signaling > > > 2
Clathrin−coated vesicle uncoating > > 2 >
Brain natriuretic peptide receptor signaling > > > 2
Atrial natriuretic peptide receptor signaling > > > 2
Ubiquitin conjugation 3 > > >
Ubiquitin ligation 4 > > >
no1stSVD
MSN01 MSN05 MSN06 MSN09
Myoglobin synthesis 6 4 4 7
Ketone utilization 6 4 4 7
Autophagosome elongation 2 > 1 4
Mitochondrial beta−oxidation 4 > 2 2
Precursor oligosaccharide transfer from dolichol to protein > 1 > 3
Ubiquitin ligation 4 > > 5
Isoleucine catabolism > 4 > 7
Valine catabolism > > > 1
Actin filament depolymerization 1 > > >
Glycolysis 3 > > >
Triacylglycerol transport by chylomicrons > 4 > >
Catecholamine inactivation > > 4 >
MSN01 MSN05 MSN06 MSN09
Ubiquitin activation 2 1 1 >
Ubiquitin conjugation 3 2 4 >
Glycogen synthesis 1 4 > >
Ubiquitin ligation 4 3 > >
C−type natriuretic peptide receptor signaling > > > 2
Brain natriuretic peptide receptor signaling > > > 2
Atrial natriuretic peptide receptor signaling > > > 2
Electron transport chain for electrons from NADH > > 2 >
Clathrin−coated vesicle uncoating > > 2 >
decomposed
MSN01 MSN05 MSN06 MSN09
Myoglobin synthesis 6 4 4 7
Ketone utilization 6 4 4 7
Autophagosome elongation 2 > 1 4
Mitochondrial beta−oxidation 4 > 2 2
Precursor oligosaccharide transfer from dolichol to protein > 1 > 3
Ubiquitin ligation 4 > > 5
Isoleucine catabolism > 4 > 7
Valine catabolism > > > 1
Actin filament depolymerization 1 > > >
Glycolysis 3 > > >
Triacylglycerol transport by chylomicrons > 4 > >
Catecholamine inactivation > > 4 >
MSN01 MSN05 MSN06 MSN09
Ubiquitin activation 2 1 1 >
Ubiquitin conjugation 3 2 4 >
Glycogen synthesis 1 4 > >
Ubiquitin ligation 4 3 > >
C−type natriuretic peptide receptor signaling > > > 2
Brain natriuretic peptide receptor signaling > > > 2
Atrial natriuretic peptide receptor signaling > > > 2
Electron transport chain for electrons from NADH > > 2 >
Clathrin−coated vesicle uncoating > > 2 >

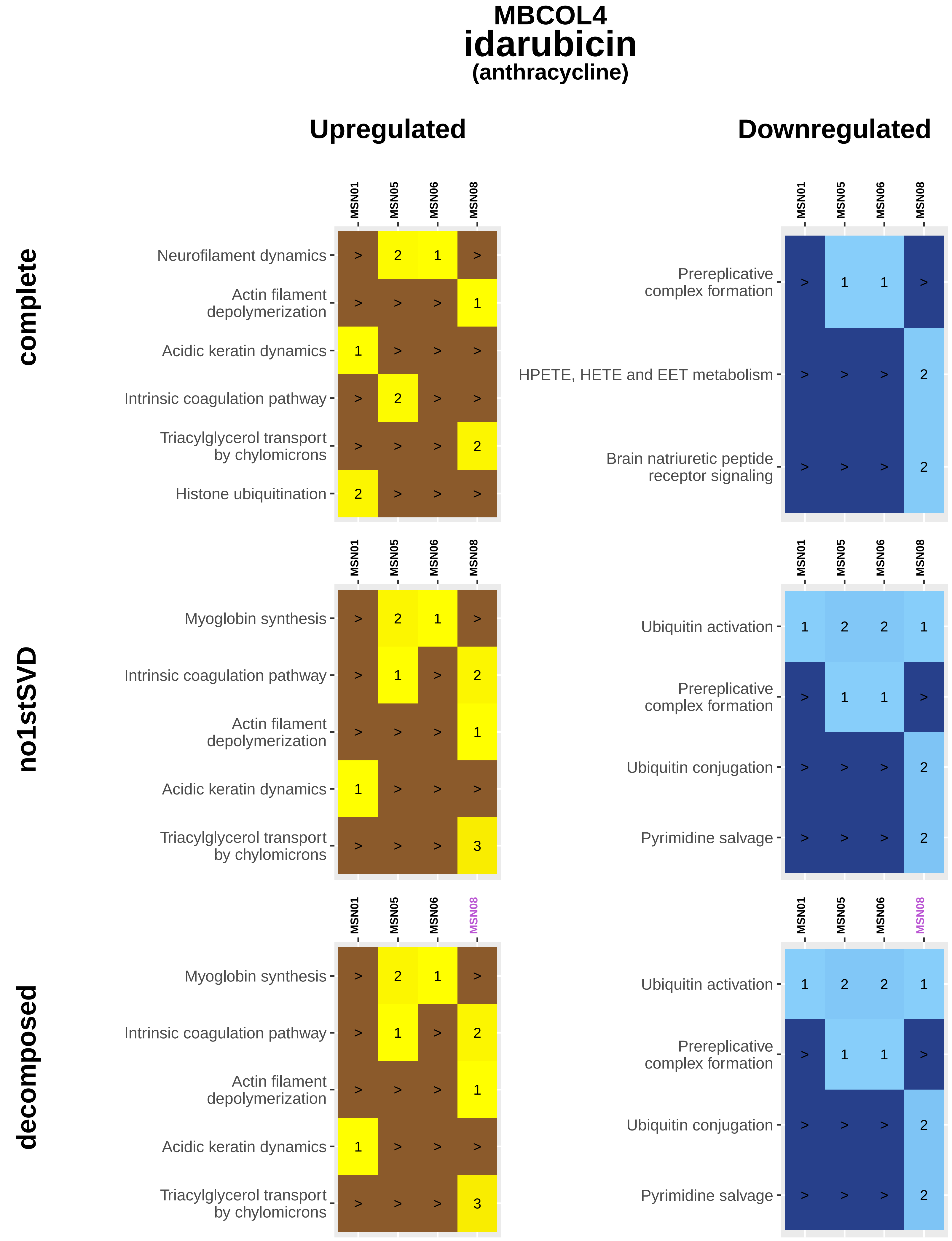
MBCOL4
idarubicin
(anthracycline)
Upregulated
Downregulated
complete
no1stSVD
decomposed
MSN01
MSN05
MSN06
MSN08
Neurofilament dynamics
Actin filament depolymerization
Acidic keratin dynamics
Intrinsic coagulation pathway
Triacylglycerol transport by chylomicrons
Histone ubiquitination
Prereplicative complex formation
HPETE, HETE and EET metabolism
Brain natriuretic peptide receptor signaling
Myoglobin synthesis
Ubiquitin activation
Ubiquitin conjugation
Pyrimidine salvage

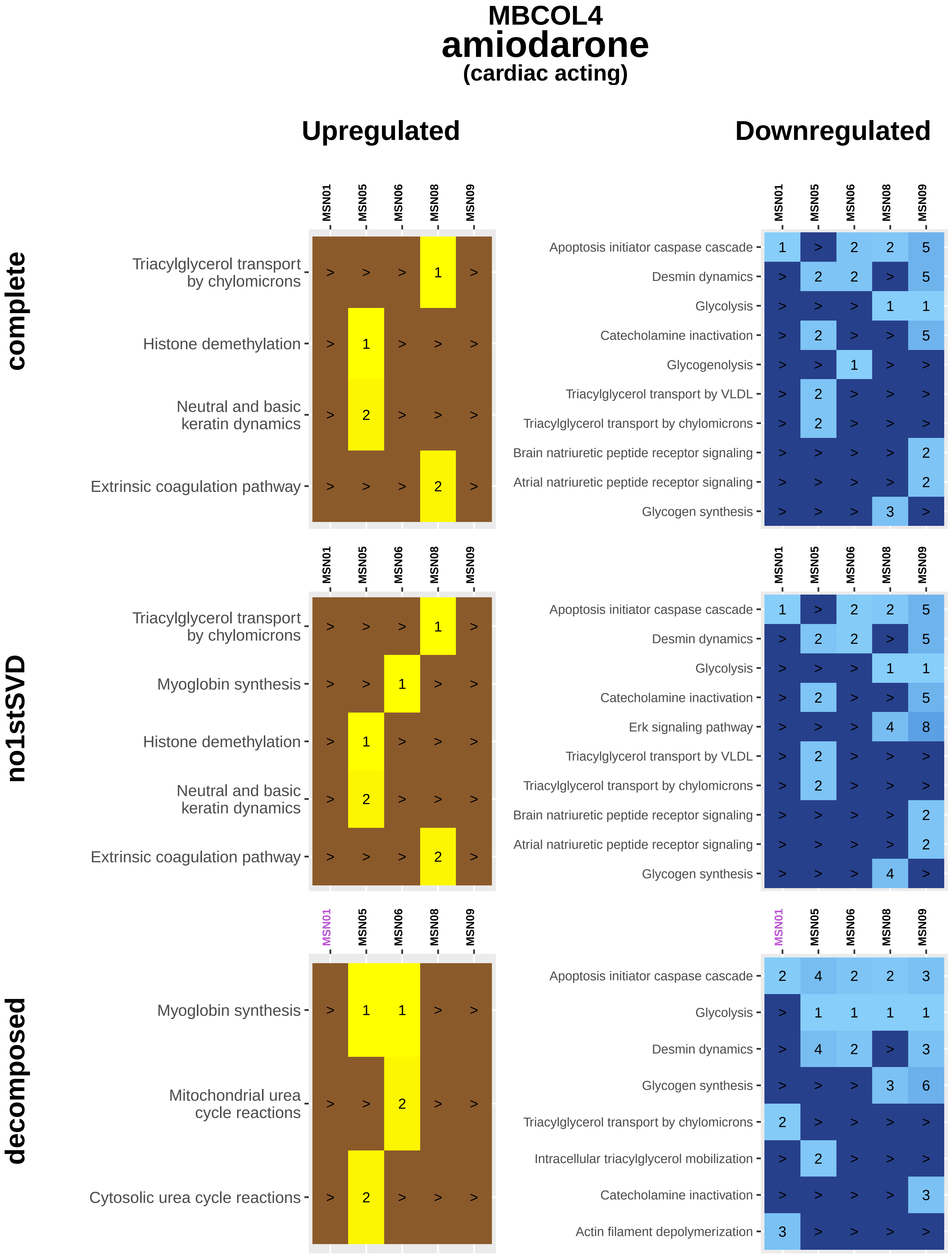

MBCOL4
amiodarone
(cardiac acting)
Upregulated
Downregulated
complete
no1stSVD
decomposed
MSN01 MSN05 MSN06 MSN08 MSN09
Triacylglycerol transport by chylomicrons > > > 1 >
Histone demethylation > 1 > > >
Neutral and basic keratin dynamics > 2 > > >
Extrinsic coagulation pathway > > > 2 >
Apoptosis initiator caspase cascade 1 > 2 2 5
Desmin dynamics > 2 2 > 5
Glycolysis > > > 1 1
Catecholamine inactivation > 2 > > 5
Glycogenolysis > > 1 > >
Triacylglycerol transport by VLDL > 2 > > >
Triacylglycerol transport by chylomicrons > 2 > > >
Brain natriuretic peptide receptor signaling > > > > 2
Atrial natriuretic peptide receptor signaling > > > > 2
Glycogen synthesis > > > 3 >
Triacylglycerol transport by chylomicrons > > > 1 >
Myoglobin synthesis > > 1 > >
Histone demethylation > 1 > > >
Neutral and basic keratin dynamics > 2 > > >
Extrinsic coagulation pathway > > > 2 >
Apoptosis initiator caspase cascade 1 > 2 2 5
Desmin dynamics > 2 2 > 5
Glycolysis > > > 1 1
Catecholamine inactivation > 2 > > 5
Erk signaling pathway > > > 4 8
Triacylglycerol transport by VLDL > 2 > > >
Triacylglycerol transport by chylomicrons > 2 > > >
Brain natriuretic peptide receptor signaling > > > > 2
Atrial natriuretic peptide receptor signaling > > > > 2
Glycogen synthesis > > > 4 >
Myoglobin synthesis > 1 1 > >
Mitochondrial urea cycle reactions > > 2 > >
Cytosolic urea cycle reactions > 2 > > >
Apoptosis initiator caspase cascade 2 4 2 2 3
Glycolysis > 1 1 1 1
Desmin dynamics > 4 2 > 3
Glycogen synthesis > > > 3 6
Triacylglycerol transport by chylomicrons 2 > > > >
Intracellular triacylglycerol mobilization > 2 > > >
Catecholamine inactivation > > > > 3
Actin filament depolymerization 3 > > > >

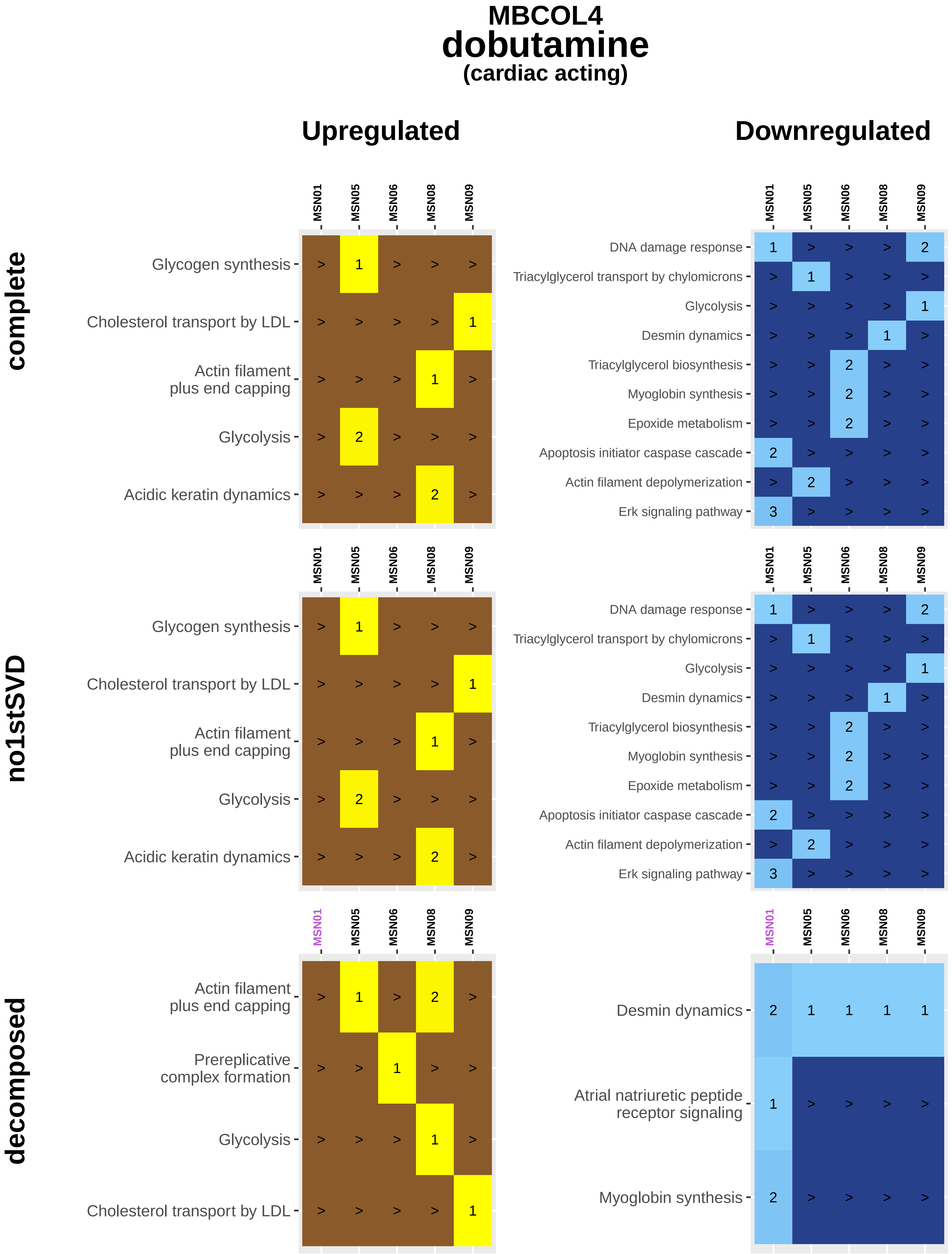

MBCOL4
dobutamine
(cardiac acting)
Upregulated
Downregulated
complete
no1stSVD
decomposed
MSN01
MSN05
MSN06
MSN08
MSN09
Glycogen synthesis
Cholesterol transport by LDL
Actin filament plus end capping
Glycolysis
Acidic keratin dynamics
DNA damage response
Triacylglycerol transport by chylomicrons
Desmin dynamics
Triacylglycerol biosynthesis
Myoglobin synthesis
Epoxide metabolism
Apoptosis initiator caspase cascade
Actin filament depolymerization
Erk signaling pathway
Prereplicative complex formation
Atrial natriuretic peptide receptor signaling

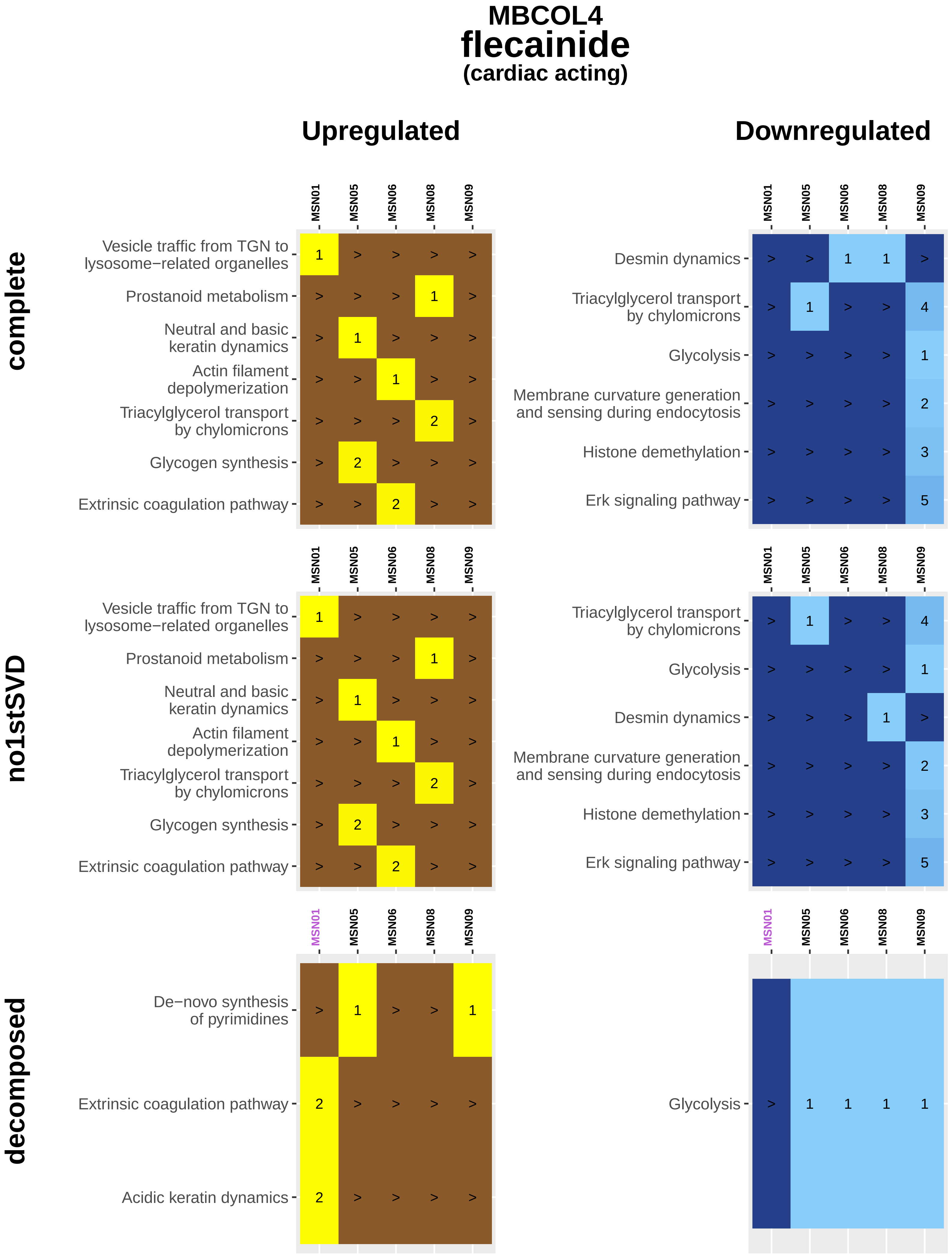
MBCOL4
flecainide
(cardiac acting)
Upregulated
Downregulated
complete
no1stSVD
decomposed
MSN01
MSN05
MSN06
MSN08
MSN09
Vesicle traffic from TGN to lysosome−related organelles
Prostanoid metabolism
Neutral and basic keratin dynamics
Actin filament depolymerization
Triacylglycerol transport by chylomicrons
Glycogen synthesis
Extrinsic coagulation pathway
Desmin dynamics
Glycolysis
Membrane curvature generation and sensing during endocytosis
Histone demethylation
Erk signaling pathway
De−novo synthesis of pyrimidines
Acidic keratin dynamics

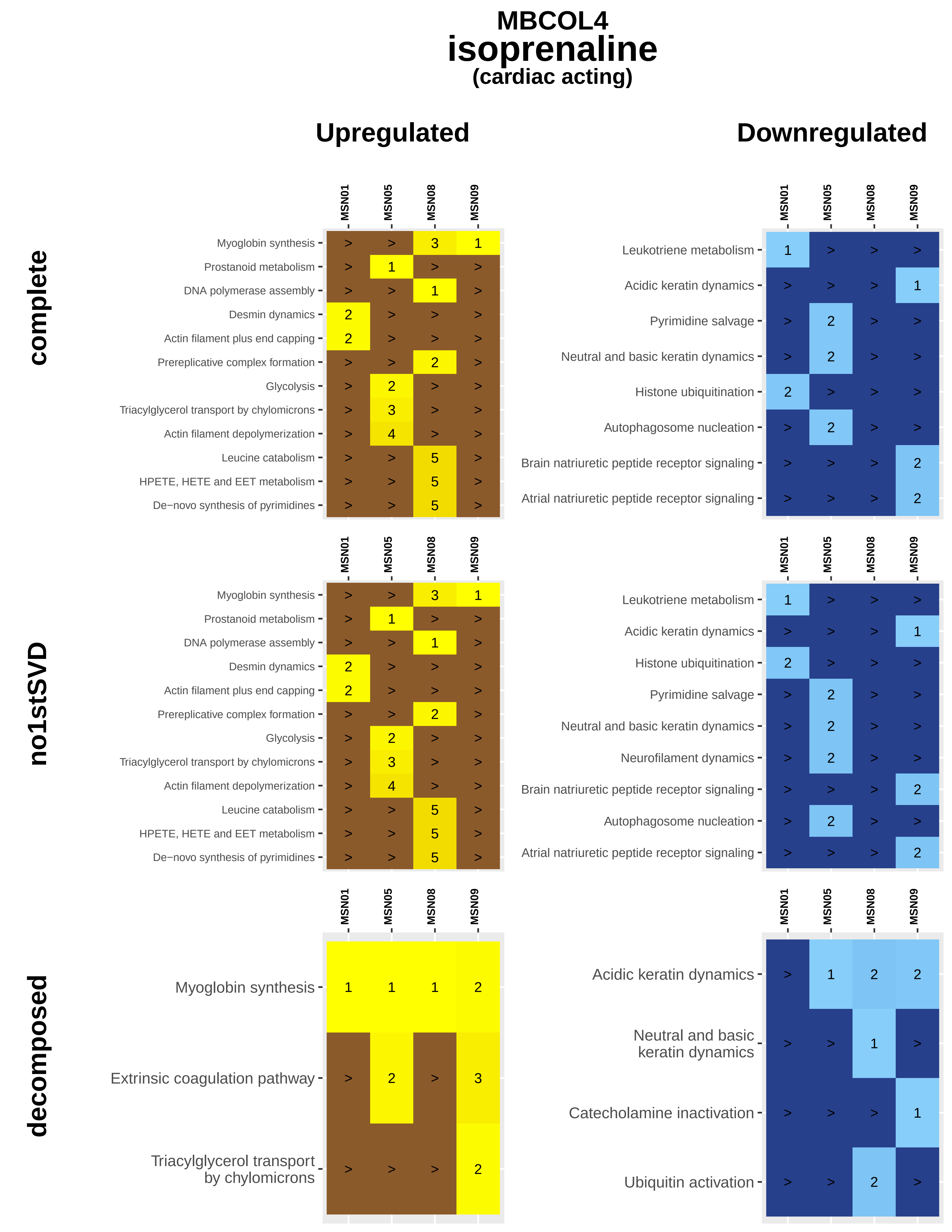

MBCOL4
isoprenaline
(cardiac acting)
Upregulated
Downregulated
complete
no1stSVD
decomposed
MSN01
MSN05
MSN08
MSN09
Myoglobin synthesis
Prostanoid metabolism
DNA polymerase assembly
Desmin dynamics
Actin filament plus end capping
Prereplicative complex formation
Glycolysis
Triacylglycerol transport by chylomicrons
Actin filament depolymerization
Leucine catabolism
HPETE, HETE and EET metabolism
De−novo synthesis of pyrimidines
Leukotriene metabolism
Acidic keratin dynamics
Pyrimidine salvage
Neutral and basic keratin dynamics
Histone ubiquitination
Autophagosome nucleation
Brain natriuretic peptide receptor signaling
Atrial natriuretic peptide receptor signaling
Neurofilament dynamics
Extrinsic coagulation pathway
Catecholamine inactivation
Ubiquitin activation

**MBCOL4**

### milrinone

**(cardiac acting)**

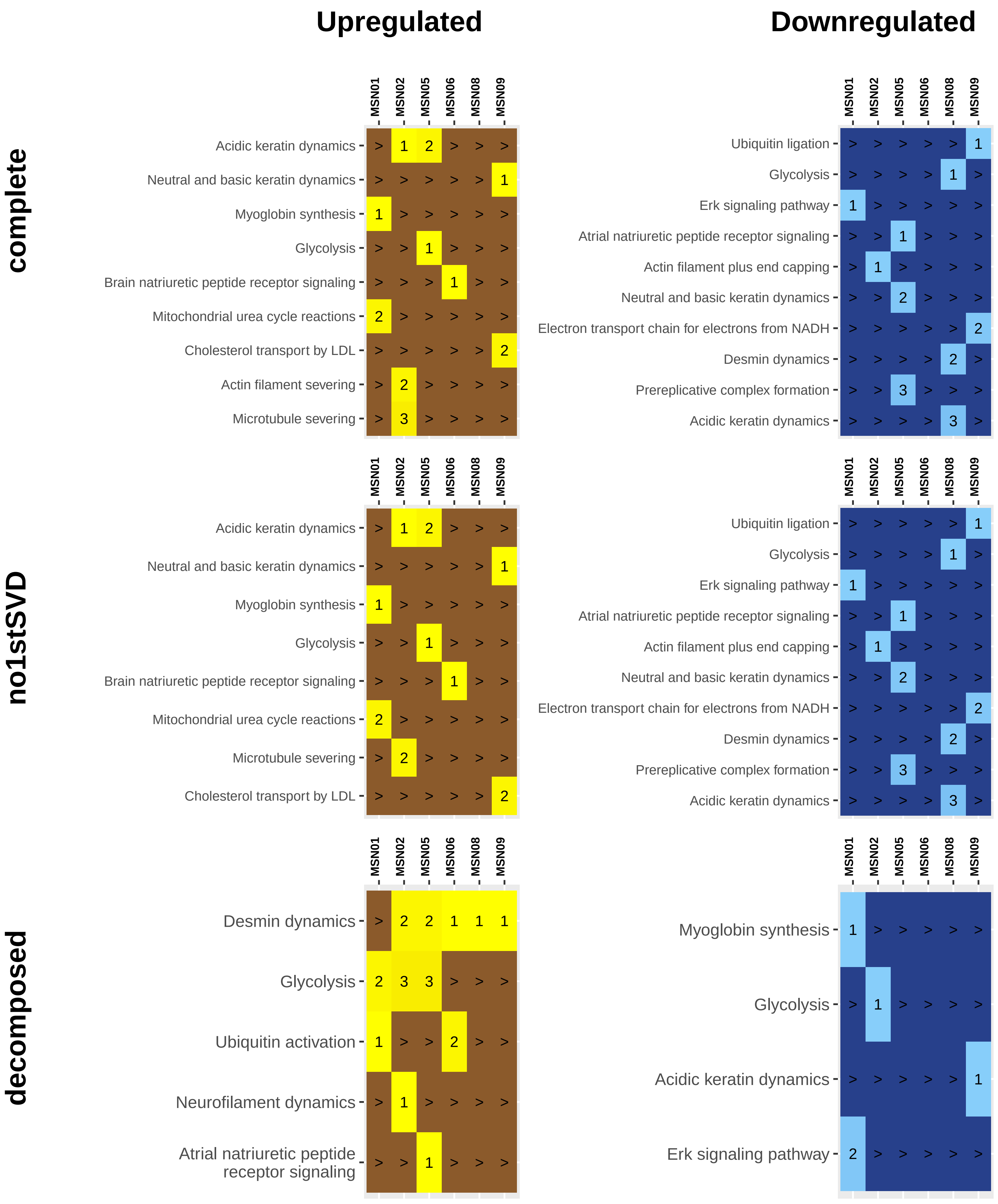

### MBCOL4
### phenylephrine
#### (cardiac acting)

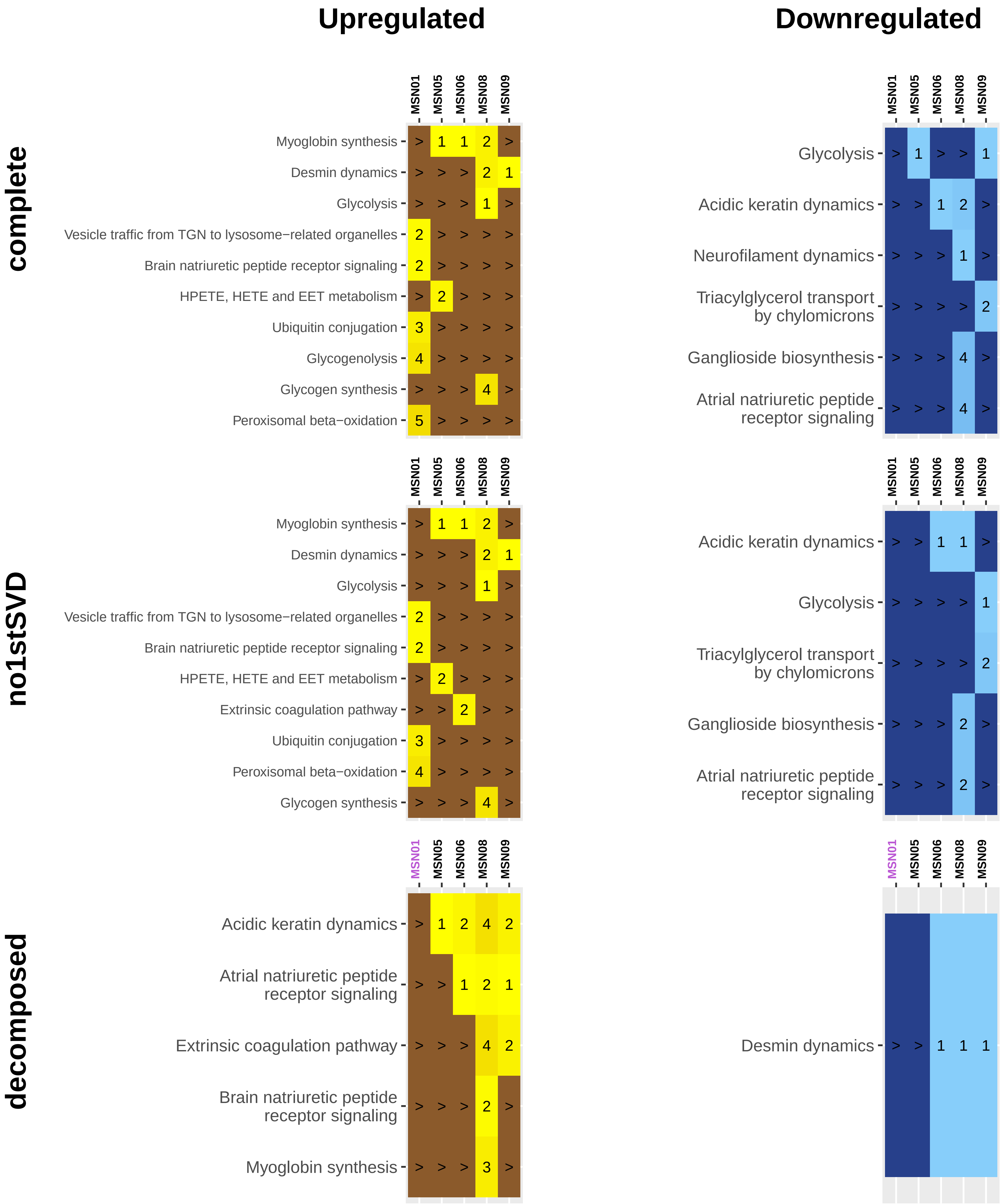

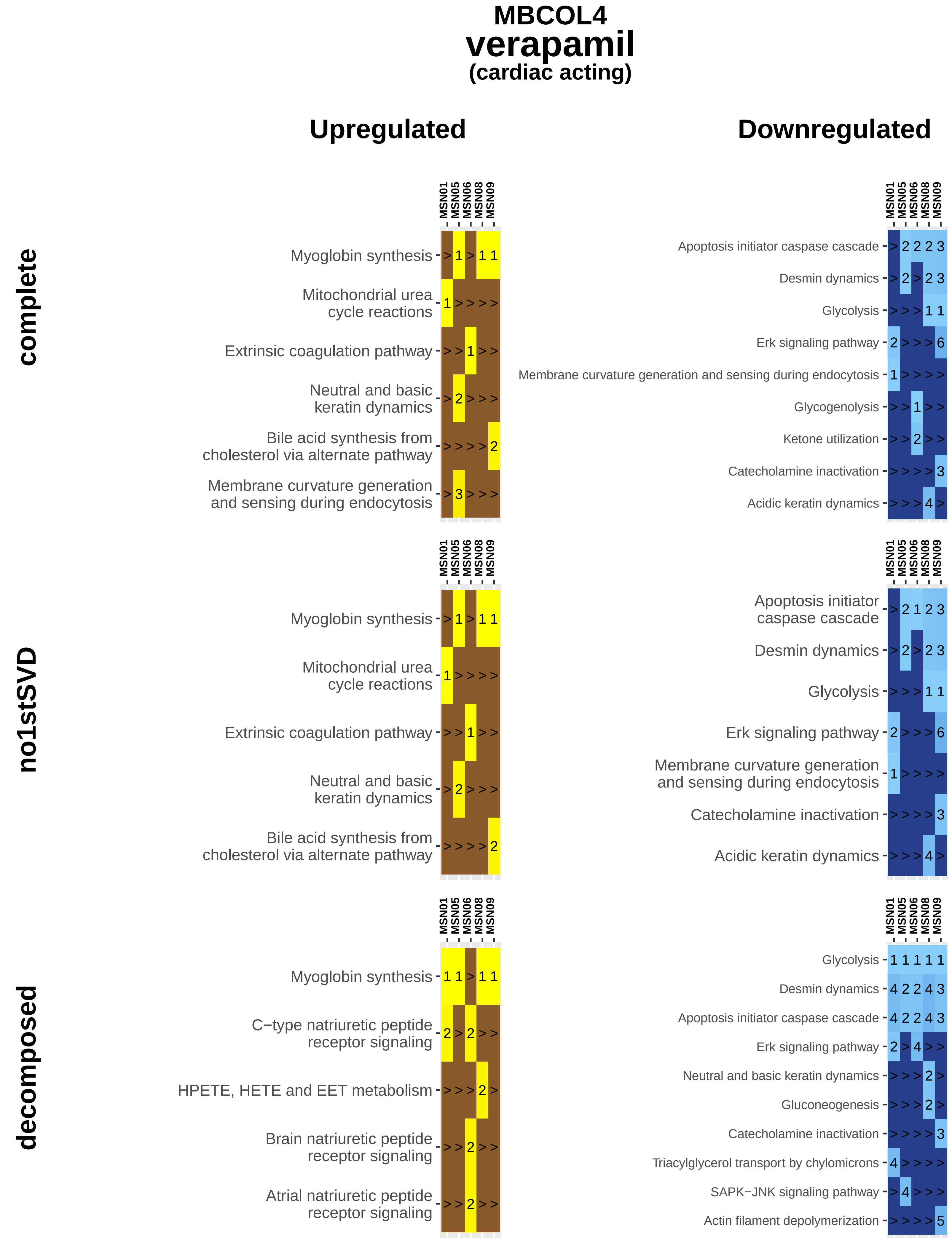

MBCOL4
verapamil
(cardiac acting)
Upregulated
Downregulated
complete
no1stSVD
decomposed
MSN01
MSN05
MSN06
MSN08
MSN09
Myoglobin synthesis
Mitochondrial urea cycle reactions
Extrinsic coagulation pathway
Neutral and basic keratin dynamics
Bile acid synthesis from cholesterol via alternate pathway
Membrane curvature generation and sensing during endocytosis
Apoptosis initiator caspase cascade
Desmin dynamics
Glycolysis
Erk signaling pathway
Glycogenolysis
Ketone utilization
Catecholamine inactivation
Acidic keratin dynamics
C−type natriuretic peptide receptor signaling
HPETE, HETE and EET metabolism
Brain natriuretic peptide receptor signaling
Atrial natriuretic peptide receptor signaling
Gluconeogenesis
Triacylglycerol transport by chylomicrons
SAPK−JNK signaling pathway
Actin filament depolymerization

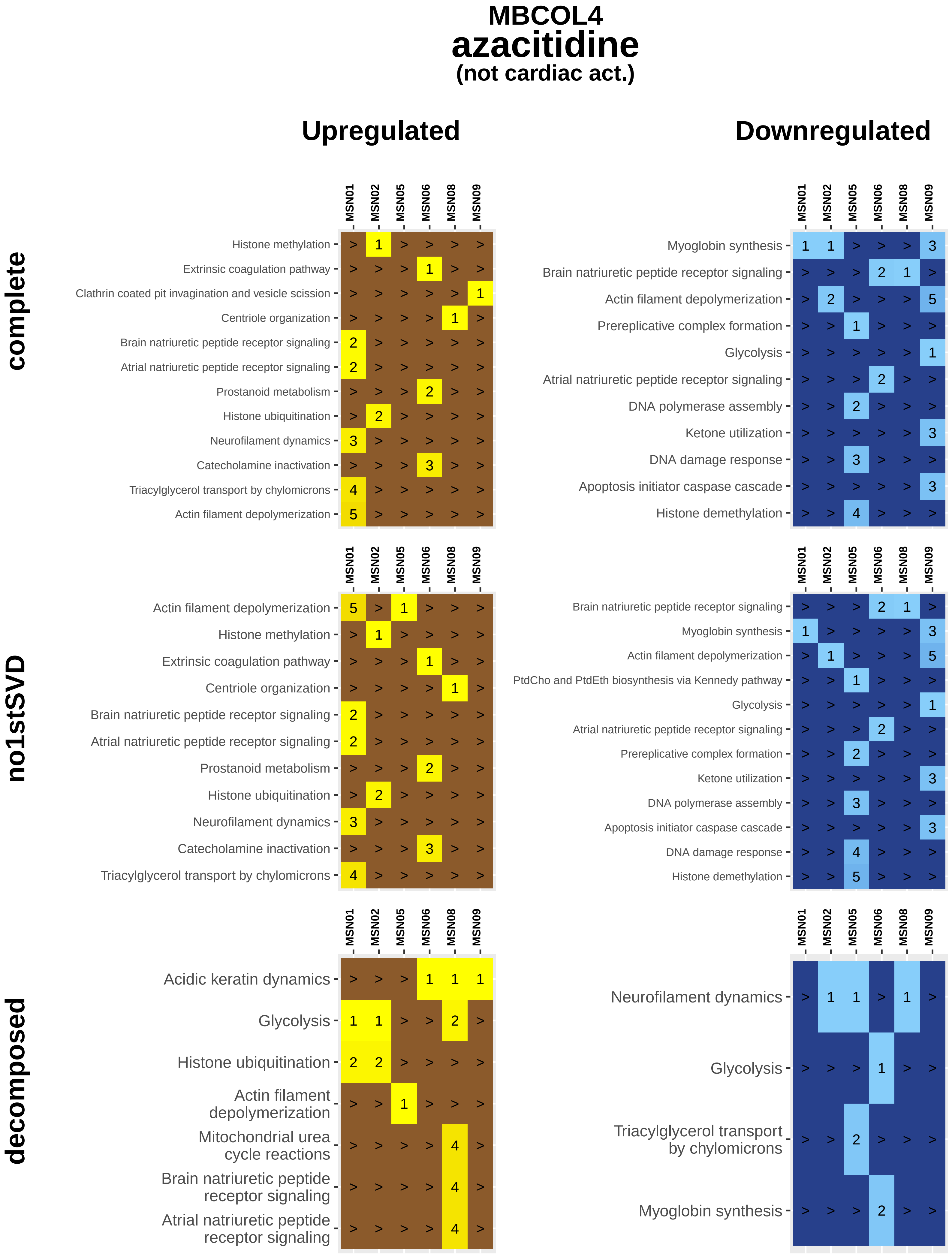
MBCOL4
azacitidine
(not cardiac act.)
Upregulated
Downregulated
complete
no1stSVD
decomposed
MSN01
MSN02
MSN05
MSN06
MSN08
MSN09
Histone methylation
Extrinsic coagulation pathway
Clathrin coated pit invagination and vesicle scission
Centriole organization
Brain natriuretic peptide receptor signaling
Atrial natriuretic peptide receptor signaling
Prostanoid metabolism
Histone ubiquitination
Neurofilament dynamics
Catecholamine inactivation
Triacylglycerol transport by chylomicrons
Actin filament depolymerization
Myoglobin synthesis
Prereplicative complex formation
Glycolysis
DNA polymerase assembly
Ketone utilization
DNA damage response
Apoptosis initiator caspase cascade
Histone demethylation
PtdCho and PtdEth biosynthesis via Kennedy pathway
Acidic keratin dynamics
Mitochondrial urea cycle reactions

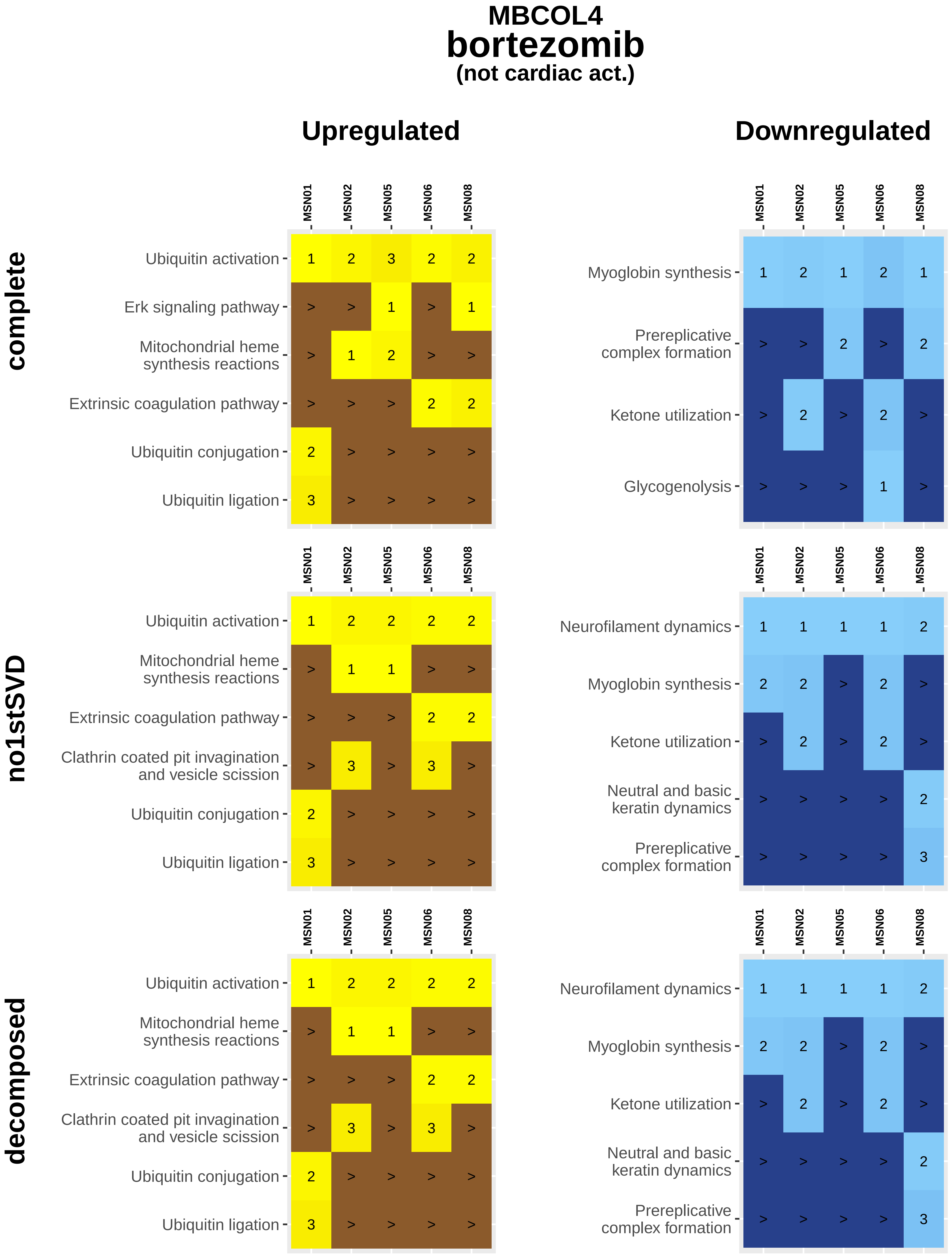
MBCOL4
bortezomib
(not cardiac act.)
Upregulated
Downregulated
complete
no1stSVD
decomposed
MSN01
MSN02
MSN05
MSN06
MSN08
Ubiquitin activation
Erk signaling pathway
Mitochondrial heme synthesis reactions
Extrinsic coagulation pathway
Ubiquitin conjugation
Ubiquitin ligation
Clathrin coated pit invagination and vesicle scission
Myoglobin synthesis
Prereplicative complex formation
Ketone utilization
Glycogenolysis
Neurofilament dynamics
Neutral and basic keratin dynamics

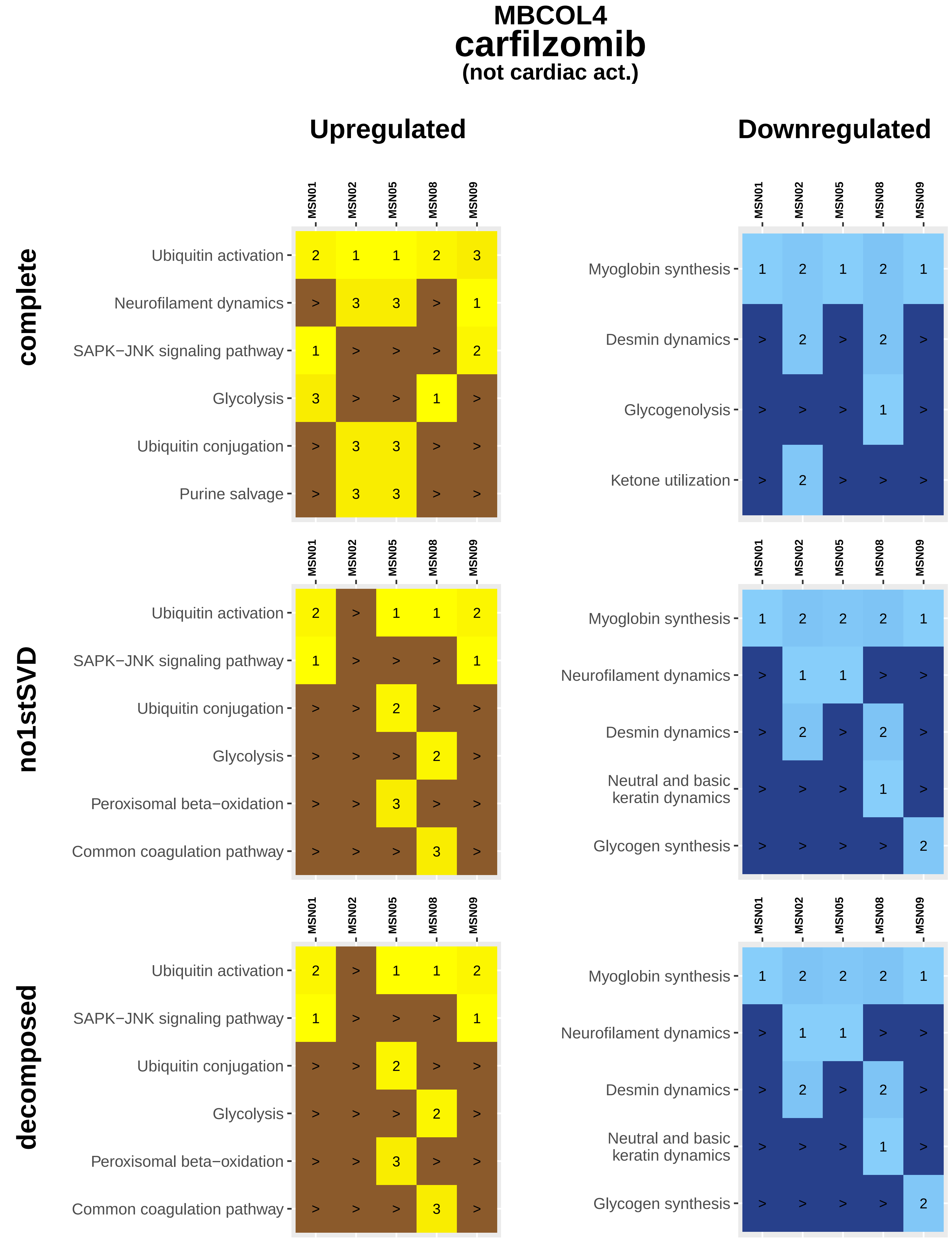
MBCOL4
carfilzomib
(not cardiac act.)
Upregulated
Downregulated
complete
no1stSVD
decomposed
MSN01
MSN02
MSN05
MSN08
MSN09
Ubiquitin activation
Neurofilament dynamics
SAPK−JNK signaling pathway
Glycolysis
Ubiquitin conjugation
Purine salvage
Myoglobin synthesis
Desmin dynamics
Glycogenolysis
Ketone utilization
Peroxisomal beta−oxidation
Common coagulation pathway
Neutral and basic keratin dynamics
Glycogen synthesis

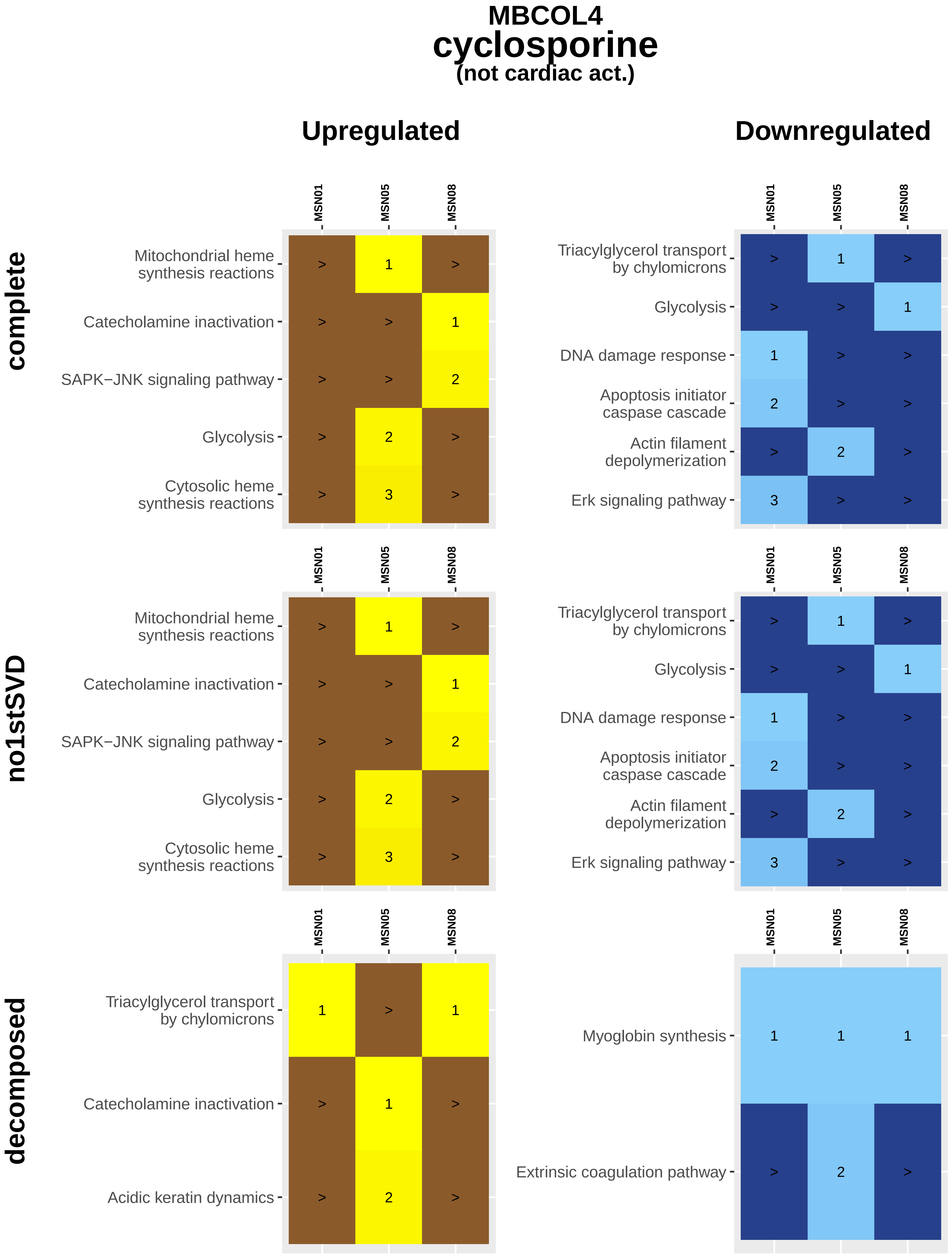

MBCOL4
cyclosporine
(not cardiac act.)
Upregulated
Downregulated
complete
no1stSVD
decomposed
MSN01
MSN05
MSN08
Mitochondrial heme synthesis reactions
Catecholamine inactivation
SAPK−JNK signaling pathway
Glycolysis
Cytosolic heme synthesis reactions
Triacylglycerol transport by chylomicrons
DNA damage response
Apoptosis initiator caspase cascade
Actin filament depolymerization
Erk signaling pathway
Acidic keratin dynamics
Myoglobin synthesis
Extrinsic coagulation pathway

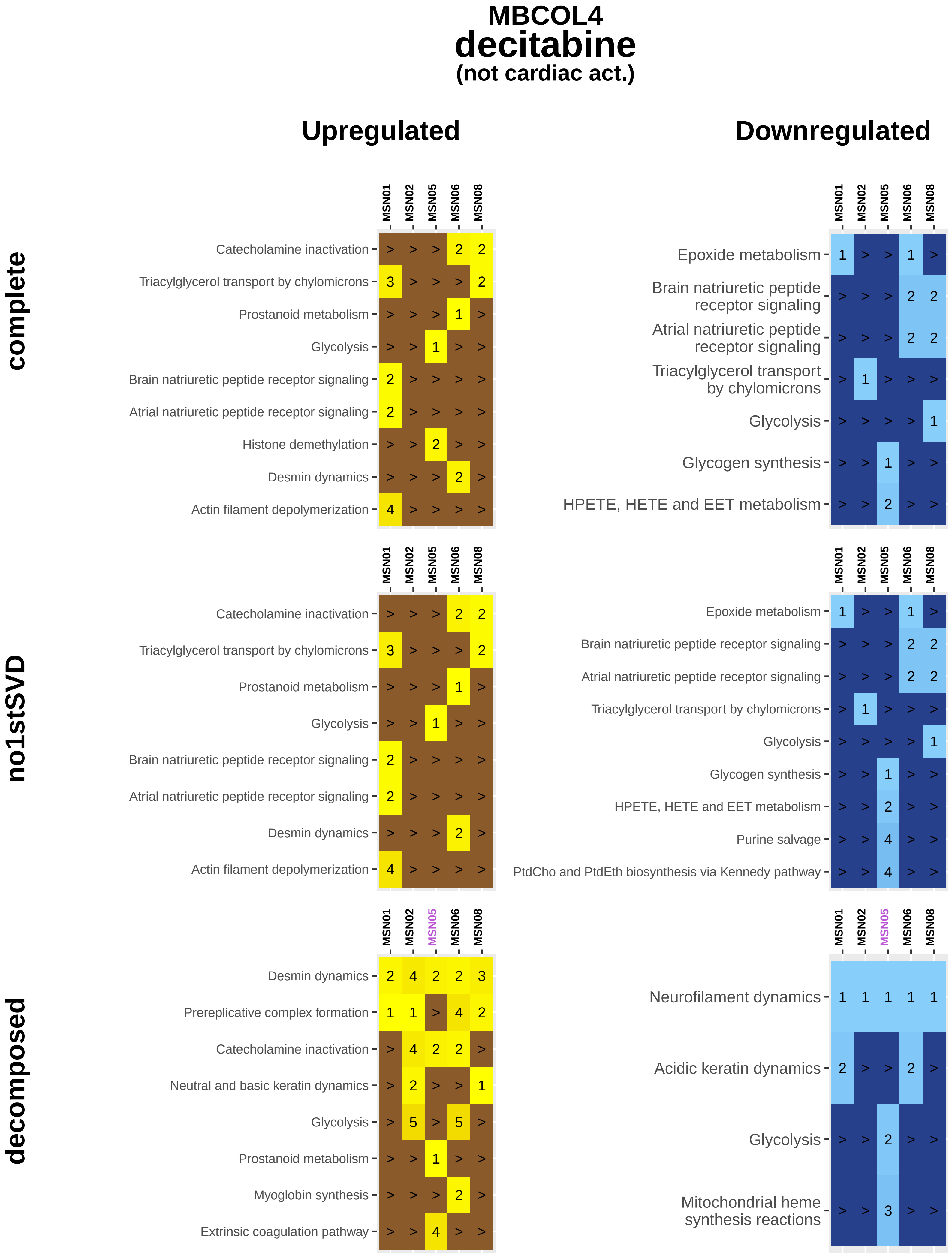
MBCOL4
decitabine
(not cardiac act.)
Upregulated
Downregulated
complete
no1stSVD
decomposed
MSN01
MSN02
MSN05
MSN06
MSN08
Catecholamine inactivation
Triacylglycerol transport by chylomicrons
Prostanoid metabolism
Glycolysis
Brain natriuretic peptide receptor signaling
Atrial natriuretic peptide receptor signaling
Histone demethylation
Desmin dynamics
Actin filament depolymerization
Epoxide metabolism
Glycogen synthesis
HPETE, HETE and EET metabolism
Purine salvage
PtdCho and PtdEth biosynthesis via Kennedy pathway
Prereplicative complex formation
Neutral and basic keratin dynamics
Myoglobin synthesis
Extrinsic coagulation pathway
Neurofilament dynamics
Acidic keratin dynamics
Mitochondrial heme synthesis reactions

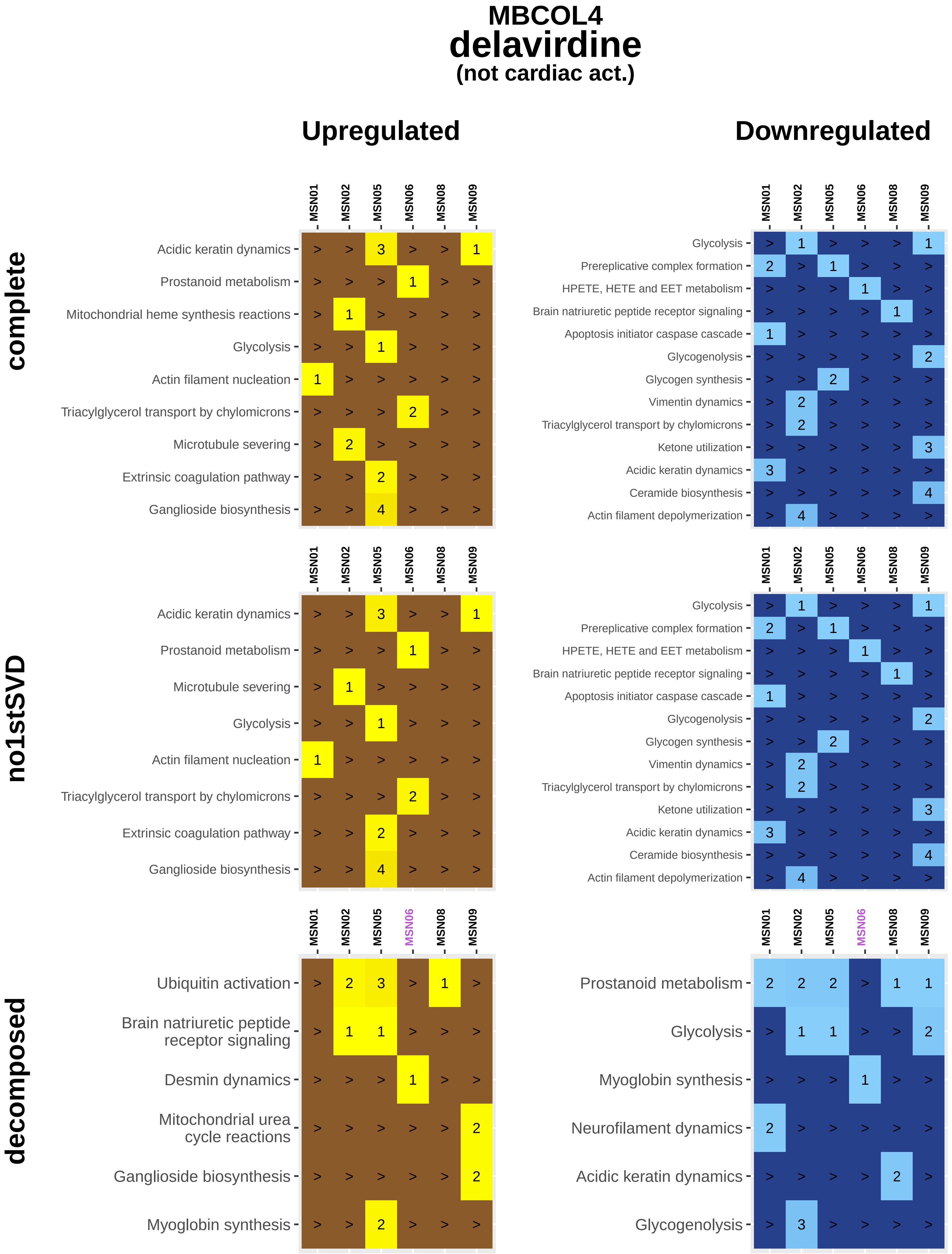
MBCOL4
delavirdine
(not cardiac act.)
Upregulated
Downregulated
complete
no1stSVD
decomposed
MSN01
MSN02
MSN05
MSN06
MSN08
MSN09
Acidic keratin dynamics
Prostanoid metabolism
Mitochondrial heme synthesis reactions
Glycolysis
Actin filament nucleation
Triacylglycerol transport by chylomicrons
Microtubule severing
Extrinsic coagulation pathway
Ganglioside biosynthesis
Prereplicative complex formation
HPETE, HETE and EET metabolism
Brain natriuretic peptide receptor signaling
Apoptosis initiator caspase cascade
Glycogenolysis
Glycogen synthesis
Vimentin dynamics
Ketone utilization
Ceramide biosynthesis
Actin filament depolymerization
Ubiquitin activation
Desmin dynamics
Mitochondrial urea cycle reactions
Myoglobin synthesis
Neurofilament dynamics

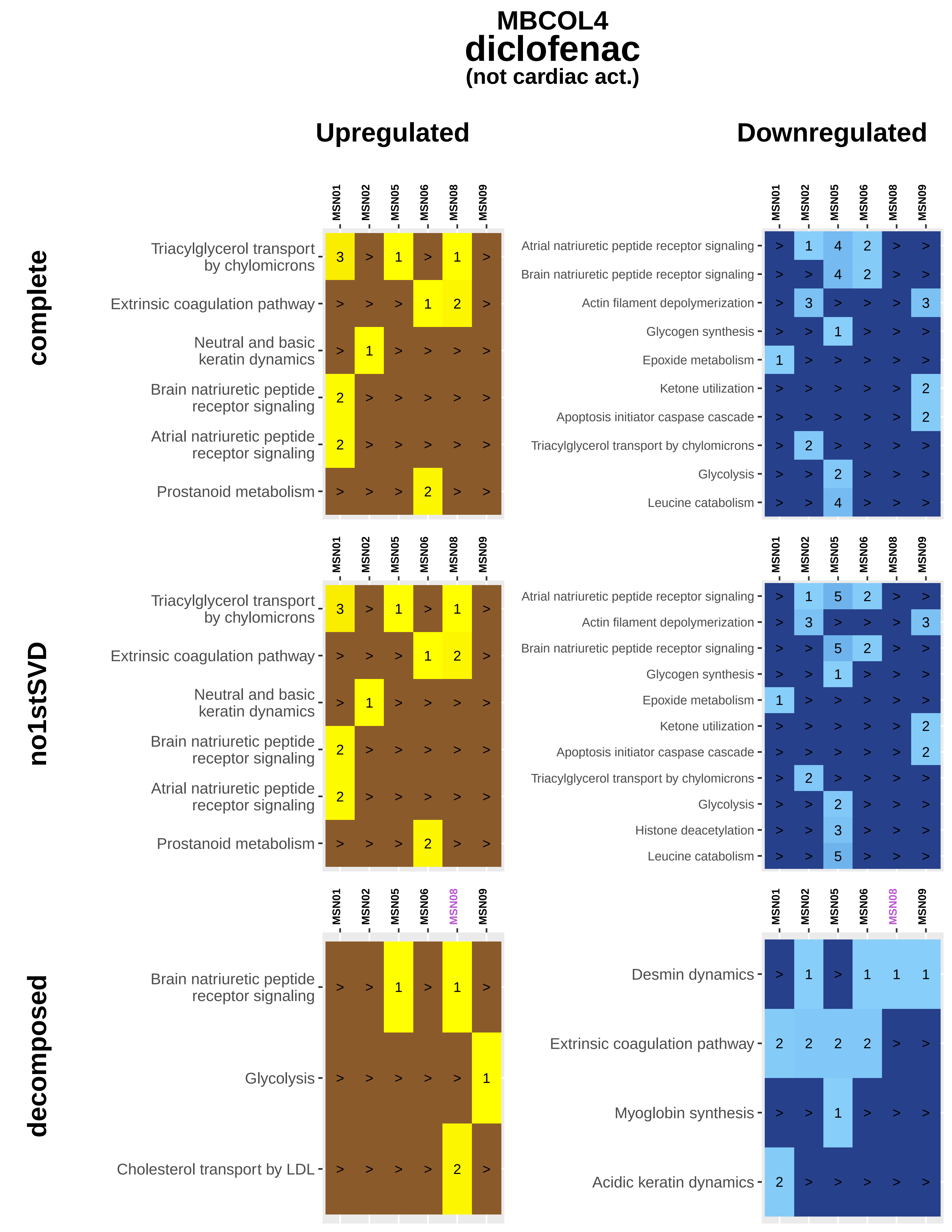
MBCOL4
diclofenac
(not cardiac act.)
Upregulated
Downregulated
complete
no1stSVD
decomposed
MSN01
MSN02
MSN05
MSN06
MSN08
MSN09
Triacylglycerol transport by chylomicrons
Extrinsic coagulation pathway
Neutral and basic keratin dynamics
Brain natriuretic peptide receptor signaling
Atrial natriuretic peptide receptor signaling
Prostanoid metabolism
Actin filament depolymerization
Glycogen synthesis
Epoxide metabolism
Ketone utilization
Apoptosis initiator caspase cascade
Glycolysis
Leucine catabolism
Histone deacetylation
Cholesterol transport by LDL
Desmin dynamics
Myoglobin synthesis
Acidic keratin dynamics

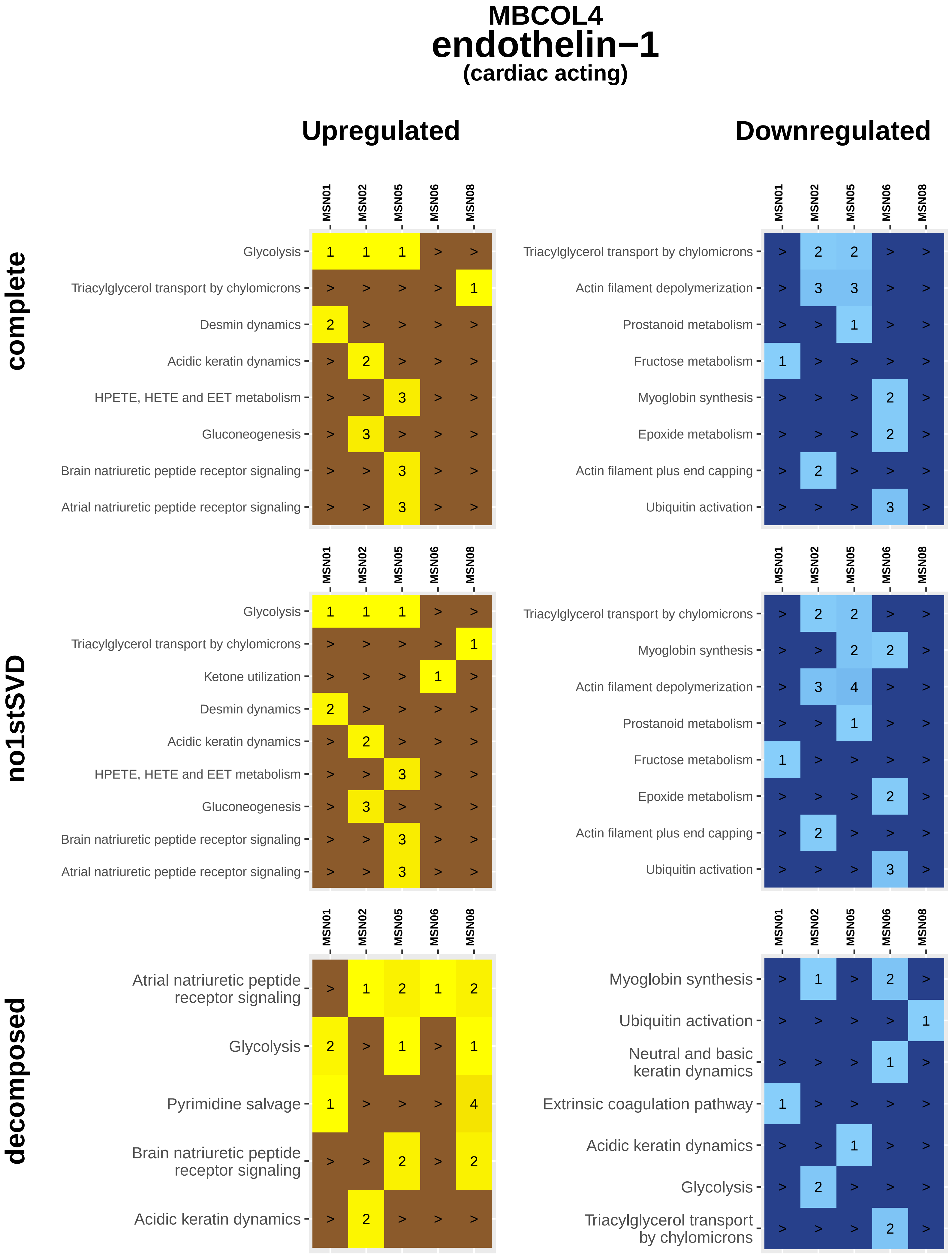
MBCOL4
endothelin−1
(cardiac acting)
Upregulated
Downregulated
complete
no1stSVD
decomposed
MSN01
MSN02
MSN05
MSN06
MSN08
Glycolysis
Triacylglycerol transport by chylomicrons
Desmin dynamics
Acidic keratin dynamics
HPETE, HETE and EET metabolism
Gluconeogenesis
Brain natriuretic peptide receptor signaling
Atrial natriuretic peptide receptor signaling
Actin filament depolymerization
Prostanoid metabolism
Fructose metabolism
Myoglobin synthesis
Epoxide metabolism
Actin filament plus end capping
Ubiquitin activation
Ketone utilization
Pyrimidine salvage
Neutral and basic keratin dynamics
Extrinsic coagulation pathway

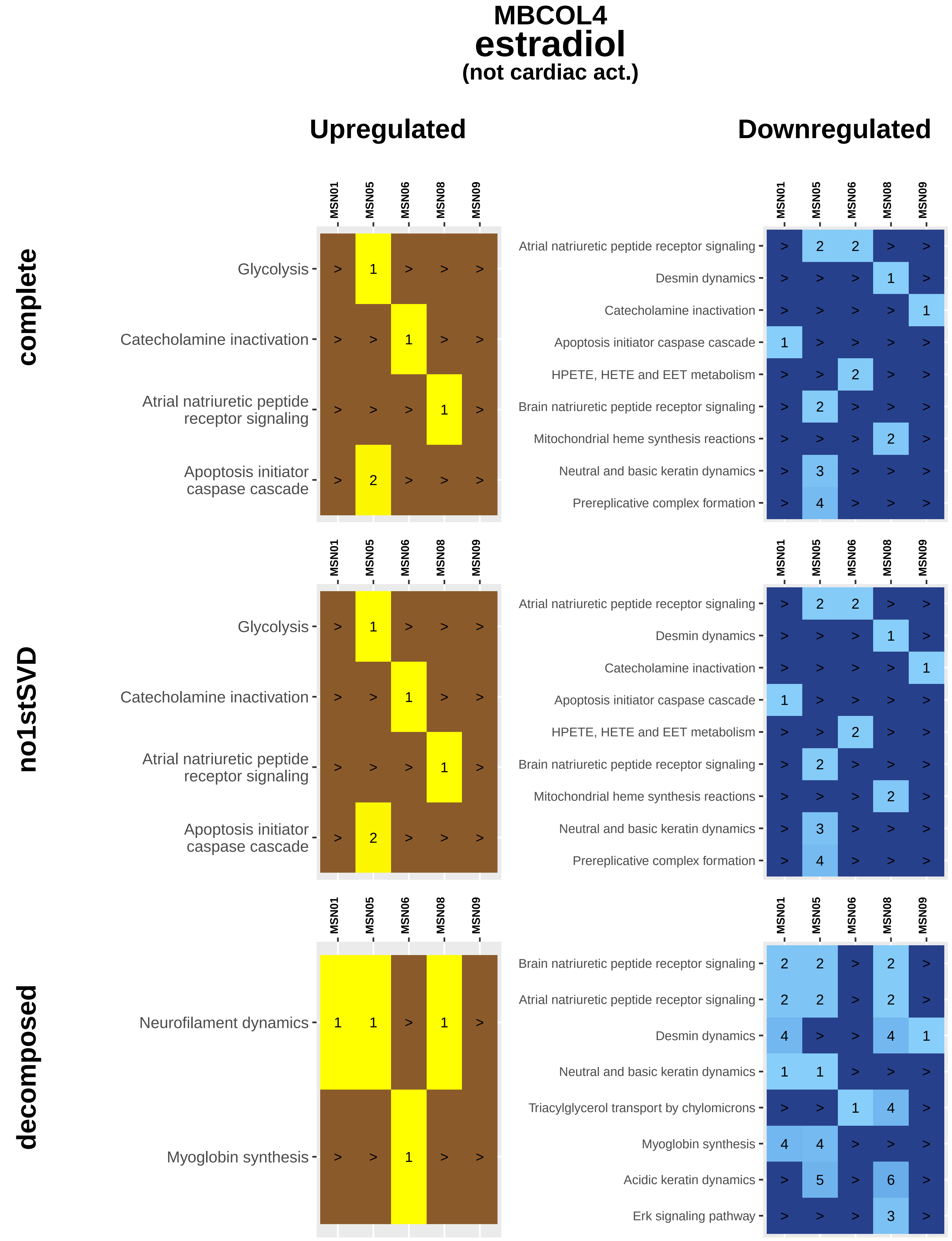

MBCOL4
estradiol
(not cardiac act.)
Upregulated
Downregulated
complete
no1stSVD
decomposed
MSN01
MSN05
MSN06
MSN08
MSN09
Glycolysis
Catecholamine inactivation
Atrial natriuretic peptide receptor signaling
Apoptosis initiator caspase cascade
Desmin dynamics
HPETE, HETE and EET metabolism
Brain natriuretic peptide receptor signaling
Mitochondrial heme synthesis reactions
Neutral and basic keratin dynamics
Prereplicative complex formation
Neurofilament dynamics
Myoglobin synthesis
Triacylglycerol transport by chylomicrons
Acidic keratin dynamics
Erk signaling pathway

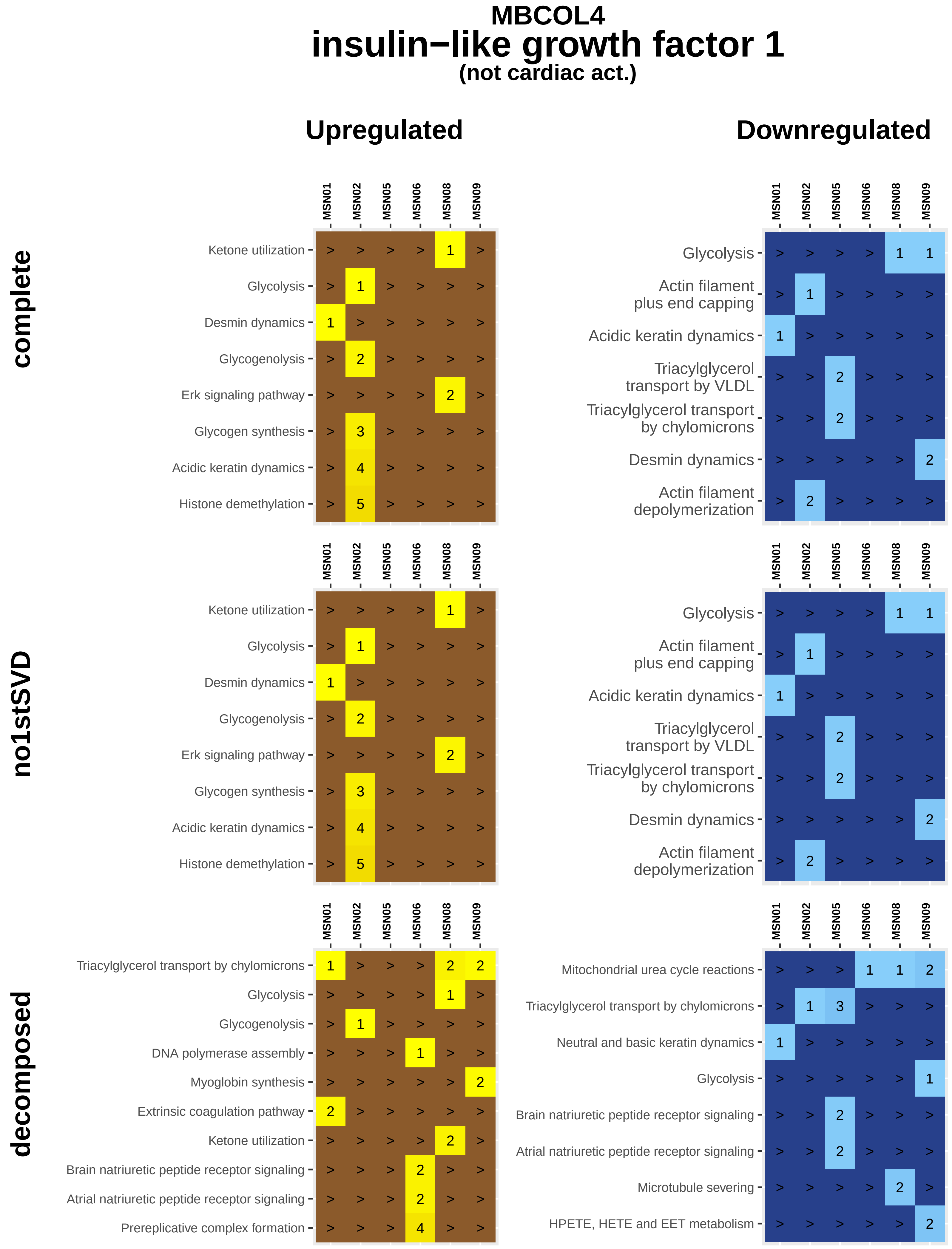
MBCOL4
insulin−like growth factor 1
(not cardiac act.)
Upregulated
Downregulated
complete
no1stSVD
decomposed
MSN01
MSN02
MSN05
MSN06
MSN08
MSN09
Ketone utilization
Glycolysis
Desmin dynamics
Glycogenolysis
Erk signaling pathway
Glycogen synthesis
Acidic keratin dynamics
Histone demethylation
Actin filament plus end capping
Triacylglycerol transport by VLDL
Triacylglycerol transport by chylomicrons
Actin filament depolymerization
DNA polymerase assembly
Myoglobin synthesis
Extrinsic coagulation pathway
Brain natriuretic peptide receptor signaling
Atrial natriuretic peptide receptor signaling
Prereplicative complex formation
Mitochondrial urea cycle reactions
Neutral and basic keratin dynamics
Microtubule severing
HPETE, HETE and EET metabolism

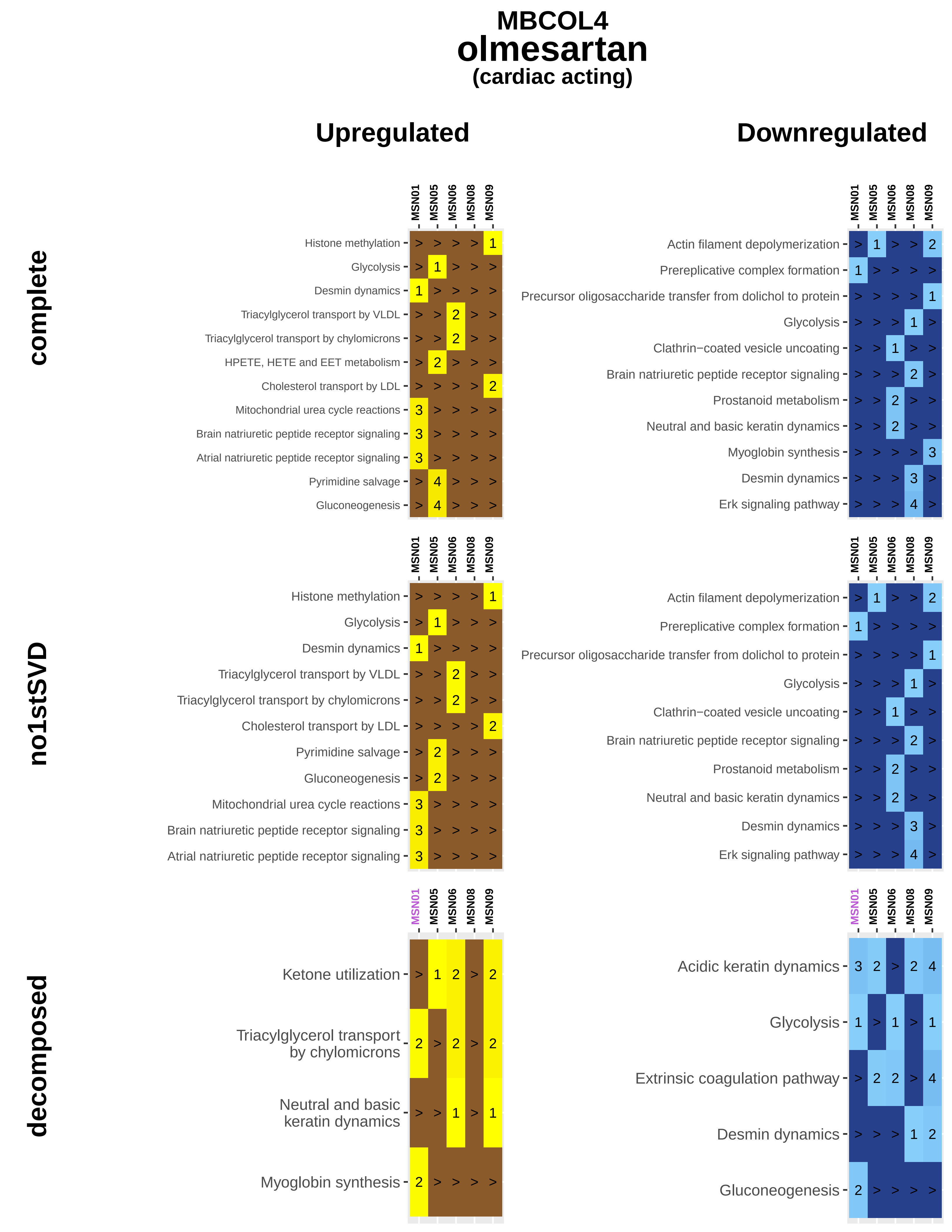

MBCOL4
olmesartan
(cardiac acting)
Upregulated
Downregulated
complete
no1stSVD
decomposed
MSN01
MSN05
MSN06
MSN08
MSN09
Histone methylation
Glycolysis
Desmin dynamics
Triacylglycerol transport by VLDL
Triacylglycerol transport by chylomicrons
HPETE, HETE and EET metabolism
Cholesterol transport by LDL
Mitochondrial urea cycle reactions
Brain natriuretic peptide receptor signaling
Atrial natriuretic peptide receptor signaling
Pyrimidine salvage
Gluconeogenesis
Actin filament depolymerization
Prereplicative complex formation
Precursor oligosaccharide transfer from dolichol to protein
Clathrin−coated vesicle uncoating
Prostanoid metabolism
Neutral and basic keratin dynamics
Myoglobin synthesis
Erk signaling pathway
Ketone utilization
Triacylglycerol transport by chylomicrons
Neutral and basic keratin dynamics
Acidic keratin dynamics
Extrinsic coagulation pathway

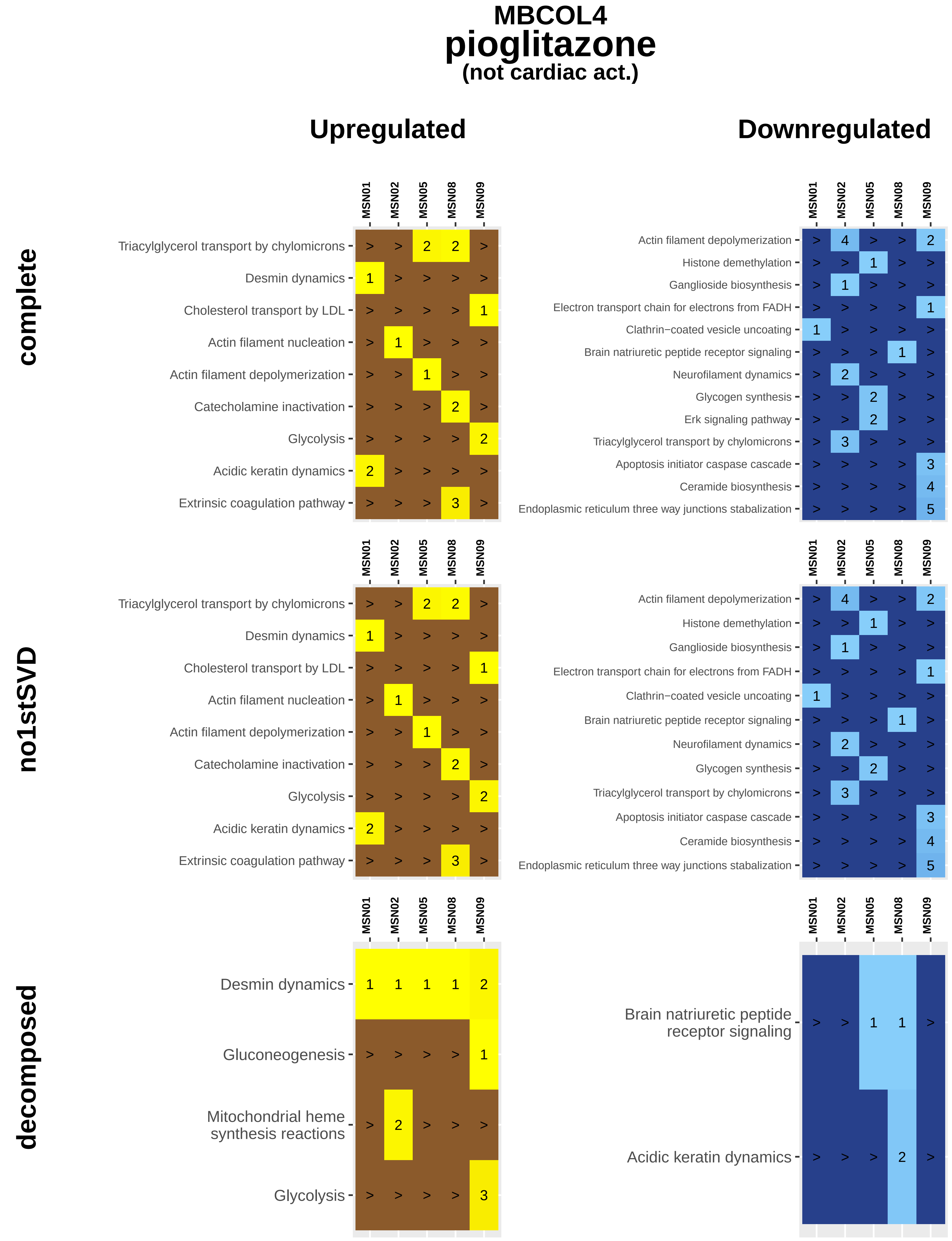
MBCOL4
pioglitazone
(not cardiac act.)
Upregulated
Downregulated
complete
no1stSVD
decomposed
MSN01
MSN02
MSN05
MSN08
MSN09
Triacylglycerol transport by chylomicrons
Desmin dynamics
Cholesterol transport by LDL
Actin filament nucleation
Actin filament depolymerization
Catecholamine inactivation
Glycolysis
Acidic keratin dynamics
Extrinsic coagulation pathway
Histone demethylation
Ganglioside biosynthesis
Electron transport chain for electrons from FADH
Clathrin−coated vesicle uncoating
Brain natriuretic peptide receptor signaling
Neurofilament dynamics
Glycogen synthesis
Erk signaling pathway
Apoptosis initiator caspase cascade
Ceramide biosynthesis
Endoplasmic reticulum three way junctions stabalization
Gluconeogenesis
Mitochondrial heme synthesis reactions
Brain natriuretic peptide receptor signaling

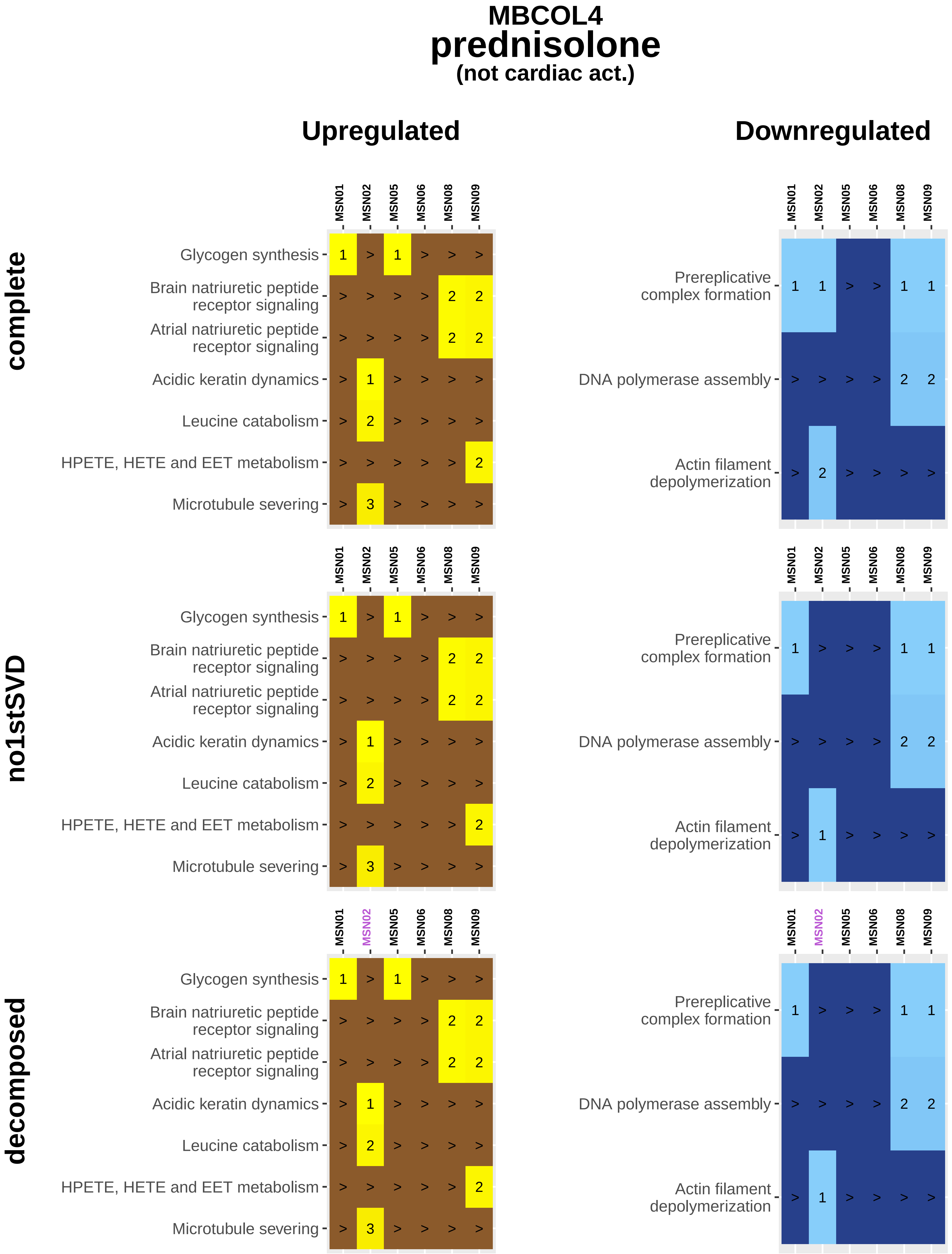
MBCOL4
prednisolone
(not cardiac act.)
Upregulated
Downregulated
complete
no1stSVD
decomposed
MSN01
MSN02
MSN05
MSN06
MSN08
MSN09
Glycogen synthesis
Brain natriuretic peptide receptor signaling
Atrial natriuretic peptide receptor signaling
Acidic keratin dynamics
Leucine catabolism
HPETE, HETE and EET metabolism
Microtubule severing
Prereplicative complex formation
DNA polymerase assembly
Actin filament depolymerization

**MBCOL4**

### rosiglitazone

**(not cardiac act.)**

#### Upregulated

#### Downregulated

complete

Triacylglycerol tra
by chylor

Actin filament nuc

Brain natriuretic
receptor si

Atrial natriuretic
receptor si

Extrinsic coagulation p

Actin f
depolyme

Neurofilament dy

Triacylglycerol transport by chylomicro

Phosphatidylcholine, phoshpatidylethanolamine and phosphatidylserine interconvers

Glycogen synthe

Desmin dynam

Apoptosis initiator caspase casca

Leucine catabol

Brain natriuretic peptide receptor signal

Atrial natriuretic peptide receptor signal

no1stSVD

Triacylglycerol transport by chyl

Extrinsic coagulation

Microtubule

G

Brain natriuretic peptide receptor

Atrial natriuretic peptide receptor

Actin filament n

Neurofilament c

Actin filament depolym

Triacylglycerol transport by chylomicro

Phosphatidylcholine, phoshpatidylethanolamine and phosphatidylserine interconvers

Glycogen synthe

Desmin dynam

Apoptosis initiator caspase casca

Leucine catabol

Brain natriuretic peptide receptor signal

Atrial natriuretic peptide receptor signal

decomposed

Myoglobin s

Triacylglycerol transport by chyl

Desmin c

G

Neurofilament c

Prereplicative complex f

Brain natriuretic peptide receptor

Atrial natriuretic peptide receptor

DNA polymerase a

Acidic keratin dynami

Neutral and bas
keratin dynami

HPETE, HETE and EET metabolis

Actin filame
depolymerizati

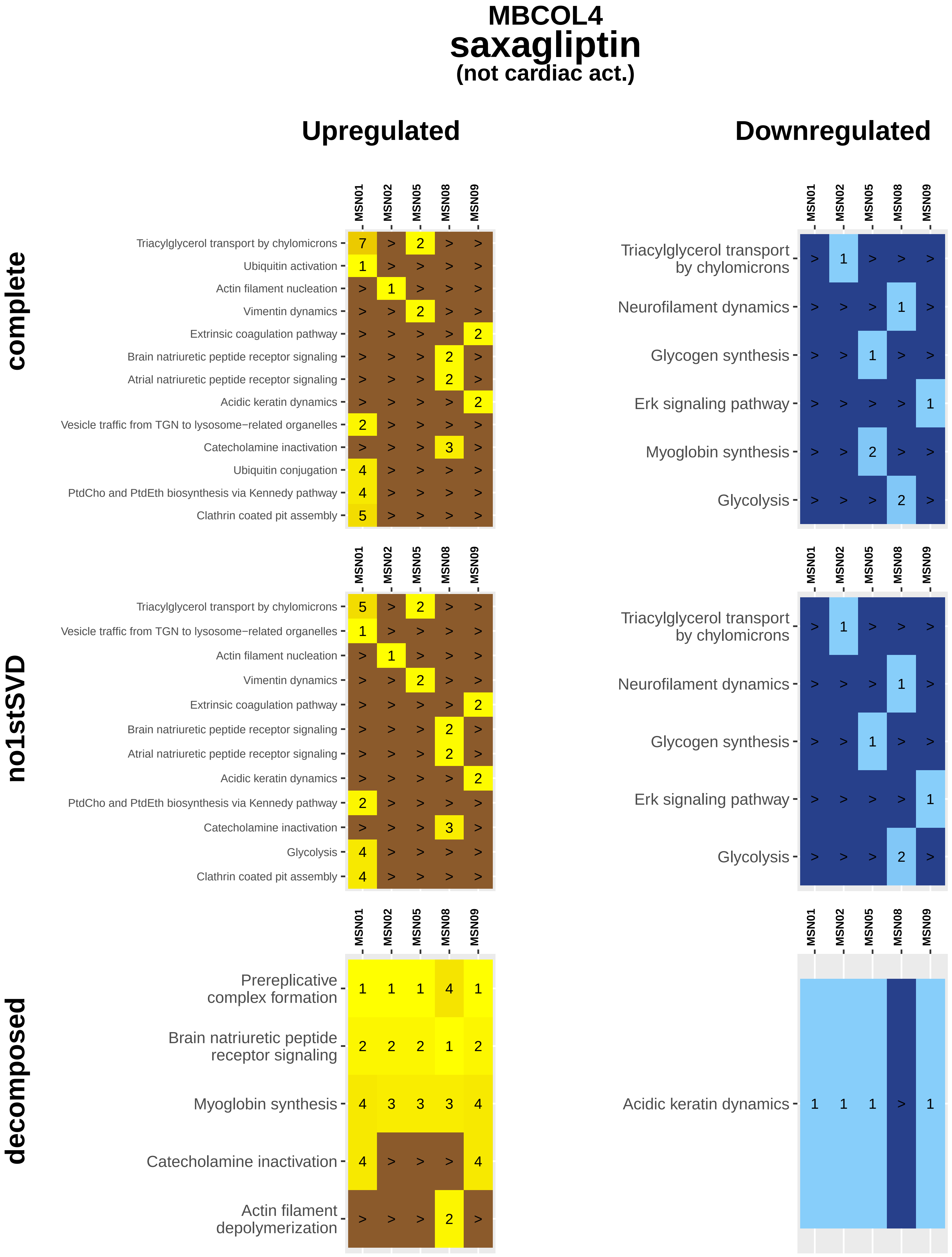
MBCOL4
saxagliptin
(not cardiac act.)
Upregulated
Downregulated
complete
no1stSVD
decomposed
MSN01
MSN02
MSN05
MSN08
MSN09
Triacylglycerol transport by chylomicrons
Ubiquitin activation
Actin filament nucleation
Vimentin dynamics
Extrinsic coagulation pathway
Brain natriuretic peptide receptor signaling
Atrial natriuretic peptide receptor signaling
Acidic keratin dynamics
Vesicle traffic from TGN to lysosome−related organelles
Catecholamine inactivation
Ubiquitin conjugation
PtdCho and PtdEth biosynthesis via Kennedy pathway
Clathrin coated pit assembly
Neurofilament dynamics
Glycogen synthesis
Erk signaling pathway
Myoglobin synthesis
Glycolysis
Prereplicative complex formation
Actin filament depolymerization

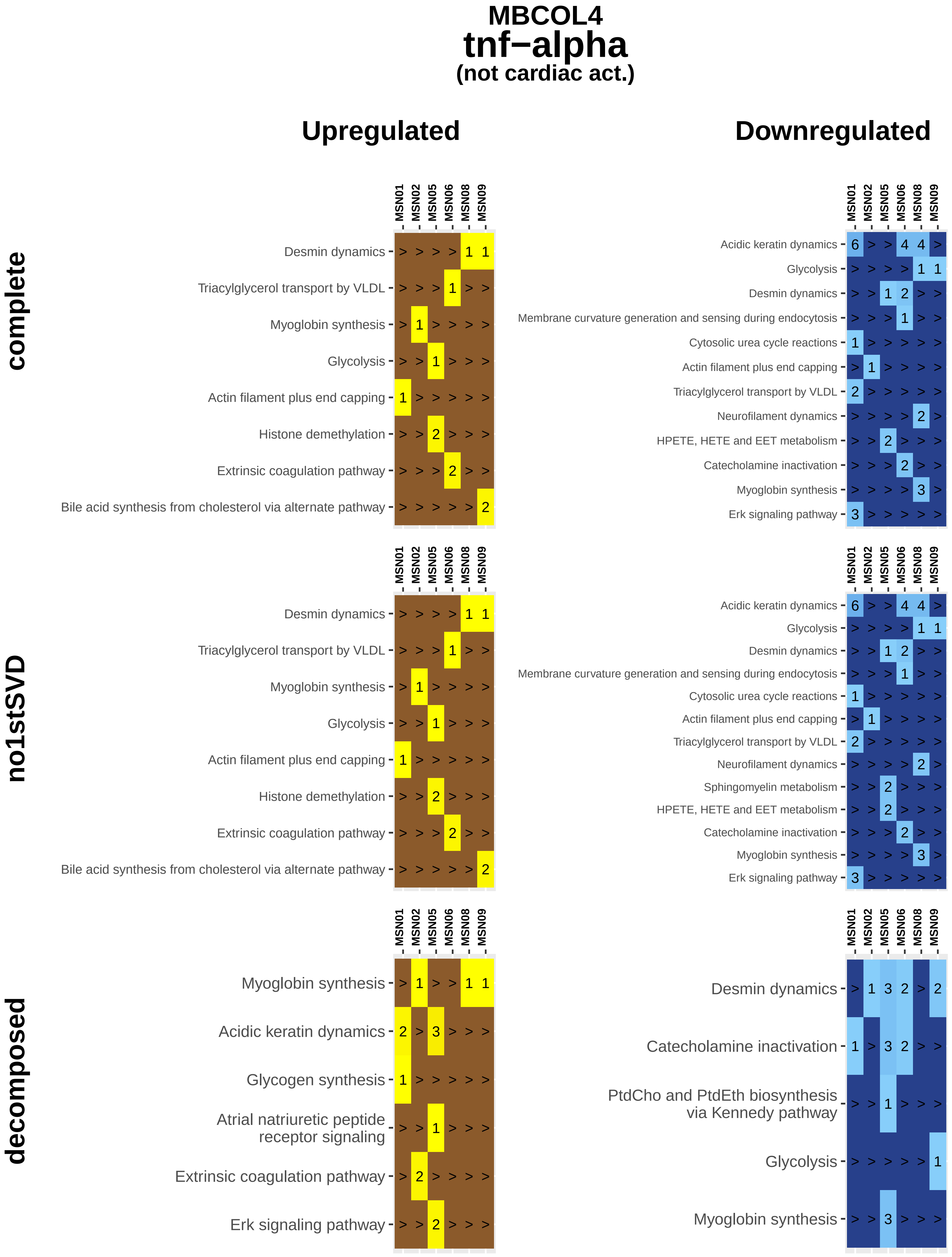
MBCOL4
tnf−alpha
(not cardiac act.)
Upregulated
Downregulated
complete
no1stSVD
decomposed
MSN01
MSN02
MSN05
MSN06
MSN08
MSN09
Desmin dynamics
Triacylglycerol transport by VLDL
Myoglobin synthesis
Glycolysis
Actin filament plus end capping
Histone demethylation
Extrinsic coagulation pathway
Bile acid synthesis from cholesterol via alternate pathway
Acidic keratin dynamics
Membrane curvature generation and sensing during endocytosis
Cytosolic urea cycle reactions
Neurofilament dynamics
HPETE, HETE and EET metabolism
Catecholamine inactivation
Erk signaling pathway
Sphingomyelin metabolism
Glycogen synthesis
Atrial natriuretic peptide receptor signaling
PtdCho and PtdEth biosynthesis via Kennedy pathway
