## Supplemental Tables 1-22 for "Multiscale mapping of transcriptomic signatures for cardiotoxic drugs": Suppl_Table_03 - Descritpion of materials used in this study.pdf

**Metadata**

1. **Materials/Reagents:** Company name, Catalogue and lot numbers

| PRODUCT | COMPANY NAME | CAT # | Storage | Usable life |
|---|---|---|---|---|
| DMEM (500 mL) | Life Technologies | 11965-118 | +4°C | M |
| IMDM | Life Technologies | 12440-053 | +4°C | M |
| DMEM/F12 | Life Technologies | 11330057 | +4°C | M |
| Penicillin/Streptomycin (100x) | Life Technologies | 15140-122 | +4°C | M |
| Sodium-Pyruvate (100 mM) | Life Technologies | 11360-070 | +4°C | M |
| L-Glutamine (200 mM) | Life Technologies | 25030-081 | +4°C | M |
| Non-Essential Amino Acids (100x) | Life Technologies | 11140-050 | +4°C | M |
| Fetal Bovine Serum (FBS; 500 mL) | Corning | 35-011-CV | +4°C | M |
| EDTA | Corning | 46-034-CI | RT | M |
| KnockOut Serum Replacement | Life Technologies | 10828-028 | -20°C<br>+4°C | M |
| mTeSR™:<br>1) mTeSR™ Basal medium<br>2) mTeSR™ 1.5X Supplement | STEMCELL Technologies | 05850 | +4°C<br>-20°C | M |
| 2-Mercaptoethanol | MP Biomedicals | 194705 | RT | N/A |
| TrypLE Express (1X) | Life Technologies | 12605010 | +4°C | M |
| ReLeSR™ | STEMCELL Technologies | 05872 | RT | M |
| Gelatin | Sigma | G1890 | RT | M |
| Matrigel | Corning | 254248 | -20°C<br>+4°C | M |
| Dulbecco's phosphate-buffered saline (DPBS) | Life Technologies | 14190-136 | +4°C | M |
| DMSO | Fisher Scientific | BP2311 | RT | M |
| FGF2 | R&D Systems | 223-FB-10 | -20°C<br>+4°C | M |
| Thiazovivin | Millipore | 420220 | -20°C<br>+4°C | M |
| Irradiated MEFs (Mouse Embryonic Fibroblasts) | Global Stem | GSC-6001G | Liquid Nitrogen | M |
| mRNA Reprogramming Kit | Stemgent | 00-0071 | -70°C | M |
| microRNA Booster Kit | Stemgent | 00-0073 | -70°C | M |
| B18R | Stemgent | 03-0071 | -70°C | M |
| Pluriton Supplement (2500x) | Stemgent | 01-0061 | -80°C | M |
| Pluriton Medium (500 mL) | Stemgent | 01-0015 | -20°C<br>+4°C | M |
| Stemfect RNA Transfection Kit:<br>1) Stemfect Transfection Buffer<br>2) Stemfect Transfection Reagent | Stemgent | 00-0069 | +4°C | M |

| | | | | |
|---|---|---|---|---|
| PureLink™ Genomic DNA Mini Kit | Life Technologies | K1820-001 | RT | N/A |
| e-Myco™ plus Mycoplasma PCR Detection KIT | iNtRON Biotechnology | 25235 | -20˚C | M |
| 50 ml conical tube | BD FALCON | 352098 | RT | N/A |
| 15 ml conical tube | BD FALCON | 352099 | RT | N/A |
| 75 $cm^2$ Cell Culture Flask | Corning | 430641 | RT | N/A |
| 50 mL Rapid Flow Conical Filter with a 0.2 µm aPES membrane | Thermo Scientific | 564-0020 | RT | N/A |
| 6 well Tissue Culture (TC) plate | Corning | 353046 | RT | N/A |
| 2 ml Aspirating pipettes | BD FALCON | 357558 | RT | N/A |
| 5 ml Serological Pipettes | BD FALCON | 357543 | RT | N/A |
| 10 ml Serological Pipettes | BD FALCON | 357551 | RT | N/A |
| Hemocytomter | Thermo Scientific | 0267110 | RT | N/A |
| 1 mL filter tips | USA Scientific | 1126-7810 | RT | N/A |
| 200 µL filter tips | USA Scientific | 1120-8810 | RT | N/A |
| 20 µL filter tips | USA Scientific | 1123-1810 | RT | N/A |
| 0.1-10 µL filter tips | USA Scientific | 1121-3810 | RT | N/A |
| Cryovials | Nunc | 377367 | RT | N/A |
| Cryo 1˚C freezing container | Nalgene | 5100-0001 | RT | N/A |
| 2-Propanol | Thermo Scientific | A417-4 | RT | N |

M; according to manufacture's shelf-life information; RT, room temperature

2. **Subject(s):** subject ID (de-identified), age, gender/sex, race/ethnicity

3. **Fibroblast(s):** subject ID (de-identified), passage number, dates of each passage, date of biopsy, date of initial plating, date of freezing.

4. **Microscopy pictures:** subject ID (de-identified), passage number, date, microscope name (company, catalogue number), magnification.

This table is taken from the Supplementary Materials for Schaniel et al. (Stem Cell Reports. 2021 Dec 14;16(12):3036-3049), since the two studies were conducted concurrently.
